# Proteomic Ligandability Maps of Phosphorus(V) Stereoprobes Identify Covalent TLCD1 Inhibitors

**DOI:** 10.1101/2025.01.31.635883

**Authors:** Hayden A. Sharma, Michael Bielecki, Meredith A. Holm, Ty M. Thompson, Yue Yin, Jacob B. Cravatt, Timothy B. Ware, Alex Reed, Molham Nassir, Tamara El-Hayek Ewing, Bruno Melillo, J Fernando Bazan, Phil S. Baran, Benjamin F. Cravatt

**Affiliations:** Department of Chemistry, Scripps Research, 10550 North Torrey Pines Road, La Jolla, California, 92037, US; ℏ bioconsulting llc, Stillwater, Minnesota, 55082, US

## Abstract

Activity-based protein profiling (ABPP) of stereoisomerically defined sets of electrophilic compounds (‘stereoprobes’) offers a versatile way to discover covalent ligands for proteins in native biological systems. Here we report the synthesis and chemical proteomic characterization of stereoprobes bearing a P(V)-oxathiaphospholane (OTP) reactive group. ABPP experiments identified numerous proteins in human cancer cells that showed stereoselective reactivity with OTP stereoprobes, and we confirmed several of these liganding events with recombinant proteins. OTP stereoprobes engaging the poorly characterized transmembrane protein TLCD1 impaired the incorporation of monounsaturated fatty acids into phosphatidylethanolamine lipids in cells, a lipidomic phenotype that mirrored genetic disruption of this protein. Using AlphaFold2, we found that TLCD1 structurally resembles the ceramide synthase and fatty acid elongase families of coenzyme A-dependent lipid processing enzymes. This structural similarity included conservation of catalytic histidine residues, the mutation of which blocked the OTP stereoprobe reactivity and lipid remodeling activity of recombinant TLCD1. Taken together, these data indicate that TLCD1 acts as a lipid acyltransferase in cells, and that OTP stereoprobes function as inhibitors of this enzymatic activity. Our findings thus illuminate how the chemical proteomic analysis of electrophilic compounds can facilitate the functional annotation and chemical inhibition of a key lipid metabolic enzyme in human cells.

## Introduction

Small molecules are critical tools for probing the biological functions of proteins and comprise a large fraction of therapeutic agents to treat human disease.^1^ Most human proteins, however, lack chemical probes, and this challenge has inspired the development of new ways to assay proteins for interactions with small molecules. For instance, binding-first assays,^2^ such as fragment-based ligand discovery ^3^ and DNA-encoded libraries,^4^ offer general ways to discover ligands but typically rely on purified proteins studied outside of the environment of the cell, where post-translational mechanisms can regulate protein structure and function in diverse ways.

Chemical proteomic methods, such as activity-based protein profiling (ABPP), have emerged as a complementary and versatile approach for ligand discovery in native biological systems and, when combined with covalent (electrophilic or photoreactive) small-molecule libraries and quantitative mass spectrometry (MS), can assess small molecule-protein interactions on a global scale.^5^ The resulting ligandability maps of human cells have emphasized the broad small molecule-binding potential of proteins from diverse structural and functional classes, including protein types historically considered undruggable (e.g., RNA/DNA-binding proteins, adaptors).^6^

Original compound libraries studied by ABPP mainly featured fragment-like small molecules, which have an advantage of broadly surveying protein pockets of diverse size and shape but tend to furnish hit compounds with limited potency and selectivity.^6^ We have more recently introduced a complementary strategy where ABPP is performed with more structurally elaborated, chiral small molecules consisting of densely functionalized sp3-rich cores bearing electrophilic or photoreactive groups.^7^ All library members are synthesized stereoisomerically pure, and the resulting ‘stereoprobes’ can then be interpreted for their stereoselective interactions with proteins by ABPP, thereby providing inactive enantiomeric control compounds for subsequent biology experiments. This approach has identified cysteine-directed electrophilic stereoprobes that modulate the functions of a wide range of proteins, including transcription factors,^7f^ RNA-binding proteins,^7g^ adaptors,^7d,h^ and enzymes.^7c^

Most of the electrophilic stereoprobes described to date have incorporated an acrylamide electrophile that preferentially reacts with cysteine residues in proteins. However, the proteome is expected to contain many ligandable pockets that lack proximal cysteines, and there is accordingly a broad interest in identifying electrophilic groups that can target additional nucleophilic amino acids, including lysines, apartates, glutamates, methionines, tyrosines, arginines, and histidines.^8^ In one notable example, chiral small molecule fragments bearing a sulfonyl fluoride electrophile were found by chemical proteomics to stereoselectively engage lysines and tyrosines on a wide array of human proteins.^9^ Here, we considered whether the P(V)-oxathiaphospholane (OTP) electrophile,^10^ which we recently introduced for selective bioconjugation and found to react in the presence of base and organic solvent with exposed serine and, to a lesser extent, tyrosine, lysine, and cysteine residues on purified proteins, can be incorporated into stereoprobes to generate compounds that stereoselectively react with proteins in human cells. We describe the synthesis and chemical proteomic analysis of a focused set of OTP stereoprobes constructed on azetidine and tryptoline scaffolds. Despite showing attenuated global proteomic reactivity profiles compared to acrylamide stereoprobes, the OTP stereoprobes stereoselectively engaged a diverse set of proteins. We confirmed several stereoselective liganding events for the OTP stereoprobes with recombinant proteins, including the poorly characterized multipass transmembrane protein TLCD1, which has been shown to modulate the phospholipid composition of cell membranes.^11,12^ We found that OTP stereoprobes engaging TLCD1 stereoselectively suppress incorporation of isotopically labeled oleic acid into phosphatidylethanolamines in cells in a manner similar to the genetic disruption of TLCD1. We additionally provide evidence from AlphaFold2 predictions that TLCD1 is structurally and mechanistically related to the ceramide synthase and fatty acid elongase families of enzymes,^13,14^ including conservation of catalytic histidine residues that are required for the OTP stereoprobe reactivity and lipid remodeling activity of TLCD1. These results indicate TLCD1 functions as an acyltransferase that modulates the monounsaturated lipid content of cell membranes and designate OTP stereoprobes as inhibitors of this activity.

## Results

### Design and Initial Characterization of OTP Stereoprobes

Previous ABPP studies have found that cysteine-directed acrylamide stereoprobes constructed on azetidine and tryptoline cores exhibit numerous stereoselective and site-specific protein liganding events in human cells.^7^ We therefore designed and synthesized OTP stereoprobes that also leveraged azetidine and tryptoline scaffolds (Figure 1A, B). We additionally introduced two different (hetero)-aromatic recognition elements into the azetidine OTP stereoprobes, in part leveraging a new approach for the synthesis of azetidine scaffolds previously reported by our group,^15^ to assess how such substitutions impacted protein interactions. A total of 24 OTP stereoprobes (**1**–**24**; 12 non-alkyne; 12 alkyne) were synthesized.

**Figure 1.**
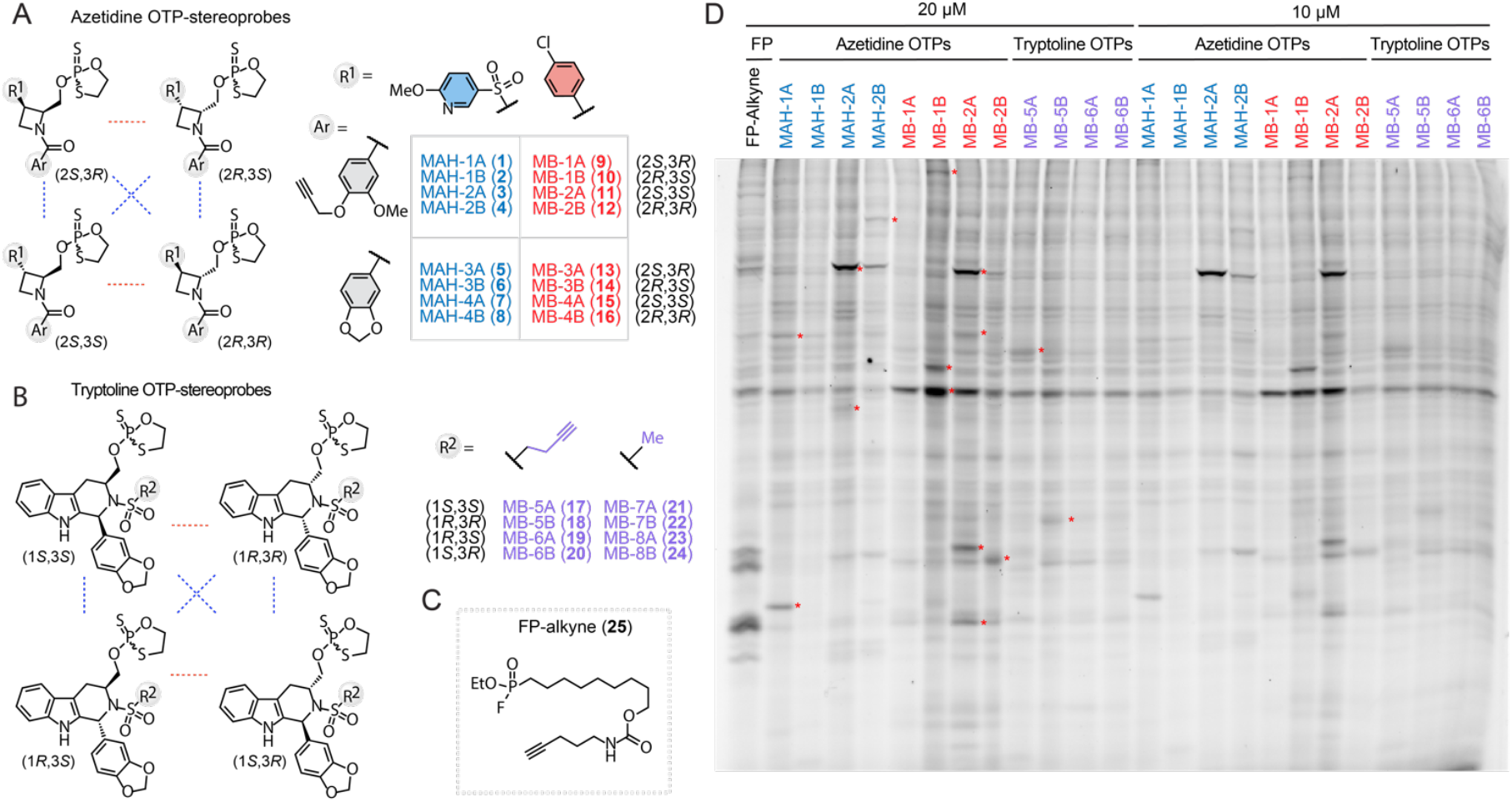
Structures and initial characterization of OTP stereoprobes. (A, B) Structures of azetidine (A) and tryptoline (B) OTP stereoprobes. Structure of a fluorophosphonate (FP)-alkyne probe. (D) Gel-ABPP data for Ramos cells treated with the indicated alkyne probes (10 or 20 µM, 3 h). Following cell lysis stereoprobe-reactive proteins were visualized by CuAAC conjugation to an azide−rhodamine reporter group, SDS-PAGE, and in-gel fluorescence scanning. Red asterisks mark representative proteins that were stereoselectively engaged by OTP stereoprobes (shown for 20 μM condition). Data are from a single experiment representative of two independent experiments.

We performed initial gel-ABPP experiments to compare the proteome-wide reactivity of each set of alkynylated OTP stereoprobes (azetidines: MB-1A/1B//2A/2B, MAH-1A/1B//2A/2B; tryptolines: MB-5A/5B//6A/6B). In this analysis, we also included a fluorophosphonate (FP)- alkyne (**25**, Figure 1C),^16,17^ as FPs are an established class of P(V) activity-based probes that preferentially target serine hydrolases.^16^ Exposure of the Ramos human B-cell cancer line to alkynylated probes (10 or 20 μM, 3 h), followed by lysis and copper-catalyzed azide−alkyne cycloaddition (CuAAC)^18^ conjugation of probe-labeled proteins to an azide−rhodamine reporter tag and in-gel fluorescence scanning,^19^ revealed numerous stereoselective OTP stereoprobe-protein interactions that were also distinct from the proteins reacting with FP-alkyne (marked with red asterisks for the 20 µM condition; Figure 1D). Additional gel-ABPP studies revealed that the OTP stereoprobes displayed much lower overall proteomic reactivity compared to acrylamide stereoprobes, even when assayed at 4–10-fold higher concentrations (Figure S1). Both sets of azetidine OTP stereoprobes displayed concentration-dependent increases in proteomic reactivity when tested in Ramos cells at up to 50 µM, whereas similar degrees of protein reactivity were observed for the tryptoline OTP stereoprobes across a concentration range of 20-50 µM (Figure S2), suggesting that these compounds may have limited solubility or exhibit saturable uptake into cells. Based on the gel-ABPP data, we selected concentrations of 50 µM and 20 μM, respectively, to evaluate azetidine and tryptoline OTP stereoprobes by mass spectrometry (MS)-ABPP.

### Protein-Directed ABPP of OTP Stereoprobes

We next evaluated the protein interactions of OTP stereoprobes by mass spectrometry (MS)- based proteomics using a protein-directed ABPP method^7h^ where Ramos cells were first treated with non-alkyne competitor stereoprobes (50 or 20 µM) or dimethyl sulfoxide (DMSO) for 2 h, followed by treatment with the corresponding alkyne stereoprobes (50 or 20 μM) for 1 h. After compound treatment, cells were lysed and alkyne stereoprobe-reactive proteins conjugated to biotin-azide by CuAAC, isolated with streptavidin beads, digested with trypsin, labeled with tandem mass tags (TMT), and identified (MS1/MS2 analysis) and quantified (MS3 analysis) by multiplexed (TMT16plex) MS-based proteomics (Figure 2A). Proteins showing greater than 2.5-fold enrichment by one stereoprobe compared to its enantiomer were assigned as *stereoselectively enriched* proteins. These proteins were further designated as being *stereoselectively liganded* if pretreatment by the corresponding competitor probe resulted in >50% blockade of stereoselective enrichment as compared to pretreatment with DMSO. While this protein-directed ABPP protocol sacrifices information on the specific amino acid residues reacting with stereoprobes, it provides, in our experience, a technically robust and straightforward method to globally map the proteins engaged by stereoprobes and to quantify the stoichiometry of these stereoprobe-protein interactions. We felt that these features were particularly advantageous for evaluating the protein targets of OTP stereoprobes, given their structural complexity (which can complicate MS2 fragmentation patterns required for assignment of probe-labeled peptides) and the potential for the OTP electrophile to react with a diverse range of amino acids.

**Figure 2.**
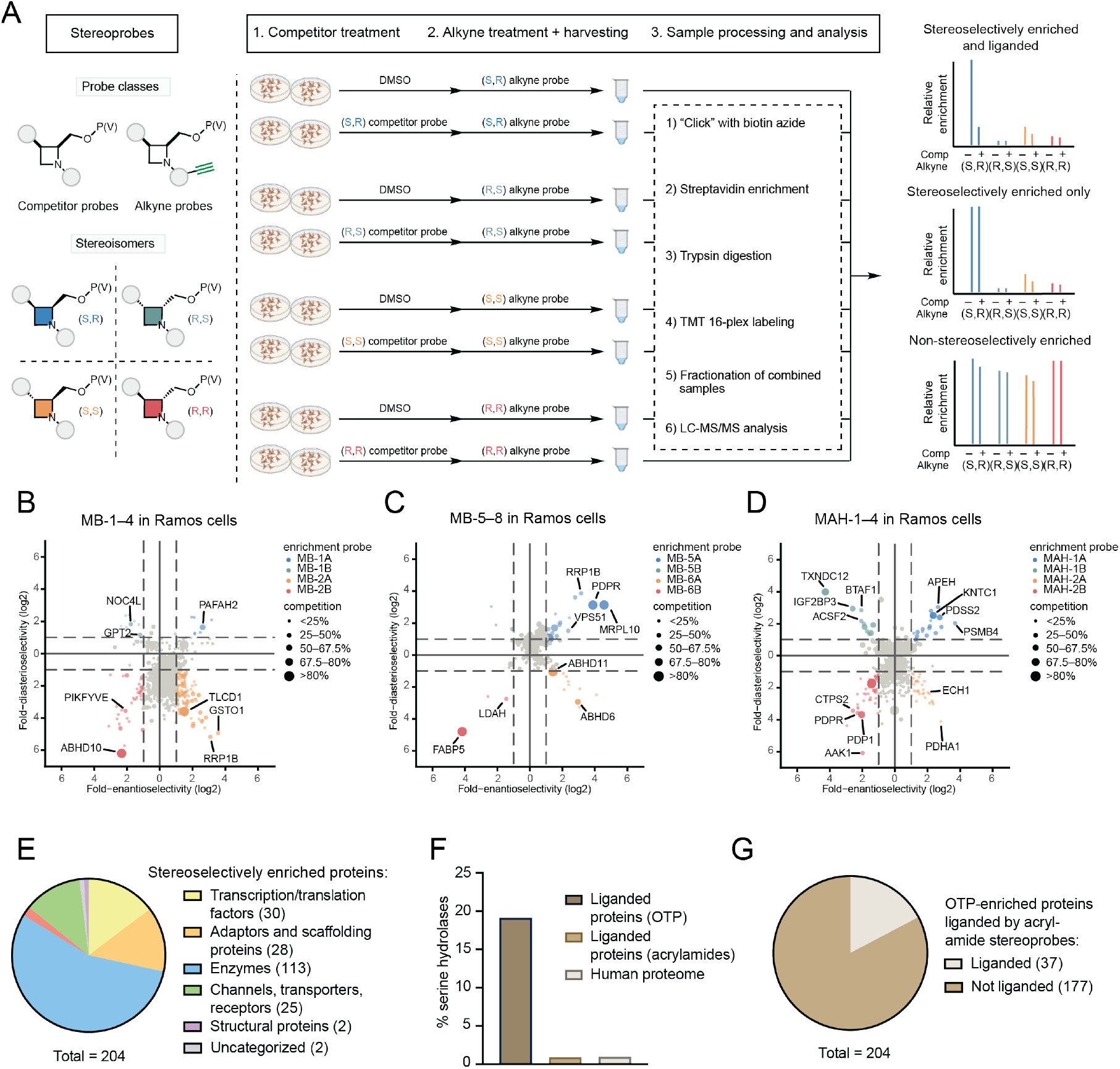
Protein-directed ABPP of OTP stereoprobes. (A) Workflow for protein-directed ABPP experiments where the stereoselective enrichment of proteins by alkyne stereoprobes and blockade of this enrichment by corresponding non-alkyne competitor stereoprobes are determined by multiplexed (TMT16plex) MS-based proteomics. (B–D) Quadrant plots highlighting enantio- and diastereo-selectively enriched proteins for each stereoconfiguration of alkyne OTP stereoprobes in Ramos cells. Cells were treated with competitor stereoprobes (20 µM MB-7/8, 50 µM MB-3/4, or 50 µM MAH-3/4) or DMSO for 2 h followed by the corresponding alkyne stereoprobes (20 µM MB-5/6, 50 µM MB-1/2, or 50 µM MAH-1/2) and then processed as summarized in (A). For each protein, enrichment enantioselectivity and diastereoselectivity (log2) are shown for the stereoprobe that produced the highest enrichment. Enantioselectivity is defined as the ratio of enrichment for one stereoisomer versus its enantiomer, and diastereoselectivity as the ratio of enrichment of one stereoisomer versus the average of its two diastereomers. A protein is shown in color if both the enantioselectivity and diastereoselectivity of enrichment (log2) are greater than 1. Competition is represented by dot size, as noted in the figure keys. Data are from two independent protein-directed ABPP experiments each performed with two technical replicates. (E) Functional class distribution of stereoselectively enriched proteins assigned by the GO (Panther), KEGG Brite, and Uniprot databases. (F) Fraction of proteins liganded by OTP or acrylamide stereoprobes annotated as serine hydrolases versus fraction of serine hydrolases annotated in the human proteome. (G) Fraction of proteins stereoselectively enriched by OTP stereoprobes that are also liganded by stereoisomeric acrylamides. Acrylamide stereoprobe data for (F, G) were obtained from refs. 7b, d, h and 15.

For each protein-directed ABPP experiment, the protein enrichment profiles can be visualized using quadrant plots (Figure 2B-D), where the positions of proteins on the x and y axes reflect enantioselective and diastereoselective enrichment, respectively, and the size of the dot represents the degree of competitive blockade of this enrichment by the corresponding non-alkyne stereoprobe. In total, we identified 204 and 22 proteins that were stereoselectively enriched and liganded by OTP stereoprobes, respectively, in Ramos cells (Figure S3 and Dataset S1). These proteins originated from diverse classes, including not only enzymes, but also transcription factors, adaptor/scaffolding proteins, and transporters (Figure 2E). Among the enzyme targets that were both stereoselectively enriched and liganded by OTP stereoprobes were several serine hydrolases (Figure 2F), which is consistent with the conserved serine nucleophile of this enzyme class showing strong reactivity with other types of P(V) electrophiles (e.g., fluoro- and aryloxy-phosph(on)ates).^16,20^ Also supportive of this relationship is the much smaller fraction of serine hydrolases liganded by stereoprobes bearing an alternative acrylamide electrophile (Figure 2G). More generally, we found that most proteins stereoselectively enriched by OTP stereoprobes have not been identified as targets of previously reported acrylamide stereoprobes (Figure 2G). Each of the three sets of OTP stereoprobes reacted with distinct proteins, and only a small fraction of these proteins cross-reacted with multiple sets of OTP stereoprobes (Figure S3 and Dataset S1). These results point to the importance of the core and recognition groups of the OTP stereoprobes in promoting stereoselective interactions with proteins. We additionally performed protein-directed ABPP experiments with the MAH-1–4 stereoprobes in a second cell line – OCI-AML-3 – which displayed a stereoprobe reactivity profile that was generally similar to Ramos cells (Figure S4 and Dataset S1), with the exception of some cell type-restricted interactions, particularly for OCI-AML-3 cells (e.g., innate immune-related proteins like STING and CD36).

### Characterization of OTP Stereoprobe–Protein Interactions

We next set out to confirm the interactions of OTP stereoprobes with representative proteins. FABP5—a member of the fatty acid-binding protein (FABP) family—was stereoselectively enriched by the tryptoline OTP MB-6B, and this enrichment was competed by MB-8B (Figure 3A). Using gel-ABPP, we confirmed stereoselective engagement of recombinantly expressed FABP5 in HEK293T cells by MB-6B and blockade of this engagement by MB-8B, but not MB-8A (Figure 3B). In attempting to deduce the site-of-reactivity for MB-6B/8B, we noticed that the distribution of TMT-reporter signals for the tryptic peptide of FABP5 containing amino acids (a.a.) 130-135 did not show the expected stereoselective enrichment profile shared by other tryptic peptides of FABP5 from MB-6B-treated cells (Figure 3C). In previous protein-directed ABPP studies, we determined that this type of “corrupted” stereo-enrichment profile for a tryptic peptide can occur when it contains the site of stereoprobe reactivity^7h^. The a.a. 130–135 peptide contains a tyrosine residue (Y131) located in the lipid-binding pocket of FABP5, and this tyrosine has been found to covalently react with sulfur(VI) compounds.^21^ Consistent with Y131 also being the site of OTP stereoprobe engagement, mutation of this tyrosine to phenylalanine (Y131F) blocked MB-6B reactivity (Figure 3D). FABP5 also reacts with azetidine acrylamide stereoprobes at a distinct residue – C120^7b^ – and we confirmed that mutation of this cysteine to alanine abolished liganding by the alkynylated azetidine acrylamide MY-11B^7b^ (Figure 3D). Interestingly, the C120A and Y131F mutants of FABP5 retained full respective reactivity with the OTP stereoprobe MB-6B and the azetidine acrylamide stereoprobe MY-11B (Figure 3D), showcasing an example of two classes of electrophilic compounds that target the same ligandable pocket through reaction with distinct nucleophilic residues.

**Figure 3.**
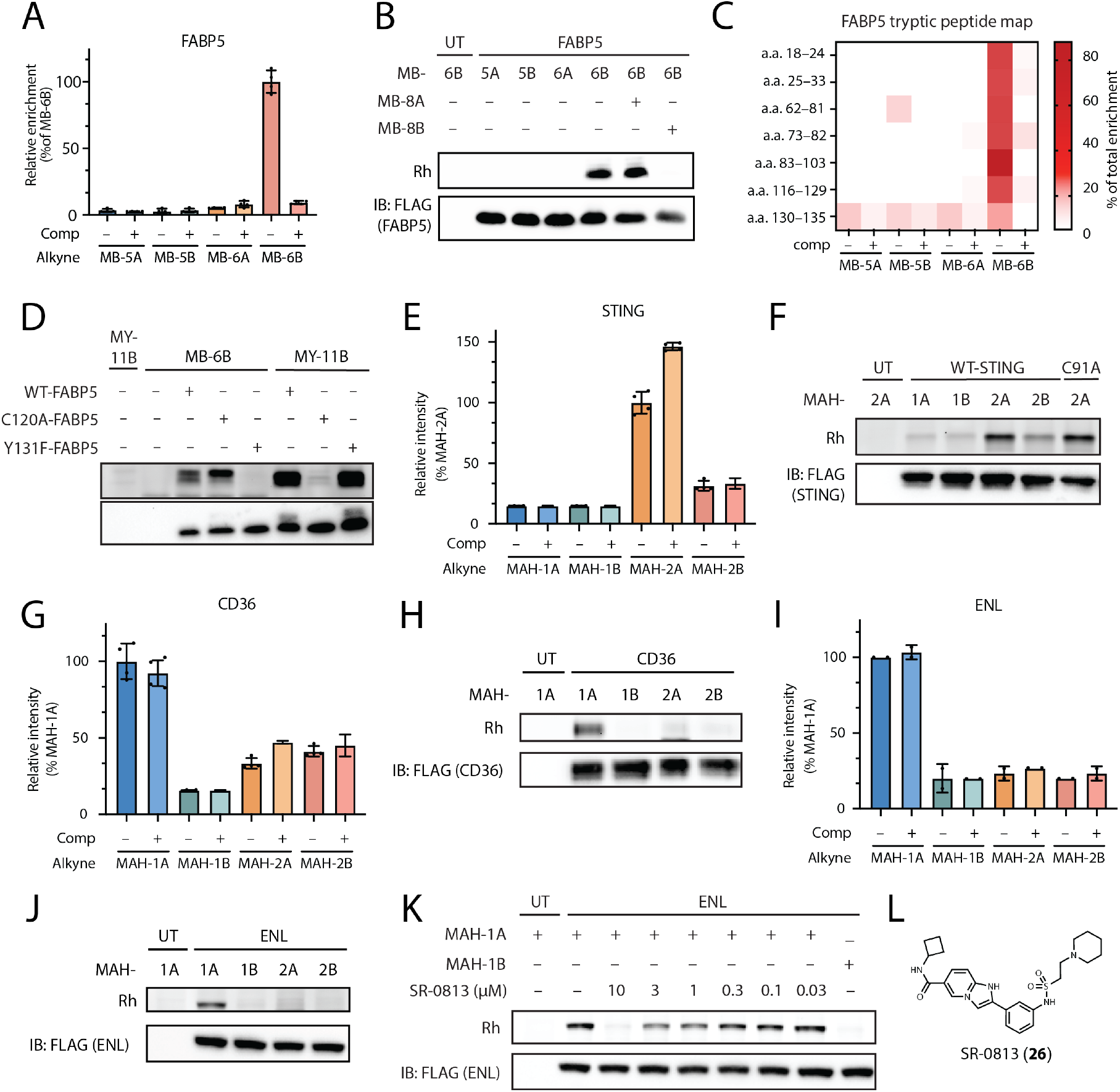
Characterization of OTP stereoprobe interactions with recombinant proteins. (A) Protein-directed ABPP data from Ramos cells showing stereoselective enrichment of FABP5 by MB-6B (20 µM, 1 h) and competition of this enrichment by MB-8B (20 µM, 2 h). (B) Gel-ABPP data confirming stereoselective engagement of recombinant FABP5 by MB-6B and blockade of this engagement by MB-8B, but not MB-8A. HEK293T cells recombinantly expressing FLAG epitope-tagged FABP5 were treated with the indicated competitor (20 µM, 2 h) or DMSO followed by the indicated alkyne stereoprobes (5 μM,1 h), after which gel-ABPP was performed as described in Figure 1D. (C) Tryptic peptide map for FABP5 from protein-directed ABPP experiments in Ramos cells showing stereoselective enrichment by MB-6B of all quantified peptides except for the Y131-containing peptide (a.a., 130-135). (D) Gel-ABPP data showing that the Y131F and C120A FABP5 mutants do not react with MY-6B (10 µM, 1 h) and MY-11B (1 µM, 1 h), respectively. Experiments were performed in HEK293T cells transiently expressing FLAG-tagged FABP5 variants. (E) Protein-directed ABPP data from OCI-AML-3 cells showing stereoselective enrichment of STING by MAH-2A (50 µM, 1 h). (F) Gel-ABPP data confirming stereoselective engagement of recombinant WT and C191A STING variants by MAH-2A (10 µM, 1 h) in transfected HEK293T cells. (G) Protein-directed ABPP data from OCI-AML-3 cells showing stereoselective enrichment of CD36 by MAH-1A (50 µM, 1 h). (H) Gel-ABPP data confirming stereoselective engagement of recombinant CD36 by MAH-1A (10 µM, 1 h) in transfected HEK293T cells. (I) Protein-directed ABPP data from OCI-AML-3 cells showing stereoselective enrichment of ENL by MAH-1A (50 µM, 1 h). (J) Gel-ABPP data confirming stereoselective engagement of recombinant ENL by MAH-1A (10 µM, 1 h) in transfected HEK293T cells and (K) the concentration-dependent blockade of this enrichment by the YEATS domain inhibitor SR-0813. (L) Structure of SR-0813. (A, C, E, G, I) Data are average values ± SD from one (I) or two (A, C, E, G) independent MS-ABPP experiments each performed with two technical replicates. (B, D, F, H, J, K) data shown are from a single replicate representative of two independent gel-ABPP experiments. IB: immunoblotting.

Among the OTP stereoprobe targets found exclusively in OCI-AML-3 cells were two innate immune proteins – i) STING (stimulator of interferon genes; or TMEM173), which is part of a signaling pathway that produces type I interferon in response to pathogenic DNA;^22^ and ii) CD36 (platelet glycoprotein 4), a monocyte-enriched integral membrane protein that binds diverse protein and lipid ligands and is implicated in immune-related and cardiovascular diseases.^23^ STING and CD36 were stereoselective enriched by MAH-2A and MAH-1A, respectively, but were not blocked in their enrichment by the corresponding competitors MAH-4A and MAH-3A (Figure 3E, F), indicating that these stereoprobe-protein interactions are specific, but low potency. Using gel-ABPP, we confirmed stereoselective engagement of recombinant STING and CD36 proteins in HEK293T cells by MAH-2A and MAH-1A, respectively (Figure 3G, H). While we do not yet know the residues that react with OTP stereoprobes in STING and CD36, we excluded C91 in STING – which is liganded by acrylamide stereoprobes^24a^ and other electrophilic compounds^24b^ – as the site of engagement for OTP stereoprobes by mutagenesis (Figure 3G). Also supporting that OTP stereoprobes engage a distinct site on STING was the preservation of stereoselective enrichment for the tryptic peptide containing C91 in our protein-directed ABPP experiments (Figure S5). Additionally, pretreatment with WX-03-60, a tryptoline acrylamide stereoprobe that we have found to engages STING_C91 by cysteine-directed ABPP,^24b^ did not block MAH-1A reactivity with recombinant STING, but did block the reactivity of an alkynylated analog of WX-03-60 (WX-03-341^7h^) (Figure S5).

Finally, we confirmed by gel-ABPP the stereoselective interaction between MAH-1A and the transcriptional coactivator protein ENL^25^ (or MLLT1) (Figure 3I, J). While MAH-1A reactivity with ENL was not blocked by the corresponding OTP stereoprobe competitor MAH-3A (Figure 3I), this interaction was disrupted in a concentration-dependent manner by SR-0813, a previously reported reversible inhibitor of the acetyl-lysine-binding YEATS domain of ENL^26^ (Figure 3K, L). We interpret these data to indicate that MAH-3A binds and reacts with a nucle-ophilic residue in the YEATS domain of ENL.

Taken together, our initial follow-up studies confirmed the veracity of diverse stereoselective enrichment and liganding events mapped for OTP stereoprobes by protein-directed ABPP, which appear to occur at both established functional (e.g., FABP5, ENL) and novel (e.g., STING) sites of small-molecule binding.

### Functional Characterization of OTP Stereoprobes Targeting the Transmembrane Protein TLCD1

Among the proteins engaged by the OTP stereoprobes, TLCD1 (TLC domain-containing protein 1) stood out as showing both strong stereoselective enrichment by MB-2A and robust blockade of this enrichment by pretreatment with MB-4A (Figure 4A, B). TLCD1 belongs to a family of transmembrane proteins that includes several paralogs in humans (TLCD2-5) and FLD1 in *C. elegans*. Previous studies have found that both TLCD1/2 and FLD1 regulate the fluidity of the plasma membrane by limiting the polyunsaturated acyl chain content of phospholipids^11^ and promoting monounsaturated fatty acid (MUFA) incorporation into phosphatidylethanolamine (PE) lipids.^12^ Additionally, mice genetically deleted for both TLCD1 and TLCD2 display attenuated development of non-alcoholic steatohepatitis.^12^ Using HepG2 cells that express TLCD1 under a tet-inducible promoter, we confirmed by gel-ABPP that recombinant TLCD1 stereoselectively reacted with MB-2A and this interaction was blocked in a stereoselective and concentration-dependent manner by MB-4A (Figure 4C, D). The low expression of recombinant TLCD1 in HepG2 cells required that we perform anti-FLAG epitope immunoprecipitation prior to gel-ABPP analysis. Attempts to measure an IC_50_ value for MB-2A with recombinant TLCD1 were complicated by impairments in anti-FLAG immunoprecipitation of the protein following engagement by MB-2A (see IB:FLAG (TLCD1) signals in MB-2A-treated samples; Figure 4C, D), but complementary protein-directed ABPP experiments of parental HepG2 cells confirmed that MB-4A stereoselectively blocked MB-2A enrichment of endogenous TLCD1 with an estimated half-maximal effect between 10-20 µM (Figure 4E). We also genetically disrupted TLCD1 expression in HepG2 cells using CRISPR/Cas9 methods and confirmed that these sgTLCD1 cells lost >90% of TLCD1 as determined by protein-directed ABPP (Figure 4E). We did not detect TLCD2 signals in any of our protein-directed ABPP experiments (Table S1), which could indicate that the OTP stereoprobes do not cross-react with this protein or that TLCD2 is poorly expressed in the cell lines examined herein.

**Figure 4.**
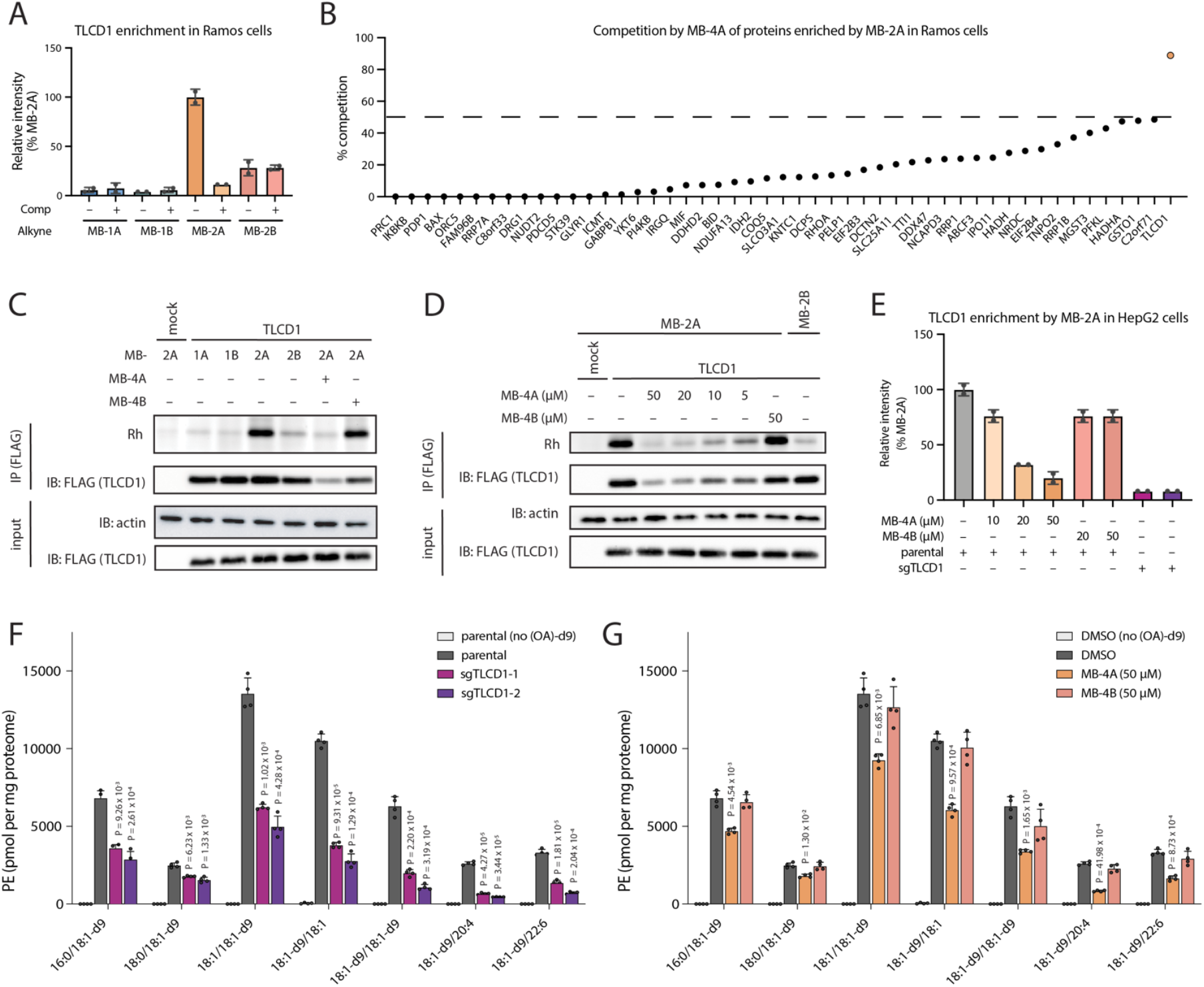
MB-4A stereoselectively engages and inhibits TLCD1. (A) Protein-directed ABPP data from Ramos cells showing stereoselective enrichment of TLCD1 by MB-2A (50 µM, 1 h) and competition of this enrichment by MB-4A (50 µM, 2 h). (B) Percentage competition by MB-4A (50 µM, 2 h) of proteins stereoselectively enriched by MB-2A (50 µM, 1 h) in Ramos cells. (C) Gel-ABPP data confirming stereoselective engagement of recombinant TLCD1 by MB-2A (5 µM, 1 h) in HepG2 cells and stereoselective blockade of this engagement by MB-4A (50 µM, 2 h). Gel-ABPP was performed after anti-FLAG immunoprecipitation to increase signal intensity for TLCD1. IB: immunoblotting.(D)Gel-ABPP data demonstrating stereoselective and concentration-dependent blockade of MB-2A–TLCD1 interactions by MB-4A (2 h) in HepG2 cells. (E) Protein-directed ABPP data from parental or sgTLCD1 HepG2 cells demonstrating stereoselective and concentration-dependent blockade of MB-2A enrichment of TLCD1 by MB-4A, as well as loss of TLCD1 enrichment in sgTLCD1 cells. HepG2 cells were treated with MB-4A/4B or DMSO for 2 h, followed by treatment with MB-2A (50 µM, 1 h). (F) LC-MS data showing decreased incorporation of 18:1-d9 (OA-d9) free fatty acid (FFA) into phosphatidylethanolamine (PE) lipids in parental or sgTLCD1 HepG2 cells. (G) LC-MS data showing decreased incorporation of 18:1-d9 FFA into PE lipids in parental HepG2 cells treated with MB-4A, but not MB-4B (50 µM, 2 h). (A, B, E) Data are average values ± SD from one (E) or two (A, B) independent MS-ABPP experiments, each with two technical replicates. (C, D) data shown are from a single replicate representative of two independent gel-ABPP experiments. (F, G) data shown are from a single experiment, with four technical replicates, representative of two independent metabolomics experiments. Data represent mean values ± s.e.m.; P values were derived using a two-sided Student’s t-test performed relative to parental HepG2 cells treated with nothing (F) or DMSO (G).

Considering that the combined genetic disruption of TLCD1 and TLCD2 in mice or primary mouse hepatocytes has been shown to decrease the MUFA content of PE lipids, ^12^ we next conducted isotopic tracing experiments using deuterated C18:1 oleic acid ((OA)-d9). We first confirmed that sgTLCD1 HepG2 cells displayed a substantial reduction in the incorporation of (OA)-d9 into a range of PE lipids in comparison to parental HepG2 cells—with PE(18:1-d9/20:4) undergoing the most dramatic decrease in abundance (Figure 4F). We next treated parental HepG2 cells with MB-4A or MB-4B (50 µM, 2 h) and observed a significant decrease in formation of (OA)-d9-containing PE species exclusively in the MB-4A-treated cells (Figure 4G). As was observed in the sgTLCD1 cells, MB-4A treatment had the most profound effect on PE(18:1-d9/20:4).

We interpreted the similar effect of genetic deletion of TLCD1 and MB-4A treatment on the MUFA content of PE lipids in HepG2 cells to indicate that MB-4A acts as an inhibitor of TLCD1. To better understand how MB-4A might impair TLCD1 function, we engaged in a complementary line of inquiry using AF2-derived models^27^ of TLCD1 to interrogate structural databases for evolutionary insights into its biochemical function—drawing on FoldSeek^28^and Dali^29^ for fast and sensitive fold comparisons and to delineate conserved residues. A structure-based dendrogram showed that TLCD1 belongs to an expansive enzyme family generally involved in regulating the lipid composition of membranes (Figure 5A). The TLCDs comprise a group of six closely related paralogs and CLN8—an ER membrane protein genetically associated with the lysosomal storage disorder neuronal ceroid lipofuscinosis 8^30^—that branches off a family of ceramide synthases (CERSs) mechanistically anchored by high-resolution complexes of human and yeast enzymes with substrates and inhibitors.^13,31^ Diverging from the CERS branch is a clan of three TRAM proteins that function in ER protein import, both as cargo receptors and lipid modulators to the Sec61 channel complex.^32^ The TLCDs and CERSs are in turn related to the ELOVL family of elongases for very long chain fatty acids^14^ and sibling TACAN/TMEM120 lipid membrane remodelers associated with ion channel and mechanosensor complexes.^33^

**Figure 5.**
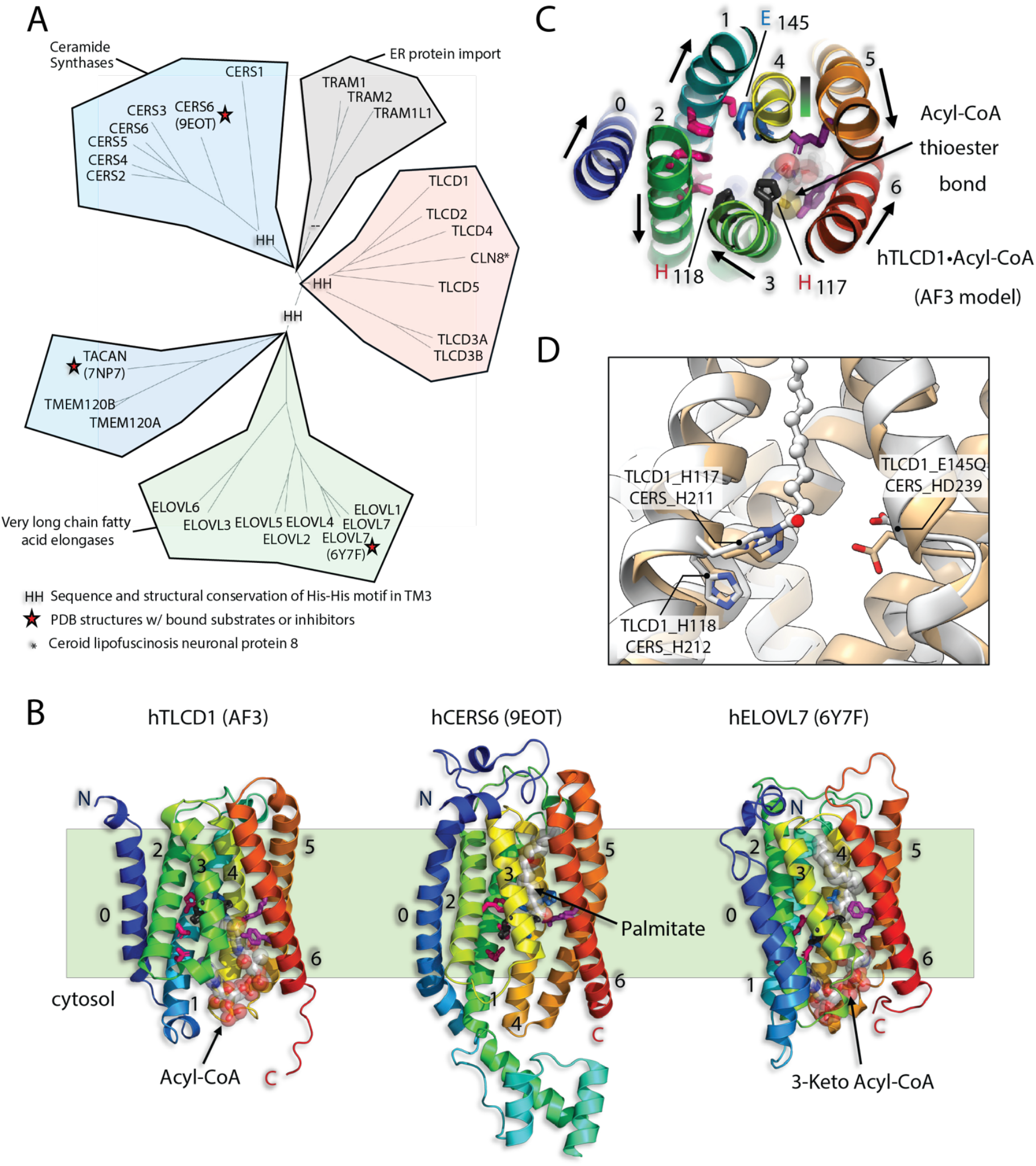
Structural modeling of TLCD1 and related proteins. (A) An unrooted structural similarity dendrogram, derived by Dali comparisons of AF2 database models, shows TLCDs clustering with CLN8, close to the ceramide synthase (CERS) branch and divergent TRAM proteins, offset from the very long chain fatty acid elongases (ELOV) and TACAN/TMEM120 families. HH motifs mark the conservation of predicted (or experimentally defined) catalytic His residues across all branches except for the divergent TRAMs. Red stars indicate PDB structures that illuminate aspects of the lipid binding and catalytic mechanism of this enzyme superfamily. (B) An AF3-derived acyl-CoA-bound model of human TLCD1 is posed alongside the superposed structures of human CERS6 and ELOVL7, respectively bound to palmitate and 3-keto acyl-CoA substrates. TM helices are numbered 1–6, with key residues involved in catalysis shown in stick form—the signature His-His motif in black. Protein chains are gradient-colored N-terminal blue to C-terminal red with Pymol (www.pymol.org). (C) A luminal view of the catalytic constellation in the hydrophobic tunnel of TLCD1 shows the spatial positioning of the predicted catalytic nucleophile H117 alongside the adjacent conserved H118 and opposite E145 over the labile acyl-CoA thioester bond. (D) Structural comparison of TLCD1 (AF-Q96CP7-F1-model_v4, light tan) to CERS6-C16:0 (PDB: 8QZ6, white) with homologous putative catalytic residues marked: TLCD1_H117/CERS6_H211, TLCD1_H118/CERS6_H212, and TLCD1_E145Q/ CERS6_D239.

All branches of this superfamily display a signature His-His motif in TM3 of a core 6TM fold (H117-H118 in TLCD1), spatially juxtaposed to a conserved Glu/Asp in TM4 (E145 in TLCD1) within a hydrophobic tunnel with side access to membrane lipids (Figure 5B-D)—save the divergent TRAMs that rely on a related but spatially distinct catalytic constellation. Interestingly, structure/function studies of both CERS6 and ELOVL7 support their reaction with fatty acyl coenzyme A (CoA) substrates to form acyl-enzyme intermediates with a conserved histidine residue as part of a catalytic mechanism that, for CERS6, involves transfer of the fatty acyl-histidine intermediate to an acceptor sphingoid base to form ceramides.^13,31C,34^ The conservation of the His-His dyad in TLCDs suggests that these proteins are also enzymes engaged in a fatty acyl (MUFA) transfer reaction with H117 as the predicted nucleophile, where the lipid acceptor species would be lyso-PE leading to the formation of MUFA PE lipids. Consistent with this premise, AF3 modeling^35^ of both MUFA-CoA and lyso-PE lipids with TLCD1 revealed a CERS-like docking of the acyl-CoA to the cytosolic vestibule of the hydrophobic tunnel, positioning the thioester bond for nucleophilic attack by H117, setting up an acyl transfer reaction to a lyso-PE lipid bound to the luminal end of the tunnel (Figure S6).

We treated dox-inducible HepG2 cell line models expressing WT-TLCD1 or H117N, H118N, and E145Q mutants with OTP stereoprobes and found that only WT-TLCD1 displayed robust stereoselective reactivity with MB-2A (Figure 6A). Having verified the importance of H117, H118, and E145 for TLCD1 reactivity with OTP stereoprobes, we next evaluated the H117N, H118N, and E145Q mutants in isotopic tracing experiments of (OA)-d9 incorporation into PE lipids in cells. We first found that reintroduction of WT-TLCD1 into sgTLCD1 HepG2 cells fully restored the incorporation of (OA)-d9 into PE(18:1-d9/20:4) (Figure 6B). In contrast, the H117N, H118N, and E145Q TLCD1 mutants were unable to rescue (OA)-d9 incorporation into PE(18:1-d9/20:4) in sgTLCD1 cells (Figure 6B).

**Figure 6.**
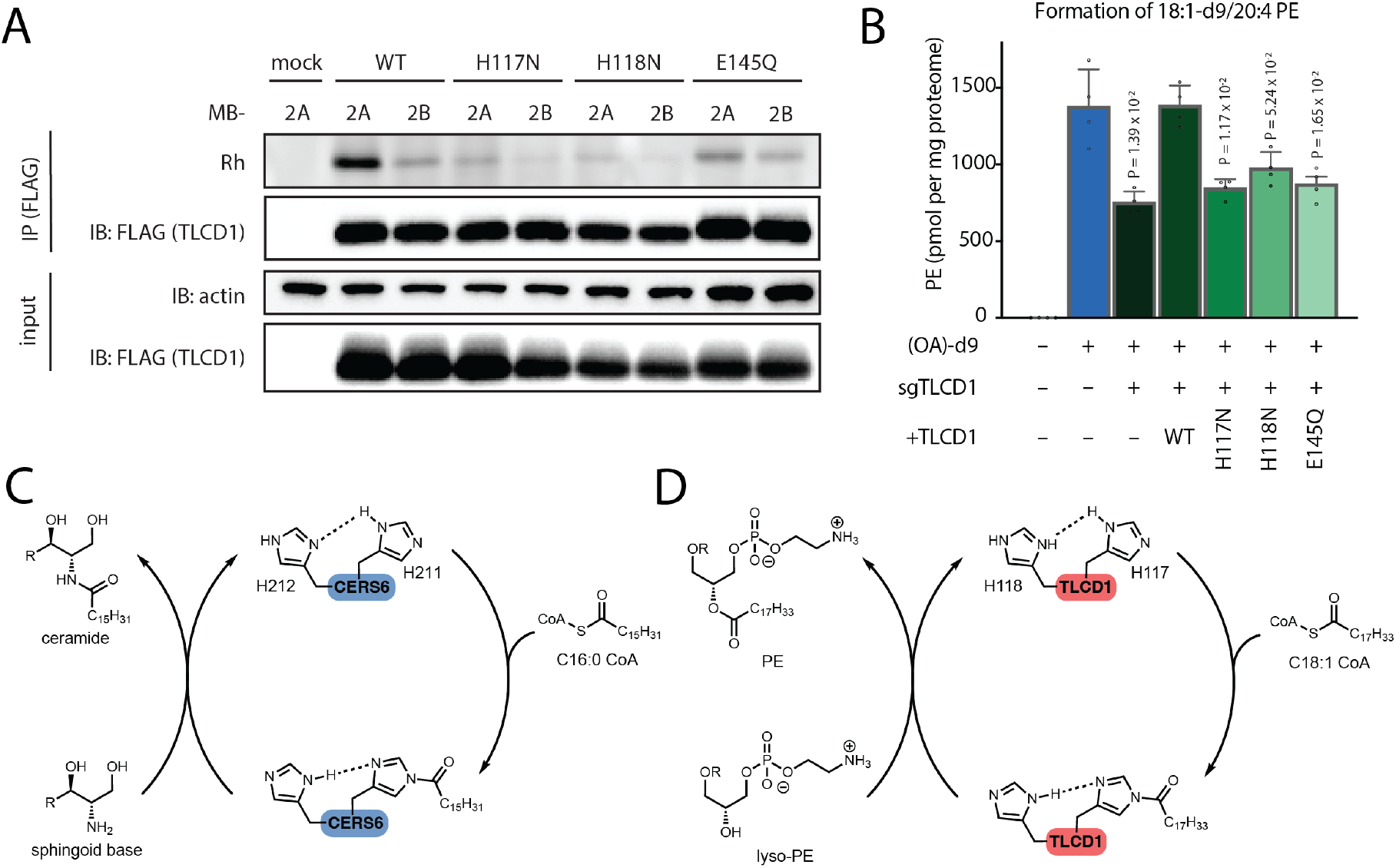
Conserved H117, H118, and E145 residues are required for the OTP stereoprobe engagement and lipid remodeling activity of TLCD1. (A) Gel-ABPP data demonstrating decreased engagement of H117N, H118N, and E145Q TLCD1 by MB-2A (10 µM, 1h) in comparison to WT TLCD1. TLCD1 proteins were expressed in HepG2 under a tet-inducible promoter. (B) LC-MS data showing that WT, but not H117N, H118N, or E145Q TLCD1 variants rescued incorporation of 18:1-d9 FFA (OA-d9) into 18:1-d9/20:4 PE species in sgTLCD1 HepG2 cells. (C) Mechanistic model of CERS6-catalyzed ceramide formation involving reaction with a fatty acyl-CoA substrate to form a covalent acyl-H211 adduct, followed by transfer of the fatty acyl group to a sphingoid base acceptor to form ceramide. (D) Proposed mechanism for TLCD1-catalyzed acyltransferase activity involving a preferential acceptance of MUFA-CoA substrates (e.g., C18:1 CoA) that react to form a covalent acyl-H117 adduct and are then transferred to lyso-PE as the acceptor co-substrate to form PE lipids. (A, B) Data are from a single experiment representative of two independent experiments involving one (A) or four (B) technical replicates. Dox concentrations: HepG2 cells with tet-inducible promoter for WT and H117N TLCD1 induced with 125 ng/mL, for H118N TLCD1 induced with 1000 ng/mL, and for E145Q TLCD1 induced with 33 ng/mL. Data represent mean values ± s.e.m.; P values were derived using a two-sided Student’s t-test performed relative to WT cells with no dox induction

Taken together, our chemical proteomic, AF modeling, lipid metabolic labeling, and mutagenesis data point to i) TLCD1 being an enzyme that transfers the acyl chain from MUFA-CoAs to lyso-PE to form MUFA PE lipids in cells via a catalytic mechanism involving a histidine (H117) nucleophile that is shared with ceramide synthase and elongase enzymes (Figure 6C, D); and ii) this acyltransferase activity being inhibited by OTP stereoprobes through covalent modification of one of the conserved catalytic residues in TLCD1.

## Discussion

The use of stereochemically defined electrophilic or photoreactive small molecules (‘stereoprobes’) in combination with ABPP has accelerated the assignment of ligandable sites on diverse proteins in human cells.^7,9,36^ So far, only a limited diversity of reactive groups has been explored with stereoprobes (e.g., acrylamides, sulfonyl fluorides, diazirines), and our protein-directed ABPP studies with the OTP electrophile support that it engages a distinct set of proteins that shares some commonality with other P(V) electrophiles (e.g., serine hydrolases that also crossreact with fluorophosphonates) in addition to proteins that have not, to our knowledge, been liganded in previous chemical proteomic studies. We also found that two sets of azetidine OTPs differed substantially in their stereoselective protein interactions, emphasizing the importance of recognition groups appended to the stereoprobe core. We believe our synthetic routes for both the azetidine and tryptoline OTPs should be compatible with generating structurally diverse sub-libraries of these stereoprobes for further chemical proteomic studies. Future goals would include both expanding the scope of proteins that can be liganded by OTP stereoprobes and improving the potency of individual OTP stereoprobe-protein interactions. Indeed, we note that many of the proteins that were stereoselectively enriched by alkynylated OTP stereoprobes in protein-directed ABPP experiments were not blocked in their enrichment by non-alkyne competitors, which we interpret to reflect mostly specific, but low potency interactions. It will also be important to develop MS protocols capable of mapping the amino acid residues that are modified by OTP stereoprobes in chemical proteomic experiments, especially since our limited mutagenesis studies are supportive of previous findings that the OTP electrophile can react with diverse amino acids.^10^ In the case of FABP5, we discovered that OTP and acrylamide stereoprobes react with orthogonal residues in the lipid-binding pocket of this protein (Y131 and C120, respectively).

The OTP-interacting proteins included the integral membrane protein TLCD1 that regulates acyl chain composition of PE lipids in cells.^11,12^ Our combined data leveraging genetic disruption systems, AF2/AF3 structural modeling, mutagenesis, and targeted lipidomic experiments support that TLCD1 is a lipid metabolic enzyme mechanistically related to the CERS and ELOVL families. Notably, one of the catalytic histidines in this enzyme superfamily has been found in both CERS6 and ELOVL7 to form an acyl-enzyme intermediate with fatty acid substrates^13,14b^ that is analogous to the acyl-enzyme intermediate formed with the catalytic serine in serine hydrolases.^37^ Considering further that the catalytic serine of serine hydrolases is the site of covalent modification of P(V) electrophiles like fluorophosphonates,^38^ we hypothesize that the corresponding putative catalytic histidine in TLCD1 (H117) may be the site of reactivity with OTP stereoprobes. We cannot, however, exclude that the OTP stereoprobes react elsewhere on TLCD1. Regardless, our data support that OTP stereoprobes act as inhibitors of TLCD1 that largely mirror the lipid metabolic effects of genetic disruption of TLCD1 in human cells. What enzymatic reaction might TLCD1 perform in cells to promote MUFA incorporation into PE lipids? The simplest model based on the catalytic activity of CERSs is that TLCD1 functions as an acyltransferase preferentially reacting with MUFA-CoAs to form a histidine-acyl enzyme intermediate and, in a subsequent catalytic step, transfers the bound MUFA to lyso-PE acceptor substrates to form MUFA-PEs (Figure 6C). Our attempts so far, however, to establish a MUFA acyltransferase activity with lysates from cells recombinantly expressing TLCD1 have been unsuccessful, which may suggest that additional cellular factors are required to support TLCD1 enzymatic activity. Future structural studies of TLCD1 may also help to clarify more details on its catalytic mechanism and the mode of inhibition for OTP stereoprobes. If confirmed to react with the catalytic histidine in TLCD1, OTP compounds would, in principle, have the potential to act as a general class of covalent inhibitors for not only other TLCDs, but also CERS and ELOVL enzymes. In this scenario, ABPP experiments performed with a larger and more structurally diverse library of OTPs may identify covalent inhibitors of additional members of the TLCD/CERS/ELOVL enzyme families.

In conclusion, through synthesizing and investigating the proteomic reactivity of stereoprobes bearing an OTP electrophile, we have discovered stereoselective covalent ligands for a wide array of proteins involved in, for instance, innate immunity (STING, CD36), transcriptional regulation (ENL), and lipid metabolism (TLCD1). As the potency, selectivity, and drug-like properties of such OTP ligands are improved, we believe they will hold great promise as chemical probes for diverse biological applications.

## Supporting information

Supplementary Biology Information

Supplementary Chemistry Information

## ASSOCIATED CONTENT

The Supporting Information is available free of charge at xxxx. Detailed biological experimental procedures and analytical data (PDF) Detailed chemical experimental procedures and analytical data (PDF) Protein-directed ABPP of stereoisomeric OTP probes (XLSX)

MS-ABPP data are available via ProteomeXchange^39^ with identifier PXD060339.

## AUTHOR INFORMATION

**Authors:**

**Hayden A. Sharma** – Department of Chemistry, Scripps Research, La Jolla, California, 92037, United States;

**Michael Bielecki**– Department of Chemistry, Scripps Research, La Jolla, California, 92037, United States;

**Meredith A. Holm** – Department of Chemistry, Scripps Research, La Jolla, California, 92037, United States;

**Ty M. Thompson** – Department of Chemistry, Scripps Research, La Jolla, California, 92037, United States;

**Yue Yin** – Department of Chemistry, Scripps Research, La Jolla, California, 92037, United States;

**Jacob B. Cravatt** – Department of Chemistry, Scripps Research, La Jolla, California, 92037, United States;

**Timothy B. Ware** – Department of Chemistry, Scripps Research, La Jolla, California, 92037, United States;

**Alex Reed** – Department of Chemistry, Scripps Research, La Jolla, California, 92037, United States;

**Molham Nassir –** Department of Chemistry, Scripps Research, La Jolla, California, 92037, United States;

**Tamara El-Hayek Ewing –** Department of Chemistry, Scripps Research, La Jolla, California, 92037, United States;

**Bruno Melillo –** Department of Chemistry, Scripps Research, La Jolla, California, 92037, United States;

**Fernando Bazan –** ħ Bioconsulting, LLC, Stillwater, MN 55082, United States;

## ACKNOWLEDGMENT

This work was supported by Bridge Biotherapeutics and the NIH (GM-118176 for P. S. B. and R35CA231991 for B. F. C). M. N. thanks the Council for Higher Education and Fulbright Israel. We are grateful to Dr. G. J. Kroon and Dr. L. Pasternack (Scripps Research) for NMR spectroscopic assistance; J. B. Brenneman, J. Jin, L. Revollo, S. Paudyal, and C. Kim (Bridge Biotherapeutics) and W. Goetzke (Scripps Research) for helpful discussions; and S. Barbas (Scripps Research) for technical assistance.

