## Supplementary Biology Information for "Proteomic Ligandability Maps of Phosphorus(V) Stereoprobes Identify Covalent TLCD1 Inhibitors"

### Biological Supporting Information

**12. Metabolomics data………………………………………………………………………………...........15**
**13. Uncropped gels and blots……………………………………………………………………………...17**

#### Supplementary Figures

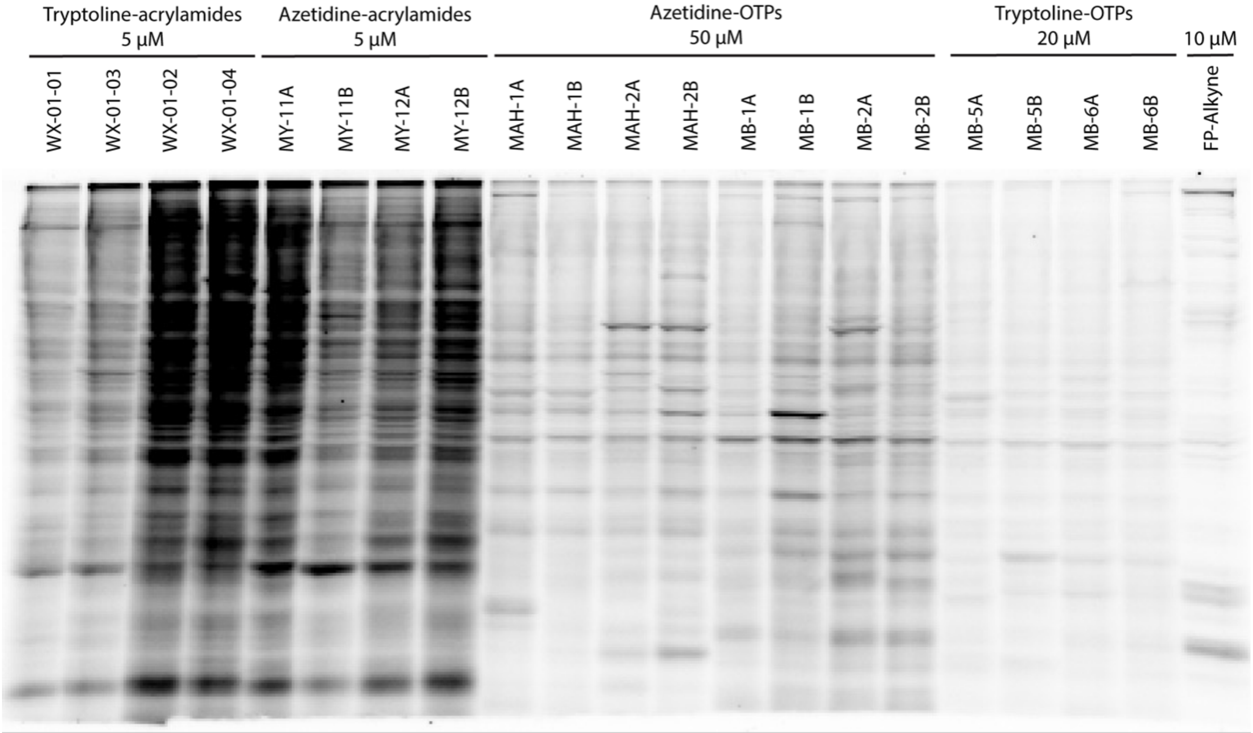

**Figure S1**. Gel-ABPP data comparing the proteomic reactivity of acrylamide- and OTP-stereoprobes, as well as the FP-alkyne probe, in Ramos cells. Cells were treated with the indicated concentrations of alkyne probes, followed by lysis and gel-ABPP analysis as summarized in the legend to Figure 1. Azetidine and tryptoline acrylamide stereoprobes are described in ref. 7h and ref 7b, respectively. Data are from a single experiment representative of multiple experiments. Samples run on 10% SDS page gels.

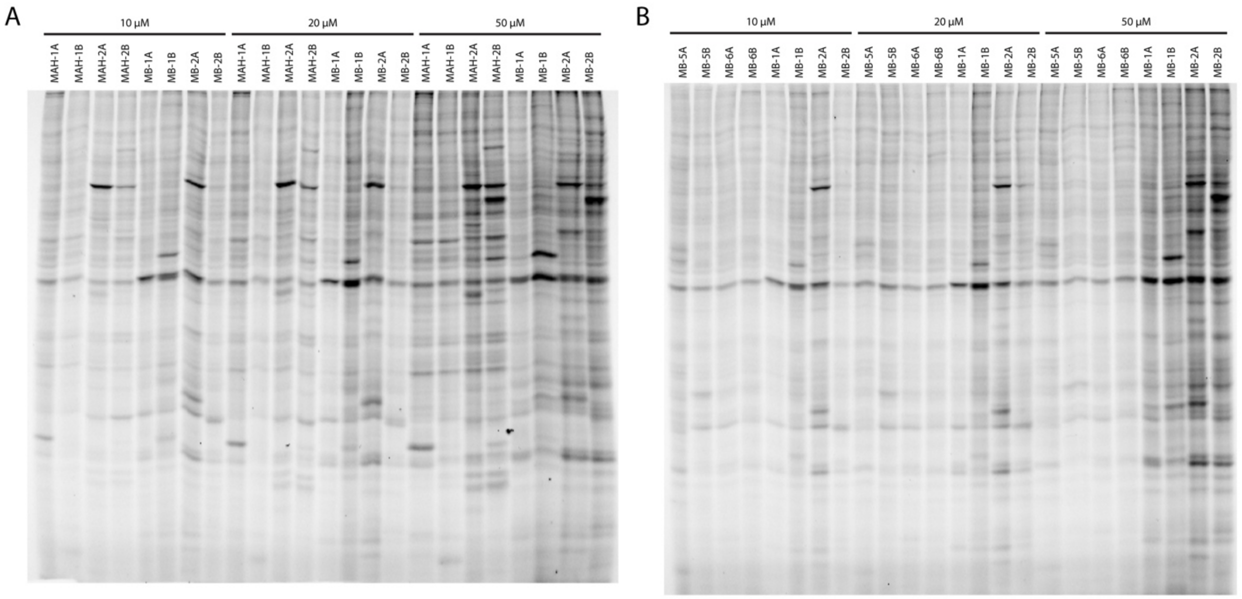

**Figure S2**. Gel-ABPP data comparing the concentration-dependent proteomic reactivity of azetidine (A) and tryptoline (B) OTP stereoprobes in Ramos cells. Data are from a single experiment representative of multiple experiments. Samples run on 10% SDS page gels.

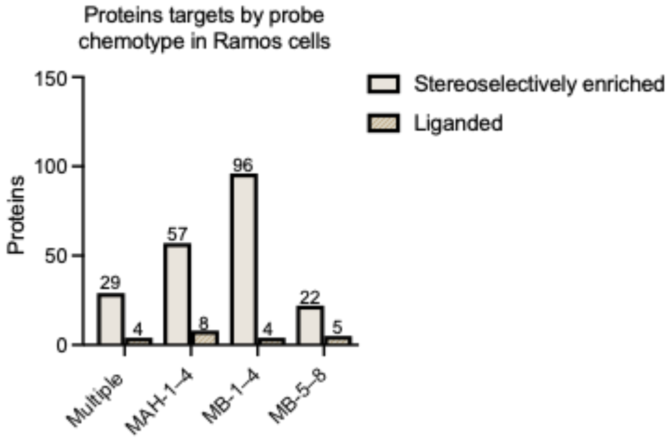

**Figure S3.** Total numbers and distributions of proteins that were stereoselectively enriched (lighter tan) and liganded (darker tan) by OTP-stereoprobes in Ramos cells. A protein was deemed stereoselectively enriched if it was enriched with >2.5-fold enantioselectivity; a stereoselectively enriched protein was considered "shared" between chemotypes if it was enriched with >2.5-fold enantioselectivity by one stereoprobe chemotype and >2-fold enantioselectivity by the other stereoprobe chemotype. A liganded protein was considered "shared" if it was stereoselectively enriched and competed by >50% by one stereoprobe chemotype and stereoselectively enriched by an alternate chemotype with >25% competition. All other stereoselectively enriched or liganded targets of the OTP-stereoprobes were considered selective for an individual OTP stereoprobe chemotype. Data are from two independent protein-directed ABPP experiments each performed with two technical replicates.

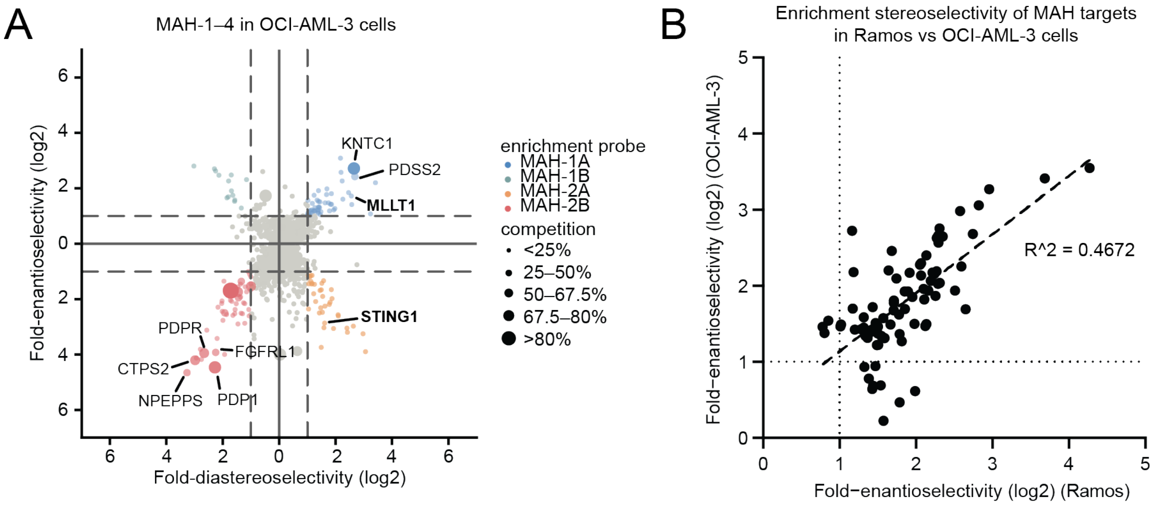

**Figure S4.** Protein-directed ABPP data for OCI-AML-3 cells treated with the MAH-1-4 OTP stereoprobes. (A) OCI-AML3 ells were treated with competitor stereoprobes (50 µM MAH-3/4) or DMSO for 2 h followed by the corresponding alkyne stereoprobes (50 µM MAH-1/2) and then processed as summarized in Figure 2A. Quadrant plot highlighting enantio- and diastereo-selectively enriched proteins for each stereoconfiguration of alkyne OTP stereoprobes MAH-1-4 in OCI-AML-3 cells. For each protein, enrichment enantioselectivity and diastereoselectivity (log2) are shown for the probe leading to the highest enrichment. Enantioselectivity is defined as the ratio of enrichment for one stereoisomer versus its enantiomer, and diastereoselectivity as the ratio of enrichment of one stereoisomer versus the average of its two diastereomers. A protein is shown in color if both the enantioselectivity and diastereoselectivity of enrichment (log2) are greater than 1. Competition is represented by dot size, with the largest circle indicating >80% competition, the medium large circle indicating 67.5–80% competition, the medium circle indicating 50–67.5% competition, the medium small circle indicating 25–50% competition, and the smallest circle indicating <25% competition. Data are from two independent protein-directed ABPP experiments each performed with two technical replicates. (B) Comparison of protein-directed ABPP data for MAH1-2 in Ramos and OCI-AML-3 cells. For each protein, enantioselective enrichment values (log2(ER)) are shown for the probe leading to the highest enrichment.

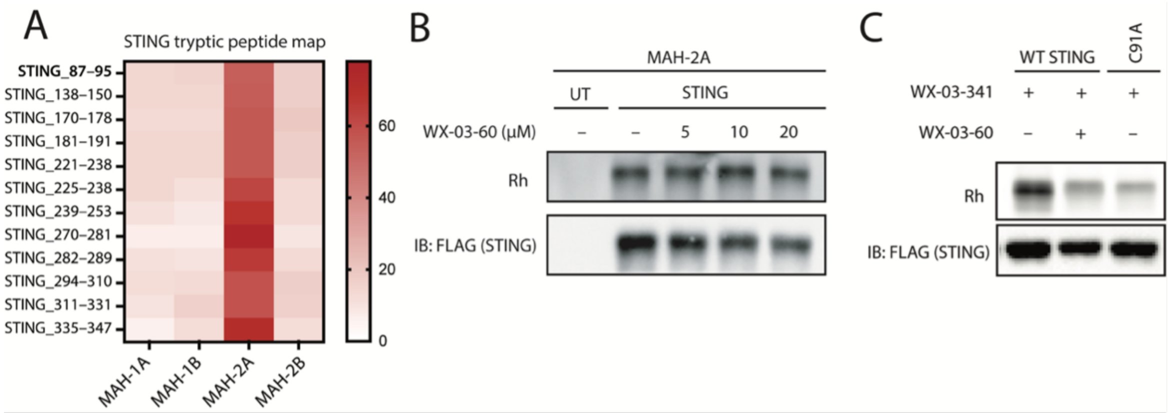

**Figure S5.** Characterization of STING-stereoprobe interactions. (A) Tryptic peptide map for STING from protein-directed ABPP experiments showing stereoselective enrichment of the tryptic peptide containing C91 (a.a. 87-95) in OCI-AML3 cells treated with MAH-2A (20 µM, 3 h) alongside other stereoselectively enriched peptides. (B) Gel-ABPP data showing that pretreatment with tryptoline acrylamide stereoprobe WX-03-60 (5–20 µM, 2h) does not block engagement of recombinant STING by OTP-stereoprobes (10 µM, 1 h). (C) Gel-ABPP data showing that pretreatment with WX-03-60 (20 µM, 2 h) and mutation of C91 to alanine (C91A) block engagement of STING by alkyne tryptoline acrylamide stereoprobe WX-03-341 (5 µM, 1 h). For (B, C), experiments were performed in transiently transfected HEK293T cells (UT = untransfected cells); shown is a single replicate from experiments performed in duplicate.

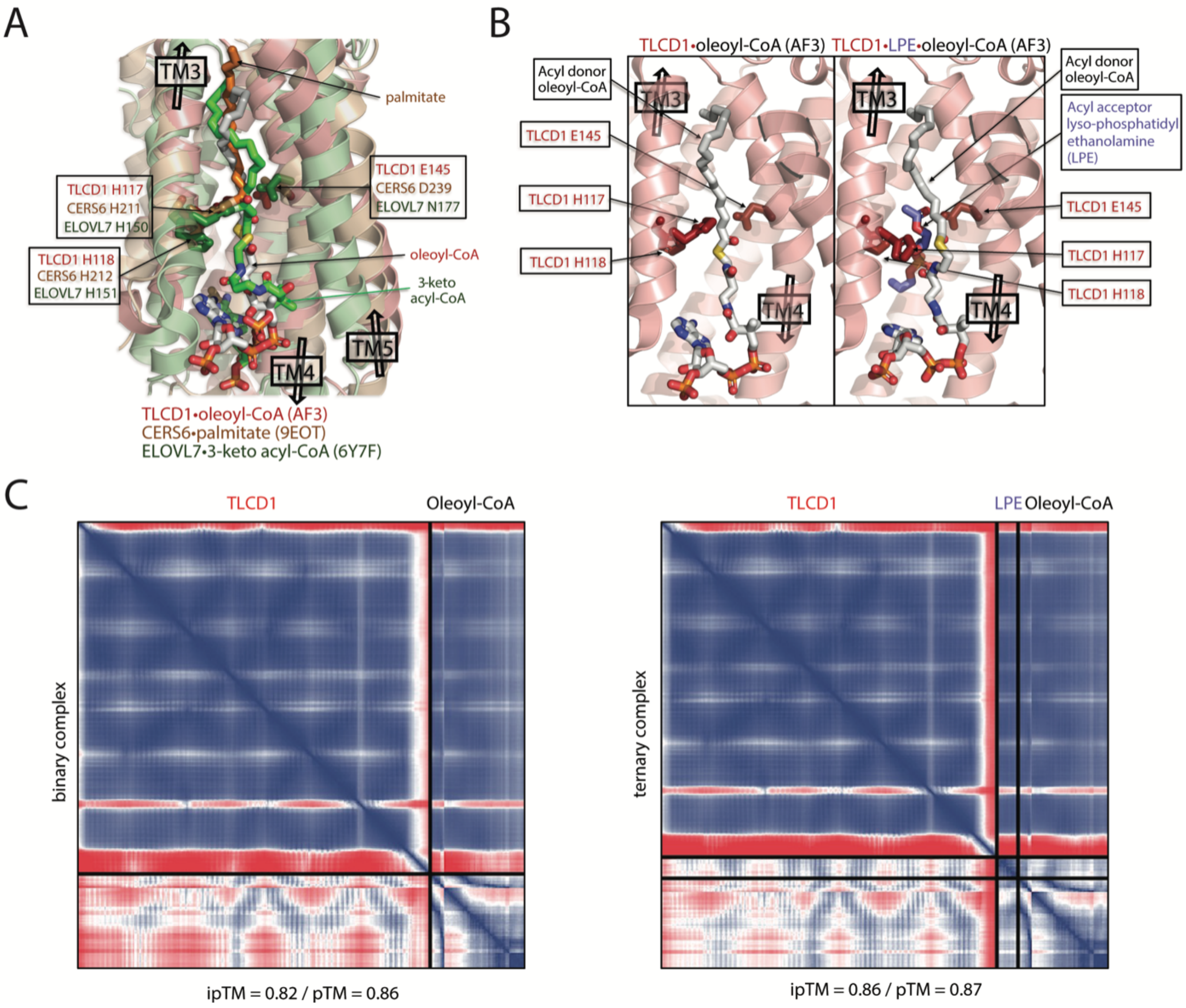

**Figure S6.** Structural analysis of computationally predicted TLCD1-lipid interactions. A) Comparison of catalytic residue constellations for substrate-bound CERS6 and ELOVL7, respectively drawn from PDB structures 9EOT and 6Y7F), with TLCD1 bound to (monounsaturated C18:1) acyl donor substrate oleoyl-CoA. The latter complex structure is derived from AF3 (https://alphafoldserver.com), taking advantage of a brief period of open-access testing by DeepMind of Chemical Component Dictionary or CCD-coded ligand entities for protein multimer predictions (see https://www.wwpdb.org/data/ccd); in the present case, the CCD code for oleoyl-CoA is 3VV. Omitting TM6 for clarity in all three structures, the TLCD cartoon helices are colored salmon, with dark red stick residues, bound to light grey oleoyl-CoA, superposed on CERS6 (wheat helices, brown residues, orange palmitate ligand) and ELOVL7 (light green helices, green residues, green 3-keto acyl-CoA ligand). The view into the catalytic cavity emphasizes the superposition of nucleophilic His residues (the first His in the 'HH' dyad, H117-H118 in TLCD1, H211-H212 in CERS6, and H150-H151 in ELOVL7), juxtaposed with a conserved acidic residue in TLCD1 and CERS6 (E145 and D239, respectively), and N177 in ELOVL7. The acyl-CoA part of the bound fatty acid acyl-CoA chain in TLCD1 and CERS6 docks into the cytosolic vestibule of the hydrophobic tunnel that leads to the centrally located active site. B) Side-by-side comparison of TLCD1 bound to substrates in binary form, with the oleoyl-CoA acyl donor in the left panel, or in a ternary complex in the right panel, with oleoyl-CoA acyl donor alongside the lyso-phosphatidylethanolamine or LPE acyl acceptor (where this latter substrate molecule has an official CCD code of LSP, for recognition by AF3). The same catalytic machinery from (A) is marked accordingly, drawing attention to the subtle displacement or distortion of the acyl chain in the donor oleoyl-CoA, when co-bound with LPE that is positioned in a side cavity; in addition, the sidechain position of the nucleophilic H117 rotates outward, concomitantly with oleoyl-CoA. C) The Predicted Aligned Error or PAE plots of the AF3 multimer models in (B), are shown side-by-side. The dark blue areas in the 2D graphs are a measure of high confidence in the positioning of paired residues in the model. The pTM and ipTM scores are AF3 multimer-provided metrics that reflect the confidence in the overall conformation of the complex, and the accuracy or fit of the predicted interface of molecules in the multimer, respectively. Scores above 0.8 for both metrics indicate high confidence.

**Reagents and materials.**

| ABPP reagents | | |
|---|---|---|
| **Reagents** | **Source** | **Catalog number** |
| Biotin-PEG4-azide | BroadPharm | BP-22119 |
| Urea | Fisher Scientific | M1084871000 |
| Iodoacetamide | Sigma-Aldrich | I1149-25G |
| Dithiothreitol (DTT) | Fisher Bioreagents | BP172-25 |
| Sequencing Grade Modified Trypsin | Promega | V5111 |
| Streptavidin Agarose Resin | Fisher Scientific | 20349 |
| Micro Bio-Spin column | Bio-Rad | 7326204 |
| NonidetTM P40 substitute (IgepalTM CA-630) | USB Corporation | 19628 |
| Triethylammonium bicarbonate buffer | Sigma-Aldrich | T7408-500ML |
| TMT10plex™ Isobaric Label Reagent Set | ThermoFisher Scientific | 90406 |
| TMTpro™ 16plex Label Reagent Set | ThermoFisher Scientific | A44520 |
| Ammonium bicarbonate | Sigma-Aldrich | 09830-500G |
| Hydroxylamine solution | Sigma-Aldrich | 467804-10ML |
| Formic acid, ~98%, for mass spectrometry | Honeywell Fluka | 94318-250ML-F |
| methanol | Fisher Scientific | A452SK-4 |
| Acetonitrile LC-MS Grade | Fisher Scientific | A955 |
| Water LC-MS Grade | Fisher Scientific | W6-4 |
| Protein LoBind Tubes 1.5 mL | Fisher Scientific | E925000090 |
| DMSO | Corning | 25-950-CQC |
| EDTA (0.5M, pH 8.0) | Invitrogen | AM9260G |

**Cell lines.** All cell lines tested negative for mycoplasma contamination. Cells were maintained at 37 °C with 5% CO2. Adherent cells were washed with Dulbecco's PBS (DPBS, Corning, 21-031-CV) and detached with trypsin-EDTA (0.05% or 0.25%, Gibco) for cell subculturing or cell-based assays. HEK293T (American Type Culture Collection (ATCC), CRL-3216) and HepG2 (ATCC, HB-8065) cells were cultured in DMEM (Corning, 15-013-CV) supplemented with 10% (v/v) FBS (Omega Scientific), penicillin (100 U mL−1), streptomycin (100 µg mL−1) and l-glutamine (2 mM). Ramos (ATCC, CRL-1596) and OCI-AML-3 (DSMZ, ACC-582) cells were grown in Roswell Park Memorial Institute medium (RPMI) supplemented with 10% (v/v) FBS (Omega Scientific), penicillin (100 U mL−1), streptomycin (100 µg mL−1) and l-glutamine (2 mM).

#### Gel-ABPP for proteome-wide reactivity

Ramos (1 mL of 3 million cells $mL^{-1}$) cells were treated with 10, 20, or 50 µM alkyne probes for 1 h or 3 h. The cells were collected and washed three times with chilled Dulbecco's phosphate-buffered saline (DPBS). The cell pellets were resuspended in 120 µl of cold DPBS and lysed by pulse sonication (1 × 15 pulses, 10% power output). The total protein content of the whole-cell lysates was measured using a Pierce

BCA protein assay kit. Samples were normalized to 1–2 mg mL$^{-1}$ and an 100µl sample was treated with 12 µl of click mix (45 µl of 1.7 mM tris((1-benzyl-4-triazolyl)methyl) amine (TBTA) in 4:1 *t*-BuOH:DMSO, 15 µl of 50 mM $CuSO_4$ in $H_2O$, 15 µl of 1.25 mM rhodamine–polyethylene glycol–azide in DMSO, 15 µl of freshly prepared 50 mM tris(2-carboxyethyl)phosphine in DPBS) for 1 h at RT with vigorous vortexing every 20 min. The click reaction was quenched by the addition of 36 µl of 4× SDS gel loading buffer, and the samples were resolved on 10% SDS–PAGE and imaged by in-gel fluorescent scanning using a BioRad imager with Image Lab software version 6.1. Gels were stained with Instant Blue and visualized by Coomassie scanning using the aforementioned imager and software.

#### Protein-directed ABPP

**In situ treatment and sample processing.** Protein-directed ABPP was carried out as previously reported (Njomen et, al.). Ramos, OCI-AML-3, HepG2, or HEK293T cells (Ramos: 8 mL at 3 million cells mL$^{-1}$ seeded 1 h before treatment. OCI-AML-3: 6 mL at 3 million cells mL$^{-1}$ seeded 1 h before treatment. HepG2: 15 million cells seeded 24 h before treatment in 15 cm$^2$ dish. HEK293T: 5 million cells seeded 48 h before treatment in a 15 cm$^2$ dish and transfected 24 h before treatment with 9 µg plasmid) were treated with DMSO or 20–50 µM non-alkyne competitor stereoprobes for 2 h. The cells were further treated with 10–50 µM stereochemically matched alkyne probes for 1 h. For suspension cell lines, the cells were washed three times with chilled DPBS. For adherent cell lines, the cells were scraped and transferred to tubes, then washed three times with chilled DPBS. In both cases, the cell pellet was immediately processed or stored at −80 °C. The cell pellets were resuspended in 500 µl of cold DPBS and lysed by pulse sonication (2 × 15 pulses, 10% power output). The total protein content of the whole-cell lysates was measured using a Pierce BCA protein assay kit. The samples were normalized to 2 mg mL$^{-1}$ and 500 µl (1 mg of proteome) treated with 55 µl of click mix (30 µl of 1.7 mM TBTA in 4:1 *t*-BuOH:DMSO, 10 µl of 50 mM $CuSO_4$ in $H_2O$, 5 µl of 10 mM biotin-PEG4-azide in DMSO, 10 µl of freshly prepared 50 mM tris(2-carboxyethyl)phosphine in DPBS) for 1 h at RT with vigorous vortexing every 20 min. Proteins were precipitated out of solution by the addition of chilled HPLC-grade methanol (600 µl), chloroform (200 µl) and water (100 µl), followed by vigorous vortexing and centrifugation at 16,000*g* for 10 min, to create a disk. Without disrupting the protein disk, both top and bottom layers were aspirated, and the protein disk re-sonicated in 500 µl of methanol and centrifuged at 16,000*g* for 10 min. After complete aspiration of the methanol, protein pellets were frozen at −80 °C or immediately resuspended in 500 µl of freshly made 8 M urea in DPBS, followed by the addition of 10 µl of 10% SDS, and pellet was sonicated to clarity. The samples were reduced with 25 µl of 200 mM dithiothreitol (DTT) at 65 °C for 15 min, followed by alkylation with 25 µl of 400 mM iodoacetamide at 37 °C for 30 min. The samples were quenched with 130 µl of 10% SDS, transferred to a 15-ml tube, and the total volume was brought to 6 mL with DPBS (0.2% final SDS). Washed streptavidin beads (Thermo, catalogue number 20353; 100 µl of 50% slurry/ sample) was then added and the probed labelled protein enriched for 1.5 h at RT with rotation. After incubation, the beads were pelleted (2 min × 2,000*g*) and washed with 0.2% SDS in DPBS (2 × 10 mL), DPBS (1 × 5 mL, then transferred to a protein low-bind Eppendorf safe-lock tube), high-performance liquid chromatography (HPLC) water (2 × 1 mL) and 200 mM 4-(2-hydroxyethyl)-1-piperazinepropanesulfonic acid (EPPS; 1 × 1 mL) at RT Enriched proteins were digested on-bead overnight with 200 µl of trypsin mix (2 M urea, 1 mM $CaCl_2$, 10 µg mL$^{-1}$ trypsin, 200 mM EPPS, pH 8.0). The beads were spun down, supernatant was collected and 100 µl of acetonitrile (30% final) was added, followed by 6 µl of 20 mg mL$^{-1}$ (in dry acetonitrile) of the corresponding TMT$^{16plex}$ tag (for competitive protein-directed ABPP or TMT$^{10plex}$ for non-competitive protein-directed ABPP) for 1.5 h at RT with vortexing every 30 min. TMT labelling was quenched by the addition of hydroxylamine (6 µl 5% solution in $H_2O$) and incubated for 15 min at RT Samples were then acidified with 20 µl 100% formic acid, combined and dried with a SpeedVac. Samples were desalted with a Sep-Pak column and then high pH fractionated into ten fractions using peptide desalting spin columns (as described in below).

**Data processing.** ***Stereoselective enrichment and liganding.*** Enrichment ratios (probe versus probe) were calculated for each peptide–spectra match by dividing each TMT reporter ion intensity by the sum intensity for all the channels. Peptide–spectra matches were then grouped based on protein ID and (excluding peptides with summed reporter ion intensities <22,500) coefficient of variation of >0.5, and <2 distinct peptides. Replicate channels were grouped across each experiment, and average values were computed for each protein. A variability metric was also computed across replicate channels, which equaled the ratio of the median absolute deviation to the average for channels corresponding to the highest enriching probe and was expressed as a percentage. A protein was considered stereoselectively enriched if the average enrichment by the alkyne probe was greater than 2.5-fold that of its enantiomer and if the variability corresponding to the alkyne probe leading to highest enrichment did not exceed 9%. A protein was considered stereoselectively liganded if enantioselective enrichment was greater than 1.42-fold the enrichment observed following treatment with a stereochemically matched non-alkyne competitor (i.e. 30% blockade of enrichment in competitor-treated channel as compared to DMSO-pretreated channels). ***Comparison to stereoisomeric acrylamides.*** For comparison to the stereoisomeric azetidines, a protein was considered stereoselectively liganded in a chemotype-selective manner if it was enantioselectively enriched >2.5-fold by a stereoprobe with >50% competition from the stereomatched competition from one chemotype and enriched with <2-fold enantioselectivity or <25% competition by the other chemotype. Conversely, a protein was considered a shared liganded target if it was enantioselectively enriched >2.5-fold by a stereoprobe with >50% competition from the stereomatched competition from one chemotype and enantioselectively enriched >2-fold and with >25% competition by a stereoprobe from the other chemotype.

**Cloning and mutagenesis**

All full-length plasmids were obtained from GenScript in pcDNA3.1-C-(k) DYK (FLAG) or were acquired from the human ORFeome V8.1 library. Mutagenesis was carried out using a Q5 site-directed mutagenesis kit (New England BioLabs, E0554S) using primers as detailed in each individual section.

#### Generation of CRISPR-mediated TLCD1-null HepG2 cells

sgTLCD1 HepG2-Cas9 cells were generated by the transduction of cells with LentiCRISPR v2-Puro (Addgene, 52961) carrying sgTLCD1. The sequences for the sgRNAs are described below (sequences are 5′ to 3′):

sgTLCD1-1-fwd: caccgGGGTGTCTTAACACTACTGG
sgTLCD1-1-rev: aaacGTGGGGGTGTCTTAACACTACTGGTGGAAG
sgTLCD1-2-fwd:caccATGGACAAGGTATTCCCAAG
sgTLCD1-2-rev:aaacCGTGATGGACAAGGTATTCCCAAGAGGCTC

SgRNAs were annealed and cloned into LentiCRISPR v2 Puro using BsmBI-v2 (NEB Golden Gate Assembly Kit, NEB #E1602). To generate lentivirus, 500,000 293TV cells were seeded in 6-cm$^2$ dishes overnight. CRISPR v2-Puro-sgRNA was co-transfected with dr8.9 envelope and VSV-G packaging plasmids using the FuGENE6 transfection reagent (Promega). Virus-containing supernatants were collected 48 hours after transfection and used to infect target HepG2 cells in the presence of 10 mg mL$^{-1}$ polybrene (Santa Cruz Biotechnology). Cells were spin-infected at 930g and 30 °C for 1 h and incubated for 48 h at 37 °C. Puromycin (2 μg mL$^{-1}$) was then added to the cells, and the cells were expanded. Selection was conducted for 10 days, after which cells were cultured with 0.8 μg mL$^{-1}$ Puromycin. Genetic alteration in sgTLCD1 cells was confirmed by protein-directed MS-ABPP experiments involving comparative enrichment of TLCD1 by MB-2A in WT vs sgTLCD1 HepG2 cells, which confirmed >90% decrease in protein abundance as compared to WT HepG2 cells.

#### Generation of sgTLCD1 base-edited HepG2 cells with a tet-inducible promoter for recombinant TLCD1 wild type and H117N, H118N, and E145Q mutants.

Full length TLCD1 (WT, H117N, H118N, and E145Q) carrying a C-terminal FLAG tag with silent mutation in the PAM NGG sequences was cloned into pIND20 (addgene 44012) vector by Gateway cloning. Viral supernatants were generated as described above using dr8.9 envelope and VSV-G packaging plasmids and used to transduce TLCD1 CRISPR KO cells (sgTLCD1-1 and sgTLCD1-2). After 48 h, cells were selected with 2000 μg $ml^{-1}$ of G418 for two weeks. Selected stable pools were evaluated after 24 h induction with 0, 40, 125, 250, 500, 1000, and 2000 ng/mL doxycycline by immunoblot with anti-FLAG antibody (1:750 dilution). Cells were cultured with 0.8 μg $mL^{-1}$ puromycin and 800 μg $mL^{-1}$ G418.

H117N-fwd: ATACCTTGTCAATCACGTCATGGCC
H117N-rev: TCCCAAGAGGCTCGCGTC
H118N-fwd: CCTTGTCCATAACGTCATGGCCA
H118N-rev: TATTCCCAAGAGGCTCGC
E145Q-fwd: ACTACTGGTGCAGGTCAGCAACATC
E145Q-rev: GTTAAGACACCCCCAC
PAM1-mutation-fwd: AACACTACTGGTTGAAGTCAGCAACATC
PAM1-mutation-rev: AAGACACCCCCACCGACA
PAM2-mutation-fwd: ACAGACGCGAGCGTCTTGGGAATA
PAM2-mutation-rev: CCGCTAGCCACGATGTCC

#### Gel-ABPP with recombinant proteins

For stereoprobe labeling involving transient expression of recombinant proteins (STING, CD36, FABP5, ENL), HEK293T cells ($3 \times 10^5$) were seeded in a six-well plate overnight and transfected with 1 μg of FLAG-epitope tag plasmids using polyethylenimine (PEI) at a ratio of 1:3 DNA:PEI (pre-incubated in opti-MEM for 15 min) for 48 h. The cells were treated with alkyne probe only for 1 h or with competitor probe for 1 h, followed by alkyne probe for a further 1 h, processed, run on 8% (CD36), 16% (FABP5), or 4-20% (all others) SDS page gels and analyzed by in-gel fluorescent scanning as described above.

After in-gel fluorescent scanning, proteins were blotted onto nitrocellulose membrane, blocked with 5% non-fat milk in TBST for 30 min at RT and immunoprobed with anti-FLAG antibody (1:2,000 dilution) in 5% non-fat milk overnight at 4 °C. The membranes were washed five times for 5 min with TBST and incubated in the corresponding anti-rabbit HRP conjugated secondary antibody for 2 h at RT in 5% non-fat milk and washed three times for 5 min. Membranes were developed with ECL western blot substrate (Thermo Fisher Scientific, catalogue number PI32106) and images taken with BioRad Image Lab version 6.1.

*Primers for study of recombinant FABP5 and STING*
FABP_C120A-fwd: AGTGGTGGAGgctGTCATGAACAATG
FABP_C120A-rev: AATTTCCCATCTTTCAATTTTC
FABP_Y131F-fwd: TACTCGGATCTTTGAAAAAGTAG
FABP_Y131F-rev: CAGGTGACATTGTTCATG
STING_C91A-fwd: CTGCCTGGGCgcgCCCCTCCGCC
STING_C91A-rev: GCCCGCACAGTCCTC

#### Gel-ABPP involving immunoprecipitation of recombinant proteins

For stereoprobe labeling of TLCD1, sgTLCD1 HepG2 cells with a tet-inducible promoter for expression of recombinant TLCD1 WT, H117N, H118N, or E145Q variants (6 x $10^6$) were seeded in a 15cm$^2$ dish overnight. The next day, cells were induced with 125 ng mL$^{-1}$ (WT and H117N), 1000 ng mL$^{-1}$ (WT and H117N), 40 ng mL$^{-1}$ (WT and H117N), or DMSO (mock). 24 h later, the cells were treated with alkyne probe only for 1 h or with competitor probe for 1 h, followed by alkyne probe for a further 1 h. Cells were washed 3x with chilled DPBS, harvested, and lysed by pulse sonication (2 × 10 pulses, 10% power output) in chilled DPBS containing complete protease inhibitor (EDTA-free, Roche). The particulate fraction was then isolated by centrifugation (45 min at 100,000 rcf) and resuspended with pulse sonication (1 × 10 pulses, 10% power output) in 400 µL chilled DPBS containing complete protease inhibitor (EDTA-free, Roche). The total protein content of the membrane lysates was measured using a Pierce BCA protein assay kit. Samples were normalized to 1–1.8 mg mL$^{-1}$ in 400µl DPBS with protease inhibitor and 1% Nonidet-P40 and let rotate at 4 °C for 1 h. 30µL of the lysate was removed for analysis of the input, and then 20 µL of anti-FLAG magnetic beads (Cell Signaling, D6W5B, #82103—very important) washed with 1% Nonidet-P40 in DPBS and resuspended in 40 µL of DPBS containing 1% Nonidet-P40 was added to the lysate. Samples were rotated overnight at 4 °C, and then the beads were washed with chilled DPBS containing 1% Nonidet-P40 (3x1 mL). The beads were resuspended in 30 µL chilled DPBS, and then both the beads and the input were treated with 3 µl of click mix (45 µl of 1.7 mM tris((1-benzyl-4-triazolyl)methyl) amine (TBTA) in 4:1 *t*-BuOH:DMSO, 15 µl of 50 mM $CuSO_4$ in $H_2O$, 15 µl of 1.25 mM rhodamine–polyethylene glycol–azide in DMSO, 15 µl of freshly prepared 50 mM tris(2-carboxyethyl)phosphine in DPBS) for 1 h at RT with vigorous vortexing every 20 min. The click reaction was quenched by the addition of 30 µl of 4× SDS gel loading buffer, followed by vigorous vortexing. 10 minutes after addition of SDS gel loading buffer, the eluant was transferred to fresh vials and resolved on 16% SDS–PAGE and imaged by in-gel fluorescent scanning using a BioRad imager with Image Lab software version 6.1.

After in-gel fluorescent scanning, proteins were blotted onto PVDF membrane activated with methanol (60V, 90 minutes), blocked with 5% 5% non-fat milk in TBST for 30 min at RT and immunoprobed with anti-FLAG (1:750 dilution) or anti-beta-actin (1:2,000 dilution) in 5% non-fat milk overnight at 4 °C. The membranes were washed five times for 5 min with TBST and incubated in the corresponding anti-mouse or anti-rabbit HRP conjugated secondary antibody in 5% non-fat milk for 2 h at RT and washed three times for 5 min. Membranes were developed with ECL western blot substrate (Thermo Fisher Scientific, catalogue number PI32106) and images taken with BioRad Image Lab version 6.1.

#### In situ metabolic labeling assay

HepG2 WT and sgTLCD1 cells were plated at 1,000,000 cells in a 6-cm$^2$ dish the day before the assay. For experiments involving OTP-stereoprobe treatment, cells were treated with DMSO, MB-4A, or MB-4B (50µM) for 2 h, then chased with DMSO or C18:1-d9 FFA (Cayman Chemical) (25 µM) for 1 h. For experiments not involving compound treatment, cells were treated with DMSO or C18:1-d9 FFA (Cayman Chemical) (25 µM) for 1 h. After treatment, cells were collected and washed twice with ice-cold PBS, and pellets were stored at −80 °C for further analysis. Cells were washed twice with cold DPBS, and the total cell metabolome was extracted in 4 mL of 2:1:1 $CHCl_3$/MeOH/DPBS (v/v/v) solution containing the internal standard of 100 pmol PE(12:0/12:0). The mixture was vortexed vigorously and centrifuged at 2,000g for 5 min at 4 °C. The bottom organic phase was collected, and the remaining aqueous phase was acidified with 100 µl of formic acid and re-extracted by the addition of 2 mL of $CHCl_3$. Both organic extracts were pooled, dried down under $N_2$ stream and reconstituted in 150 µl of 2:1 $CHCl_3$/MeOH (v/v) for LC–MS analysis.

Metabolites analyzed in this study were quantified using LC– MS-based MRM methods (Agilent Technologies, 6460 or 6470 Triple Quad). MS analysis was performed using ESI with the following parameters: drying gas temperature, 350 °C; drying gas flow, 9 L min$^{-1}$; nebulizer pressure, 45 Ψ; sheath gas temperature, 375 °C; sheath gas flow, 12 L $min^{-1}$; fragmentor voltage, 100 V; and capillary voltage, 3.5 kV. The MRM transitions for the targeted LC–MS analysis are presented in the compiled lipidomics data spreadsheet. The separation of metabolites was achieved using a 50 mm × 4.6 mm 5-µm Gemini C18 column (Phenomenex) coupled to a guard column (Gemini: C18: 4 × 3 mm). For negative mode analysis, $H_2O$:MeOH (95:5, v/v) with 0.1% NH4OH (v/v) and iPrOH:MeOH:$H_2O$ (60:35:5, v/v) with 0.1% NH4OH (v/v) were used as solvent A and solvent B, respectively. The LC gradient for negative mode analysis was the following after injection: 20% B at 0.1 mL $min^{-1}$ for 5 minutes; increase to 85% B at 0.4 mL $min^{-1}$ for 15 minutes; increase to 100% B at 0.5 mL $min^{-1}$ for 5 minutes; run at 100% B at 0.5 mL $min^{-1}$ for 2 minutes; and then go back to 20% B and equilibrate at 0.5 mL $min^{-1}$ for 5 minutes. Lipid species were quantified by measuring areas under the curve in comparison to the corresponding internal standards and then normalizing to the proteome content (assessed with BCA protein assay).

PE(18:0/18:1-d9) (753.5 m/z → 290.3 m/z), PE(16:0/18:1-d9) (725.4 m/z → 290.3 m/z), LPE(18:1-d9) (487.29 m/z → 290.23 m/z), PE(18:1-d9/22:6) (797.5 m/z → 327.3 m/z), PE(18:1-d9/20:4) (773.5 m/z → 303.24 m/z), PE(18:1-d9/18:1-d9) (760.5 m/z → 290.3 m/z), PE(18:1-d9/18:1) (751.5 m/z → 281.3 m/z), PE(18:1/18:1-d9) (751.5 m/z → 290.3 m/z), and PE(12:0/12:0) (internal standard, 578.3 m/z → 199.2 m/z) were measured.

For addback experiments, HepG2 WT and sgTLCD1 cells with a tet-inducible promoter for TLCD1 wild type and mutants were plated at 700,000 cells in a 6-$cm^2$ dish two days before the assay. The next day, cells were treated with DMSO or doxycycline (125 ng/mL for WT and H117N, 1000 ng/mL for H118N, and 40 ng/mL for E145Q). 24 h later, the media was refreshed and cells were treated with DMSO or C18:1-d9 FFA (Cayman Chemical) (20 µM) for 1 h. After treatment, cells were collected and washed twice with ice-cold PBS, and pellets were stored at −80 °C for further analysis as described above.

#### Metabolomics Data

**Table 1. (OA)-d9 incorporation into PE lipids in parental and sgTLCD1 cells (Figure 4F)**

| | parental | | Parental +(OA)-d9 | | sgTLCD1-1+(OA)-d9 | | | sgTLCD1-2+(OA)-d9 | | |
|---|---|---|---|---|---|---|---|---|---|---|
| Lipid | Avg | SEM | Avg | SEM | Avg | SEM | p-value (vs. parental +(OA)-d9) | Avg | SEM | p-value (vs. parental +(OA)-d9) |
| 16:0/18:1-d9 | 2.45 | 0.69 | 6818 | 235 | 3598 | 110 | 9.26E-04 | 2875 | 251 | 2.61E-04 |
| 18:0/18:1-d9 | 1.25 | 0.43 | 2494 | 78 | 1770 | 28 | 6.23E-03 | 1553 | 85 | 1.33E-03 |
| 18:1/18:1-d9 | 2.45 | 0.73 | 13548 | 501 | 6241 | 90 | 1.02E-03 | 4979 | 339 | 4.28E-04 |
| 18:1-d9/18:1 | 49.8 | 17.62 | 10512 | 214 | 3794 | 86 | 9.31E-05 | 2782 | 222 | 1.29E-04 |
| 18:1-d9/18:1-d9 | 1.52 | 0.66 | 6306 | 302 | 2000 | 107 | 2.20E-04 | 1082 | 86 | 3.19E-04 |
| 18:1-d9/20:4 | 2.42 | 0.60 | 2611 | 56 | 663 | 29 | 4.27E-05 | 452 | 6 | 3.44E-05 |
| 18:1-d9/22:6 | 2.20 | 0.87 | 3322 | 97 | 1364 | 85 | 1.81E-05 | 712 | 36 | 2.04E-04 |

**Table 2. (OA)-d9 incorporation into PE lipids in parental OTP-stereoprobe-treated cells (Figure 4G)**

| | parental | | parental+(OA)-d9 | | parental+MB-4A+(OA)-d9 | | | parental+MB-4B+(OA)-d9 | |
|---|---|---|---|---|---|---|---|---|---|
| Lipid | Avg | SEM | Avg | SEM | AVG | SEM | p-value (vs. parental +(OA)-d9) | Avg | SEM |
| 16:0/18:1-d9 | 2.45 | 0.69 | 6818 | 235 | 4703 | 92 | 4.54E-03 | 6575 | 231 |
| 18:0/18:1-d9 | 1.25 | 0.43 | 2494 | 78 | 1811 | 55 | 1.30E-02 | 2442 | 123 |
| 18:1/18:1-d9 | 2.45 | 0.73 | 13548 | 501 | 9265 | 202 | 6.85E-03 | 12680 | 654 |
| 18:1-d9/18:1 | 49.8 | 17.62 | 10512 | 214 | 6060 | 176 | 9.57E-04 | 10077 | 492 |
| 18:1-d9/18:1-d9 | 1.52 | 0.66 | 6306 | 302 | 3388 | 57 | 1.65E-03 | 5028 | 538 |
| 18:1-d9/20:4 | 2.42 | 0.60 | 2611 | 56 | 838 | 25 | 1.98E-04 | 2285 | 127 |
| 18:1-d9/22:6 | 2.2 | 0.87 | 3322 | 97 | 1660 | 78 | 8.73E-04 | 2935 | 228 |

**Table 3. (OA)-d9 incorporation into PE(18:1-d9/20:4) in addback experiments (Figure 6A)**

| | parental | | parental+(OA)-d9 | | sgTLCD1-1+(OA)-d9 | | | sgTLCD1-1+WT-TLCD1+(OA)-d9 | |
|---|---|---|---|---|---|---|---|---|---|
| Lipid | Avg | SEM | Avg | SEM | Avg | SEM | p-value (vs. parental +(OA)-d9) | Avg | SEM |
| PE(18:1-d9/20:4) | 2 | 0.2 | 1292 | 122.5 | 1198 | 36.5 | 1.39E-02 | 1457 | 66.3 |

| | sgTLCD1-1+H117N-TLCD1 +(OA)-d9 | | | sgTLCD1-1+H118N-TLCD1 +(OA)-d9 | | | sgTLCD1-1+E145Q-TLCD1 +(OA)-d9 | | |
|---|---|---|---|---|---|---|---|---|---|
| Lipid | Avg | SEM | p-value (vs. parental +(OA)-d9) | Avg | SEM | p-value (vs. parental +(OA)-d9) | Avg | SEM | p-value (vs. parental +(OA)-d9) |
| PE(18:1-d9/20:4) | 1376 | 30.5 | 1.17E-02 | 1613 | 53.5 | 5.24E-02 | 1334 | 51.2 | 1.65E-02 |

**Uncropped gels (main figures)**

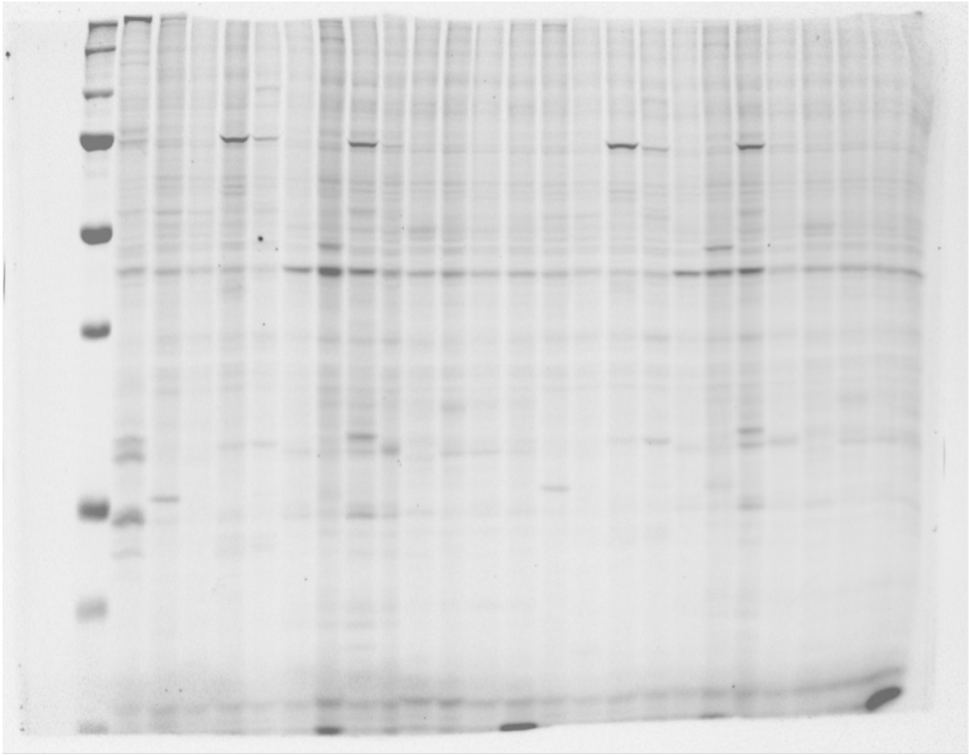

**Figure 1 rhodamine**

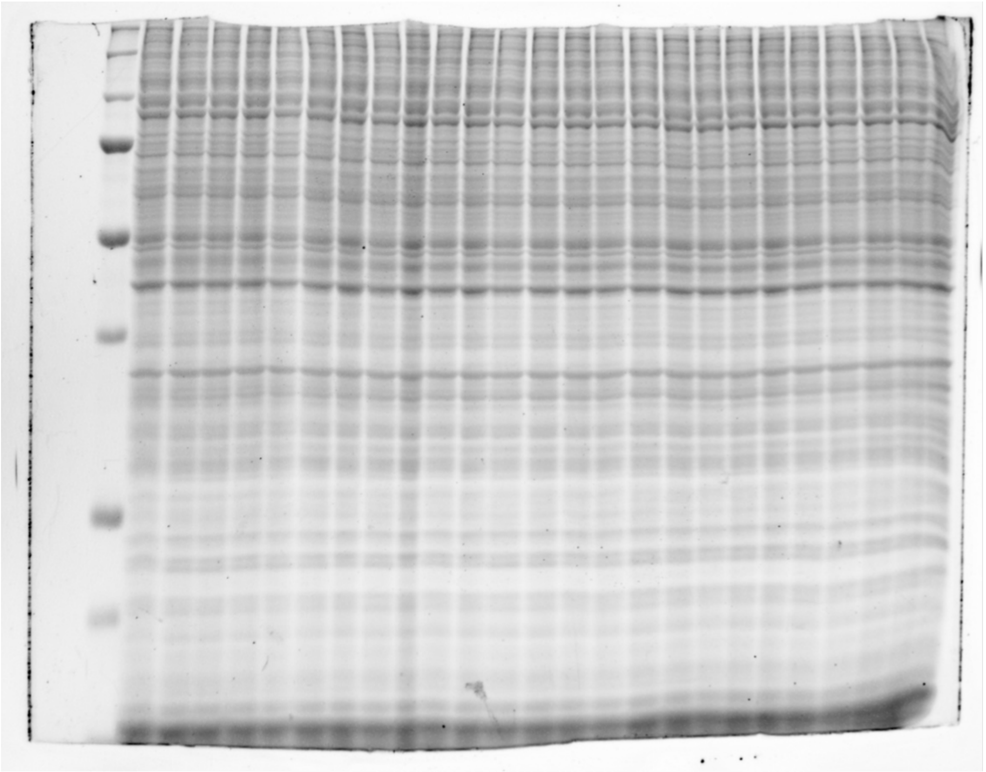

**Figure 1 instant blue**

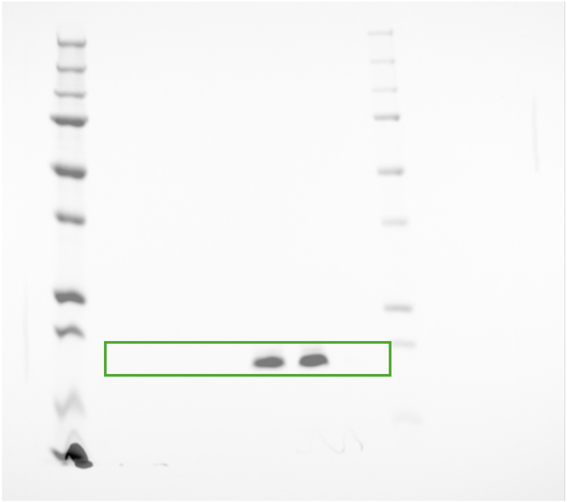

**Figure 3B rhodamine**

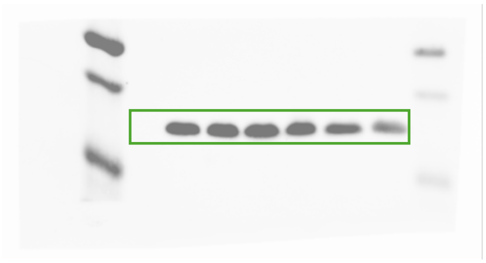

**Figure 3B western blot**

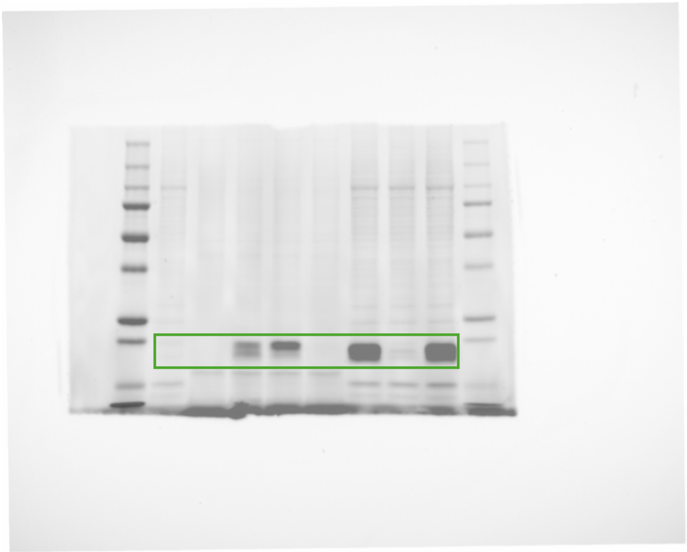

**Figure 3D rhodamine**

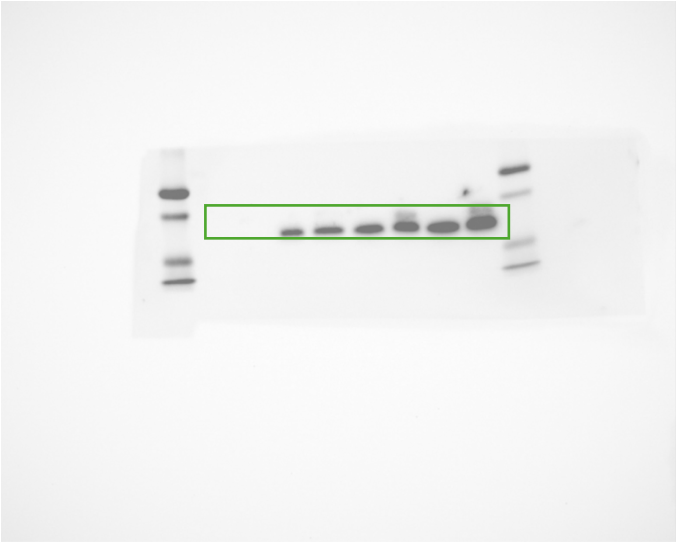

**Figure 3D western blot**

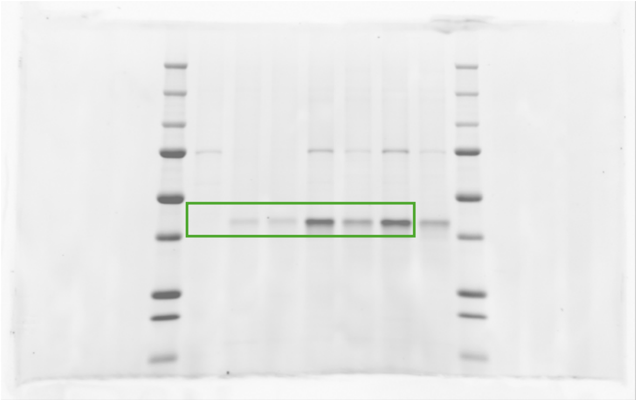

**Figure 3F rhodamine**

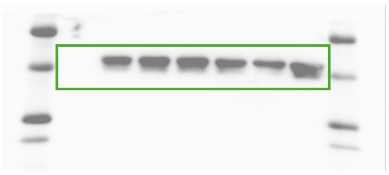

**Figure 3F western blot**

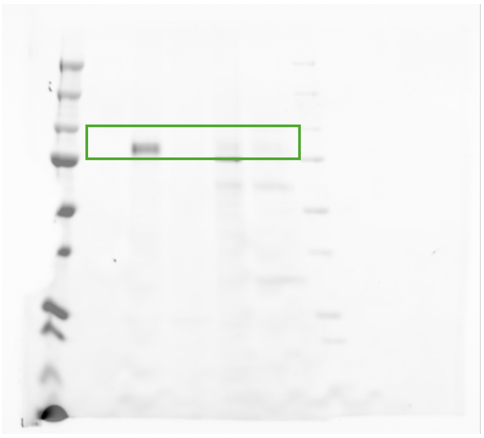

**Figure 3H rhodamine**

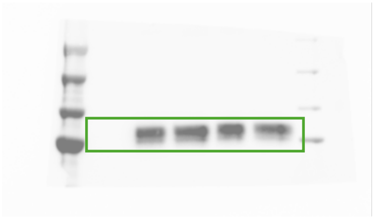

**Figure 3H western blot**

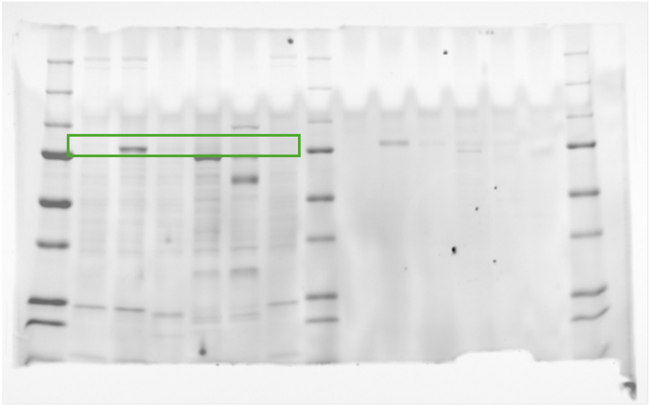

**Figure 3J rhodamine**

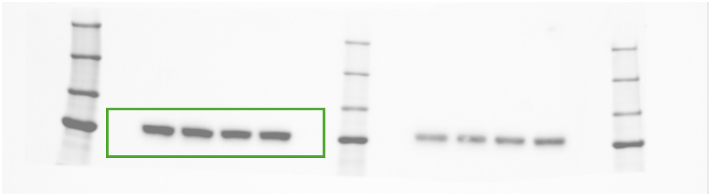

**Figure 3J western blot**

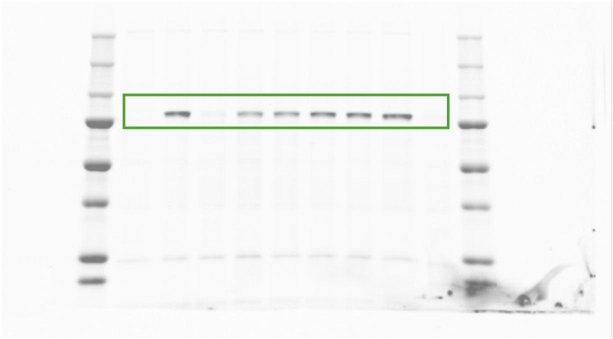
**Figure 3K rhodamine**

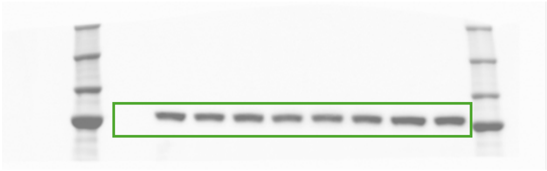
**Figure 3K western blot**

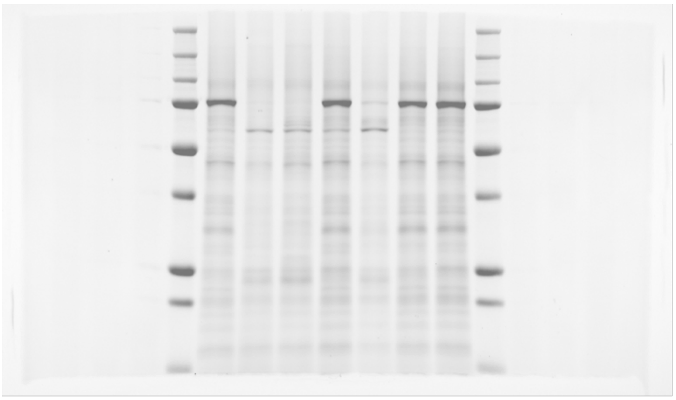
**Figure 4C input rhodamine**

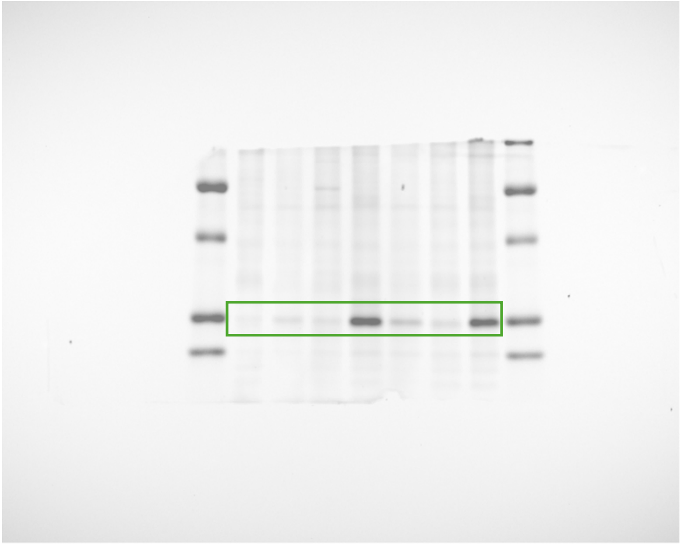
**Figure 4C IP rhodamine**

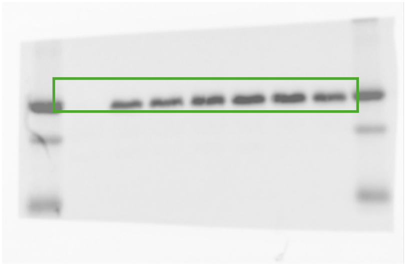

**Figure 4C input anti-FLAG western blot**

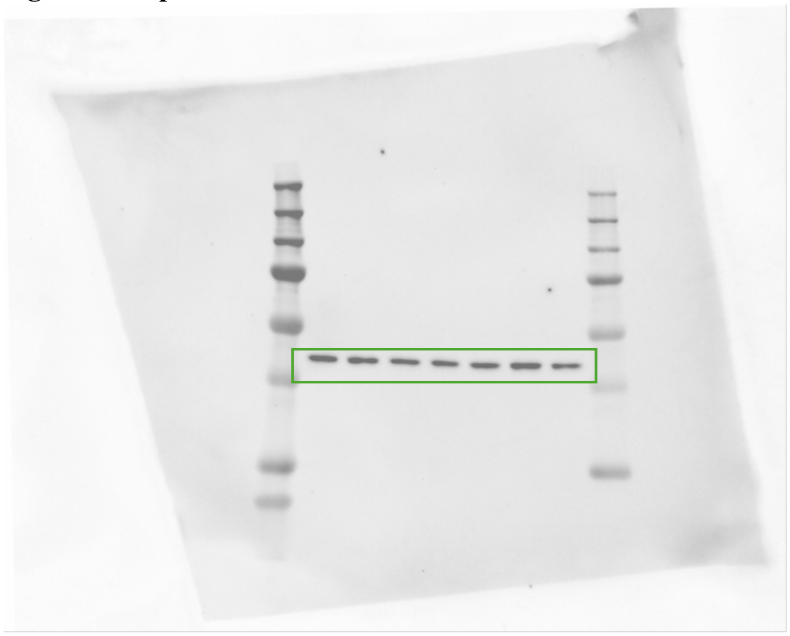

**Figure 4C input anti-actin western blot**

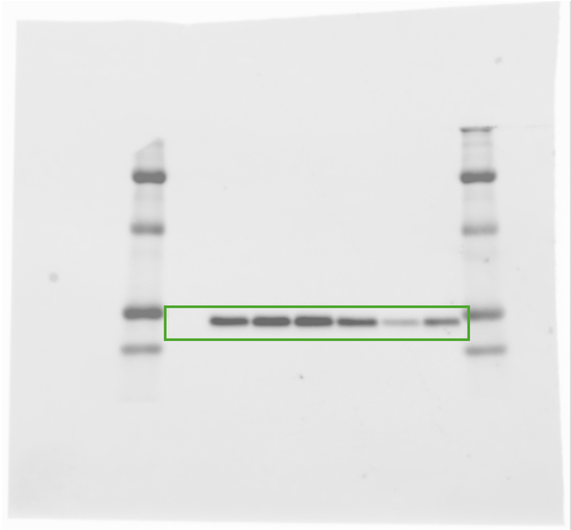

**Figure 4C IP anti-FLAG western blot**

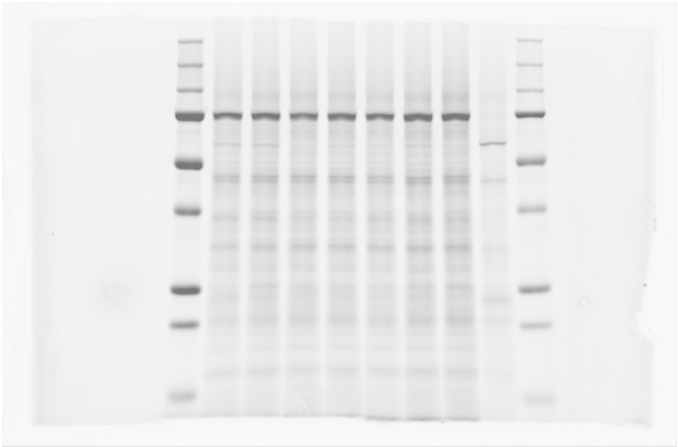
**Figure 4D input rhodamine**

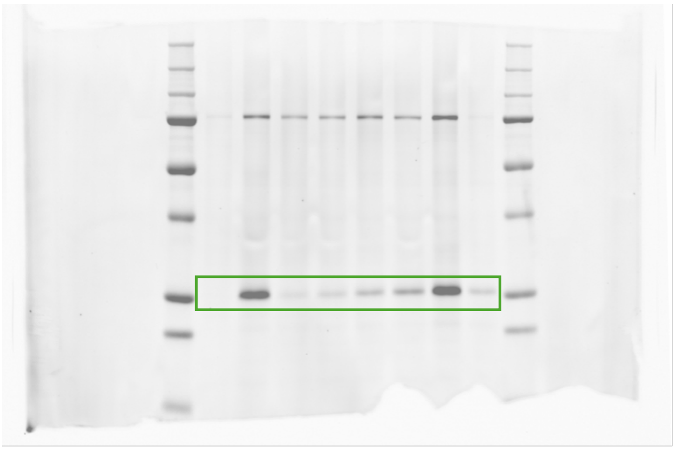
**Figure 4D IP rhodamine**

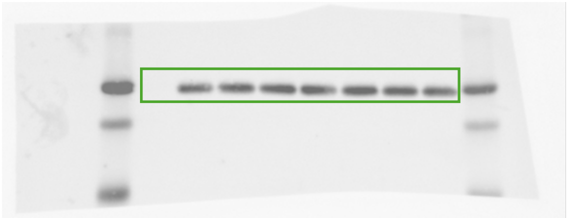
**Figure 4D input anti-FLAG western blot**

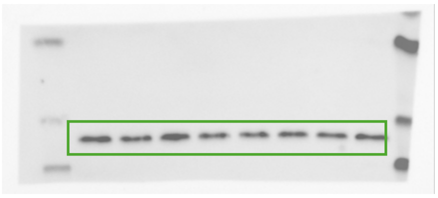
**Figure 4D input anti-actin western blot**

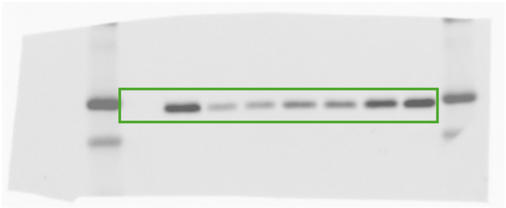

**Figure 4D IP anti-FLAG western blot**

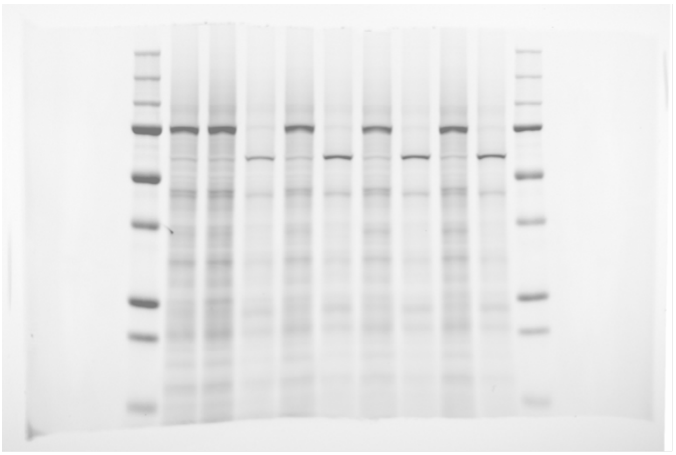

**Figure 5D input rhodamine**

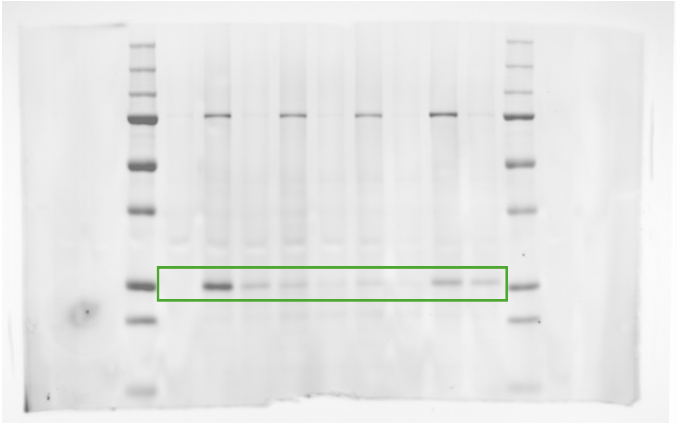

**Figure 5D IP rhodamine**

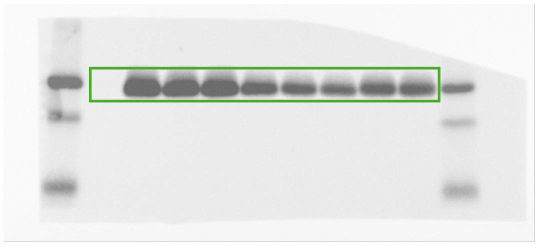

**Figure 5D input anti-FLAG western blot**

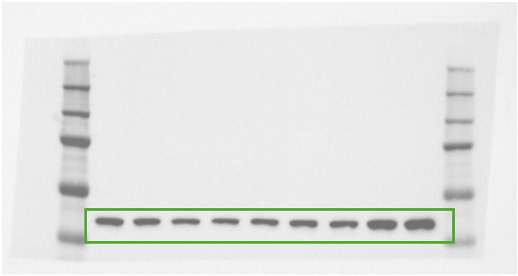

**Figure 5D input anti-actin western blot**

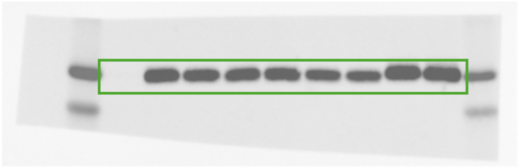

**Figure 5D IP anti-FLAG western blot**

**Uncropped gels (supplementary figures)**

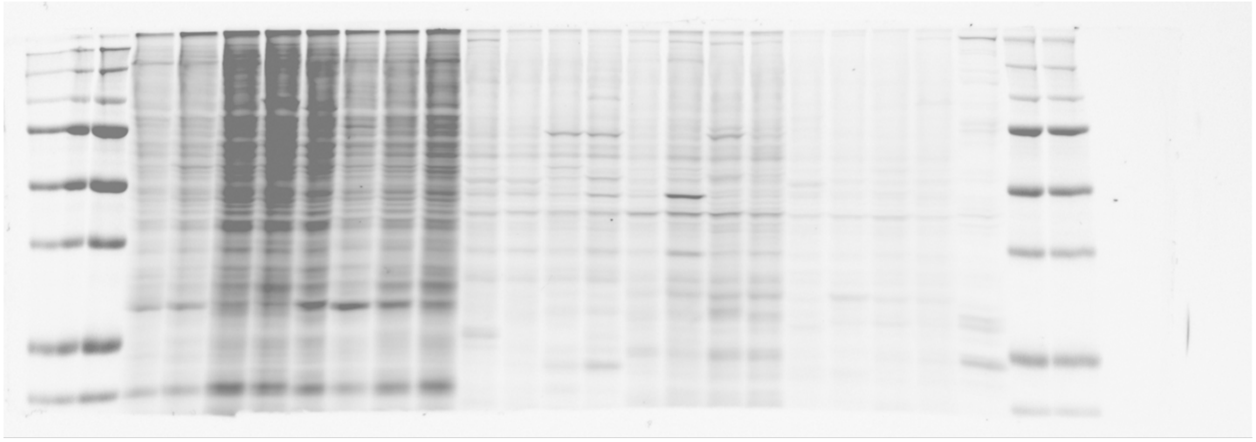

**Figure S1 rhodamine**

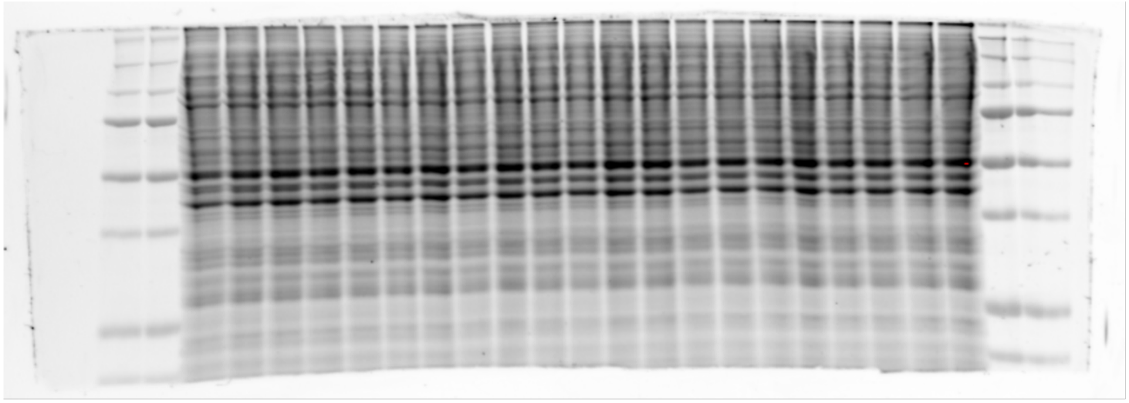

**Figure S1 instant blue**

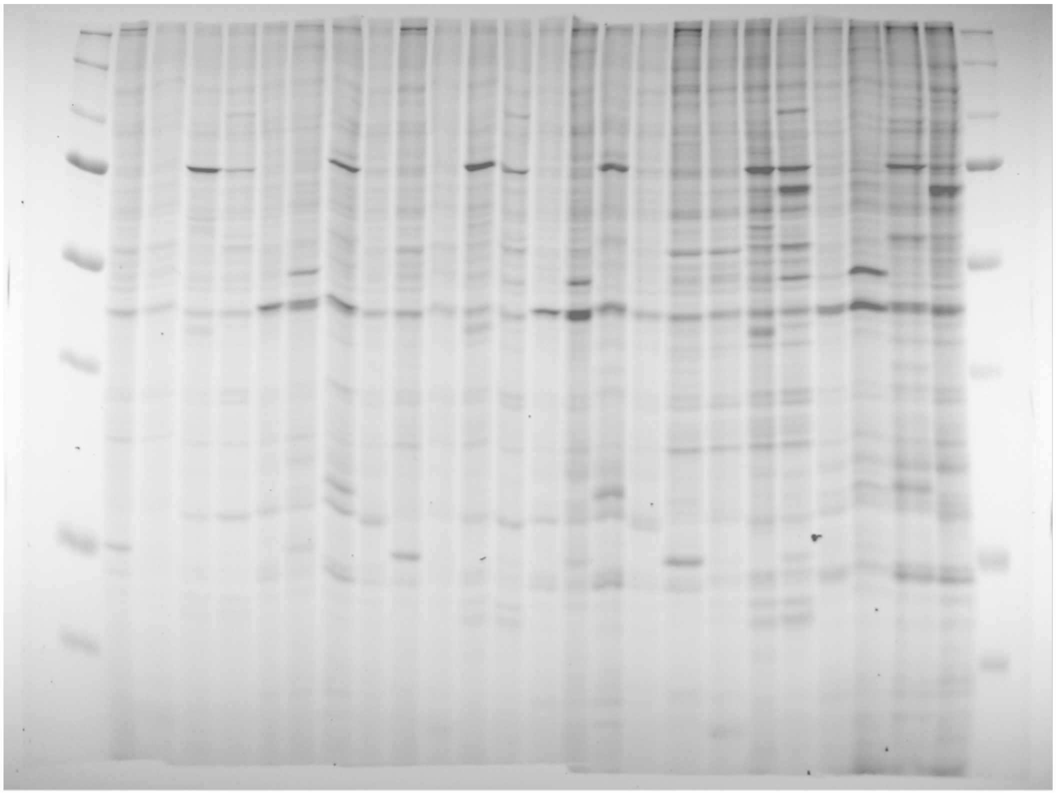

**Figure S2A rhodamine**

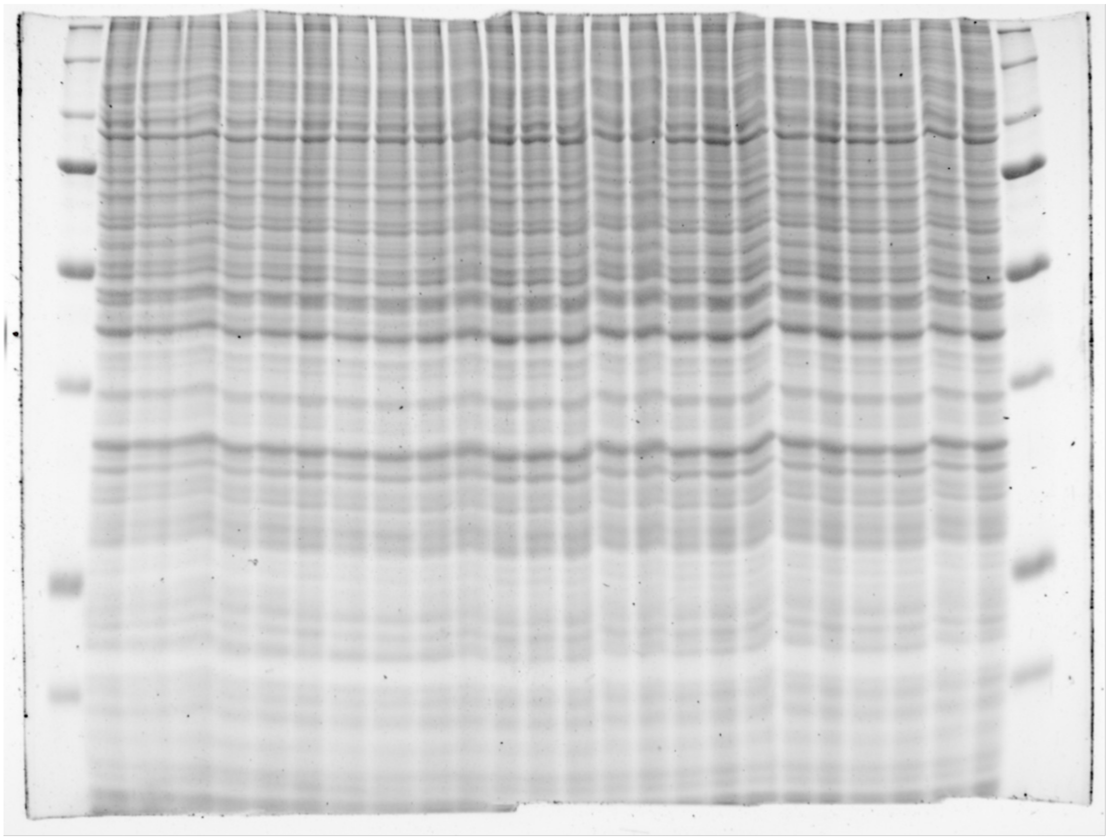

**Figure S2A instant blue**

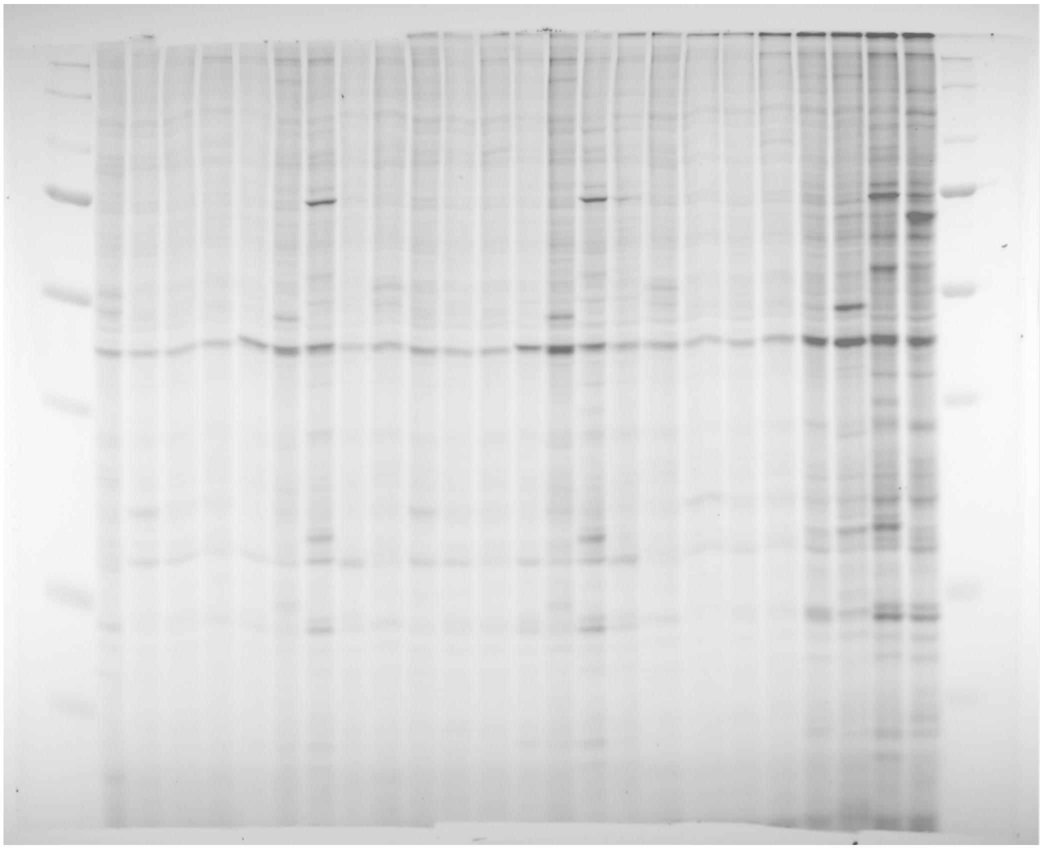

**Figure S2B rhodamine**

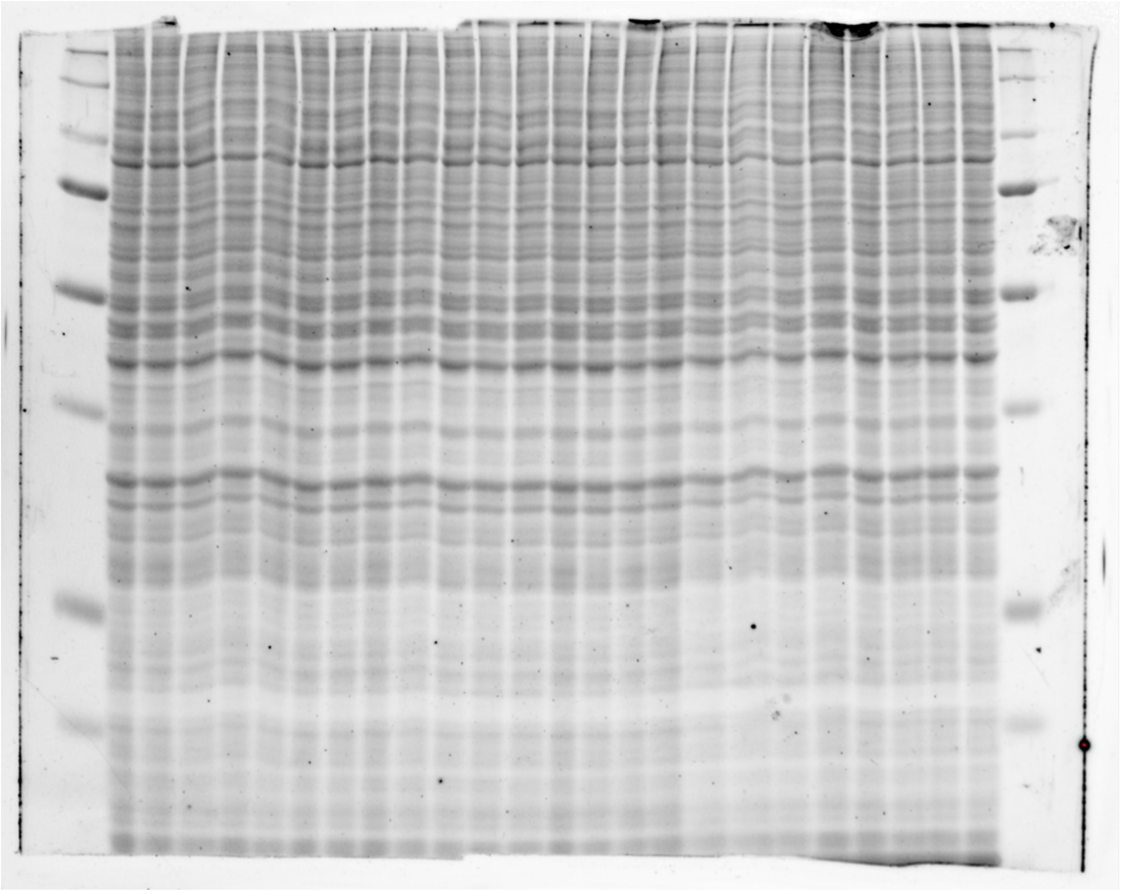

**Figure S2B instant blue**

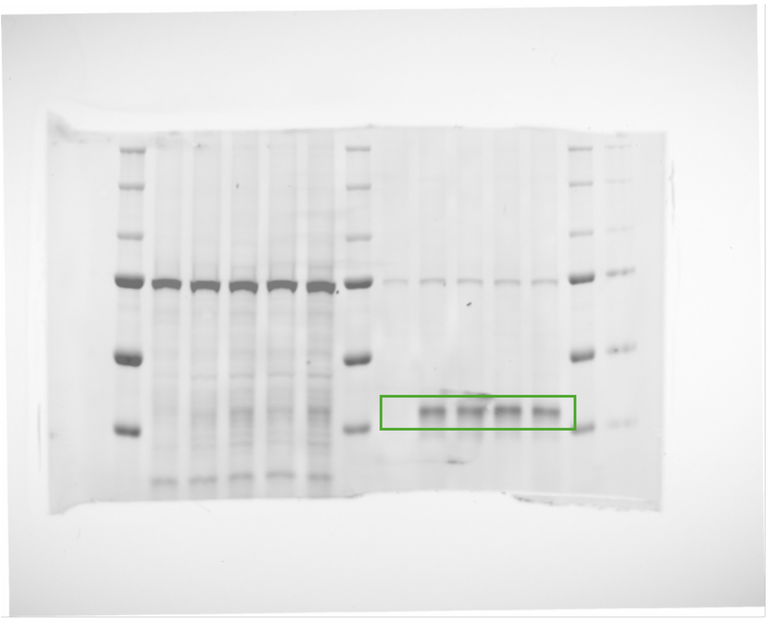

**Figure S5B rhodamine**

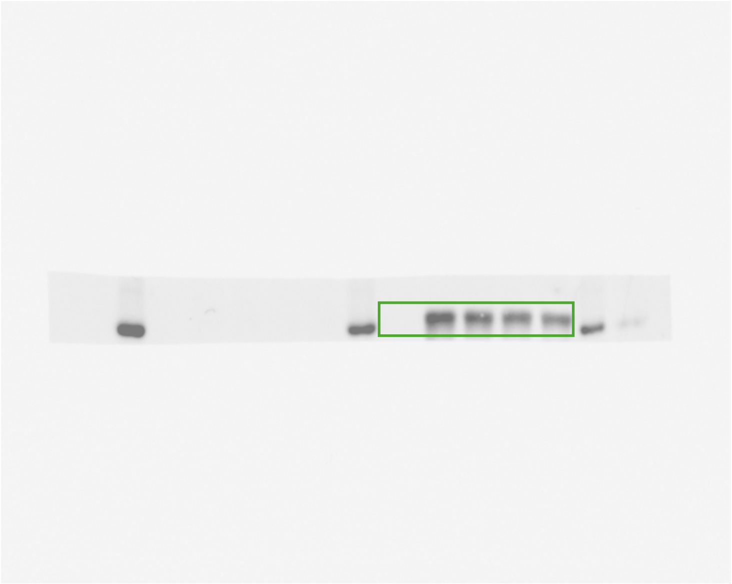

**Figure S5B anti-FLAG western blot**

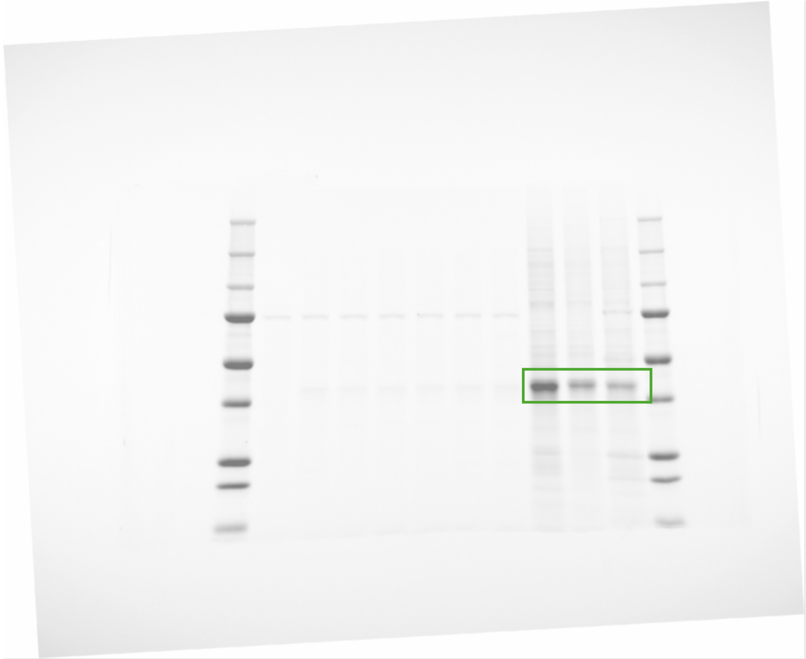

**Figure S5C rhodamine**

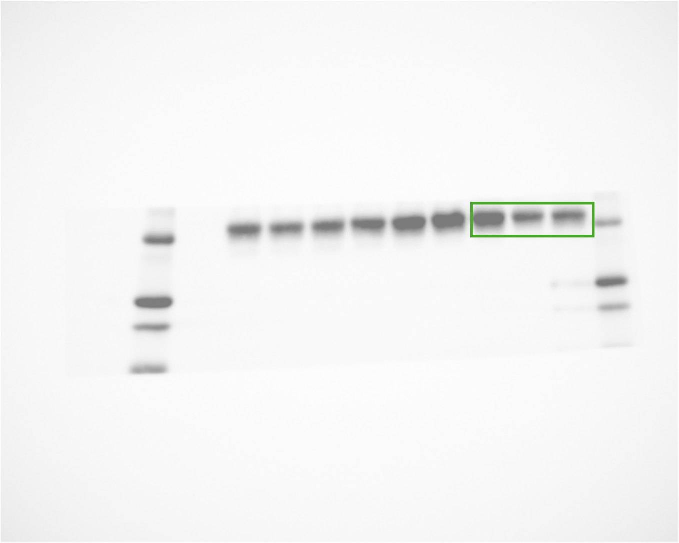

**Figure S5C anti-FLAG western blot**
