## Supplementary Chemistry Information for "Proteomic Ligandability Maps of Phosphorus(V) Stereoprobes Identify Covalent TLCD1 Inhibitors"

### Chemical Supporting Information

### 1. General Experimental

Tetrahydrofuran (THF), *N*,*N*-dimethylformamide (DMF), acetonitrile (MeCN), and toluene (PhMe) were obtained by passing the previously degassed solvents through an activated alumina column. Commercial anhydrous solvents: 1,2-dichloroethane (DCE) (Sigma-Aldrich), dioxane (Sigma-Aldrich), 2-methyltetrahydrofuran (Acros Organics), α,α,α-trifluorotoluene ($PhCF_3$) (Sigma-Aldrich) were used without any further purification. All reagents were purchased of the highest commercial quality and used without further purification unless otherwise stated. Isolated yields refer to chromatographically and spectroscopically ($^1H$ NMR) pure material, unless otherwise stated. Thin layer chromatography (TLC) was performed on 0.25 mm E. Merck silica plates (60F-254), using short-wave UV light as the visualizing agent, and cerium ammonium molybdate (CAM), *p*-anisaldehyde or $KMnO_4$ and heat as the developing agents. Column chromatography was performed using E. Merck silica gel (60, particle size 0.043–0.063 mm). Automated reverse phase flash chromatography was performed on Teledyne ISCO CombiFlash NextGen 300+ using RediSep Gold® Reverse-phase C18 cartridges. NMR spectra were recorded on Bruker DRX-600, DRX-500, and AMX-400 instruments and are calibrated using residual solvent ($CHCl_3$, MeOH, DMSO, MeCN, HDO at 7.26, 3.31, 2.50, 1.94, 4.79 ppm for $^1H$ NMR, respectively, and 77.16, 49.00, 39.52, 1.32 ppm for $^{13}C$ NMR, respectively). The $^{31}P$ NMR and $^{19}F$ chemical shifts are proton decoupled referenced to the deuterium solvent calibrated in $^1H$ NMR. High-resolution mass spectra (HRMS) were recorded either on an Agilent LC/MSD TOF mass spectrometer by electrospray ionization time of flight reflection experiments or Waters Xevo G2-XS TOF calibrated against sodium formate clusters and using a LeuEnk lockmass. Expected monoisotopic masses were calculated using MassLynx 4.1 and the m/z values for calibrant and lockmass were MassLynx-default values. Melting points were recorded on a Fisher-Johns 12-144 melting point apparatus and are uncorrected. Optical rotations were recorded on a Rudolph Research Analytical Autopol III Automatic Polarimeter. HPLC analyses were conducted on a Agilent 1220 Infinity LC equipped with a photodiode array detector and Agilent 1260 Infinity SFC. UPLC analyses were performed on a Waters I-Class LC and a Waters Acquity QDA MS with a Waters Cortecs C18 column (1.6 µm, 2.1x55 mm) using a 5-99% B gradient lasting 2.5 minutes (A: 0.1% aqueous formic acid, B: 0.06% formic acid in acetonitrile, 0.8 mL/min flow rate, 35 °C column temperature). The single-crystal X-ray diffraction studies were carried out on a Bruker D8 Venture Ultra diffractometer equipped with Mo Kα radiation (λ = 0.71073 Å).

#### 2. General scheme for synthesis of Probes and Competitors

##### 2.1 General Scheme for synthesis of Tryptoline-based Probes and Competitors.

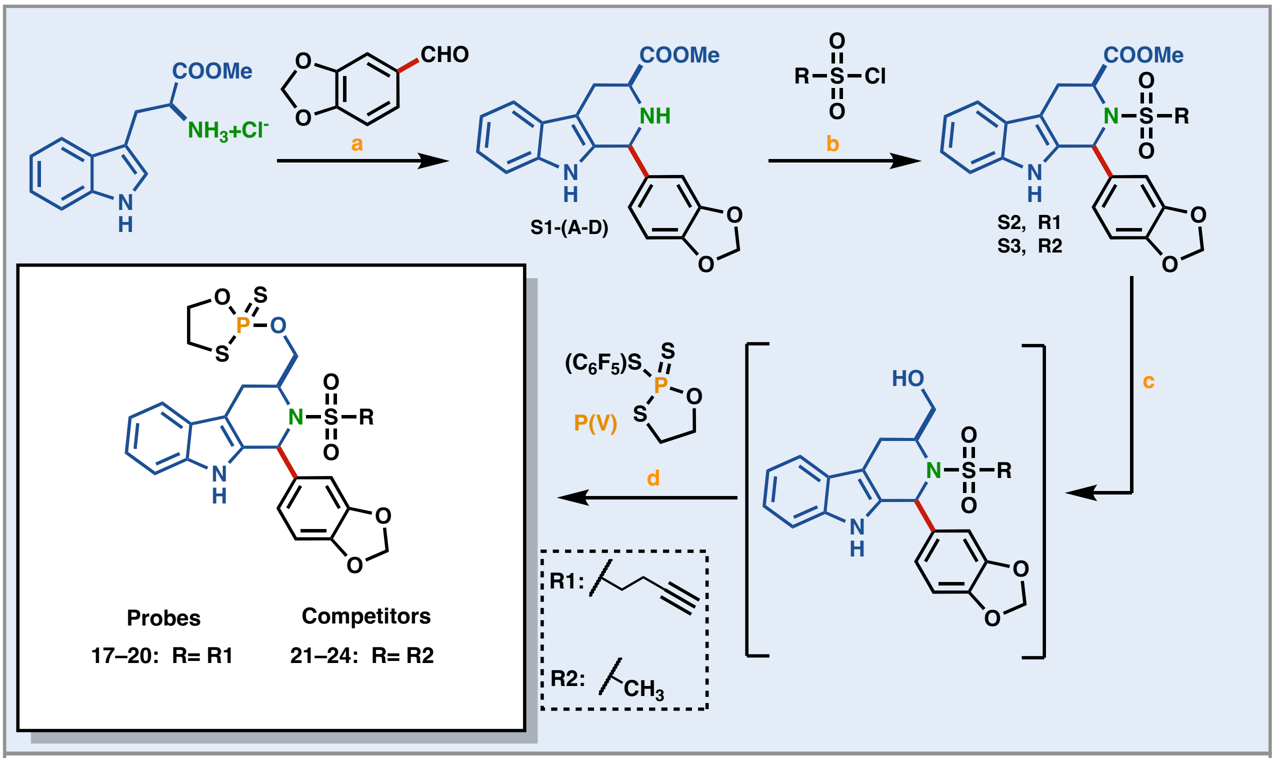

Reaction conditions: a. MeOH, 65 °C, 18 h; b. DMAP, MeCN, -40 °C to rt; c. $LiBH_4$, PhMe, 2-3 hrt; d. P(V), DBU, $CH_2ClCOOH$, MeCN, rt, 30 min

##### Syntheses of P(V)-Reagent:

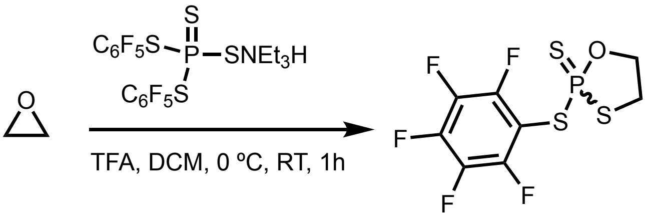

**Ψ**

**2-((perfluorophenyl)thio)-1,3,2-oxathiaphospholane 2-sulfide (Ψ)**

In a 100 mL round bottom flask equipped with a stir bar was charged with dry DCM (49 mL), **bis(2,3,4,5,6-pentafluorophenyl) phosphorotetrathioate triethylammonium salt** (10.0 g, 17.3 mmol, 1 equiv) and **ethylene oxide** (10.3 mL of a ca. 3.0 M solution in THF, 26 mmol, 1.3 equiv). The formed solution was chilled to 0 °C in an ice bath and dry **TFA** (2.97 g, 2.0 mL, 25.9 mmol, 1.3 equiv) was added dropwise with stirring. The flask was removed from the cooling bath and its contents were left stirring at room temperature for 1 h. The volatiles were removed *in vacuo* and the oily residue was dissolved in $Et_2O$ and washed with aq. $NaHCO_3$-brine (1:1, ×2) followed by brine-$H_2O$ (×2); the organic layer was dried with $MgSO_4$, and the solvent was removed by rotary evaporation. The oily residue was dissolved in MeOH (50 mL) and chilled to –78 °C in solid $CO_2$/acetone bath. Once the white precipitate formed it was filtered off and washed with three small portions of –78 °C-cold MeOH to afford 3.4 g of the titled compound after drying (**Yield** = 58%).

*Note: bis(2,3,4,5,6-pentafluorophenyl) phosphorotetrathioate triethylammonium salt was synthesized according to a previously reported procedure (Knouse, K. W.; deGruyter, J. N.; Schmidt, M. A.; Zheng, B.; Vantourout, J. C.; Kingston, C.; Mercer, S. E.; McDonald, I. M.; Olson, R. E.; Zhu, Y.; Hang, C.; Zhu, J.; Yuan, C.; Wang, Q.; Park, P.; Eastgate, M. D.; Baran, P. S., Unlocking P(V): Reagents for chiral phosphorothioate synthesis. Science **2018**, 361, 1234–1238.)*

**Physical state:** white powder

**$^{1}$H NMR (400 MHz, $CDCl_3$) δ** 4.64 (ddt, *J* = 28.5, 10.1, 5.1 Hz, 1H), 4.48 (dddd, *J* = 14.8, 10.0, 8.1, 5.0 Hz, 1H), 3.63 (dtd, *J* = 11.3, 8.4, 5.4 Hz, 1H), 3.41 (ddt, *J* = 13.2, 11.3, 5.0 Hz, 1H).

**$^{13}$C NMR (151 MHz, $CDCl_3$, $^{13}$C {$^{1}$H, $^{19}$F}) δ** δ 148.0 (d, *J* = 4.9 Hz), 143.2 (d, *J* = 4.2 Hz), 137.9 (d, *J* = 3.8 Hz), 104.8 (d, *J* = 7.5 Hz), 71.0 (d, *J* = 3.6 Hz), 37.5.

**$^{31}$P NMR (162 MHz, $CDCl_3$) δ** 106.49 (m).

**HRMS (ESI-TOF) m/z**: n/a

#### Compounds (S1A and S1B).

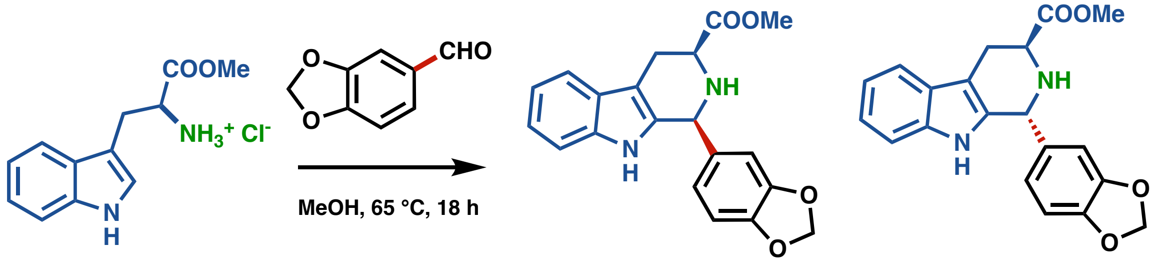

S1A S1B

**Methyl (1*S*,3*S*)-1-(benzo[*d*][1,3]dioxol-5-yl)-2,3,4,9-tetrahydro-1*H*-pyrido[3,4-*b*]indole-3-carboxylate (S1A) and methyl (1*R*,3*S*)-1-(benzo[*d*][1,3]dioxol-5-yl)-2,3,4,9-tetrahydro-1*H*-pyrido[3,4-*b*]indole-3-carboxylate (S1B)**

**Methyl *L*-tryptophan hydrochloride** (5.00 g, 19.6 mmol, 1 equiv) and **piperonal** (3.24 g, 21.6 mmol, 1.1 equiv) were combined in 50 mL of MeOH in a 100 mL round bottom flask equipped with a stir bar and condenser. The reaction vessel was purged and kept under Ar and stirred at reflux for 18 h. **KOH** (1.10 g, 19.6 mmol, 1 equiv) was dissolved in the reaction mixture and the solvent was removed *in vacuo*. The obtained residue was triturated with EtOAc, the organic layers were combined, filtered and the solvent removed via rotary evaporation. TLC (40% EtOAc in hexanes) indicated a roughly 1:1 product ratio. The *trans*- and *cis*- products were isolated by silica chromatography with 35% EtOAc in hexanes to afford 2.30 g (**Yield** = 35%, **d.r.** > 20:1) and 1.70 g (**Yield** = 25%, **d.r.** > 20:1) of **S1A** and **S1B** compounds, respectively.

*Note: this was prepared according to a previously reported method in Bioorganic & Medicinal Chemistry (2004), 12(6), 1505-1515*

#### Compounds (S1A).

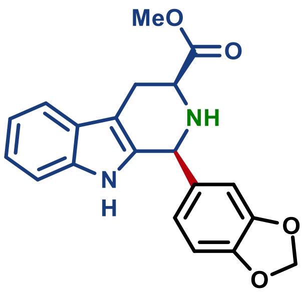

**Methyl (1*S*,3*S*)-1-(benzo[*d*][1,3]dioxol-5-yl)-2,3,4,9-tetrahydro-1*H*-pyrido[3,4-*b*]indole-3-carboxylate (S1A)**

**Physical state:** off-white solid

**$R_f$:** 0.45 (40% EtOAc in hexanes), UV active

**Optical rotation:** $[\alpha]_D^{25}$ −24.0° (*c* 1.2, $CHCl_3$)

**$^1$H NMR (500 MHz, $CDCl_3$) δ** 7.53 (d, *J* = 7.4 Hz, 1H), 7.47 (s, 1H), 7.22 (d, *J* = 7.7 Hz, 1H), 7.13 (m, i2H), 6.88 (d, *J* = 7.9 Hz, 1H), 6.83 (s, 1H), 6.80 (d, *J* = 7.9 Hz, 1H), 5.95 (s, 2H), 5.17 (s, 1H), 3.95 (dd, *J* = 11.1, 4.2 Hz, 1H), 3.82 (d, *J* = 0.9 Hz, 3H), 3.21 (dd, *J* = 15.2, 3.7 Hz, 1H), 3.00 (ddd, *J* = 14.4, 11.0, 2.6 Hz, 1H).

**$^{13}$C NMR (125 MHz, $CDCl_3$) δ** 173.3, 148.4, 148.0, 136.3, 134.8, 134.7, 127.3, 122.15, 122.14, 119.8, 118.4, 111.1, 109.0, 108.9, 108.5, 101.4, 58.6, 57.0, 52.4, 25.8.

**HRMS (ESI-TOF) m/z**: Calcd for $C_{20}H_{18}N_2O_4$ $[M+H]^+$ 351.1345; Found 351.1349

**Compounds (S1B).**

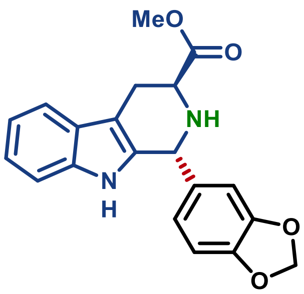

**Methyl (1*R*,3*S*)-1-(benzo[*d*][1,3]dioxol-5-yl)-2,3,4,9-tetrahydro-1*H*-pyrido[3,4-b]indole-3-carboxylate (S1B)**

**Physical state:** light yellow solid

**$R_f$:** 0.33 (40% EtOAc in hexanes), UV active

**Optical rotation:** $[\alpha]_D^{25}$ −35.3° (*c* 1.2, $CHCl_3$)

**$^1$H NMR (400 MHz, $CDCl_3$) δ** 7.66 (s, 1H), 7.55 (d, *J* = 7.6 Hz, 1H), 7.23 (d, *J* = 7.8 Hz, 1H), 7.14 (m, 2H), 6.75 (s, 3H), 5.92 (s, 2H), 5.31 (s, 1H), 3.98 (t, *J* = 6.0 Hz, 1H), 3.72 (s, 3H), 3.25 (dd, *J* = 15.4, 5.4 Hz, 1H), 3.12 (dd, *J* = 15.3, 6.6 Hz, 1H).

**$^{13}$C NMR (151 MHz, $CDCl_3$) δ** 174.3, 148.2, 147.6, 136.3, 136.1, 133.4, 127.1, 122.1, 121.9, 119.7, 118.4, 111.0, 108.8, 108.5, 108.3, 101.3, 54.8, 52.8, 52.3, 24.6.

**HRMS (ESI-TOF) m/z**: Calcd for $C_{20}H_{18}N_2O_4$ $[M+H]^+$ 351.1345; Found 351.1351

#### Compound (S1C).

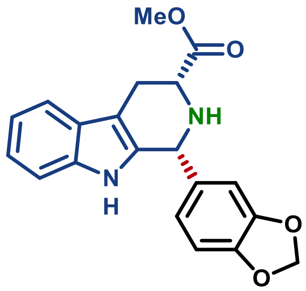

**Methyl (1*R*,3*R*)-1-(benzo[*d*][1,3]dioxol-5-yl)-2,3,4,9-tetrahydro-1*H*-pyrido[3,4-*b*]indole-3-carboxylate (S1C)**

Utilizing procedure reported for **S1A**, compound **S1B** was prepared from **methyl *D*-tryptophan hydrochloride** (5.00 g, 19.6 mmol, 1 equiv) to afford 2.70 g of the titled product (**Yield** = 39%,%, **d.r.** > 20:1).

**Optical rotation:** $[\alpha]_D^{25}$ +23.9° (*c* 1.2, $CHCl_3$)

*Note: other characterizations were identical to compound **S1A***

#### Compounds (S1D).

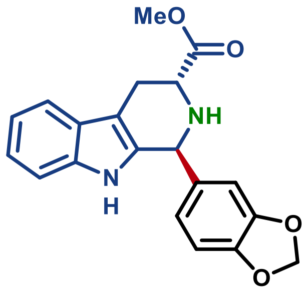

**Methyl (1*S*,3*R*)-1-(benzo[*d*][1,3]dioxol-5-yl)-2,3,4,9-tetrahydro-1*H*-pyrido[3,4-*b*]indole-3-carboxylate (S1D)**

Utilizing procedure reported for **S2A**, compound **S2B** was prepared from **methyl *D*-tryptophan hydrochloride** (5.00 g, 19.6 mmol, 1 equiv) to afford 2.10 g of the titled product (**Yield** = 30%, %, **d.r.** > 20:1).

**Optical rotation:** $[\alpha]_D^{25}$ +32.2° (*c* 1.2, $CHCl_3$)

*Note: other characterizations were identical to compound **S1B***

#### Compound (S2A).

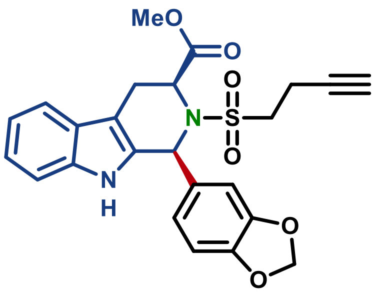

**Methyl (1*S*,3*S*)-1-(benzo[*d*][1,3]dioxol-5-yl)-2-(but-3-yn-1-ylsulfonyl)-2,3,4,9-tetrahydro-1*H*-pyrido[3,4-*b*]indole-3-carboxylate (S2A)**

A 4 dram glass vial equipped with a stir bar was charged with **S1A** (200 mg, 0.571 mmol, 1 equiv), **DMAP** (188 mg, 1.54 mmol, 2.7 equiv), **TEA** (231 mg, 318 µ, 2.28 mmol, 4 equiv), powdered *3Å molecular sieves*, dry MeCN (4.4 mL) and purged with Ar. The mixture was chilled to −40 °C and **3-butyne-1-sulfonyl chloride** (109 µL, 0.713 mmol, 2.5 equiv) (see *note*) was added. After stirring for 15 min. at −40 °C the vial was placed in ice and left stirring for 18 h letting the ice melt. The reaction mixture was diluted with EtOAc, filtered and washed with aq. $NaH_2PO_4$ (×1), brine (×1), dried with $MgSO_4$ and the solvent was removed *in vacuo*. The crude product was isolated by silica chromatography with 80% DCM in hexanes to afford 120 mg of the titled compound (**Yield** = 45%, **d.r.** > 20:1).

*Note*: **3-butyne-1-sulfonyl chloride** was synthesized *according to a previously reported procedure: Org. Lett.* **2021**, *23, 20, 7713–7717.*

**Physical state:** off-white solid

**$R_f$:** 0.41 (40% EtOAc in hexanes) UV active

**Optical rotation:** $[\alpha]_D^{25}$ +72.2° (*c* 1.1, $CHCl_3$)

**$^1$H NMR (400 MHz, $CDCl_3$) δ** 7.64 – 7.59 (m, 2H), 7.28 – 7.27 (m, 1H), 7.22 (t, *J* = 7.2 Hz, 1H), 7.17 (t, *J* = 7.5 Hz, 1H), 6.80 – 6.68 (m, 2H), 6.27 (s, 1H), 5.96 (s, 2H), 5.03 (d, *J* = 6.6 Hz, 1H), 3.69 (d, *J* = 15.9 Hz, 1H), 3.35 (s, 3H), 3.26 (ddd, *J* = 15.1, 9.4, 6.4 Hz, 1H), 3.10 (dd, *J* = 15.8, 6.4 Hz, 1H), 3.03 (ddd, *J* = 14.9, 9.2, 6.7 Hz, 1H), 2.76 – 2.56 (m, 2H), 2.00 (s, 1H).

**$^{13}$C NMR (151 MHz, $CDCl_3$) δ** 171.1, 148.12, 148.08, 136.4, 132.0, 129.2, 126.5, 123.4, 123.0, 120.1, 118.9, 111.2, 109.9, 108.6, 108.1, 101.5, 80.1, 70.5, 55.2, 53.3, 52.62, 52.60, 22.3, 13.8.

**HRMS (ESI-TOF) m/z**: Calcd for $C_{24}H_{23}N_2O_6S$ $[M+H]^+$ 467.1277; Found 467.1275

**Compound (S2B).**

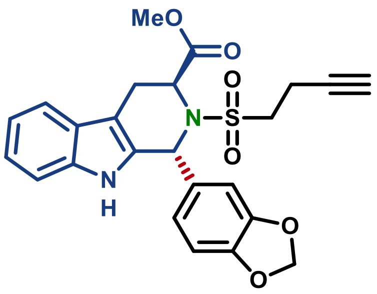

**Methyl (1*R*,3*S*)-1-(benzo[*d*][1,3]dioxol-5-yl)-2-(but-3-yn-1-ylsulfonyl)-2,3,4,9-tetrahydro-1*H*-pyrido[3,4-*b*]indole-3-carboxylate (S2B)**

A 4 dram glass vial equipped with a stir bar was charged with **S1B** (100 mg, 0.285 mmol, 1 equiv), **DMAP** (94 mg, 0.77 mmol, 2.7 equiv), **TEA** (115 mg, 159 µ, 1.14 mmol, 4 equiv), powdered *3Å molecular sieves*, dry MeCN (2.2 mL) and purged with Ar. The mixture was chilled to −40 °C and **3-butyne-1-sulfonyl chloride** (109 µL, 0.713 mmol, 2.5 equiv) (see *note*) was added. After stirring for 15 min. at −40 °C the vial was placed in ice and left stirring for 18 h letting the ice melt. The reaction mixture was diluted with EtOAc, filtered and washed with 1 M HCl (×1), brine (×3), dried with $MgSO_4$ and the solvent was removed *in vacuo*. The crude product was isolated by silica chromatography with 100% DCM to afford 69 mg of the titled compound (**Yield** = 52%, **d.r.** > 20:1).

*Note*: **3-butyne-1-sulfonyl chloride** was synthesized *according to a previously reported procedure: Org. Lett.* **2021***, 23, 20, 7713–7717.*

**Physical state:** off-white solid

**$R_f$:** 0.36 (40% EtOAc in hexanes), UV active

**Optical rotation:** $[\alpha]_D^{25}$ −40.4° (*c* 0.94, $CHCl_3$)

**$^1$H NMR (400 MHz, $CDCl_3$) δ** 7.56 – 7.52 (m, 2H), 7.22 (d, *J* = 8.0 Hz, 1H), 7.17 (t, *J* = 7.5 Hz, 1H), 7.13 (t, *J* = 7.3 Hz, 1H), 6.96 (d, *J* = 8.0 Hz, 1H), 6.79 (d, *J* = 8.0 Hz, 1H), 6.72 (s, 1H), 6.17 (s, 1H), 5.94 (s, 2H), 4.96 (s, 1H), 3.65 (s, 3H), 3.52 (d, *J* = 15.7 Hz, 1H), 3.36 (dd, *J* = 15.3, 4.0 Hz, 1H), 2.82 (dt, *J* = 14.8, 7.1 Hz, 1H), 2.58 – 2.46 (m, 2H), 2.32 (dt, *J* = 15.1, 8.1 Hz, 1H), 2.00 (s, 1H).

**$^{13}$C NMR (151 MHz, $CDCl_3$) δ** 172.0, 148.29, 148.28, 136.9, 133.5, 132.5, 126.3, 123.8, 122.7, 120.0, 118.6, 111.2, 109.4, 108.5, 106.1, 101.6, 77.4, 70.4, 57.9, 57.8, 53.5, 52.9, 24.7, 14.0.

**HRMS (ESI-TOF) m/z**: Calcd for $C_{24}H_{23}N_2O_6S$ $[M+H]^+$ 467.1277; Found 467.1278

#### Compound (S2C).

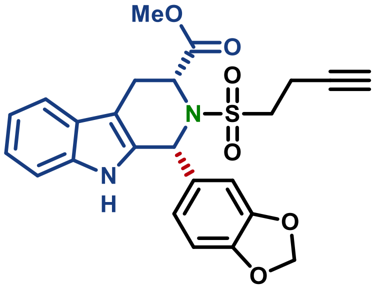

**Methyl (1*R*,3*R*)-1-(benzo[*d*][1,3]dioxol-5-yl)-2-(but-3-yn-1-ylsulfonyl)-2,3,4,9-tetrahydro-1*H*-pyrido[3,4-*b*]indole-3-carboxylate (S2C)**

Utilizing procedure reported for **S2B**, compound **S2C** was prepared from **S1C** (100 mg, 0.285mmol, 1 equiv) to afford 180 mg of the titled product (**Yield** = 68%).

**Optical rotation:** $[\alpha]_D^{25}$ −74.4° (*c* 1.1, $CHCl_3$)

*Note: other characterizations were identical to* ***S2A***

**Compound (S2D).**

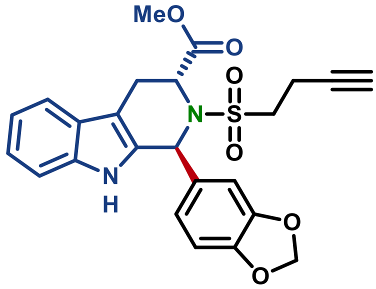

**Methyl (1*S*,3*R*)-1-(benzo[*d*][1,3]dioxol-5-yl)-2-(but-3-yn-1-ylsulfonyl)-2,3,4,9-tetrahydro-1*H*-pyrido[3,4-*b*]indole-3-carboxylate (S2D)**

Utilizing procedure reported for **S2A**, compound **S2D** was prepared from **S1D** (100 mg, 0.285mmol, 1 equiv) to afford 66 mg of the titled product (**Yield** = 50%).

**Optical rotation:** $[\alpha]_D^{25}$ +42.4° (*c* 0.8, $CHCl_3$)

*Note: other characterizations were identical to* ***S2B***

**Compound (S3A).**

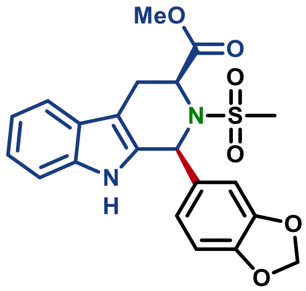

**Methyl (1*R*,3*R*)-1-(benzo[*d*][1,3]dioxol-5-yl)-2-(methylsulfonyl)-2,3,4,9-tetrahydro-1*H*-pyrido[3,4-*b*]indole-3-carboxylate (S3A)**

A 4 dram glass vial equipped with a stir bar was charged with **S1A** (100 mg, 0.285 mmol, 1 equiv), **DMAP** (37 mg, 0.30 mmol, 1.1 equiv), **TEA** (101 mg, 79 µL, 0.57 mmol, 2 equiv), powdered *3Å molecular sieves*, dry MeCN (2.2 mL) and purged with Ar. The mixture was chilled to −40 °C and **mesyl chloride** (34 mg, 23 µL, 0.30 mmol, 1.1 equiv) was added. After stirring for 15 min. at −40 °C the vial was placed in ice and left stirring for 18 h, letting the ice slowly melt. The reaction mixture was diluted with EtOAc, filtered and washed with 1 M HCl (×1), brine (×3), dried with $MgSO_4$ and the solvent was removed *in vacuo*. The crude product was isolated by silica chromatography with 10% EtOAc in PhMe to afford 55 mg of the titled compound (**Yield** = 49%, **d.r.** > 20:1).

**Physical state:** off-white solid

**R$_f$:** 0.15 (40% EtOAc in hexanes), UV active

**Optical rotation:** $[\alpha]_D^{25}$ +59.1° (*c* 1.0, $CHCl_3$)

**$^1$H NMR (400 MHz, $CDCl_3$) δ** 7.65 (s, 1H), 7.61 (d, *J* = 7.8 Hz, 1H), 7.29 – 7.27 (m, 1H)(overlapped with solvent peak), 7.21 (t, *J* = 7.6 Hz, 1H), 7.17 (t, *J* = 7.6 Hz, 1H), 6.80 – 6.73 (m, 3H), 6.26 (s, 1H), 5.95 (s, 2H), 5.14 (d, *J* = 5.5 Hz, 1H), 3.68 (d, *J* = 15.9 Hz, 1H), 3.36 (s, 3H), 3.09 (dd, *J* = 16.4, 6.0 Hz, 1H), 2.84 (s, 3H).

**$^{13}$C NMR (151 MHz, $CDCl_3$) δ** 171.4, 148.08, 148.06, 136.5, 131.8, 129.4, 126.6, 123.5, 122.9, 120.1, 118.9, 111.1, 109.9, 108.6, 108.0, 101.5, 55.4, 53.4, 52.5, 41.7, 22.3.

**HRMS (ESI-TOF) m/z**: Calcd for $C_{21}H_{21}N_2O_6S$ $[M+H]^+$ 429.112; Found 429.1123

#### Compound (S3B).

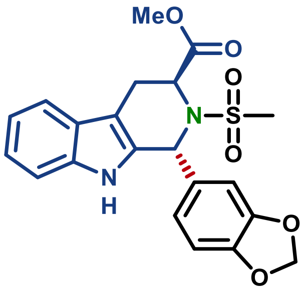

**Methyl (1*R*,3*S*)-1-(benzo[*d*][1,3]dioxol-5-yl)-2-(methylsulfonyl)-2,3,4,9-tetrahydro-1*H*-pyrido[3,4-*b*]indole-3-carboxylate (S3B)**

Utilizing procedure reported for **S3A**, compound **S3B** was prepared from **S1B** (100 mg, 0.285mmol, 1 equiv) to afford 55 mg of the titled product (**Yield** = 45%, **d.r.** > 20:1).

**Physical state:** off-white solid

**R$_f$:** 0.28 (40% EtOAc in hexanes), UV active

**Optical rotation:** $[\alpha]_D^{25}$ −37.0° (*c* 1.0, $CHCl_3$)

**$^1$H NMR (400 MHz, $CDCl_3$) δ** 7.58 (s, 1H), 7.54 (d, *J* = 7.8 Hz, 1H), 7.23 (d, *J* = 8.0 Hz, 1H), 7.17 (t, *J* = 7.4 Hz, 1H), 7.13 (t, *J* = 7.3 Hz, 1H), 6.93 (d, *J* = 7.1 Hz, 1H), 6.77 (d, *J* = 7.9 Hz, 1H), 6.74 (s, 1H), 6.16 (s, 1H), 5.96 (s, 1H), 5.93 (s, 1H), 4.95 (s, 1H), 3.67 (s, 3H), 3.52 (dd, *J* = 15.6, 2.1 Hz, 1H), 3.33 (dd, *J* = 15.7, 4.6 Hz, 1H), 2.57 (s, 3H).

**$^{13}$C NMR (151 MHz, $CDCl_3$) δ** 171.9, 148.3, 148.2, 136.8, 133.3, 133.0, 126.4, 123.5, 122.7, 120.0, 118.6, 111.2, 109.3, 108.4, 106.5, 101.5, 57.8, 57.2, 52.8, 43.8, 24.7.

**HRMS (ESI-TOF) m/z**: Calcd for $C_{21}H_{21}N_2O_6S$ $[M+H]^+$ 429.112; Found 429.1121

#### Compound (S3C).

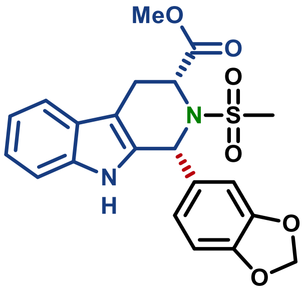

**Methyl (1*R*,3*R*)-1-(benzo[*d*][1,3]dioxol-5-yl)-2-(methylsulfonyl)-2,3,4,9-tetrahydro-1*H*-pyrido[3,4-*b*]indole-3-carboxylate (S3C)**

Utilizing procedure reported for **S3A**, compound **S3C** was prepared from **S1C** (100 mg, 0.285mmol, 1 equiv) to afford 82 mg of the titled product (**Yield** = 67%, **d.r.** > 20:1).

**Optical rotation:** $[\alpha]_D^{25}$ −59.2 ° (*c* 1.2, $CHCl_3$)

*Note: other characterizations were identical to* ***S3A***

#### Compound (S3D).

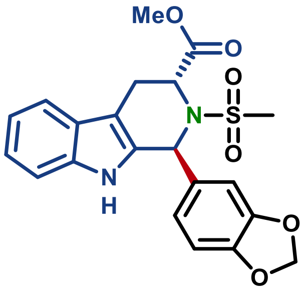

**Methyl (1*S*,3*R*)-1-(benzo[*d*][1,3]dioxol-5-yl)-2-(methylsulfonyl)-2,3,4,9-tetrahydro-1*H*-pyrido[3,4-*b*]indole-3-carboxylate (S3D)**

Utilizing procedure reported for **S3A**, compound **S3D** was prepared from **S1D** (100 mg, 0.285mmol, 1 equiv) to afford 37 mg of the titled product (**Yield** = 30%, **d.r.** > 20:1).

**Optical rotation:** $[\alpha]_D^{25}$ +38.9° (*c* 1.0, $CHCl_3$)

*Note: other characterizations were identical to* ***S3B***

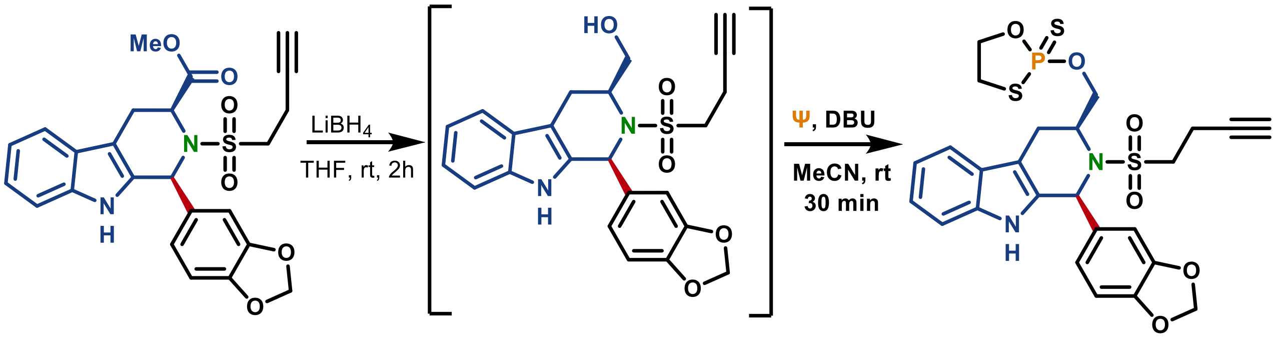

**2-(((1*S*,3*S*)-1-(benzo[*d*][1,3]dioxol-5-yl)-2-(but-3-yn-1-ylsulfonyl)-2,3,4,9-tetrahydro-1*H*-pyrido[3,4-*b*]indol-3-yl)methoxy)-1,3,2-oxathiaphospholane 2-sulfide (17, MB-5A)**

**S3A** (54 mg, 0.130 mmol, 1 equiv) was dissolved in dry PhMe (6.5 mL) in a glass scintillation vial equipped with a stir bar. 2 M $LiBH_4$ solution in THF (261 µL, 4 equiv) added and the mixture was stirred for 2 h at room temperature at which point complete conversion was observed. The reaction was diluted with EtOAc and washed with water (×1), 1 M HCl (×1), brine (×3), dried with $MgSO_4$ and the solvent was removed *in vacuo* to afford 42 mg of the crude alcohol that was used directly in the next step. The obtain residue (42 mg, 0.096 mmol, 1 equiv) was combined with **Ψ** (47 mg, 0.14 mmol, 1.5 equiv) and 2-chloroacetamide (13 mg, 0.14 mmol, 1.5 equiv) in dry MeCN (0.96 mL). A portion of powdered 3Å *molecular sieves* was added and stirred for 5 min at room temperature before adding dry **DBU** (21 mg, 20 µL, 0.14 mmol, 1.5 equiv) and the mixture was stirred for 30 min at room temperature. The reaction mixture was diluted with EtOAc and washed with aq. $KH_2PO_4$ (×2), brine (×2), dried with $MgSO_4$ and the solvent was removed *in vacuo*. The product was isolated by silica PTLC with 40% EtOAc in hexanes and further purified via PTLC with 10% $Et_2O$ in DCM to afford 24 mg of the titled compound (**Yield** = 44%, **d.r.** ~1:1, >98% e.e.).

**Physical state:** light yellow film

**$R_f$:** 0.70 (50% EtOAc in hexanes), UV active

**Optical rotation:** $[\alpha]_D^{25}$ +31.4° (*c* 0.43, $CHCl_3$)

**$^1$H NMR (400 MHz, $CDCl_3$) δ** 7.80 (d, *J* = 6.9 Hz, 1H), 7.56 (d, *J* = 7.8 Hz, 1H), 7.34 (dd, *J* = 8.1, 3.1 Hz, 1H), 7.28 – 7.23 (m, 1H)(*overlapped with $CHCl_3$ peak*), 7.18 (t, *J* = 7.5 Hz, 1H), 6.92 (dd, *J* = 5.8, 1.8 Hz, 1H), 6.82 – 6.78 (m, 1H), 6.77 (d, *J* = 2.8 Hz, 1H), 6.76 (d, *J* = 2.8 Hz, 1H), 6.27 (d, *J* = 13.8 Hz, 1H), 6.02 – 5.94 (m, 2H), 4.69 (dq, *J* = 14.4, 7.4 Hz, 1H), 4.47 – 4.24 (m, 2H), 4.15 – 3.99 (m, 1H), 3.93 – 3.83 (m, 1H), 3.44 – 3.27 (m, 2H), 3.26 – 3.13 (m, 2H), 3.11 – 2.98 (m, 2H), 2.72 – 2.63 (m, 2H), 1.98 (t, *J* = 2.6 Hz, 1H).

**$^{13}$C NMR (151 MHz, $CDCl_3$) δ** 148.5, 148.1, 136.4, 133.4, 133.3, 128.9, 128.8, 126.84, 126.82, 123.2, 122.6, 122.4, 120.3, 118.74, 118.70, 111.32, 111.29, 109.4, 109.3, 108.6, 108.53, 107.51, 107.4, 101.57, 101.56, 80.1, 80.0, 70.7, 70.65, 70.61, 68.8, 68.7, 68.65, 68.6, 54.0, 53.89, 52.3, 52.22, 52.19, 52.12, 52.08, 52.06, 36.7, 36.6, 22.31, 22.27, 14.0, 13.9.

**$^{31}$P NMR (162 MHz, $CDCl_3$) δ** 103.8, 103.6.

**HRMS (ESI-TOF) m/z**: Calcd for $C_{25}H_{26}N_2O_6PS_3$ $[M+H]^+$ 577.0691; Found 577.0695

#### Compound (19, MB-6A).

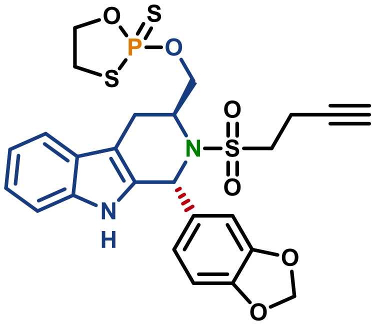

**2-(((1*R*,3*S*)-1-(benzo[*d*][1,3]dioxol-5-yl)-2-(but-3-yn-1-ylsulfonyl)-2,3,4,9-tetrahydro-1*H*-pyrido[3,4-*b*]indol-3-yl)methoxy)-1,3,2-oxathiaphospholane 2-sulfide (19, MB-6A)**

Utilizing procedure reported for **17,** compound **19** was prepared from **S2B**, obtained from reduction of **S4,** to afford 15 mg of the titled product (**Yield** = 29%, **d.r.** ~1:1, 96% e.e.).

*Note: Additional purification with 10% $Et_2O$ in DCM was not required.*

**Physical state:** light yellow film

**$R_f$:** 0.74 (50% EtOAc in hexanes), UV active

**Optical rotation:** $[\alpha]_D^{25}$ −21.3° (*c* 0.38, $CHCl_3$)

**$^1$H NMR (400 MHz, $CDCl_3$) δ** 7.60 – 7.51 (m, 2H), 7.28 – 7.24 (m, 1H)(*overlap with $CHCl_3$ peak*), 7.20 (t, *J* = 7.0 Hz, 1H), 7.15 (t, *J* = 7.4 Hz, 1H), 6.86 – 6.82 (m, 1H), 6.77 (app d, *J* = 8.1 Hz, 1H), 6.74 (s, 1H), 5.98 – 5.94 (m, 2H), 5.90 (d, *J* = 8.9 Hz, 1H), 4.54 – 4.17 (m, 5H), 3.62 – 3.33 (m, 2H), 3.26 – 3.05 (m, 2H), 2.94 – 2.78 (m, 1H), 2.67 – 2.38 (m, 3H), 1.98 (dd, *J* = 2.8 Hz, 1H).

**$^{13}$C NMR (151 MHz, $CDCl_3$) δ** 148.45, 148.44, 148.38, 136.84, 136.82, 131.89, 131.85, 131.83, 131.77, 126.69, 126.68, 123.79, 123.76, 122.94, 122.93, 120.22, 120.20, 118.80, 118.71, 111.30, 111.26, 109.66, 108.45, 108.44, 107.78, 107.71, 101.63, 101.62, 79.77, 79.72, 70.72, 70.71, 70.67, 70.66, 70.51, 67.94, 67.89, 67.69, 67.64, 57.50, 57.48, 54.47, 54.40, 54.33, 54.27, 53.42, 53.40, 36.64, 36.63, 36.53, 36.52, 23.55, 23.49, 14.17, 14.16.

**$^{31}$P NMR (162 MHz, $CDCl_3$) δ** 103.8, 103.7.

**HRMS (ESI-TOF) m/z**: Calcd for $C_{25}H_{26}N_2O_6PS_3$ $[M+H]^+$ 577.0691; Found 577.0696

#### Compound (18, MB-5B).

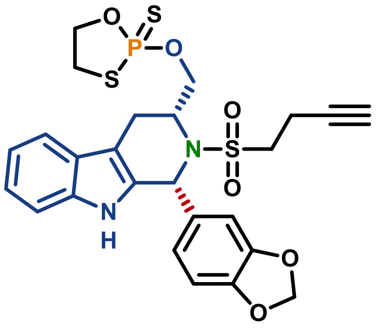

**2-(((1*R*,3*R*)-1-(benzo[*d*][1,3]dioxol-5-yl)-2-(but-3-yn-1-ylsulfonyl)-2,3,4,9-tetrahydro-1*H*-pyrido[3,4-*b*]indol-3-yl)methoxy)-1,3,2-oxathiaphospholane 2-sulfide (18, MB-5B)**

Utilizing procedure reported for **17**, compound **18** was prepared from the **S2C** obtained from reduction of **S3B,** to afford 23 mg of the titled product (**Yield** = 34%, **d.r.** ~1:1, >99% e.e.).

**Optical rotation:** $[\alpha]_D^{25}$ −35.8° (*c* 0.43, $CHCl_3$)

*Note: other characterizations were identical to **17***

#### Compound (20, MB-6B).

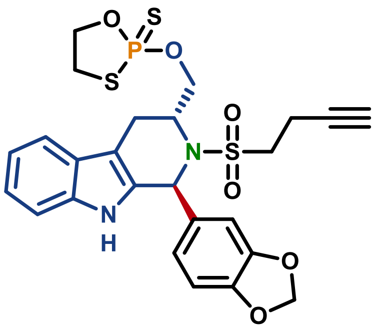

**2-(((1*S*,3*R*)-1-(benzo[*d*][1,3]dioxol-5-yl)-2-(but-3-yn-1-ylsulfonyl)-2,3,4,9-tetrahydro-1*H*-pyrido[3,4-*b*]indol-3-yl)methoxy)-1,3,2-oxathiaphospholane 2-sulfide (20, MB-6B)**

Utilizing procedure reported for **17**, compound **20** was prepared from **S2D**, to afford 21 mg of the titled product (**Yield** = 30%, ~1:1 d.r., 98% e.e.).

**Optical rotation:** $[\alpha]_D^{25}$ +25.5° (*c* 0.44, $CHCl_3$)

*Note: other characterizations were identical to **19***

#### Compound (21, MB-7A).

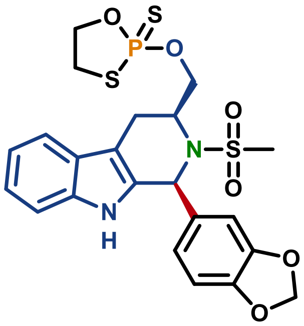

**2-(((1*S*,3*S*)-1-(benzo[*d*][1,3]dioxol-5-yl)-2-(methylsulfonyl)-2,3,4,9-tetrahydro-1*H*-pyrido[3,4-*b*]indol-3-yl)methoxy)-1,3,2-oxathiaphospholane 2-sulfide (21, MB-7A)**

Utilizing procedure reported for **17**, compound **21** was prepared from the corresponding crude alcohol (37 mg, 0.092 mmol, 1 equiv), obtained from reduction of **S3A**, to afford 39 mg of the titled product (**Yield** = 78%, ~1:1 d.r., >99%% e.e.).

*Note: Product was isolated via silica PTLC with 20% EtOAc in PhMe.*

**Physical state:** light yellow film

**$R_f$:** 0.55 (50% EtOAc in hexanes), UV active

**Optical rotation:** $[\alpha]_D^{25}$ +18.2° (*c* 0.85, $CHCl_3$)

**$^1$H NMR (400 MHz, $CDCl_3$) δ** 7.87 (s, 1H), 7.56 (d, *J* = 7.8 Hz, 1H), 7.35 (dd, *J* = 8.1, 1.0 Hz, 1H), 7.28 – 7.22 (m, 1H)(*overlapped with $CHCl_3$ peak*), 7.21 – 7.15 (m, 1H), 6.95 (t, *J* = 2.0 Hz, 1H), 6.82 – 6.78 (m, 1H), 6.75 (dd, *J* = 8.1, 1.5 Hz, 1H), 6.24 (d, *J* = 6.6 Hz, 1H), 5.99 – 5.95 (m, 2H), 4.78 (dq, *J* = 14.4, 7.3 Hz, 1H), 4.44 – 4.27 (m, 2H), 4.03 (dddd, *J* = 26.8, 11.1, 10.4, 7.7 Hz, 1H), 3.85 (ddt, *J* = 11.9, 10.4, 7.7 Hz, 1H), 3.46 – 3.27 (m, 2H), 3.15 (dd, *J* = 16.2, 6.7 Hz, 1H), 3.02 (dd, *J* = 16.6, 5.7 Hz, 1H), 2.85 (s, 1.5H), 2.84 (s, 1.5H).

**$^{13}$C NMR (151 MHz, $CDCl_3$) δ** 148.44, 148.42, 148.05, 148.01, 136.42, 133.26, 128.78, 126.87, 126.86, 123.14, 123.12, 122.27, 122.26, 120.23, 120.22, 118.72, 118.67, 111.33, 111.30, 109.29, 108.46, 108.45, 107.37, 107.26, 101.54, 70.62, 68.78, 68.73, 54.04, 53.93, 52.06, 52.02 – 52.01 (m), 51.99, 51.95, 41.02, 40.94, 36.64, 36.63, 36.60, 36.60, 21.97, 21.92.

**$^{31}$P NMR (162 MHz, $CDCl_3$) δ** 103.6, 103.4.

**HRMS (ESI-TOF) m/z**: Calcd for $C_{22}H_{24}N_2O_6PS_3$ $[M+H]^+$ 539.0534; Found 539.0545

#### Compound (23, MB-8A).

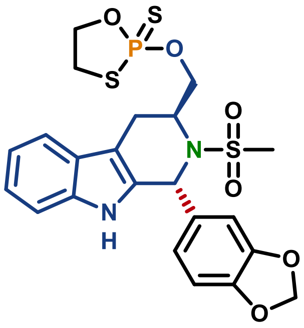

**2-(((1*R*,3*S*)-1-(benzo[*d*][1,3]dioxol-5-yl)-2-(methylsulfonyl)-2,3,4,9-tetrahydro-1*H*-pyrido[3,4-*b*]indol-3-yl)methoxy)-1,3,2-oxathiaphospholane 2-sulfide (23, MB-8A)**

Utilizing procedure reported for **17**, compound **23** was prepared from the corresponding crude alcohol (40 mg, 0.100 mmol, 1 equiv), obtained from reduction of **S3B,** to afford 40 mg of the titled product (**Yield** = 74%, **d.r.** ~1:1, >99% e.e.).

*Note: Product was isolated via silica PTLC with 20% EtOAc in PhMe.*

**Physical state:** light yellow film

**$R_f$:** 0.58 (50% EtOAc in hexanes), UV active

**Optical rotation:** $[\alpha]_D^{25}$ −26.8° (*c* 0.72, $CHCl_3$)

**$^1$H NMR (400 MHz, $CDCl_3$) δ** 7.74 (d, *J* = 9.1 Hz, 1H), 7.56 (dd, *J* = 7.9, 4.3 Hz, 1H), 7.30 (dd, *J* = 8.1, 2.5 Hz, 1H), 7.24 – 7.20 (m, 1H), 7.16 (t, *J* = 7.5 Hz, 1H), 6.82 – 6.70 (m, 3H), 5.98 – 5.91 (m, 3H), 4.57 – 4.21 (m, 5H), 3.57 – 3.30 (m, 2H), 3.18 (td, *J* = 15.2, 4.6 Hz, 1H), 3.12 – 3.01 (m, 1H), 2.64 (s, 1.5H), 2.62 (s, 1.5H).

**$^{13}$C NMR (151 MHz, $CDCl_3$) δ** 148.31, 148.24, 148.23, 136.73, 136.71, 132.21, 132.20, 131.39, 131.31, 126.65, 126.64, 123.22, 123.19, 122.98, 122.97, 120.24, 120.23, 118.79, 118.71, 111.34, 111.31, 109.44, 109.43, 108.44, 108.42, 108.31, 108.30, 101.58, 101.56, 70.70, 70.64, 68.64, 68.59, 68.41, 68.36, 57.88, 53.80, 53.74, 53.68, 42.87, 42.84, 36.69, 36.59, 23.64, 23.63.

**$^{31}$P NMR (162 MHz, $CDCl_3$) δ** 103.7, 103.5.

**HRMS (ESI-TOF) m/z**: Calcd for $C_{22}H_{24}N_2O_6PS_3$ $[M+H]^+$ 539.0534; Found 539.0538

#### Compound (22, MB-7B).

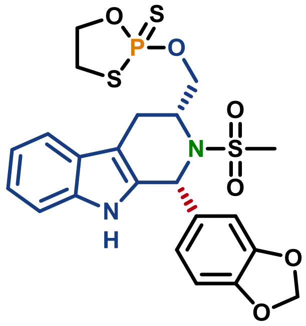

**2-(((1*R*,3*R*)-1-(benzo[*d*][1,3]dioxol-5-yl)-2-(methylsulfonyl)-2,3,4,9-tetrahydro-1*H*-pyrido[3,4-*b*]indol-3-yl)methoxy)-1,3,2-oxathiaphospholane 2-sulfide (22, MB-7B)**

Utilizing procedure reported for **17**, compound **22** was prepared from the corresponding crude alcohol (40 mg, 0.100 mmol, 1 equiv), obtained from reduction of **S3C**, to afford 44 mg of the titled product (**Yield** = 82%, **d.r.** ~1:1, >99% e.e.).

**Optical rotation:** $[\alpha]_D^{25}$ −21.3° (*c* 0.38, $CHCl_3$)

*Note: other characterizations were identical to **20***

#### Compound (24, MB-8B).

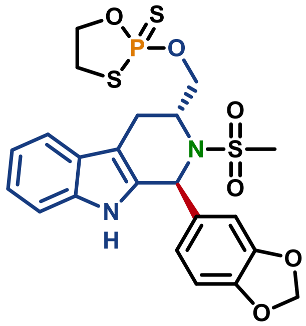

**2-(((1*S*,3*R*)-1-(benzo[*d*][1,3]dioxol-5-yl)-2-(methylsulfonyl)-2,3,4,9-tetrahydro-1*H*-pyrido[3,4-*b*]indol-3-yl)methoxy)-1,3,2-oxathiaphospholane 2-sulfide (24, MB-8A)**

Utilizing procedure reported for **17**, compound **24** was prepared from the corresponding crude alcohol (40 mg, 0.100 mmol, 1 equiv), obtained from reduction of **S3D**, to afford 37 mg of the titled product (**Yield** = 69%, ~1:1 d.r., 99% e.e.).

*Note: Product was isolated via silica PTLC with 20% EtOAc in PhMe.*

**Optical rotation:** $[\alpha]_D^{25}$ +25.0° (*c* 0.34, $CHCl_3$)

*Note: other characterizations were identical to **22***

##### 2.2 General Scheme for synthesis of 3-aryl-azetidine based Probes and Competitors.

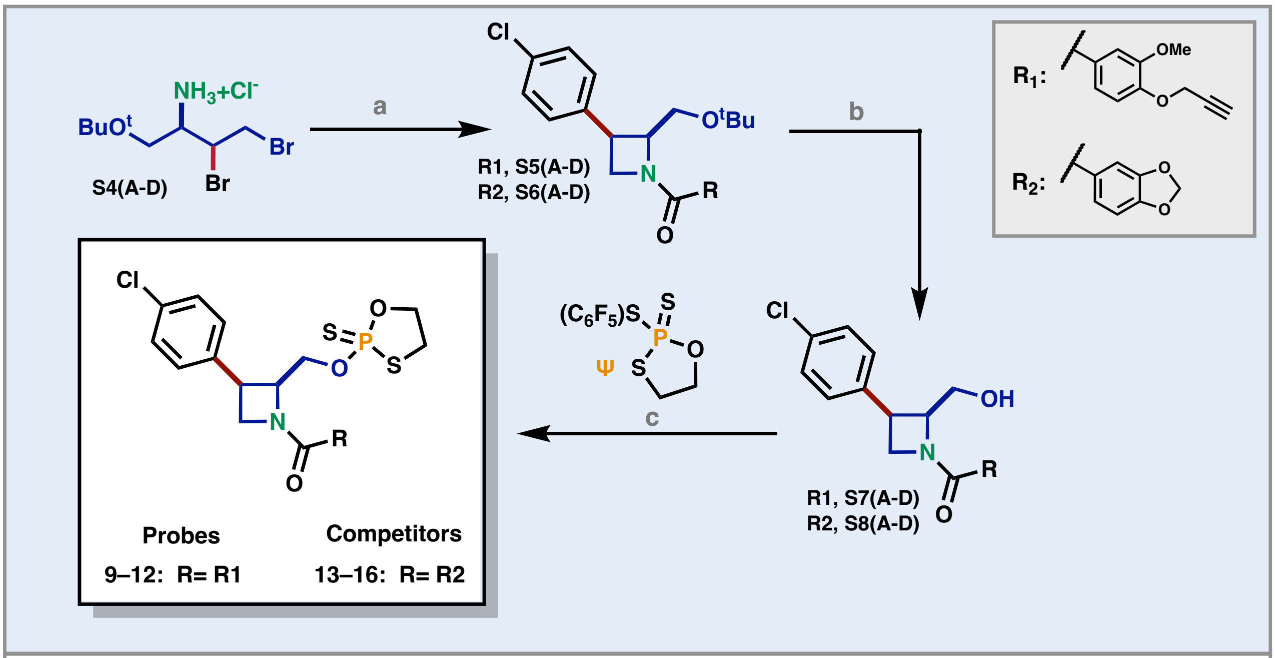

**Reaction conditions:** a. (i) MeLi, THF, -78 °C, 1.5 h; (ii) 4-ClPhMgCl, -78 °C —> rt, 18h; (iii) HATU b. 85% $H_3PO_4$, DCM, 35 °C, 16 h; c. DBU, MeCN, 30 min;

##### General Scheme for synthesis of S5 and S6.

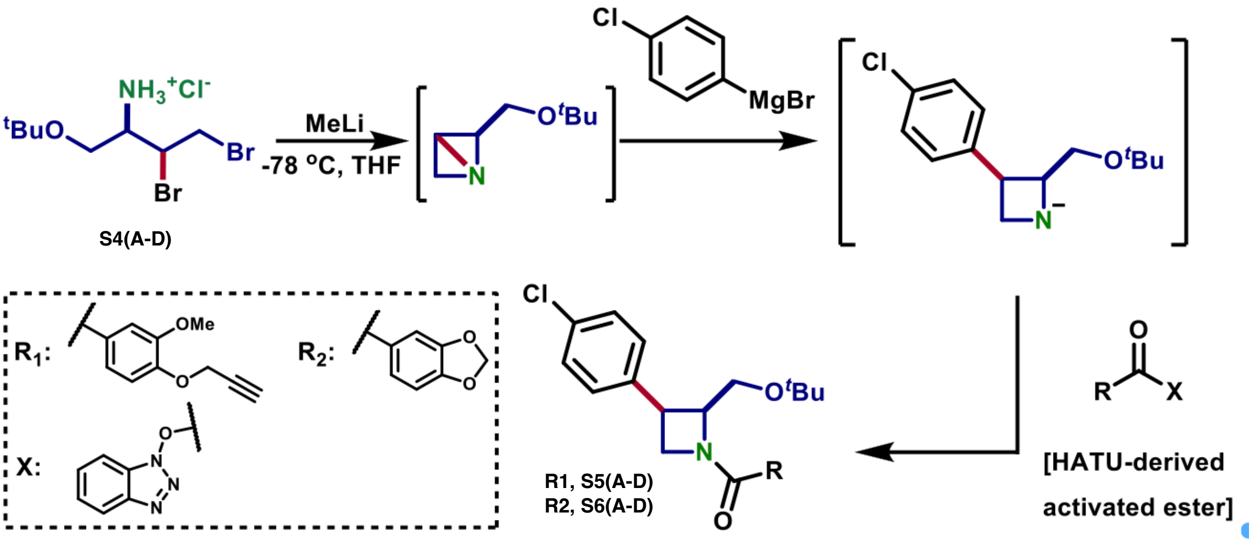

#### Compound (S4A).

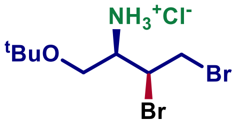

**(2*R*,3*S*)-2-amino-3,4-dibromobutan-1-ol *Tert*-Butyl Ether Hydrochloride (S4A)**

Compound **S4A** was prepared according to the procedure described in Bielecki et al. 2024 (10.26434/chemrxiv-2024-cmvd4). All characterizations matched the previously reported.

**Physical state:** white solid

**$R_t$:** 0.78 min

**$^1$H NMR (400 MHz, $CDCl_3$)** δ 8.87 (s, 3H), 4.58 (s, 1H), 4.36 (t, *J* = 10.3 Hz, 1H), 3.95 – 3.83 (m, 2H), 3.82 – 3.76 (m, 1H), 3.67 – 3.59 (m, 1H), 1.25 (s, 3H)

**LRMS (ESI-TOF) m/z**: Calcd for $C_8H_{18}Br_2NO$ $[M+H]^+$ 303.97; Found 303.98

#### Compound (S4B).

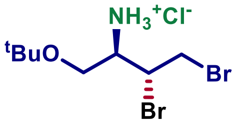

**(2*R*,3*R*)-2-amino-3,4-dibromobutan-1-ol *Tert*-Butyl Ether Hydrochloride (S4B)**

Compound **S4B** was prepared according to the procedure described in the in Bielecki et al. 2024 (10.26434/chemrxiv-2024-cmvd4). Characterizations matched the previously reported.

**Physical state:** white solid

**$R_t$:** 0.79 min

**$^1$H NMR (400 MHz, $CDCl_3$) δ** 8.81 (s, 3H), 4.82 (dt, *J* = 9.5, 5.1 Hz, 1H), 4.01 – 3.87 (m, 3H), 3.80 (dd, *J* = 9.7, 5.0 Hz, 1H), 3.73 (dd, *J* = 9.7, 6.4 Hz, 1H).

**LRMS (ESI-TOF) m/z**: Calcd for $C_8H_{18}Br_2NO$ $[M+H]^+$ 303.97; Found 303.98

#### Compound (S4C).

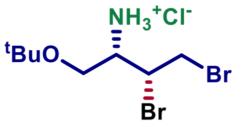

**(2$S$,3$R$)-2-amino-3,4-dibromobutan-1-ol *Tert*-Butyl Ether Hydrochloride (S4C)**

Compound **S4C** was prepared according to the procedure described in Bielecki et al. 2024 (10.26434/chemrxiv-2024-cmvd4). All characterizations matched the previously reported.

**Physical state:** white solid

**$R_t$:** 0.79 min

**$^1$H NMR (600 MHz, $CDCl_3$) δ** 8.88 (s, 3H), 4.58 (ddd, $J$ = 9.6, 5.2, 3.0 Hz, 1H), 4.36 (dd, $J$ = 11.3, 9.6 Hz, 1H), 3.93 – 3.86 (m, 2H), 3.78 (dd, $J$ = 9.4, 5.4 Hz, 1H), 3.64 (dd, $J$ = 9.4, 6.7 Hz, 1H), 1.24 (s, 9H).

**LRMS (ESI-TOF) m/z**: Calcd for $C_8H_{18}Br_2NO$ $[M+H]^+$ 303.97; Found 303.98

#### Compound (S4D).

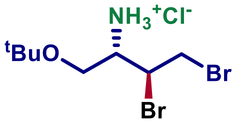

**(2$S$,3$S$)-2-amino-3,4-dibromobutan-1-ol *Tert*-Butyl Ether Hydrochloride (S4D)**

Compound **S8B** was prepared according to the procedure described in Bielecki et al. 2024 (10.26434/chemrxiv-2024-cmvd4). All characterizations matched the previously reported.

**Physical state:** white solid

**$R_t$:** 0.79 min

**$^1$H NMR (600 MHz, $CDCl_3$) δ** 8.82 (s, 3H), 4.81 (dtd, $J$ = 13.7, 5.4, 5.1, 1.1 Hz, 1H), 3.97 (dd, $J$ = 11.1, 5.5 Hz, 1H), 3.94 – 3.89 (m, 2H), 3.80 (dd, $J$ = 9.7, 5.2 Hz, 1H), 3.73 (dd, $J$ = 9.7, 6.3 Hz, 1H), 1.26 (s, 9H).

**LRMS (ESI-TOF) m/z**: Calcd for $C_8H_{18}Br_2NO$ $[M+H]^+$ 303.97; Found 303.97

#### Compound (S5A).

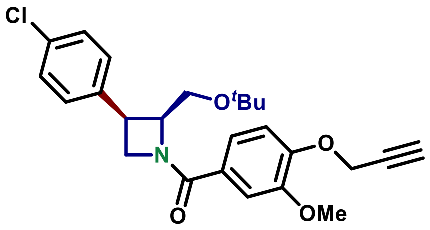

**((2$S$,3$R$)-3-(4-chlorophenyl)-2-(hydroxymethyl)azetidin-1-yl)(3-methoxy-4-(prop-2-yn-1-yloxy)phenyl)methanone *Tert*-butyl Ether (S5A)**

A flamed dried 10 mL glass culture tube equipped with a stir bar was charged with **S4A** (100 mg, 0.30 mmol, 1 equiv) and dry PhMe (3.20 mL). The tube was evacuated and back filled with Ar three times and chilled –78°C in solid $CO_2$ bath. 2.5 M ***n*BuLi** solution in hexanes (0.38 mL, 0.94 mmol, 3.2 equiv) was added dropwise and the mixture was stirred for 2 h at –78°C. **(4-chlorophenyl)magnesium bromide** as 0.95M THF solution (0.62 mL, 0.59 mmol, 2 equiv)(see *note #1*) was added and the reaction mixture was removed from the cooling bath and left stirring overnight at room temperature. The reaction was quenched with a 0.2 M **HATU**-derived ester of **3-methoxy-4-(prop-2-yn-1-yloxy)benzoic acid** (see *note #2*) solution in THF (2.2 mL, 0.44 mmol, 1.5 equiv) (see *note #3*). The solvent was evaporated and replaced with MeOH. $K_2CO_3$ (excess) was added and the suspension was stirred vigorously at room temperature for 1 h to decompose the unreacted activated ester (see *note #4*). The solvent was removed on a rotary evaporator and the yellow orange residue was taken up in EtOAc and filtered. The filtrate was washed with 1 M HCl (×1) followed by brine-$H_2O$ (1:1, ×2). The organic layer was dried with $MgSO_4$, and the solvent was removed *in vacuo.* The crude product was isolated by silica chromatography (30% EtOAc in hexanes) to give 33 mg (**Yield** = 25%, **d.r.** > 20:1)

*Note #1: (4-chlorophenyl)magnesium bromide was prepared according to previously reported procedure in Bielecki et al..*

*Note #2: 3-methoxy-4-(prop-2-yn-1-yloxy)benzoic acid was prepared according to a previously reported procedure in Joseph et al..*

*Note #3: The activated ester was prepared by combining HATU (0.92 g, 2.4 mmol, 1 equiv) with 3-methoxy 3-methoxy-4-(prop-2-yn-1-yloxy)benzoic acid (0.50 g, 2.4 mmol, 1 equiv) in dry THF (10 mL). Dry DIPEA (0.31 g, 0.42 mL, 2.4 mmol, 1 equiv) was added dropwise followed by dry THF to a final volume of ~12 mL.*

*Note #4: Otherwise, the unreacted activated ester partially co-eluted with product during separation.*

**Physical state:** colorless film

**$^1$H NMR (600 MHz, $CDCl_3$) δ** 7.37 – 7.28 (m, 3H), 7.28 – 7.21 (m, 3H)(*overlapped with $CHCl_3$ peak*), 7.02 (d, $J$ = 8.3 Hz, 1H), 4.89 (dt, $J$ = 9.4, 4.7 Hz, 1H), 4.80 (d, $J$ = 2.5 Hz, 2H), 4.58 (t, $J$ = 8.1 Hz, 1H), 4.36 (br s, 1H), 4.10 (br s, 1H), 3.91 (s, 3H), 3.72 – 2.83 (m, 2H), 2.52 (s, 1H), 0.78 (s, 9H).

**$^{13}$C NMR (151 MHz, $CDCl_3$) δ** 170.5, 149.6, 149.1, 136.0, 132.7, 130.1, 128.2, 120.7, 113.0, 111.9, 78.1, 76.3, 72.7, 64.6(broad), 58.2, 56.8, 56.2, 37.1, 27.0.

**LRMS (ESI-TOF) m/z**: Calcd for $C_{25}H_{29}ClNO_4$ $[M+H]^+$ 442.18; Found 442.24

#### Compound (S5B).

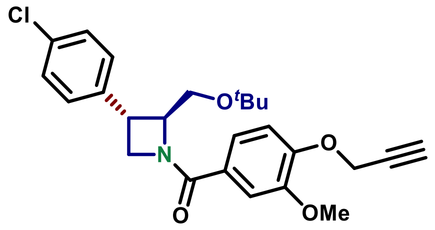

**((2*S*,3*S*)-3-(4-chlorophenyl)-2-(hydroxymethyl)azetidin-1-yl)(3-methoxy-4-(prop-2-yn-1-yloxy)phenyl)methanone *Tert*-butyl Ether (S5B)**

Utilizing procedure reported for **S5A**, compound **S5B** was prepared from **S4B** (100 mg, 0.30 mmol, 1 equiv) to afford 52 mg (**Yield** = 40%, **d.r.** > 20:1).

*Note #1: The product was isolated by silica PTLC (50% EtOAc in hexanes)*

**Physical state:** colorless film

**Optical rotation:** $[\alpha]_D^{25}$ –47.9° (*c* 2.5, $CHCl_3$)

**$^1$H NMR (600 MHz, $CDCl_3$) δ** 7.39 – 7.27 (m, 3H), 7.26 – 7.14 (m, 2H)(*overlapped with $CHCl_3$ peak*), 7.01 (d, *J* = 8.3 Hz, 1H), 4.80 (d, *J* = 2.4 Hz, 2H), 4.74 – 4.49 (m, 2H), 4.10 (br s, 1H), 4.00 – 3.64 (m, 5H), 2.52 (s, 1H), 1.22 (br s, 9H).

**$^{13}$C NMR (151 MHz, $CDCl_3$) δ** 170.1, 149.7, 149.2, 140.4, 132.8, 129.0, 128.4, 127.2, 120.7, 113.0, 111.9, 78.1, 76.4, 73.3, 69.2, 61.7, 58.5, 56.8, 56.2, 37.3, 27.7.

**HRMS (ESI-TOF) m/z**: Calcd for $C_{25}H_{29}ClNO_4$ $[M+H]^+$ 442.1785; Found 442.1795

#### Compound (S5C).

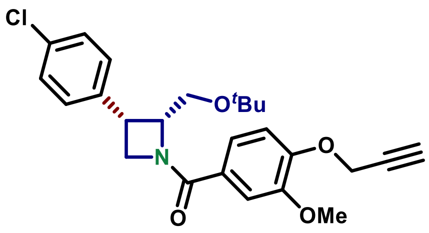

**((2*R*,3*S*)-3-(4-chlorophenyl)-2-(hydroxymethyl)azetidin-1-yl)(3-methoxy-4-(prop-2-yn-1-yloxy)phenyl)methanone *Tert*-butyl Ether (S5C)**

Utilizing procedure reported for **S5A**, compound **S5C** was prepared from **S4C** (100 mg, 0.30 mmol, 1 equiv) to afford 45 mg of the titled product (**Yield** = 35%, **d.r.** 20:1).

**Physical state:** colorless film

**$^1$H NMR (600 MHz, $CDCl_3$) δ** 7.35 – 7.29 (m, 3H), 7.28 – 7.21 (m, 3H) )(*overlapped with $CHCl_3$ peak*), 7.02 (d, *J* = 8.3 Hz, 1H), 4.89 (dt, *J* = 9.3, 4.8 Hz, 1H), 4.80 (d, *J* = 2.4 Hz, 2H), 4.58 (t, *J* = 8.1 Hz, 1H), 4.36 (br s, 1H), 4.10 (br s, 1H), 3.91 (s, 3H), 3.64 – 2.83 (m, 2H), 2.52 (s, 1H), 0.78 (s, 9H).

**$^{13}$C NMR (151 MHz, $CDCl_3$) δ** 170.5, 149.6, 149.1, 136.0, 132.7, 130.1, 128.2, 120.7, 113.0, 111.9, 78.1, 76.6 (d, *J* = 90.9 Hz), 72.7, 64.8, 58.2, 56.8, 56.2, 37.1, 27.0.

**LRMS (ESI-TOF) m/z**: Calcd for $C_{25}H_{29}ClNO_4$ $[M+H]^+$ 442.18; Found 442.23

#### Compound (S5D).

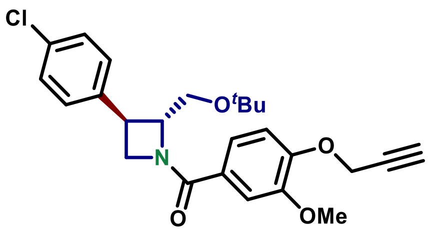

**((2*R*,3*R*)-3-(4-chlorophenyl)-2-(hydroxymethyl)azetidin-1-yl)(3-methoxy-4-(prop-2-yn-1-yloxy)phenyl)methanone *Tert*-butyl Ether (S5D)**

Utilizing procedure reported for **S5A**, compound **S5D** was prepared from **S4D** (100 mg, 0.30 mmol, 1 equiv) to afford 65 mg of the titled product (**Yield** = 50%, **d.r.** 20:1).

**Optical rotation:** $[\alpha]_D^{25}$ +50.9° (*c* 1.4, $CHCl_3$)

*Note: Other characterizations were identical to compound **S5B.***

#### Compound (S6A).

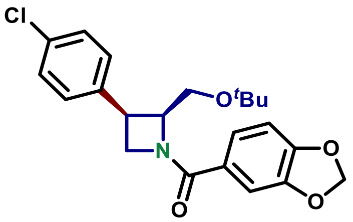

**Benzo[*d*][1,3]dioxol-5-yl((2*S*,3*R*)-3-(4-chlorophenyl)-2-(hydroxymethyl)azetidin-1-yl)methanone *Tert*-Butyl Ether (S6A)**

Utilizing procedure reported for **S5A**, compound **S5A** was prepared from **S4A** (100 mg, 0.30 mmol, 1 equiv) and 0.2 M THF stock solution of the **HATU**-derived ester of **piperonylic acid** (2.2 mL, 0.44 mmol, 1.5 equiv) (see *note #1*) to afford 28 mg of the titled product (**Yield** = 24%, **d.r.** > 20:1).

*Note #1: The activated ester was prepared by combining HATU (0.92 g, 2.4 mmol, 1 equiv) with piperonylic acid (0.40 g, 2.4 mmol, 1 equiv) in dry THF (10 mL). Dry DIPEA (0.31 g, 0.42 mL, 2.4 mmol, 1 equiv) was added dropwise, followed by dry THF to a final volume of ~11.0 mL.*

*Note # 2: The titled compound was isolated by PTLC with 50% EtOAc in hexanes.*

**Physical state:** colorless film

**$^1$H NMR (600 MHz, $CDCl_3$) δ** 7.31 (d, *J* = 8.1 Hz, 2H), 7.28 – 7.21 (m, 3H)(*overlapped with $CHCl_3$ peak*), 7.18 (s, 1H), 6.83 (d, *J* = 8.6 Hz, 1H), 6.01 (d, *J* = 3.3 Hz, 2H), 4.87 (dt, *J* = 9.3, 4.8 Hz, 1H), 4.56 (t, *J* = 8.1 Hz, 1H), 4.32 (br s, 1H), 4.09 (br s, 1H), 3.77 – 2.87 (br m, 2H), 0.78 (s, 9H)

**$^{13}$C NMR (151 MHz, $CDCl_3$) δ** 170.5, 150.0, 147.8, 136.0, 132.7, 130.1, 128.2, 122.8, 108.5, 108.1, 101.7, 72.7, 64.8, 58.2, 56.9, 37.0, 27.0.

**LRMS (ESI-TOF) m/z**: Calcd for $C_{22}H_{25}ClNO_4$ $[M+H]^+$ 402.15; Found 402.21

#### Compound (S6B).

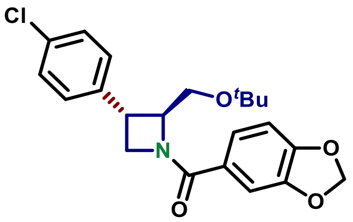

**Benzo[*d*][1,3]dioxol-5-yl((2*S*,3*S*)-3-(4-chlorophenyl)-2-(hydroxymethyl)azetidin-1-yl)methanone *Tert*-Butyl Ether (S6B)**

Utilizing procedure reported for **S5A**, compound **S6B** was prepared from **S4B** (100 mg, 0.30 mmol, 1 equiv) and 0.2 M THF stock solution of the **HATU**-derived ester of **piperonylic acid** (2.2 mL, 0.44 mmol, 1.5 equiv) to afford 61 mg of the titled product (**Yield** = 52%, **d.r.** > 20:1).

**Physical state:** colorless film

**$^1$H NMR (600 MHz, $CDCl_3$) δ** 7.39 – 7.10 (m, 6H) (*overlapped with $CHCl_3$ peak*), 6.82 (d, *J* = 7.9 Hz, 1H), 6.01 (d, *J* = 2.3 Hz, 2H), 4.64 (br s, 1H), 4.55 (br s, 1H), 4.05 (br s, 1H), 3.95 – 3.13 (br m, 3H), 1.19 (s, 9H).

**$^{13}$C NMR (151 MHz, $CDCl_3$) δ** 170.1, 150.0, 147.8, 140.5, 132.8, 129.0, 128.4, 122.8, 108.5, 108.1, 101.7, 73.3, 69.1, 61.6, 58.4, 37.1, 27.7.

**LRMS (ESI-TOF) m/z**: Calcd for $C_{22}H_{25}ClNO_4$ $[M+H]^+$ 402.15; Found 402.21

#### Compound (S6C).

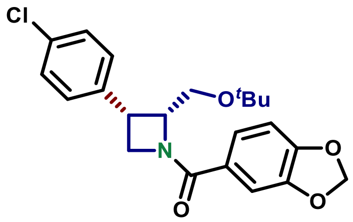

**Benzo[*d*][1,3]dioxol-5-yl((2*S*,3*R*)-3-(4-chlorophenyl)-2-(hydroxymethyl)azetidin-1-yl)methanone *Tert*-Butyl Ether (S6C)**

Utilizing procedure reported for **S5A**, compound **S6C** was prepared from **S4C** (100 mg, 0.30 mmol, 1 equiv) to afford 33 mg of the titled product (**Yield** = 28%, **d.r.** > 20:1).

**Physical state:** colorless film

**$^1$H NMR (600 MHz, $CDCl_3$) δ** 7.31 (d, *J* = 8.1 Hz, 2H), 7.28 – 7.21 (m, 3H) )(*overlapped with $CHCl_3$ peak*), 7.18 (s, 1H), 6.83 (d, *J* = 8.1 Hz, 1H), 6.01 (d, *J* = 3.3 Hz, 2H), 4.87 (dt, *J* = 9.3, 4.8 Hz, 1H), 4.56 (t, *J* = 8.1 Hz, 1H), 4.32 (br s, 1H), 4.09 (br s, 1H), 3.66 – 2.83 (br m, 2H), 0.78 (s, 9H).

**$^{13}$C NMR (151 MHz, $CDCl_3$) δ** 170.5, 150.0, 147.8, 136.0, 132.7, 130.1, 128.2, 122.8, 108.5, 108.1, 101.7, 72.7, 58.2, 37.0, 27.0.

#### Compound (S6D).

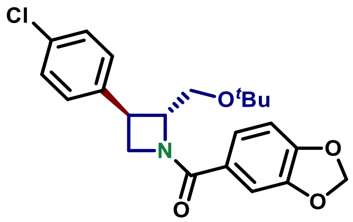

**Benzo[*d*][1,3]dioxol-5-yl((2*R*,3*S*)-3-(4-chlorophenyl)-2-(hydroxymethyl)azetidin-1-yl)methanone *Tert*-Butyl Ether (S6D)**

Utilizing procedure reported for **S5A**, compound **S6D** was prepared from **S4D** (100 mg, 0.30 mmol, 1 equiv) to afford 63 mg (**Yield** = 53%, **d.r.** > 20:1) of the titled product

**Physical state:** colorless film

**$^1$H NMR (600 MHz, CDCl$_3$) δ** 7.35 – 7.10 (m, 6H)(*overlapped with CHCl$_3$ peak*), 6.82 (d, *J* = 8.1 Hz, 1H), 6.01 (d, *J* = 2.3 Hz, 1H), 4.64 (br s, 1H), 4.55 (br s, 1H), 4.04 (br s, 1H), 3.96 – 3.19 (br m, 3H), 1.19 (s, 9H).

**$^{13}$C NMR (151 MHz, CDCl$_3$) δ** 170.1, 150.0, 147.8, 140.5, 132.8, 129.0, 128.4, 122.9, 108.5, 108.1, 101.7, 73.3, 69.1, 61.6, 58.4, 37.1, 27.7.

**LRMS (ESI-TOF) m/z**: Calcd for $C_{22}H_{25}ClNO_4$ $[M+H]^+$ 402.15; Found 402.23

#### Compound (S7A).

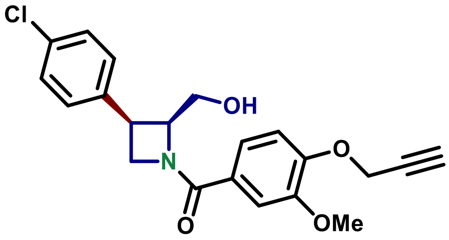

**((2*S*,3*R*)-3-(4-chlorophenyl)-2-(hydroxymethyl)azetidin-1-yl)(3-methoxy-4-(prop-2-yn-1-yloxy)phenyl)methanone (S7A)**

A 4 dram glass vial equipped with a stir bar was charged with **S5A** (33 mg, 0.075 mmol, 1 equiv), DCM (0.20 mL) and 85% $H_3PO_4$ (0.20 mL). The reaction mixture was vigorously stirred overnight at 35 °C. The reaction mixture was diluted with EtOAc, washed with brine-water (1:1, ×1), aq. $NaHCO_3$ (×2), dried with $MgSO_4$ and the solvent was removed *in vacuo*. The crude product was purified by PTLC with 30% acetone in DCM to afford 16 mg of the titled product (**Yield** = 56%, **d.r.** > 20:1)

**Physical state:** colorless film

**R$_f$:** 0.15 (50% EtOAc in hexanes), UV active

**$^1$H NMR (600 MHz, CDCl$_3$) δ** 7.38 (d, *J* = 2.0 Hz, 1H), 7.36 – 7.34 (m, 1H), 7.34 – 7.32 (m, 1H), 7.28 – 7.26 (m, 1H), 7.26 – 7.25 (m, 1H), 7.25 – 7.24 (m, 1H), 7.06 (d, *J* = 8.4 Hz, 1H), 5.00 (t, *J* = 9.3 Hz, 1H), 4.83 (d, *J* = 2.4 Hz, 2H), 4.77 – 4.69 (m, 2H), 4.56 (ddd, *J* = 9.4, 5.6, 1.1 Hz, 1H), 4.00 – 3.93 (m, 1H), 3.93 (s, 3H), 3.62 (t, *J* = 11.0 Hz, 1H), 3.24 (t, *J* = 10.9 Hz, 1H), 2.54 (t, *J* = 2.4 Hz, 1H).

**$^{13}$C NMR (151 MHz, CDCl$_3$) δ** 170.8, 149.8, 149.7, 135.0, 133.7, 129.4, 129.0, 126.2, 120.8, 112.9, 112.1, 78.0, 76.5, 68.8, 64.5, 57.0, 56.8, 56.2, 36.5.

**HRMS (ESI-TOF) m/z**: Calcd for $C_{21}H_{21}ClNO_4$ $[M+H]^+$ 386.1159; Found 386.1165

#### Compound (S7B).

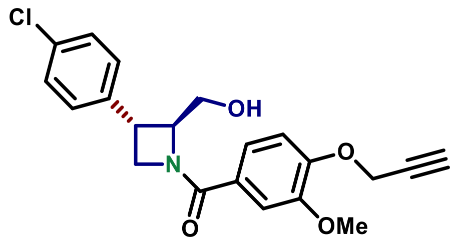

**((2*S*,3*S*)-3-(4-chlorophenyl)-2-(hydroxymethyl)azetidin-1-yl)(3-methoxy-4-(prop-2-yn-1-yloxy)phenyl)methanone (S7B)**

Utilizing procedure reported for **S7A**, compound **S7B** was prepared from **S5B** (52 mg, 0.12 mmol) to afford 32 mg of the titled product (**Yield** = 70%, **d.r.** > 20:1).

**Physical state:** colorless film

**$R_f$:** 0.13 (50% EtOAc in hexanes), UV active

**$R_t$:** 1.44 min

**Optical rotation:** $[\alpha]_D^{25}$ –62.7° (*c* 0.38, $CHCl_3$)

**$^1$H NMR (600 MHz, $CDCl_3$) δ** 7.35 (s, 1H), 7.33 (d, *J* = 8.4 Hz, 2H), 7.23 (d, *J* = 8.2 Hz, 3H), 7.03 (d, *J* = 8.3 Hz, 1H), 4.88 – 4.72 (m, 4H), 4.66 (t, *J* = 9.0 Hz, 1H), 4.32 (t, *J* = 8.0 Hz, 1H), 4.00 – 3.94 (m, 1H), 3.94 – 3.83 (m, 4H), 3.54 (q, *J* = 7.7 Hz, 1H), 2.53 (t, *J* = 2.4 Hz, 1H).

7.35 (s, 1H), 7.33 (d, *J* = 8.4 Hz, 2H), 7.23 (d, *J* = 8.2 Hz, 3H), 7.03 (d, *J* = 8.3 Hz, 1H), 4.88 – 4.72 (m, 4H), 4.66 (t, *J* = 9.0 Hz, 1H), 4.32 (t, *J* = 8.0 Hz, 1H), 4.00 – 3.94 (m, 1H), 3.94 – 3.83 (m, 4H), 3.54 (q, *J* = 7.7 Hz, 1H), 2.53 (t, *J* = 2.4 Hz, 1H).

**$^{13}$C NMR (151 MHz, $CDCl_3$) δ** 170.6, 149.8, 149.6, 138.2, 133.5, 129.2, 128.4, 126.2, 120.8, 112.9, 112.0, 78.0, 76.5, 72.9, 66.0, 58.3, 56.8, 56.2, 36.8.

**HRMS (ESI-TOF) m/z**: Calcd for $C_{21}H_{21}ClNO_4$ $[M+H]^+$ 386.1159; Found 386.1165

#### Compound (S7C).

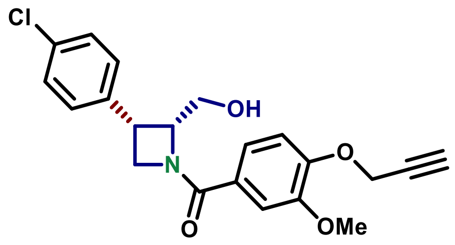

**((2*R*,3*S*)-3-(4-chlorophenyl)-2-(hydroxymethyl)azetidin-1-yl)(3-methoxy-4-(prop-2-yn-1-yloxy)phenyl)methanone (S7C)**

Utilizing procedure reported for **S7A**, compound **S7C** was prepared from **S5C** (14 mg, 0.032 mmol, 1 equiv) to afford 9 mg of the titled product (**Yield** = 74%, **d.r.** > 20:1).

**Physical state:** colorless film

**$R_f$:** 0.15 (50% EtOAc in hexanes), UV active

**$R_t$:** 1.48 min

**$^1$H NMR (600 MHz, $CDCl_3$) δ** 7.38 (d, *J* = 1.5 Hz, 1H), 7.35 – 7.34 (m, 1H), 7.34 – 7.32 (m, 1H), 7.28 – 7.26 (m, 1H), 7.27 – 7.24 (m, 1H), 7.25 – 7.23 (m, 1H), 7.06 (d, *J* = 8.4 Hz, 1H), 5.00 (t, *J* = 9.1 Hz, 1H), 4.83 (d, *J* = 2.4 Hz, 2H), 4.77 – 4.70 (m, 1H), 4.56 (dd, *J* = 9.4, 5.6 Hz, 1H), 4.00 – 3.94 (m, 2H), 3.93 (s, 1H), 3.62 (t, *J* = 10.9 Hz, 1H), 3.24 (t, *J* = 10.9 Hz, 1H), 2.55 (t, *J* = 2.4 Hz, 1H).

**$^{13}$C NMR (151 MHz, $CDCl_3$) δ** 170.8, 149.8, 149.7, 135.0, 133.7, 129.4, 129.0, 126.2, 120.8, 112.9, 112.1, 78.0, 76.5, 68.8, 64.5, 57.0, 56.8, 56.2, 36.5.

**LRMS (ESI-TOF) m/z**: Calcd for $C_{21}H_{21}ClNO_4$ $[M+H]^+$ 386.12; Found 386.19

#### Compound (S7D).

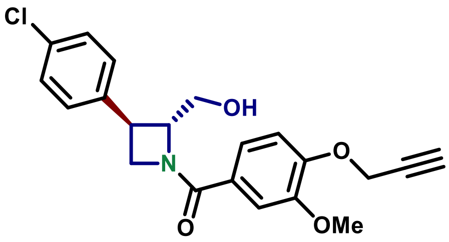

**((2*R*,3*R*)-3-(4-chlorophenyl)-2-(hydroxymethyl)azetidin-1-yl)(3-methoxy-4-(prop-2-yn-1-yloxy)phenyl)methanone (S7D)**

Utilizing procedure reported for **S7A**, compound **S7D** was prepared from **S5D** (65 mg, 0.147 mmol, 1 equiv) to afford 34 mg of the titled product (**Yield** = 60%, **d.r.** > 20:1).

**Optical rotation:** $[\alpha]_D^{25}$ +57.1° (*c* 0.56, $CHCl_3$)

*Note: Other characterizations were identical to compound **S7B.***

#### Compound (S8A).

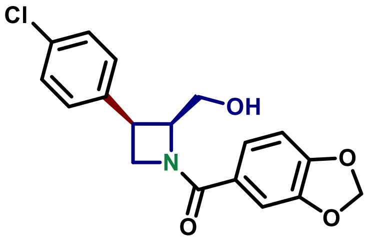

**Benzo[*d*][1,3]dioxol-5-yl((2*S*,3*R*)-3-(4-chlorophenyl)-2-(hydroxymethyl)azetidin-1-yl)methanone (S8A)**

Utilizing procedure reported for **S7A**, compound **S8A** was prepared from **S6A** (28 mg, 0.15 mmol, 1 equiv) to afford 12 mg of the titled product (**Yield** = 60%, **d.r.** > 20:1).

**Physical state:** colorless film

**$^1$H NMR (600 MHz, $CDCl_3$) δ** 7.35 – 7.34 (m, 1H), 7.34 – 7.32 (m, 1H), 7.27 (dd, *J* = 8.1, 1.7 Hz, 1H), 7.25 – 7.25 (m, 1H), 7.24 – 7.23 (m, 1H), 7.21 (d, *J* = 1.7 Hz, 1H), 6.87 (d, *J* = 8.1 Hz, 1H), 6.04 (s, 2H), 4.98 (t, *J* = 9.2 Hz, 1H), 4.70 – 4.62 (m, 2H), 4.52 (ddd, *J* = 9.4, 5.6, 1.1 Hz, 1H), 4.00 – 3.88 (m, 1H), 3.60 (t, *J* = 10.8 Hz, 1H), 3.24 (t, *J* = 11.1 Hz, 1H).

**$^{13}$C NMR (151 MHz, CDCl$_3$) δ** 170.8, 150.5, 148.0, 135.0, 133.7, 129.4, 129.0, 126.4, 123.1, 108.5, 108.3, 101.9, 68.6, 64.5, 56.9, 36.4.

**LRMS (ESI-TOF) m/z**: Calcd for $C_{18}H_{17}ClNO_4$ $[M+H]^+$ 346.08; Found 346.11

#### Compound (S8B).

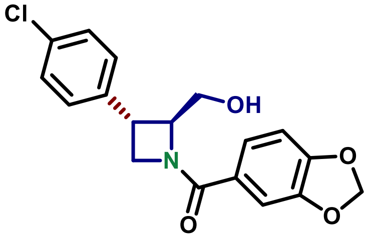

**Benzo[*d*][1,3]dioxol-5-yl((2*S*,3*S*)-3-(4-chlorophenyl)-2-(hydroxymethyl)azetidin-1-yl)methanone (S8B)**

Utilizing procedure reported for **S7A**, compound **S8B** was prepared from **S6B** (61 mg, 0.15 mmol, 1 equiv) to afford 52 mg of the titled product (**Yield** = 65%, **d.r.** > 20:1).

**Physical state:** colorless film

**$R_t$:** 1.41 min

**$R_f$:** 0.19 (50% EtOAc in hexanes), UV active

**Optical rotation:** $[\alpha]_D^{25}$ –58.4° (*c* 0.46, CHCl$_3$)

**$^1$H NMR (600 MHz, CDCl$_3$) δ** 7.33 (d, *J* = 8.4 Hz, 2H), 7.24 – 7.17 (m, 4H), 6.83 (d, *J* = 8.1 Hz, 1H), 6.02 (s, 2H), 4.83 – 4.71 (m, 2H), 4.61 (t, *J* = 9.0 Hz, 1H), 4.27 (t, *J* = 8.1 Hz, 1H), 4.03 – 3.82 (m, 2H), 3.57 – 3.49 (m, 1H).

**$^{13}$C NMR (151 MHz, CDCl$_3$) δ** 170.6, 150.5, 148.0, 138.2, 133.5, 129.2, 128.4, 126.4, 123.1, 108.5, 108.2, 101.9, 72.8, 65.9, 58.2, 36.6.

**HRMS (ESI-TOF) m/z**: Calcd for $C_{18}H_{17}ClNO_4$ $[M+H]^+$ 346.0846; Found 346.0851

#### Compound (S8C).

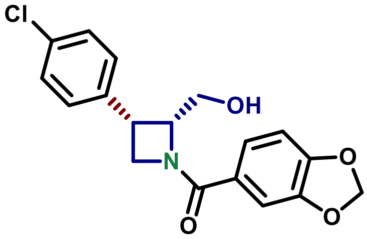

**Benzo[*d*][1,3]dioxol-5-yl((2*R*,3*S*)-3-(4-chlorophenyl)-2-(hydroxymethyl)azetidin-1-yl)methanone (S8C)**

Utilizing procedure reported for **S7A**, compound **S8C** was prepared from **S6C** (28 mg, 0.11 mmol, 1 equiv) to afford 16 mg of the titled product (**Yield** = 66%, **d.r.** > 20:1).

**Physical state:** colorless film

**$^1$H NMR (600 MHz, $CDCl_3$) δ** 7.35 – 7.34 (m, 1H), 7.34 – 7.32 (m, 1H), 7.27 (dd, *J* = 8.2, 1.7 Hz, 1H), 7.25 – 7.25 (m, 1H), 7.24 – 7.23 (m, 1H), 7.21 (d, *J* = 1.7 Hz, 1H), 6.87 (d, *J* = 8.1 Hz, 1H), 6.04 (s, 2H), 4.98 (t, *J* = 8.8 Hz, 1H), 4.71 – 4.61 (m, 2H), 4.52 (ddd, *J* = 9.4, 5.6, 1.1 Hz, 1H), 3.99 – 3.91 (m, 1H), 3.64 – 3.56 (m, 1H), 3.24 (t, *J* = 10.2 Hz, 1H).

**$^{13}$C NMR (151 MHz, $CDCl_3$) δ** 170.8, 150.5, 148.0, 135.0, 133.7, 129.4, 129.0, 126.5, 123.1, 108.5, 108.3, 101.9, 68.6, 64.5, 56.9, 36.4.

**LRMS (ESI-TOF) m/z**: Calcd for $C_{18}H_{17}ClNO_4$ $[M+H]^+$ 346.08; Found 346.14

#### Compound (S8D).

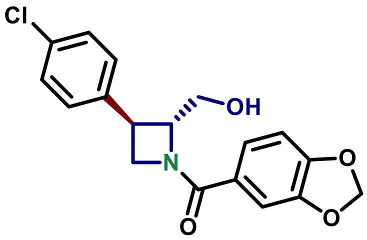

**Benzo[*d*][1,3]dioxol-5-yl((2*R*,3*R*)-3-(4-chlorophenyl)-2-(hydroxymethyl)azetidin-1-yl)methanone (S8D)**

Utilizing procedure reported for **S7A**, compound **S8D** was prepared from **S6D** (45 mg, 0.11 mmol, 1 equiv) to afford 22 mg of the titled product (**Yield** = 57%, **d.r.** > 20:1).

**Optical rotation:** $[\alpha]_D^{25}$ +63.3° (*c* 0.60, $CHCl_3$)

*Note: Other characterizations were identical to compound **S8B.***

#### Compound (9, MB-1A).

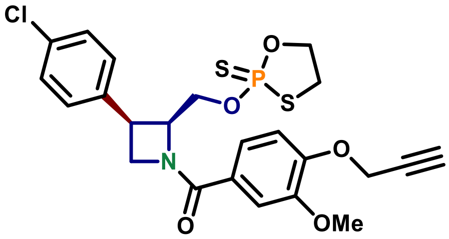

**((2*S*,3*R*)-3-(4-chlorophenyl)-2-(((2-sulfido-1,3,2-oxathiaphospholan-2-yl)oxy)methyl)azetidin-1-yl)(3-methoxy-4-(prop-2-yn-1-yloxy)phenyl)methanone (9, MB-1A)**

In a 4 dram glass vial equipped with a stir bar, **S7A** (20 mg, 0.052 mmol, 1 equiv), **Ψ** (19 mg, 0.057 mmol, 1.5 equiv) and powdered *3 Å molecular sieves* were combined in dry MeCN (0.52 mL) and stirred for 5 min. at room temperature. Dry **DBU** (8.7 mg, 8.5 µL, 0.057 mmol, 1.5 equiv) and stirred for 30 min. The reaction mixture was diluted with EtOAc, filtered through celite. The filtrate was washed with aq. $NaH_2PO_4$ (×2), aq. $Na_2HPO_4$ (×2), brine (×1) , dried with $MgSO_4$ and the solvent was removed *in vacuo*. The obtained crude product was purified by PTLC with 55% EtAcO in hexanes to afford 7.3 mg (**Yield** = 27%, **d.r.** ~1:1, 96% e.e.)

**Physical state:** colorless film

**$R_f$:** 0.22 (60% EtOAc in hexanes), UV active

**Optical rotation:** $[\alpha]_D^{25}$ +30.7° (*c* 0.73, $CHCl_3$)

**$^1$H NMR (600 MHz, $CDCl_3$) δ** 7.38 – 7.35 (m, 2H), 7.32 (d, *J* = 2.5 Hz, 1H), 7.30 – 7.23 (m, 3H), 7.04 (d, *J* = 8.3 Hz, 1H), 5.05 (ddt, *J* = 9.8, 6.9, 3.8 Hz, 1H), 4.81 (d, *J* = 2.4 Hz, 2H), 4.63 (q, *J* = 9.2, 8.2 Hz, 1H), 4.42 (t, *J* = 9.0 Hz, 1H), 4.34 – 4.02 (m, 5H), 3.92 (s, 3H), 3.34 (ddt, *J* = 19.3, 10.5, 5.3 Hz, 1H), 3.25 (s, 1H), 2.54 (t, *J* = 2.4 Hz, 1H).

**$^{13}$C NMR (151 MHz, $CDCl_3$) δ** 170.9, 170.7, 149.73, 149.7, 149.5, 134.8, 133.5, 129.8, 129.6, 128.97, 128.9, 126.9, 120.9, 113.0, 111.9, 78.0, 76.5, 70.5, 65.42, 65.36, 65.2, 65.1, 63.6, 56.9, 56.24, 56.23, 37.0, 36.8, 36.57, 36.56.

**$^{31}$P NMR (162 MHz, $CDCl_3$) δ** 103.4, 103.0

**HRMS (ESI-TOF) m/z**: Calcd for $C_{23}H_{24}ClNO_5PS_2$ $[M+H]^+$ 524.0522; Found 524.0527

#### Compound (11, MB-2A).

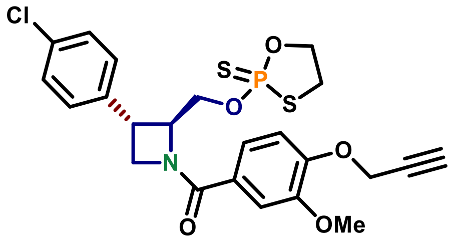

**((2*S*,3*S*)-3-(4-chlorophenyl)-2-(((2-sulfido-1,3,2-oxathiaphospholan-2-yl)oxy)methyl)azetidin-1-yl)(3-methoxy-4-(prop-2-yn-1-yloxy)phenyl)methanone (11, MB-2A)**

Utilizing procedure reported for **9**, compound **11** was prepared from **S7B** (20 mg, 0.052 mmol, 1 equiv) to afford 10 mg of the titled product (**Yield** = 50 %, **d.r.** ~1:1, 99% e.e.).

*Note: the titled product was isolated by PTLC with 30% Et2O in hexanes.*

**Physical state:** colorless film

**$R_t$:** 1.77 min

**$R_f$:** 0.37 (60% EtOAc in hexanes), UV active

**Optical rotation:** $[\alpha]_D^{25}$ –68.4° (*c* 0.58, $CHCl_3$)

**$^1$H NMR (600 MHz, $CDCl_3$) δ** 7.37 – 7.29 (m, 3H), 7.28 – 7.20 (m, 3H), 7.03 (d, *J* = 8.3 Hz, 1H), 4.81 (d, *J* = 2.4 Hz, 2H), 4.78 – 4.59 (m, 3H), 4.55 – 4.34 (m, 3H), 4.15 – 4.09 (m, 1H), 3.96 – 3.80 (m, 4H), 3.59 – 3.39 (m, 2H), 2.53 (t, *J* = 2.4 Hz, 1H).

**$^{13}$C NMR (151 MHz, CDCl$_3$) δ** 170.9, 170.8, 149.7, 149.6, 139.1, 133.39, 133.36, 129.20, 129.18, 128.44, 128.41, 126.5, 121.0, 113.0, 111.9, 78.0, 76.5, 70.9, 68.0, 67.9, 67.65, 67.60, 58.6, 56.8, 56.23, 56.22, 37.1, 36.97, 36.86, 36.8, 36.6.

**$^{31}$P NMR (162 MHz, CDCl$_3$) δ** 105.2, 104.9.

**HRMS (ESI-TOF) m/z**: Calcd for $C_{23}H_{24}ClNO_5PS_2$ [M+H]$^+$ 524.0522; Found 524.0529

#### Compound (10, MB-1B).

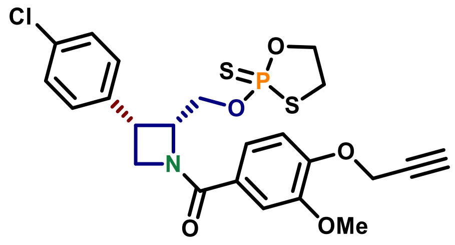

**((2*R*,3*S*)-3-(4-chlorophenyl)-2-(((2-sulfido-1,3,2-oxathiaphospholan-2-yl)oxy)methyl)azetidin-1-yl)(3-methoxy-4-(prop-2-yn-1-yloxy)phenyl)methanone (10, MB-1B)**

Utilizing procedure reported for **9,** compound **3C** was prepared from **S7C (**10 mg, 0.026 mmol, 1 equiv) to afford 3.5 mg of the titled product (**Yield** = 26%, **d.r.** ~1:1, 96% e.e.).

**Optical rotation:** $[\alpha]_D^{25}$ –31.4° (*c* 0.035, CHCl$_3$)

*Note: Other characterizations were identical to compound **9.***

#### Compound (12, MB-2B).

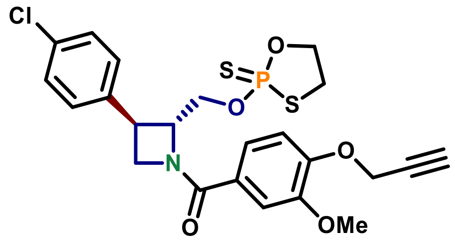

**((2*R*,3*R*)-3-(4-chlorophenyl)-2-(((2-sulfido-1,3,2-oxathiaphospholan-2-yl)oxy)methyl)azetidin-1-yl)(3-methoxy-4-(prop-2-yn-1-yloxy)phenyl)methanone (12, MB-2B)**

Utilizing procedure reported for **9,** compound **12** was prepared from **S7D** (57 mg, 0.052 mmol, 1 equiv) to afford 46 mg of the titled product (**Yield** = 59%, **d.r.** ~1:1, 96% e.e.).

**Optical rotation:** $[\alpha]_D^{25}$ +69.5° (*c* 1.0, CHCl$_3$)

*Note: Other characterizations were identical to compound **11.***

#### Compound (13, MB-3A).

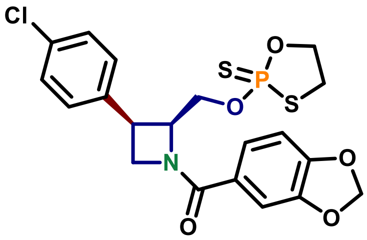

**Benzo[*d*][1,3]dioxol-5-yl((2*S*,3*R*)-2-(*tert*-butoxymethyl)-3-(4-chlorophenyl)azetidin-1-yl)methanone (13, MB-3A)**

Utilizing a modified (see *notes*) procedure reported for **9**, compound **13** was prepared from **S8A** (20 mg, 0.052 mmol, 1 equiv) to afford 16 mg of the titled product (**Yield** = 63%, **d.r.** ~1:1, 93% e.e.).

*Note #1: 2-chloroacetamide (15 mg, 0.16 mmol, 3 equiv) was added before the addition of DBU to improve the yield by scavenging the formed perfluorobenzenethiol.*

*Note #2: The amount of the **Ψ** reagent and **DBU** was doubled: (52 mg, 0.16 mmol, 3 equiv) and (24 mg, 23 μL, 0156 mmol, 3 equiv), respectively.*

**Physical state:** colorless film

**$R_f$:** 0.32 (60% EtOAc in hexanes), UV active

**Optical rotation:** $[\alpha]_D^{25}$ +27.9° (*c* 0.43, $CHCl_3$)

**$^1$H NMR (600 MHz, $CDCl_3$) δ** 7.40 – 7.34 (m, 2H), 7.29 – 7.22 (m, 3H), 7.19 – 7.16 (m, 1H), 6.85 (d, *J* = 8.1 Hz, 1H), 6.03 (d, *J* = 1.4 Hz, 1H), 6.02 (d, *J* = 1.4 Hz, 1H), 5.07 – 4.96 (m, 1H), 4.65 – 4.57 (m, 1H), 4.38 (t, *J* = 9.0 Hz, 1H), 4.32 – 3.99 (m, 5H), 3.42 – 3.31 (m, 1H), 3.31 – 3.21 (m, 1H).

**$^{13}$C NMR (151 MHz, $CDCl_3$) δ** 170.8, 170.6, 150.4, 148.0, 134.8, 133.5, 129.8, 129.6, 129.98, 128.95, 127.1, 123.1, 108.52, 108.51, 108.30, 108.29, 101.8, 70.54, 70.53, 70.52, 65.4, 65.3, 65.11, 65.07, 36.9, 36.7, 36.58, 36.57, 36.5.

**$^{31}$P NMR (162 MHz, $CDCl_3$) δ** 103.45, 103.00.

**HRMS (ESI-TOF) m/z**: Calcd for $C_{20}H_{20}ClNO_5PS_2$ $[M+H]^+$ 484.0209; Found 484.0211

#### Compound (15, MB-4A).

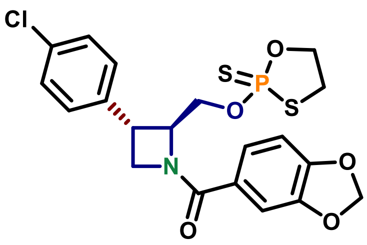

**Benzo[*d*][1,3]dioxol-5-yl((2*S*,3*S*)-2-(*tert*-butoxymethyl)-3-(4-chlorophenyl)azetidin-1-yl)methanone (15, MB-4A)**

Utilizing a modified (see *notes*) procedure reported for **9**, compound **15** was prepared from **S8B** (20 mg, 0.052 mmol, 1 equiv) to afford 14 mg of the titled product (**Yield** = 33%, **d.r.** ~1:1, 95% e.e.).

*Note #1: 2-chloroacetamide (12 mg, 0.13 mmol, 1.5 equiv) was added before the addition of DBU to improve the yield by scavenging the formed perfluorobenzenethiol.*

*Note #2: Crude product was purified by PTLC with 50% EtOAc in hexanes.*

**Physical state:** colorless film

**$R_f$:** 0.22 (50% EtOAc in hexanes), UV active

**Optical rotation:** $[\alpha]_D^{25}$ –79.7° (*c*, 0.33, $CHCl_3$)

**$^1$H NMR (600 MHz, $CDCl_3$) δ** 7.32 (d, *J* = 6.5 Hz, 2H), 7.25 – 7.22 (m, 3H)(overlapped with $CHCl_3$ peak), 7.18 (s, 1H), 6.84 (d, *J* = 8.1 Hz, 1H), 6.04 – 6.01 (m, 2H), 4.67 (br s, 3H), 4.55 – 4.31 (m, 3H), 4.10 – 4.02 (m, 1H), 3.85 (br app d, *J* = 39.1 Hz, 1H), 3.60 – 3.42 (m, 2H).

**$^{13}$C NMR (151 MHz, $CDCl_3$) δ** 170.74, 170.63, 150.46, 148.00, 147.99, 139.09, 133.39, 133.36, 129.21, 129.18, 128.44, 128.42, 126.87, 123.14, 108.53, 108.30, 101.82, 70.92, 70.76, 67.82, 67.51, 58.52, 36.97, 36.88, 36.87, 36.73, 36.49.

**$^{31}$P NMR (162 MHz, $CDCl_3$) δ** 105.3, 104.9.

**HRMS (ESI-TOF) m/z**: Calcd for $C_{20}H_{20}ClNO_5PS_2$ $[M+H]^+$ 484.0209; Found 484.0212

#### Compound (14, MB-3B).

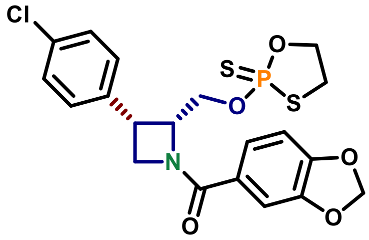

**Benzo[d][1,3]dioxol-5-yl((2R,3S)-3-(4-chlorophenyl)-2-(((2-sulfido-1,3,2-oxathiaphospholan-2-yl)oxy)methyl)azetidin-1-yl)methanone (14, MB-3B)**

Utilizing the procedure reported for **9**, compound **14** was prepared from **S8C** (20 mg, 0.052 mmol, 1 equiv) to afford 17 mg of the titled product (**Yield** = 63%, **d.r.** ~1:1, 97% e.e.).

**Optical rotation:** $[\alpha]_D^{25}$ –28.3° (*c* 0.66, $CHCl_3$)

*Note: Other characterizations were identical to compound **13**.*

#### Compound (16, MB-4B).

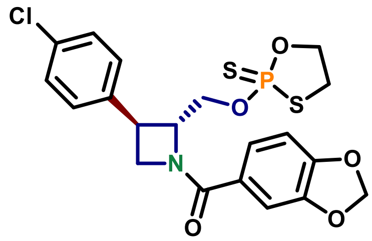

**Benzo[d][1,3]dioxol-5-yl((2R,3R)-2-(tert-butoxymethyl)-3-(4-chlorophenyl)azetidin-1-yl)methanone (16. MB-4B)**

Utilizing procedure reported for **9**, compound **16** was prepared from **S8D** (33 mg, 0.095 mmol, 1 equiv) to afford 20 mg of the titled product (**Yield** = 43%, **d.r.** ~1:1, 95% e.e.).

**Optical rotation:** $[\alpha]_D^{25}$ +73.6° ($c$ 0.66, $CHCl_3$)

*Note: Other characterizations were identical to compound **15**.*

#### 2.3 General Scheme for synthesis of alternative azetidine alkyne probes and competitors.

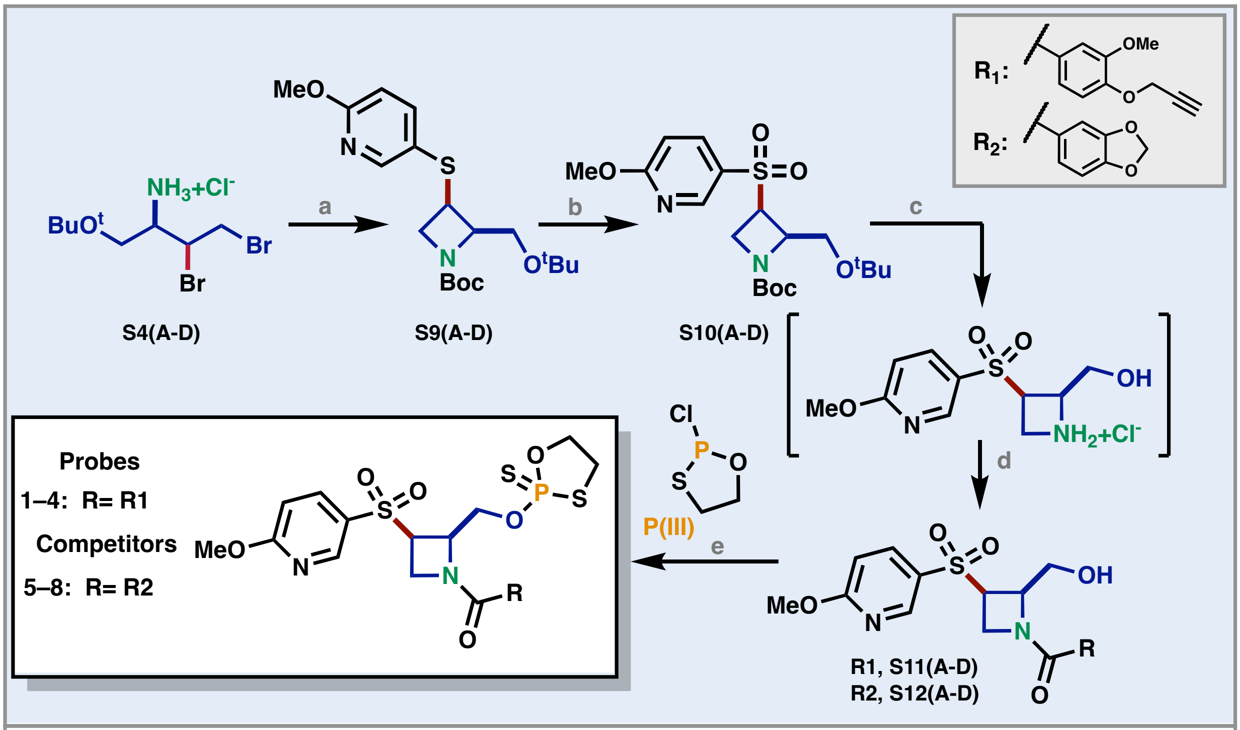

a. i. nBuLi, PhMe, -78 °C, 3 h; iii. ArSH, -78 °C —> rt, 20 min; iii. $Boc_2O$, 1 h; b. TFA, mCPBA, DCM, 0 °C 45 min h; c. 85% $H_3PO_4$, DCM, rt, 2 h; d. RCOOH, HATU, DIPEA, DCM, rt; e. P(III), $S_8$, pyridine, 1 h

#### Compound (S9A).

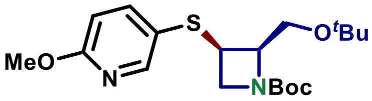

***Tert*-butyl (2*R*,3*R*)-2-(*tert*-butoxymethyl)-3-((6-methoxypyridin-3-yl)thio)azetidine-1-carboxylate (S9A)**

A flamed dried 10 mL glass culture tube equipped with a stir bar was charged with **S4A** (300 mg, 0.885 mmol, 1 equiv) and dry PhMe (8.8 mL). The tube was evacuated and back filled with argon three times and chilled –78°C in a solid $CO_2$ bath. 2.5 M ***n*BuLi** solution in hexanes (1.1 mL, 2.8 mmol, 3.2 equiv) was added dropwise and the mixture was stirred for 3 h at –78°C. **6-methoxypyridine-3-thiol** (150 µL, 1.06 mmol, 1.2 equiv)(see *note*) was added and the reaction mixture was removed from the cooling bath and left stirring at room temperature for 30 min. $Boc_2O$ (306 µL, 1.3 mmol, 1.5 equiv) was added and stirred for 1h. The solvent was removed *in vacuo* and the product was isolated by silica chromotagraphy (5% EtOAc in hexanes, then 15% EtOAc in hexanes once product started to elute) to give 170 mg of the titled product (**Yield** = 50 %, d.r. >20:1).

*Note: Prepared according to previously reported procedure in Bielecki et. al.*

**Physical state:** colorless film

**$R_t$:** 2.14 min

**$^1$H NMR (600 MHz, $CDCl_3$) δ** 8.16 (d, $J$ = 2.3 Hz, 1H), 7.63 (dd, $J$ = 8.6, 2.5 Hz, 1H, 6.66 (d, $J$ = 8.6 Hz, 1H), 4.49 – 4.41 (m, 1H), 4.17 (t, $J$ = 8.8 Hz, 1H), 4.01 – 3.91 (m, 2H), 3.91 (s, 3H),3.75 (dd, $J$ = 8.8, 6.2 Hz, 1H), 3.69 (dd, $J$ = 9.2, 2.9 Hz, 1H), 1.43 (s, 9H), 1.20 (d, $J$ = 0.7 Hz, 9H).

**$^{13}$C NMR (151 MHz, $CDCl_3$) δ** 163.6, 155.9, 149.4, 142.3, 124.5, 111.4, 79.9, 73.6, 65.5, 59.5, 55.7, 53.7, 41.8, 28.5, 27.6.

**LRMS (ESI-TOF) m/z**: Calcd for $C_{15}H_{23}N_2O_4S$ $[M{-}C_4H_8{+}H]^+$ 327.14; Found 327.11

**Compound (S9B).**

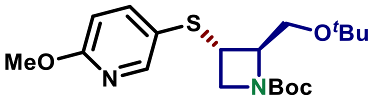

***Tert*-butyl (2*R*,3*S*)-2-(*tert*-butoxymethyl)-3-((6-methoxypyridin-3-yl)thio)azetidine-1-carboxylate (S9B)**

Utilizing procedure reported for **S9A**, compound **S9B** was prepared from **S4B** (300 mg, 0.885 mmol, 1 equiv) to afford 125 mg on the titled compound (**Yield** = 37%, **d.r.** > 20:1).

**Physical state:** colorless film

**$R_t$:** 2.11 min

**$^1$H NMR (600 MHz, $CDCl_3$) δ** 8.23 (dd, $J$ = 2.6, 0.6 Hz, 1H), 7.66 (dd, $J$ = 8.6, 2.5 Hz, 1H), 6.70 (dd, $J$ = 8.6, 0.7 Hz, 1H), 4.15 – 4.07 (m, 2H), 3.92 (s, 3H), 3.71 (dt, $J$ = 7.6, 4.9 Hz, 1H), 3.67 (dd, $J$ = 8.7, 5.3 Hz, 1H), 3.61 (br s, 2H), 3.43 (dd, $J$ = 10.2, 2.8 Hz, 1H), 1.41 (s, 9H), 1.12 (s, 9H).

**$^{13}$C NMR (151 MHz, $CDCl_3$) δ** 164.2, 155.9, 151.2, 143.8, 121. 9, 111.6, 79.8, 73.2, 69.5, 61.2, 53.8, 39.0, 28.5, 27.6.

**LRMS (ESI-TOF) m/z**: Calcd for $C_{19}H_{31}N_2O_4S$ $[M{+}H]^+$ 383.20; Found 383.26

#### Compound (S9C).

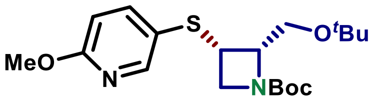

***Tert*-butyl (2*S*,3*S*)-2-(*tert*-butoxymethyl)-3-((6-methoxypyridin-3-yl)thio)azetidine-1-carboxylate (S9C)**

Utilizing procedure reported for **S9A**, compound **S9C** was prepared from **S4C** (300 mg, 0.885 mmol, 1 equiv) to afford 162 mg on the titled compound (**Yield** = 47%, **d.r.** > 20:1).

**$^1$H NMR (600 MHz, CDCl$_3$) δ** 8.16 (s, 1H), 7.63 (dd, $J$ = 8.5, 2.2 Hz, 1H), 6.67 (d, $J$ = 8.7 Hz, 1H), 4.47 – 4.42 (m, 1H), 4.17 (t, $J$ = 8.8 Hz, 1H), 3.99 – 3.92 (m, 2H), 3.91 (s, 3H), 3.75 (dd, $J$ = 8.8, 6.3 Hz, 1H), 3.69 (dd, $J$ = 9.3, 2.4 Hz, 1H), 1.43 (s, 9H), 1.20 (s, 9H).

**$^{13}$C NMR (151 MHz, CDCl$_3$) δ** 163.6, 155.9, 149.4, 142.3, 124.5, 111.4, 79.9, 73.6, 65.4, 59.5, 55.5, 53.7, 41.8, 28.5, 27.6.

**LRMS (ESI-TOF) m/z**: Calcd for $C_{15}H_{23}N_2O_4S$ [M−$C_4H_8$ +H]$^+$ 327.14; Found 327.10

#### Compound (S9D).

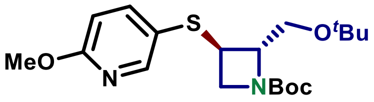

***Tert*-butyl (2*S*,3*R*)-2-(*tert*-butoxymethyl)-3-((6-methoxypyridin-3-yl)thio)azetidine-1-carboxylate (S9D)**

Utilizing procedure reported for **S9A**, compound **S9D** was prepared from **S4D** (300 mg, 0.885 mmol, 1 equiv) to afford 166 mg on the titled compound (**Yield** = 49%, **d.r.** > 20:1).

**$^1$H NMR (600 MHz, CDCl$_3$) δ** 8.23 (dd, $J$ = 2.6, 0.6 Hz, 1H), 7.66 (dd, $J$ = 8.6, 2.5 Hz, 1H), 6.70 (dd, $J$ = 8.6, 0.7 Hz, 1H), 4.15 – 4.07 (m, 2H), 3.92 (s, 3H), 3.71 (dt, $J$ = 7.6, 4.9 Hz, 1H), 3.67 (dd, $J$ = 8.7, 5.3 Hz, 1H), 3.61 (br s, 2H), 3.43 (dd, $J$ = 10.2, 2.8 Hz, 1H), 1.41 (s, 9H), 1.12 (s, 9H).

**$^{13}$C NMR (151 MHz, CDCl$_3$) δ** 164.2, 155.9, 151.2, 143.8, 121.9, 111.6, 79.8, 73.2, 69.5, 61.2, 54.15, 53.8, 39.0, 28.5, 27.6.

**LRMS (ESI-TOF) m/z**: Calcd for $C_{19}H_{31}N_2O_4S$ [M +H]$^+$ 383.20; Found 383.17

#### Compound (S10A).

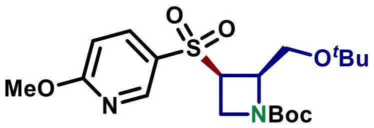

***T*ert-butyl (2*R*,3*R*)-2-(*tert*-butoxymethyl)-3-((6-methoxypyridin-3-yl)sulfonyl)azetidine-1-carboxylate (S10A)**

In a 4 dram glass vial equipped with a stir, **S9A** (161 mg, 0.421 mmol, 1 equiv) was dissolved in DCM (2.5 mL) and chilled to 0 °C. TFA (32 µLm 0.421 mmol, 1 equiv) followed by 51% mCPBA (313 mg, 0.926 mmol, 2.2 equiv) and stirred for 45 min at 0 °C. The reaction mixture was diluted with $CHCl_3$ and the organic layer was washed with aq. $Na_2SO_3$ (×3), aq. $NaHCO_3$ (×2), dried with $MgSO_4$ and the solvent was removed *in vacuo* to obtain 173 mg of the titled compound (**Yield** = 99%)

**Physical state:** white solid

**$R_f$**: 0.48 (40% EtOAc in hexanes), UV active

**$R_t$**: 1.80 min

**Optical rotation:** $[\alpha]_D^{25}$ –33.8° (*c* 2.6, $CHCl_3$)

**$^1$H NMR (600 MHz, $CDCl_3$) δ** 8.71 (dd, *J* = 2.5, 0.7 Hz, 1H), 8.01 (dd, *J* = 8.8, 2.6 Hz, 1H), 6.83 (dd, *J* = 8.8, 0.7 Hz, 1H), 4.49 (td, *J* = 8.6, 4.0 Hz, 1H), 4.27 – 4.24 (m, 1H), 4.22 (dd, *J* = 9.4, 5.4 Hz, 1H), 4.11 – 4.04 (m, 1H), 4.03 – 3.99 (m, 4H)(*$OCH_3$ overlapped with a CH*), 3.79 (dd, *J* = 9.8, 4.1 Hz, 1H), 1.44 (s, 9H), 1.17 (s, 9H).

**$^{13}$C NMR (151 MHz, $CDCl_3$) δ** 67.4, 156.0, 149.3, 138.3, 128.4, 111.6, 80.8, 73.8, 64.7, 58.9, 54.8, 54.6, 48.5, 28.5, 27.5.

**HRMS (ESI-TOF) m/z**: Calcd for $C_{19}H_{31}N_2O_6S$ $[M+H]^+$ 415.1903; Found 415.1913

#### Compound (S10B).

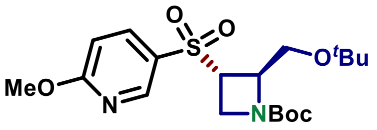

***T*ert-butyl (2*R*,3*S*)-2-(*tert*-butoxymethyl)-3-((6-methoxypyridin-3-yl)sulfonyl)azetidine-1-carboxylate (S10B)**

Utilizing procedure reported for **S10A**, compound **S10B** was prepared from **S9B** (168 mg, 0.440 mmol, 1 equiv) to afford 180 mg on the titled compound (**Yield** = 99%).

**Physical state:** white solid

**$R_f$:** 0.62 (40% EtOAc in hexanes), UV active

**Optical rotation:** $[\alpha]_D^{25}$ –13.1° (*c* 1.9, $CHCl_3$)

**$^1$H NMR (600 MHz, $CDCl_3$) δ** 8.69 (dd, *J* = 2.5, 0.7 Hz, 1H), 7.99 (dd, *J* = 8.8, 2.5 Hz, 1H), 6.86 (dd, *J* = 8.7, 0.7 Hz, 1H), 4.53 (s, 1H), 4.15 (s, 1H), 4.02 (s, 3H), 4.00 – 3.95 (m, 1H), 3.94 – 3.91 (m, 1H), 3.64 (dd, *J* = 10.4, 4.5 Hz, 1H), 3.34 (dd, *J* = 10.2, 2.8 Hz, 1H), 1.43 (s, 9H), 1.05 (s, 9H).

**$^{13}$C NMR (151 MHz, $CDCl_3$) δ** 167.6, 155.6, 149.4, 138.5, 127.0, 111.9, 80.6, 73.4, 63.0 (broad), 60.7 (broad), 54.8 (broad), 54.6, 47.8 (v broad), 28.5, 27.5.

**HRMS (ESI-TOF) m/z**: Calcd for $C_{19}H_{30}N_2NaO_6S$ $[M+Na]^+$ 437.1722; Found 437.1729

#### Compound (S10C).

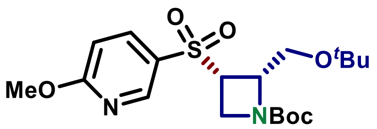

***Tert*-butyl (2*S*,3*S*)-2-(*tert*-butoxymethyl)-3-((6-methoxypyridin-3-yl)sulfonyl)azetidine-1-carboxylate (S10C)**

Utilizing procedure reported for **S10A**, compound **S10C** was prepared from **S4C** (154 mg, 0.403 mmol, 1 equiv) to afford 158 mg on the titled compound (**Yield** = 95%).

**Optical rotation:** $[\alpha]_D^{25}$ +34.5° (*c* 3.0, $CHCl_3$)

*Note: Other characterizations were identical to compound **S10A.***

#### Compound (S10D).

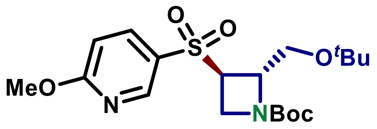

***Tert*-butyl (2*S*,3*R*)-2-(*tert*-butoxymethyl)-3-((6-methoxypyridin-3-yl)sulfonyl)azetidine-1-carboxylate (S10D)**

Utilizing procedure reported for **S10A**, compound **S10D** was prepared from **S4D** (150 mg, 0.403 mmol, 1 equiv) to afford 161 mg on the titled compound (**Yield** = 99%).

**Optical rotation:** $[\alpha]_D^{25}$ +16.2° (*c* 1.9, $CHCl_3$)

*Note: Other characterizations were identical to compound **S10B.***

#### Compound (S11A).

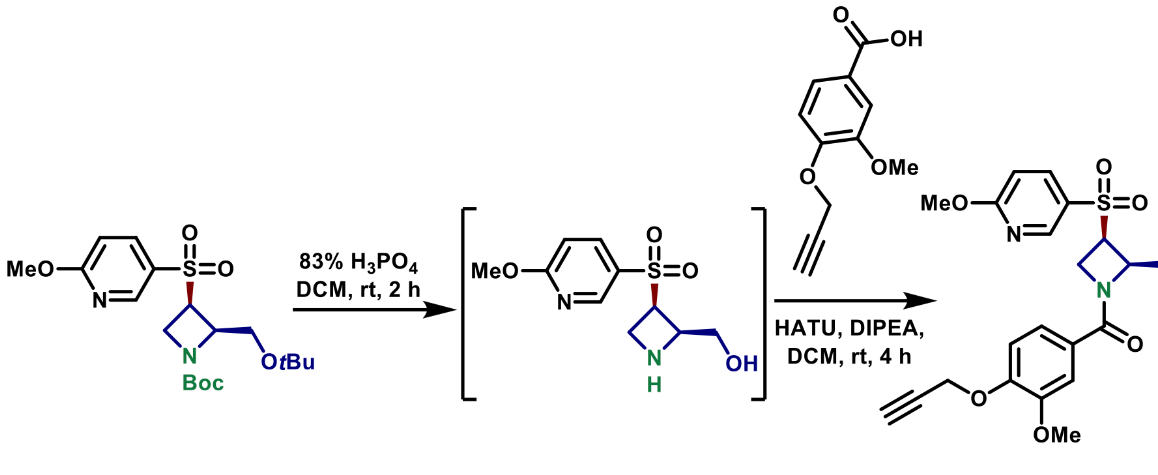

S10A

S11A

**((2*R*,3*R*)-2-(hydroxymethyl)-3-((6-methoxypyridin-3-yl)sulfonyl)azetidin-1-yl)(3-methoxy-4-(prop-2-yn-1-yloxy)phenyl)methanone (S11A)**

A 4 dram glass vial equipped with a stir bar was charged with **S10A** (93 mg, 0.22 mmol, 1 equiv), DCM (1 mL) and **85% $H_3PO_4$** (0.40 mL). The reaction mixture was vigorously stirred for 2 h. The reaction mixture was chilled to 0 °C and quenched with conc. $K_2CO_3$ and extracted with $CHCl_3$ (×4). The extracts were combined, dried with $Na_2SO_4$ and the solvent was removed *in vacuo* to obtain a white solid which was used directly in the next step. A portion of the deprotected crude product (16 mg, 0.062 mmol, 1 equiv), **3-methoxy-4-(prop-2-yn-1-yloxy)benzoic acid** (14 mg, 0.066 mmol, 1.1 equiv) and **HATU** (25 mg, 0.066 mmol, 1.1 equiv) were combined in DCM (0.80 mL) in a 1 dram glass vial equipped with a stir bar. **DIPEA** (27 µL, 0.158 mmol, 2.5 equiv) was added and the mixture was stirred for 4 h at room temperature. The reaction mixture was diluted with EtOAc and washed with aq. $NaH_2PO_4$ (×2), $NaHCO_3$ (×2), brine (×1), dried with NaSO4 and the solvent was removed on rotary evaporated. The product was isolated by PTLC with 20% acetone in DCM to give 13 mg of the titled compound (**Yield** = 47%).

**Physical state**: white solid

**$^1$H NMR (600 MHz, $CD_3CN$) δ** 8.68 (d, $J$ = 2.6 Hz, 1H), 8.05 (dd, $J$ = 8.8, 2.6 Hz, 1H), 7.27 – 7.20 (m, 2H), 7.02 (d, $J$ = 8.1 Hz, 1H), 6.93 (d, $J$ = 8.8 Hz, 1H), 5.00 – 4.93 (m, 1H), 4.80 (d, $J$ = 2.4 Hz, 2H), 4.58 – 4.45 (m, 2H), 4.40 – 4.30 (m, 2H), 4.24 (br s, 1H), 4.07 (br s, 1H), 3.98 (s, 3H), 3.84 (s, 3H), 2.84 (t, $J$ = 2.4 Hz, 1H).

**$^{13}$C NMR (151 MHz, $CD_3CN$) δ** 171.6, 168.6, 150.6, 150.4, 150.0, 139.5, 129.0, 126.7, 121.9, 113.8, 112.7, 112.5, 79.2, 77.3, 67.9, 62.0, 57.2, 56.6, 55.8, 55.2, 53.7.

**LRMS (ESI-TOF) m/z**: Calcd for $C_{21}H_{23}N_2O_7S$ $[M+H]^+$ 447.12; Found 447.20

#### Compound (S11B).

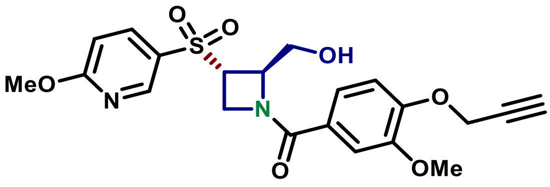

**((2*R*,3*S*)-2-(hydroxymethyl)-3-((6-methoxypyridin-3-yl)sulfonyl)azetidin-1-yl)(3-methoxy-4-(prop-2-yn-1-yloxy)phenyl)methanone (S11B)**

Utilizing procedure reported for **S11A**, compound **S11B** was prepared from corresponding crude deprotection product (21 mg, 0.081 mmol) derived from S**10B** to afford 10 mg of the titled product (**Yield** = 28%).

**Physical state:** white solid

**$^1$H NMR (600 MHz, $CD_3CN$) δ** 8.69 – 8.66 (m, 1H), 8.06 (d, $J$ = 8.1 Hz, 1H), 7.19 (br s, 2H), 7.03 (d, $J$ = 8.1 Hz, 1H), 6.93 (d, $J$ = 8.8 Hz, 1H), 4.84 – 4.74 (m, 3H), 4.51 (br s, 1H), 4.27 (br s, 1H), 4.17 – 4.13 (m, 1H), 3.99 (s, 3H), 3.85 (s, 3H), 3.78 (s, 1H), 3.54 (s, 2H), 2.84 (t, $J$ = 2.4 Hz, 1H).

**$^{13}$C NMR (151 MHz, $CD_3CN$) δ** 171.0, 168.6, 150.5, 150.4, 150.2, 139.8, 127.6, 126.7(broad), 121.9(broad), 113.9, 112.7, 112.4, 79.2, 77.3, 65.2(broad), 62.6(broad), 57.2, 56.5, 55.28, 55.19, 52.5(broad).

**LRMS (ESI-TOF) m/z**: Calcd for $C_{21}H_{23}N_2O_7S$ $[M+H]^+$ 447.12; Found 447.20

#### Compound (S11C).

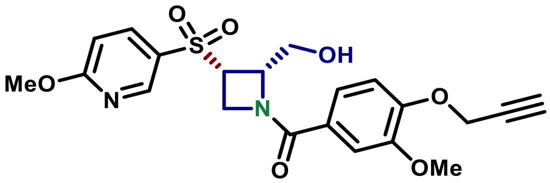

**((2*S*,3*S*)-2-(hydroxymethyl)-3-((6-methoxypyridin-3-yl)sulfonyl)azetidin-1-yl)(3-methoxy-4-(prop-2-yn-1-yloxy)phenyl)methanone (S11C)**

Utilizing procedure reported for **S11A**, compound **S11C** was prepared from the corresponding crude deprotection product (20 mg, 0.077 mmol) derived from S**10C** to afford 16 mg of the titled product (**Yield** = 47%).

**Physical state**: white solid

**$^1$H NMR (600 MHz, $CD_3CN$ ) δ** 8.68 (d, *J* = 2.5 Hz, 1H), 8.05 (dd, *J* = 8.8, 2.6 Hz, 1H), 7.25 – 7.20 (m, 2H), 7.02 (d, *J* = 8.2 Hz, 1H), 6.93 (d, *J* = 8.9 Hz, 1H), 4.98 – 4.94 (m, 1H), 4.80 (d, *J* = 2.4 Hz, 2H), 4.58 – 4.42 (m, 2H), 4.40 – 4.29 (br m, 2H), 4.24 (br s, 1H), 4.10 – 4.03 (m, 1H), 3.98 (s, 3H), 3.84 (s, 3H), 2.84 (t, *J* = 2.4 Hz, 1H).

**$^{13}$C NMR (151 MHz, $CD_3CN$) δ** 171.6, 168.6, 150.6, 150.4, 150.0, 139.5, 129.0, 126.7, 121.9, 113.8, 112.7, 112.5, 79.2, 77.3, 67.9, 62.0, 57.2, 56.6, 55.8, 55.2, 53.7.

**LRMS (ESI-TOF) m/z**: Calcd for $C_{21}H_{23}N_2O_7S$ $[M+H]^+$ 447.12; Found 447.20

#### Compound (S11D).

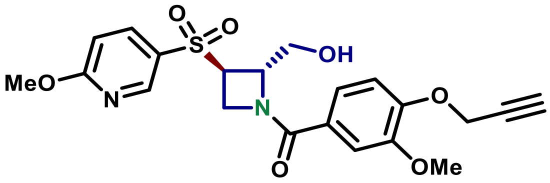

**((2*S*,3*R*)-2-(hydroxymethyl)-3-((6-methoxypyridin-3-yl)sulfonyl)azetidin-1-yl)(3-methoxy-4-(prop-2-yn-1-yloxy)phenyl)methanone (S11D)**

Utilizing procedure reported for **S11A**, compound **S11D** was prepared from corresponding crude deprotection product (19 mg, 0.074 mmol) derived from S**10D** to afford 19 mg of the titled product (**Yield** = 57%).

**Physical state**: white solid

**$^1$H NMR (600 MHz, $CD_3CN$ ) δ** 8.69 – 8.65 (m, 1H), 8.06 (d, *J* = 8.7 Hz, 1H), 7.19 (br s, 2H), 7.03 (d, *J* = 8.2 Hz, 1H), 6.93 (d, *J* = 8.8 Hz, 1H), 4.80 – 4.76 (m, 3H), 4.51 (br s, 1H), 4.27 (br s, 1H), 4.18 – 4.12 (m, 1H), 3.99 (s, 3H), 3.85 (s, 3H), 3.79 (br s, 1H), 3.54 (br s, 2H), 2.84 (t, *J* = 2.4 Hz, 1H).

**$^{13}$C NMR (151 MHz, CD$_3$CN) δ** 171.0, 168.6, 150.5, 150.4, 150.2, 139.8, 127.6, 126.7(broad), 121.9(broad), 113.9, 112.7, 112.4, 79.2, 77.3, 65.2(broad), 62.6(broad), 57.2, 56.5, 55.2, 55.2, 52.4(broad).

**LRMS (ESI-TOF) m/z**: Calcd for $C_{21}H_{23}N_2O_7S$ $[M+H]^+$ 447.12; Found 447.19

#### Compound (S12A).

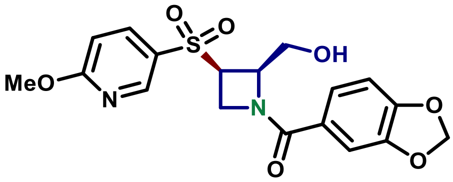

**Benzo[*d*][1,3]dioxol-5-yl((2*R*,3*R*)-2-(hydroxymethyl)-3-((6-methoxypyridin-3-yl)sulfonyl)azetidin-1-yl)methanone (S12A)**

Utilizing procedure reported for **S11A**, compound **S12A** was prepared from corresponding crude deprotection product (23 mg, 0.089 mmol) (derived from S**10A)** and **piperonylic acid** to afford 13 mg of the titled product (**Yield** = 36%).

**Physical state:** white solid

**$^1$H NMR (600 MHz, CD$_3$CN) δ** 8.68 (d, $J$ = 2.6 Hz, 1H), 8.05 (dd, $J$ = 8.8, 2.6 Hz, 1H), 7.20 (dd, $J$ = 8.1, 1.8 Hz, 1H), 7.13 (d, $J$ = 1.8 Hz, 1H), 6.93 (dd, $J$ = 8.9, 0.7 Hz, 1H), 6.88 (d, $J$ = 8.1 Hz, 1H), 6.03 (d, $J$ = 1.0 Hz, 1H), 6.03 (d, $J$ = 1.0 Hz, 1H), 4.94 (td, $J$ = 8.2, 3.7 Hz, 1H), 4.51 (dd, $J$ = 9.3, 5.5 Hz, 1H), 4.46 – 4.30 (m, 2H), 4.30 – 4.16 (m, 2H), 4.10 – 4.02 (m, 1H), 3.98 (s, 3H).

**$^{13}$C NMR (151 MHz, CD$_3$CN) δ** 171.5, 168.6, 151.5, 150.0, 148.9, 139.6, 128.9, 127.1, 124.1, 112.7, 108.92, 108.88, 103.2, 67.8(broad), 62.0(broad), 55.7, 55.2, 53.6(broad).

**LRMS (ESI-TOF) m/z**: Calcd for $C_{18}H_{19}N_2O_7S$ $[M+H]^+$ 407.09; Found 407.18

#### Compound (S12B).

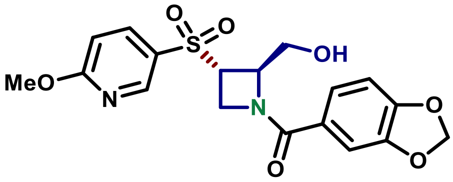

**Benzo[*d*][1,3]dioxol-5-yl((2*R*,3*S*)-2-(hydroxymethyl)-3-((6-methoxypyridin-3-yl)sulfonyl)azetidin-1-yl)methanone (S12B)**

Utilizing procedure reported for **S11A**, compound **S12B** was prepared from the corresponding crude deprotection product (25 mg, 0.097 mmol) (derived from S**10B)** and **piperonylic acid** to afford 9.3 mg of the titled product (**Yield** = 24%).

**Physical state:** white solid

**$^1$H NMR (600 MHz, CDCl$_3$) δ** 8.67 (s, 1H), 8.06 (d, $J$ = 9.1 Hz, 1H), 7.15 (br s, 1H), 7.08 (br s, 1H), 6.93 (d, $J$ = 8.9 Hz, 1H), 6.88 (d, $J$ = 8.1 Hz, 1H), 6.03 (d, $J$ = 1.1 Hz, 1H), 6.03 (d, $J$ = 1.1 Hz, 1H), 4.77 (d, $J$ = 4.2

Hz, 1H), 4.46 (br s, 1H), 4.24 (br s, 1H), 4.18 – 4.11 (m, 1H), 3.99 (s, 3H), 3.80 – 3.74 (br m, 1H), 3.52 (br s, 2H).

**$^{13}$C NMR (151 MHz, $CDCl_3$) δ** 170.8, 168.6, 151.4, 150.2, 148.9, 139.8, 127.6, 124.0 (broad), 112.7, 108.9, 108.8, 103.1, 65.2 (broad), 62.5 (broad), 55.22 (broad, overlapped), 55.19, 52.4 (broad).

**HRMS (ESI-TOF) m/z**: Calcd for $C_{18}H_{19}N_2O_7S$ $[M+H]^+$ 407.09; Found 407.17

#### Compound (S12C).

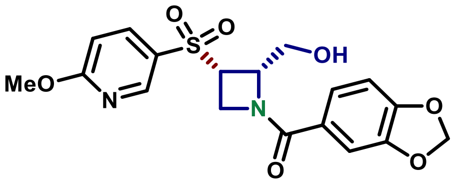

**Benzo[*d*][1,3]dioxol-5-yl((2*S*,3*S*)-2-(hydroxymethyl)-3-((6-methoxypyridin-3-yl)sulfonyl)azetidin-1-yl)methanone (S12C)**

Utilizing procedure reported for **S11A**, compound **S12C** was prepared from the corresponding crude deprotection product (34 mg, 0.13 mmol) (derived from S**10C)** and **piperonylic acid** to afford 26 mg of the titled product (**Yield** = 49%).

**Physical state**: white solid

**$R_t$**: 1.13 min

**$^1$H NMR (600 MHz, $CD_3CN$) δ** 8.68 (d, *J* = 2.6 Hz, 1H), 8.05 (dd, *J* = 8.8, 2.6 Hz, 1H), 7.21 (dd, *J* = 8.1, 1.7 Hz, 1H), 7.14 (d, *J* = 1.7 Hz, 1H), 6.94 (d, *J* = 8.8 Hz, 1H), 6.89 (d, *J* = 8.1 Hz, 1H), 6.03 (d, *J* = 0.9 Hz, 1H), 6.03 (d, *J* = 1.0 Hz, 1H), 4.94 (td, *J* = 8.3, 3.8 Hz, 1H), 4.51 (dd, *J* = 9.5, 5.4 Hz, 1H), 4.46 – 4.30 (m, 2H), 4.30 – 4.17 (m, 2H), 4.10 – 4.03 (m, 1H), 3.99 (s, 3H).

**$^{13}$C NMR (151 MHz, $CD_3CN$) δ** 171.5, 168.6, 151.5, 150.0, 148.9, 139.6, 128.9, 124.1, 112.7, 108.93, 108.88, 103.2, 67.8, 62.0, 55.7, 55.2, 53.6, 38.9.

**LRMS (ESI-TOF) m/z**: Calcd for $C_{18}H_{19}N_2O_7S$ $[M+H]^+$ 407.09; Found 407.17

#### Compound (S12D).

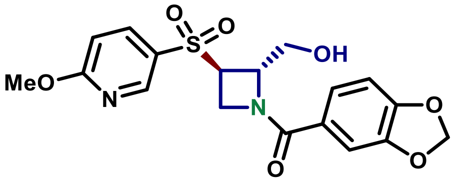

**Benzo[*d*][1,3]dioxol-5-yl((2*S*,3*R*)-2-(hydroxymethyl)-3-((6-methoxypyridin-3-yl)sulfonyl)azetidin-1-yl)methanone (S12D)**

Utilizing procedure reported for **S11A**, compound **S12D** was prepared from the corresponding crude deprotection product (24 mg, 0.093 mmol) (derived from S**10D)** and **piperonylic acid** to afford 14 mg of the titled product (**Yield** = 37%).

**Physical state**: white solid

**$R_t$**: 1.06 min

**$^1$H NMR (600 MHz, $CD_3CN$) δ** 8.67 (s, 1H), 8.05 (dd, *J* = 8.9, 2.5 Hz, 1H), 7.15 (d, *J* = 7.9 Hz, 1H), 7.08 (s, 1H), 6.93 (d, *J* = 8.8 Hz, 1H), 6.88 (d, *J* = 8.0 Hz, 1H), 6.03 (d, *J* = 1.1 Hz, 1H), 6.03 (d, *J* = 1.1 Hz, 1H), 4.82 – 4.73 (m, 1H), 4.45 (br s, 1H), 4.24 (br s, 1H), 4.14 (dt, *J* = 8.8, 5.0 Hz, 1H), 3.99 (s, 3H), 3.76 (br s, 1H), 3.52 (br s, 2H).

*Characterization otherwise matches S12B*

#### Compound (1, MAH-1A).

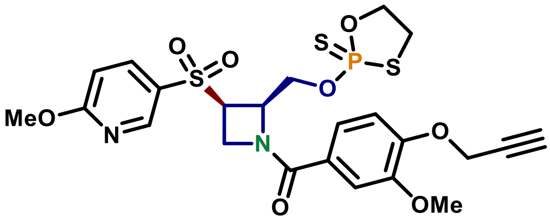

**(3-methoxy-4-(prop-2-yn-1-yloxy)phenyl)((2*R*,3*R*)-3-((6-methoxypyridin-3-yl)sulfonyl)-2-(((2-sulfido-1,3,2-oxathiaphospholan-2-yl)oxy)methyl)azetidin-1-yl)methanone (1, MAH-1A)**

A 4 dram glass vial equipped with a stir bar was charged with **S11A** (13 mg, 0.029 mmol, 1 equiv), **2-chloro-1,3,2-oxathiaphospholane** (4.4 µL, 0.044 mmol, 1.5 equiv) and **sulfur** (4.7 mg, 0.146 mmol, 5 equiv), powdered *3 Å molecular sieves* and dry pyridine (300 µL). The mixture was stirred for 1 h at room temperature. The reaction mixture was diluted with dry PhMe and filtered through celite. The solvent was removed *in vacuo* redissolved in PhMe and rotary evaporated. The crude product was purified by PTLC with 10% acetone in DCM to give 11 mg of the titled product (**Yield** = 67%, **d.r.** ~ 1:1, 96% e.e.)

**Physical state:** colorless solid

**$R_f$**: 0.27 (70% EtOAc in hexanes), UV active

**Optical rotation:** $[\alpha]_D^{25}$ –86.1° (*c* 0.46, $CHCl_3$)

**$^1$H NMR (600 MHz, $CDCl_3$) δ** 8.72 – 8.69 (m, 1H), 8.05 – 7.96 (m, 1H), 7.24 (dd, *J* = 5.0, 2.0 Hz, 1H), 7.20 – 7.16 (m, 1H), 7.04 – 7.00 (m, 1H), 6.88 (d, *J* = 8.8 Hz, 1H), 5.13 (td, *J* = 8.0, 5.3 Hz, 0.5H), 5.07 (td, *J* = 7.6, 4.9 Hz, 0.5H), 4.93 – 4.85 (m, 1H), 4.80 (d, *J* = 2.0 Hz, 2H), 4.76 – 4.65 (m, 2H), 4.65 – 4.49 (m, 1H), 4.43 – 4.33 (m, 1H), 4.30 – 4.22 (m, 1H), 4.19 (q, *J* = 8.9 Hz, 1H), 4.02 (s, 3H), 3.90 (d, *J* = 1.2 Hz, 3H), 3.73 – 3.66 (m, 0.5H), 3.64 – 3.56 (m, 0.5H), 3.48 – 3.36 (m, 1H), 2.56 – 2.51 (m, 1H).

**$^{13}$C NMR (151 MHz, $CDCl_3$) δ** 171.18, 171.04, 167.73, 167.71, 150.00, 149.83, 149.82, 149.22, 138.14, 138.09, 127.78, 127.76, 125.57, 121.11, 121.08, 112.99, 112.22, 111.76, 111.72, 77.89, 76.60, 70.89, 70.87, 64.28, 64.23, 64.08, 64.03, 62.63, 56.76, 56.26, 54.84, 54.77, 52.34, 36.67, 36.60.

**$^{31}$P NMR (162 MHz, $CDCl_3$) δ** 103.95, 103.72.

**HRMS (ESI-TOF) m/z**: Calcd for $C_{23}H_{26}N_2O_8PS_3$ $[M+H]^+$ 585.0589; Found 585.0599

#### Compound (3, MAH-2A).

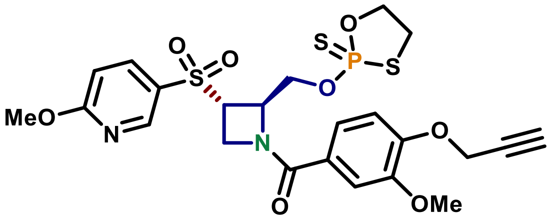

**(3-methoxy-4-(prop-2-yn-1-yloxy)phenyl)((2*R*,3*S*)-3-((6-methoxypyridin-3-yl)sulfonyl)-2-(((2-sulfido-1,3,2-oxathiaphospholan-2-yl)oxy)methyl)azetidin-1-yl)methanone (3, MAH-2A)**

Utilizing procedure reported for **1**, compound **3** was prepared from **S11B** (9.3 mg, 0.021mmol) to afford 7.4 mg of the titled product (**Yield** = 61%, **d.r. ~ 1:1, 97% e.e.**).

**Physical state:** colorless solid

**$R_f$:** 0.32 (70% EtOAc in hexanes), UV active

**Optical rotation:** $[\alpha]_D^{25}$ –60.3° (*c*. 0.13, $CHCl_3$)

**$^1$H NMR (600 MHz, $CDCl_3$) δ** 8.69 (s, 1H), 7.99 (dd, *J* = 8.8, 2.6 Hz, 1H), 7.27 (br s, 1H)(*overlapped with $CHCl_3$ peak*), 7.15 (d, *J* = 8.4 Hz, 1H), 7.03 (dd, *J* = 8.3, 0.9 Hz, 1H), 6.89 (d, *J* = 8.8 Hz, 1H), 5.01 – 4.94 (m, 1H), 4.82 (d, *J* = 2.4 Hz, 2H), 4.64 – 4.34 (m, 5H), 4.26 – 4.07 (m, 2H), 4.03 (s, 3H), 3.92 (s, 3H), 3.58 – 3.38 (m, 2H), 2.55 (t, *J* = 2.4 Hz, 1H).

**$^{13}$C NMR (151 MHz, $CDCl_3$) δ** 170.87, 167.88, 167.86, 149.91, 149.88, 149.87, 149.52, 149.48, 138.42, 138.37, 126.18, 126.11, 125.55, 120.82, 112.99, 112.30, 112.27, 111.83, 77.93, 76.59, 70.98, 70.80, 66.64, 66.60, 61.58, 56.78, 56.27, 56.25, 54.89, 54.79, 54.77, 54.47, 36.84, 36.75.

**$^{31}$P NMR (162 MHz, $CDCl_3$) δ** 105.42, 105.38.

**HRMS (ESI-TOF) m/z**: Calcd for $C_{23}H_{26}N_2O_8PS_3$ $[M+H]^+$ 585.0589; Found 585.0602

#### Compound (2, MAH-1B).

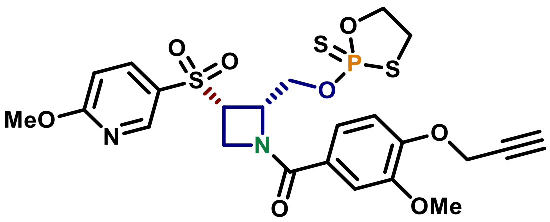

**(3-methoxy-4-(prop-2-yn-1-yloxy)phenyl)((2*S*,3*S*)-3-((6-methoxypyridin-3-yl)sulfonyl)-2-(((2-sulfido-1,3,2-oxathiaphospholan-2-yl)oxy)methyl)azetidin-1-yl)methanone (2, MAH-1B)**

Utilizing procedure reported for **1**, compound **2** was prepared from **S11C** (17.4 mg, 0.039 mmol) to afford 15.3 mg of the titled product (**Yield** = 67%, **d.r. ~ 1:1, 96% e.e.**).

**Optical rotation:** $[\alpha]_D^{25}$ +80.6° (*c*. 0.88, $CHCl_3$)

*Note: Other characterizations were identical to compound **5A.***

#### Compound (4, MAH-2B).

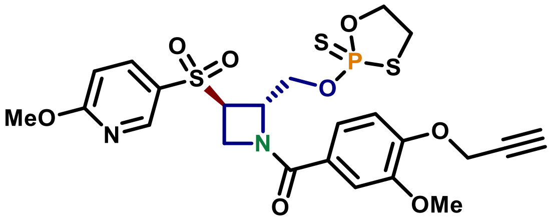

**(3-methoxy-4-(prop-2-yn-1-yloxy)phenyl)((2*S*,3*R*)-3-((6-methoxypyridin-3-yl)sulfonyl)-2-(((2-sulfido-1,3,2-oxathiaphospholan-2-yl)oxy)methyl)azetidin-1-yl)methanone (4, MAH-2B)**

Utilizing procedure reported for **1**, compound **4** was prepared from **S11D** (19.2 mg, 0.039 mmol) to afford 16.0 mg of the titled product (**Yield** = 64%, **d.r. ~ 1:1, 91% e.e.**).

**Optical rotation:** $[\alpha]_D^{25}$ +60.9° (*c* 0.87, $CHCl_3$)

*Note: Other characterizations were identical to compound **5B.***

#### Compound (5, MAH-3A).

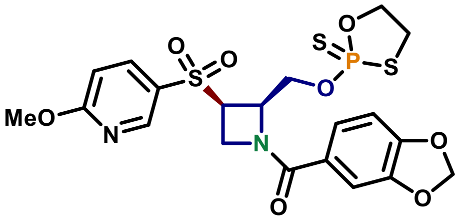

**Benzo[*d*][1,3]dioxol-5-yl((2*R*,3*R*)-3-((6-methoxypyridin-3-yl)sulfonyl)-2-(((2-sulfido-1,3,2-oxathiaphospholan-2-yl)oxy)methyl)azetidin-1-yl)methanone (5, MAH-3A)**

Utilizing procedure reported for **1**, compound **5** was prepared from **S12A** (12.2 mg, 0.030mmol) to afford 11.1 mg of the titled product (**Yield** = 68%, **d.r. ~ 1:1, 97% e.e.**).

**Physical state:** colorless solid

**$R_f$:** 0.36 (70% EtOAc in hexanes), UV active

**Optical rotation:** $[\alpha]_D^{25}$ – 89.8° (*c* 0.44, $CHCl_3$)

**$^1$H NMR (600 MHz, $CDCl_3$) δ** 8.70 (t, *J* = 2.2 Hz, 1H), 8.03 – 7.98 (m, 1H), 7.18 (dd, *J* = 8.1, 1.7 Hz, 1H), 7.10 (d, *J* = 1.7 Hz, 1H), 6.87 (d, *J* = 8.8 Hz, 1H), 6.83 (dd, *J* = 8.1, 1.7 Hz, 1H), 6.04 – 6.01 (m, 1H), 6.02 – 6.02 (m, 1H), 5.11 (td, *J* = 8.0, 5.2 Hz, 0.5H), 5.08 – 5.03 (m, 0.5H), 4.92 – 4.83 (m, 1H), 4.73 – 4.64 (m, 2H), 4.64 – 4.51 (m, 1H), 4.42 – 4.34 (m, 1H), 4.28 – 4.20 (m, 1H), 4.15 (q, *J* = 9.4 Hz, 1H), 4.02 (s, 3H), 3.76 – 3.66 (m, 0.5H), 3.65 – 3.57 (m, 0.5H), 3.48 – 3.38 (m, 1H).

**$^{13}$C NMR (151 MHz, CDCl$_3$) δ** 171.01, 170.87, 167.72, 167.70, 150.92, 149.22, 148.13, 138.14, 138.10, 127.79, 127.77, 125.86, 123.36, 112.21, 108.45, 108.44, 108.42, 101.93, 70.91, 70.90, 70.87, 70.86, 64.30, 64.25, 64.11, 64.07, 62.52, 54.76, 54.71, 52.34, 36.66, 36.58.

**$^{31}$P NMR (162 MHz, CDCl$_3$) δ** 103.97, 103.71.

**HRMS (ESI-TOF) m/z**: Calcd for $C_{20}H_{22}N_2O_8PS_3$ $[M+H]^+$ 545.0276; Found 545.0284

#### Compound (7, MAH-4A).

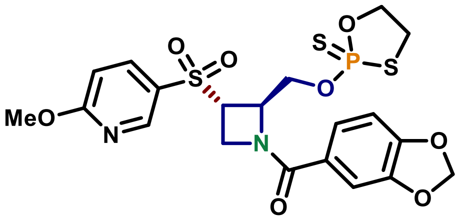

**Benzo[*d*][1,3]dioxol-5-yl((2*R*,3*S*)-3-((6-methoxypyridin-3-yl)sulfonyl)-2-(((2-sulfido-1,3,2-oxathiaphospholan-2-yl)oxy)methyl)azetidin-1-yl)methanone (7, MAH-4A)**

Utilizing procedure reported for **1**, compound **7** was prepared from **S12B** (15.5 mg, 0.038 mmol) to afford 12.3 mg of the titled product (**Yield** = 59%, **d.r. ~ 1:1, 97% e.e.**).

**Physical state:** colorless film

**R$_f$:** 0.41 (70% EtOAc in hexanes), UV active

**Optical rotation:** $[\alpha]_D^{25}$ – 64.1° (*c* 0.71, CHCl$_3$)

**$^1$H NMR (600 MHz, CDCl$_3$) δ** 8.68 (d, *J* = 2.5 Hz, 1H), 7.98 (dd, *J* = 8.7, 2.5 Hz, 1H), 7.20 – 7.12 (m, 1H), 7.12 – 7.04 (m, 1H), 6.89 (d, *J* = 8.7 Hz, 1H), 6.83 (d, *J* = 8.1 Hz, 1H), 6.08 – 5.98 (m, 2H), 5.00 – 4.90 (m, 1H), 4.63 – 4.32 (m, 5H), 4.24 – 4.06 (m, 2H), 4.03 (s, 3H), 3.56 – 3.41 (m, 2H).

**$^{13}$C NMR (151 MHz, CDCl$_3$) δ** 170.67, 167.87, 167.85, 150.82, 150.80, 149.51, 149.47, 148.11, 138.42, 138.36, 126.14, 126.06, 123.16, 112.28, 112.26, 108.38, 101.92, 101.91, 71.00, 70.81, 66.56, 66.51, 61.48, 54.78, 54.76, 54.40, 51.26, 36.83, 36.72.

**$^{31}$P NMR (162 MHz, CDCl$_3$) δ** 105.49, 105.44.

**HRMS (ESI-TOF) m/z**: Calcd for $C_{20}H_{22}N_2O_8PS_3$ $[M+H]^+$ 545.0276; Found 545.0283

#### Compound (6, MAH-3B).

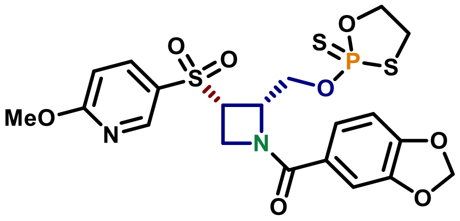

**Benzo[*d*][1,3]dioxol-5-yl((2*S*,3*S*)-3-((6-methoxypyridin-3-yl)sulfonyl)-2-(((2-sulfido-1,3,2-oxathiaphospholan-2-yl)oxy)methyl)azetidin-1-yl)methanone (6, MAH-3B)**

Utilizing procedure reported for **1**, compound **6** was prepared from **S12C** (26.3 mg, 0.064 mmol) to afford 24.4 mg of the titled product (**Yield** = 70%, **d.r. ~ 1:1, 96% e.e.**).

**Optical rotation:** $[\alpha]_D^{25}$ +86.6° (*c* 1.6, $CHCl_3$)

*Note: Other characterizations were identical to compound* ***5.***

#### Compound (8, MAH-4B).

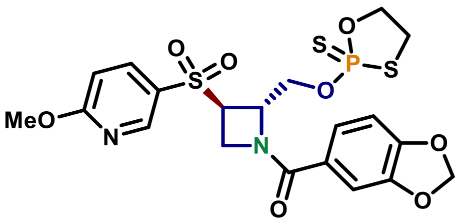

**Benzo[*d*][1,3]dioxol-5-yl((2*S*,3*R*)-3-((6-methoxypyridin-3-yl)sulfonyl)-2-(((2-sulfido-1,3,2-oxathiaphospholan-2-yl)oxy)methyl)azetidin-1-yl)methanone (8, MAH-4B)**

Utilizing procedure reported for **1**, compound **8** was prepared from **S12D** (12.5 mg, 0.031 mmol) to afford 13.0 mg of the titled product (**Yield** = 78%, **d.r. ~ 1:1, 91% e.e.**).

**Optical rotation:** $[\alpha]_D^{25}$ +69.8° (*c* 0.57, $CHCl_3$)

*Note: Other characterizations were identical to compound 7.*

#### 3. NMR Spectra

**[1]H NMR of compound (P(V)-Reagent) (600 MHz, $CDCl_3$)**

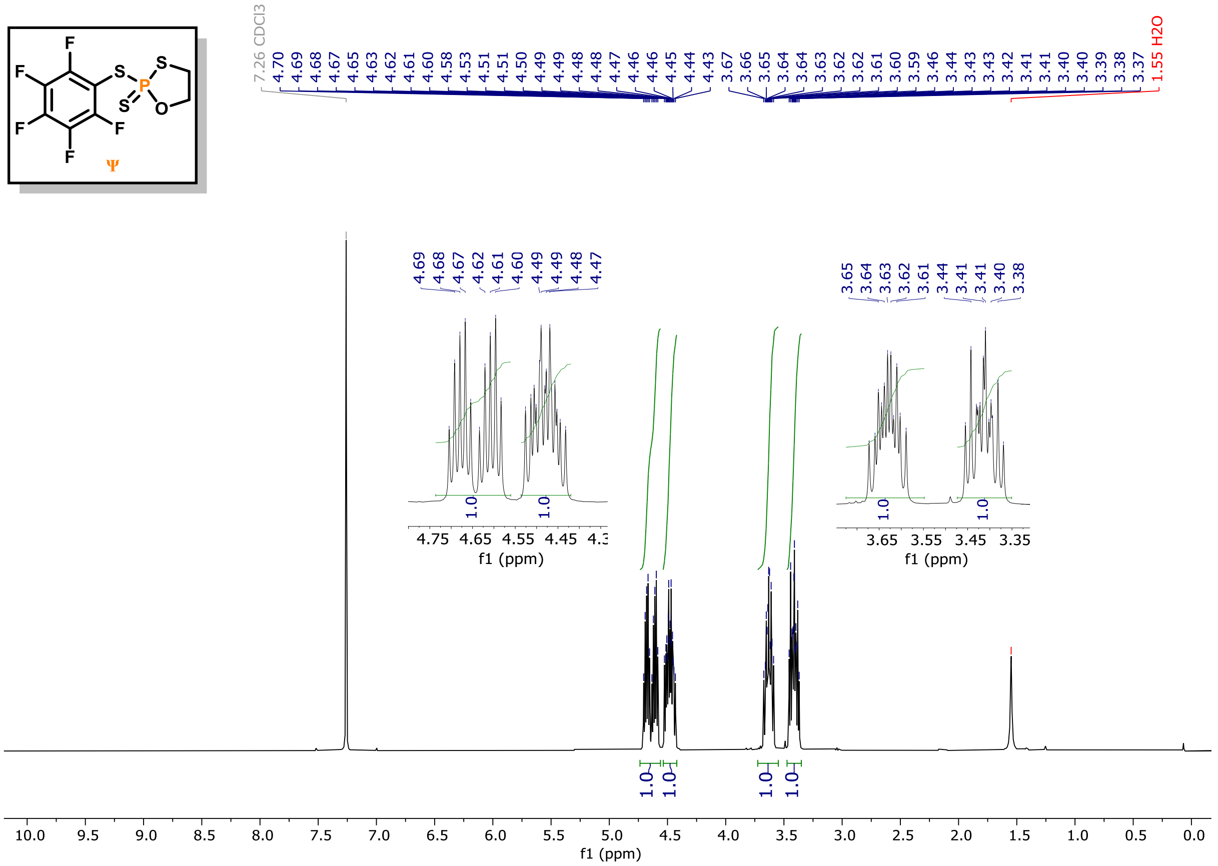

**[13]C NMR of compound (P(V)-Reagent) (150 MHz, $CDCl_3$)**

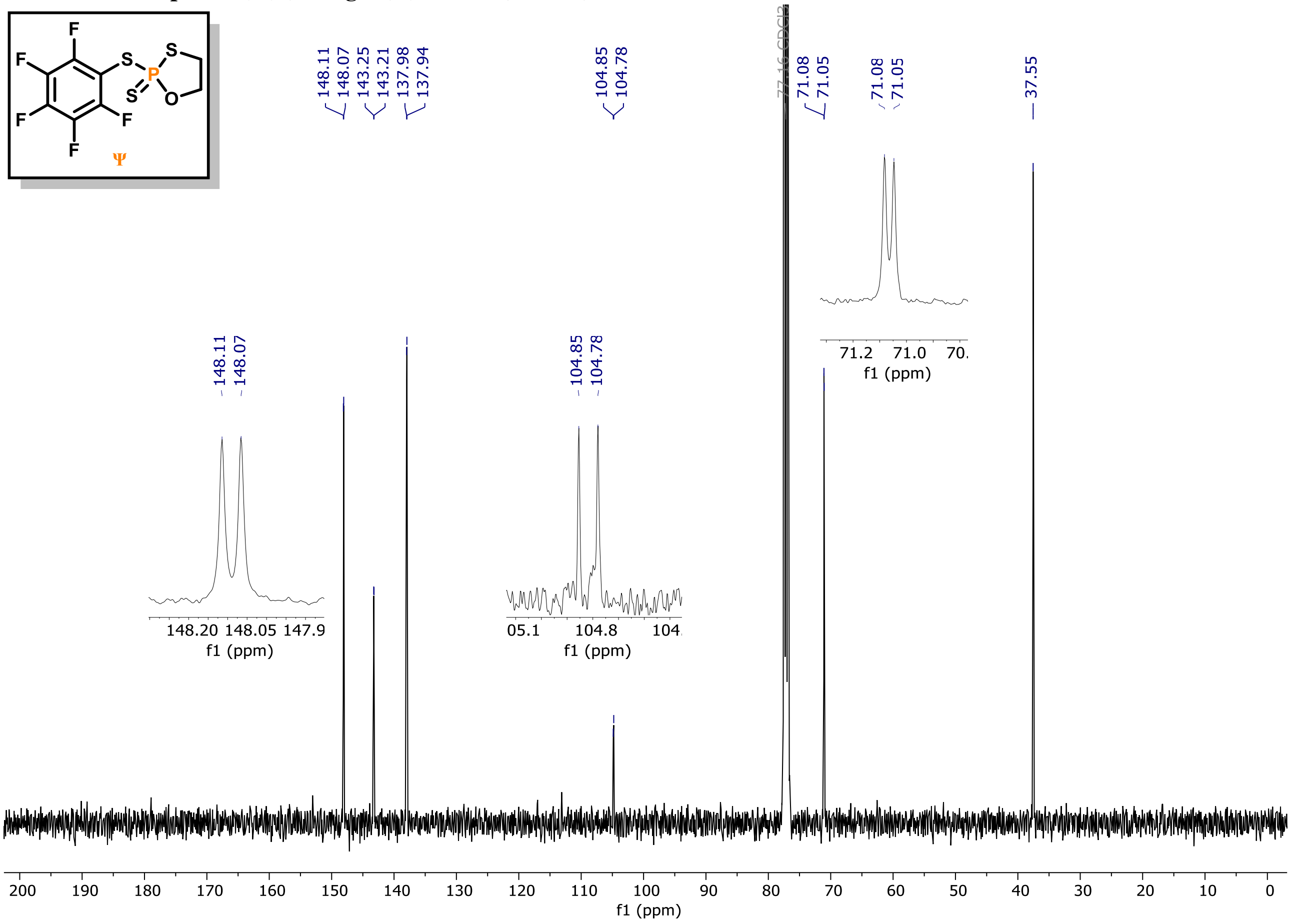

**[31]P NMR of compound (P(V)-Reagent) (162 MHz, CDCl$_3$)**

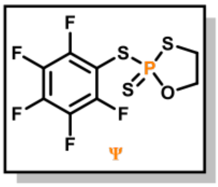

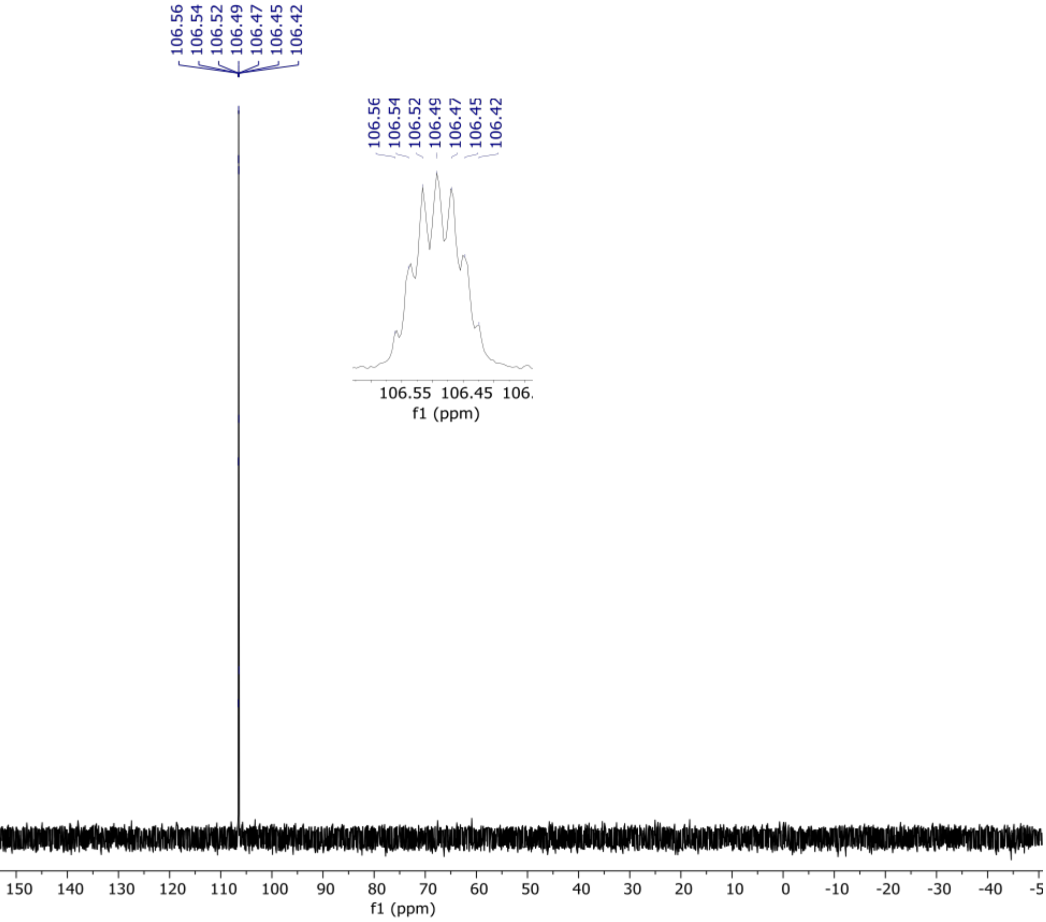

$^1$H NMR of compound (S1A) (600 MHz, $CDCl_3$)

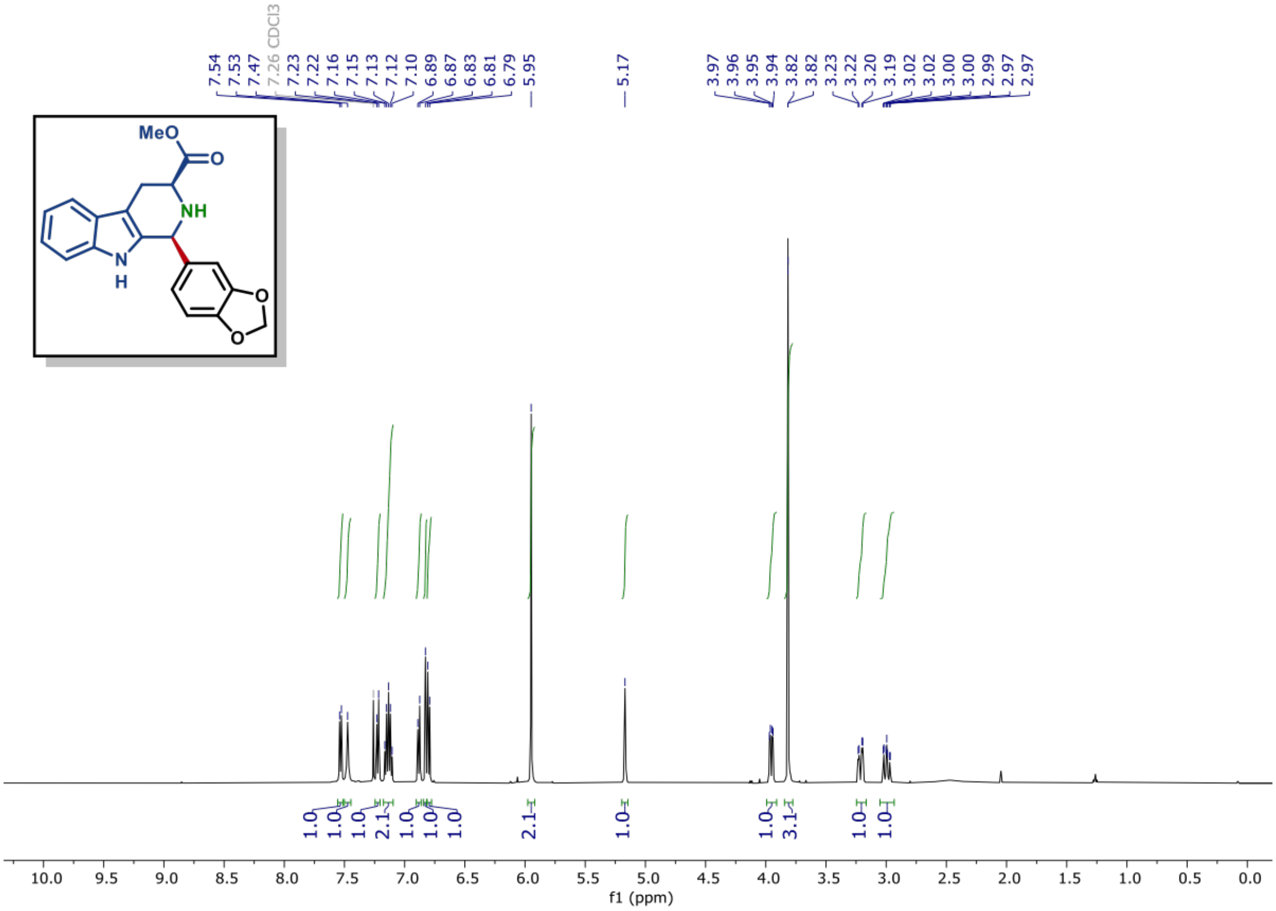

$^{13}$C NMR of compound (S1A) (150 MHz, $CDCl_3$)

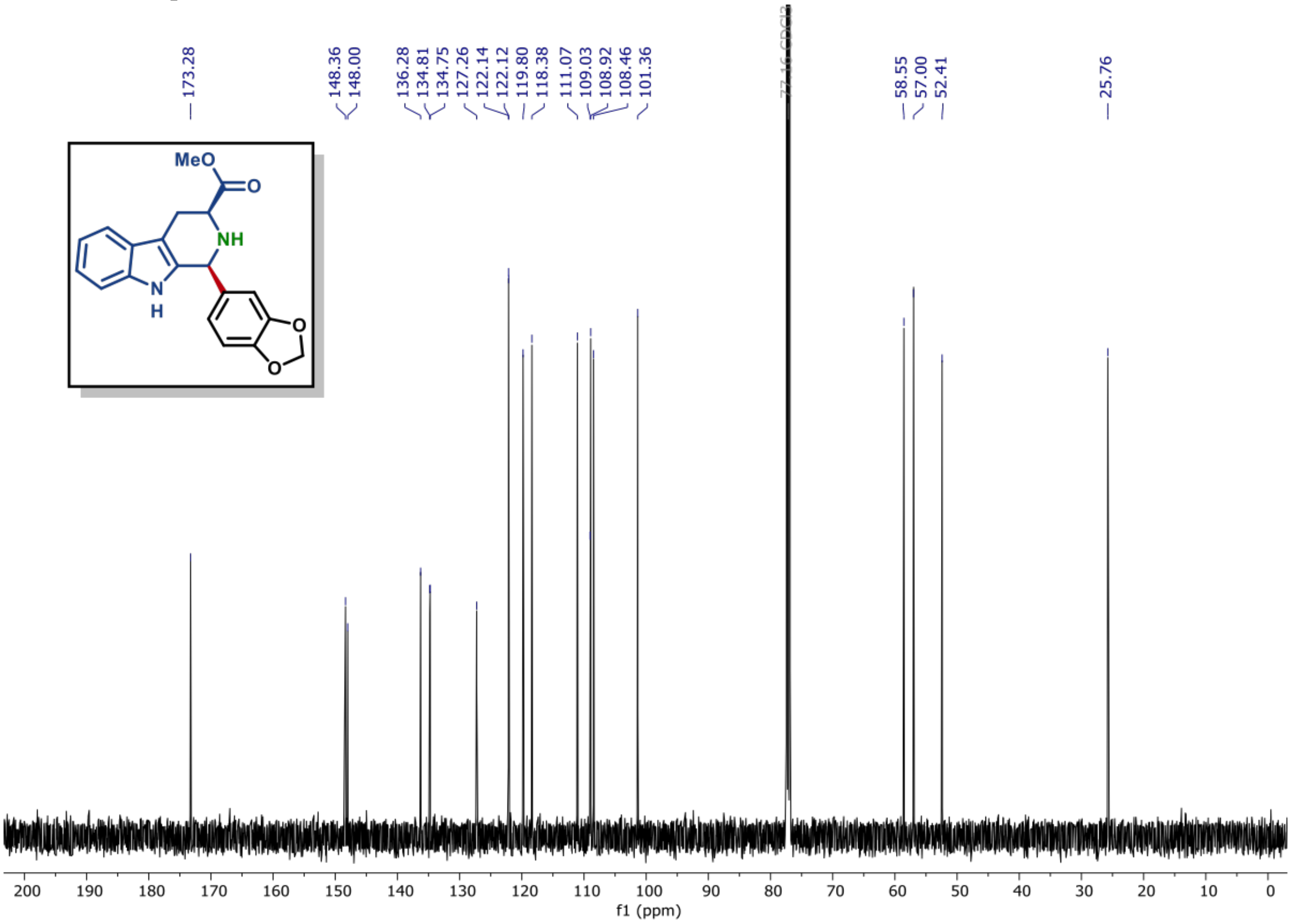

[1]H NMR of compound (S1B) (600 MHz, $CDCl_3$)

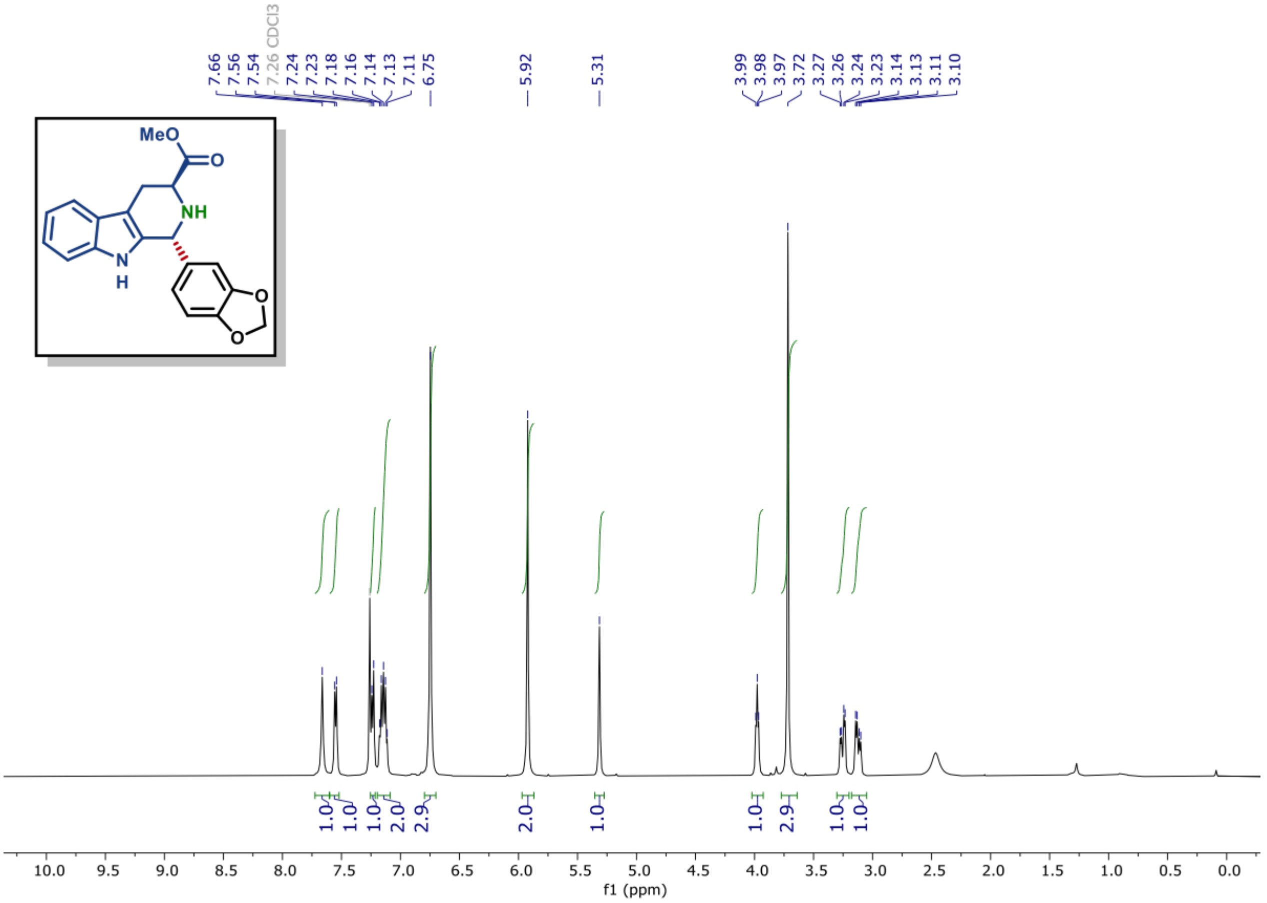

[13]C NMR of compound (S1B) (150 MHz, $CDCl_3$)

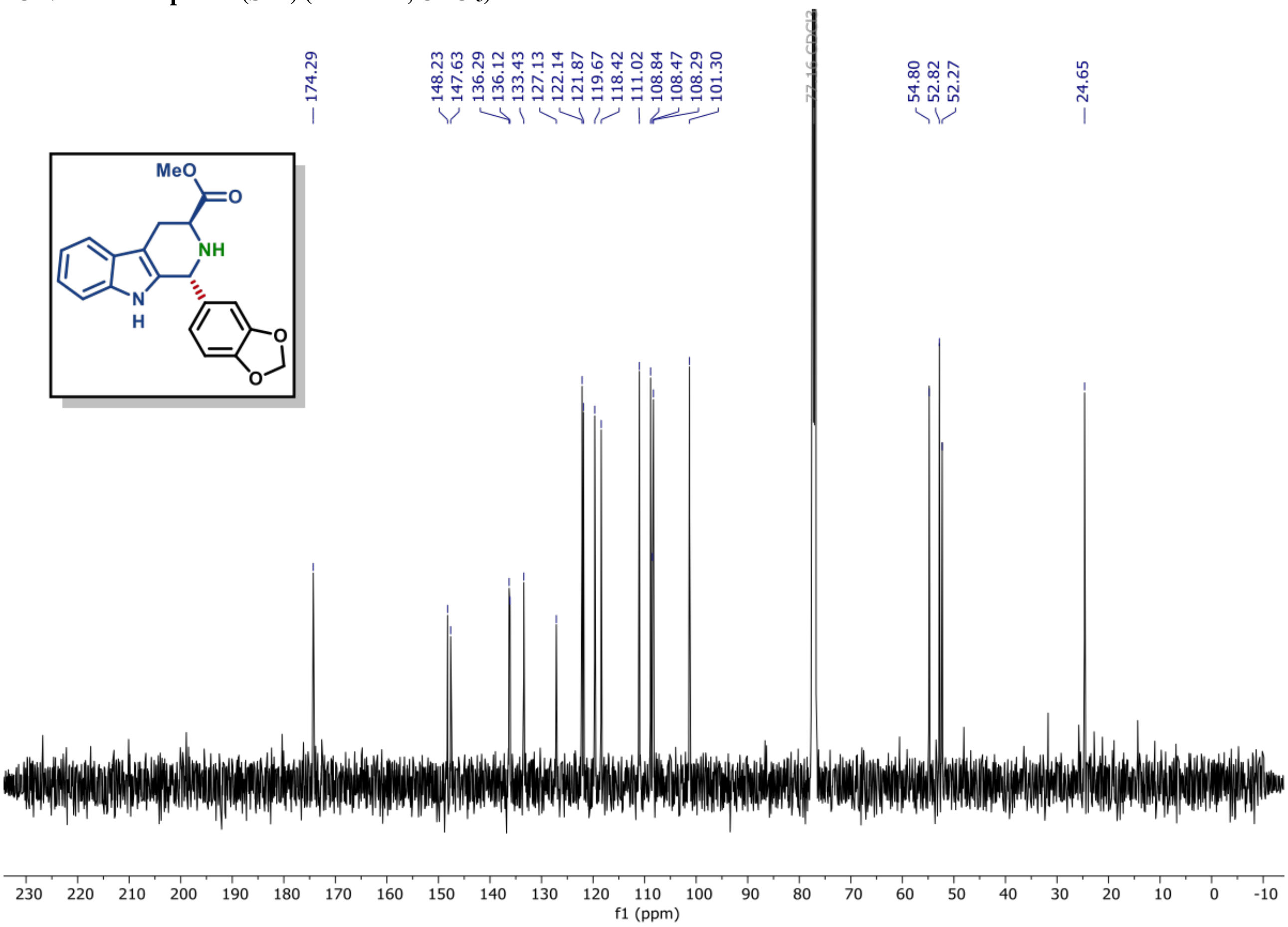

**[1]H NMR of compound (S1C) (600 MHz, CDCl$_3$)**

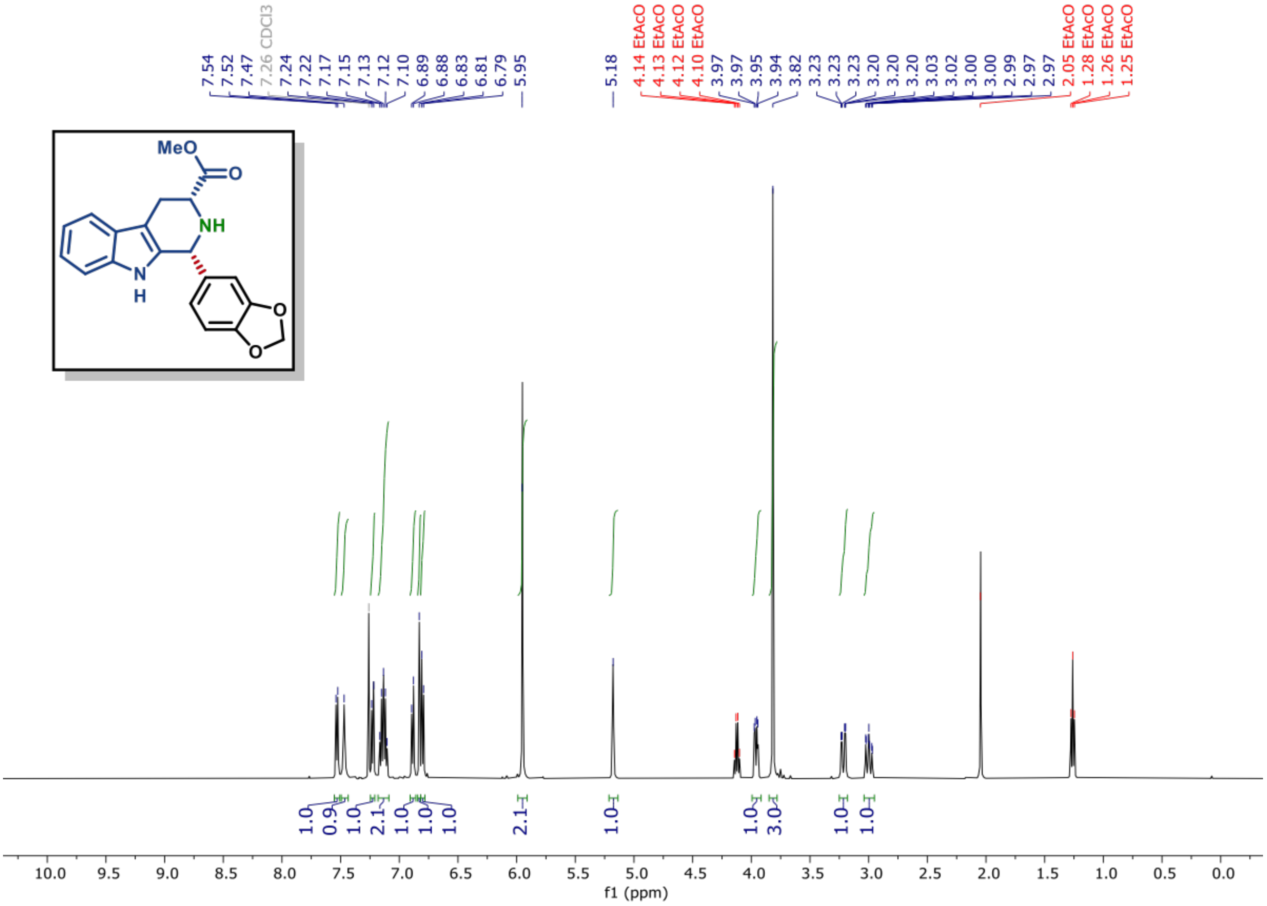

**[13]C NMR of compound (S1C) (150 MHz, CDCl$_3$)**

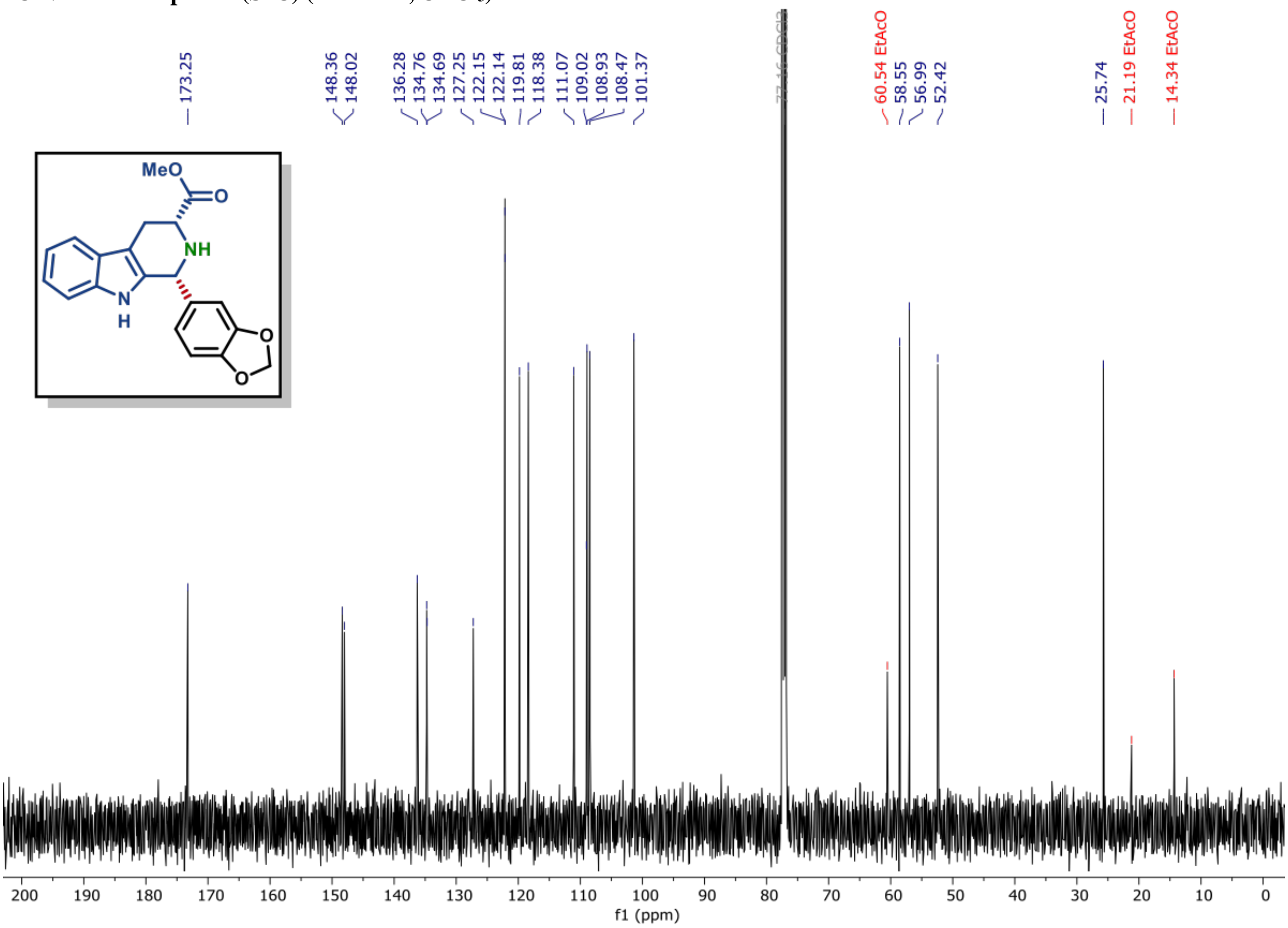

[1]H NMR of compound (S1D) (600 MHz, $CDCl_3$)

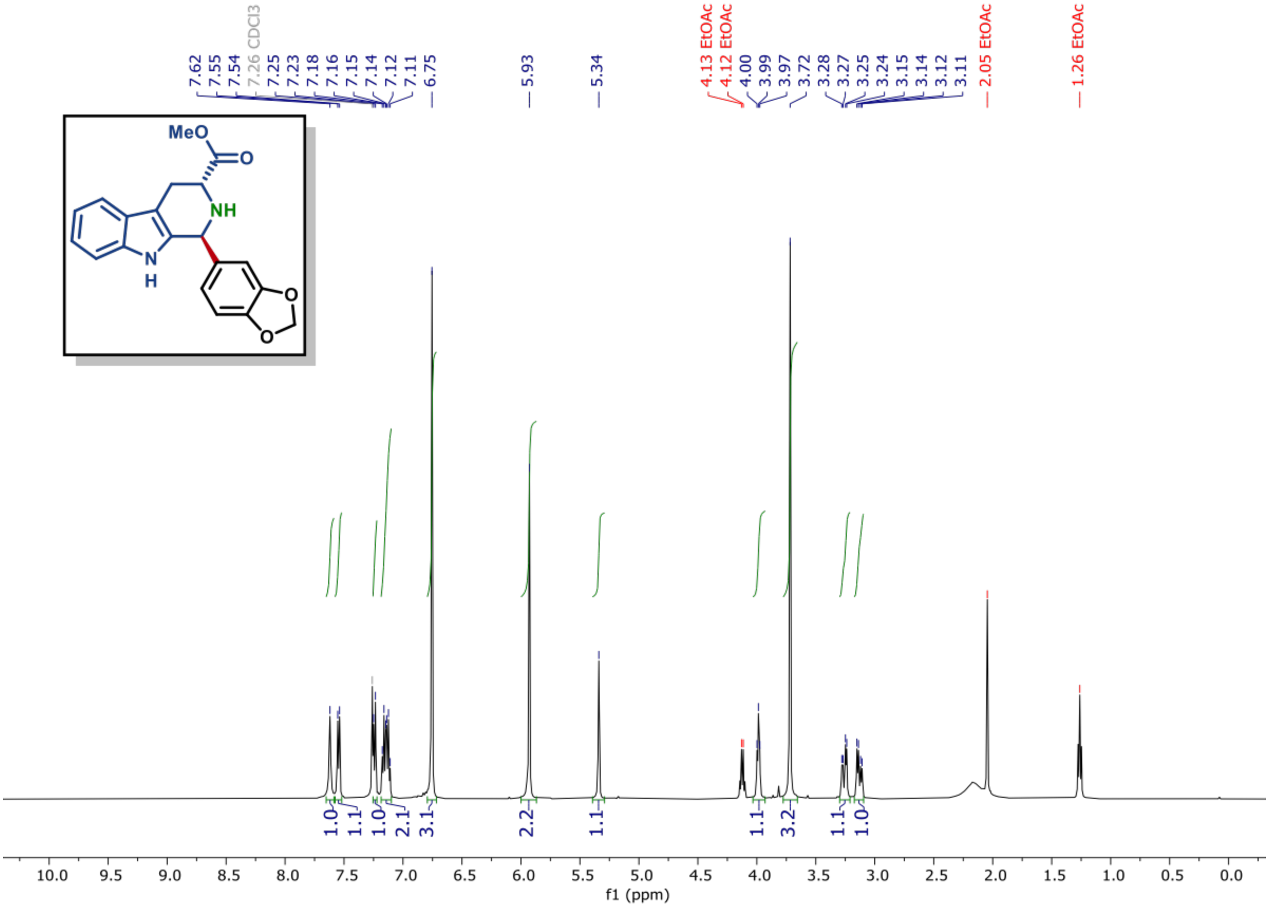

[13]C NMR of compound (S1D) (150 MHz, $CDCl_3$)

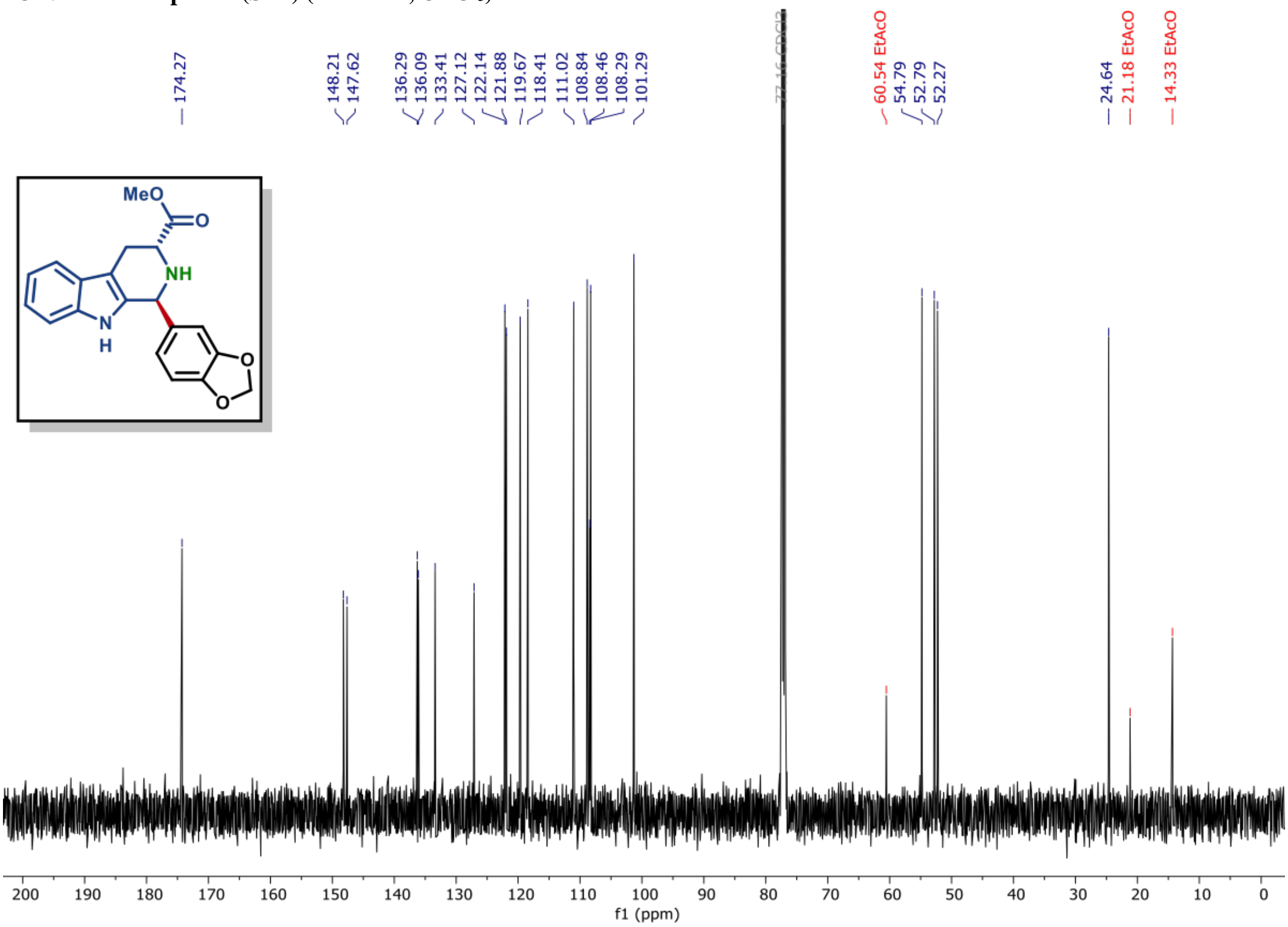

[1]H NMR of compound (S2A) (600 MHz, $CDCl_3$)

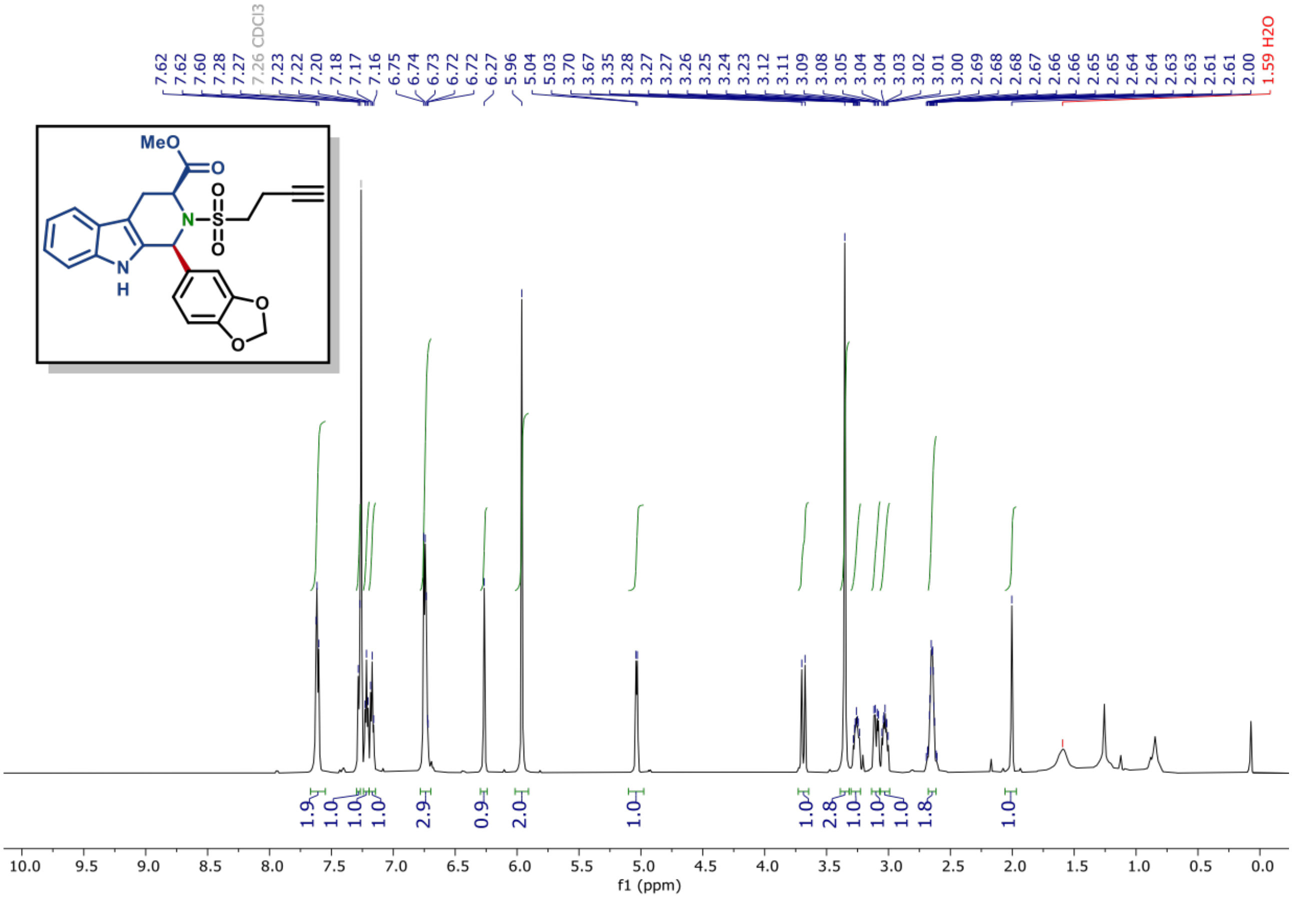

[13]C NMR of compound (S2A) (150 MHz, $CDCl_3$)

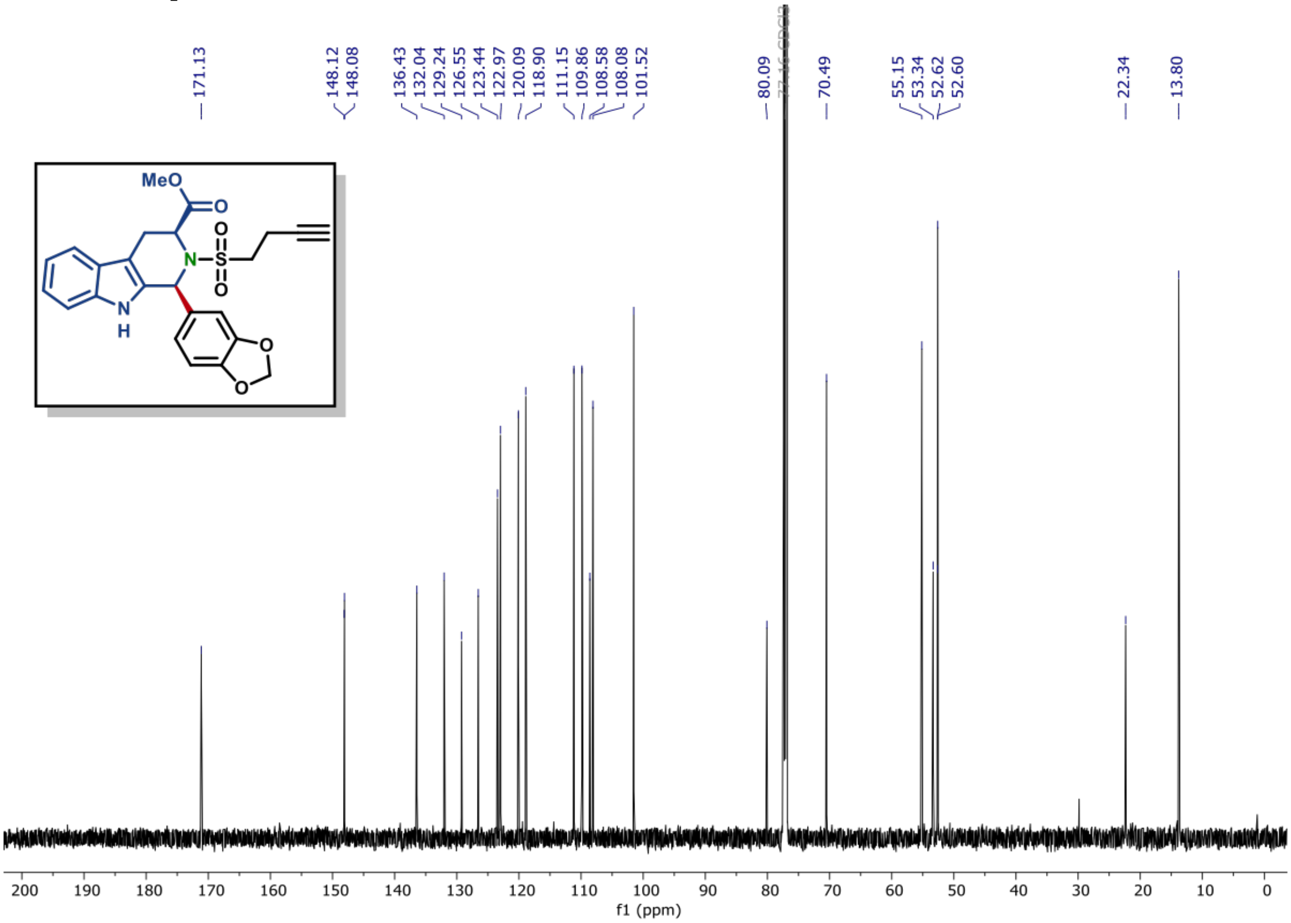

[1]H NMR of compound (S2B) (600 MHz, CDCl$_3$)

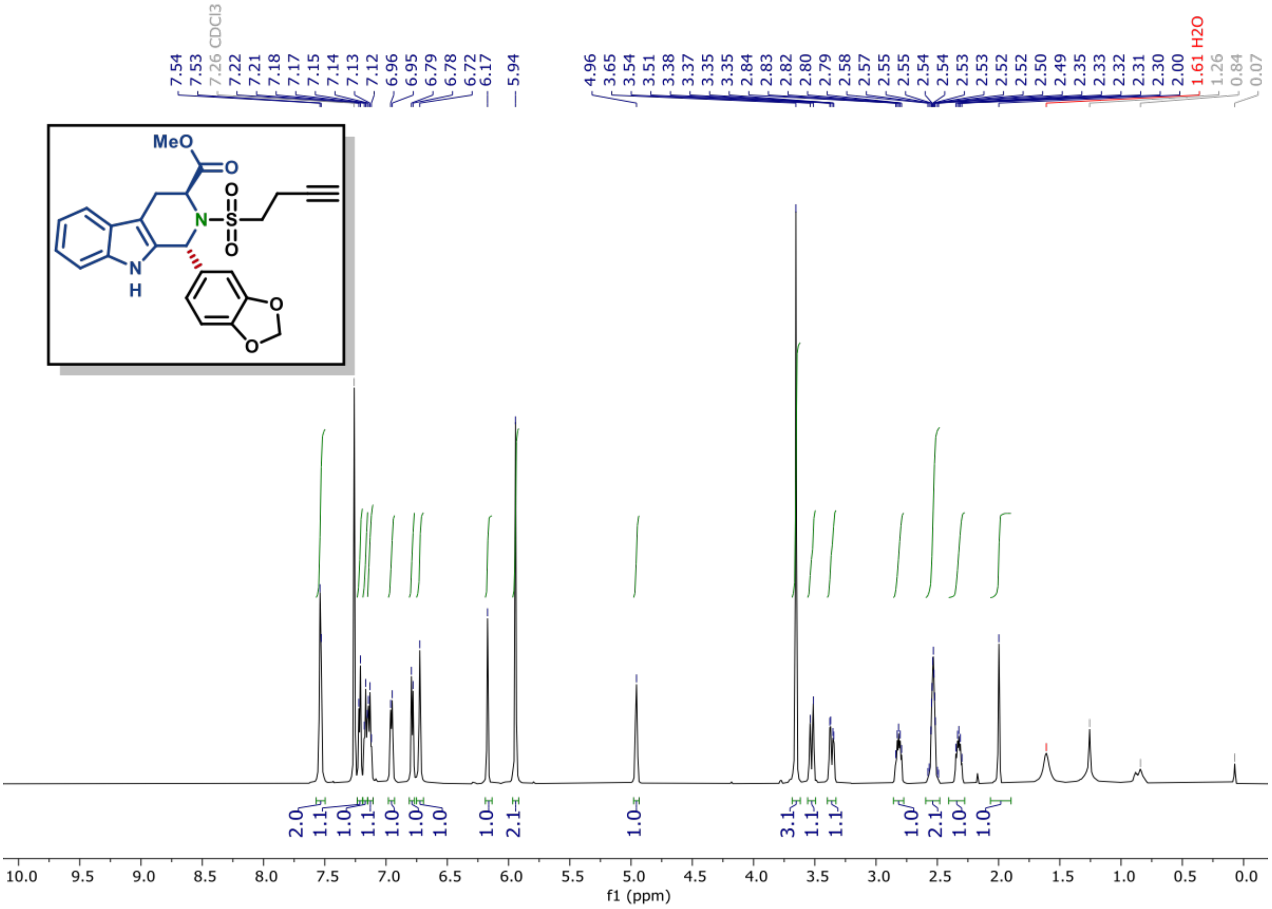

[13]C NMR of compound (S2B) (150 MHz, CDCl$_3$)

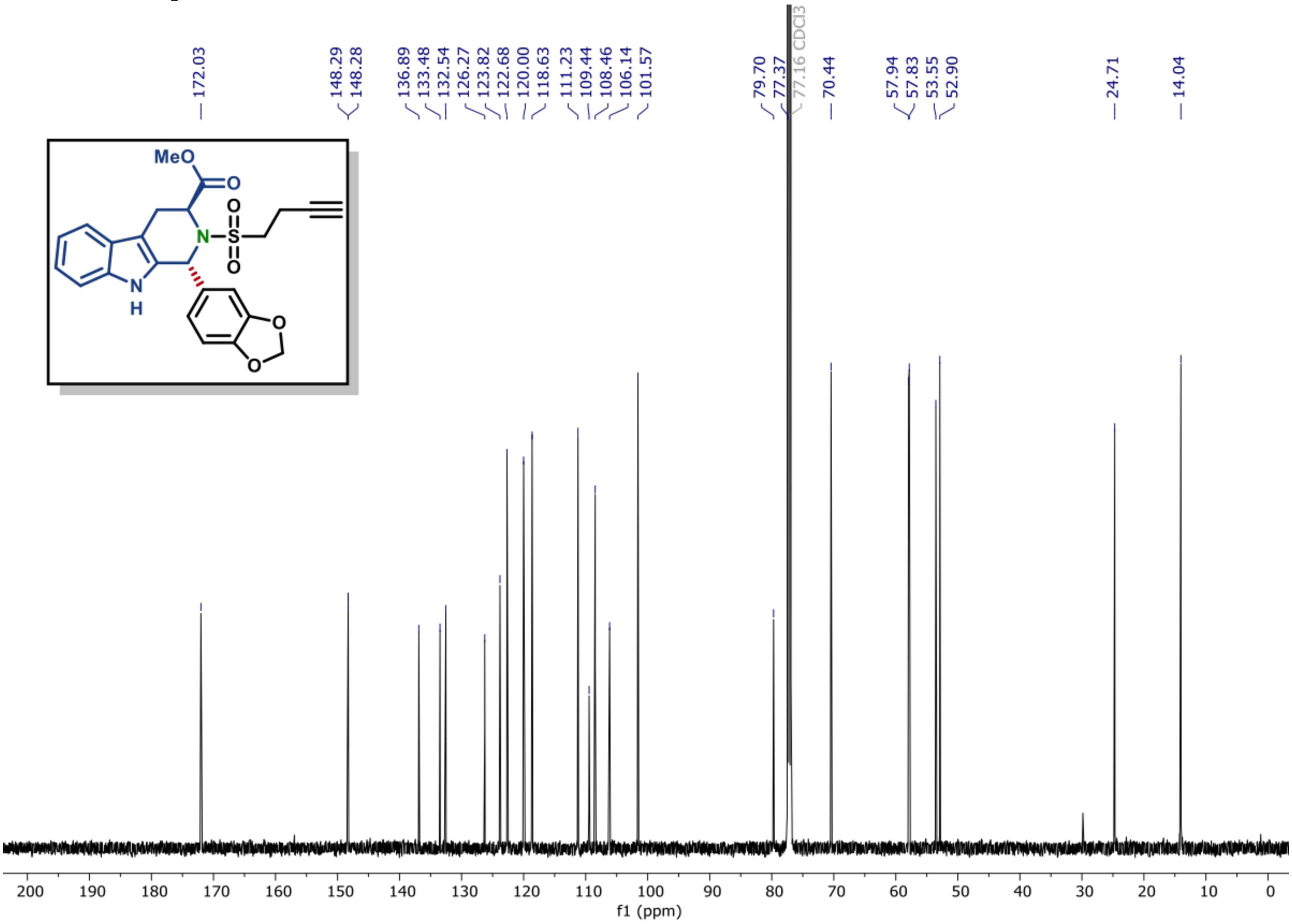

$^{1}H$ NMR of compound (S2C) (600 MHz, $CDCl_3$)

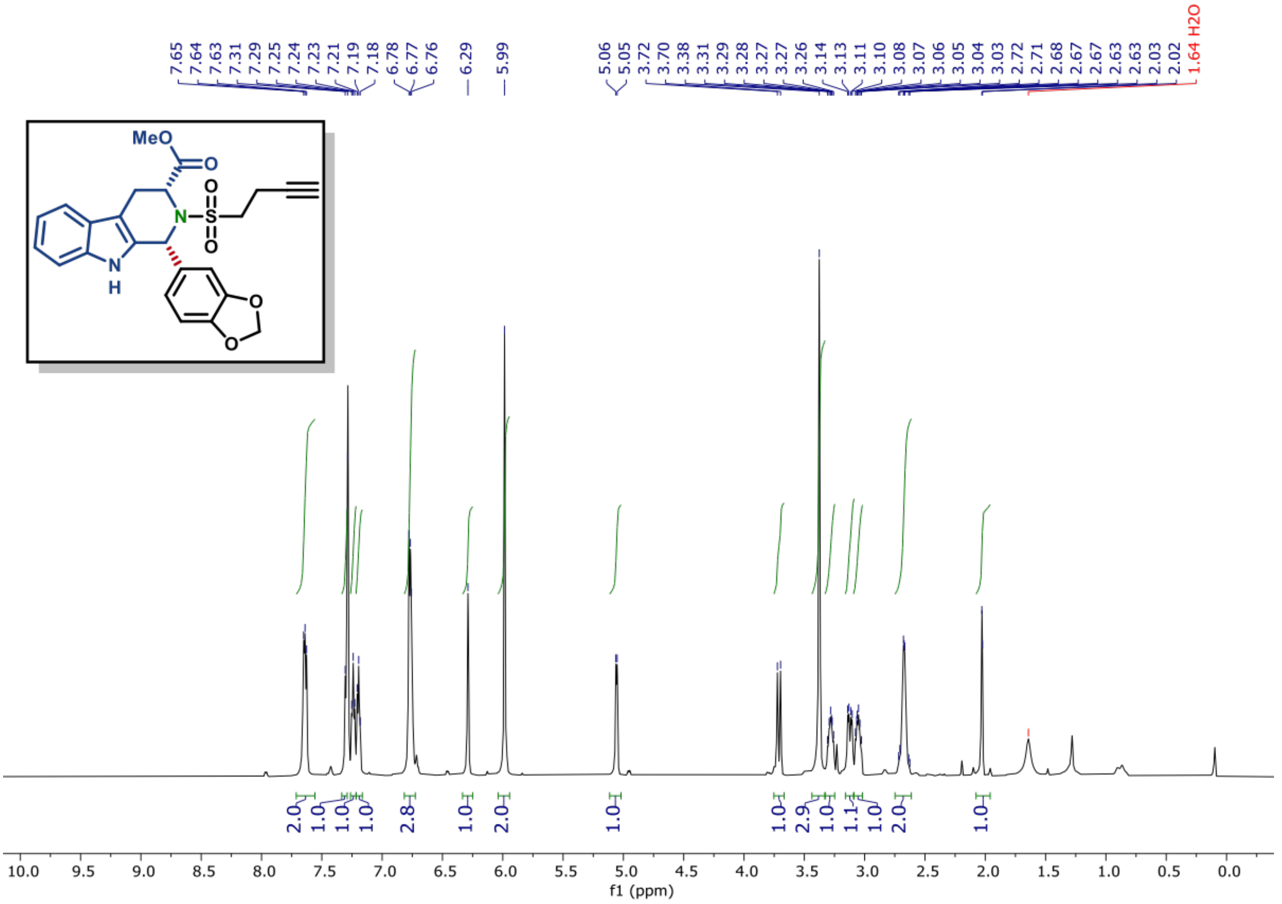

$^{13}C$ NMR of compound (S2C) (150 MHz, $CDCl_3$)

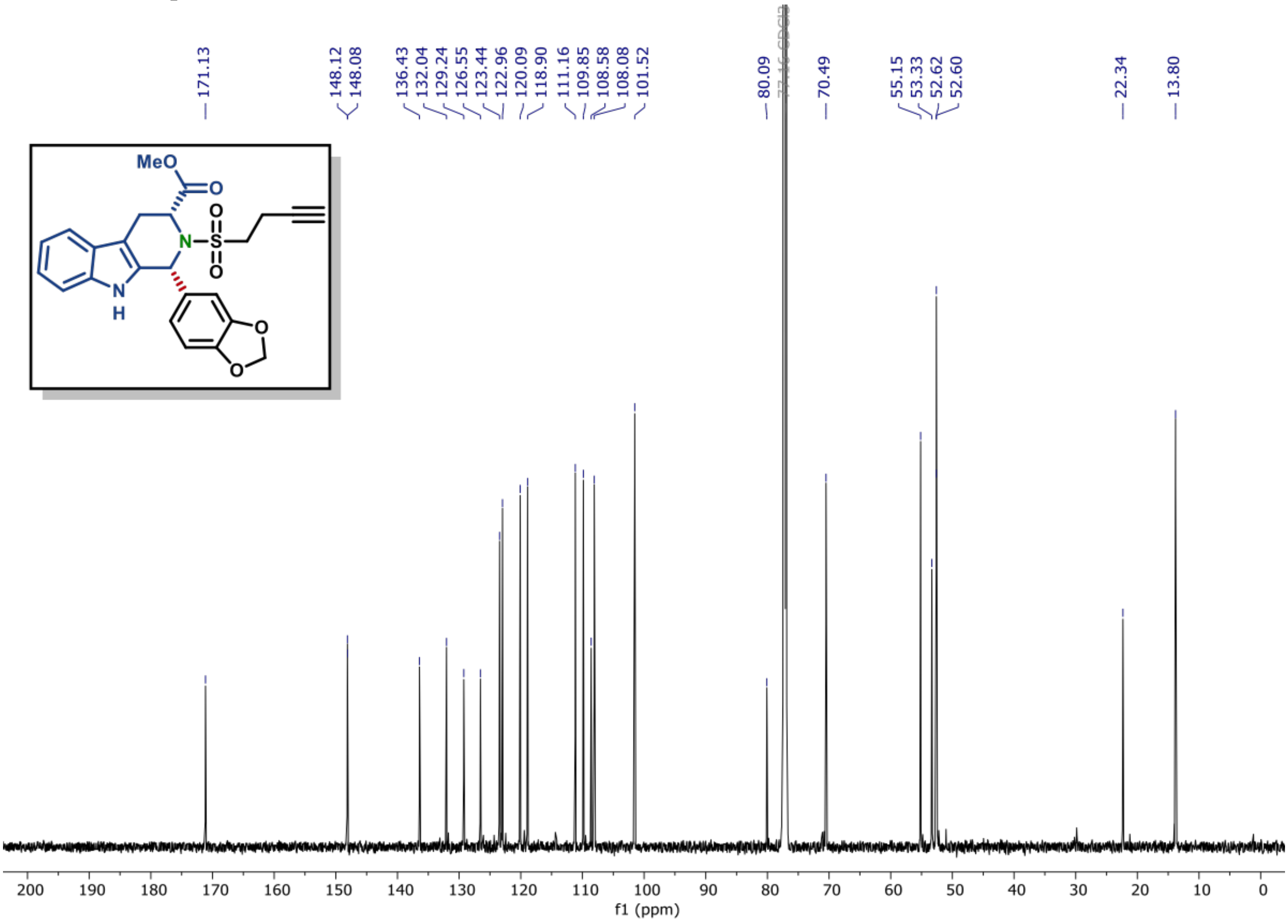

[1]H NMR of compound (S2D) (600 MHz, $CDCl_3$)

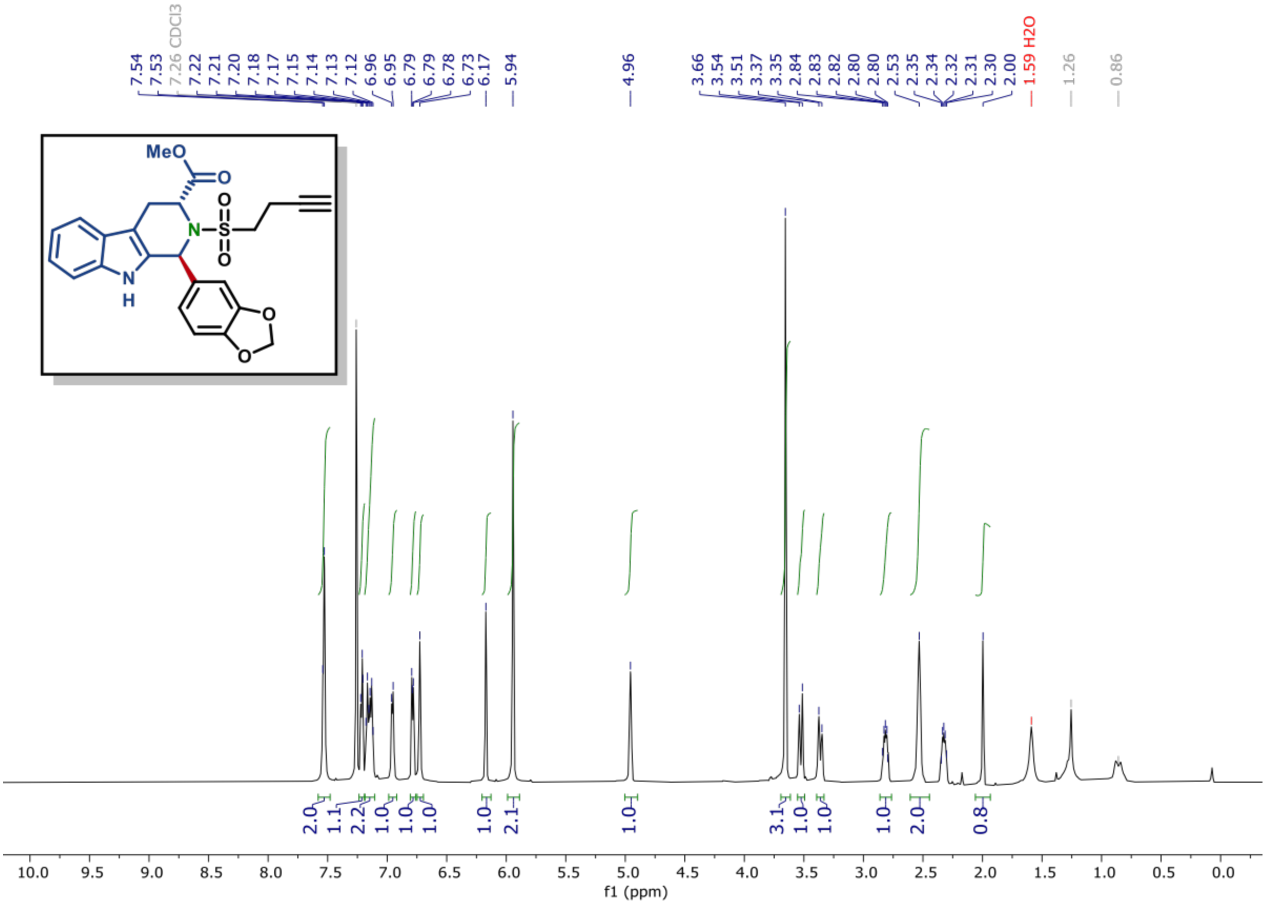

[13]C NMR of compound (S2D) (150 MHz, $CDCl_3$)

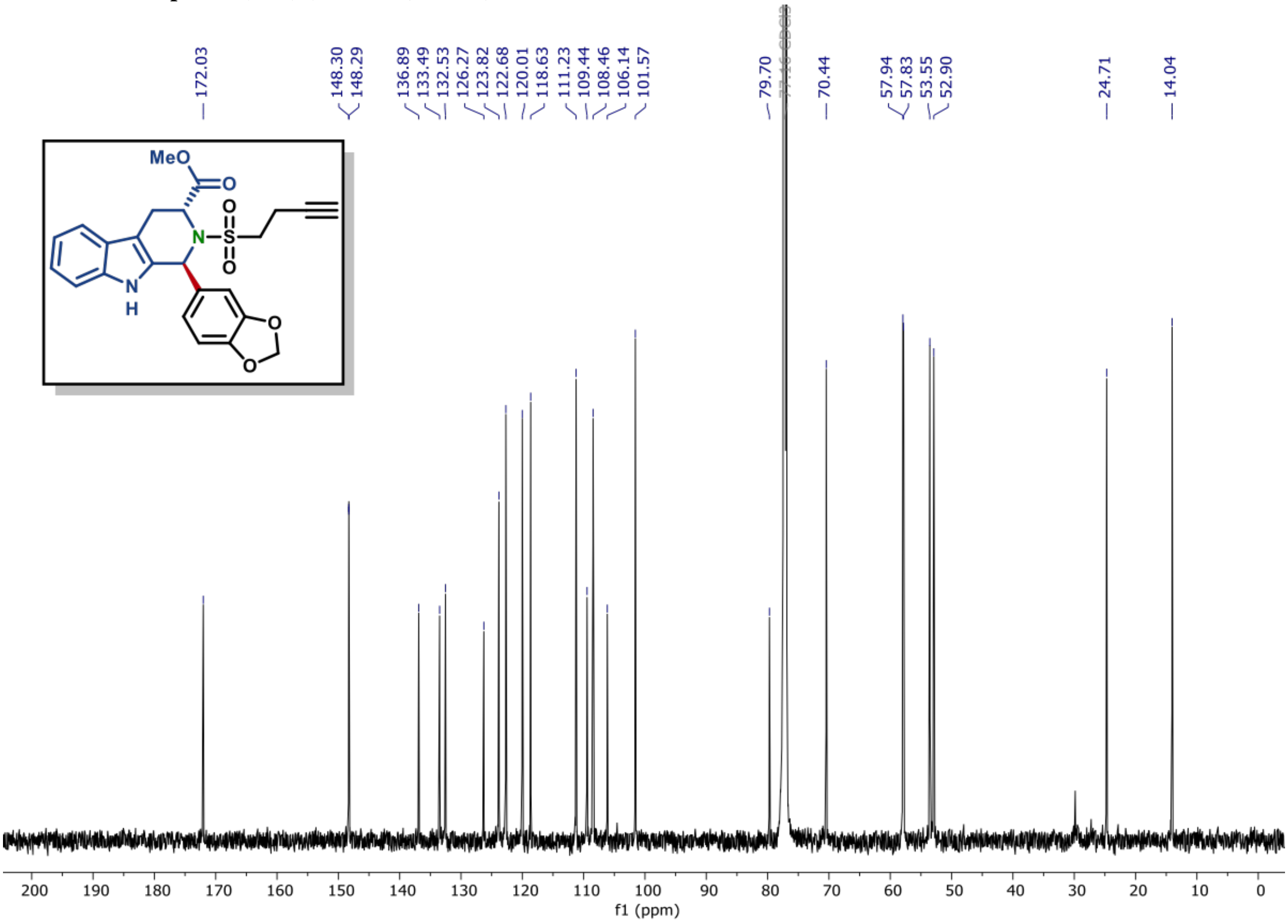

$^{1}$H NMR of compound (S3A) (600 MHz, $CDCl_3$)

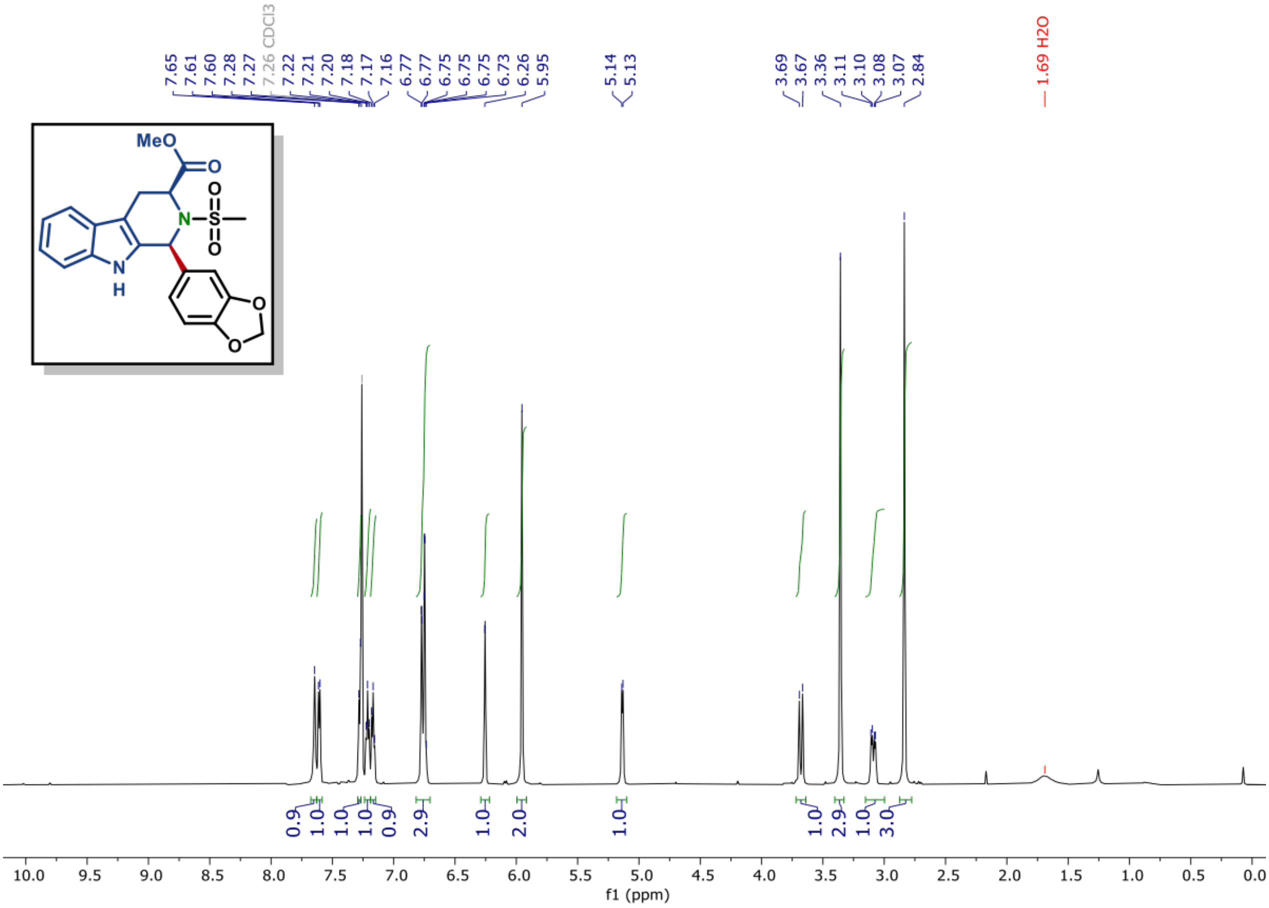

$^{13}$C NMR of compound (S3A) (150 MHz, $CDCl_3$)

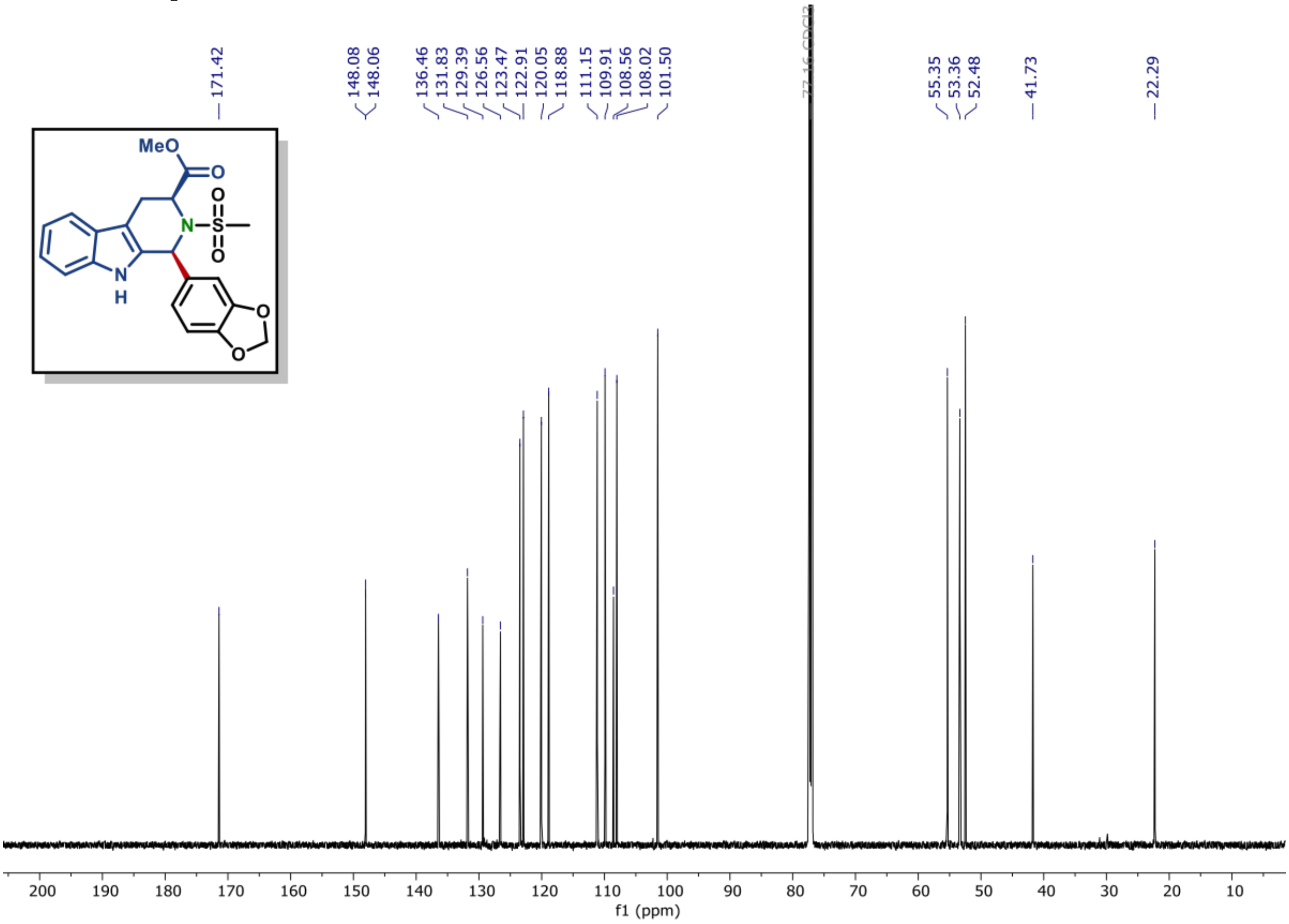

$^{1}$H NMR of compound (S3B) (600 MHz, $CDCl_3$)

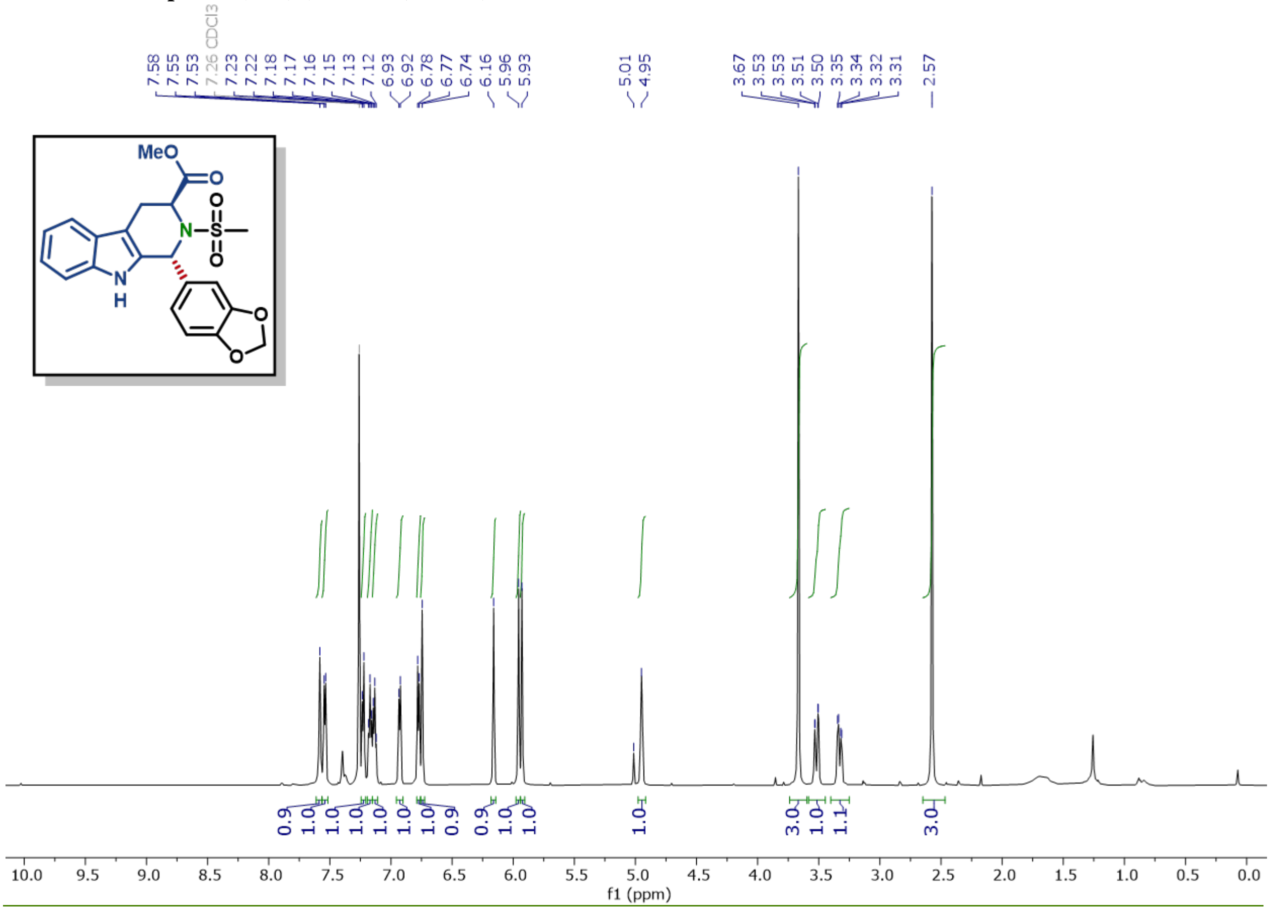

$^{13}$C NMR of compound (S3B) (150 MHz, $CDCl_3$)

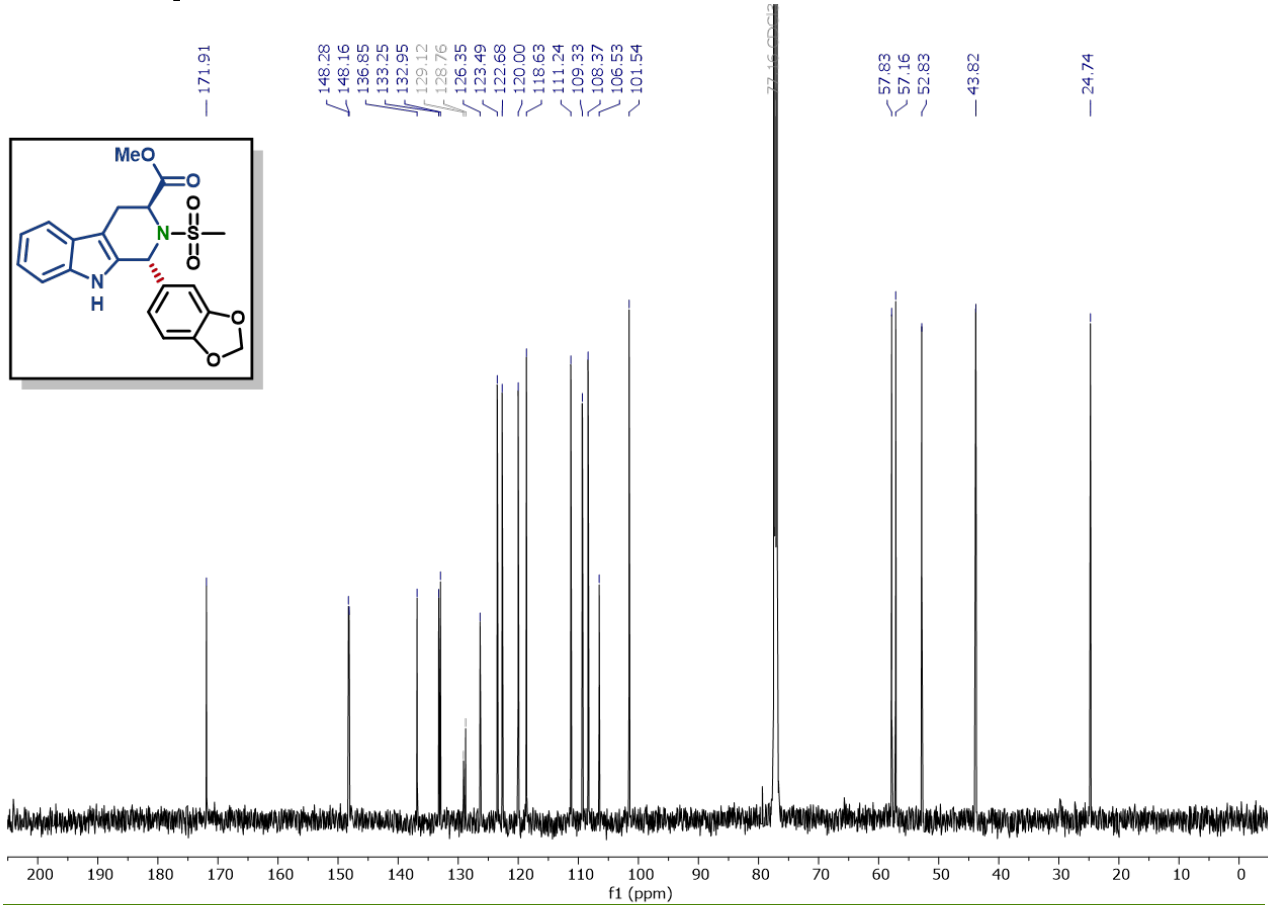

$^{1}H$ NMR of compound (S3C) (600 MHz, $CDCl_3$)

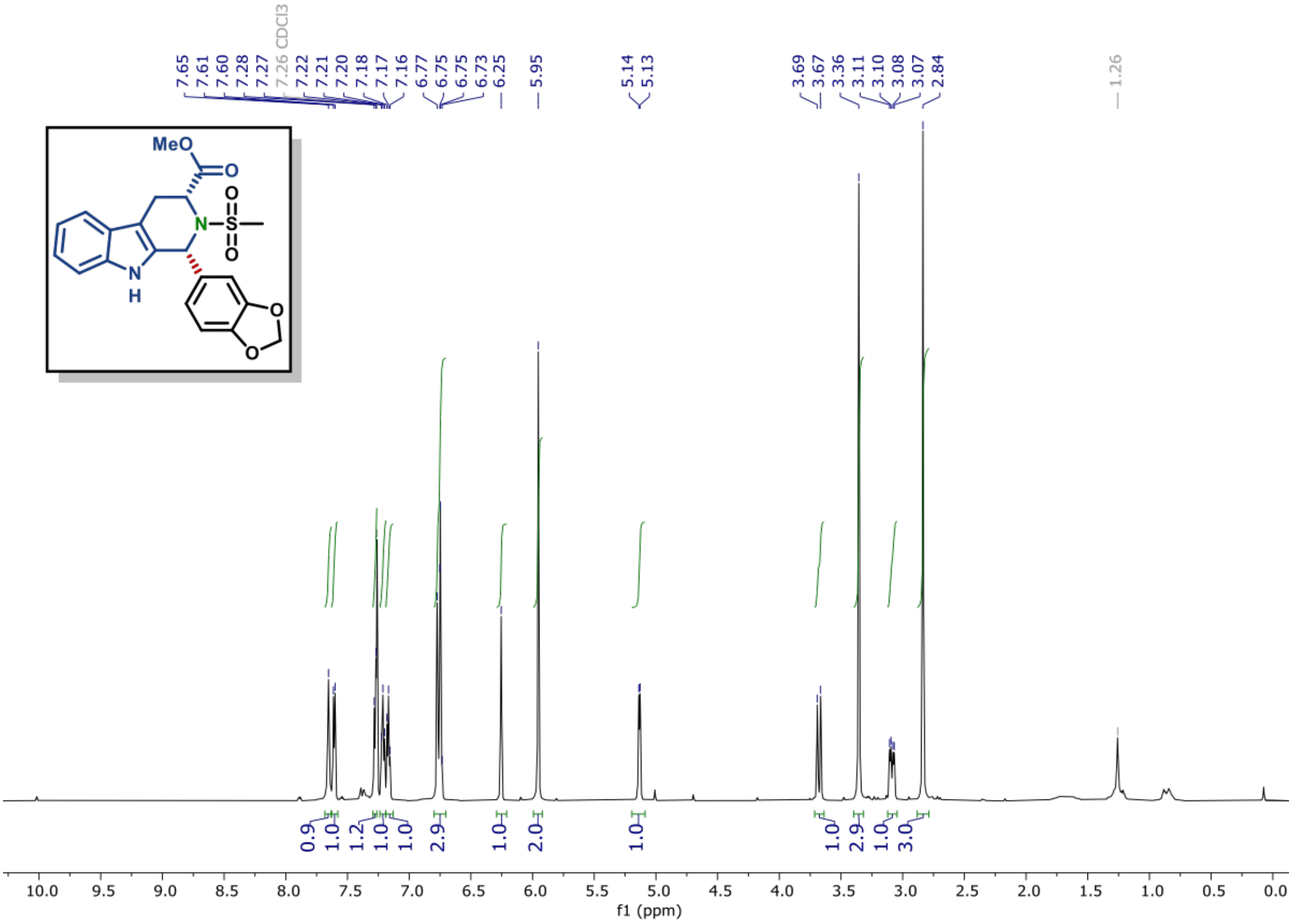

$^{13}C$ NMR of compound (S3C) (150 MHz, $CDCl_3$)

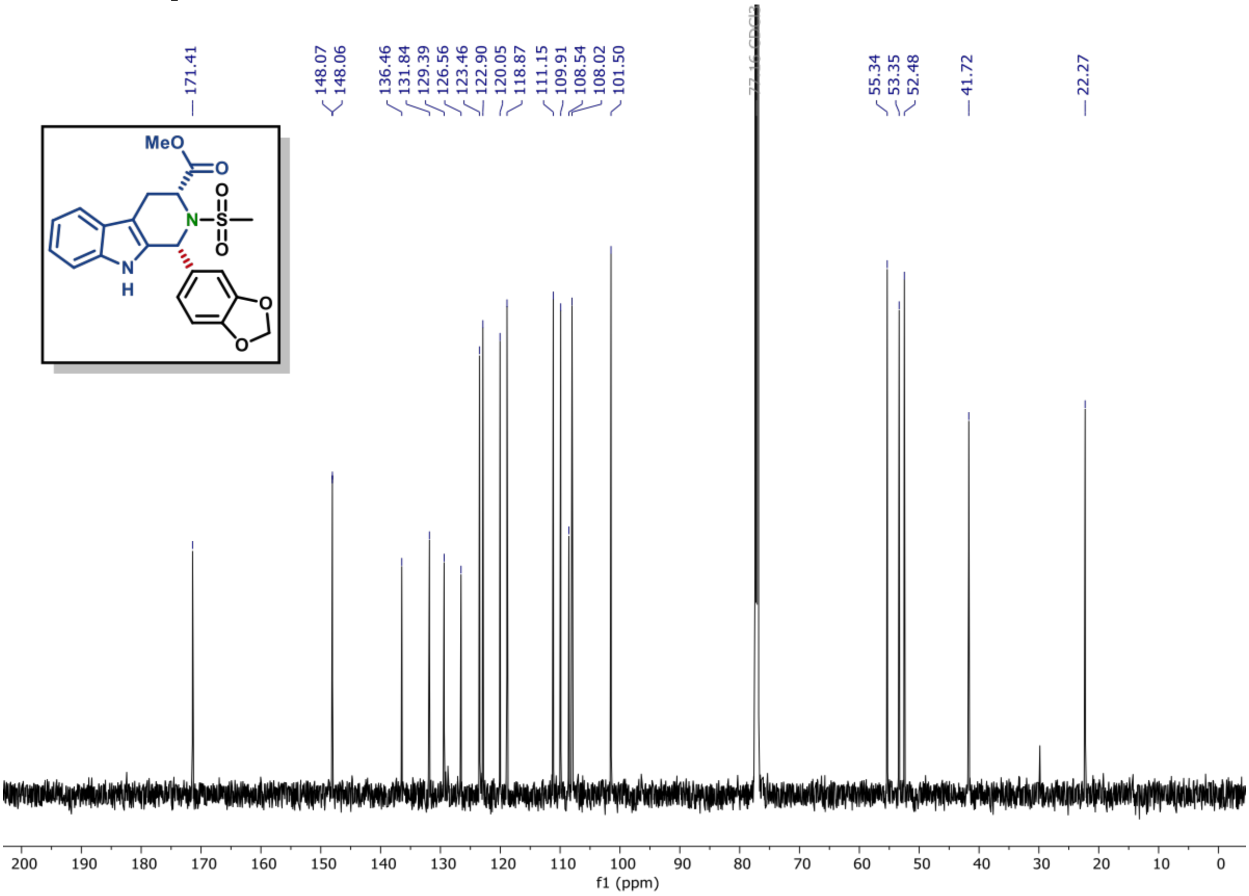

$^1H$ NMR of compound (S3D) (600 MHz, $CDCl_3$)

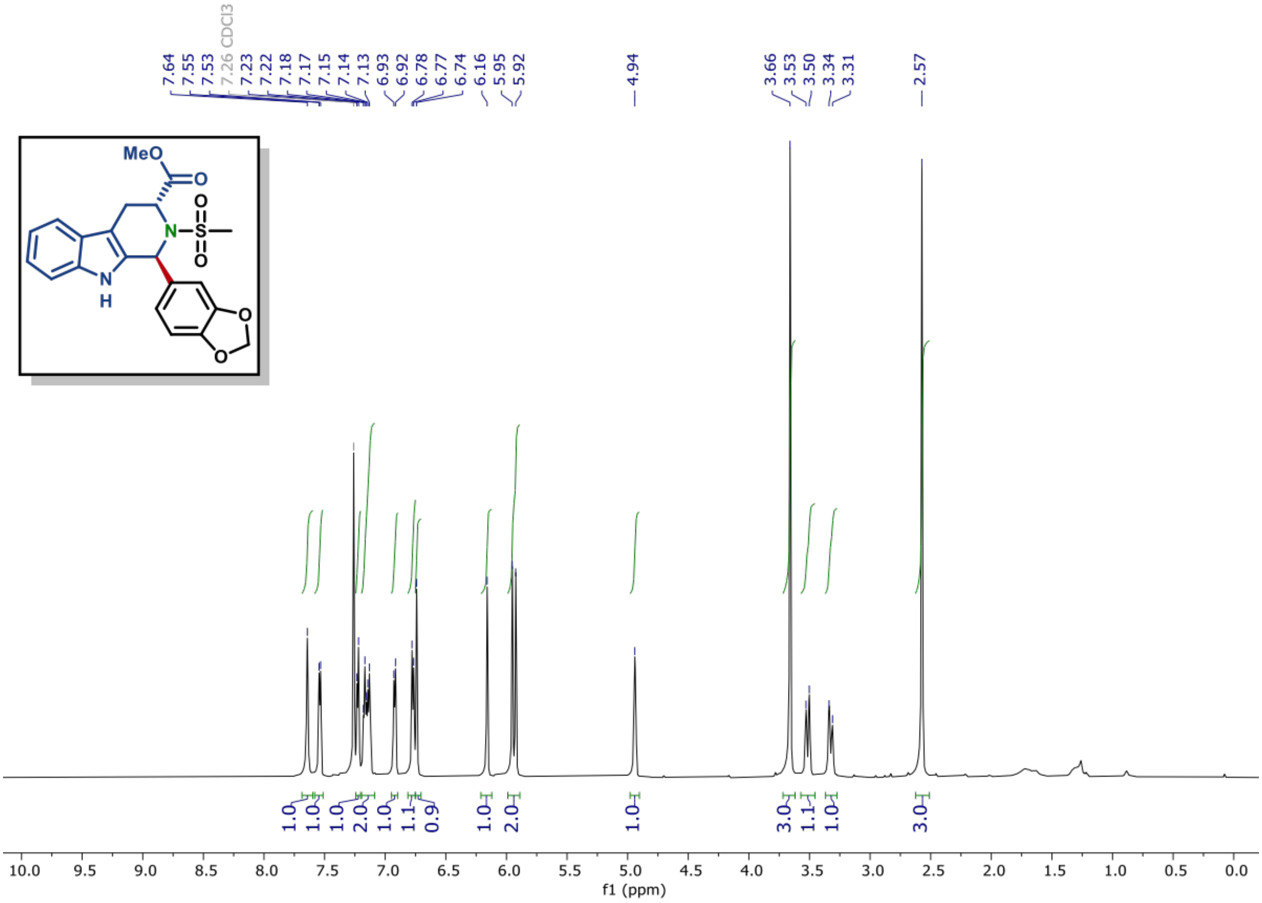

$^{13}C$ NMR of compound (S3D) (150 MHz, $CDCl_3$)

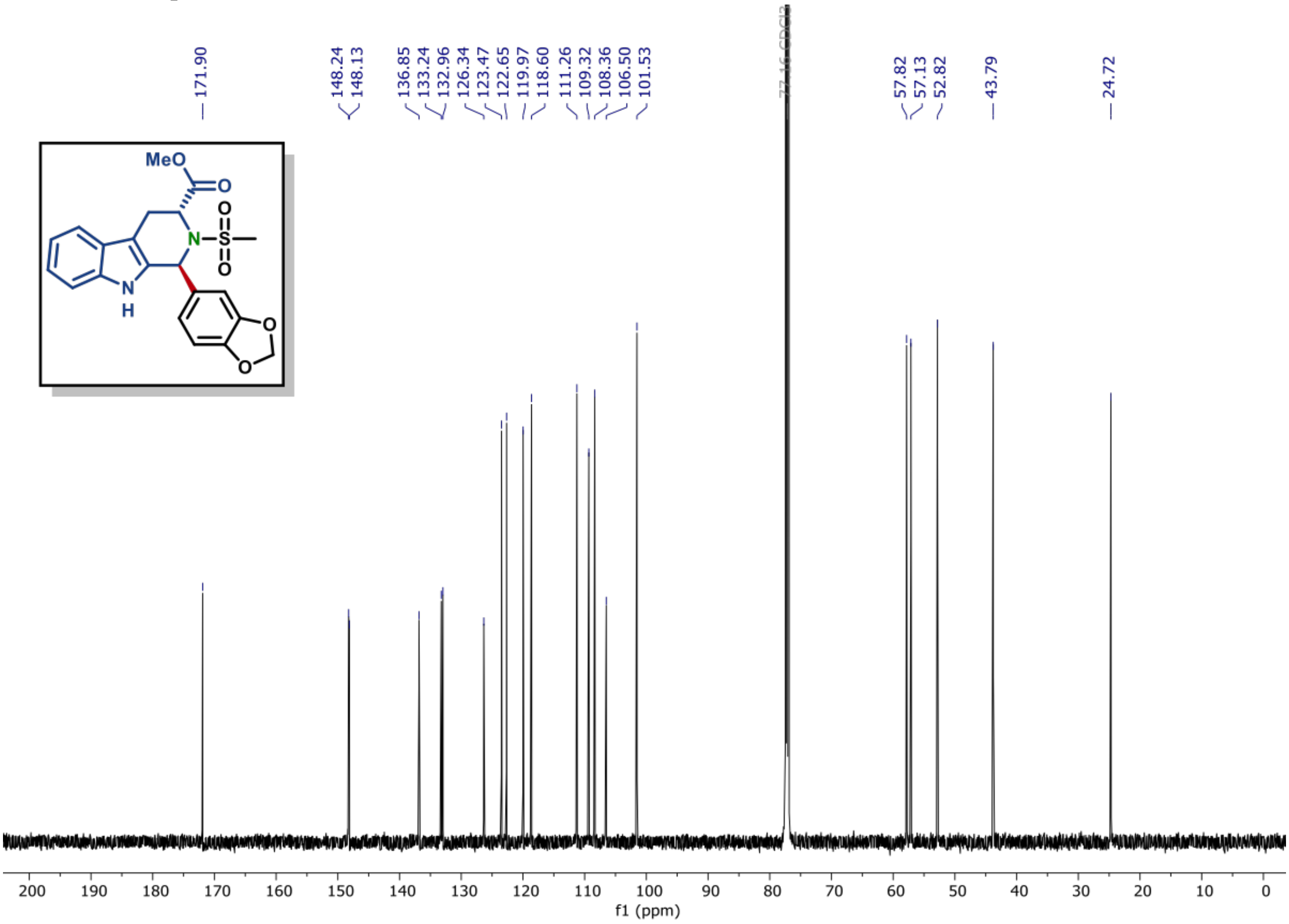

**$^{31}$P NMR of compound (17) (162 MHz, $CDCl_3$)**

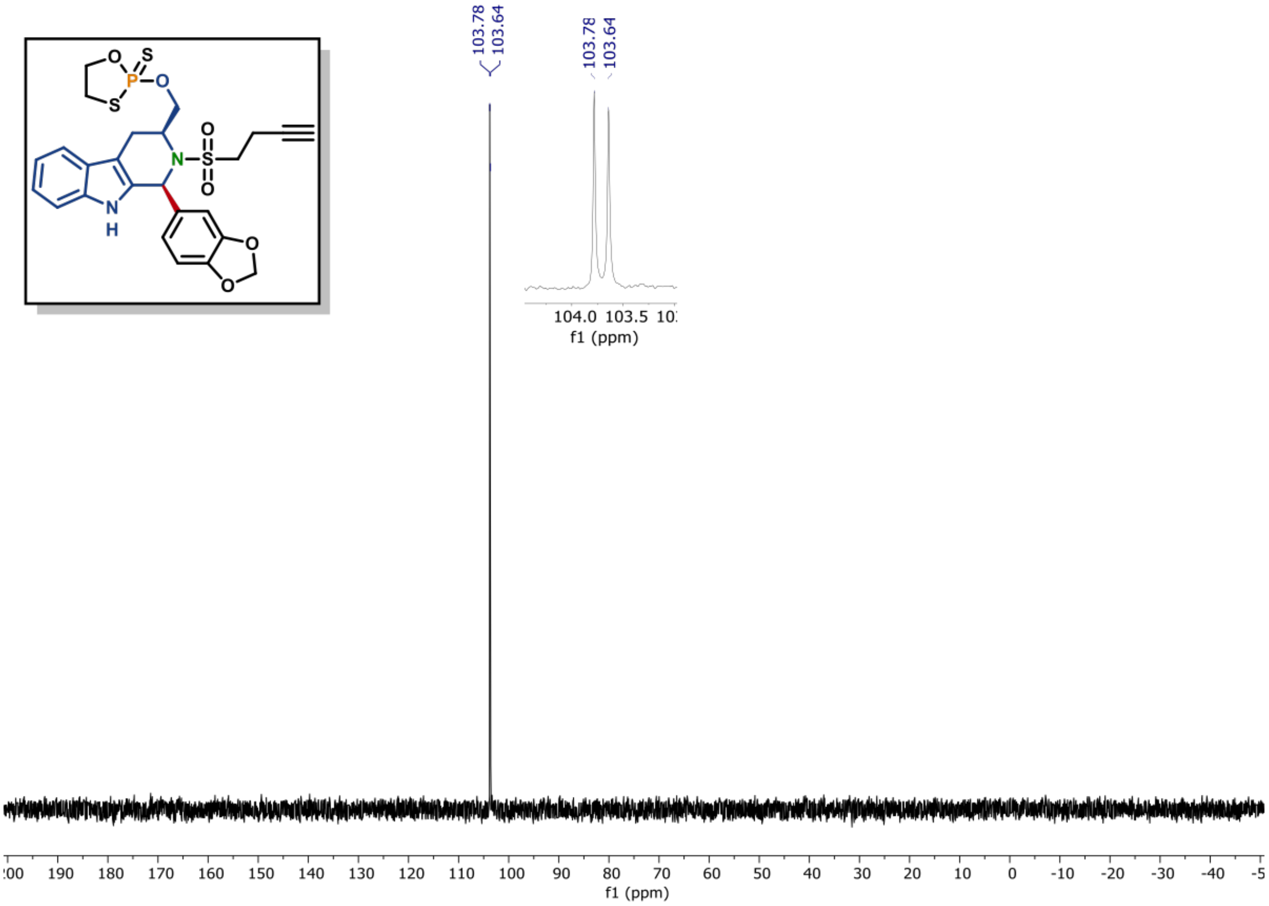

**$^{1}$H NMR of compound (17) (600 MHz, $CDCl_3$)**

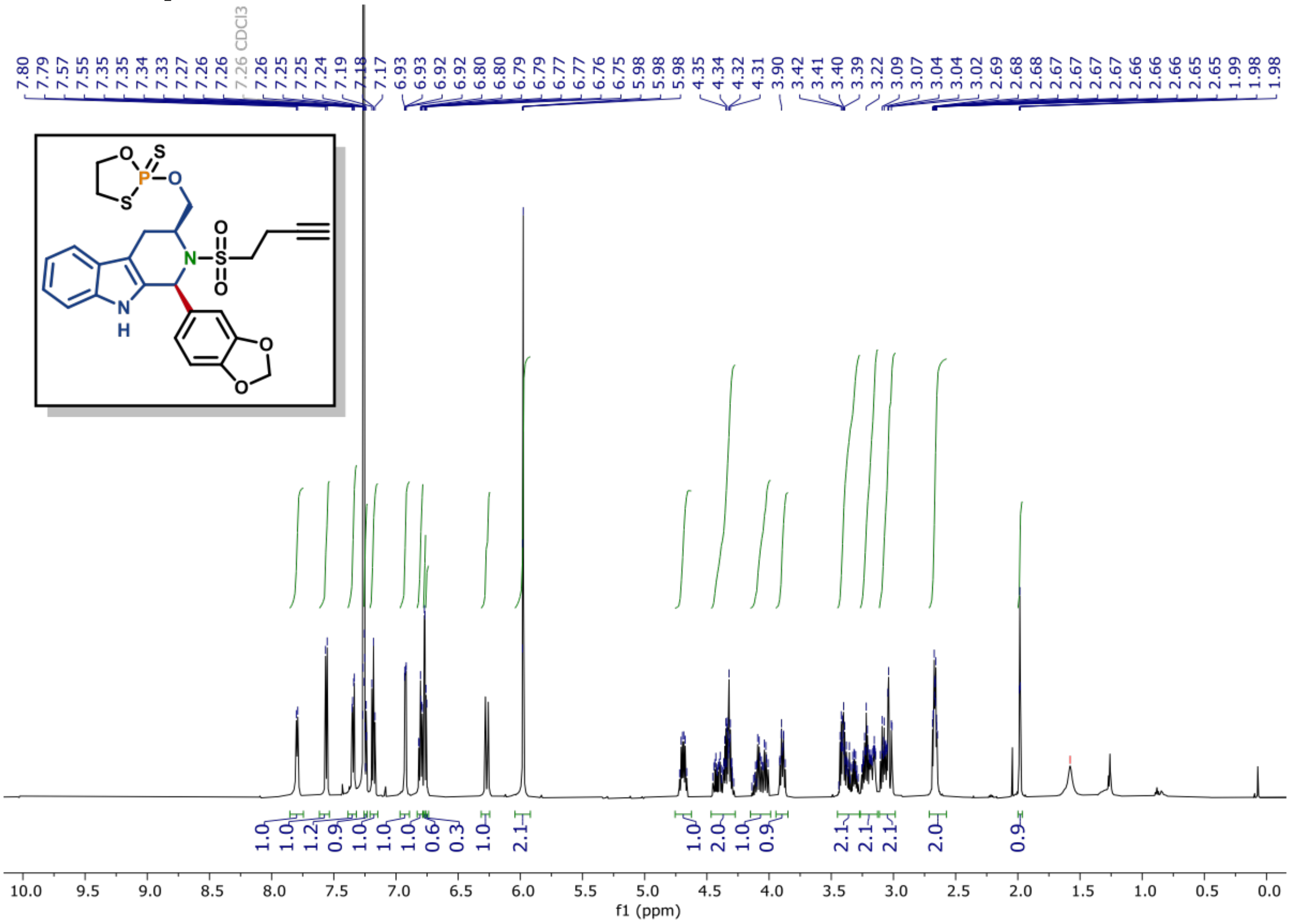

[13]C NMR of compound (17) (150 MHz, CDCl3)

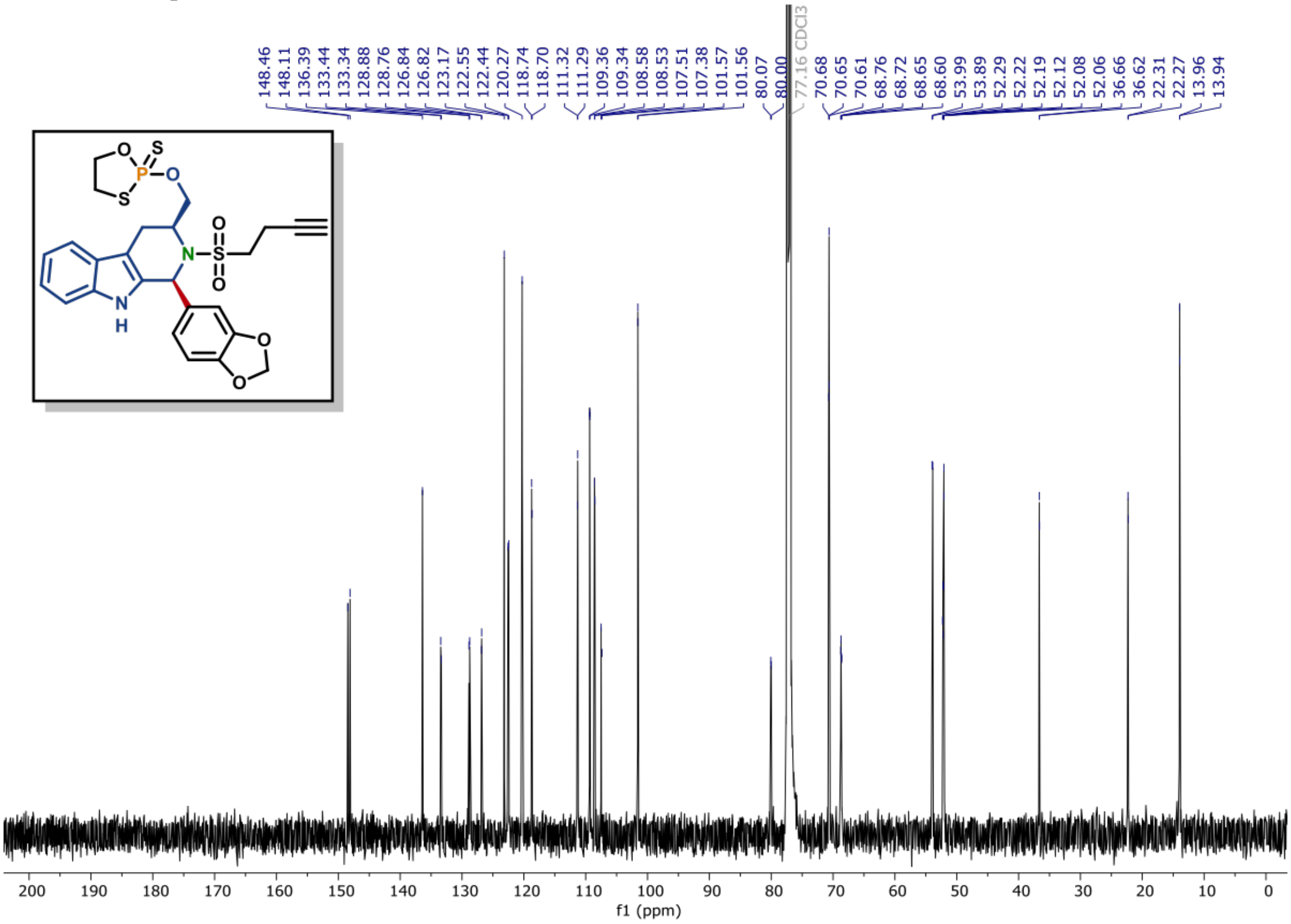

**[31]P NMR of compound (19) (162 MHz, $CDCl_3$)**

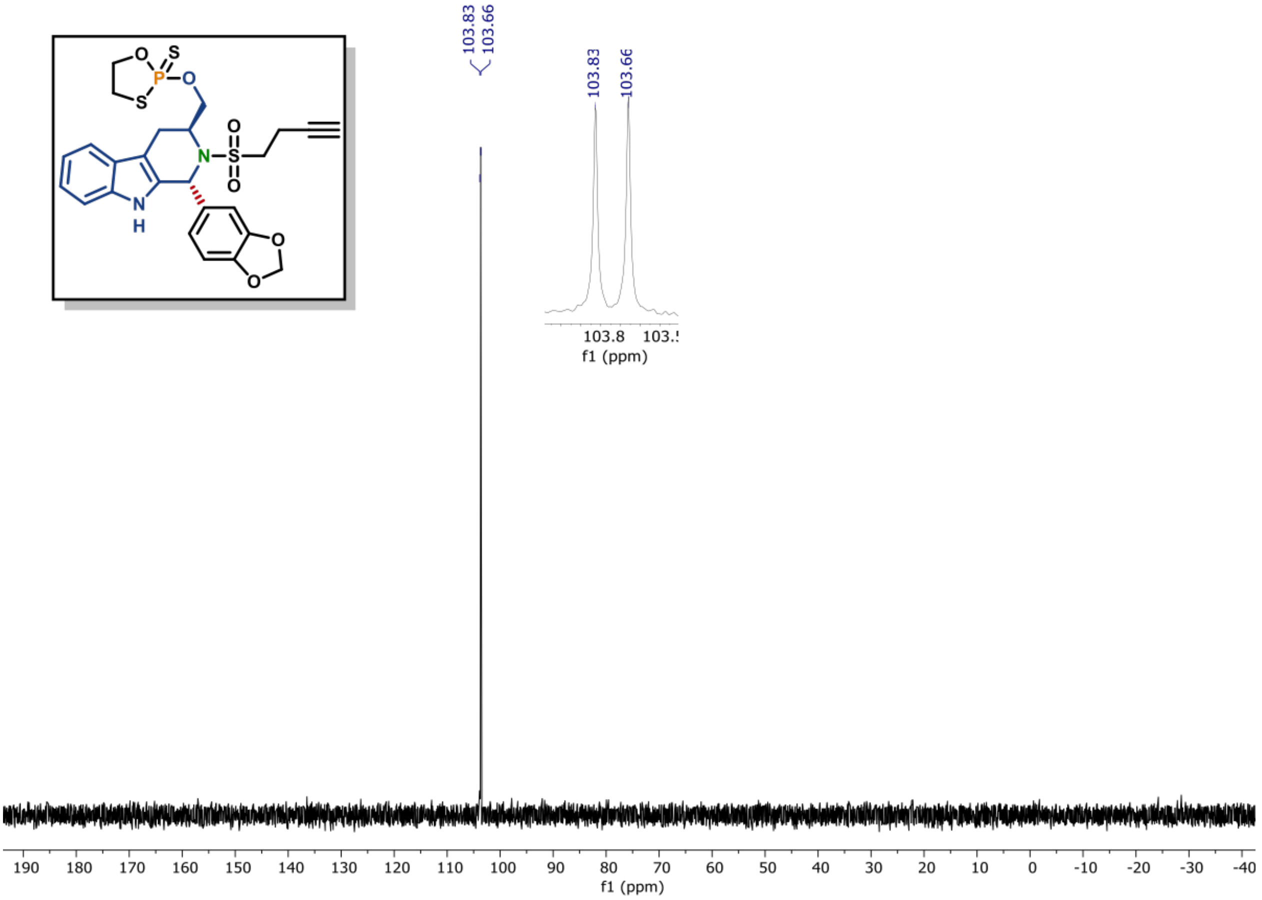

**[1]H NMR of compound (19) (600 MHz, $CDCl_3$)**

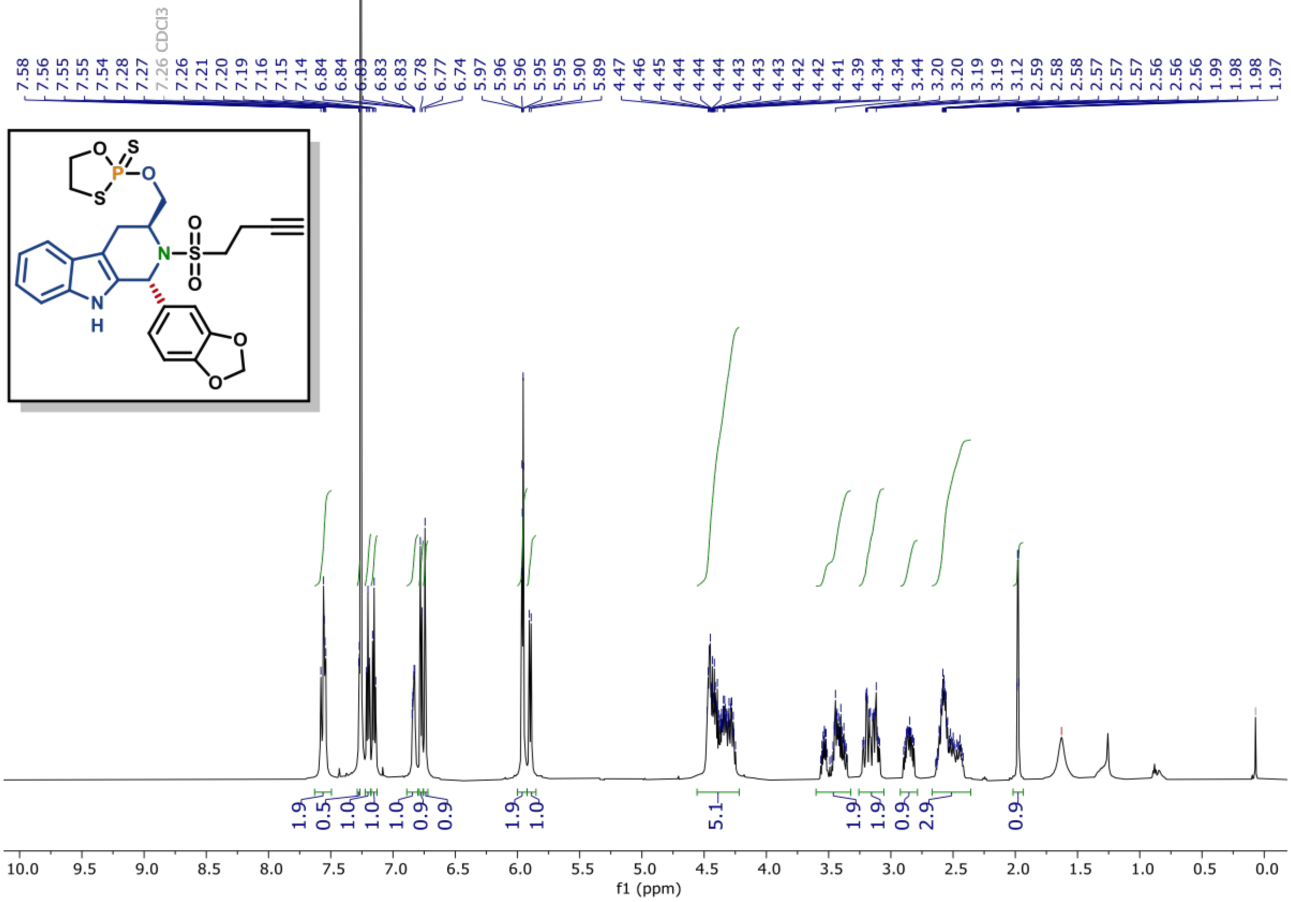

[13]C NMR of compound (19) (150 MHz, CDCl$_3$)

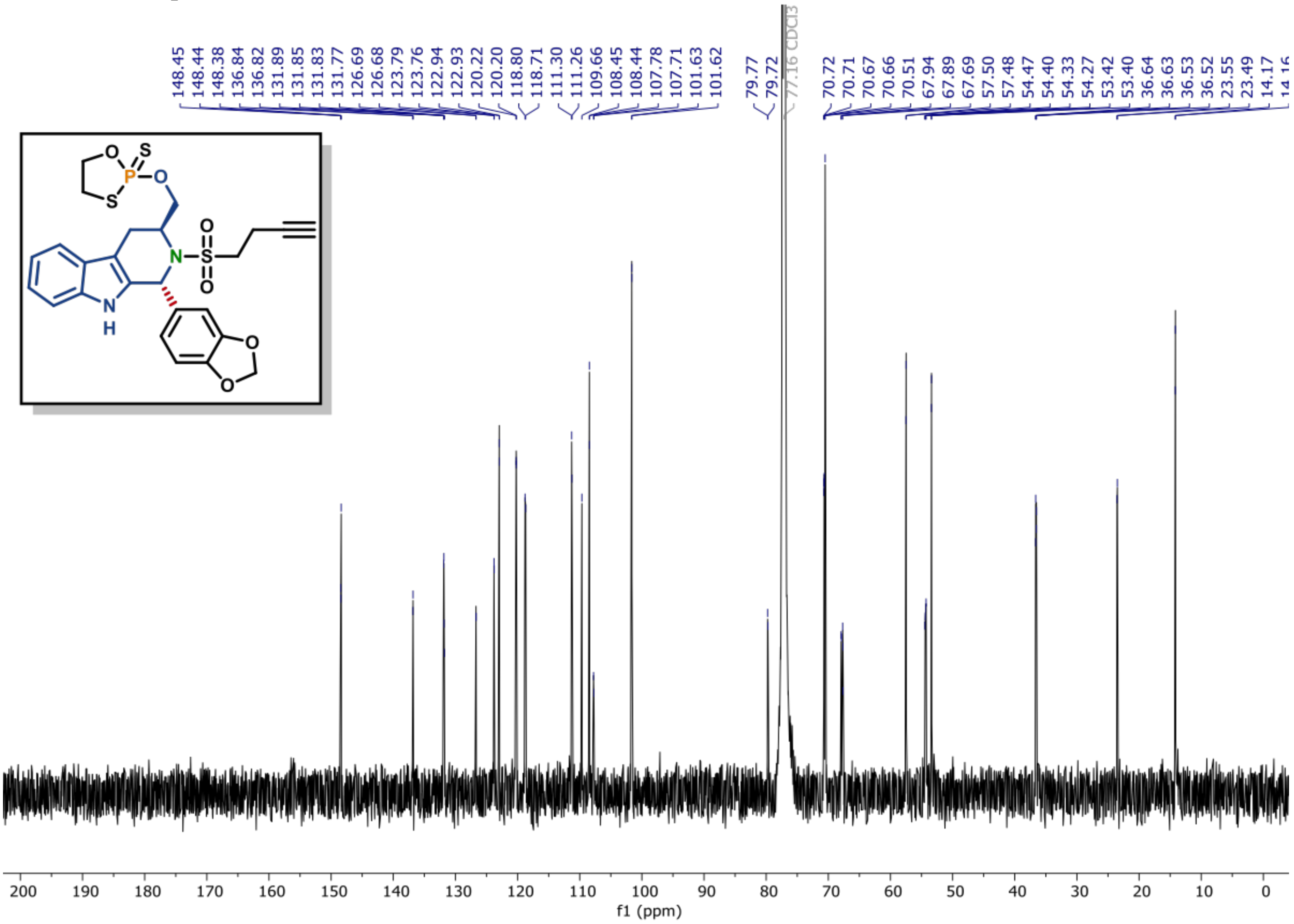

$^{31}$P NMR of compound (18) (162 MHz, $CDCl_3$)

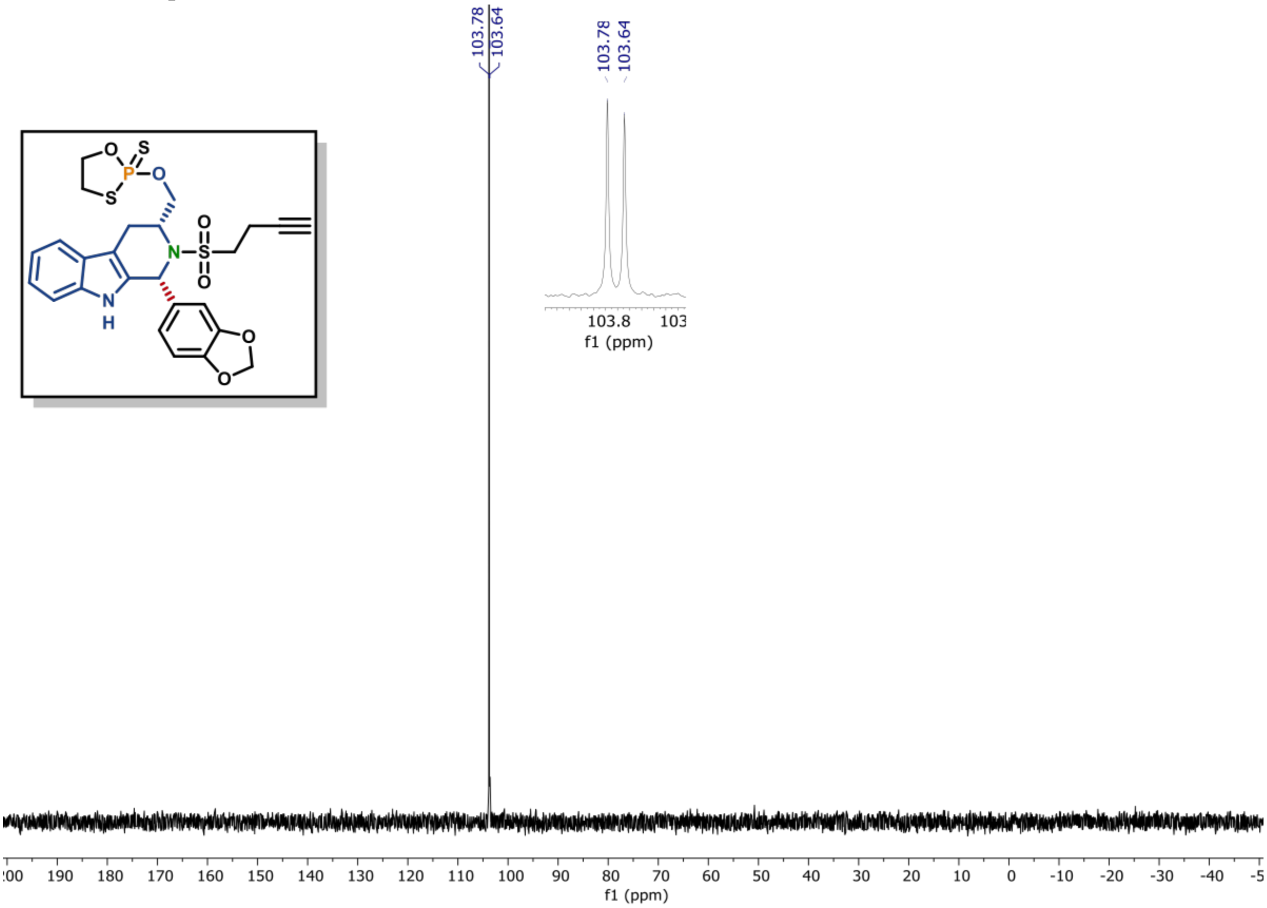

$^{1}$H NMR of compound (18) (600 MHz, $CDCl_3$)

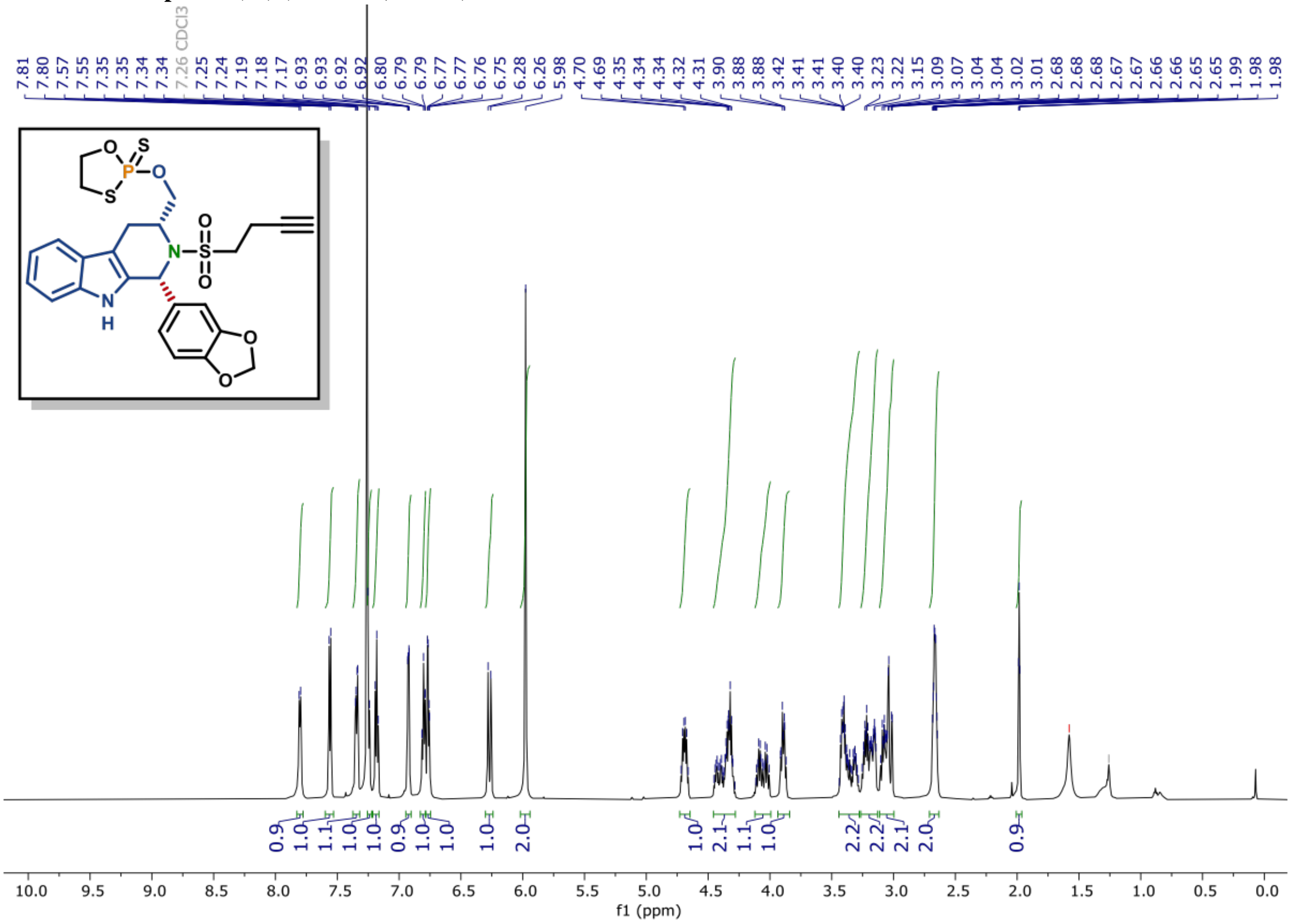

**$^{13}$C NMR of compound (18) (150 MHz, CDCl$_3$)**

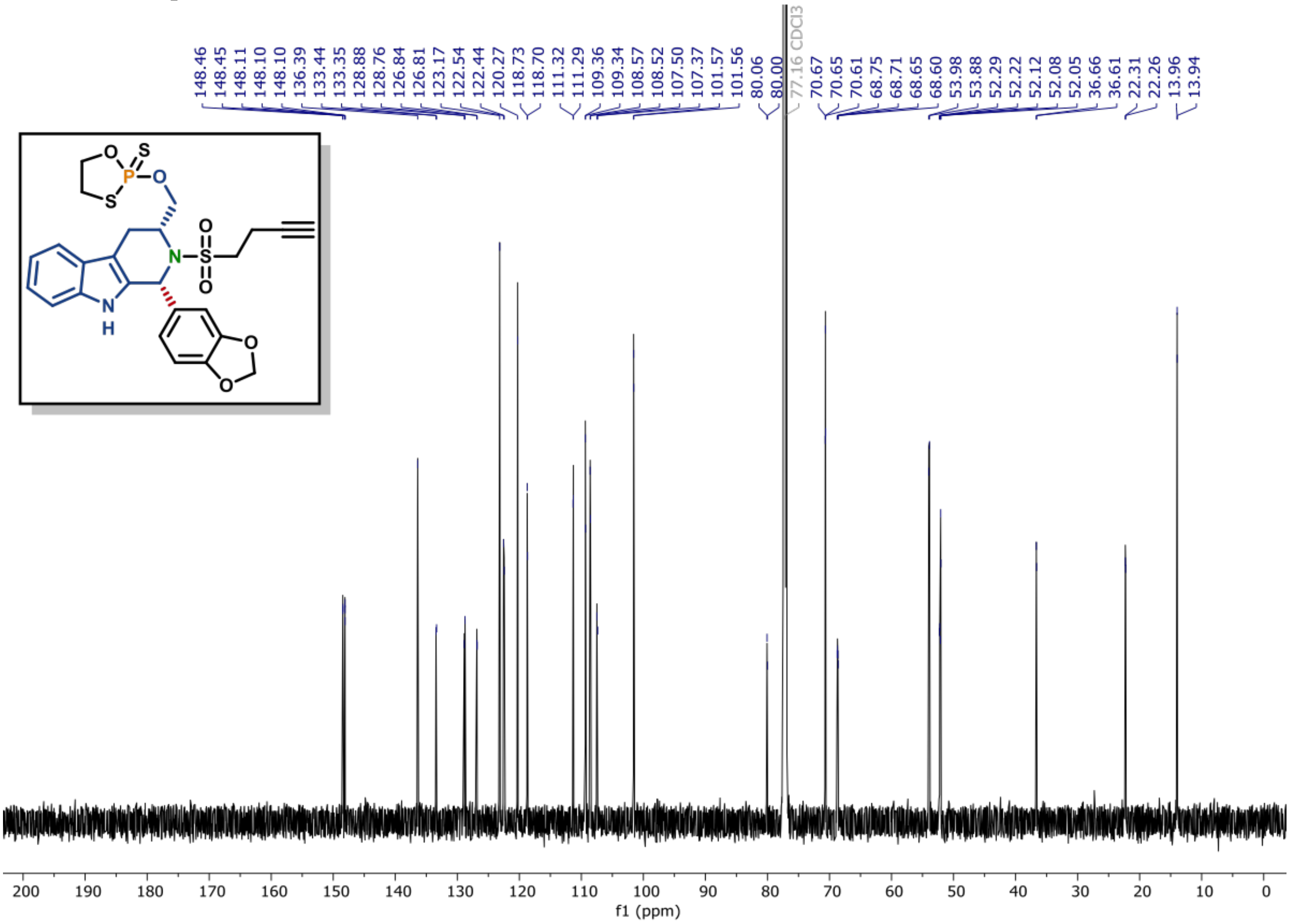

[31]P NMR of compound (20) (162 MHz, $CDCl_3$)

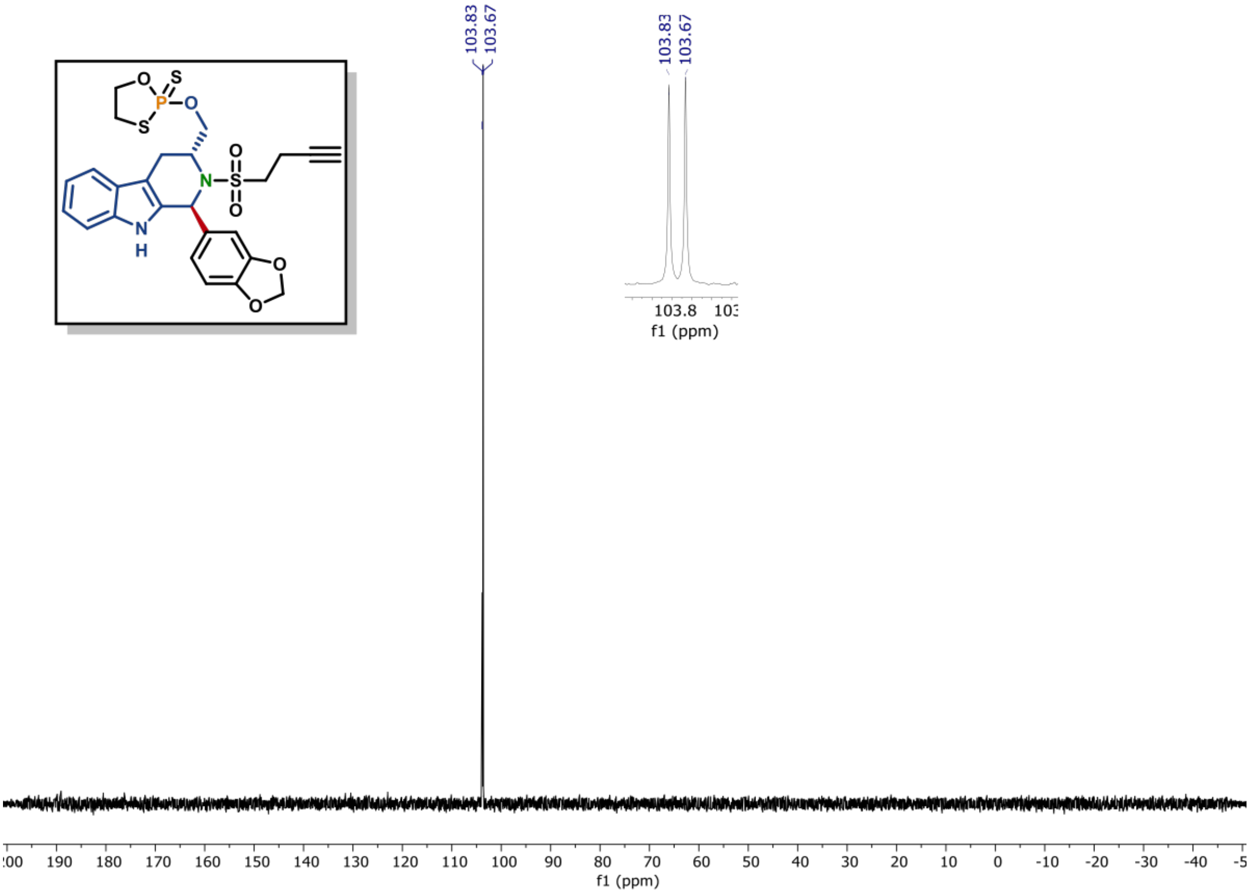

[1]H NMR of compound (20) (600 MHz, $CDCl_3$)

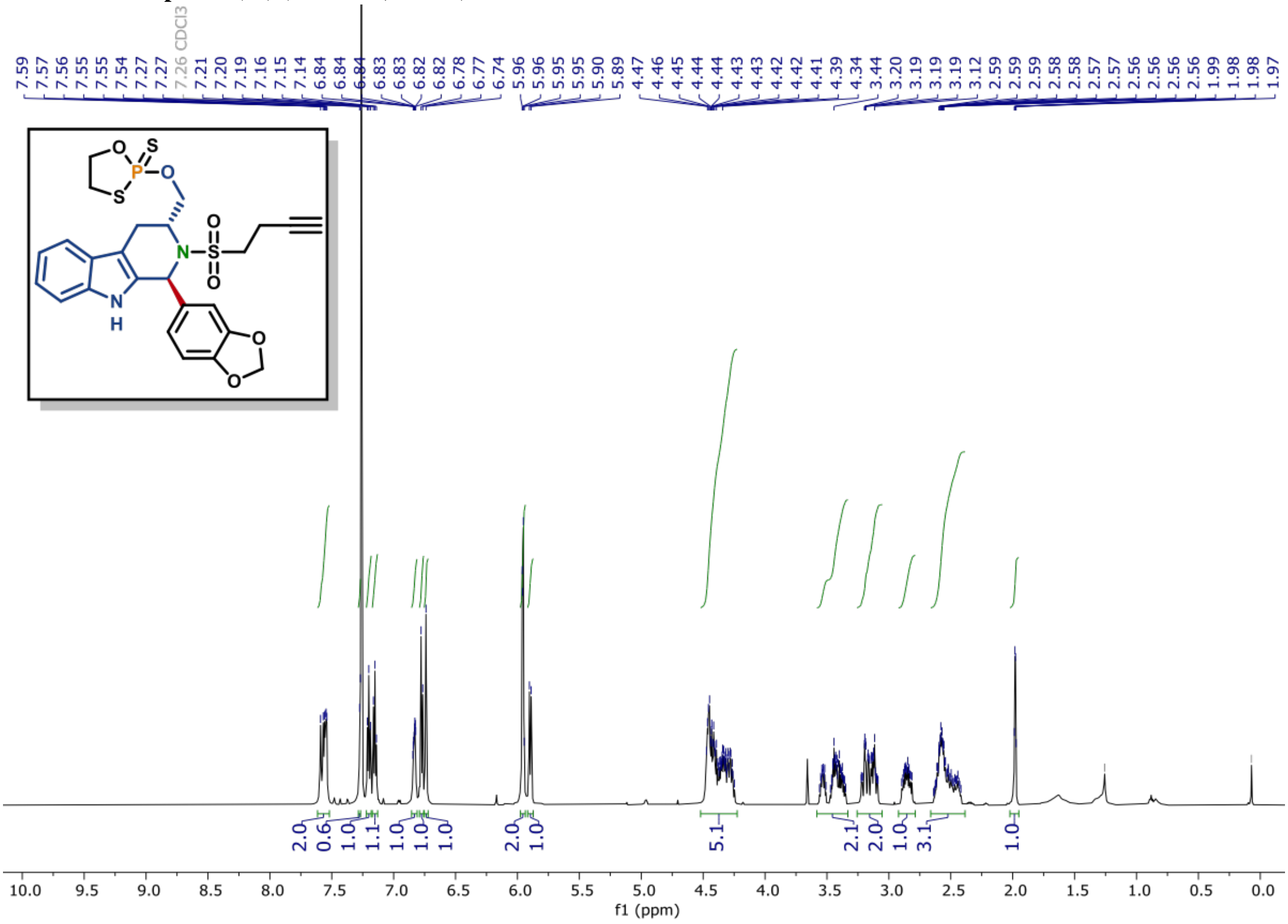

$^{13}C$ NMR of compound (20) (150 MHz, $CDCl_3$)

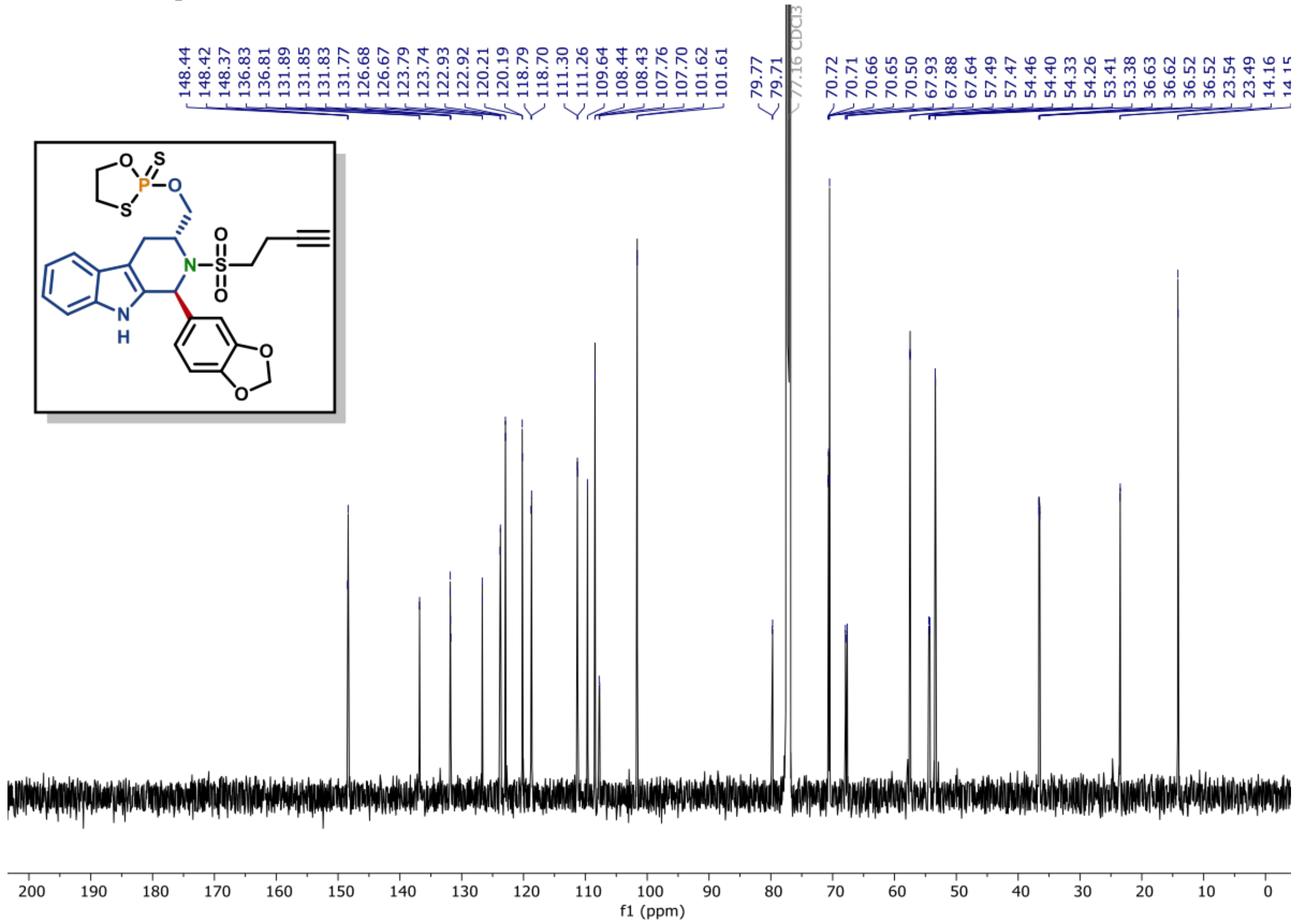

**[31]P NMR of compound (21) (162 MHz, CDCl$_3$)**

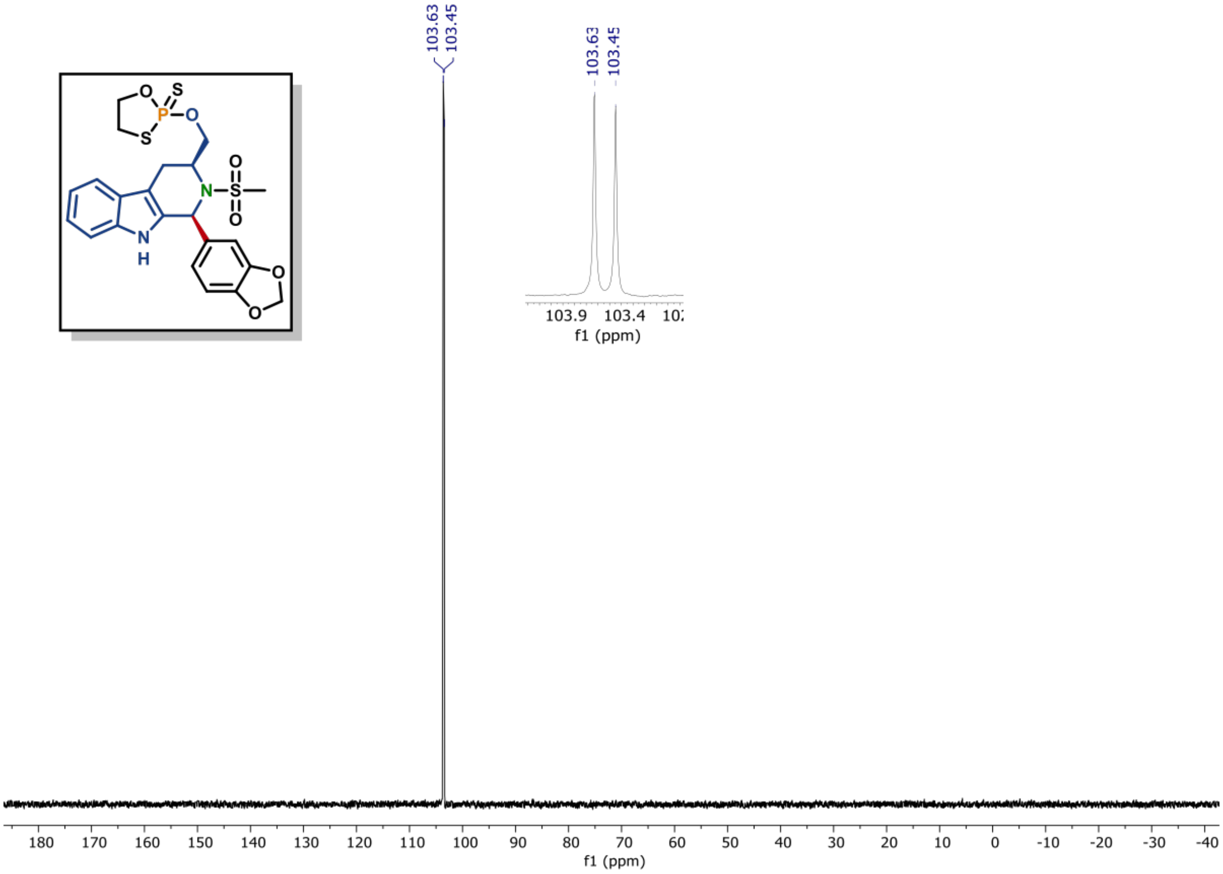

**[1]H NMR of compound (21) (600 MHz, CDCl$_3$)**

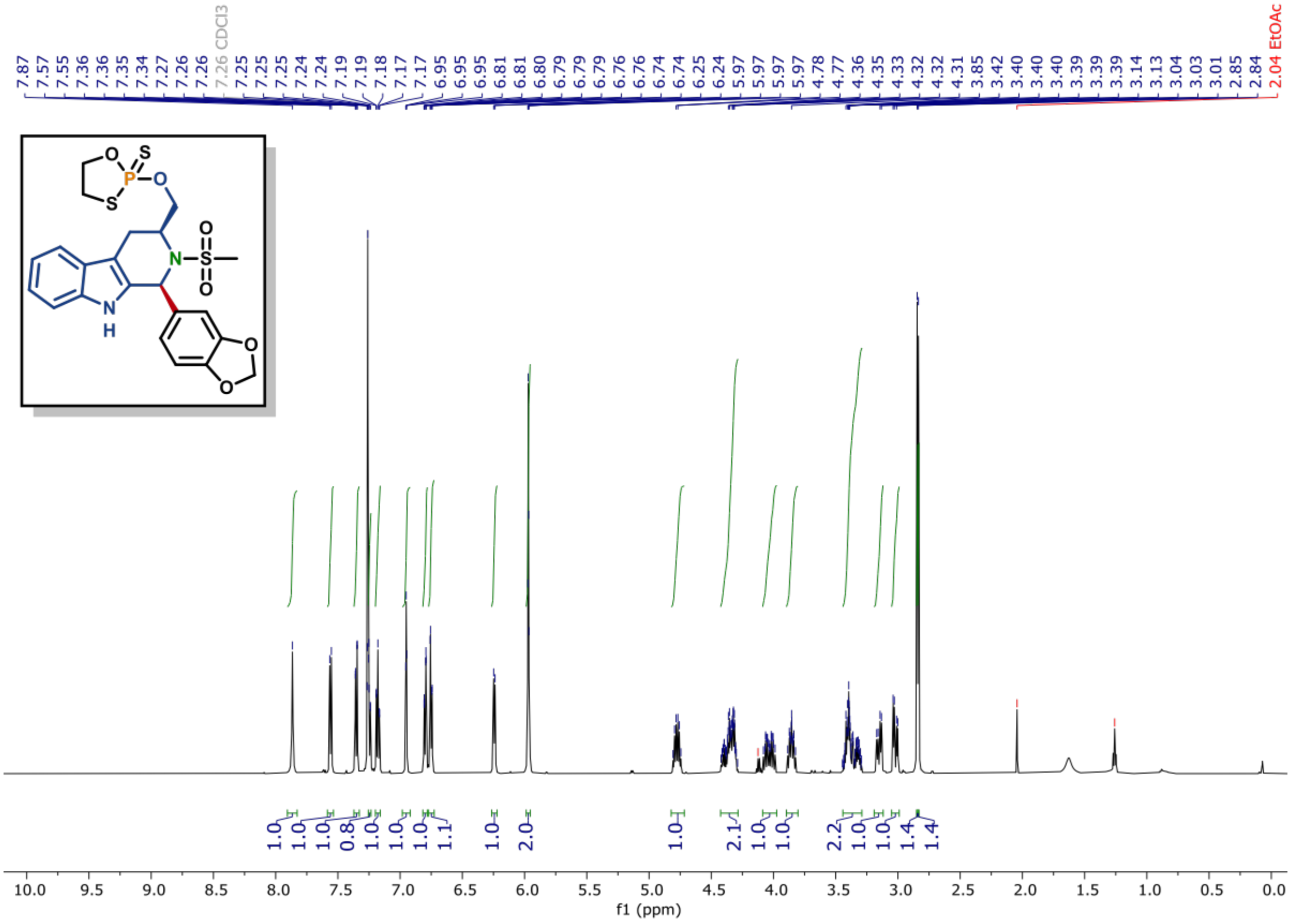

[13]C NMR of compound (21) (150 MHz, $CDCl_3$)

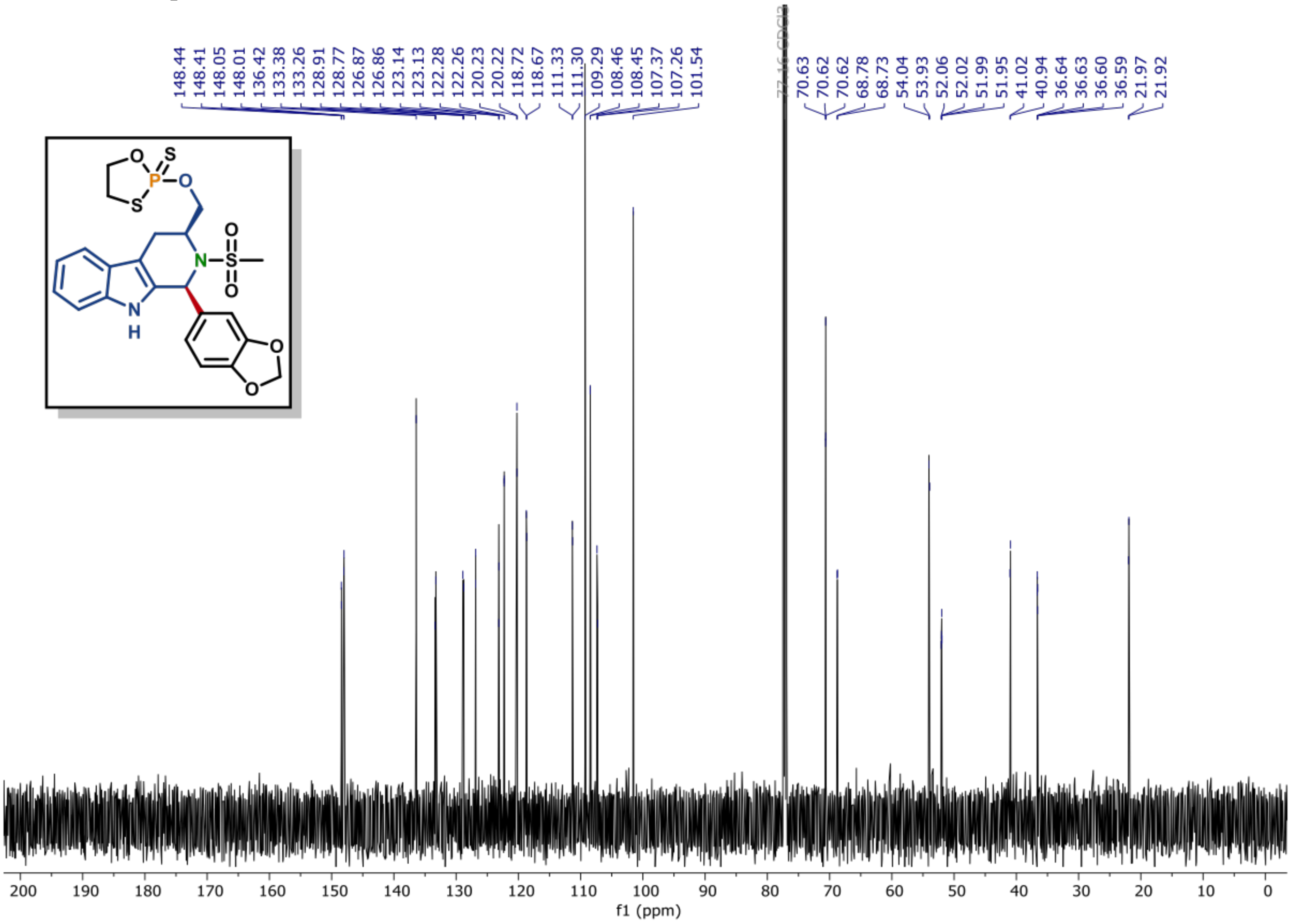

**[31]P NMR of compound (23) (162 MHz, CDCl₃)**

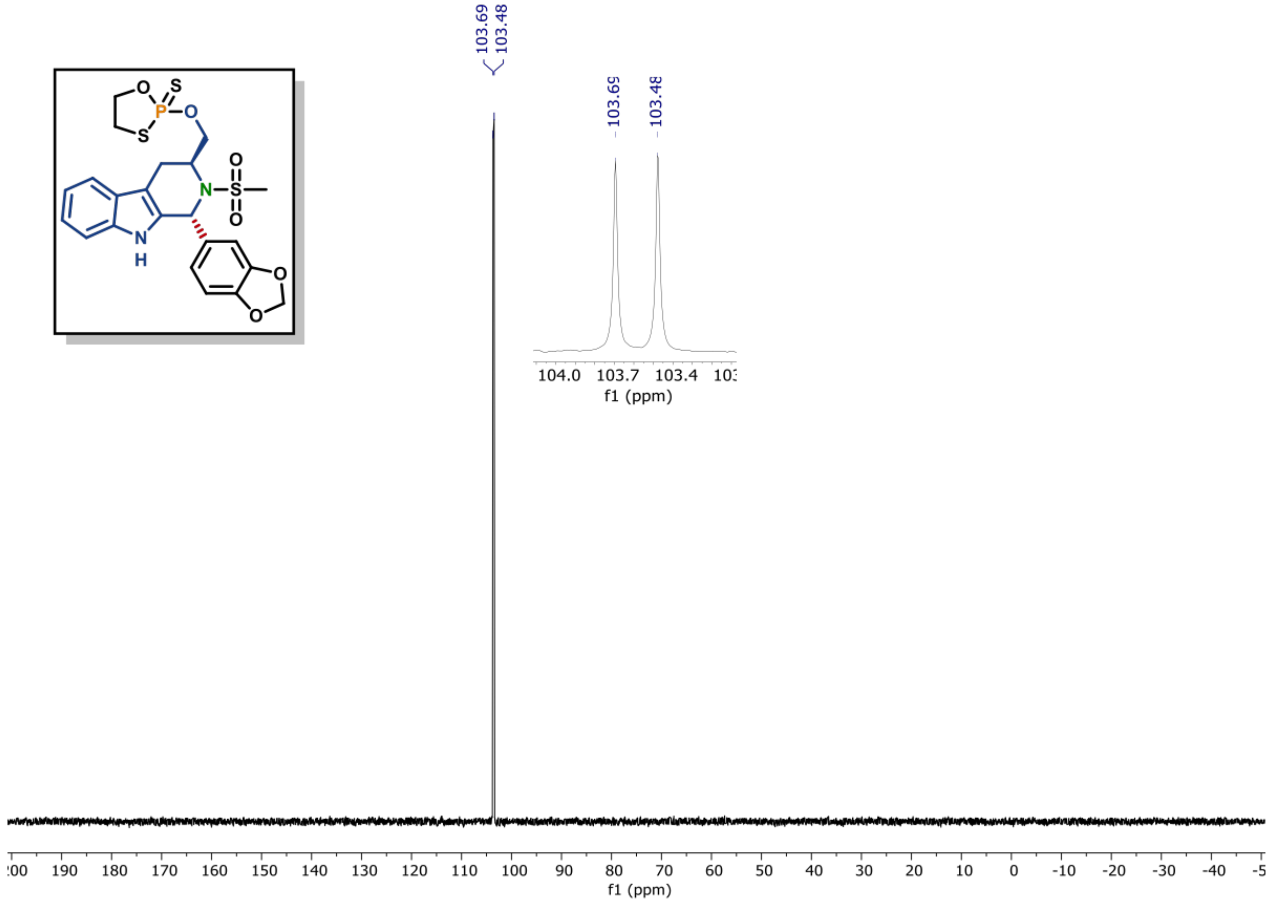

**[1]H NMR of compound (23) (600 MHz, CDCl₃)**

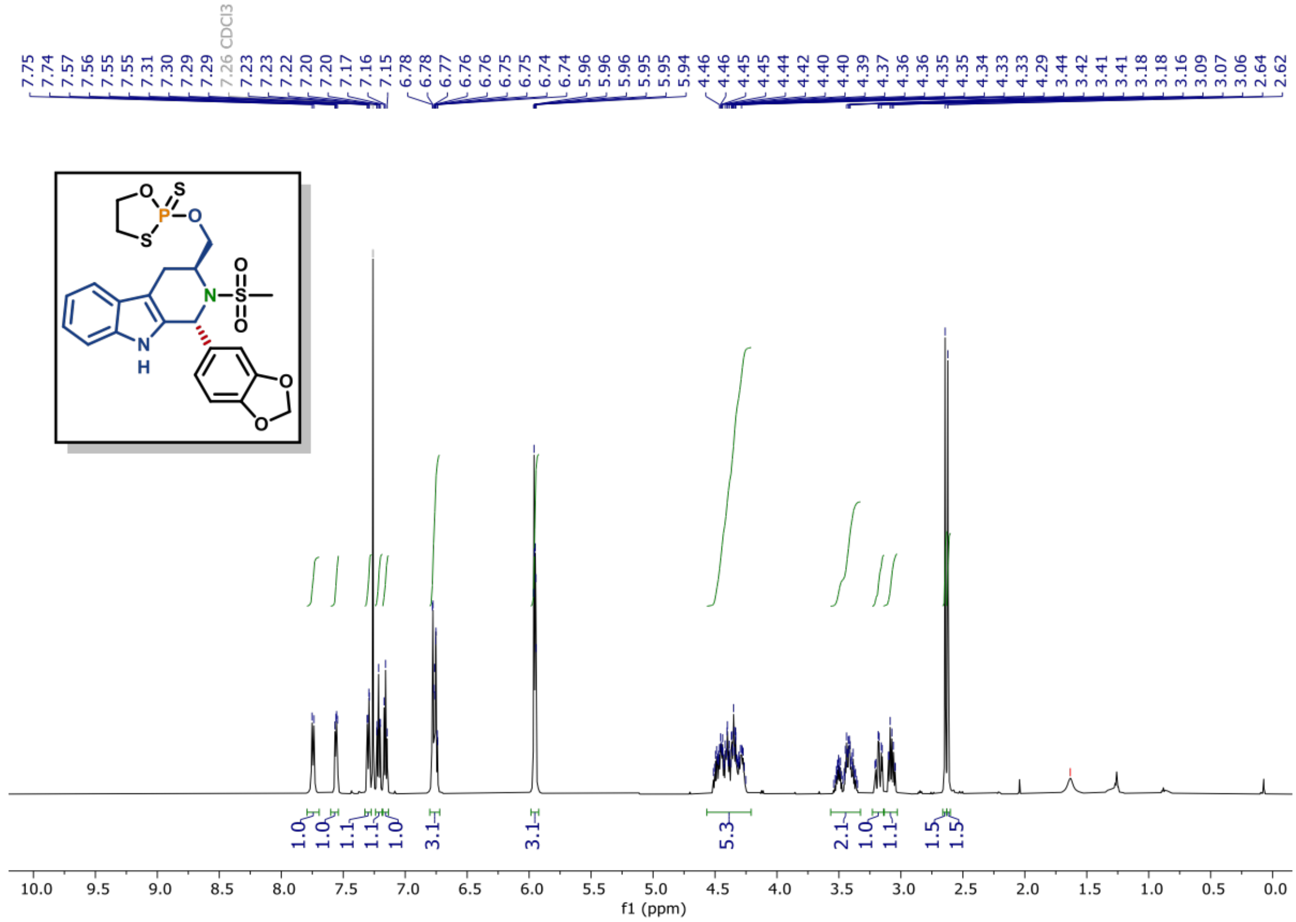

**$^{13}C$ NMR of compound (23) (150 MHz, $CDCl_3$)**

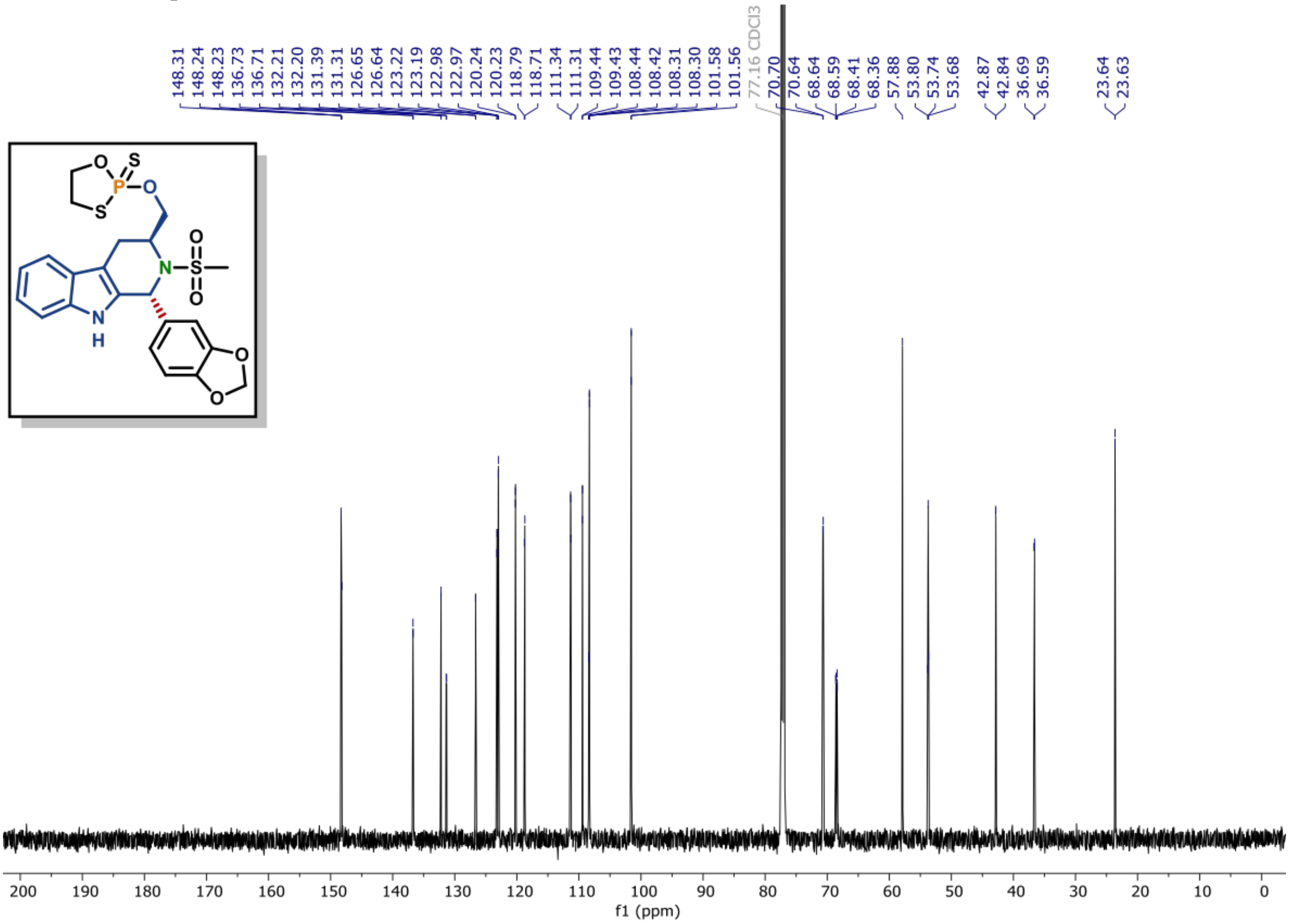

**[31]P NMR of compound (22) (162 MHz, CDCl$_3$)**

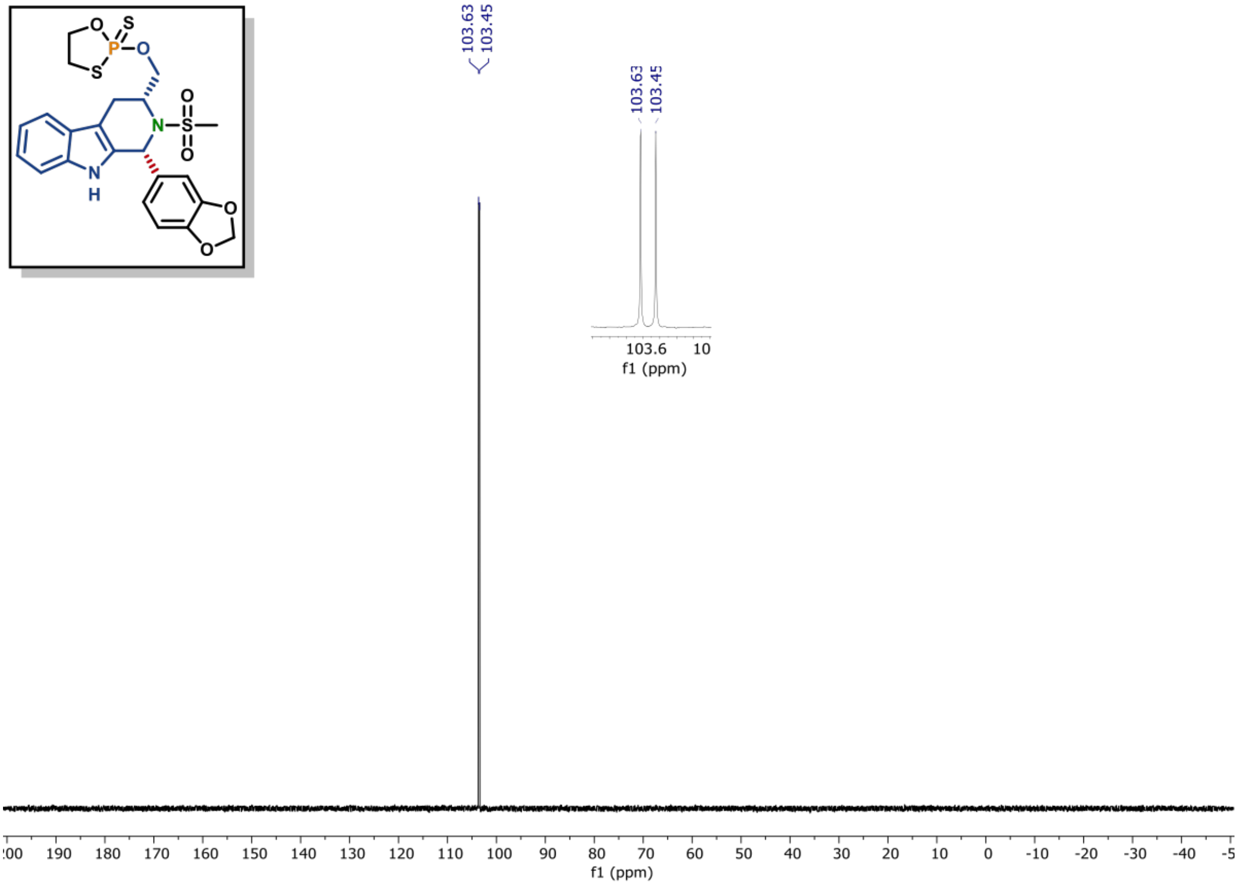

**[1]H NMR of compound (22) (600 MHz, CDCl$_3$)**

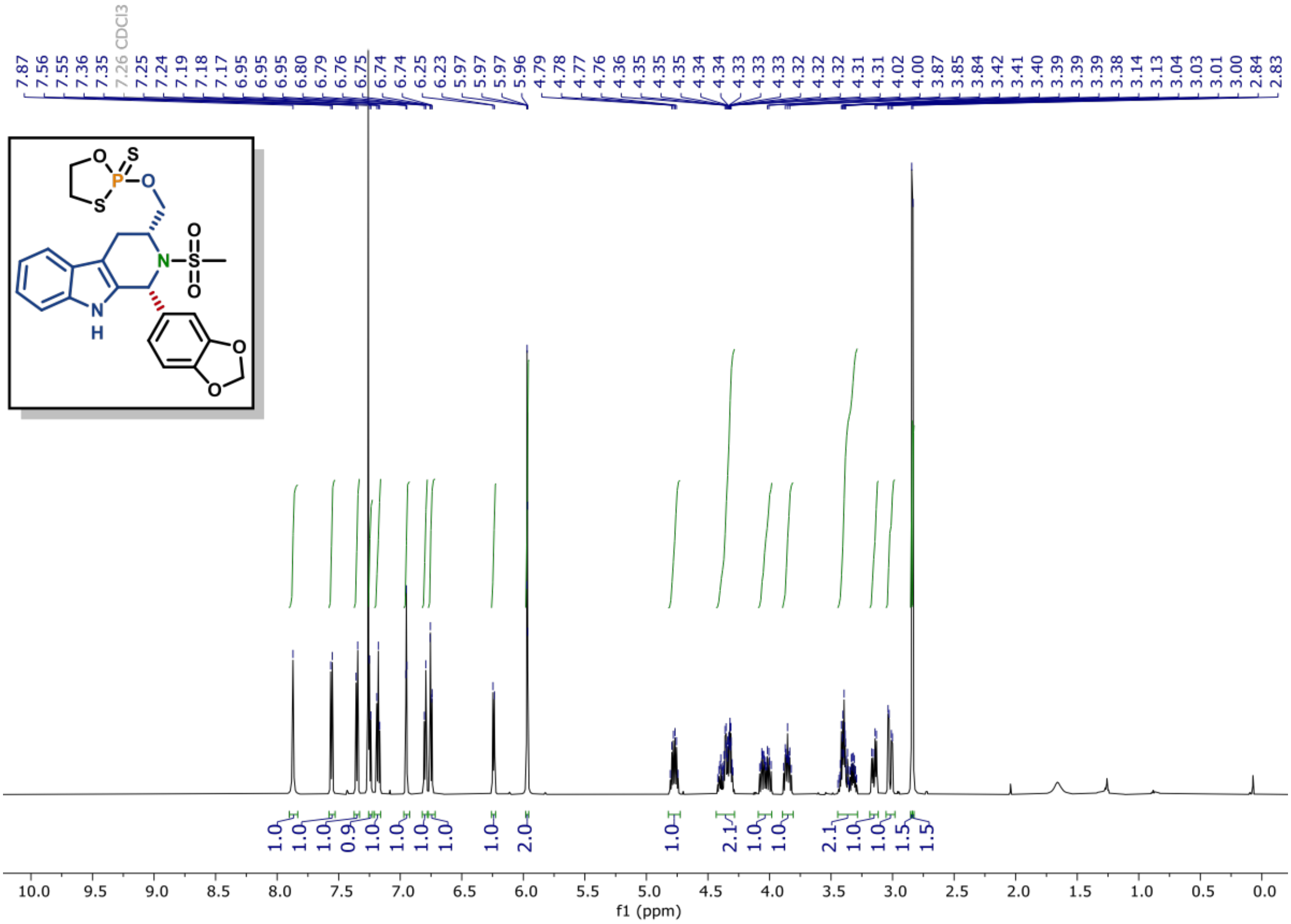

**[13]C NMR of compound (22) (150 MHz, CDCl₃)**

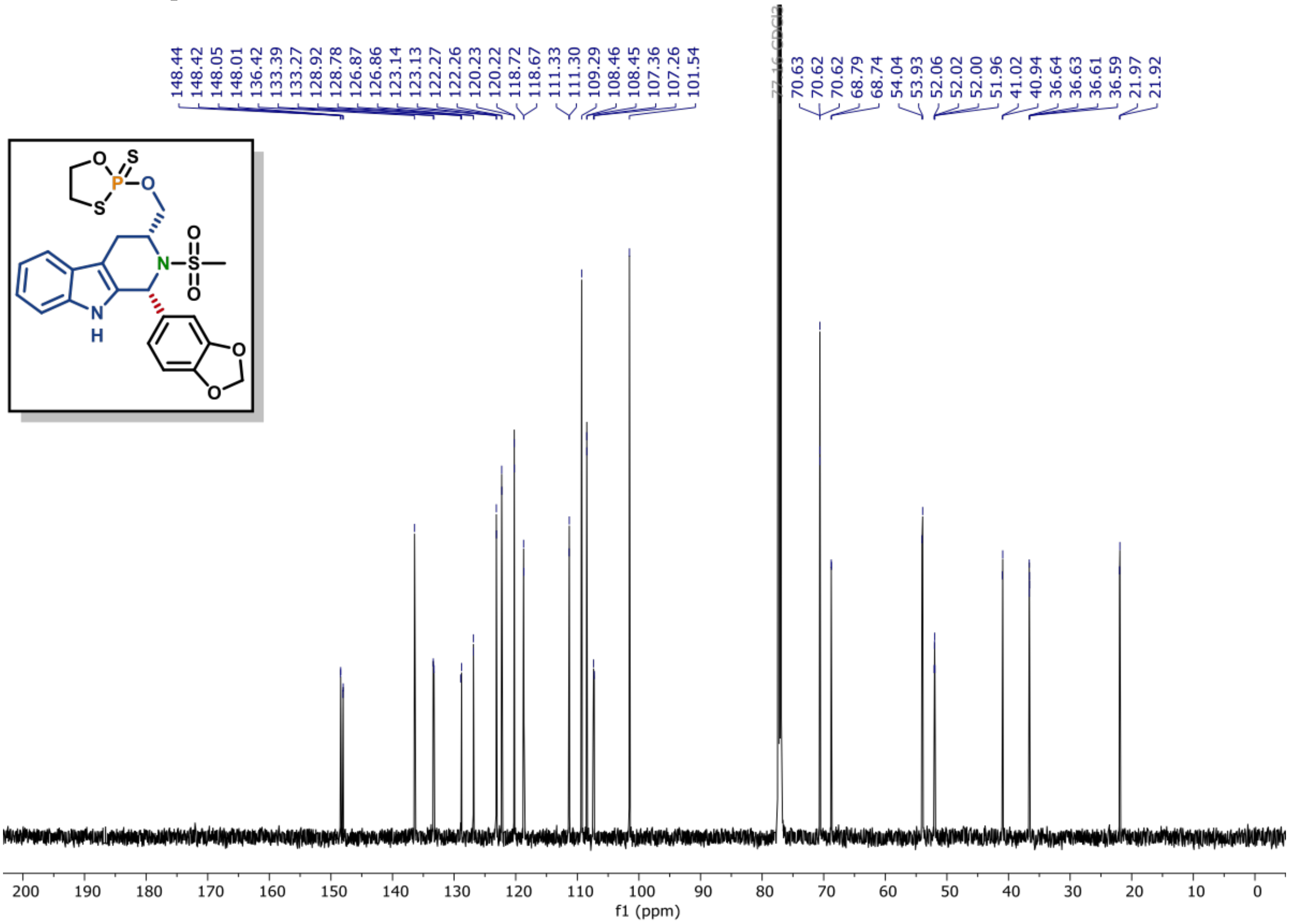

**[1]H NMR of compound (24) (600 MHz, $CDCl_3$)**

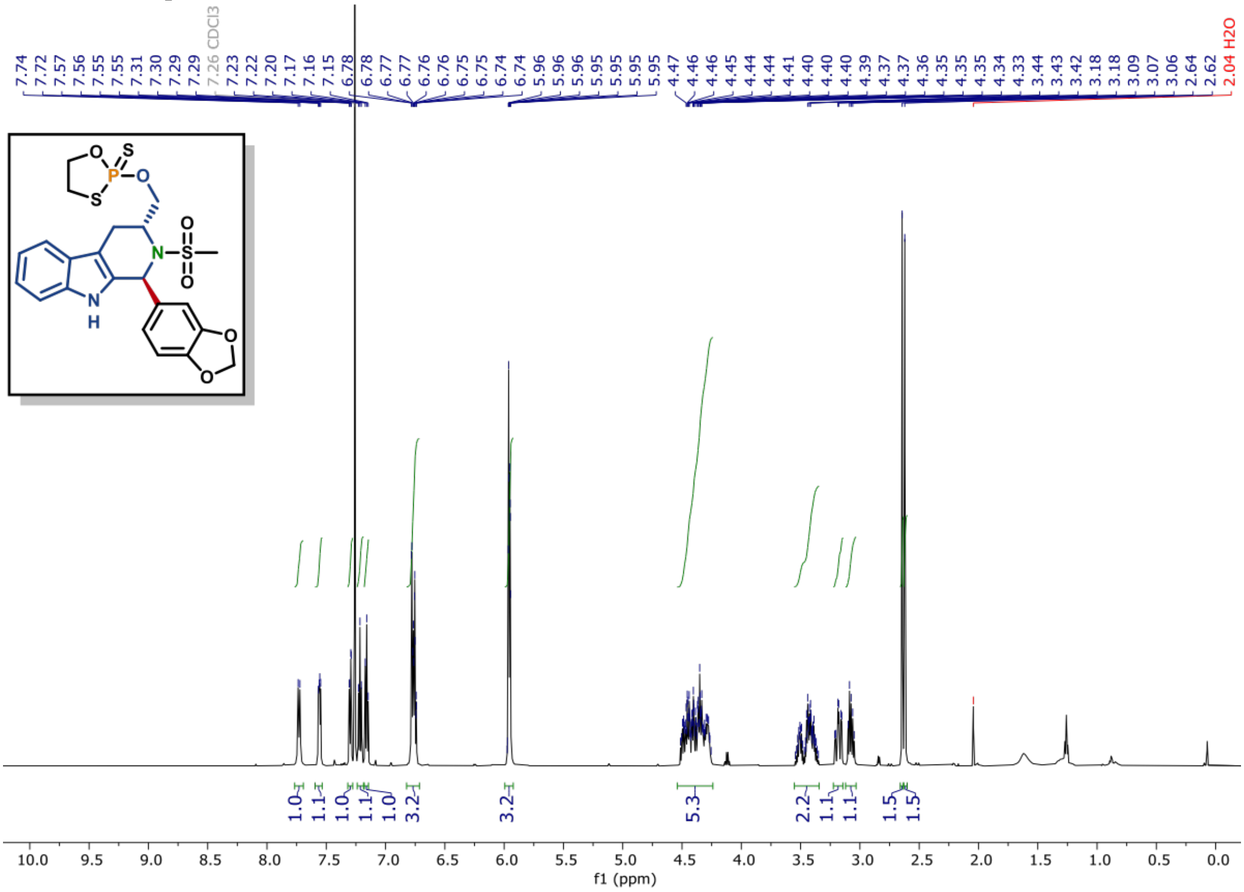

**[13]C NMR of compound (24) (150 MHz, $CDCl_3$)**

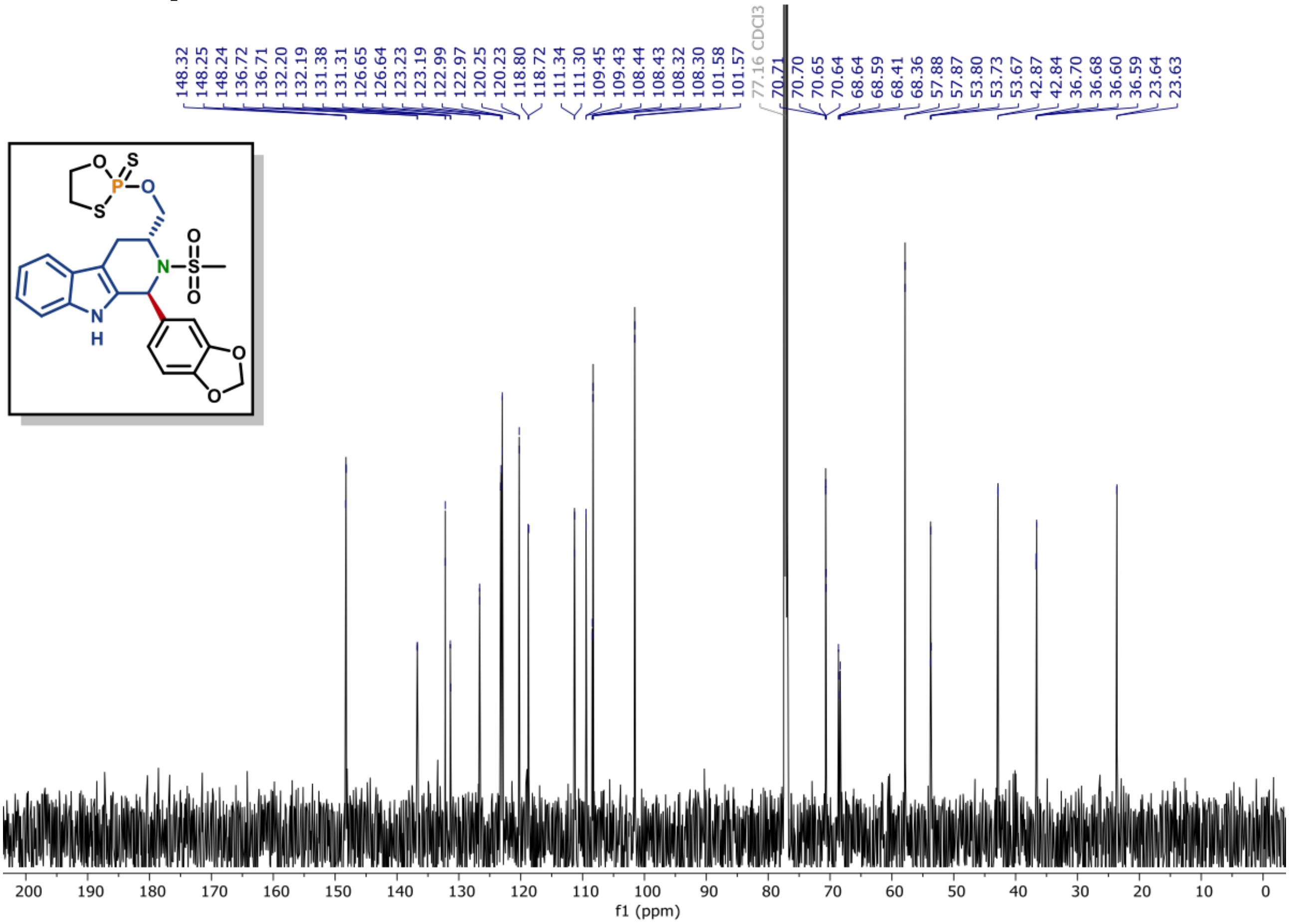

**[1]H NMR of compound (S5A) (600 MHz, CDCl3)**

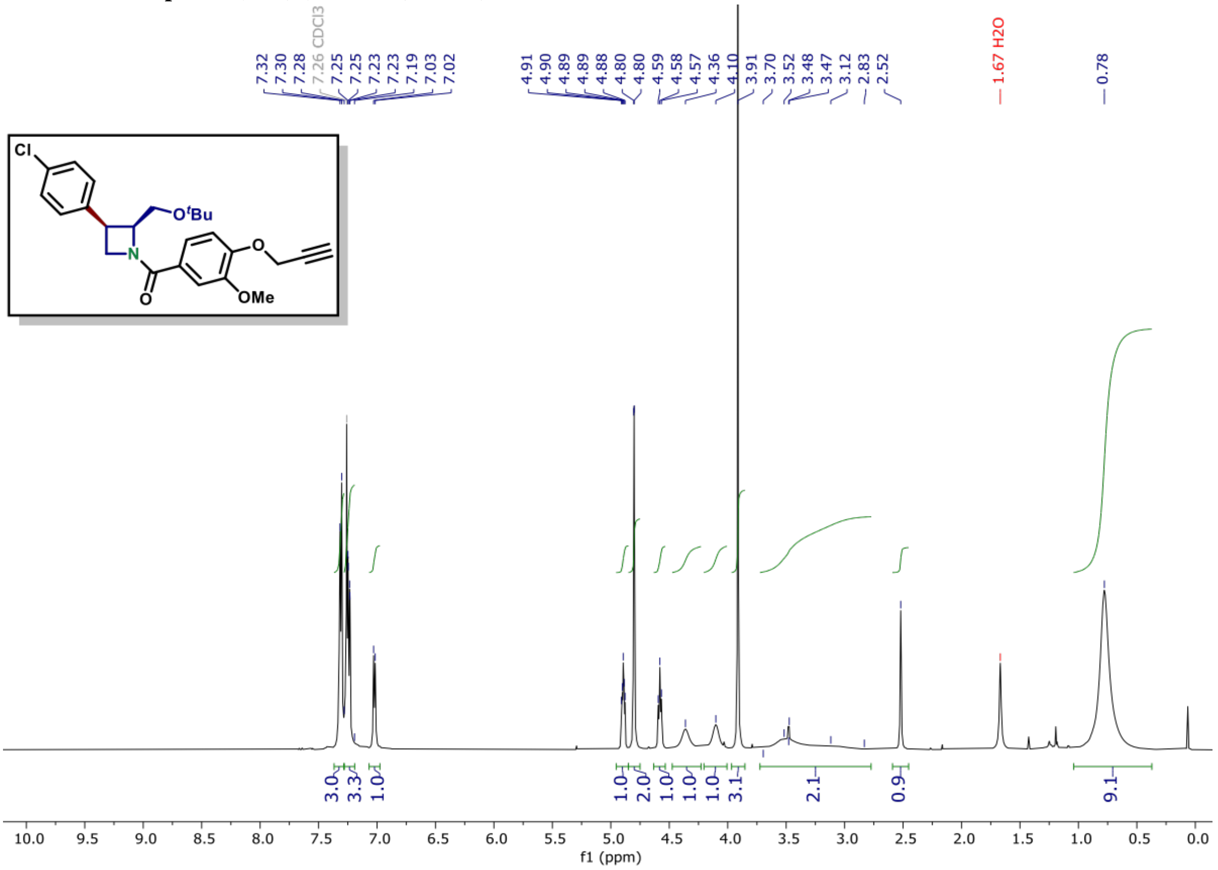

**[13]C NMR of compound (S5A) (150 MHz, CDCl3)**

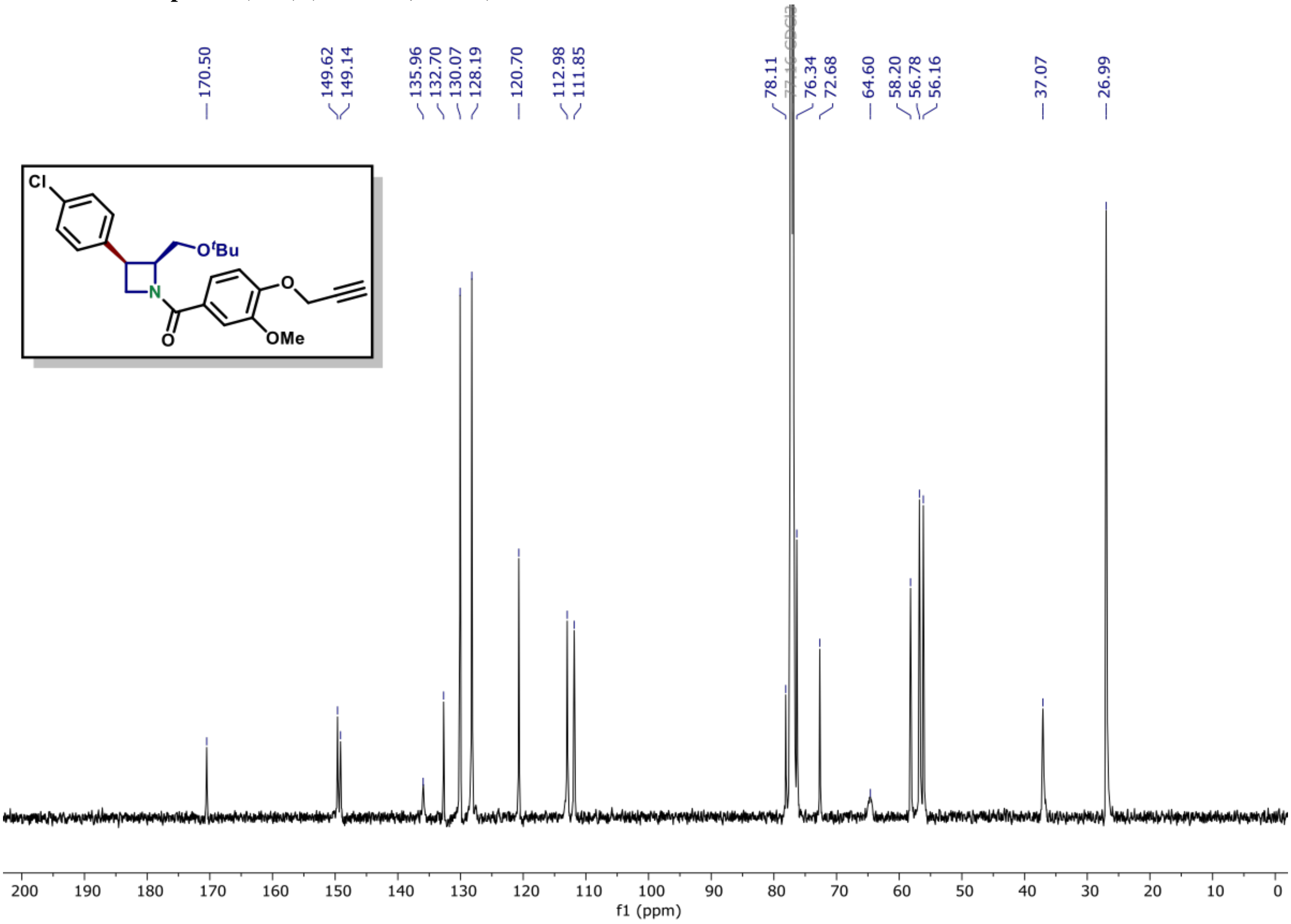

**[1]H NMR of compound (S5B) (600 MHz, CDCl₃)**

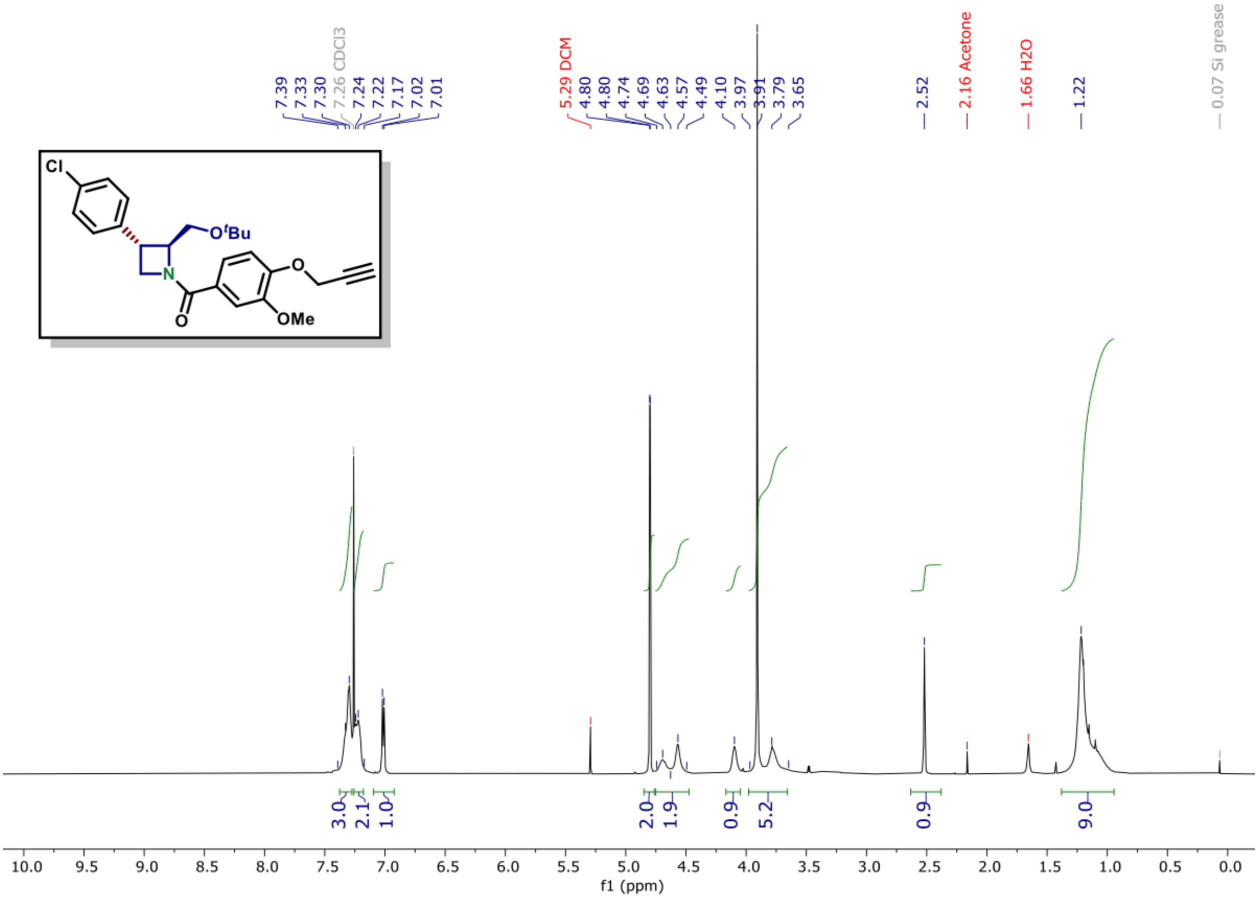

**[13]C NMR of compound (S5B) (150 MHz, CDCl₃)**

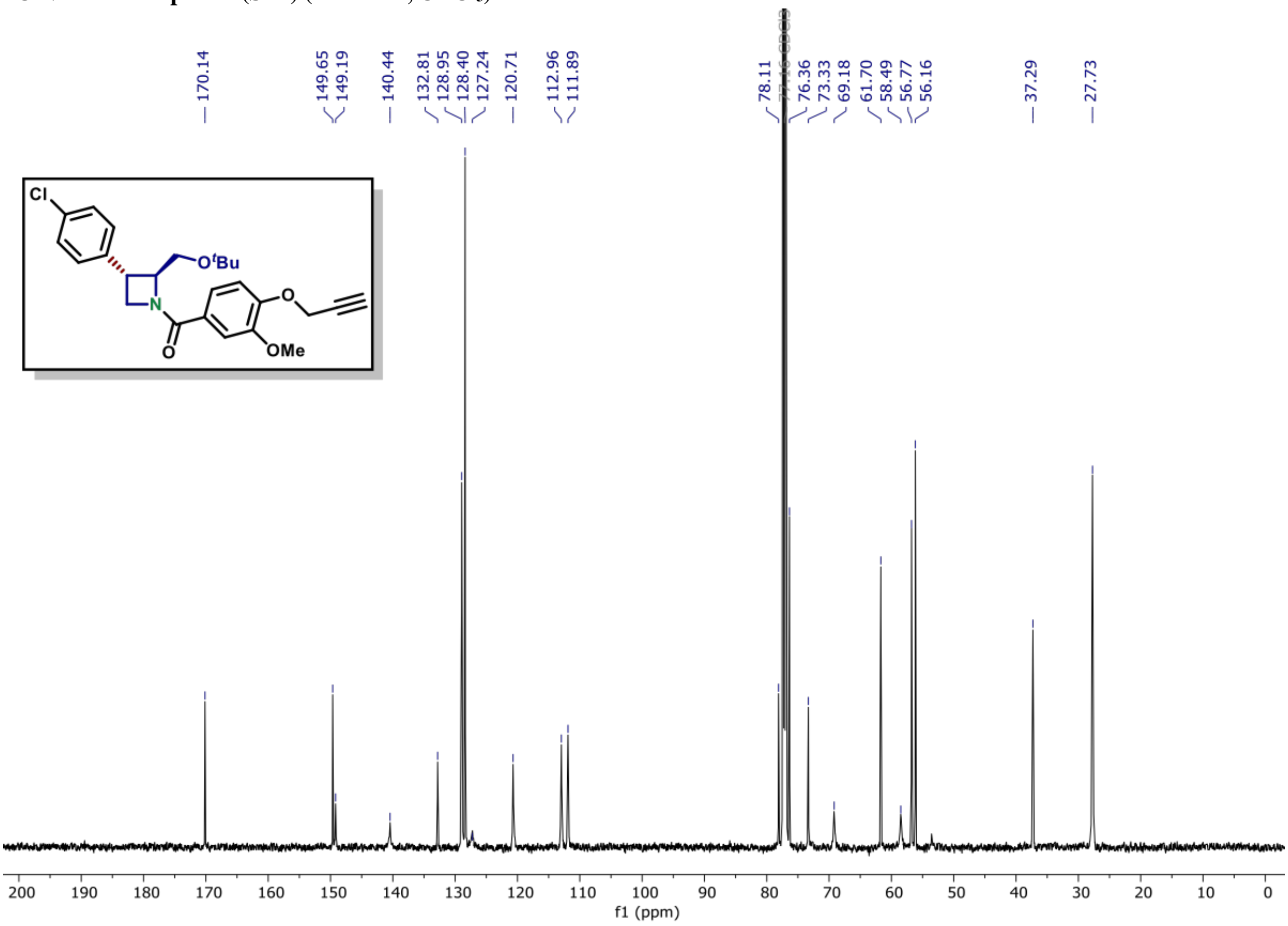

[1]H NMR of compound (S5C) (600 MHz, $CDCl_3$)

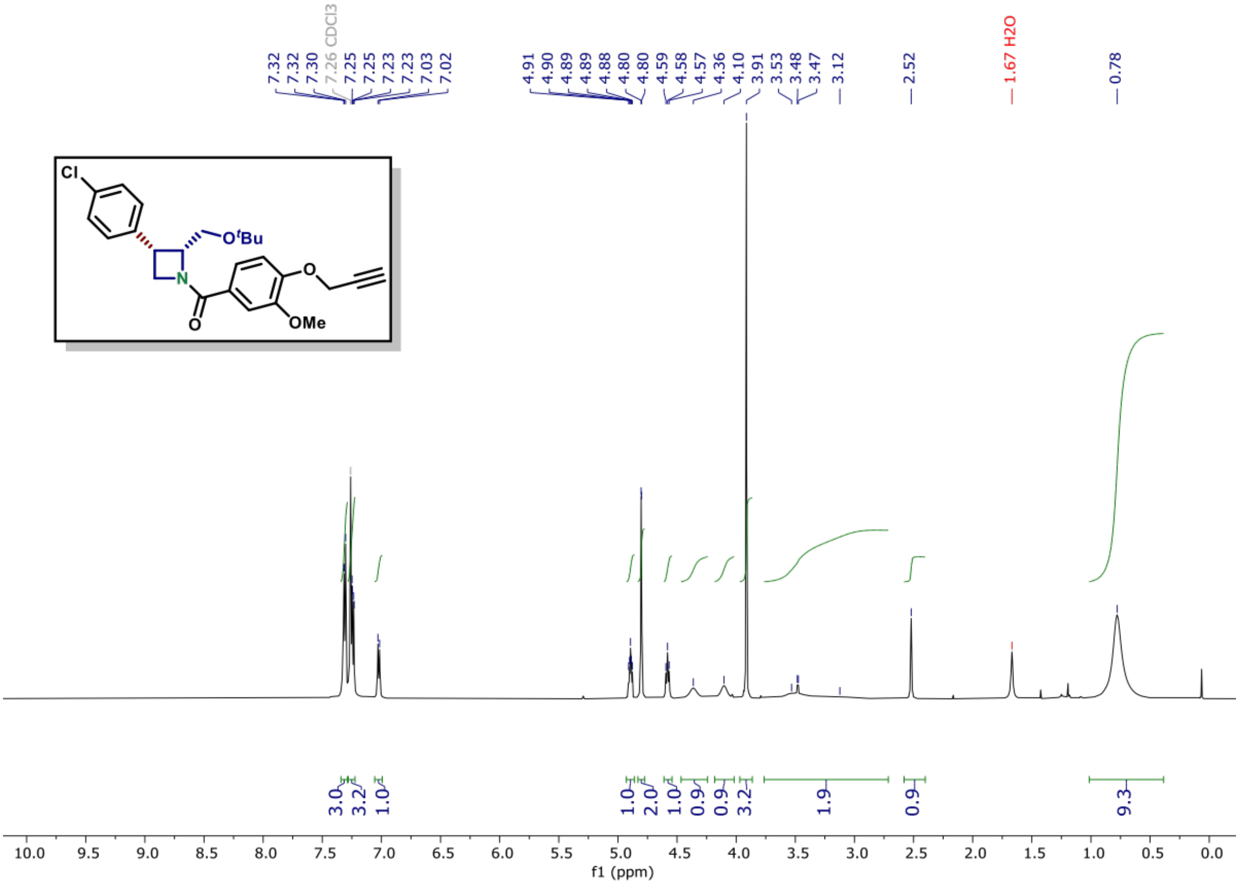

[13]C NMR of compound (S5C) (150 MHz, $CDCl_3$)

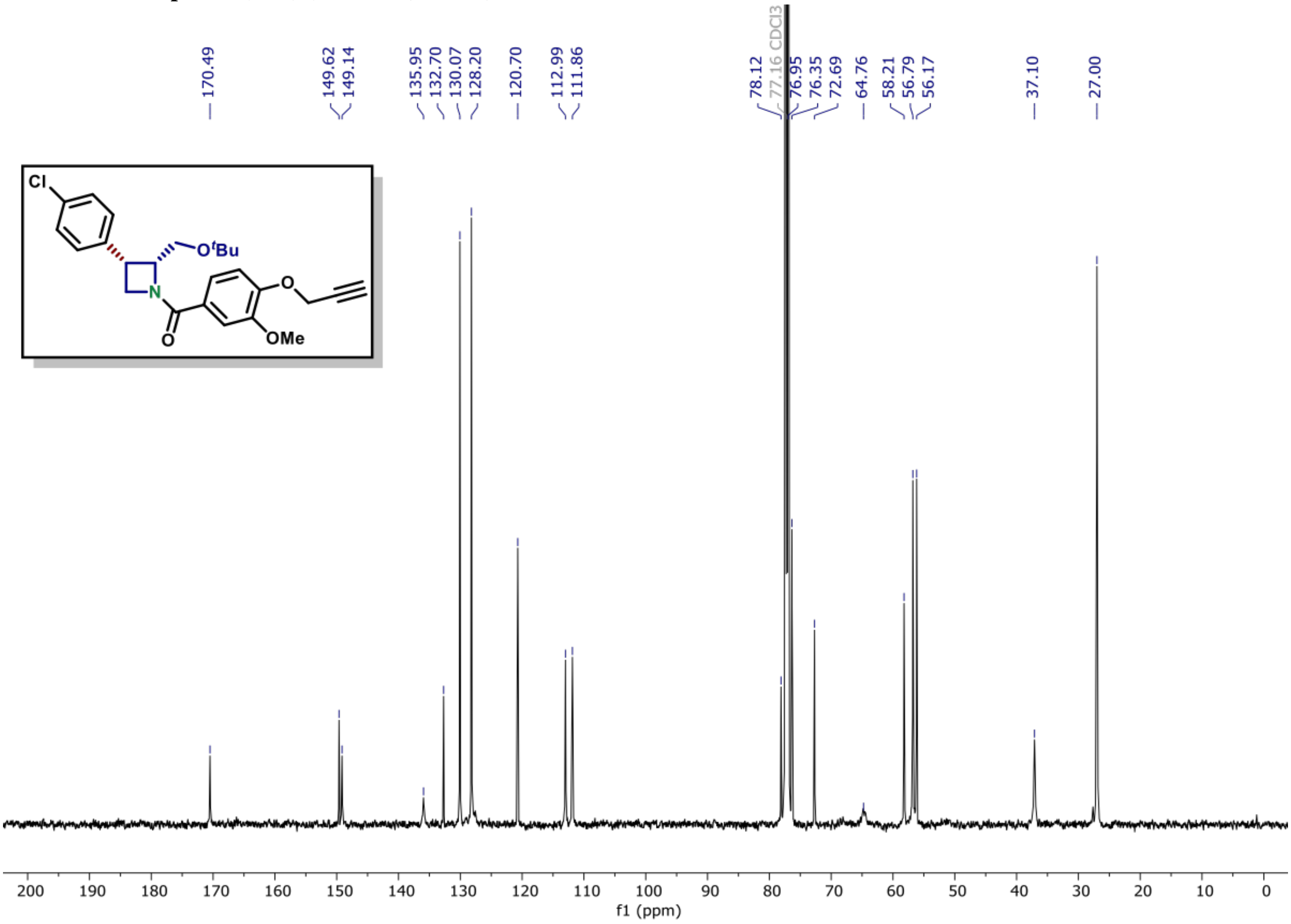

$^{1}$H NMR of compound (S5D) (600 MHz, $CDCl_3$)

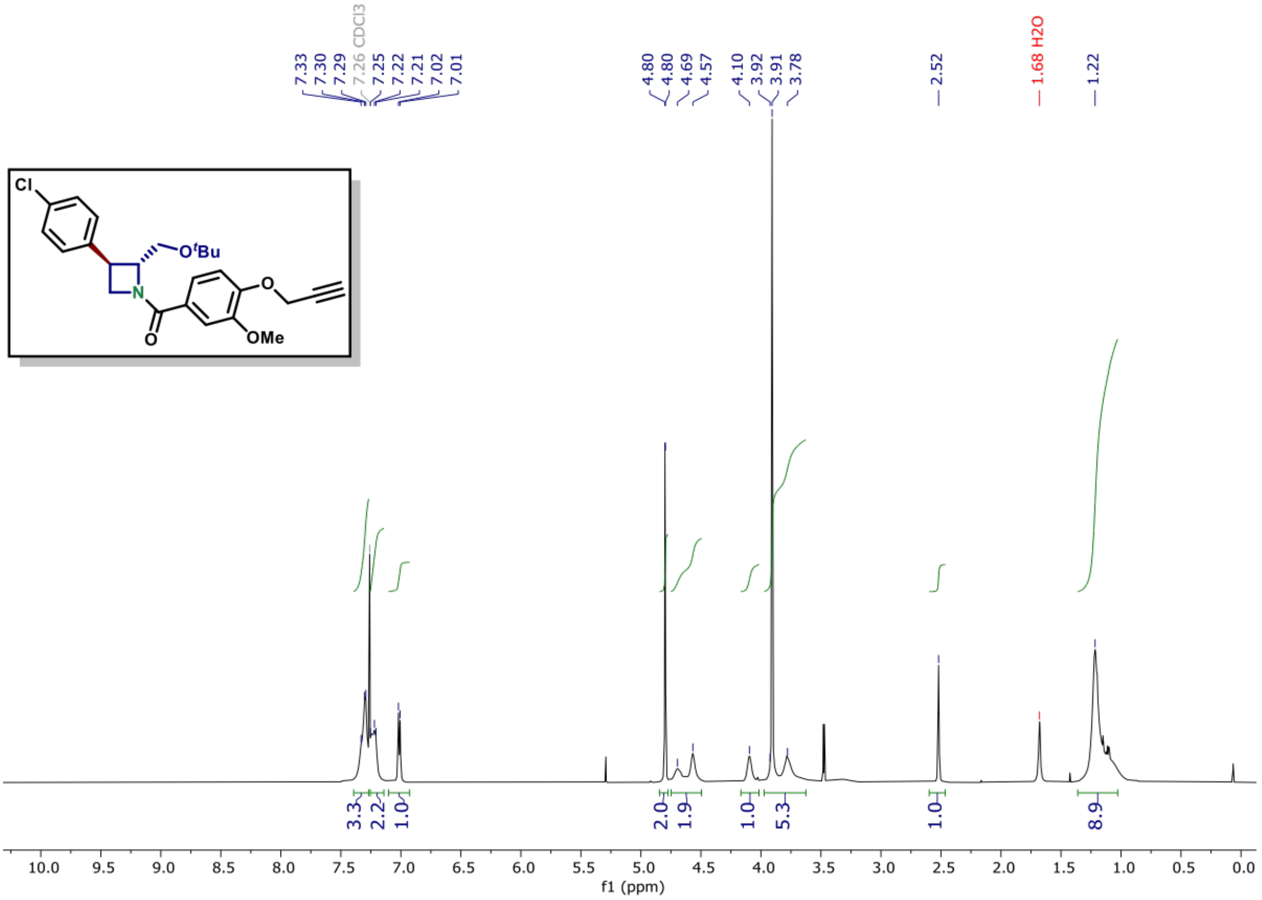

$^{13}$C NMR of compound (S5D) (150 MHz, $CDCl_3$)

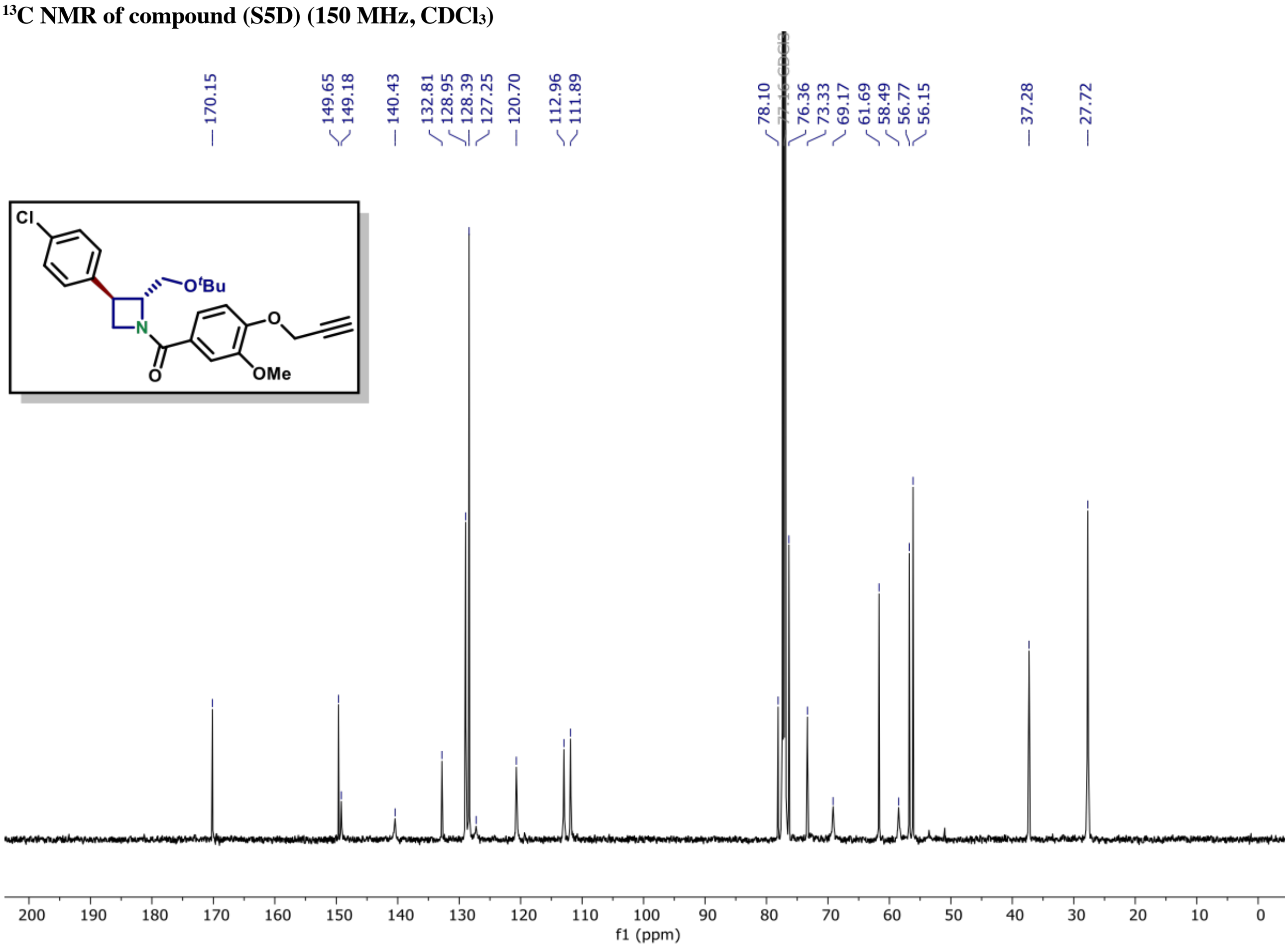

[1]H NMR of compound (S6A) (600 MHz, $CDCl_3$)

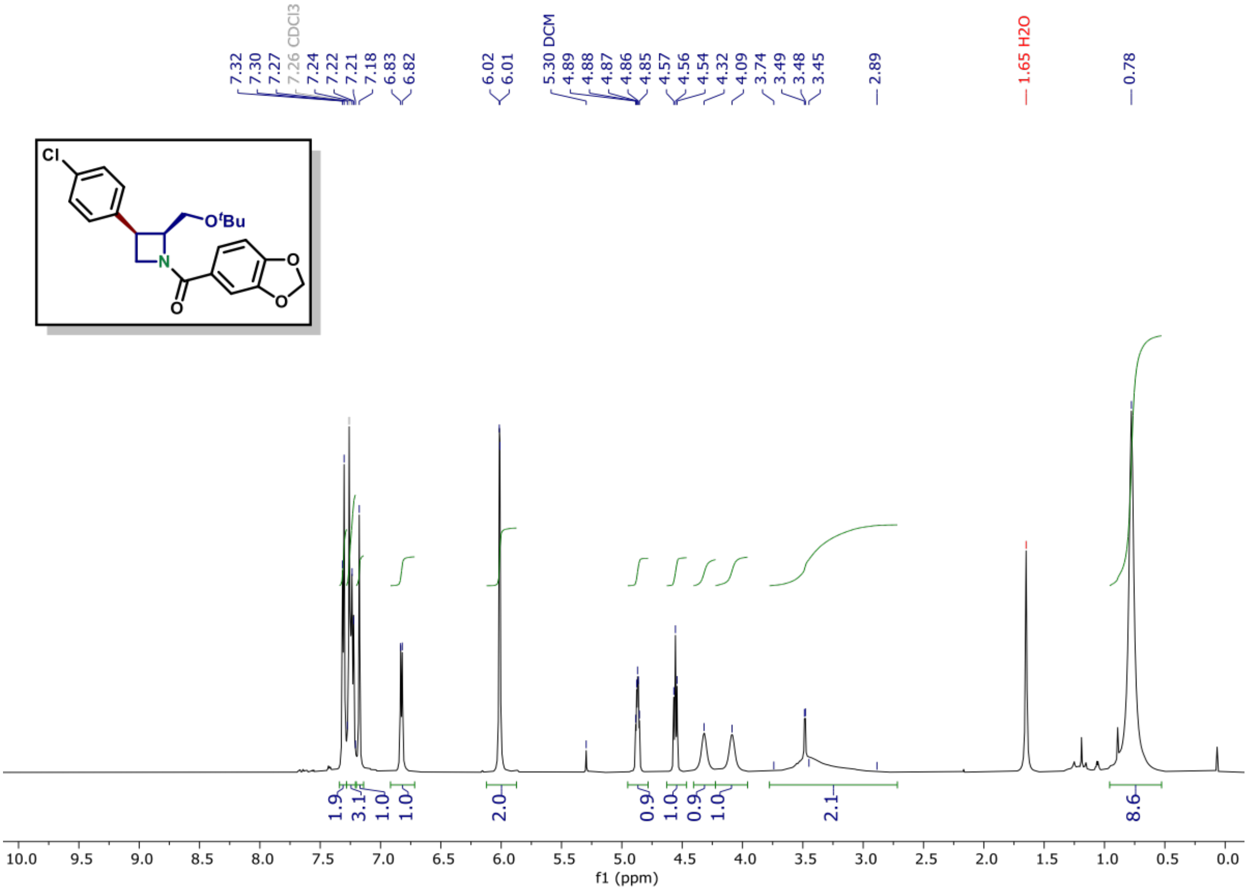

[13]C NMR of compound (S6A) (150 MHz, $CDCl_3$)

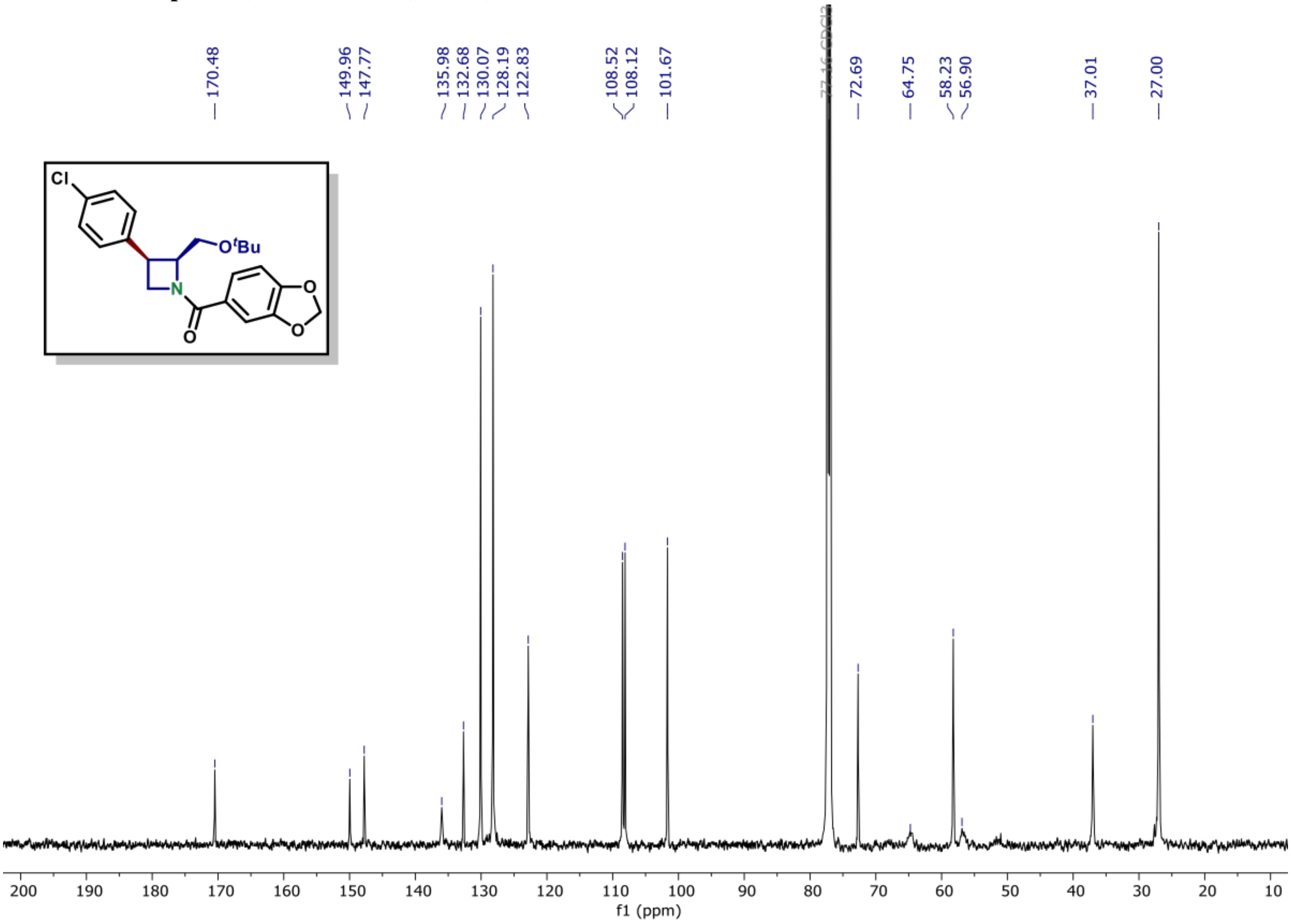

[1]H NMR of compound (S6B) (600 MHz, $CDCl_3$)

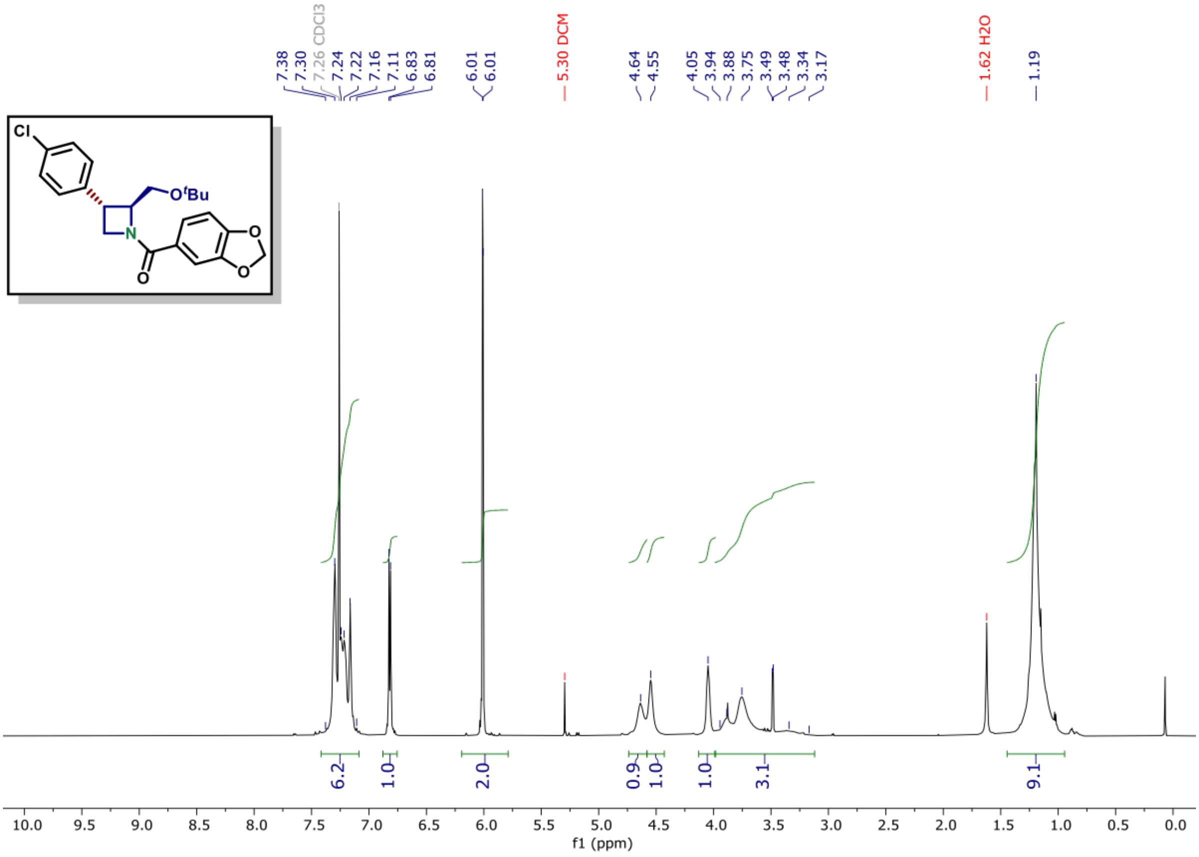

[13]C NMR of compound (S6B) (150 MHz, $CDCl_3$)

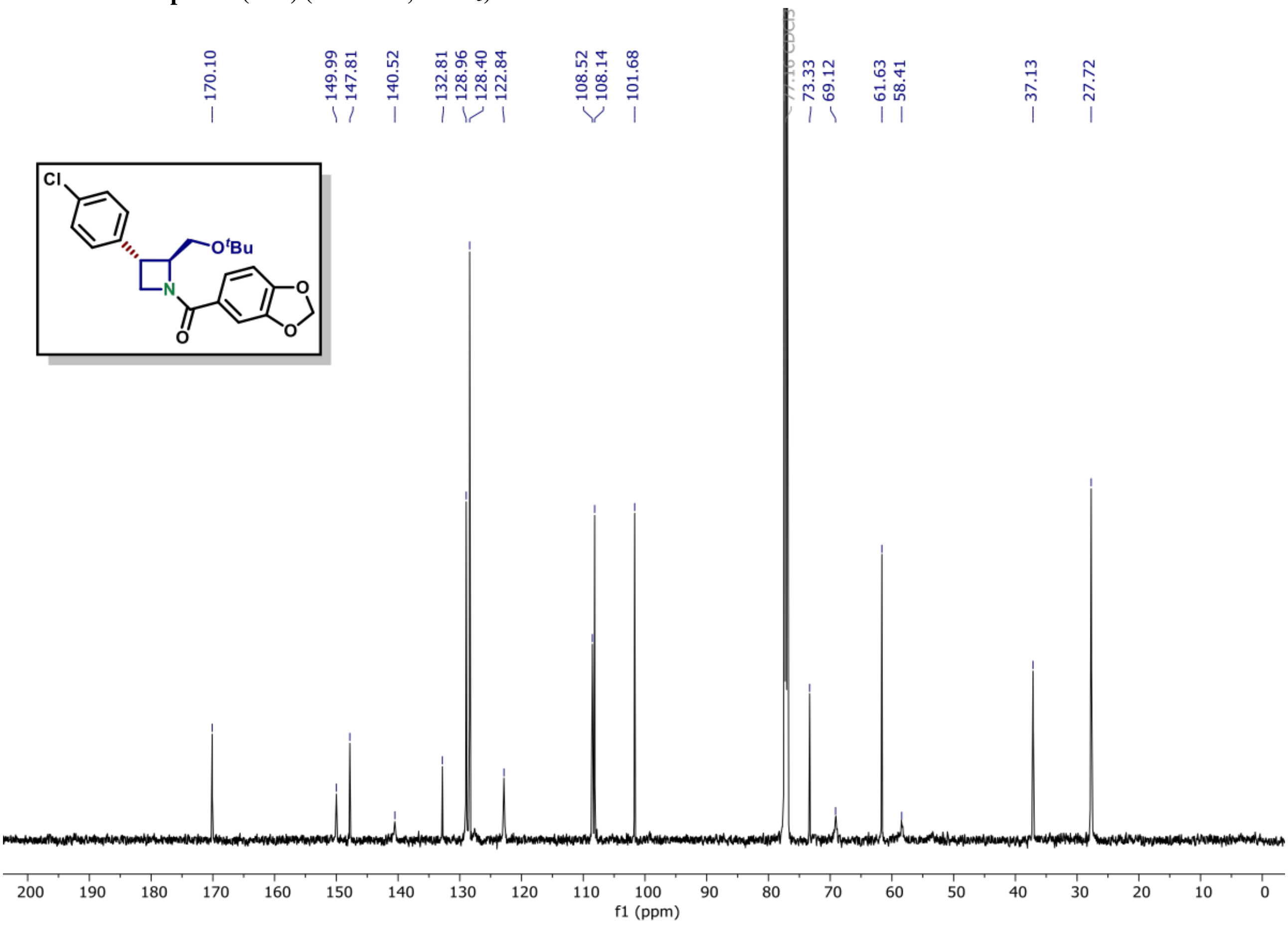

**[1]H NMR of compound (S6C) (600 MHz, $CDCl_3$)**

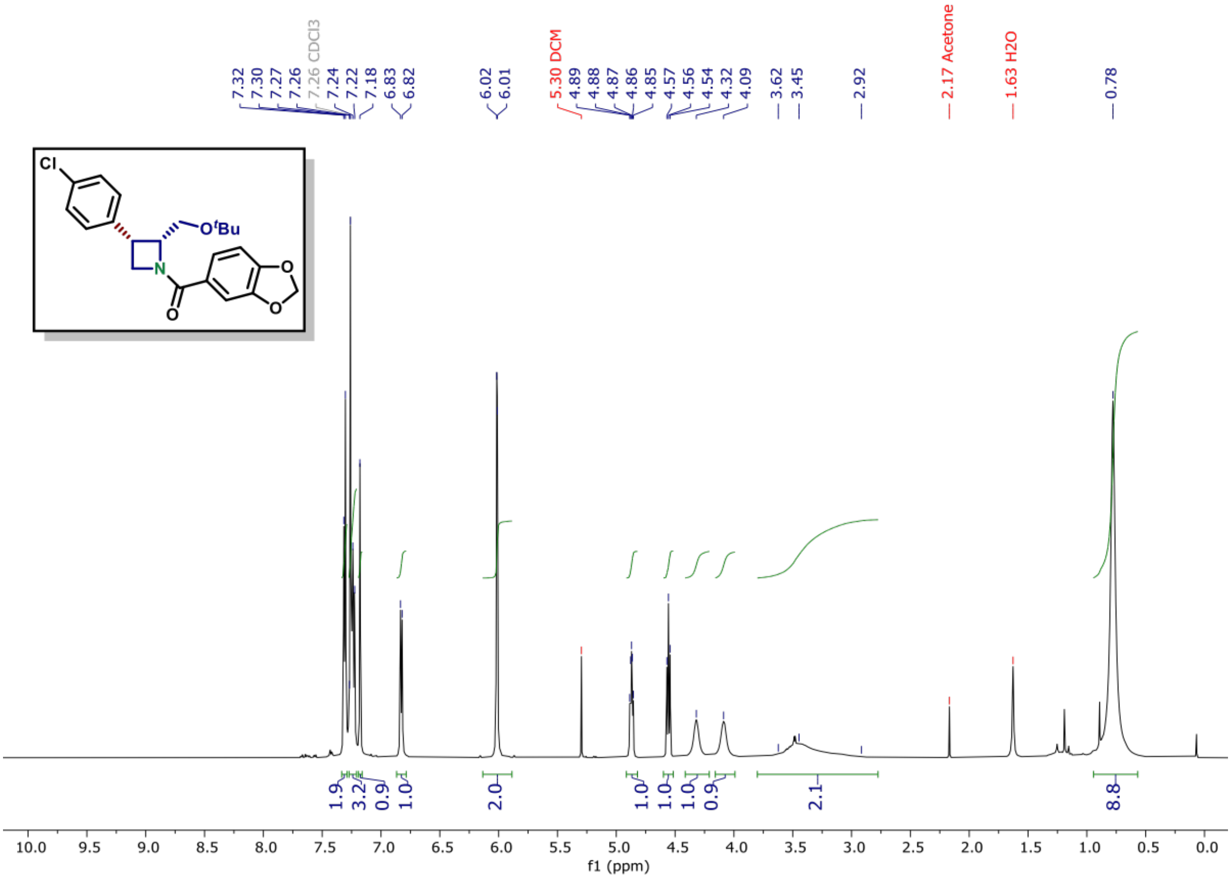

**[13]C NMR of compound (S6C) (150 MHz, $CDCl_3$)**

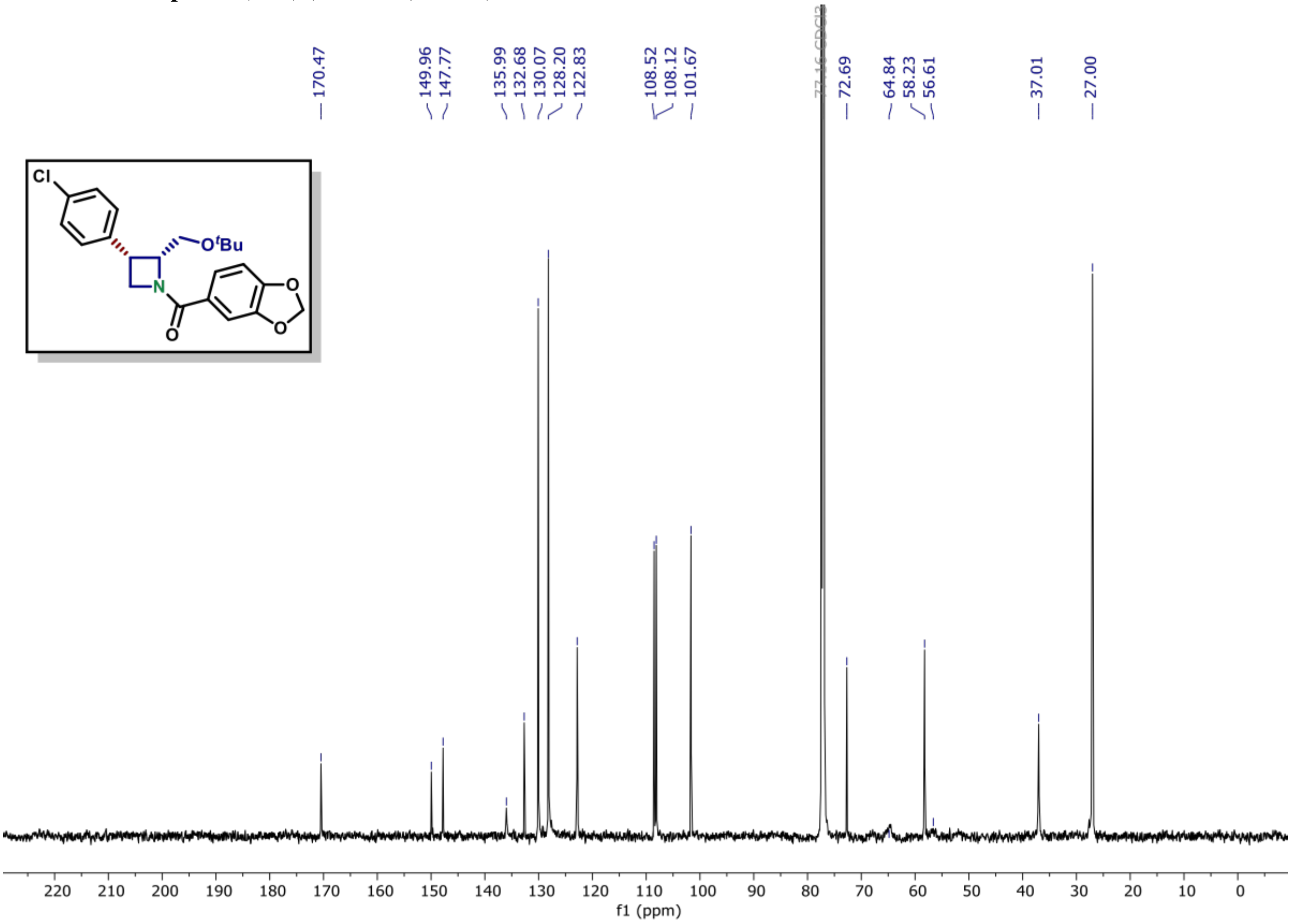

**[1]H NMR of compound (S6D) (600 MHz, $CDCl_3$)**

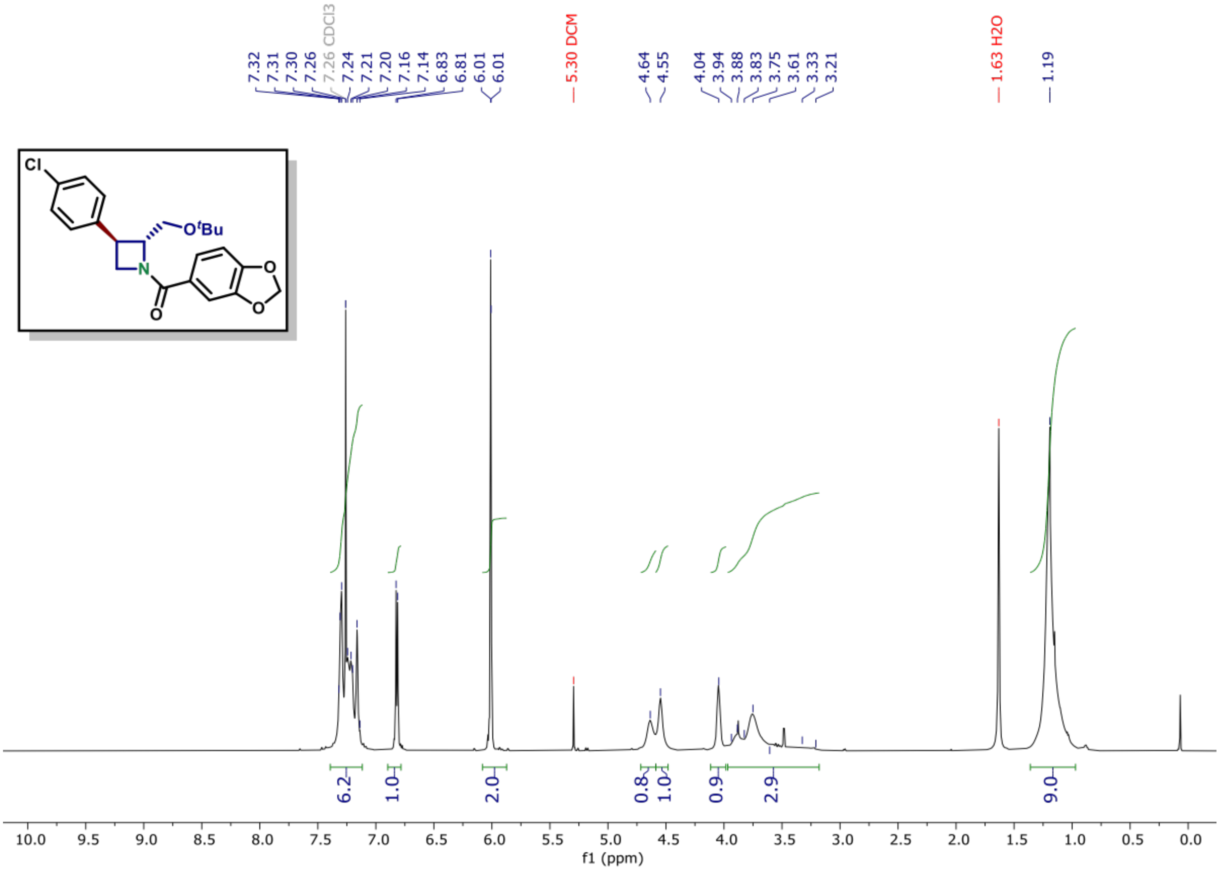

**[13]C NMR of compound (S6D) (150 MHz, $CDCl_3$)**

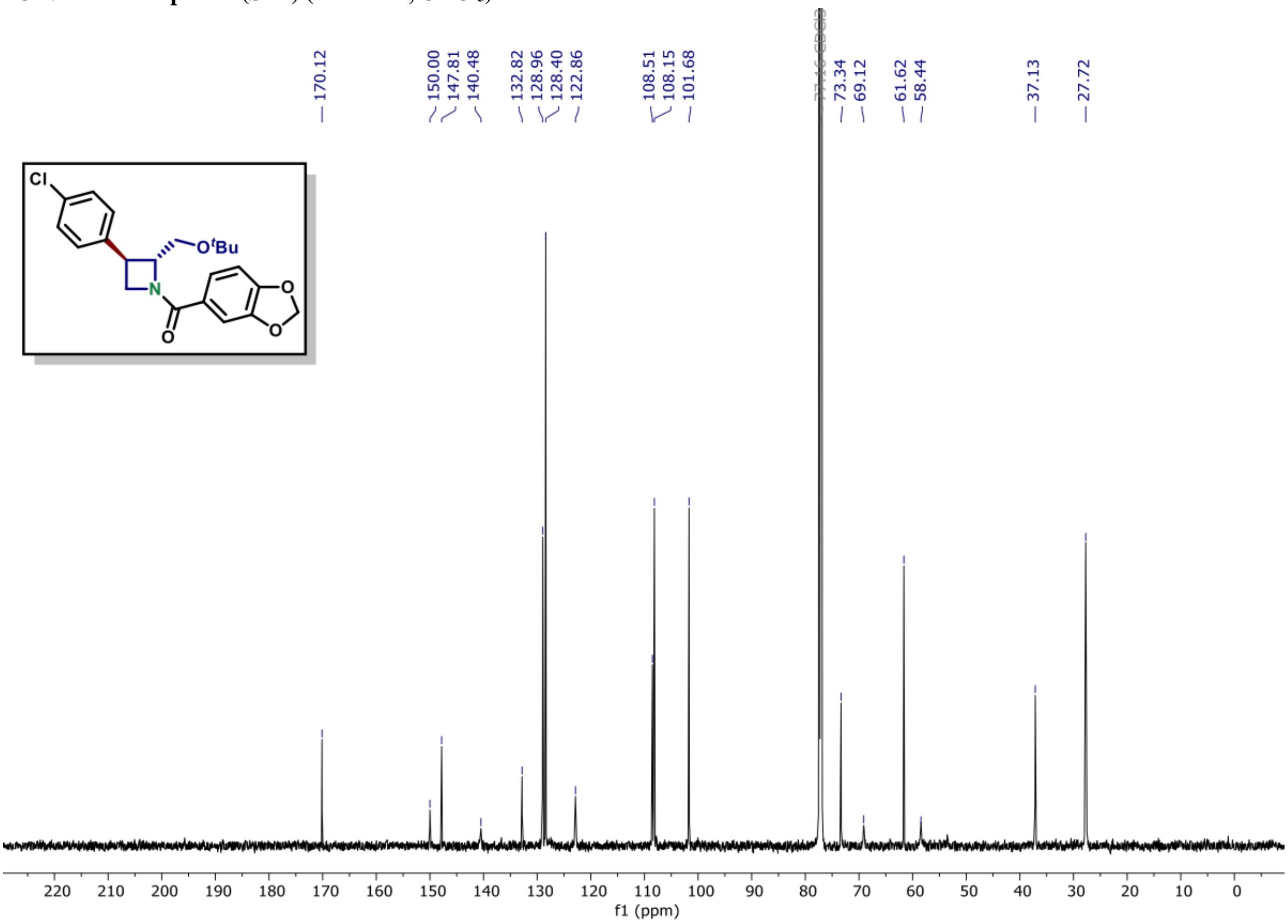

$^{1}$H NMR of compound (S7A) (600 MHz, $CDCl_3$)

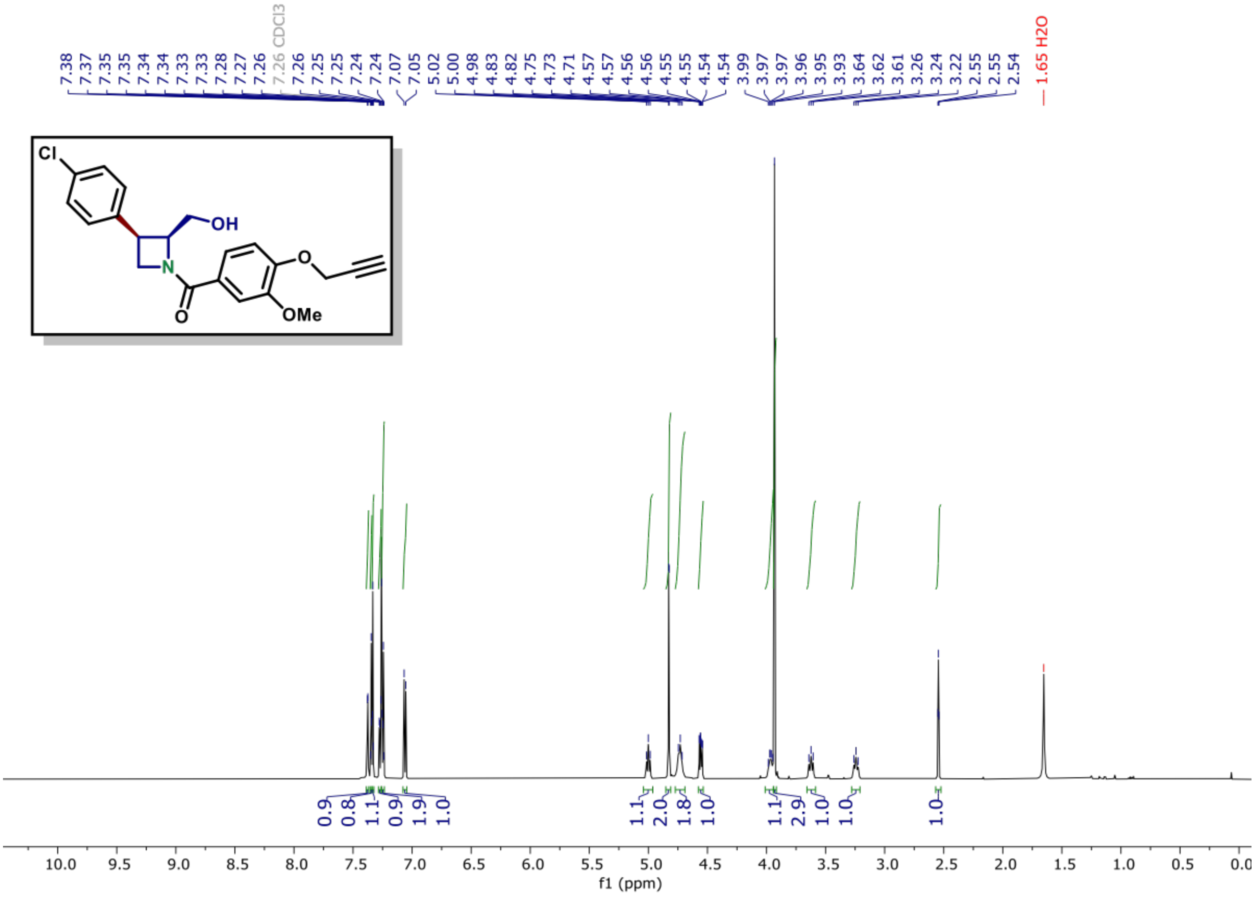

$^{13}$C NMR of compound (S7A) (150 MHz, $CDCl_3$)

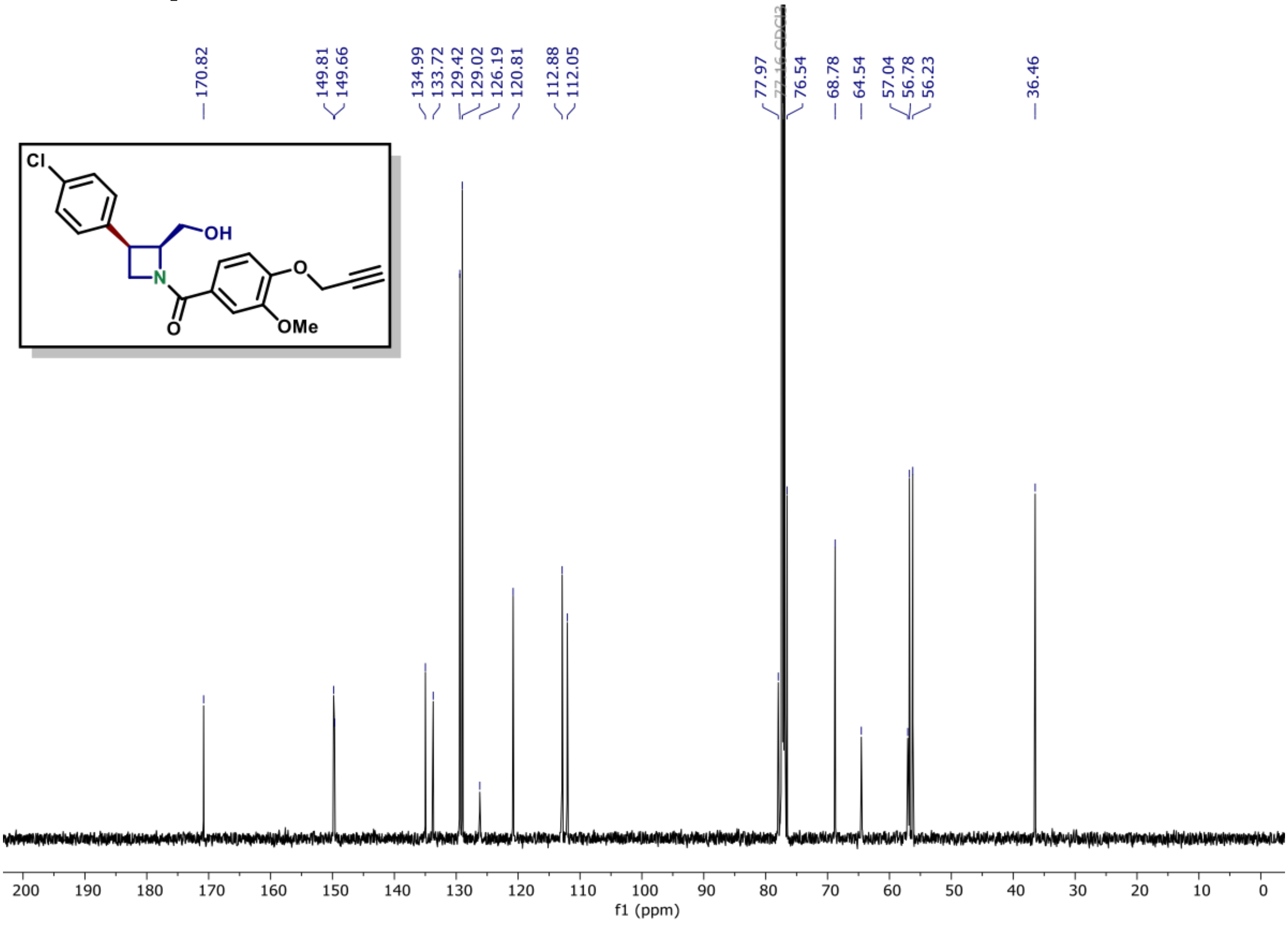

[1]H NMR of compound (S7B) (600 MHz, $CDCl_3$)

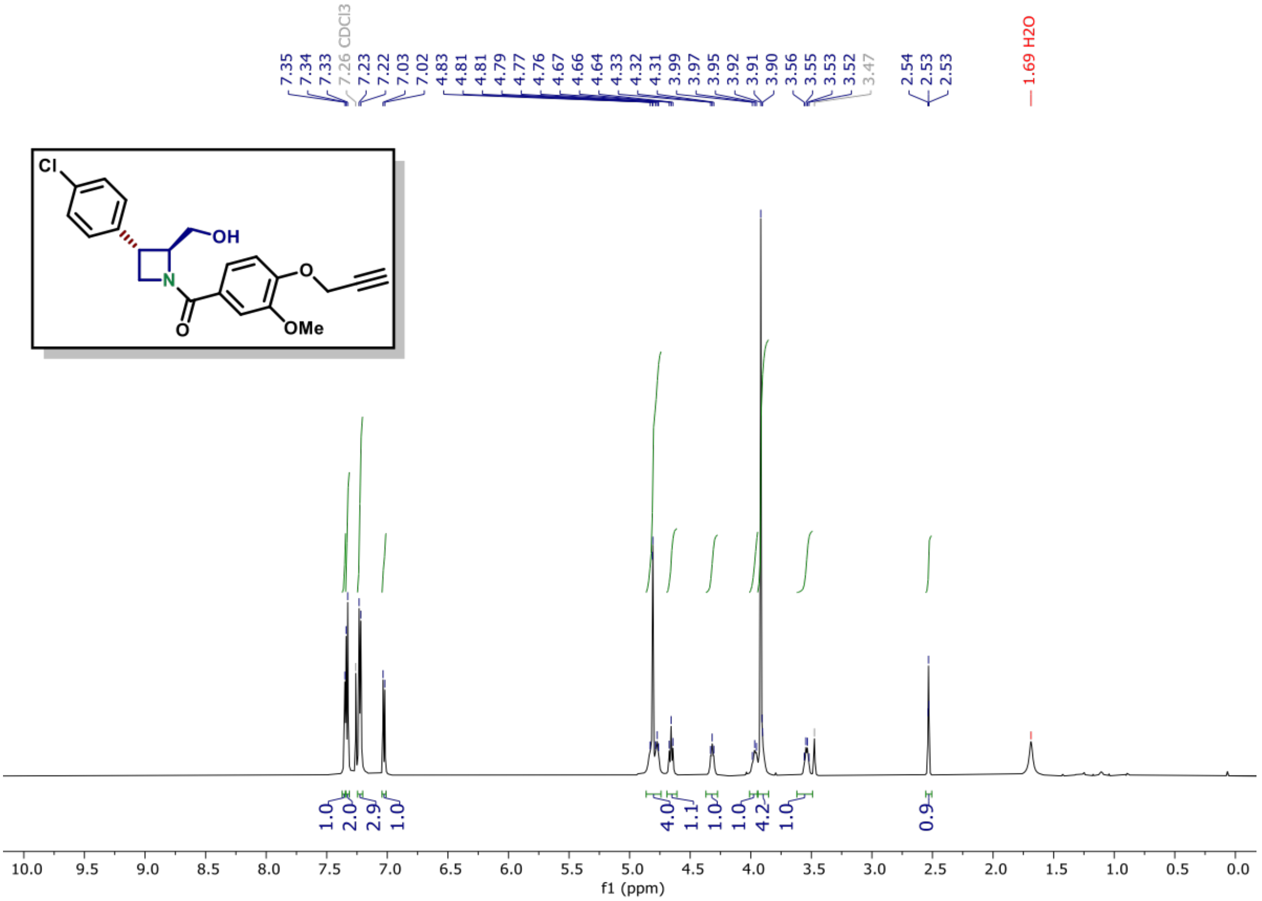

[13]C NMR of compound (S7B) (150 MHz, $CDCl_3$)

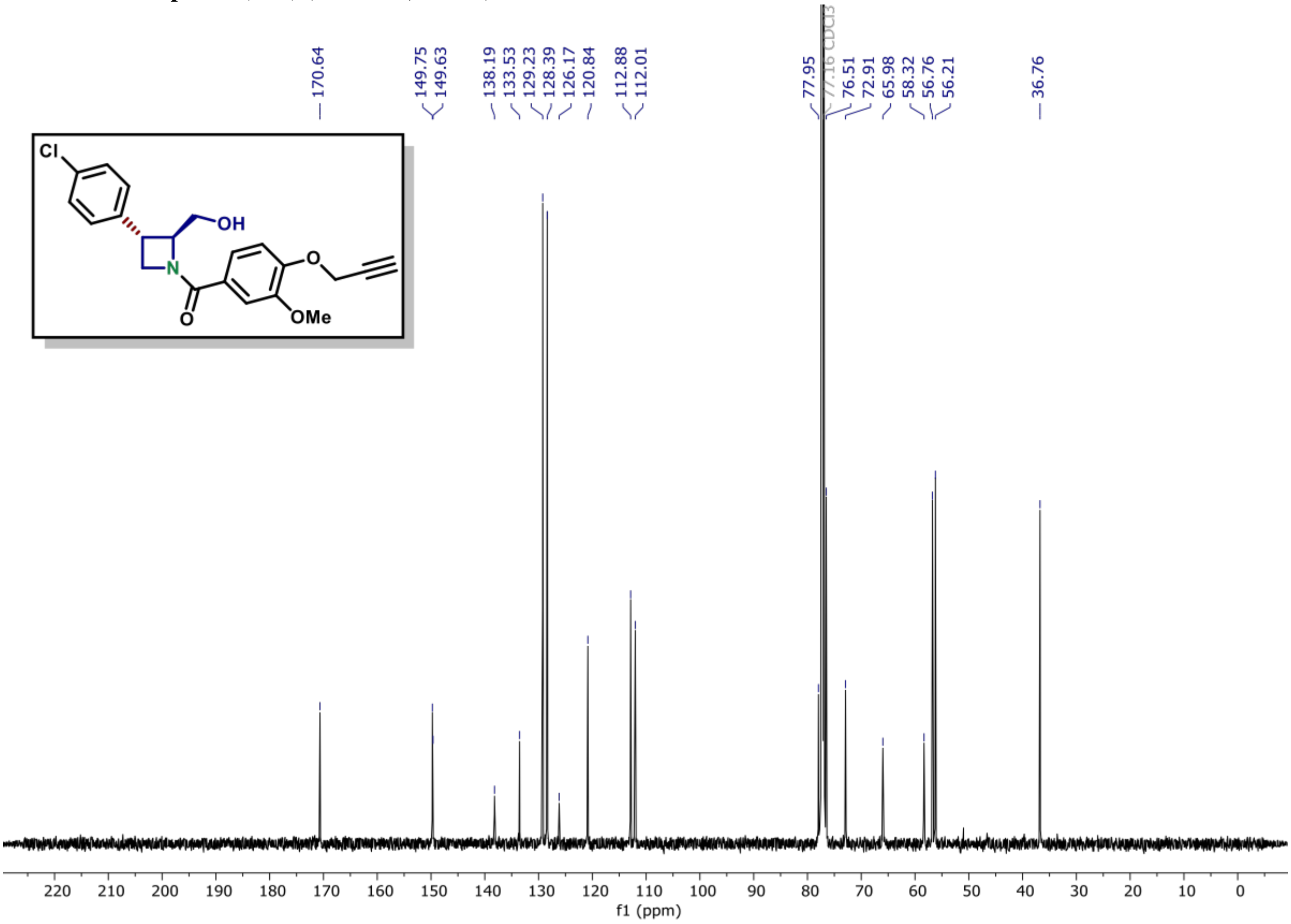

$^{1}$H NMR of compound (S7C) (600 MHz, $CDCl_3$)

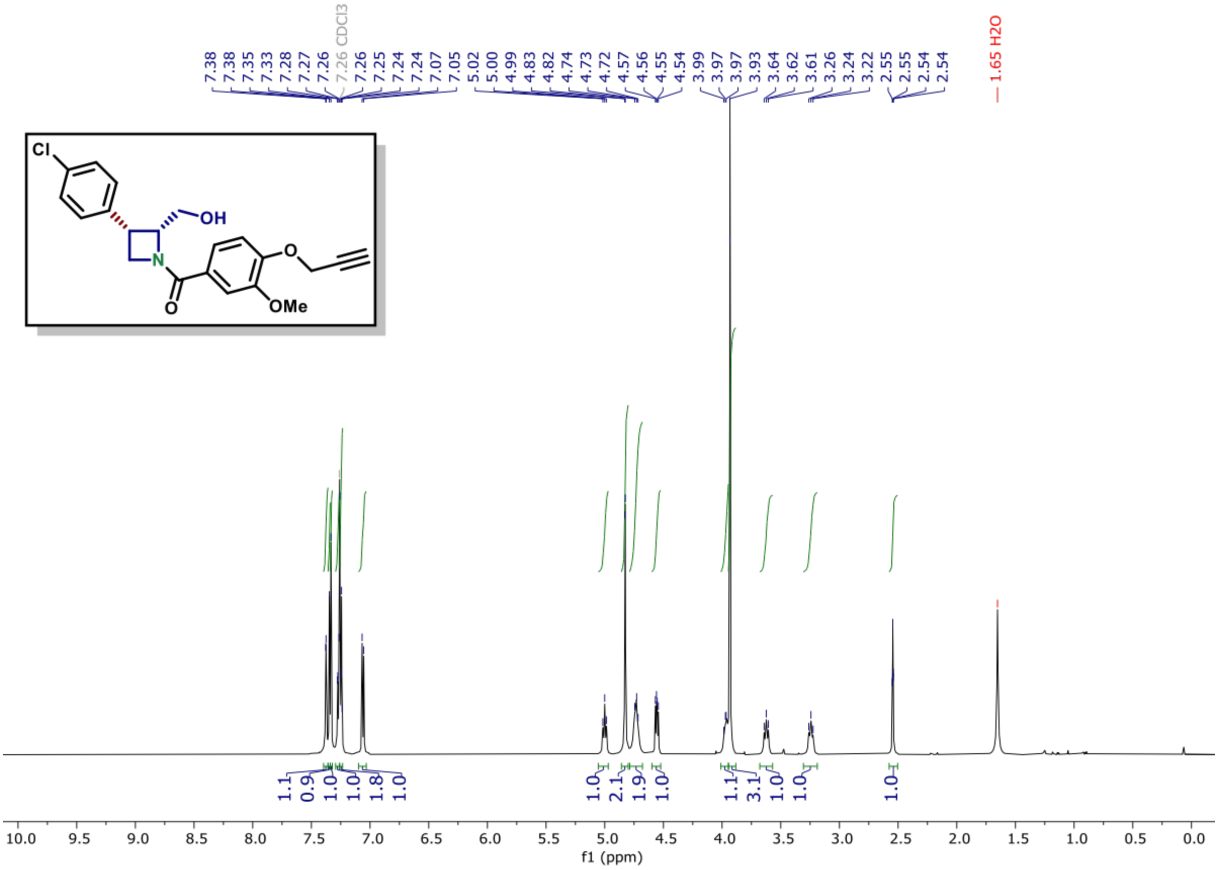

$^{13}$C NMR of compound (S7C) (150 MHz, $CDCl_3$)

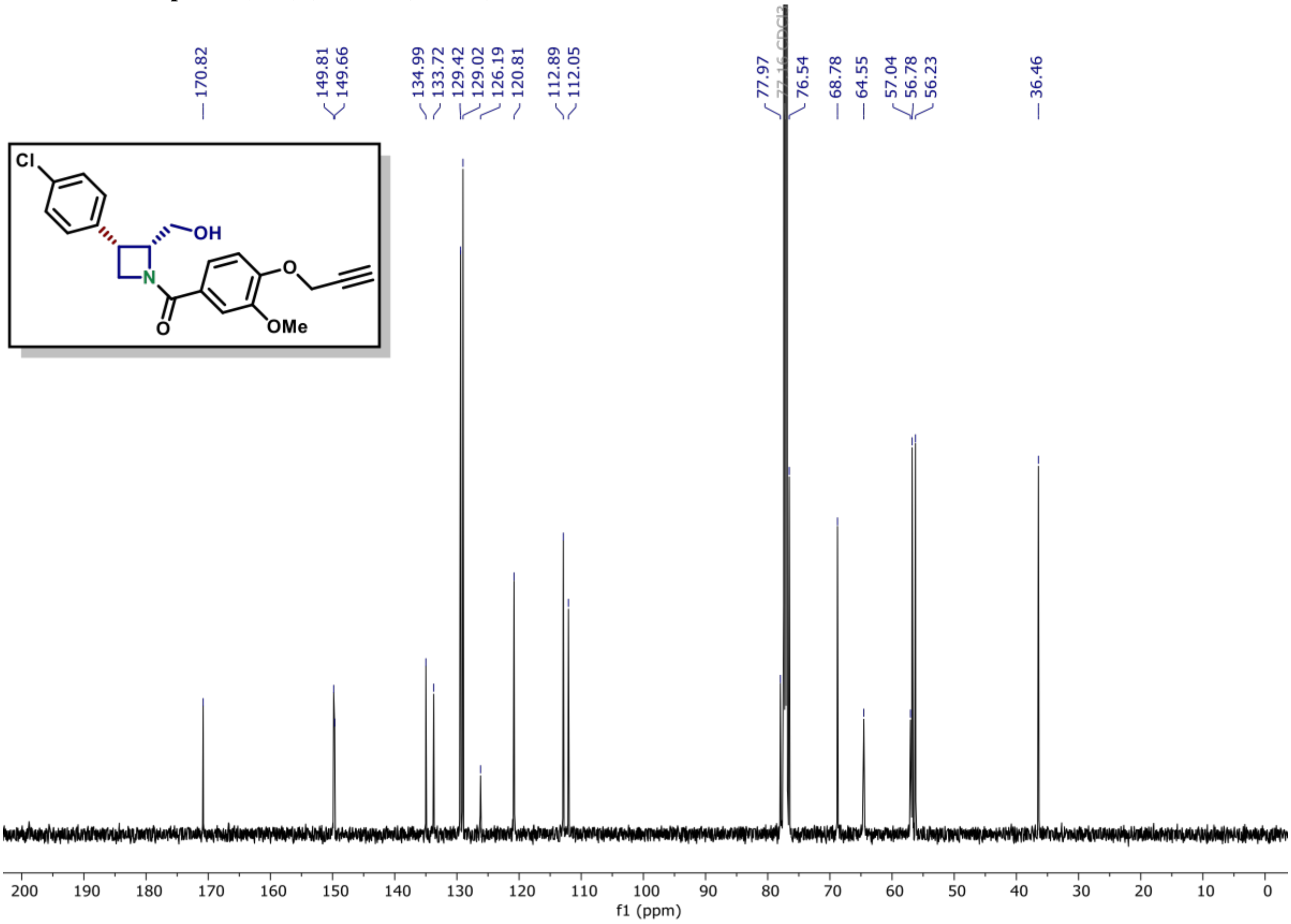

**[1]H NMR of compound (S7D) (600 MHz, $CDCl_3$)**

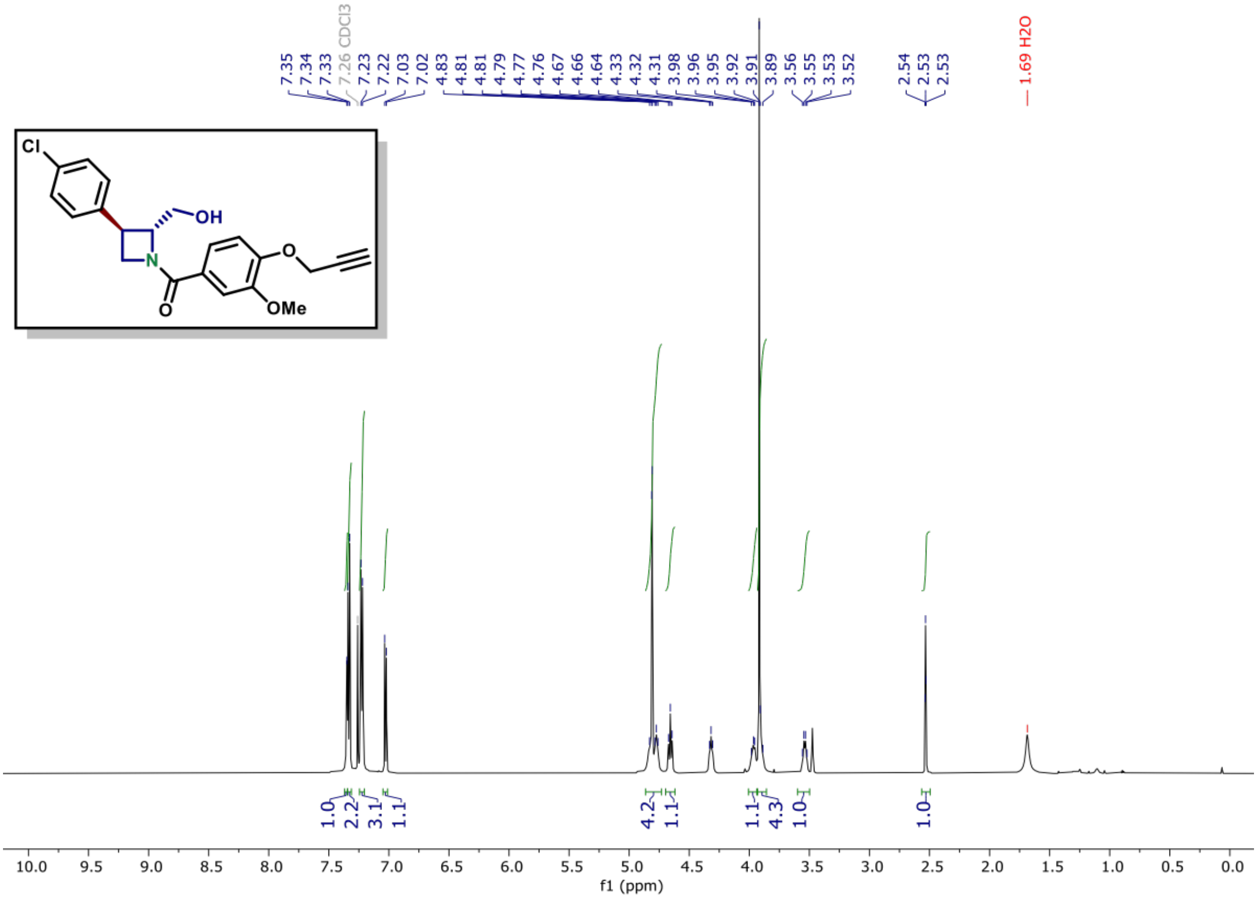

**[13]C NMR of compound (S7D) (150 MHz, $CDCl_3$)**

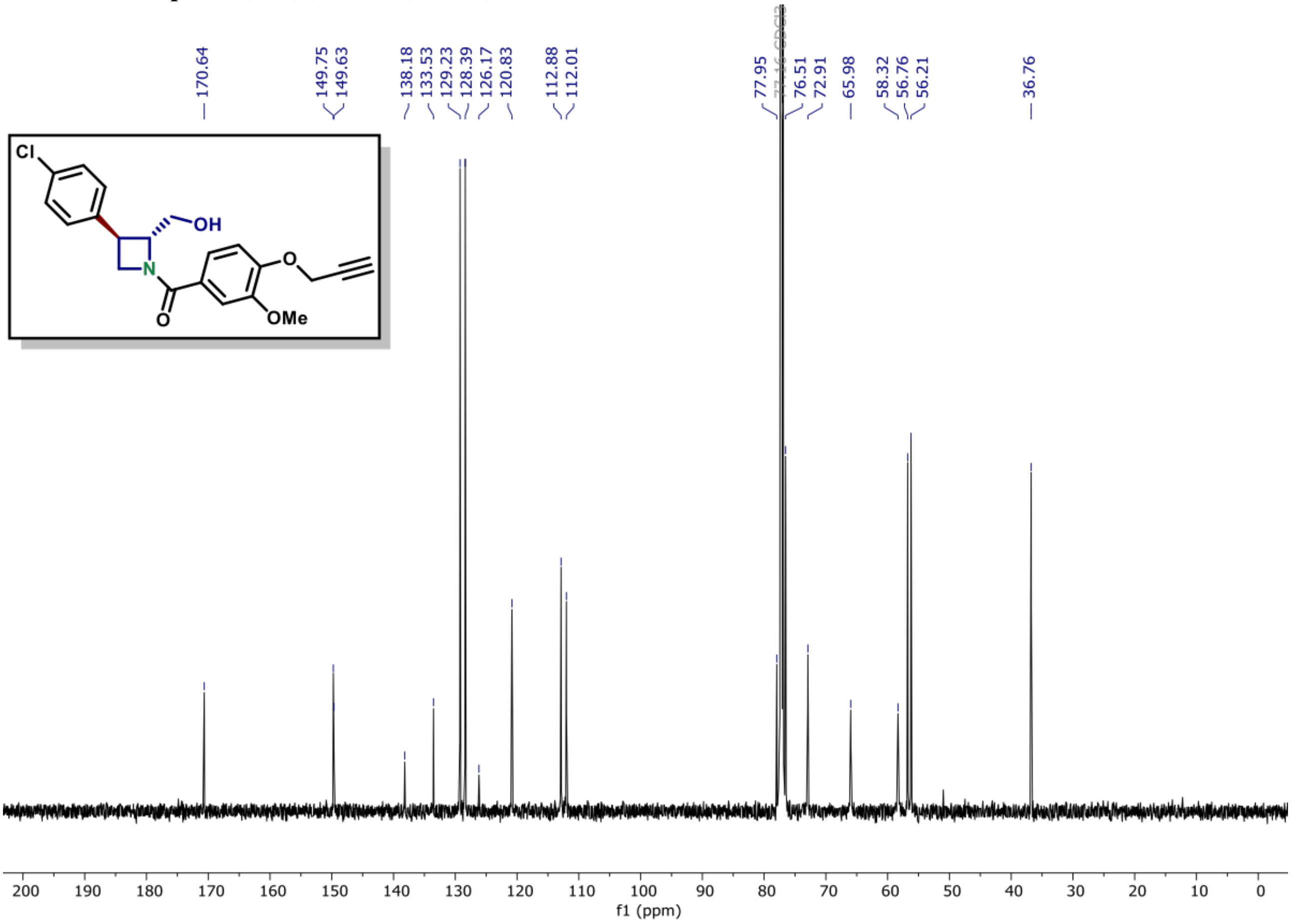

$^{1}$H NMR of compound (S8A) (600 MHz, $CDCl_3$)

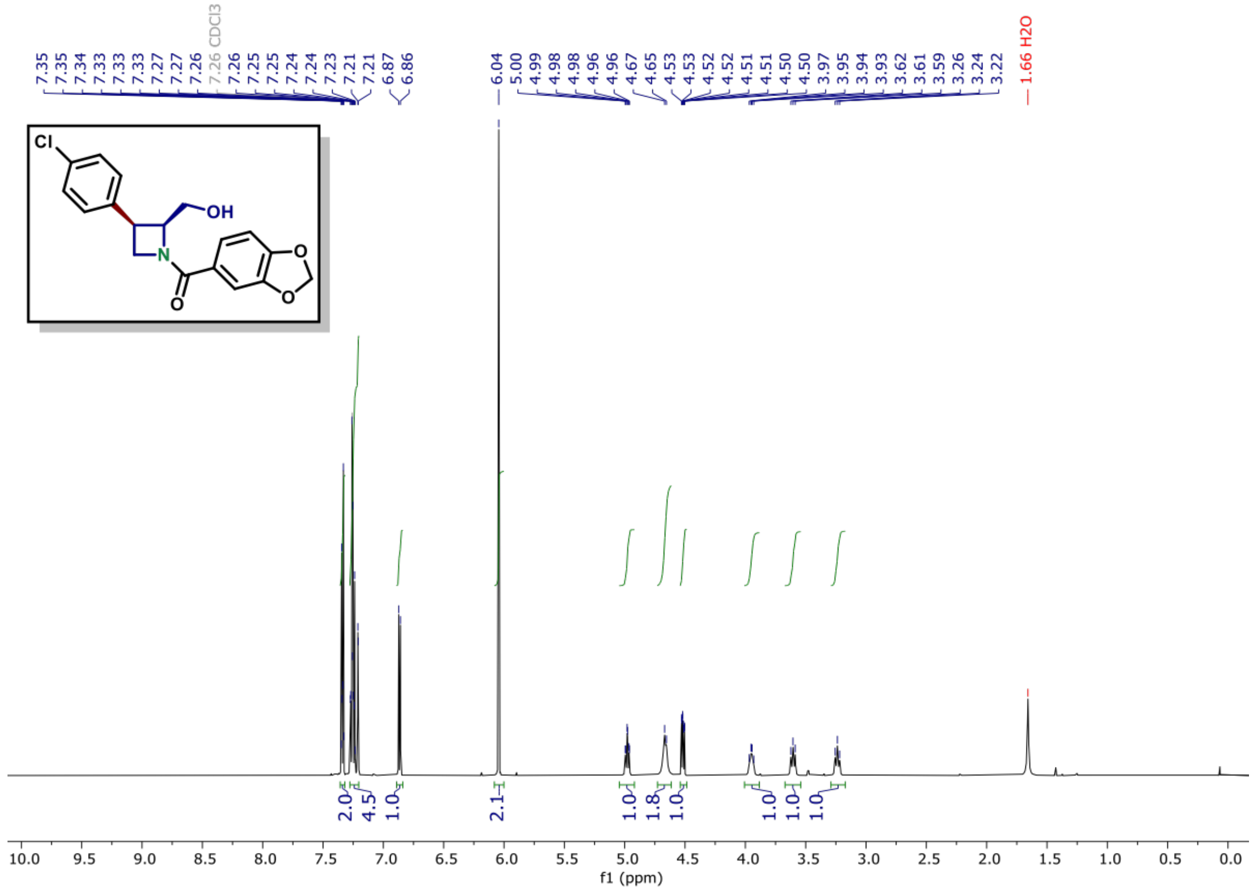

$^{13}$C NMR of compound (S8A) (150 MHz, $CDCl_3$)

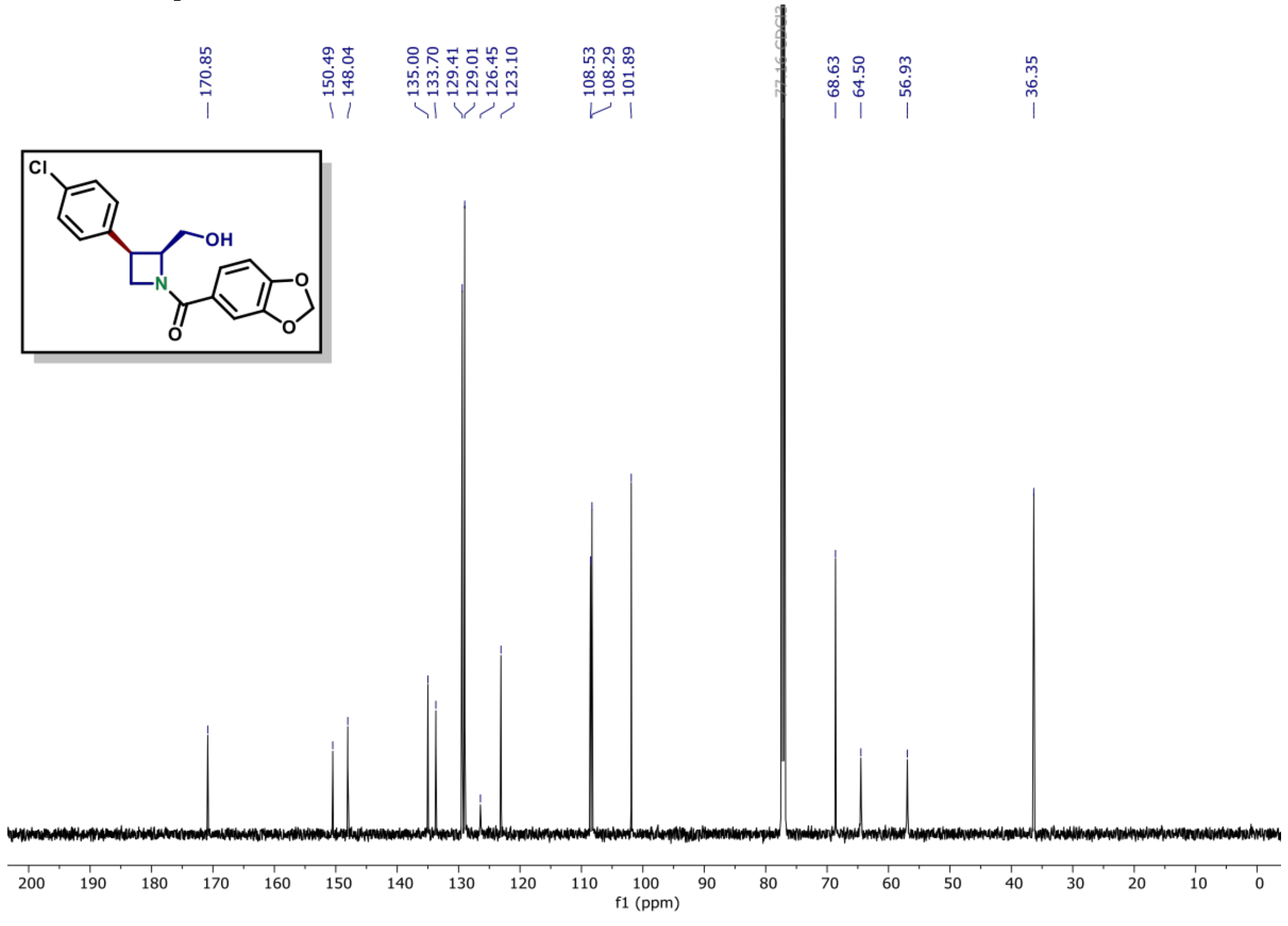

[1]H NMR of compound (S8B) (600 MHz, $CDCl_3$)

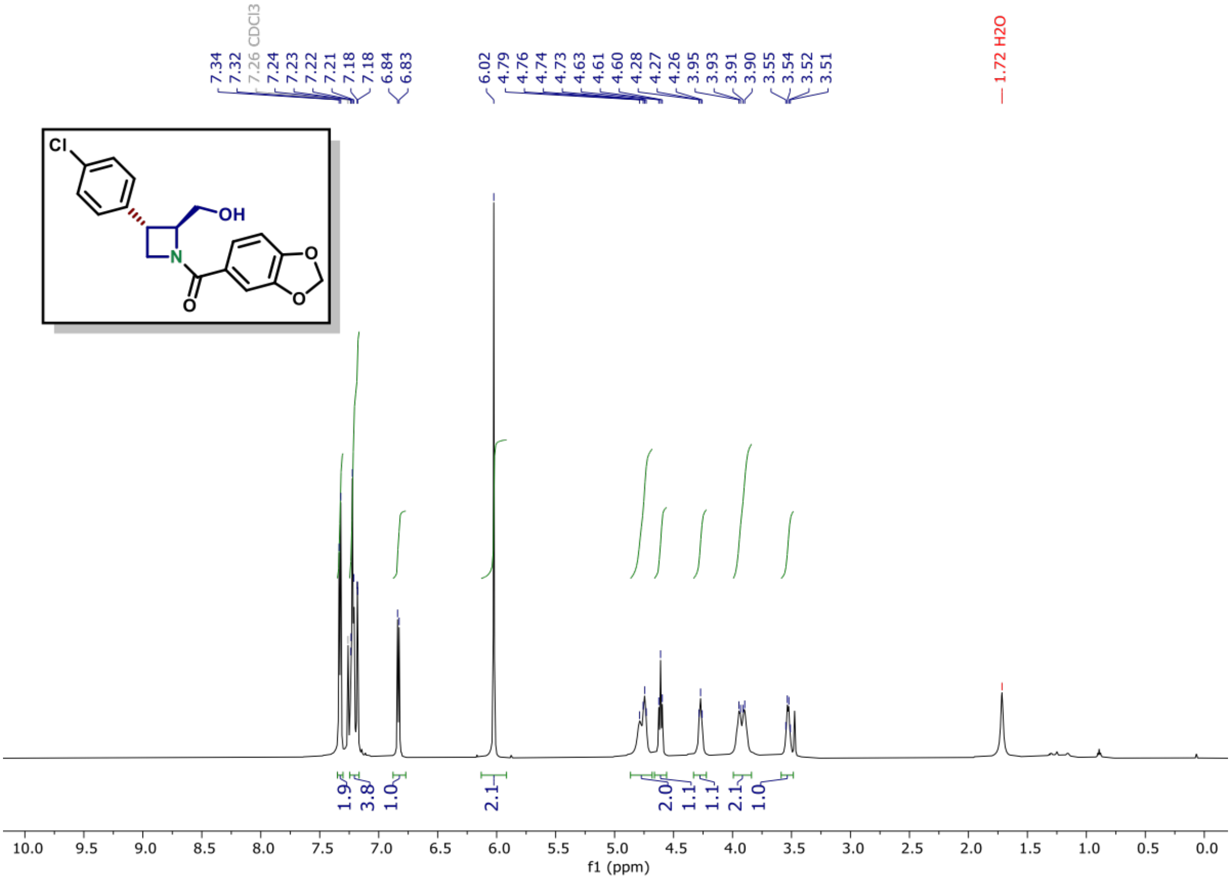

[13]C NMR of compound (S8B) (150 MHz, $CDCl_3$)

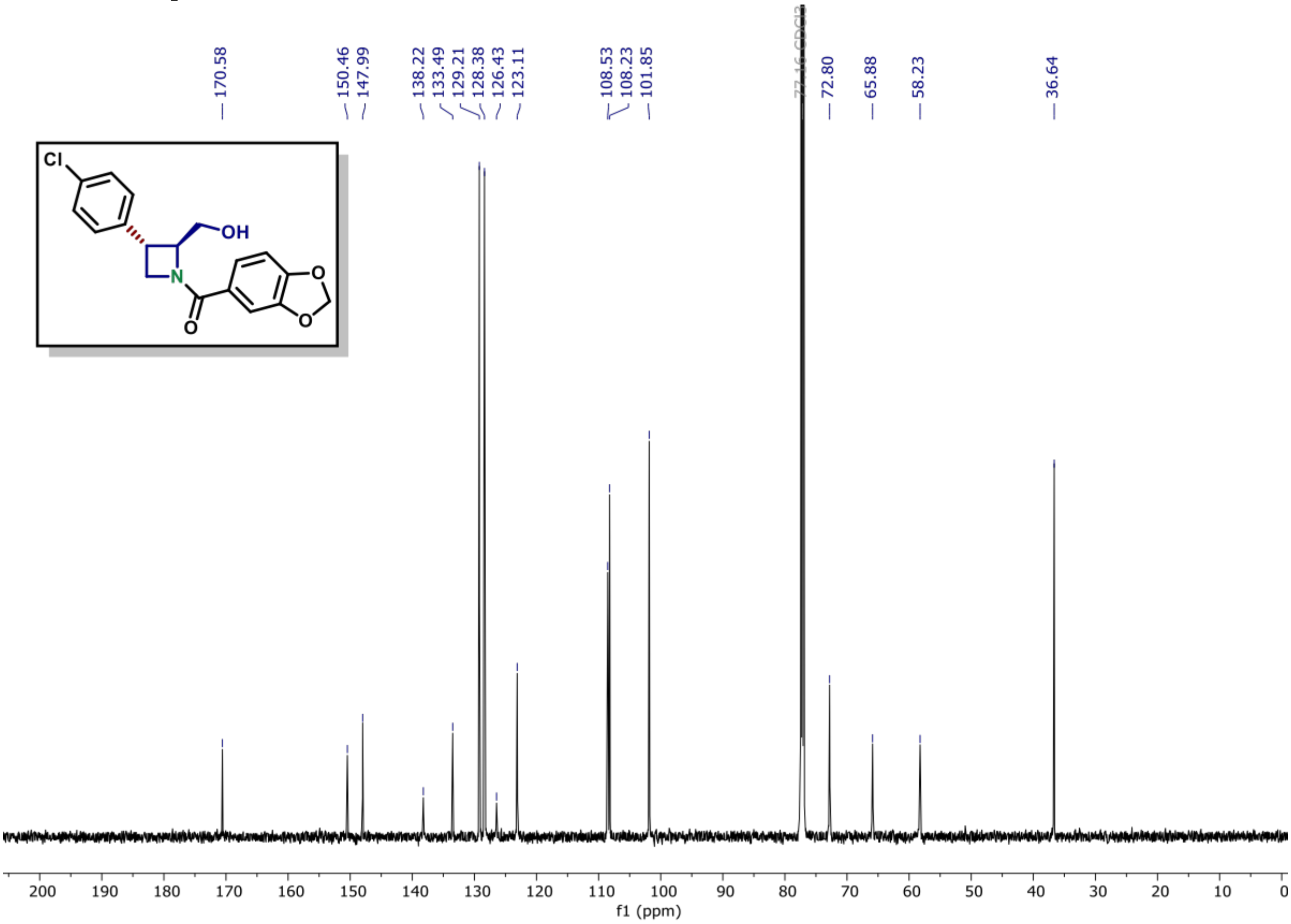

[1]H NMR of compound (S8C) (600 MHz, $CDCl_3$)

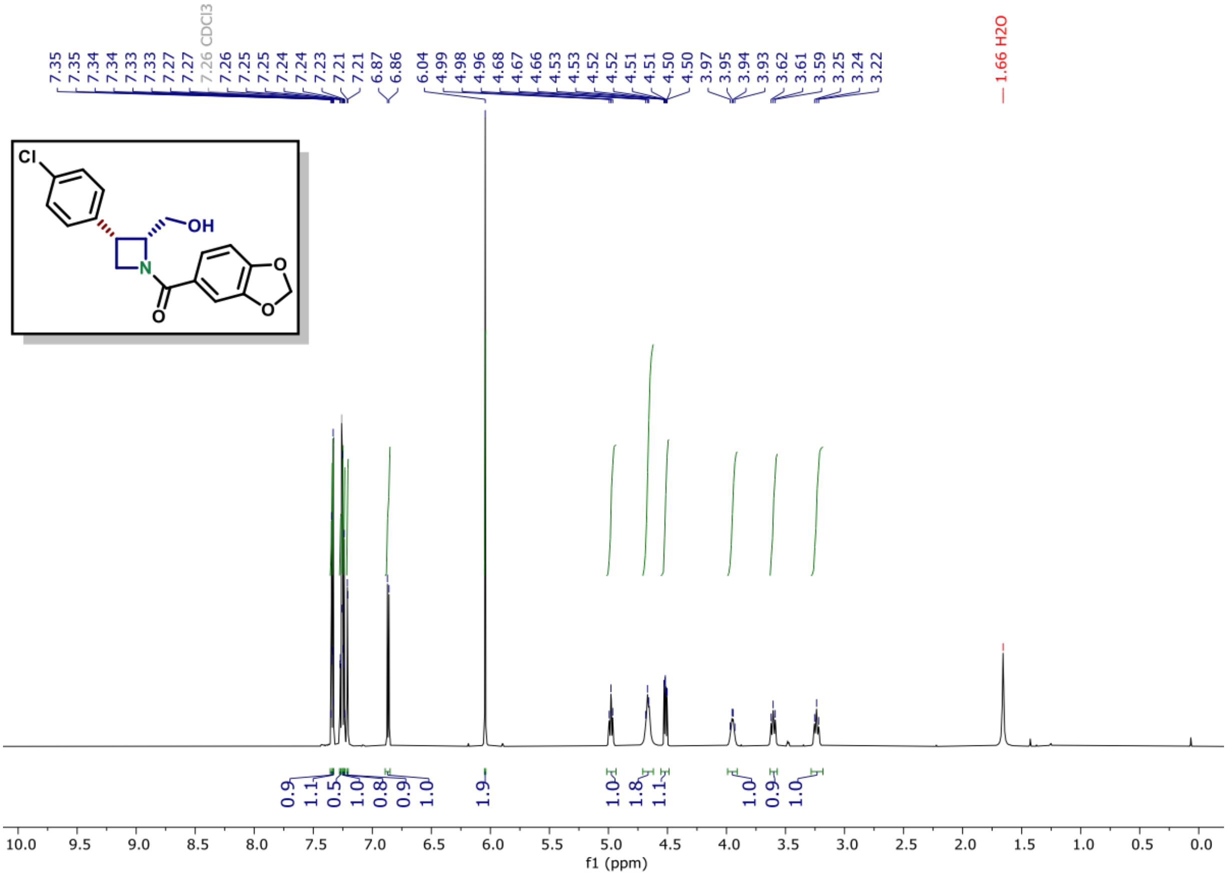

[13]C NMR of compound (S8C) (150 MHz, $CDCl_3$)

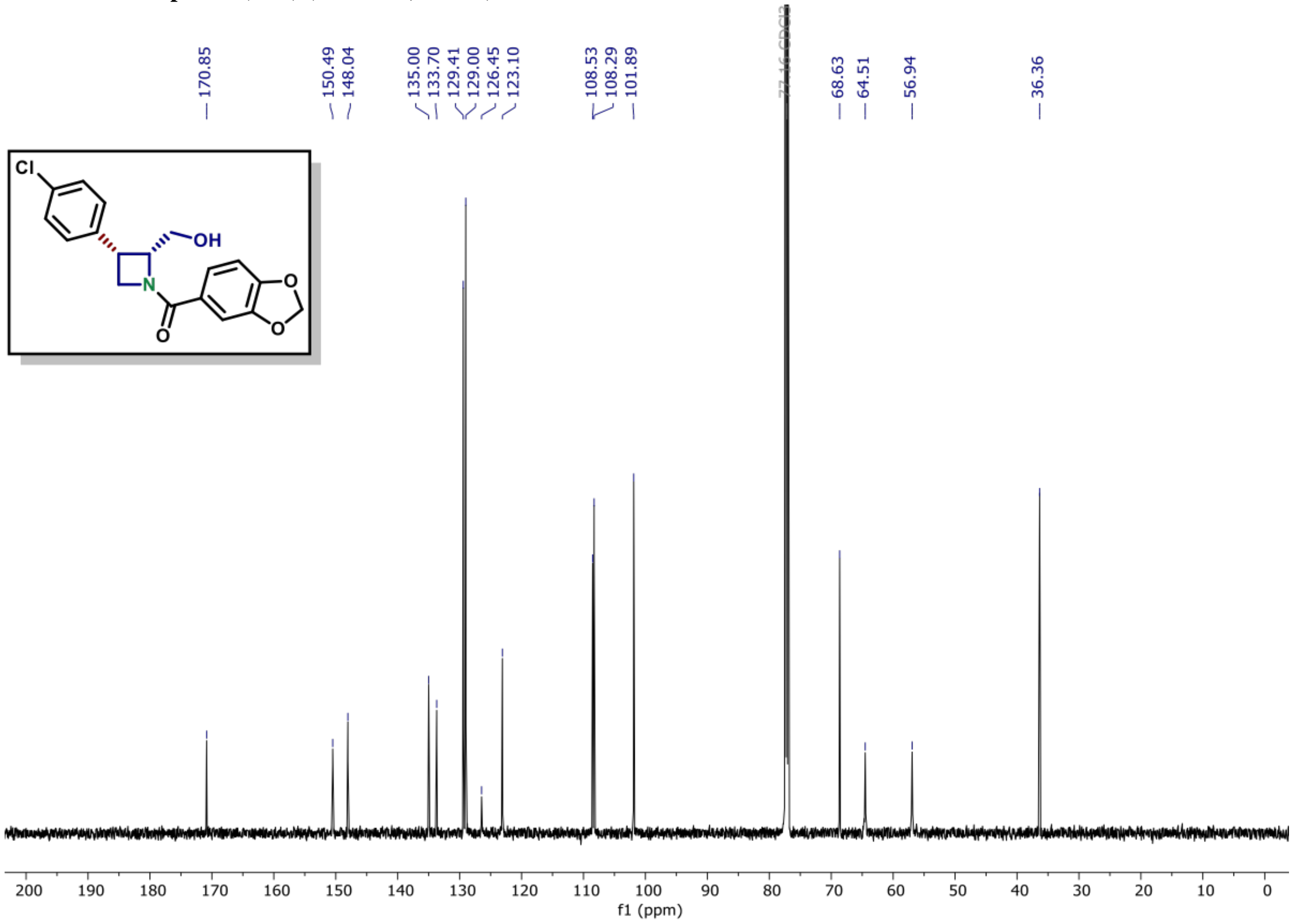

[1]H NMR of compound (S8D) (600 MHz, $CDCl_3$)

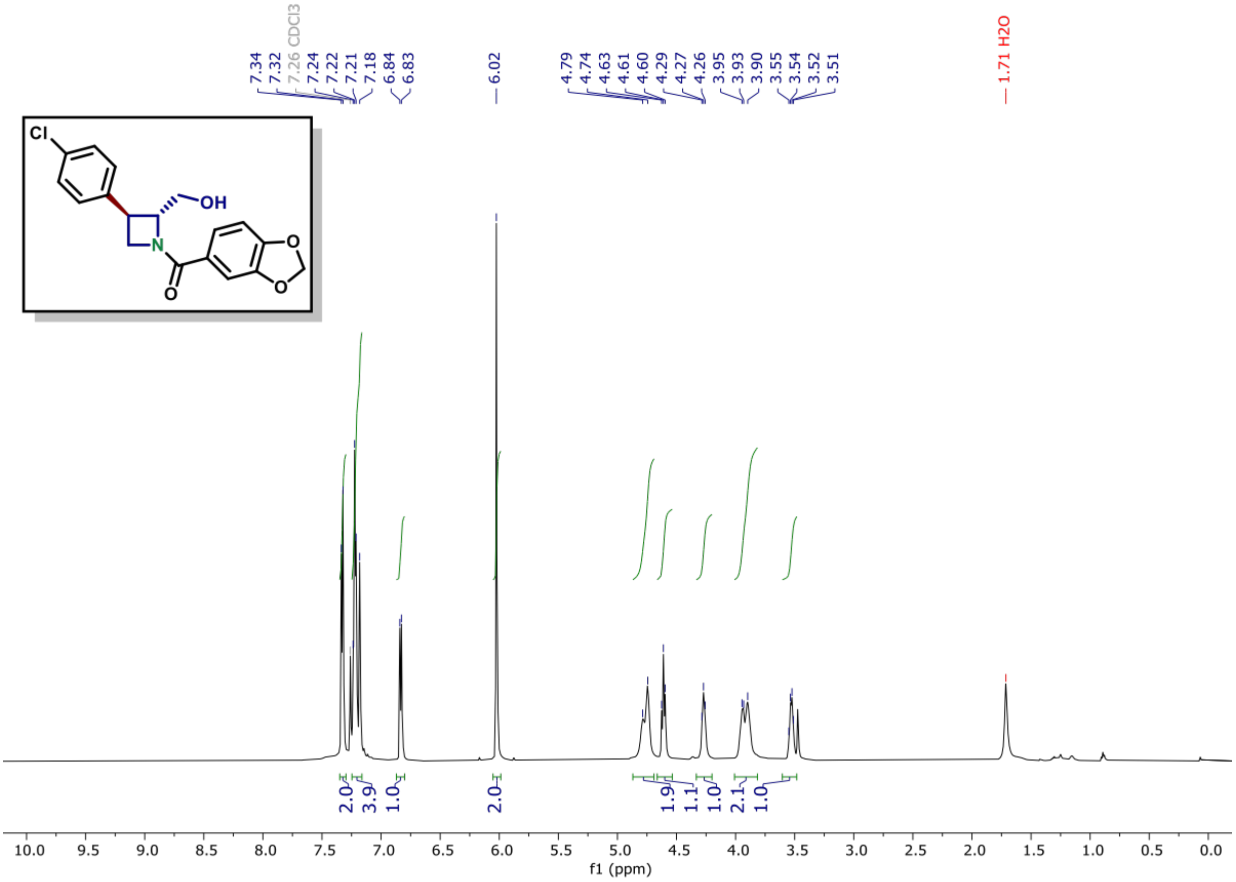

[13]C NMR of compound (S8D) (150 MHz, $CDCl_3$)

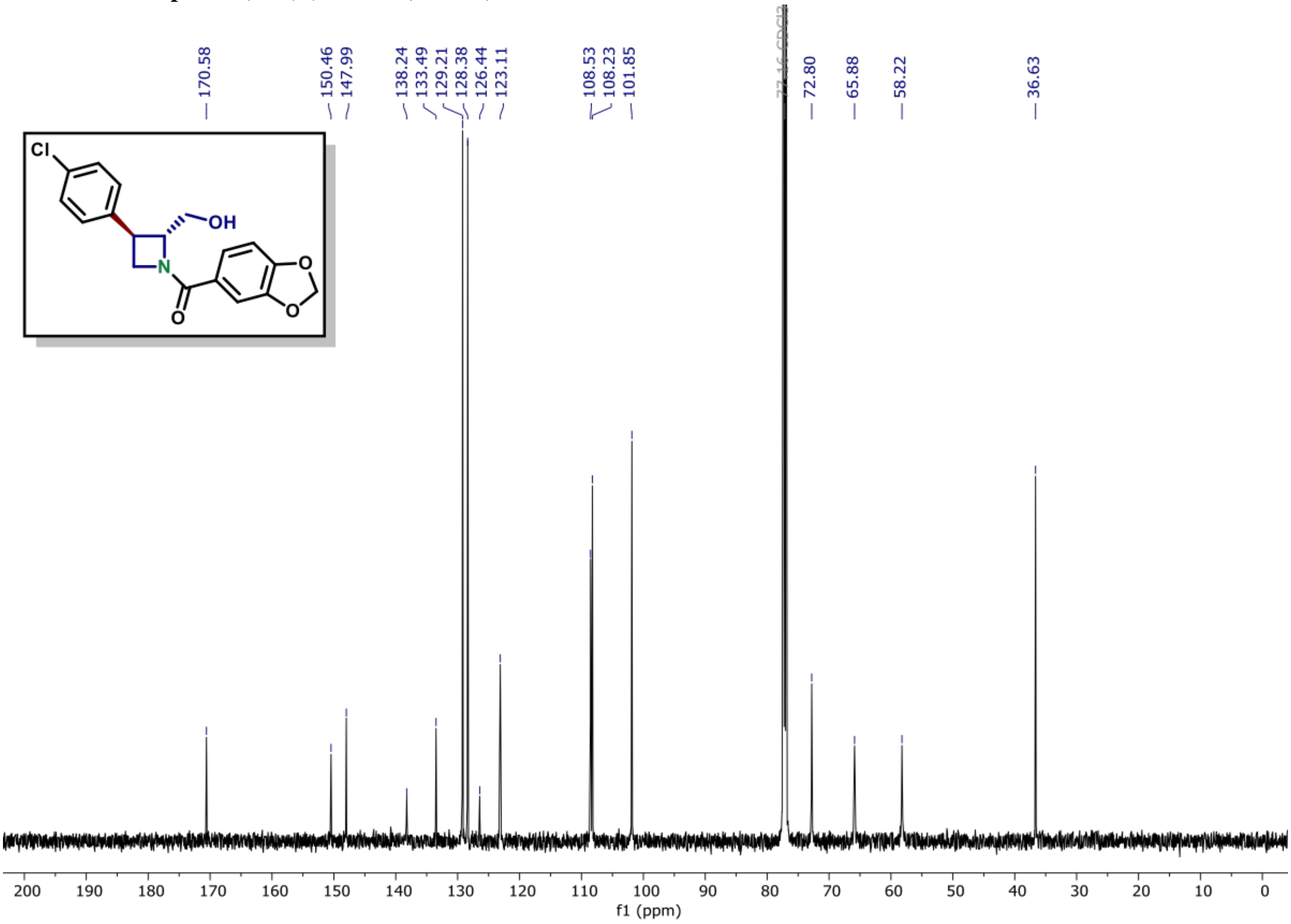

$^{31}$P NMR of compound (9) (162 MHz, $CDCl_3$)

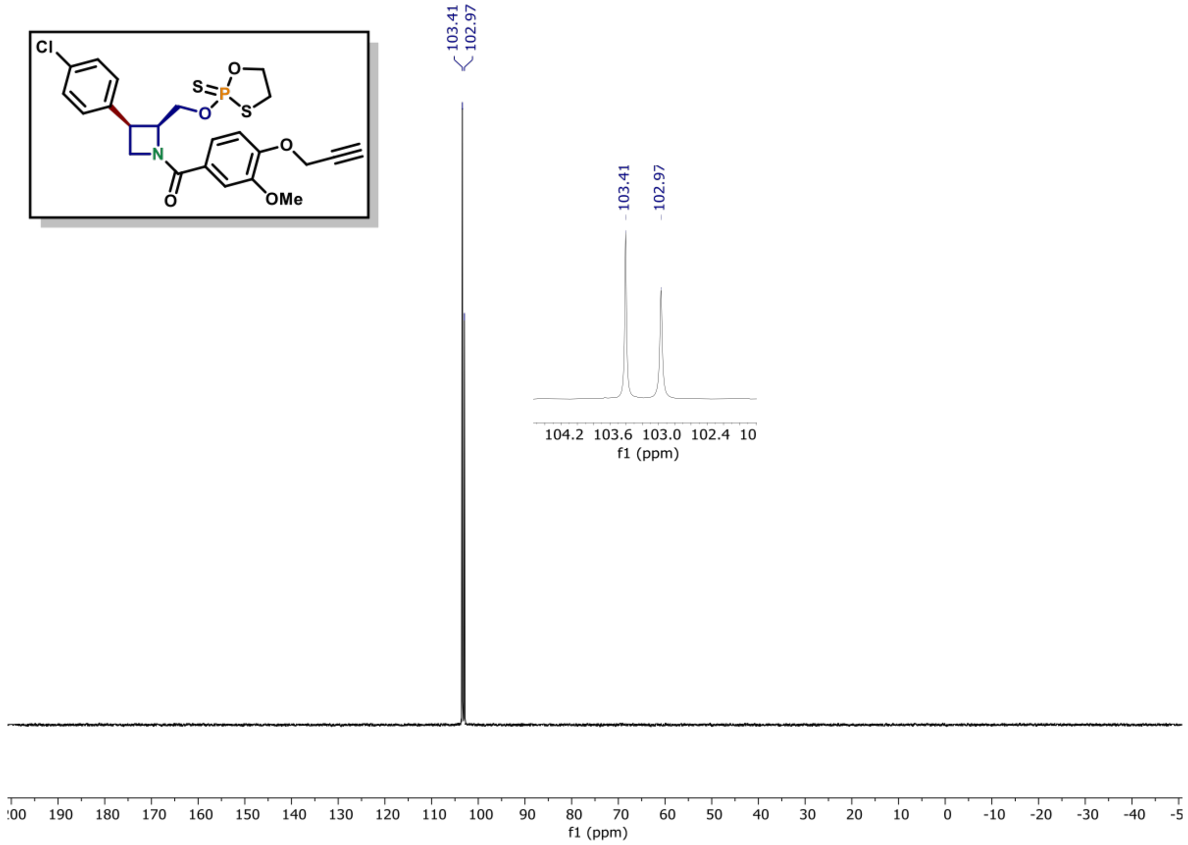

$^{1}$H NMR of compound (9) (600 MHz, $CDCl_3$)

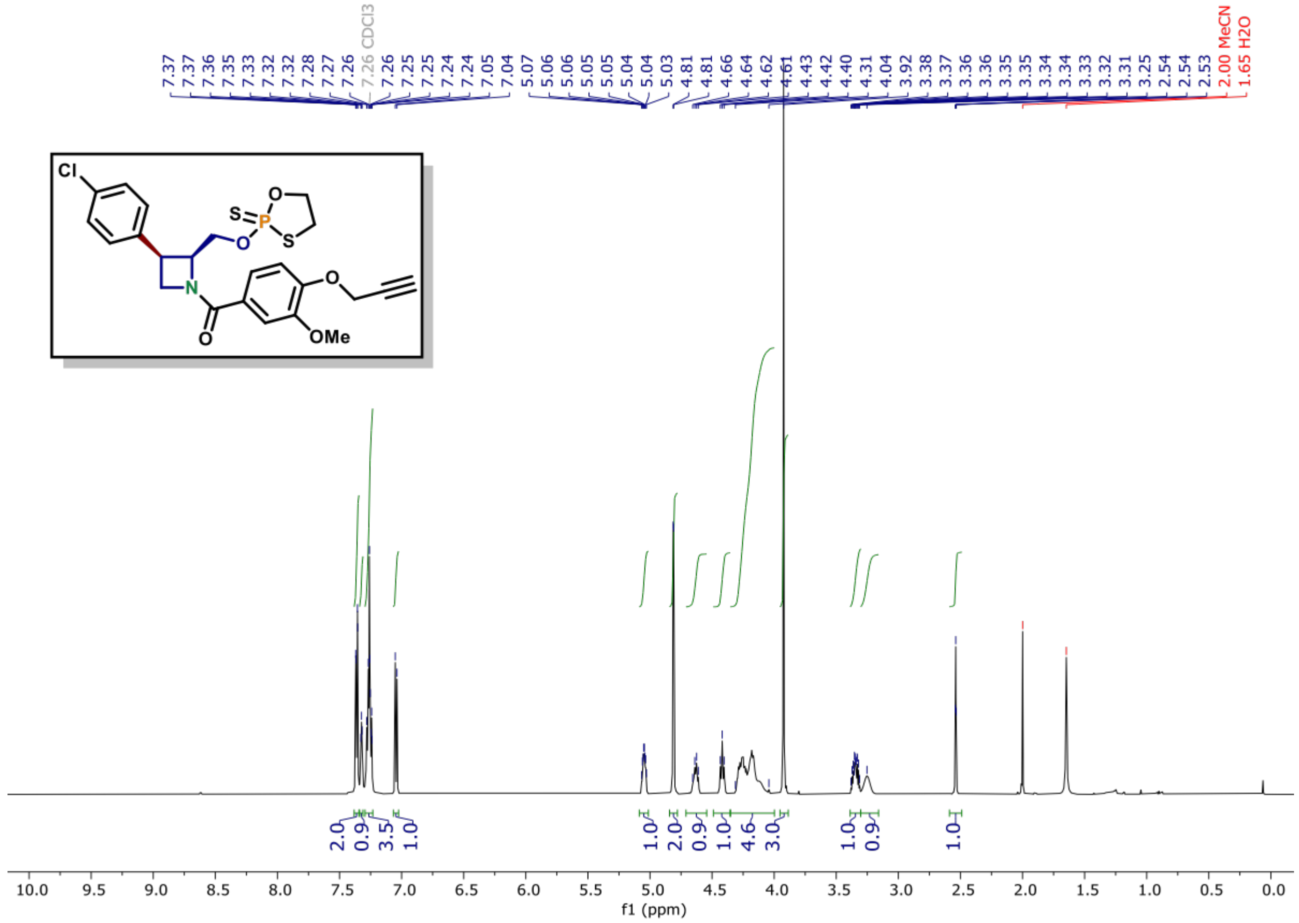

$^{13}$C NMR of compound (9) (150 MHz, CDCl$_3$)

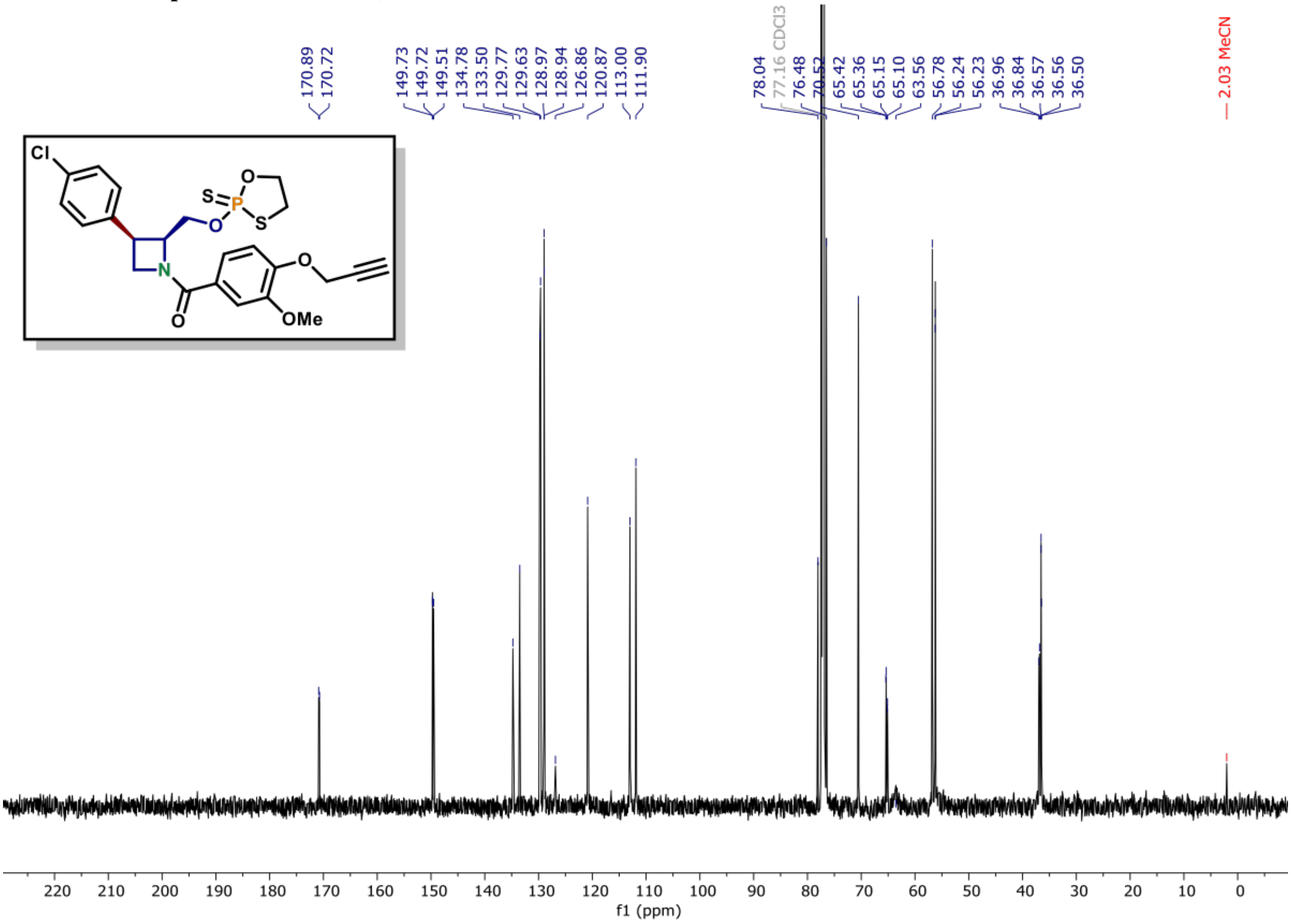

$^{31}$P NMR of compound (11) (162 MHz, $CDCl_3$)

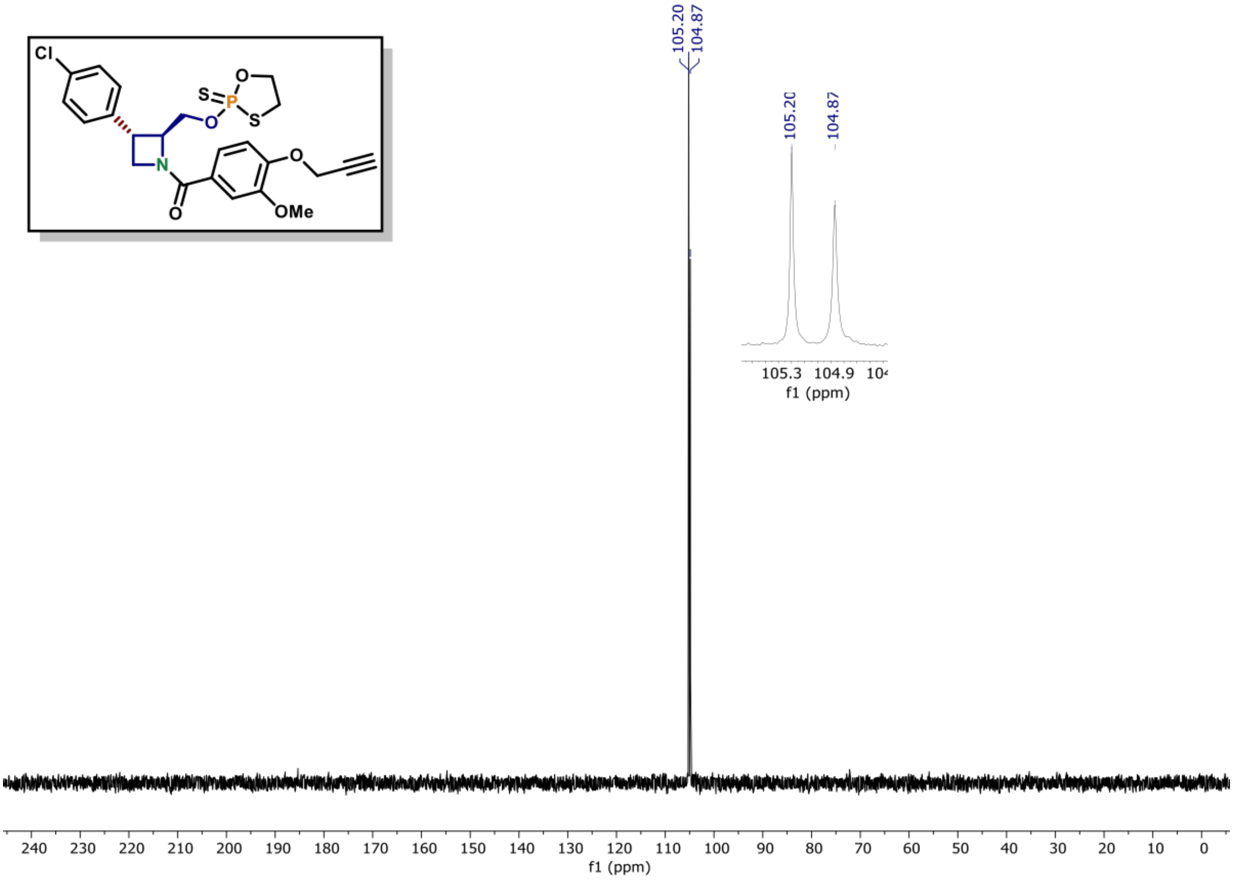

$^{1}$H NMR of compound (11) (600 MHz, $CDCl_3$)

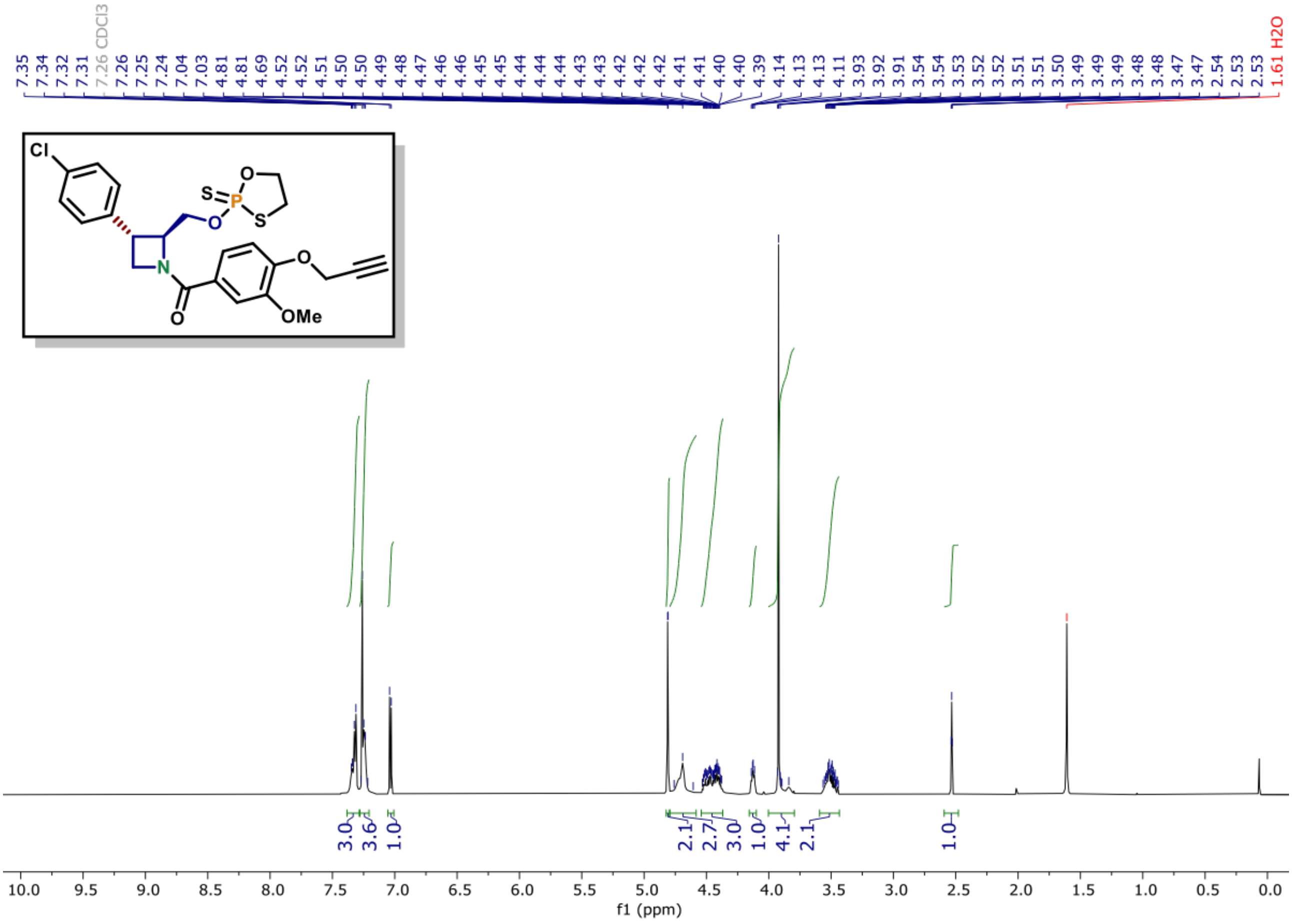

**[13]C NMR of compound (11) (150 MHz, CDCl₃)**

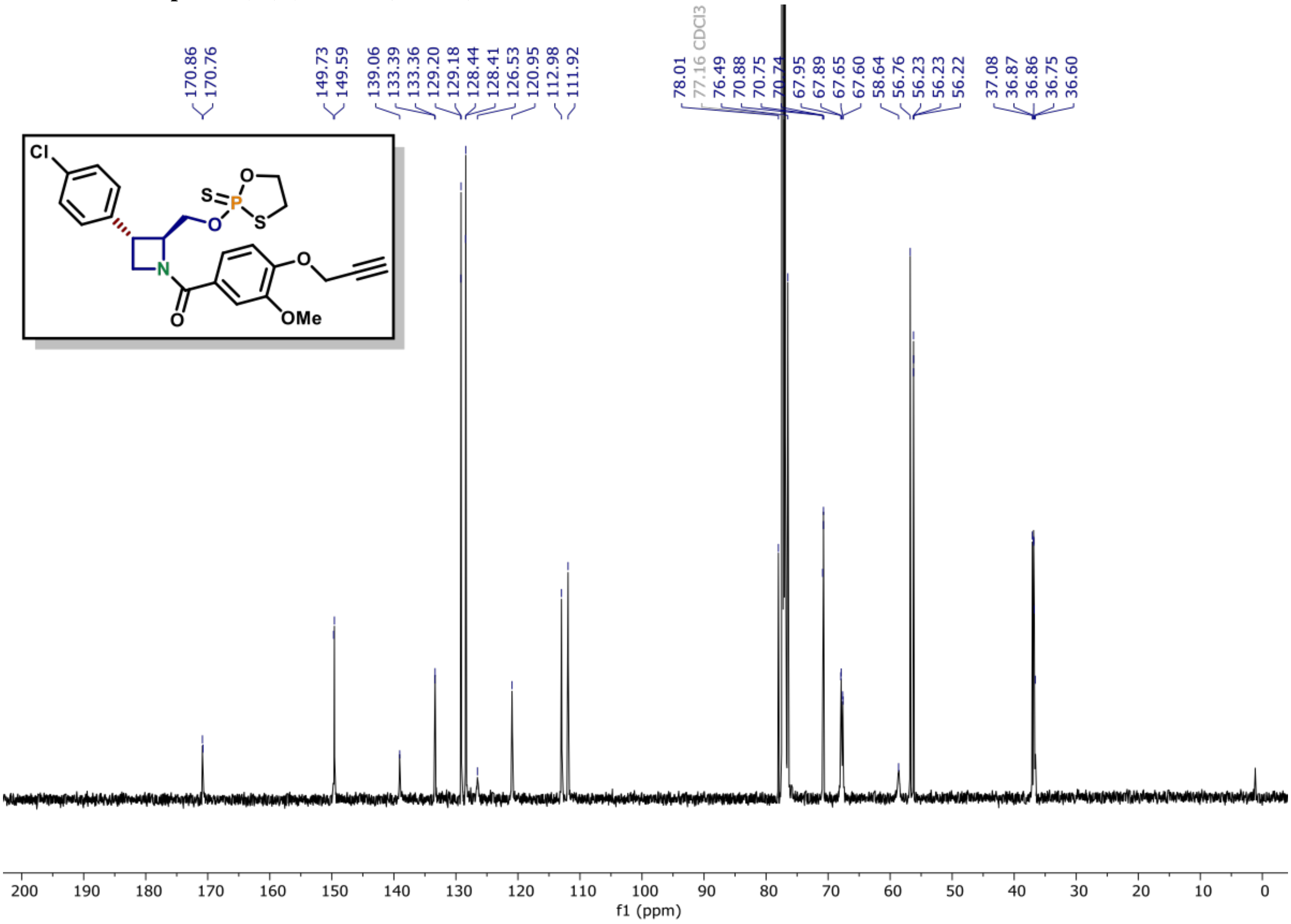

[31]P NMR of compound (10) (162 MHz, $CDCl_3$)

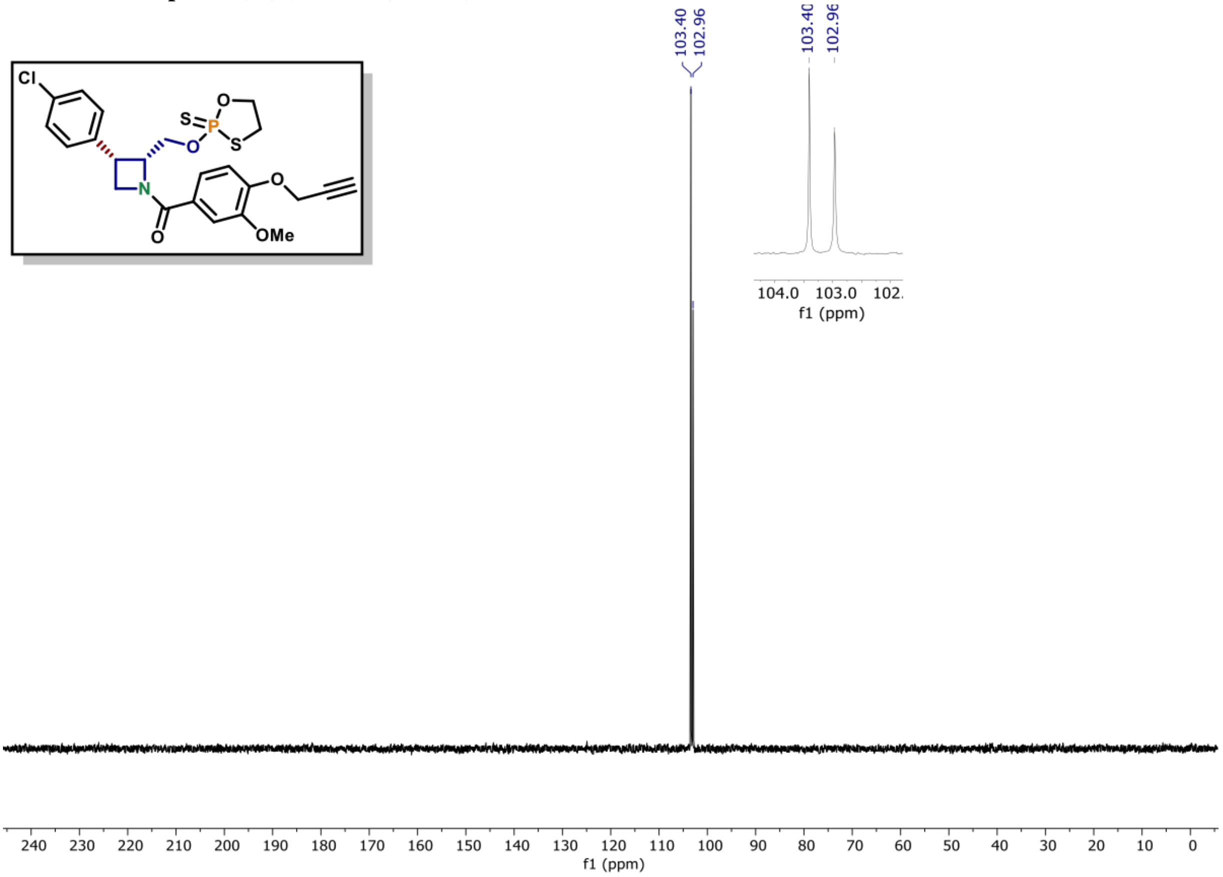

[1]H NMR of compound (10) (600 MHz, $CDCl_3$)

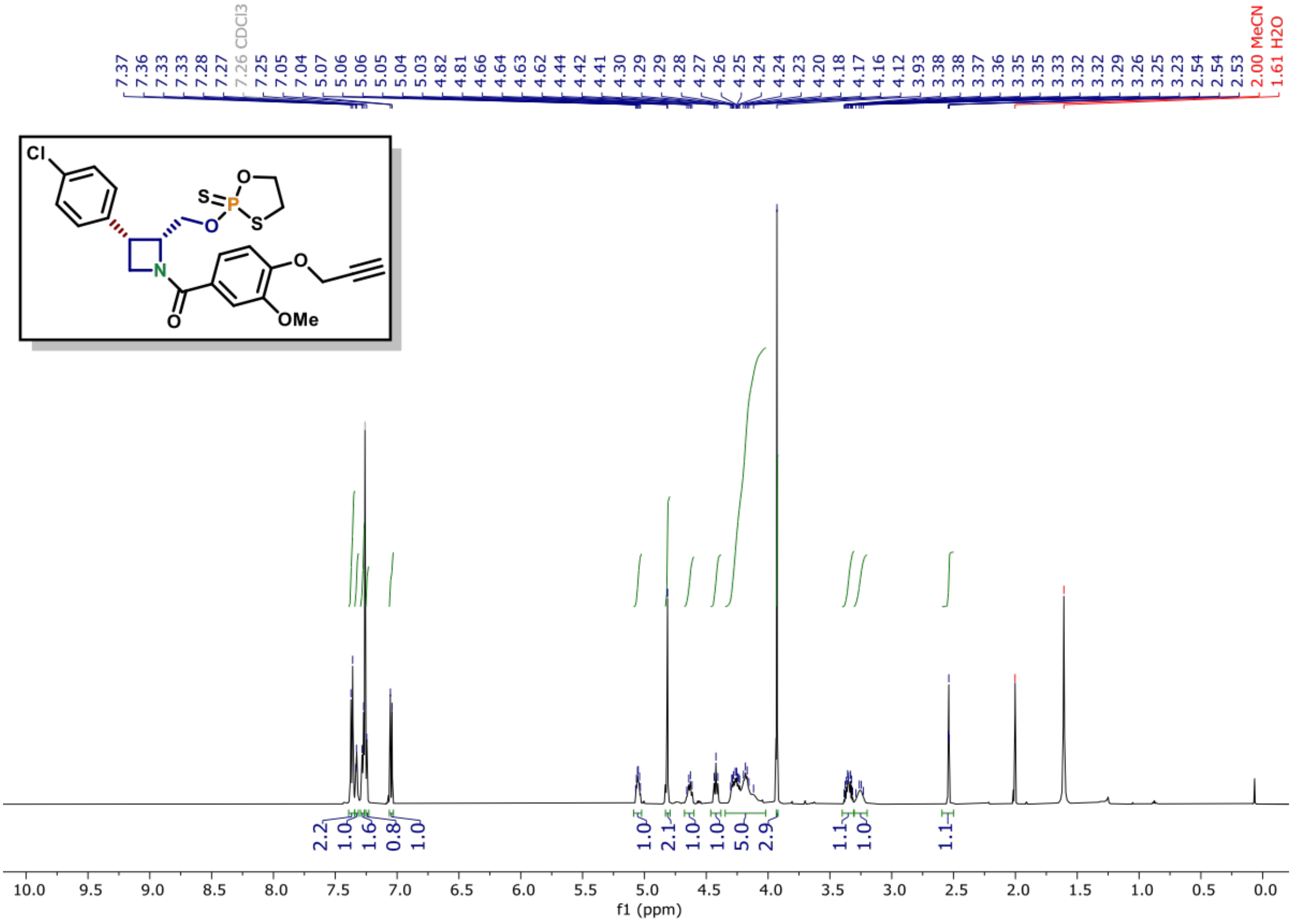

[13]C NMR of compound (10) (150 MHz, $CDCl_3$)

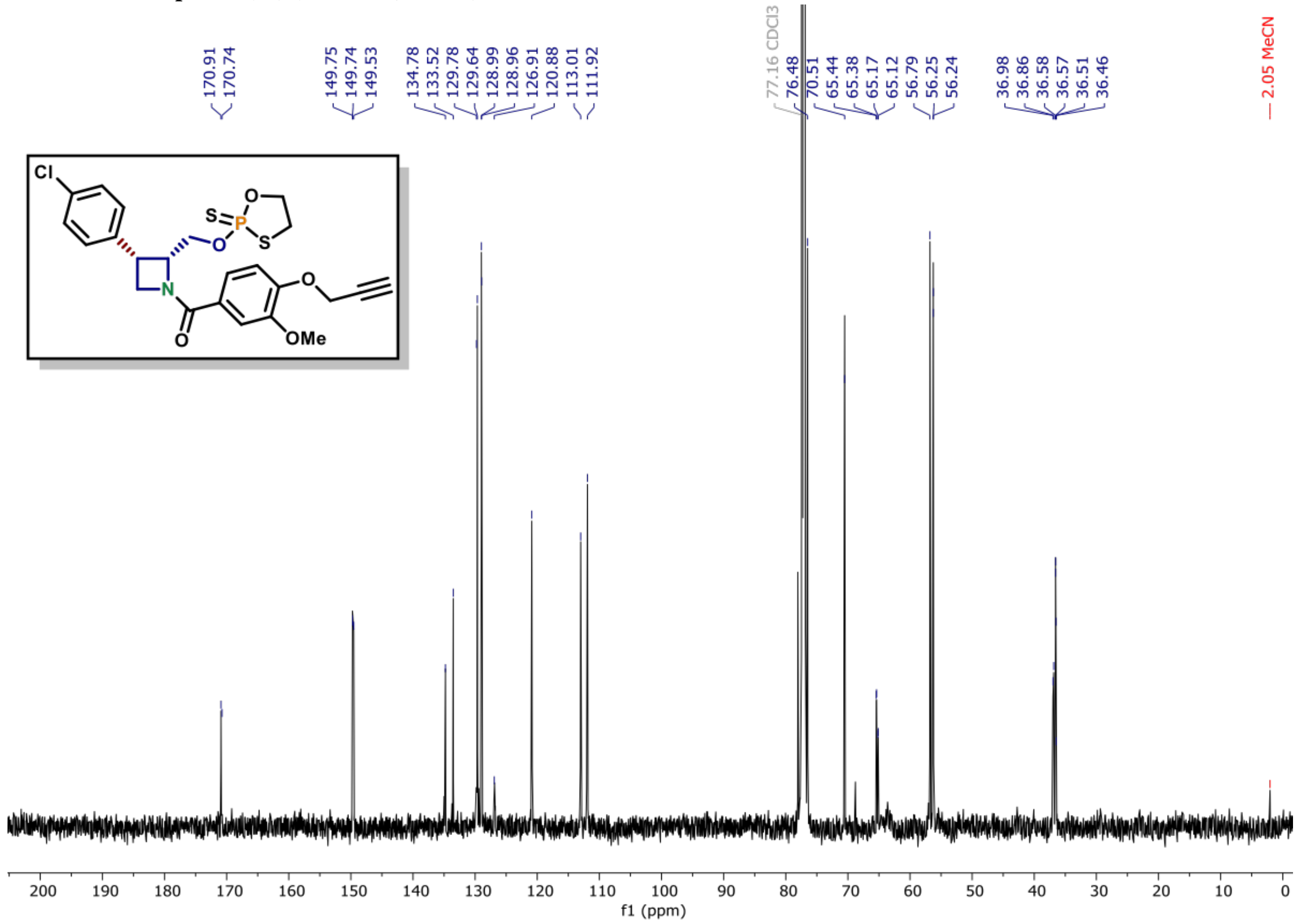

**[1]H NMR of compound (12) (600 MHz, $CDCl_3$)**

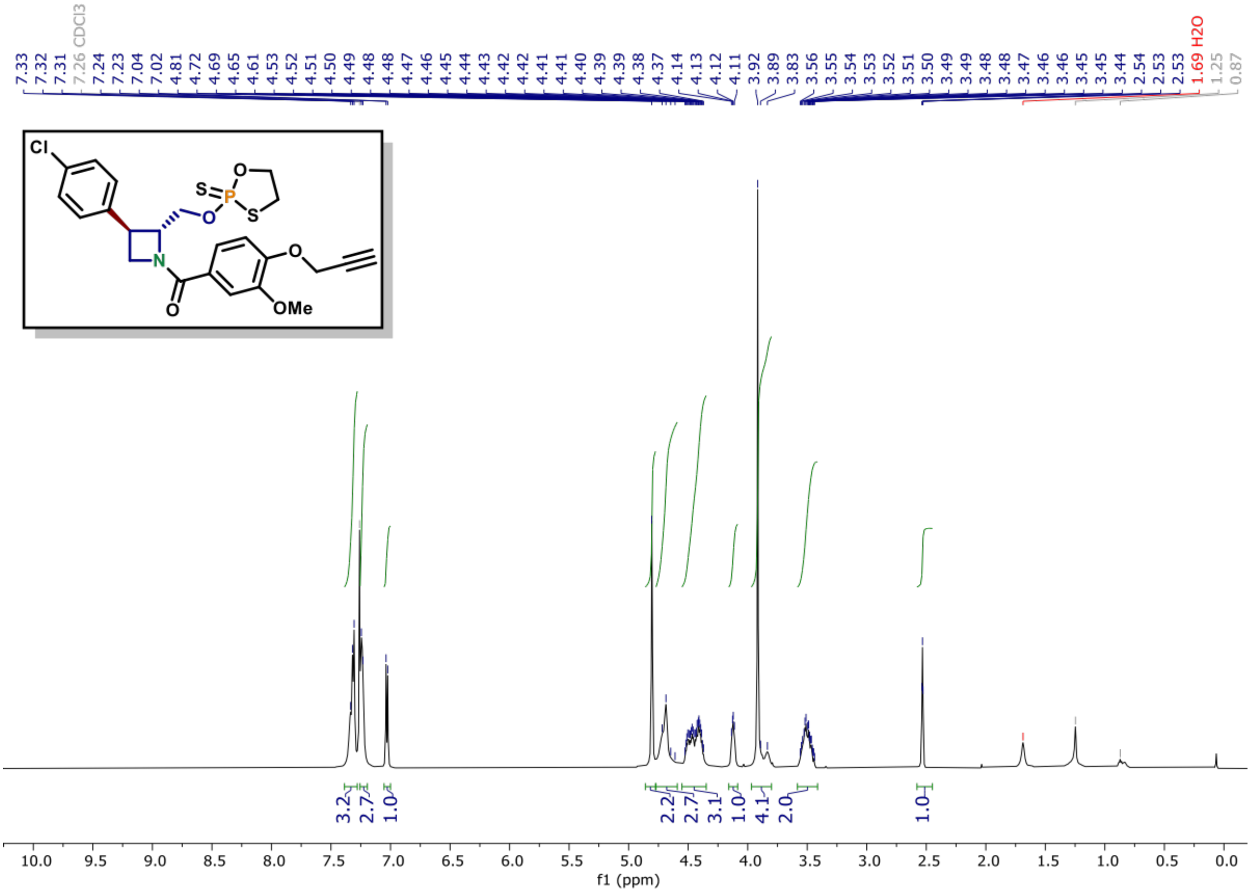

**[13]C NMR of compound (12) (150 MHz, $CDCl_3$)**

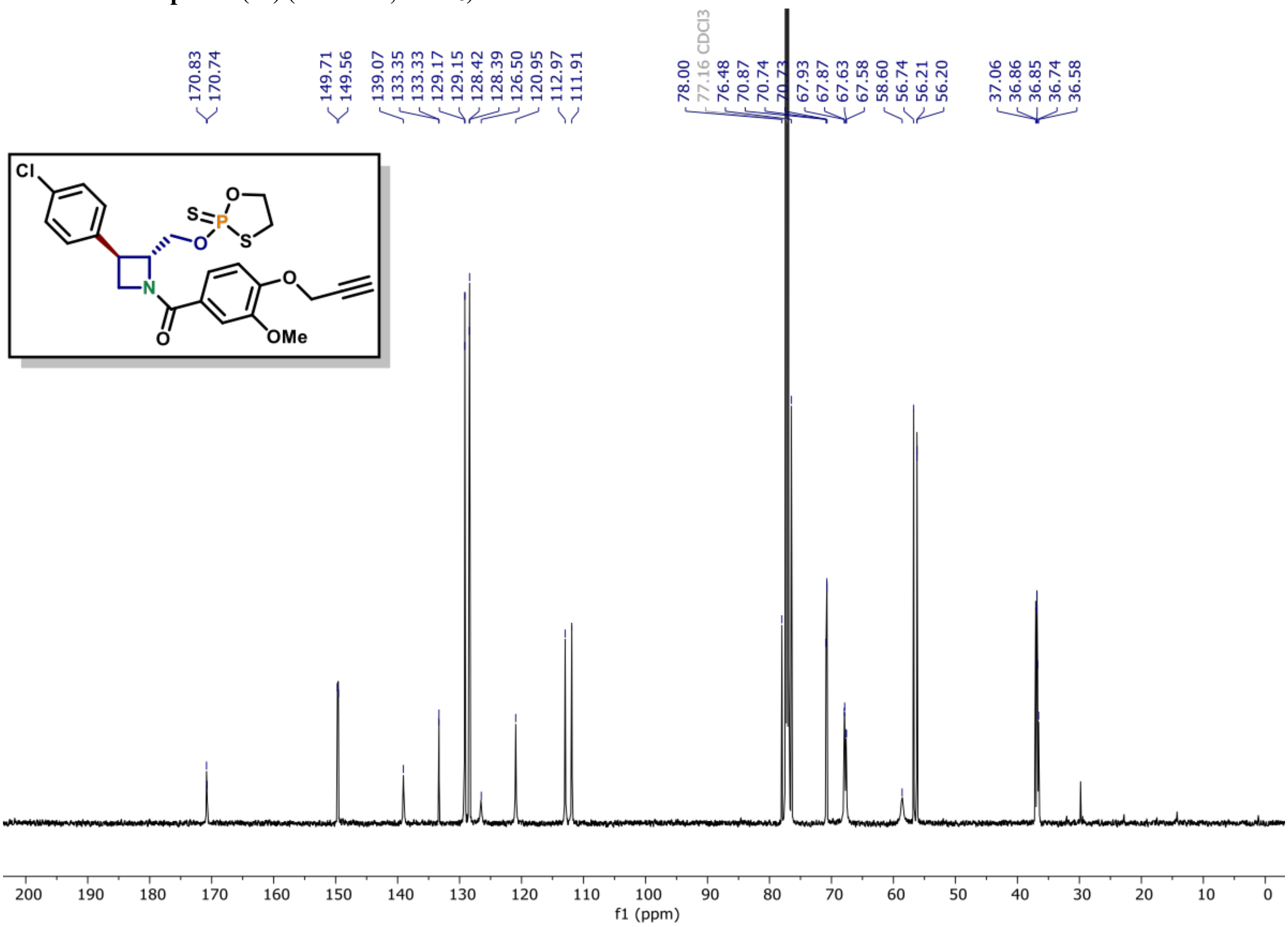

**[31]P NMR of compound (13) (162 MHz, $CDCl_3$)**

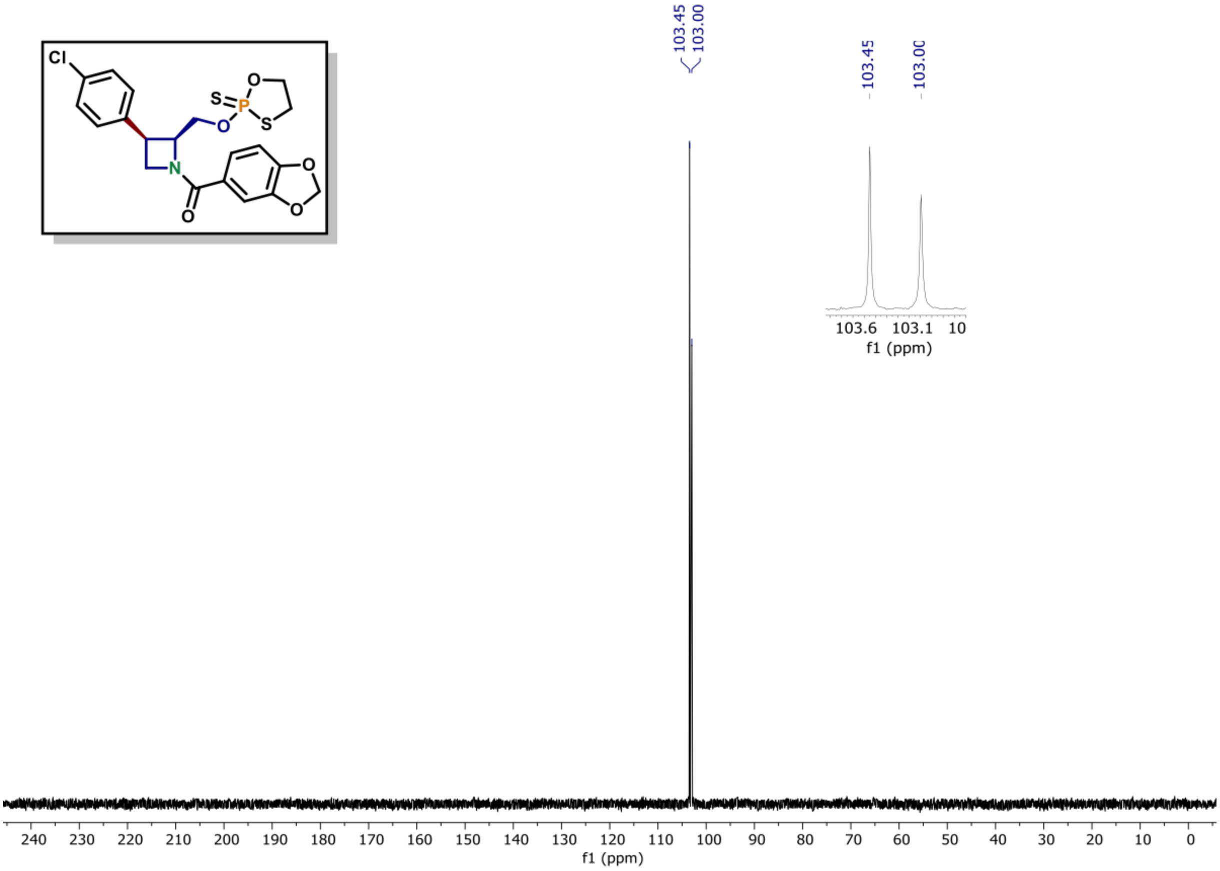

**[1]H NMR of compound (13) (600 MHz, $CDCl_3$)**

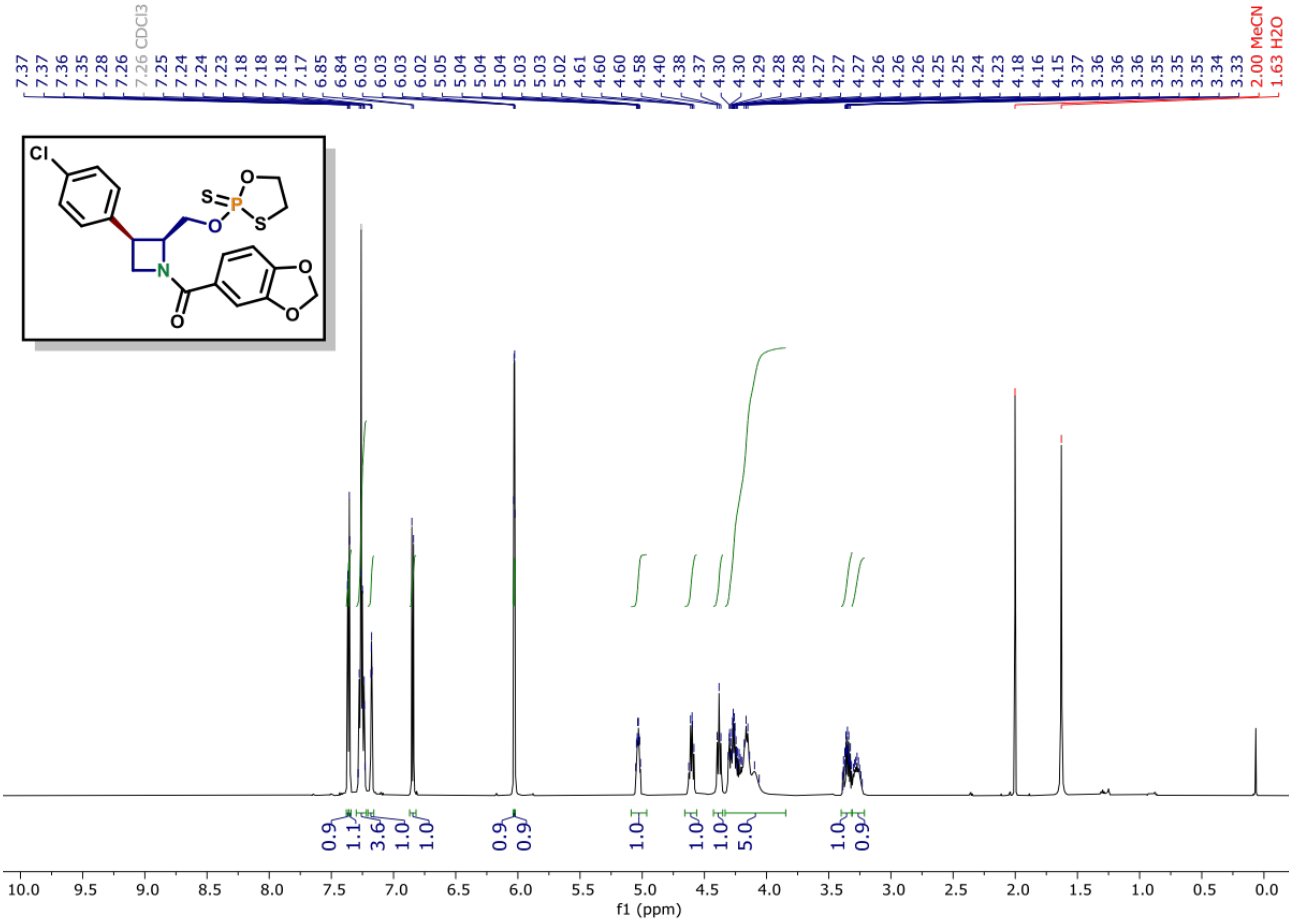

[13]C NMR of compound (13) (150 MHz, CDCl3)

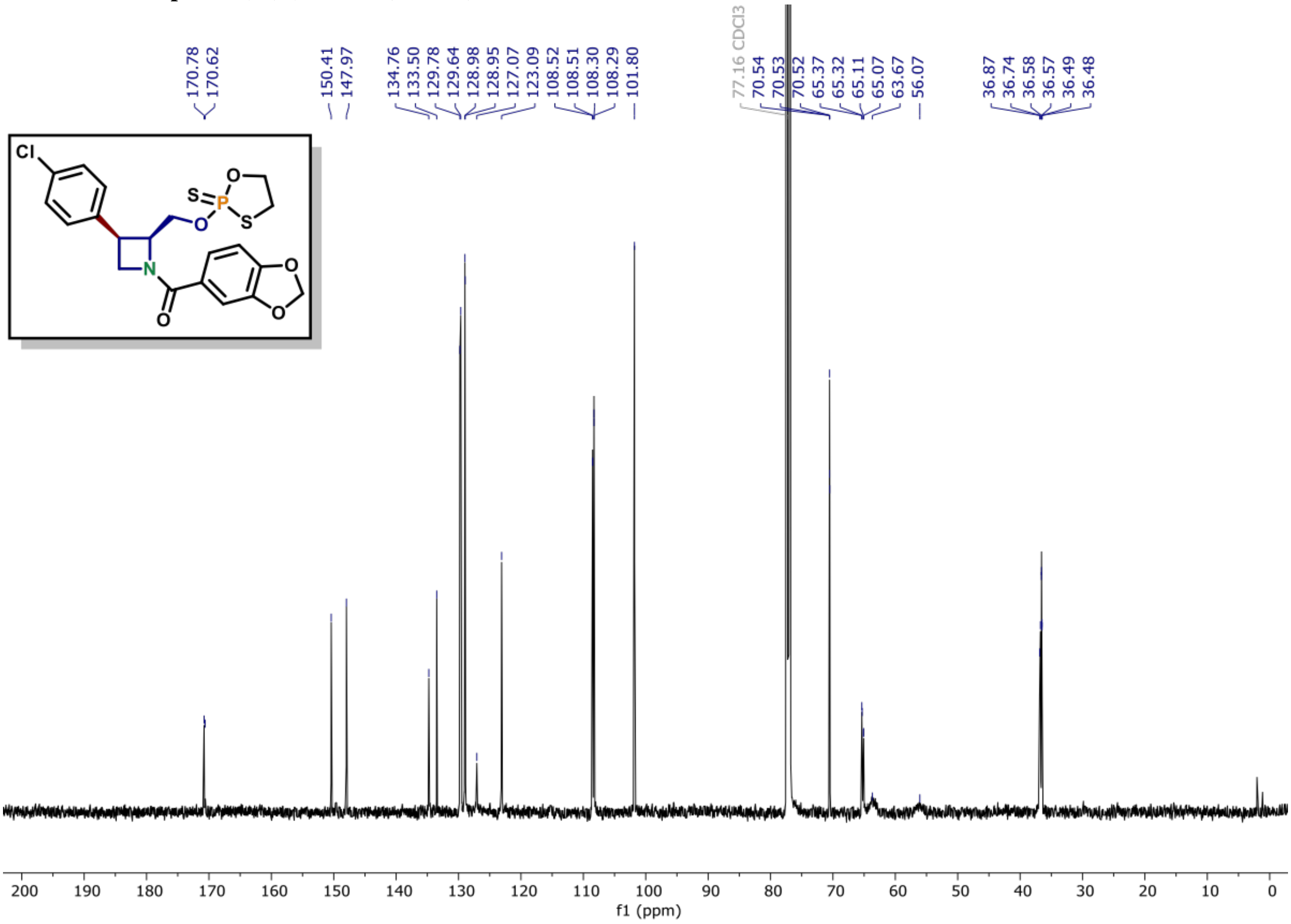

**[31]P NMR of compound (15) (162 MHz, CDCl₃)**

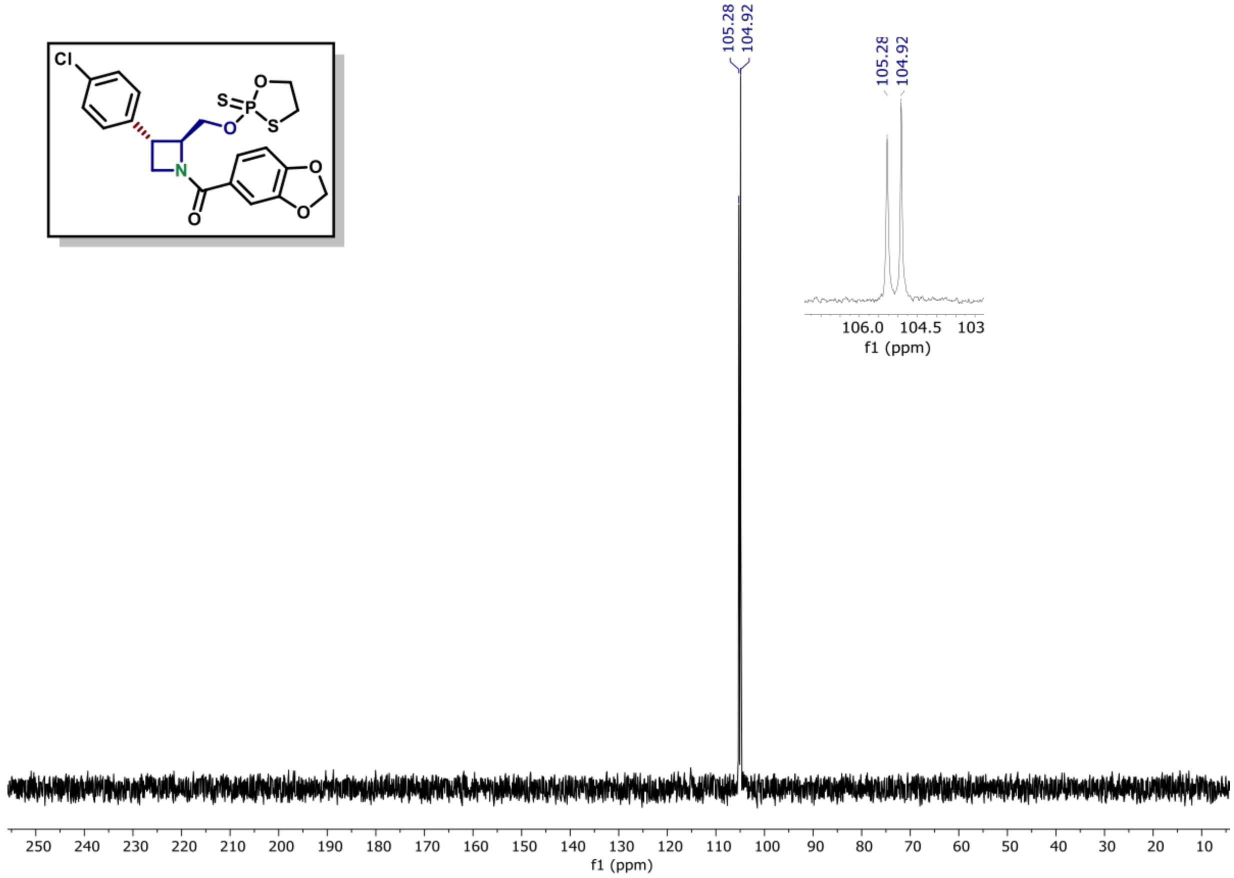

**[1]H NMR of compound (15) (600 MHz, CDCl₃)**

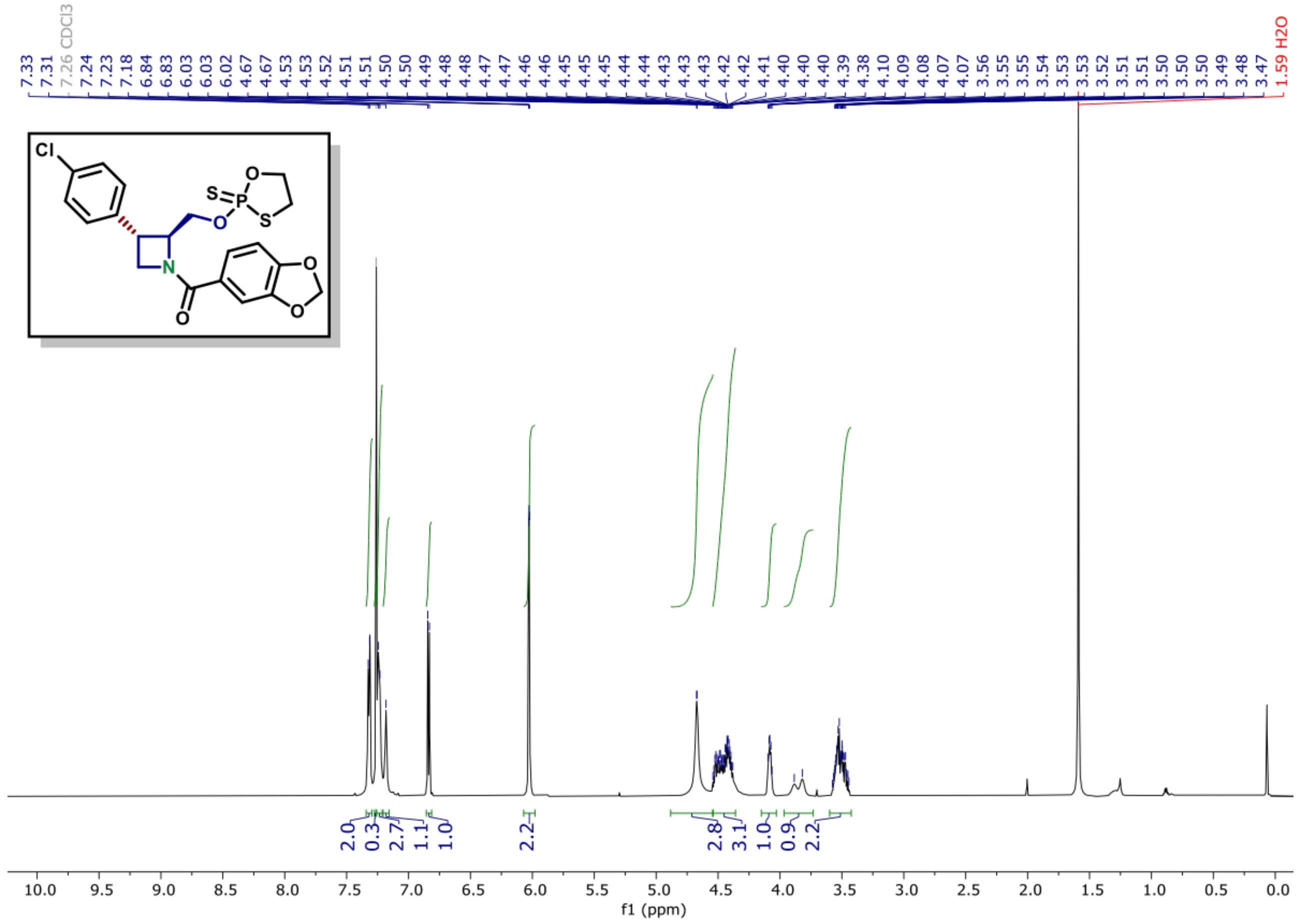

**$^{13}$C NMR of compound (15) (150 MHz, CDCl$_3$)**

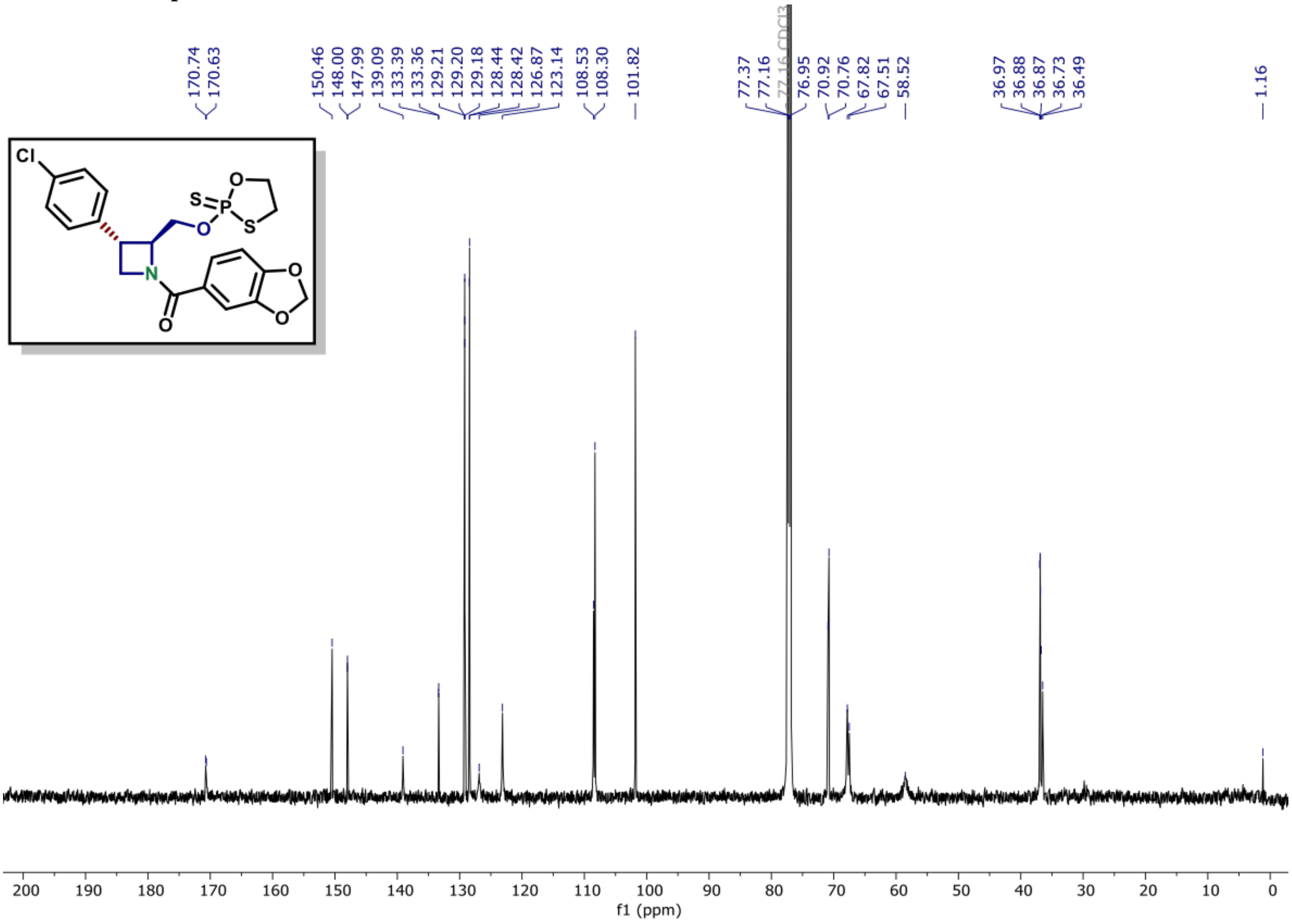

**[31]P NMR of compound (14) (162 MHz, CDCl₃)**

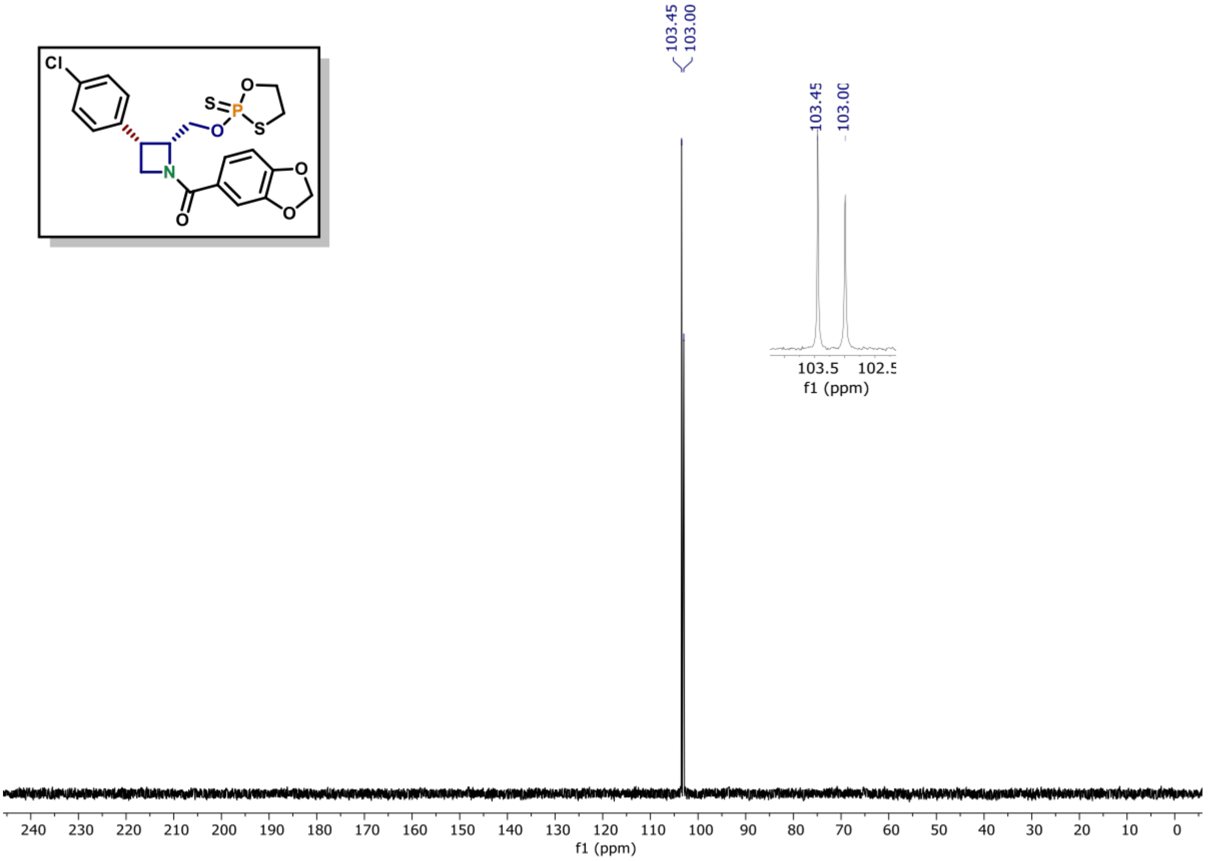

**[1]H NMR of compound (14) (600 MHz, CDCl₃)**

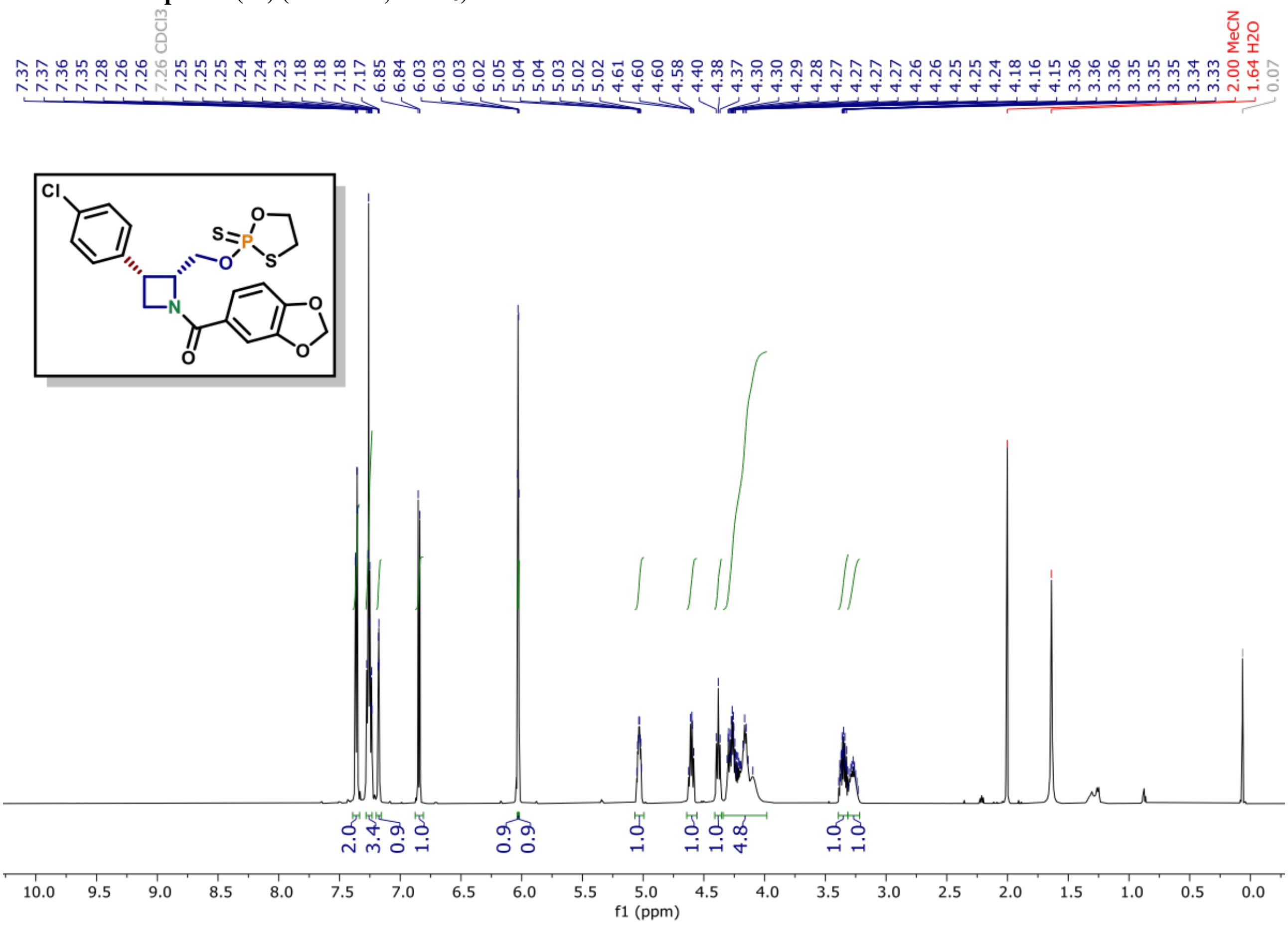

[13]C NMR of compound (14) (150 MHz, CDCl₃)

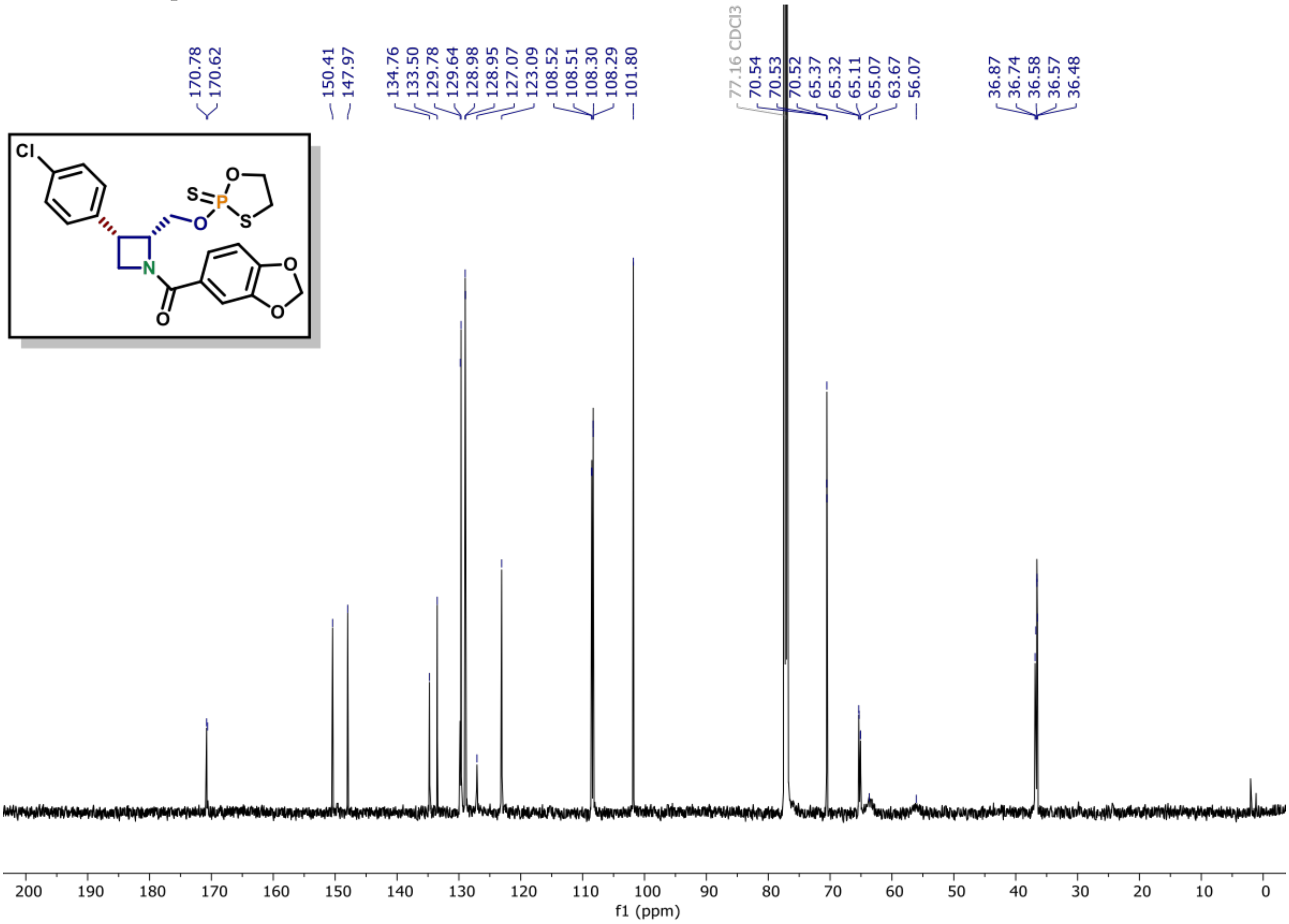

[31]P NMR of compound (16) (162 MHz, $CDCl_3$)

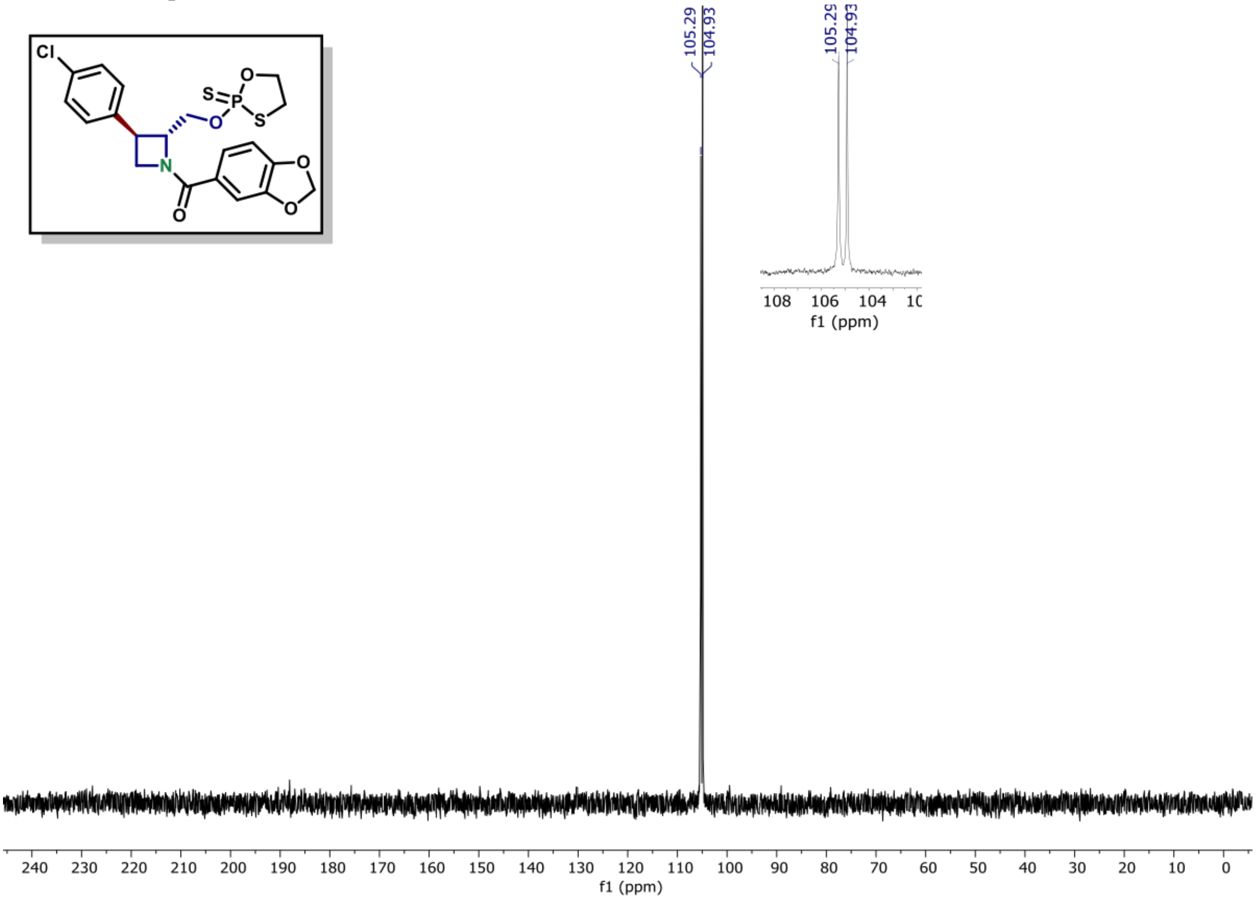

[1]H NMR of compound (16) (600 MHz, $CDCl_3$)

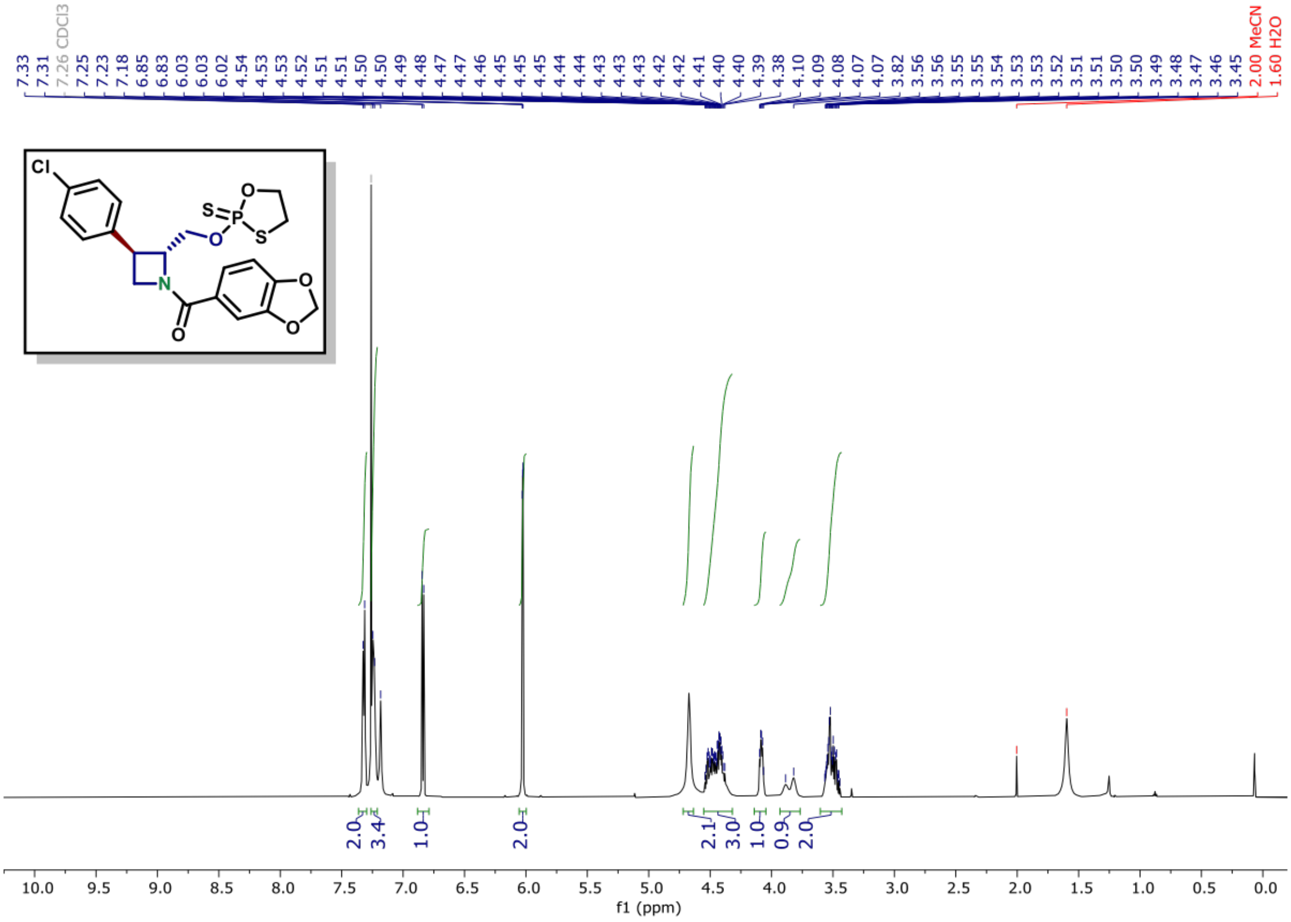

**[13]C NMR of compound (16) (150 MHz, CDCl₃)**

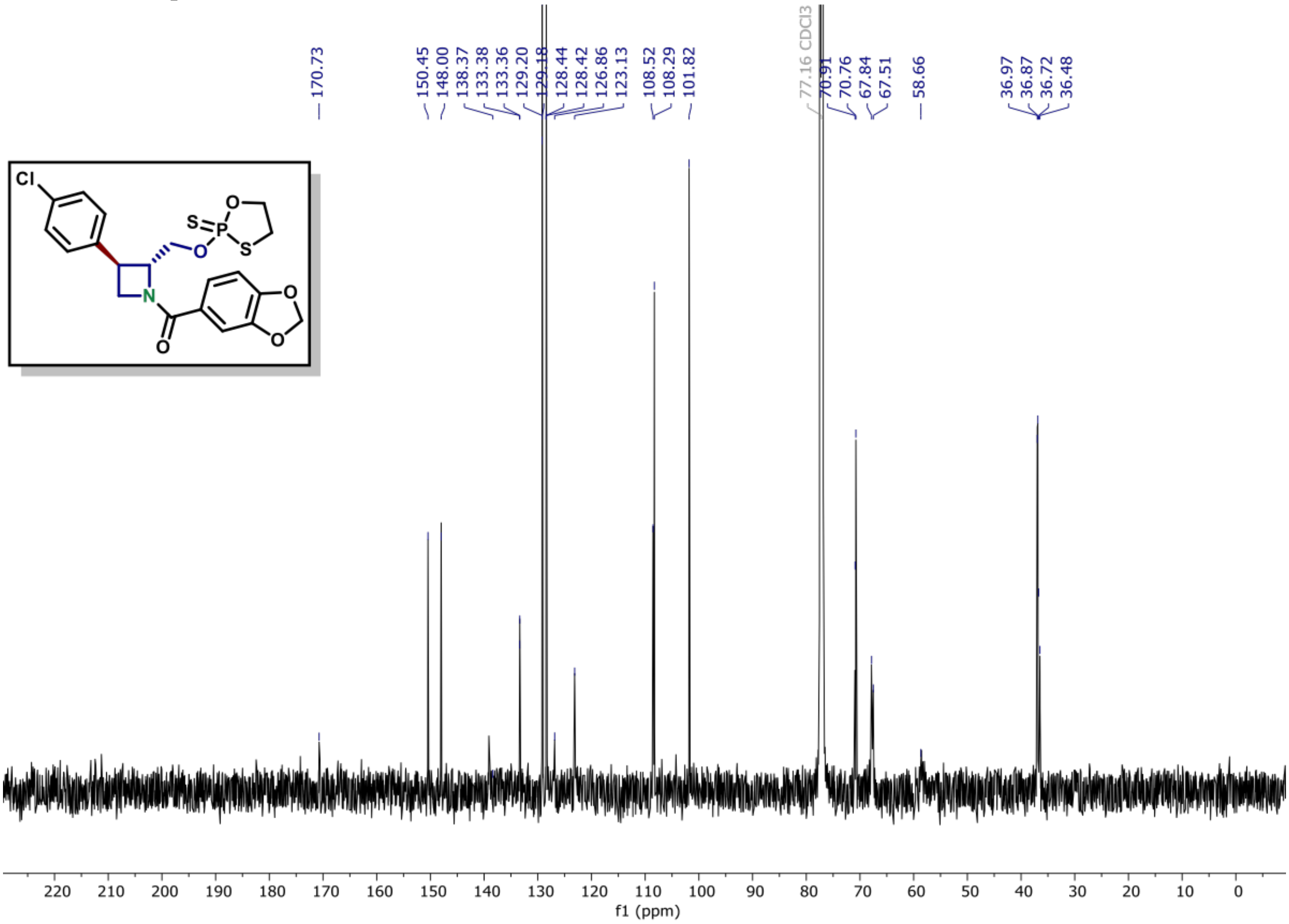

[1]H NMR of compound (S9A) (600 MHz, $CDCl_3$)

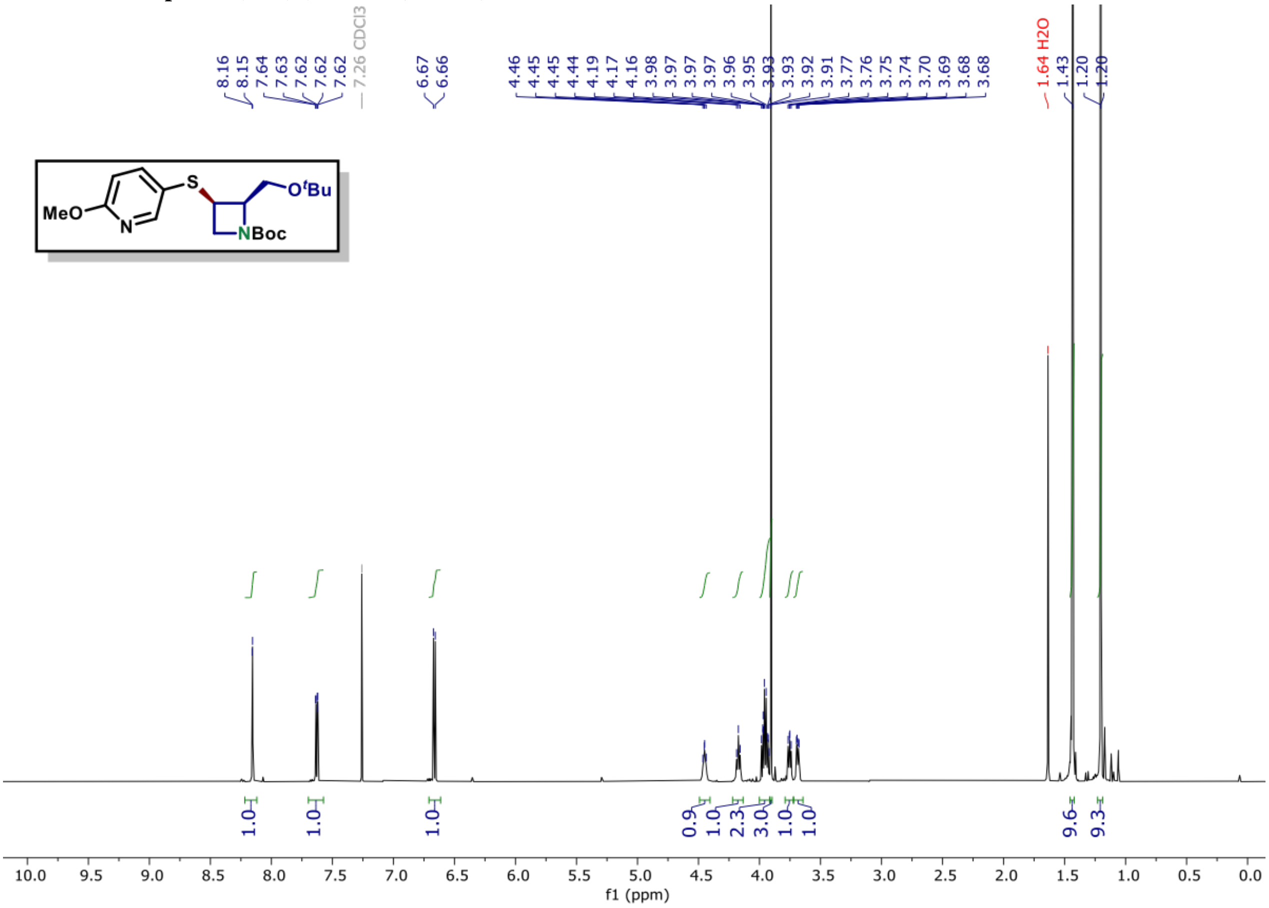

[13]C NMR of compound (S9A) (150 MHz, $CDCl_3$)

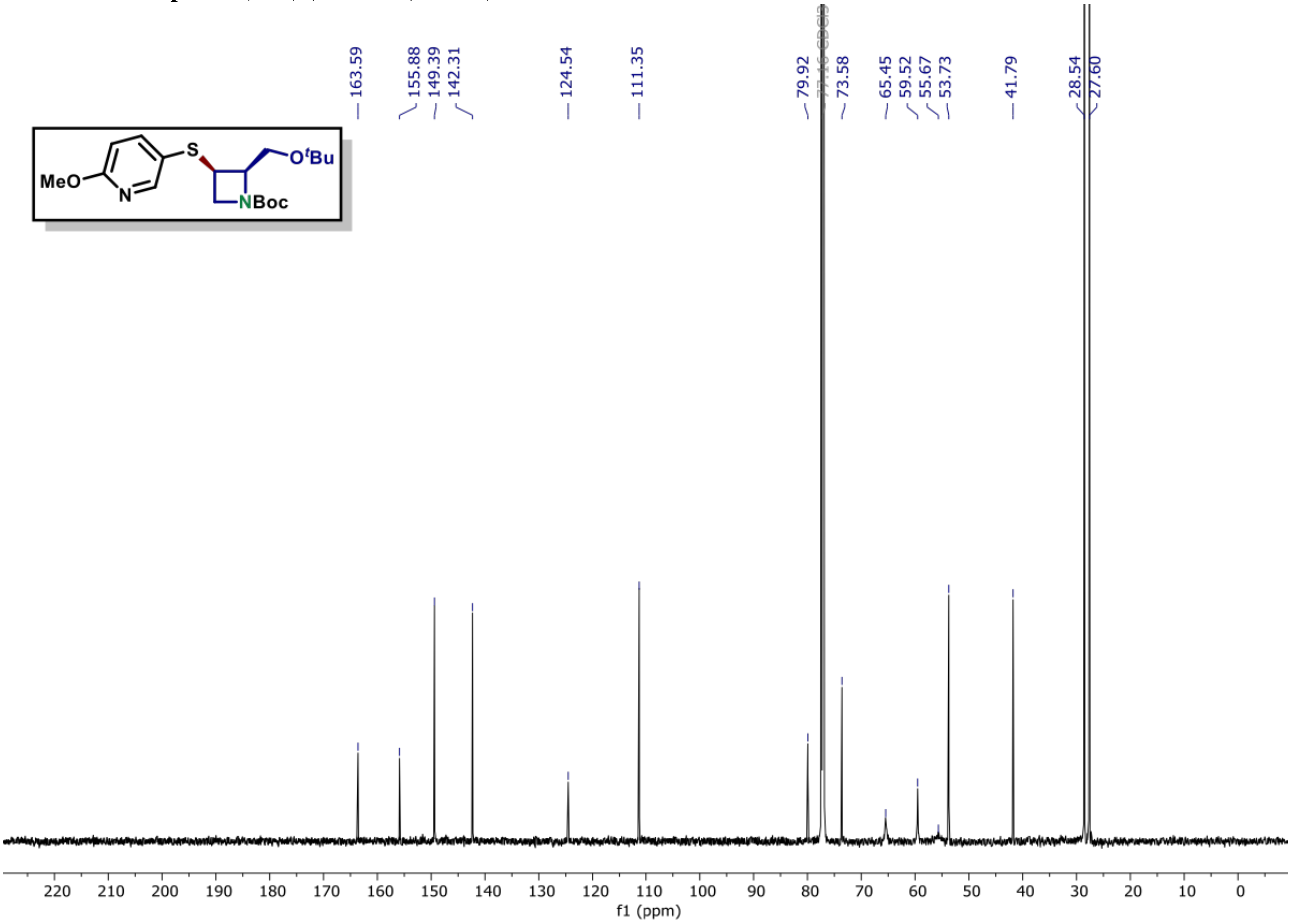

[1]H NMR of compound (S9B) (600 MHz, $CDCl_3$)

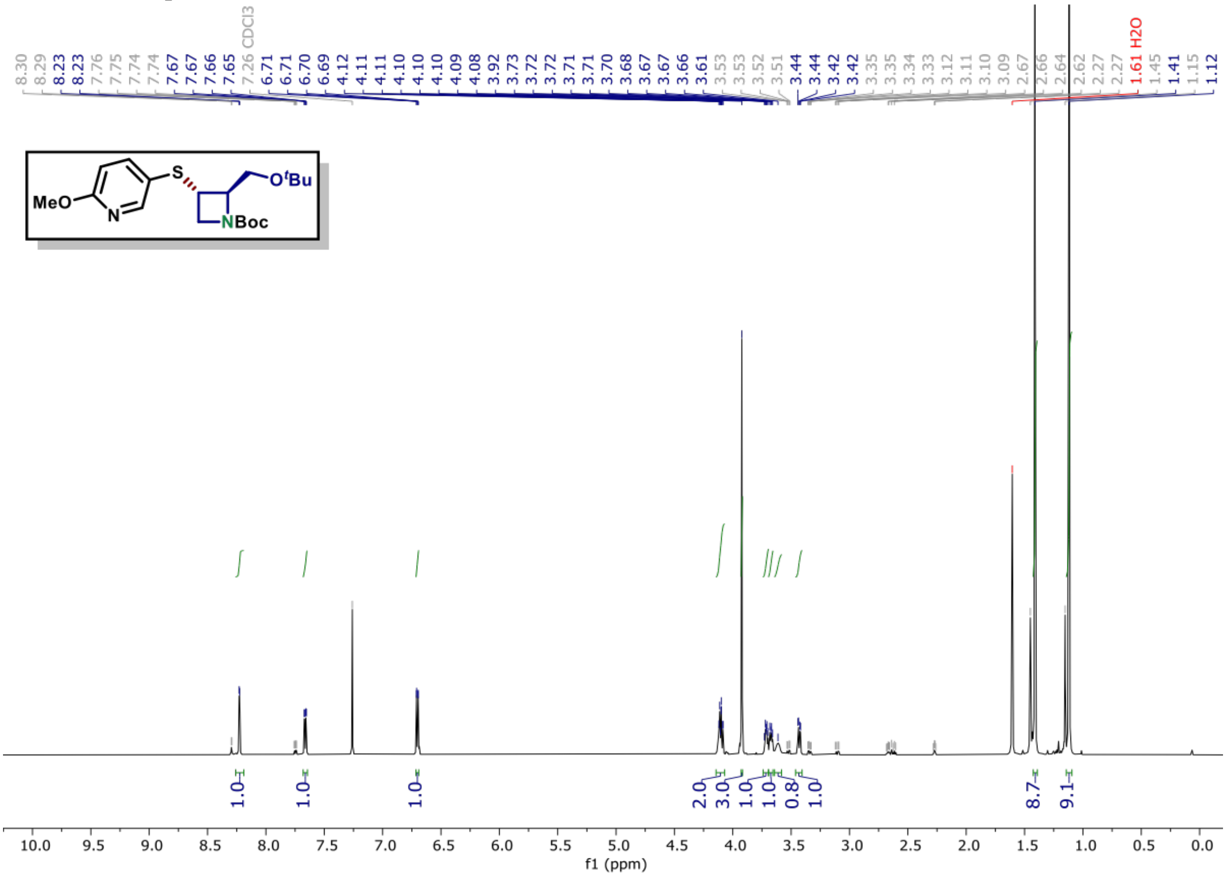

[13]C NMR of compound (S9B) (150 MHz, $CDCl_3$)

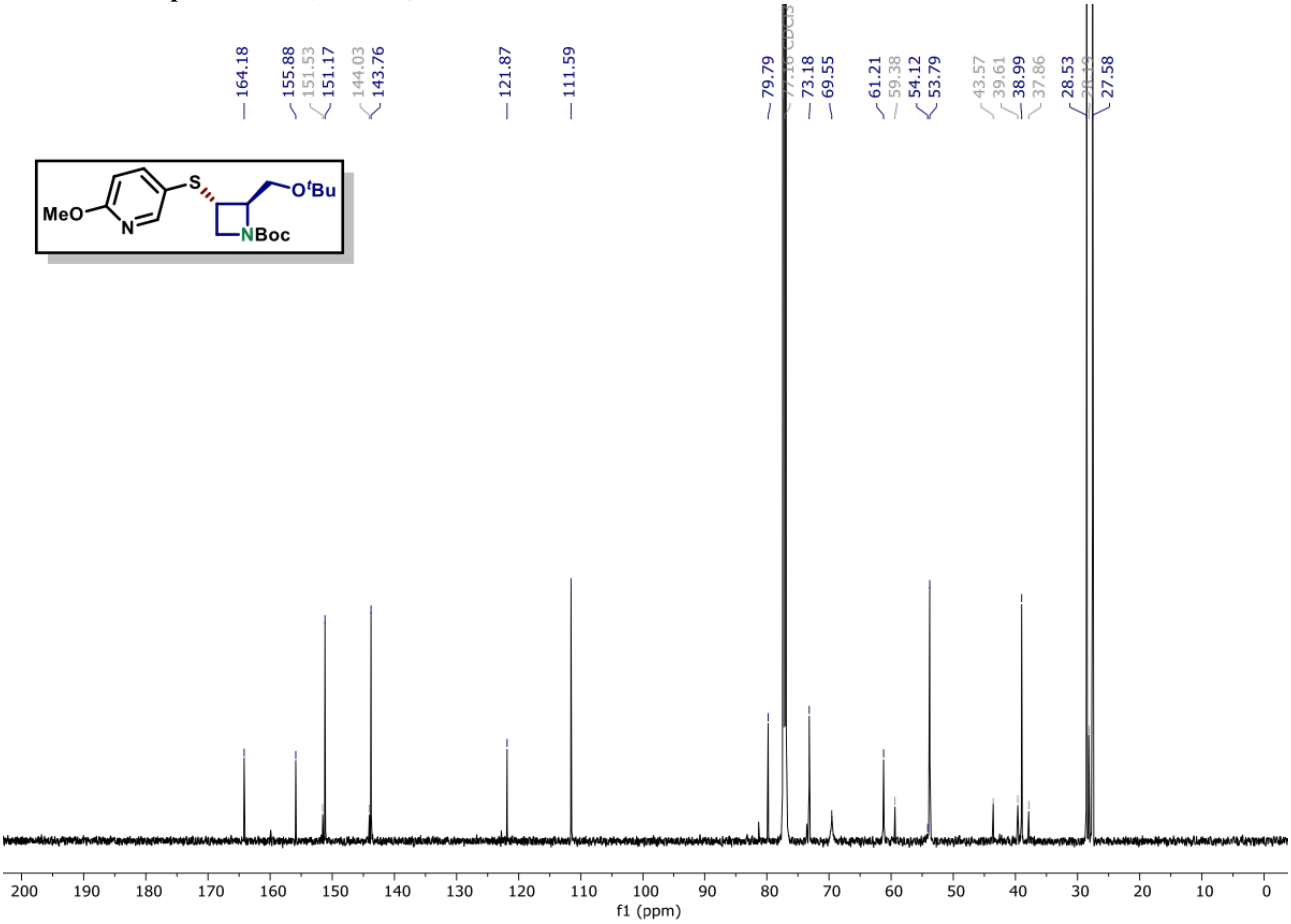

**[1]H NMR of compound (S9C) (600 MHz, CDCl$_3$)**

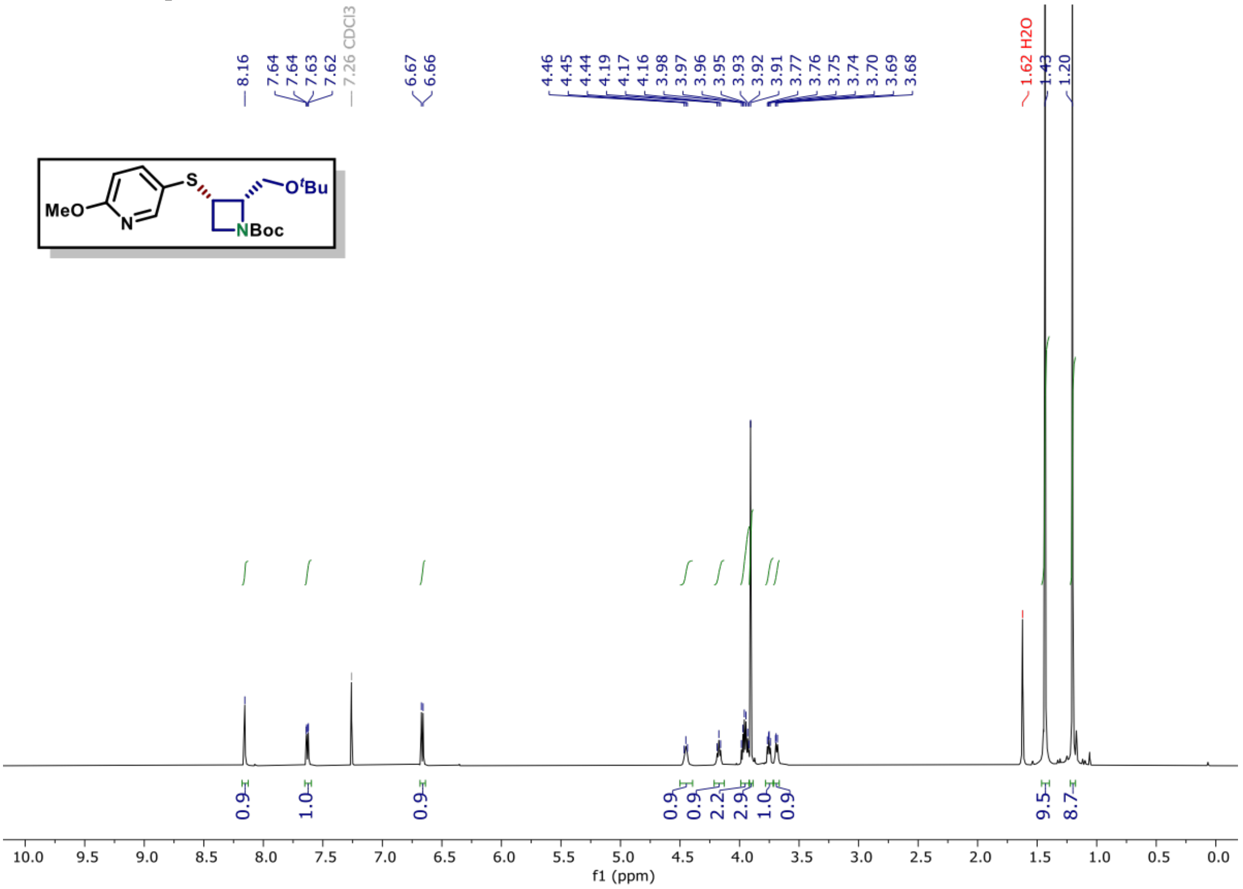

**[13]C NMR of compound (S9C) (150 MHz, CDCl$_3$)**

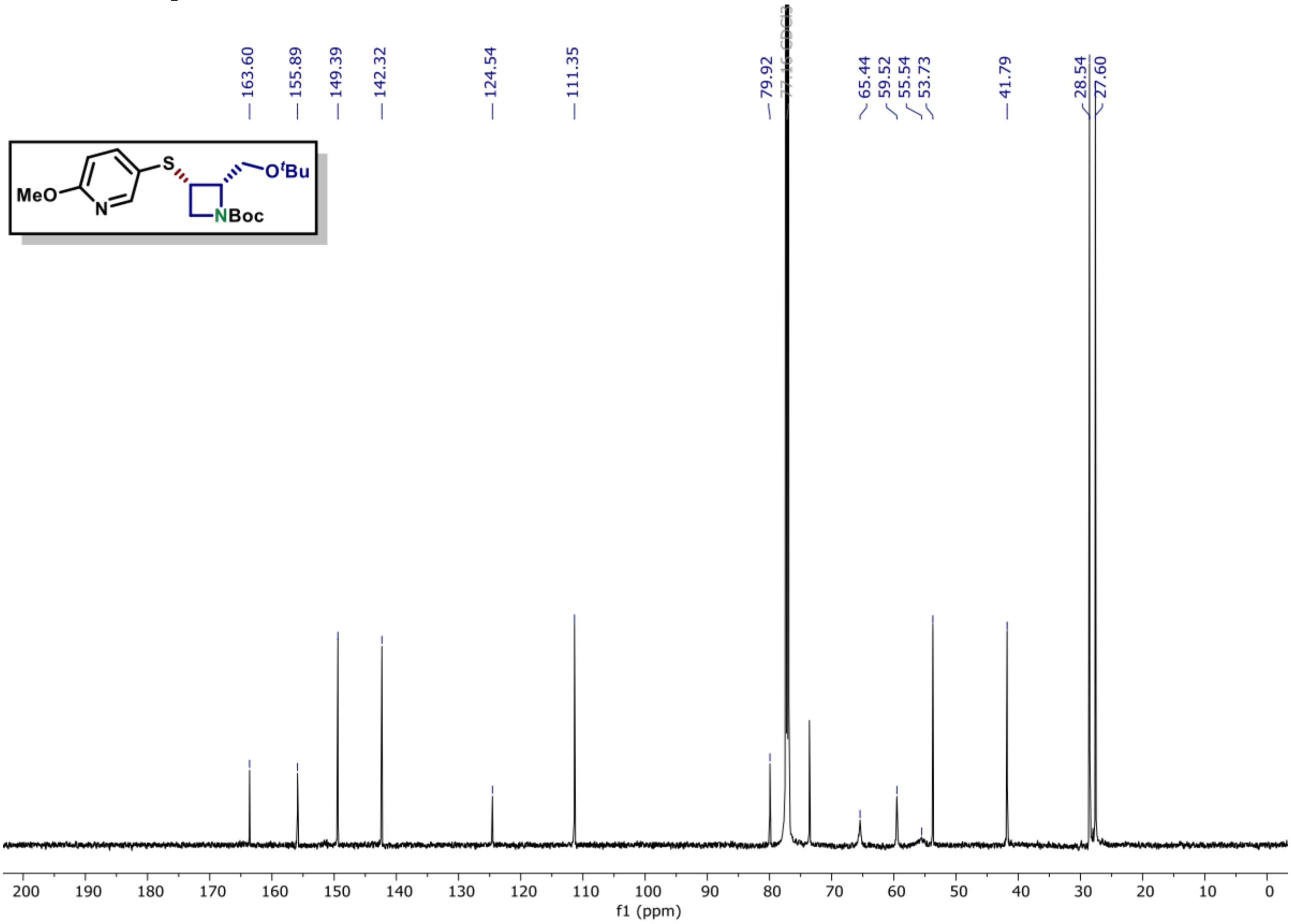

$^{1}H$ NMR of compound (S9D) (600 MHz, $CDCl_3$)

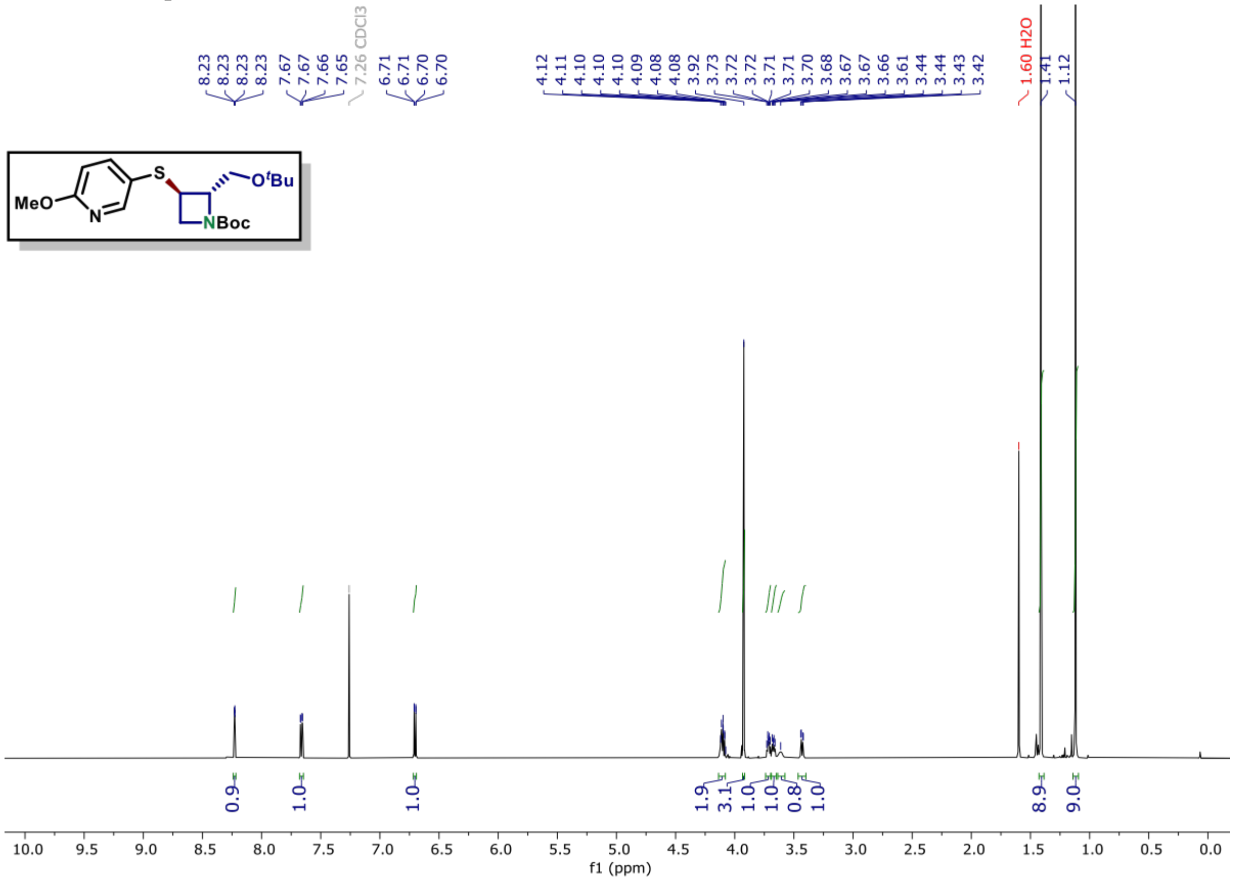

$^{13}C$ NMR of compound (S9D) (150 MHz, $CDCl_3$)

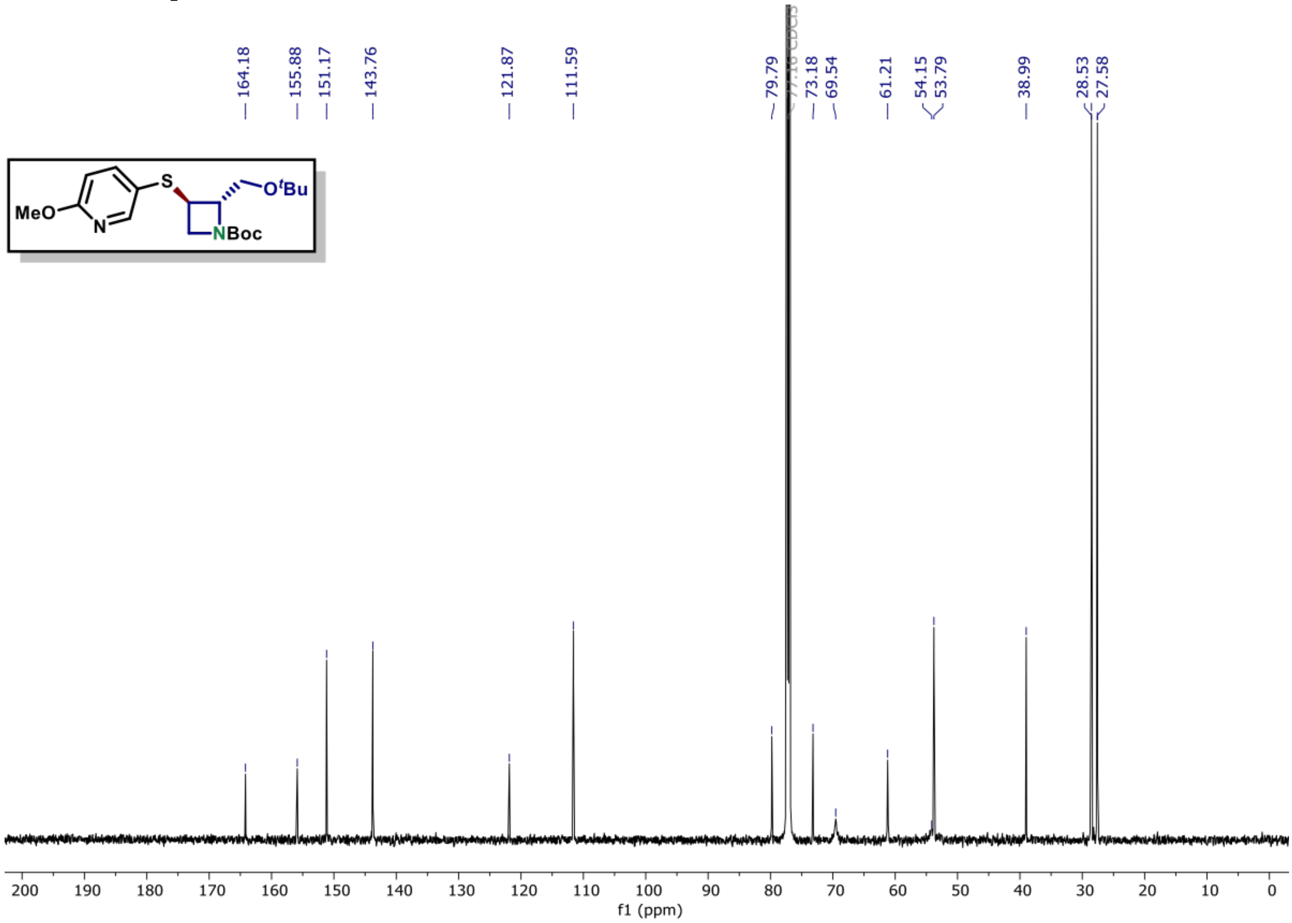

**$^{1}$H NMR of compound (S10A) (600 MHz, CDCl$_3$)**

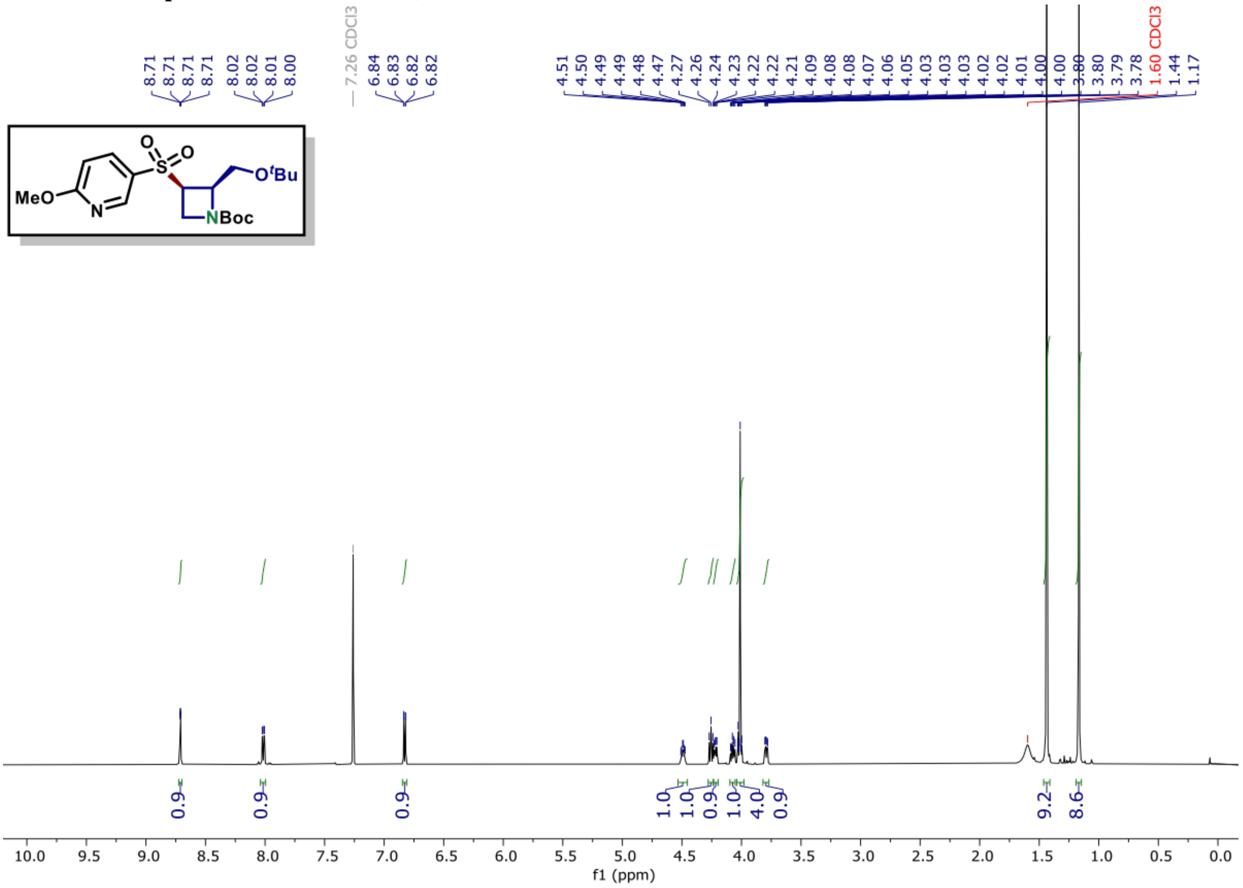

**$^{13}$C NMR of compound (S10A) (150 MHz, CDCl$_3$)**

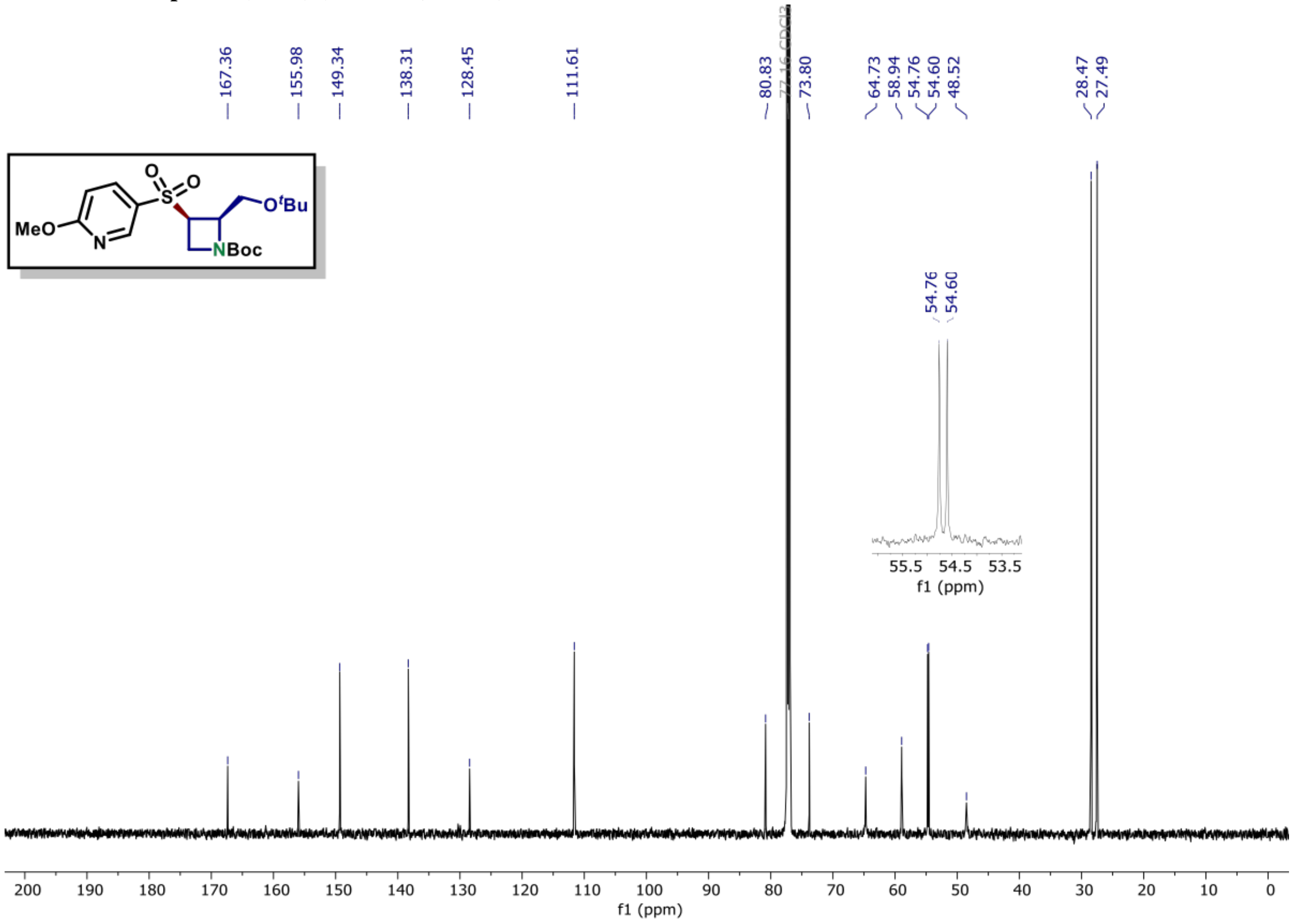

**[1]H NMR of compound (S10B) (600 MHz, CDCl₃)**

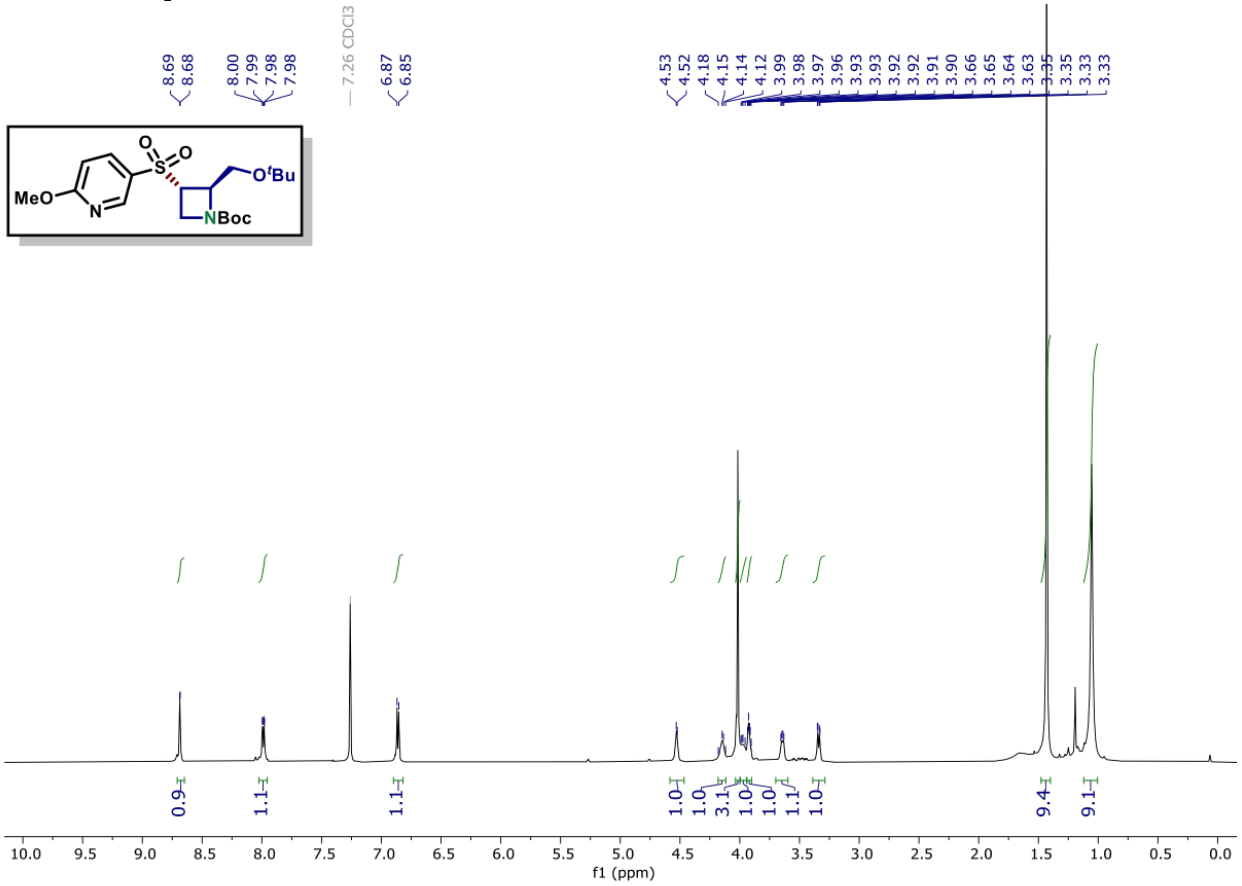

**[13]C NMR of compound (S10B) (150 MHz, CDCl₃)**

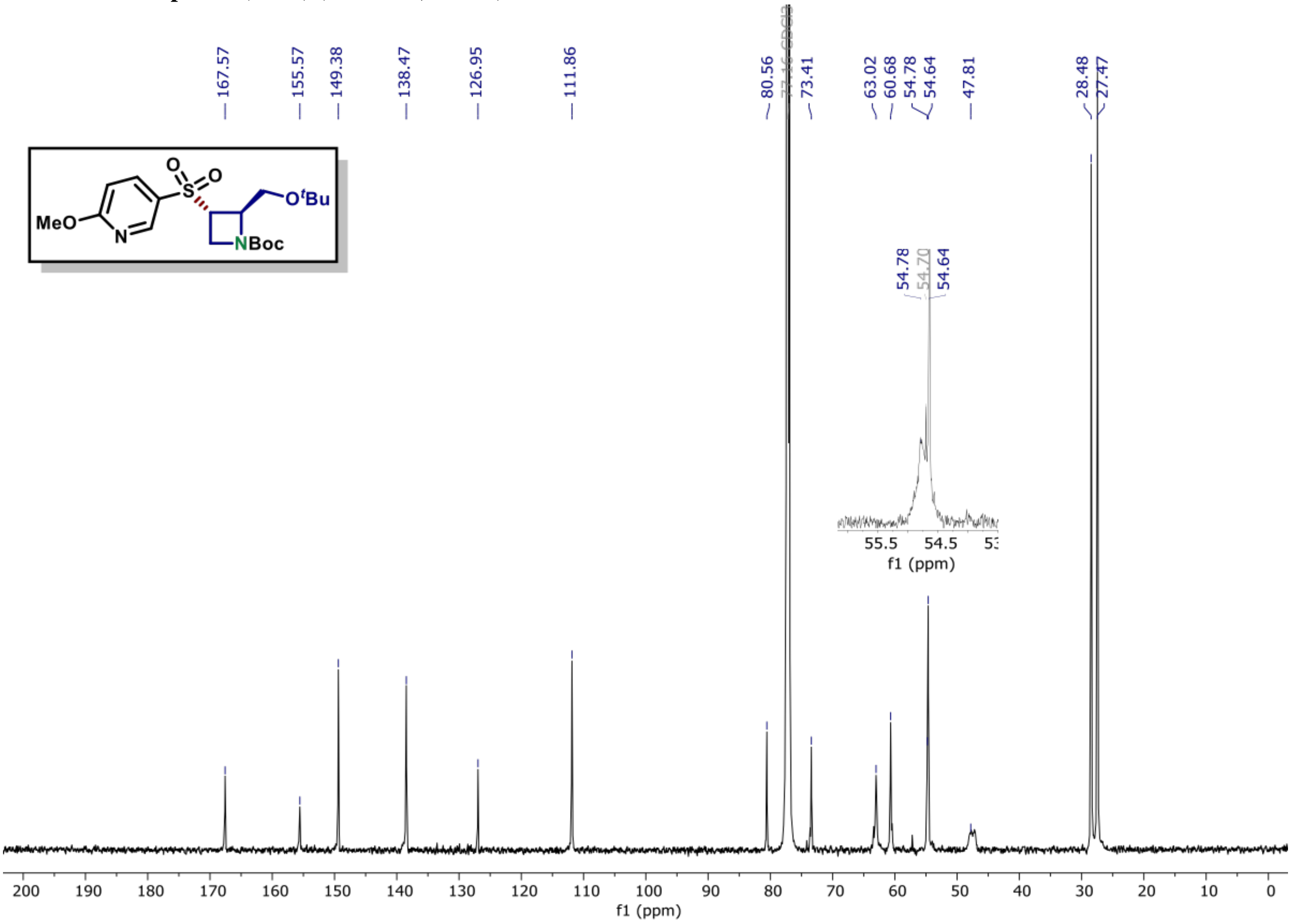

**$^{1}$H NMR of compound (S10C) (600 MHz, $CDCl_3$)**

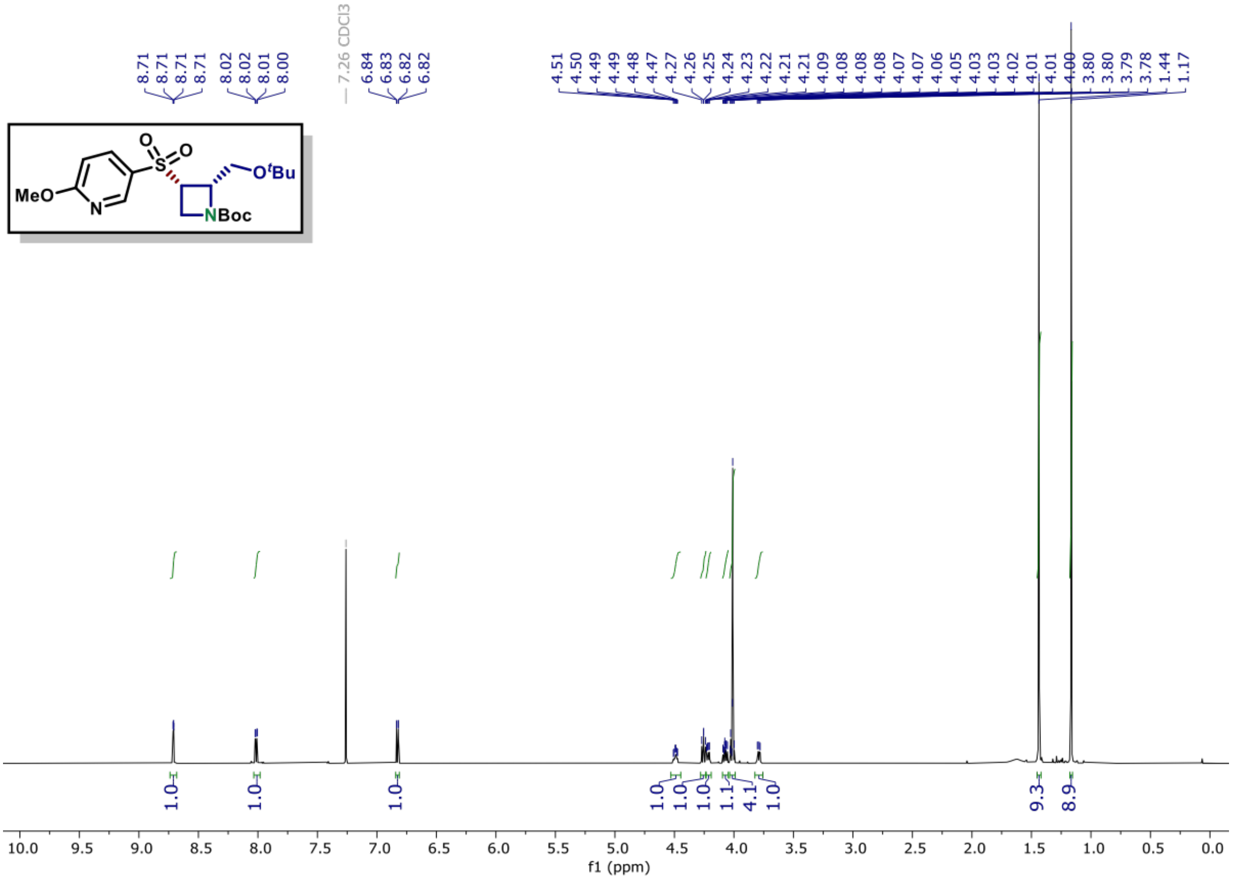

**$^{13}$C NMR of compound (S10C) (150 MHz, $CDCl_3$)**

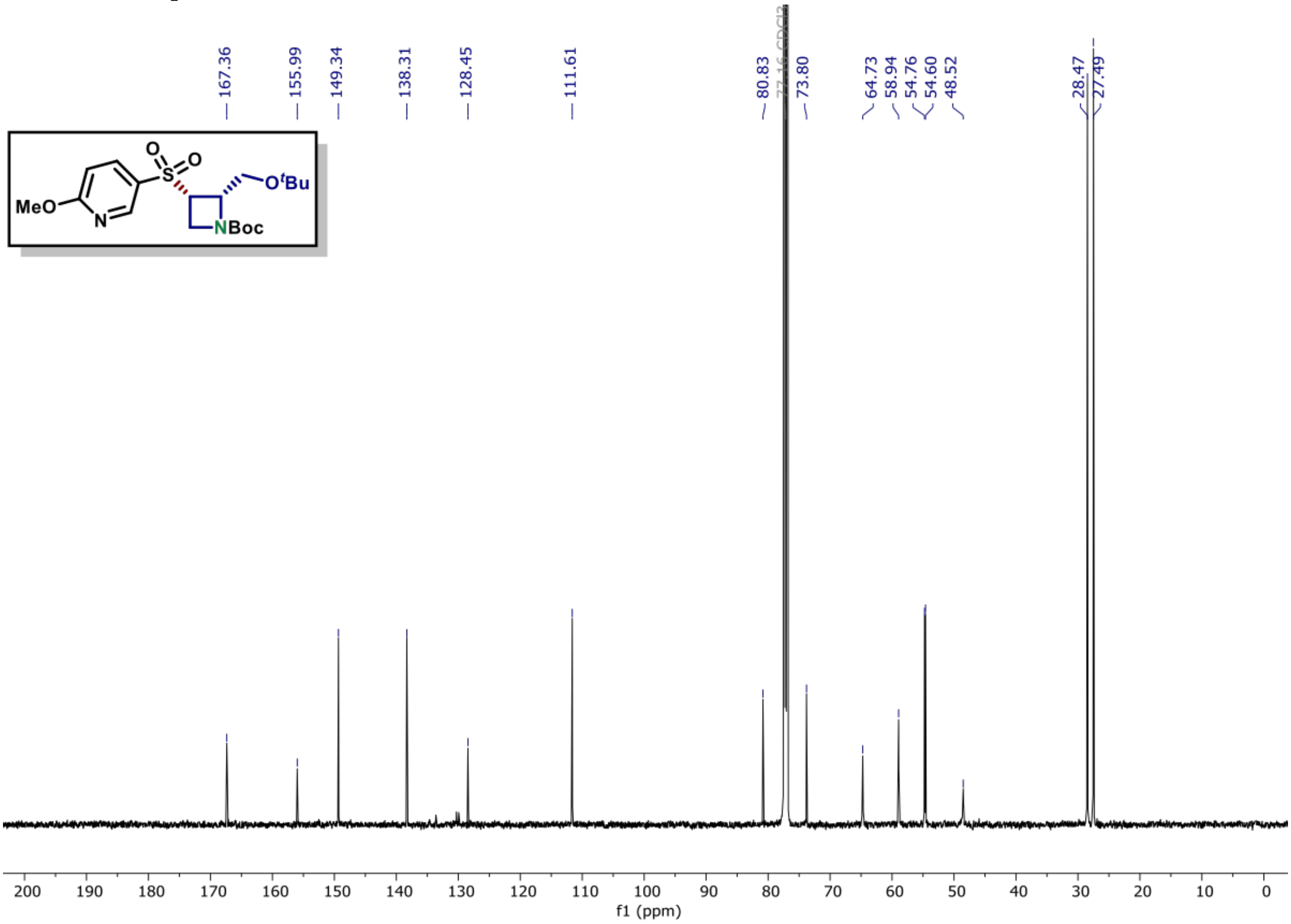

[1]H NMR of compound (S10D) (600 MHz, $CDCl_3$)

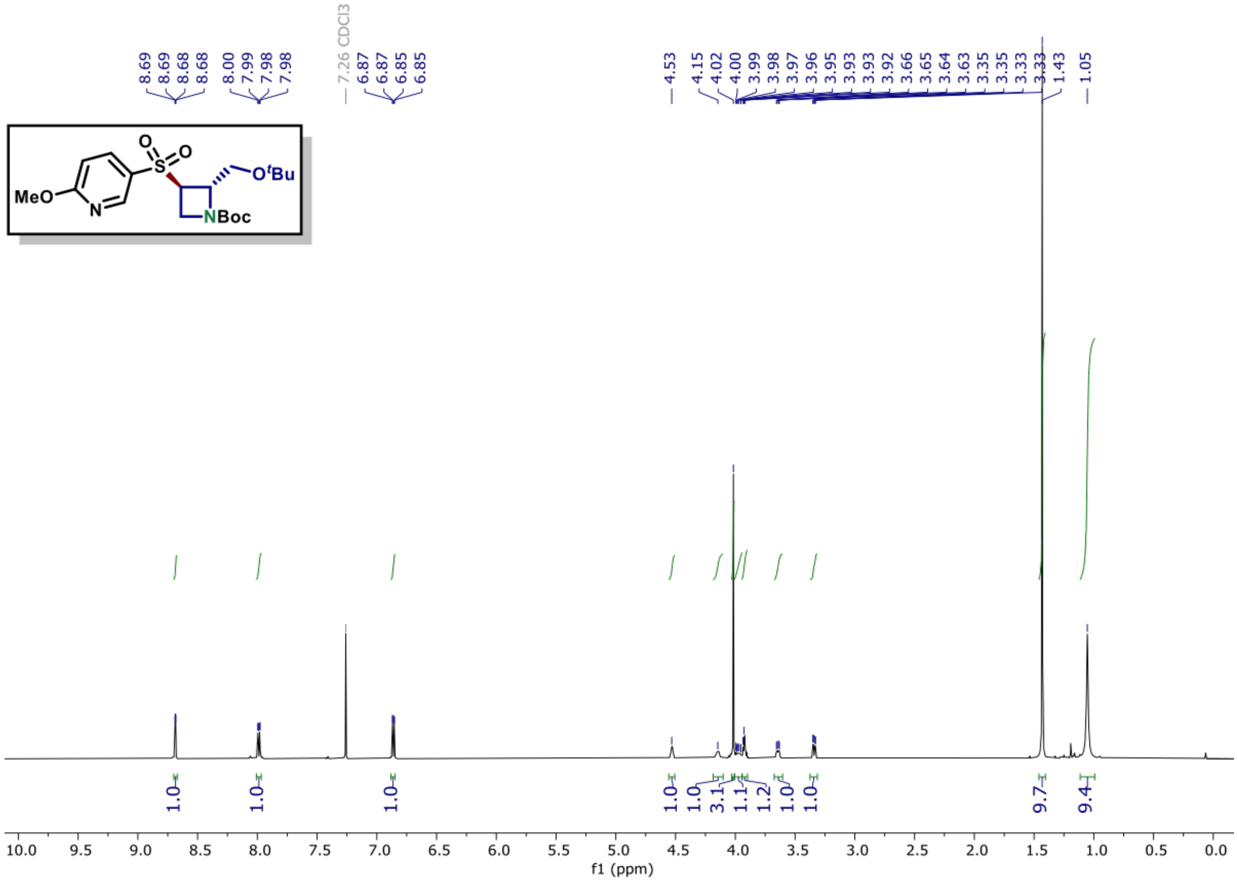

[13]C NMR of compound (S10D) (150 MHz, $CDCl_3$)

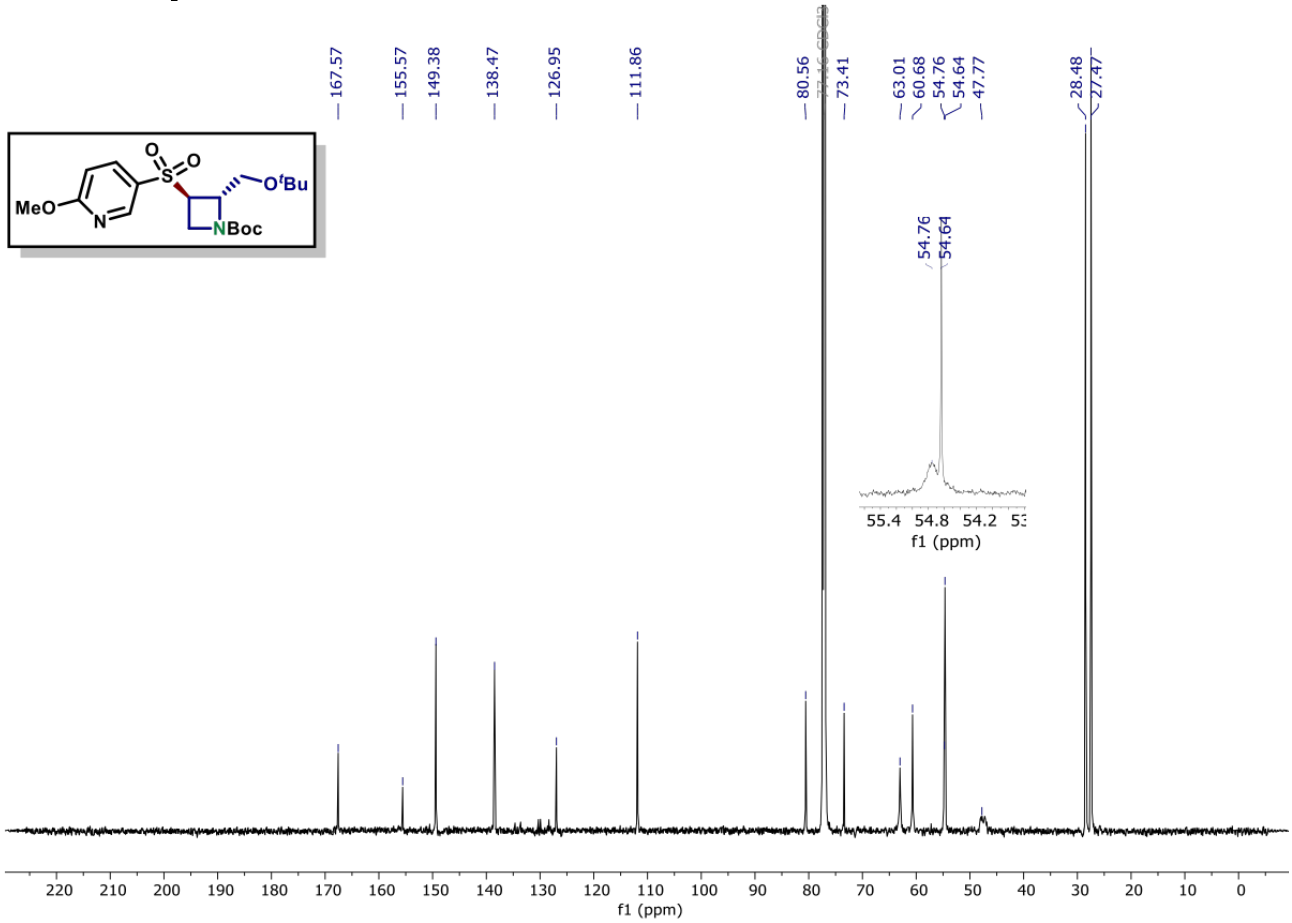

[1]H NMR of compound (11A) (600 MHz, $CDCl_3$)

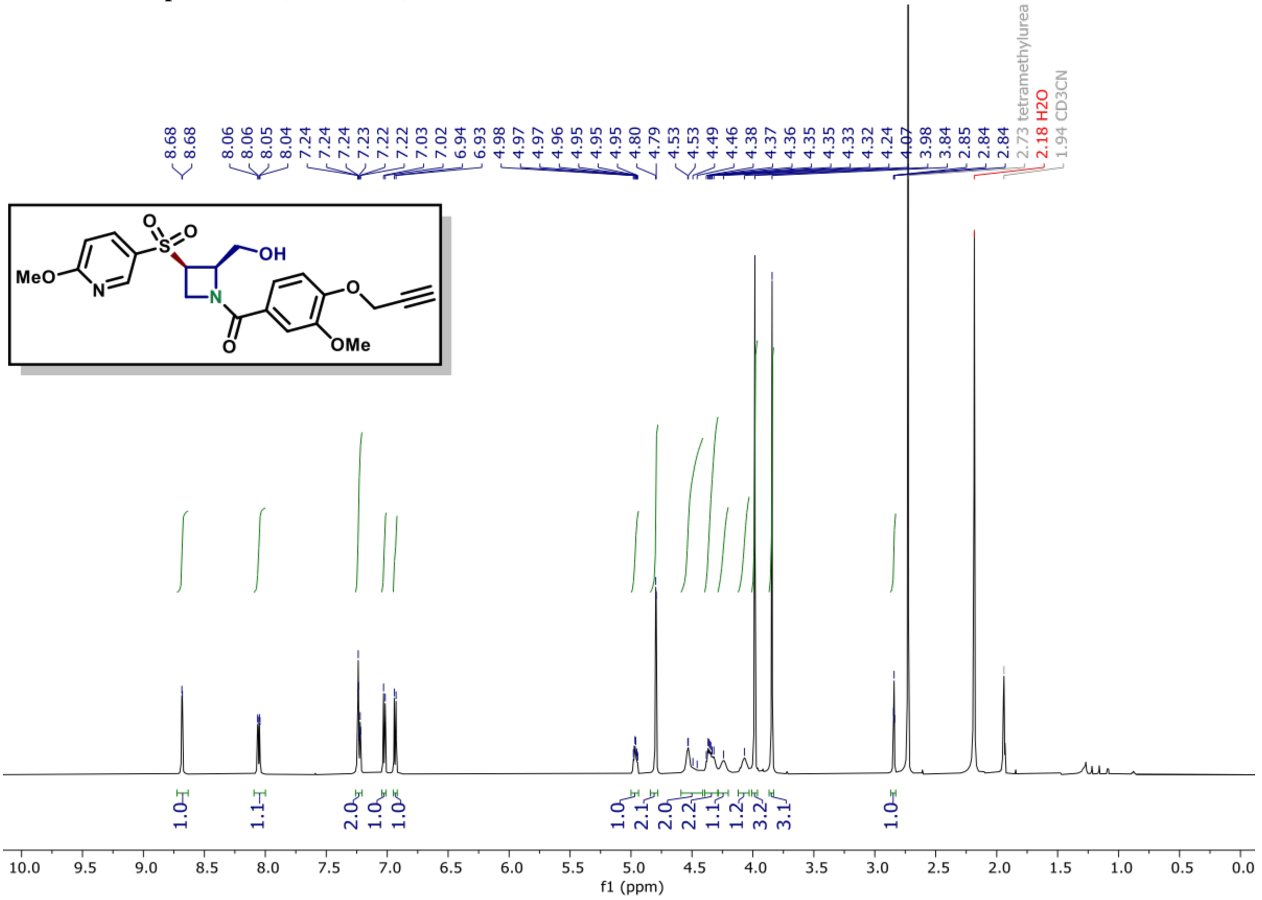

[13]C NMR of compound (11A) (150 MHz, $CDCl_3$)

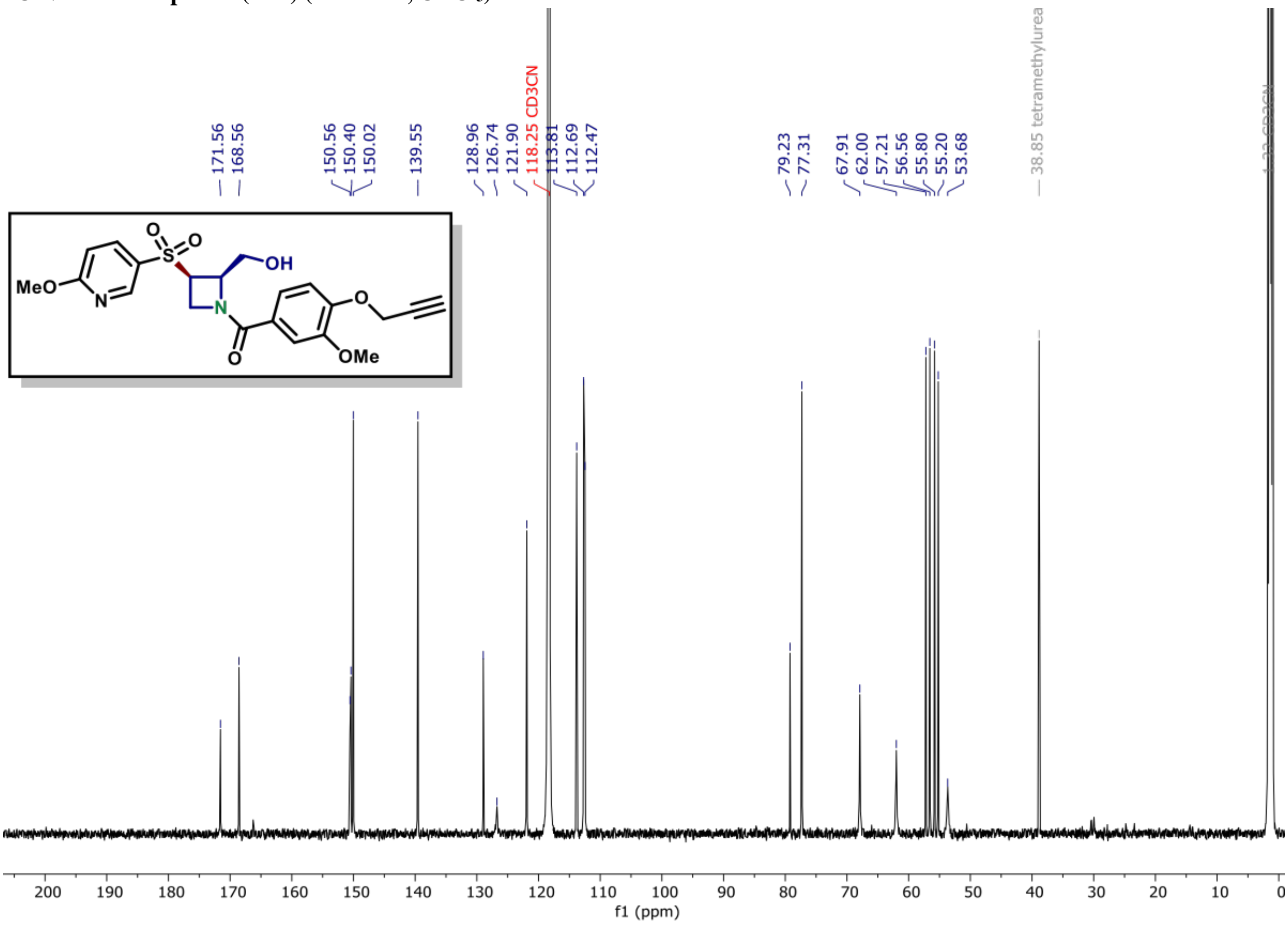

$^{1}H$ NMR of compound (11B) (600 MHz, $CDCl_3$)

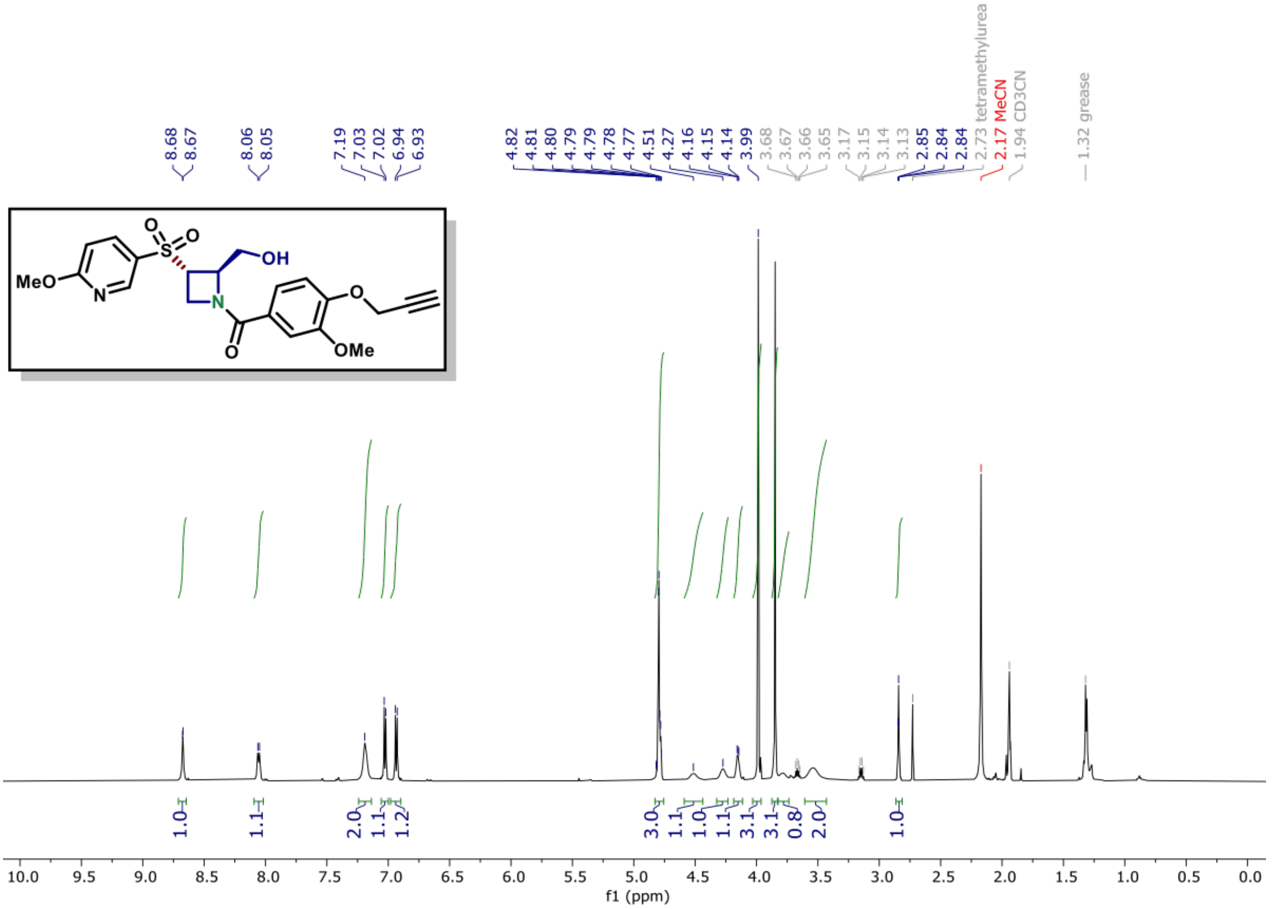

$^{13}C$ NMR of compound (11B) (150 MHz, $CDCl_3$)

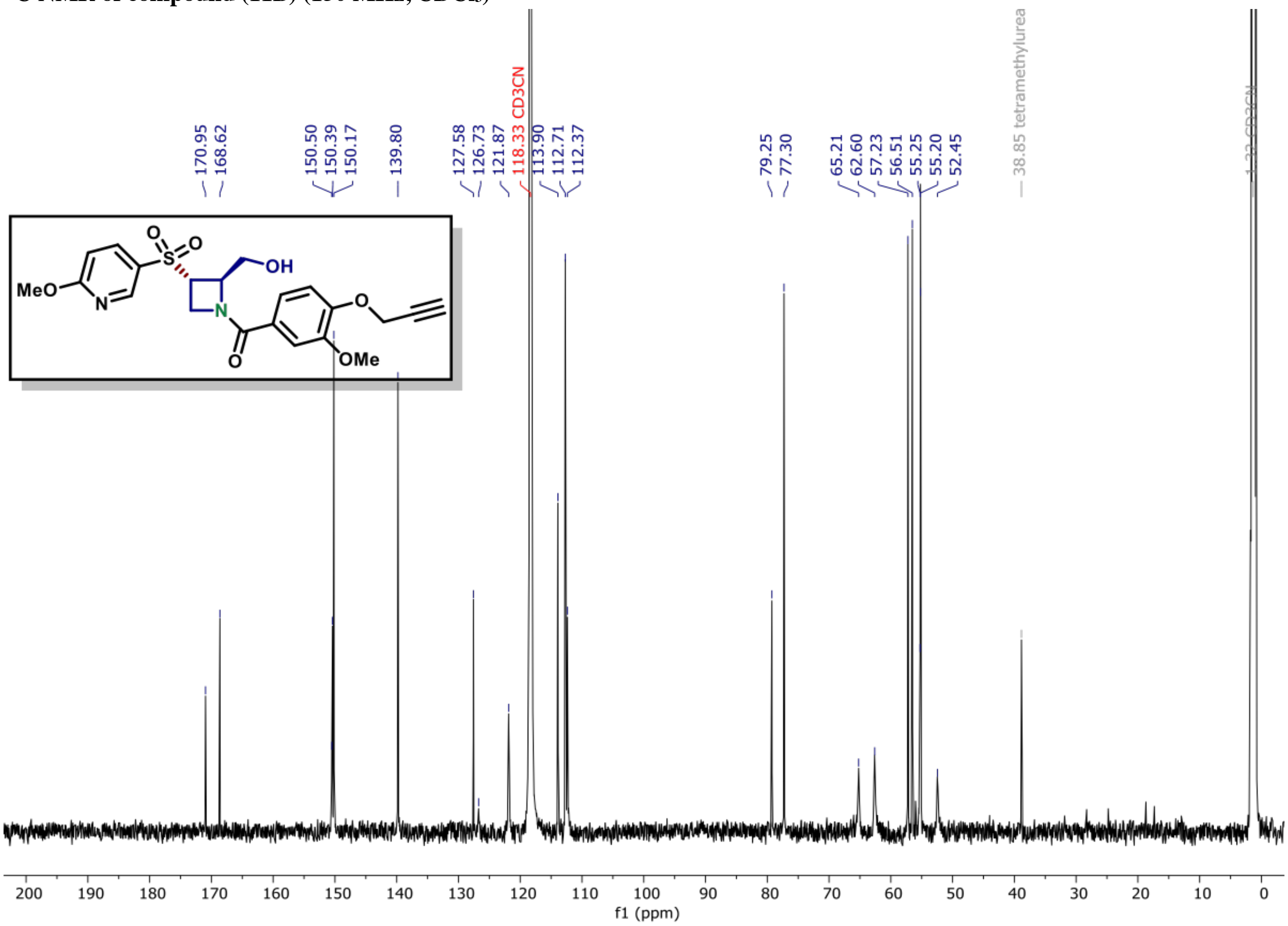

**[1]H NMR of compound (11C) (600 MHz, $CDCl_3$)**

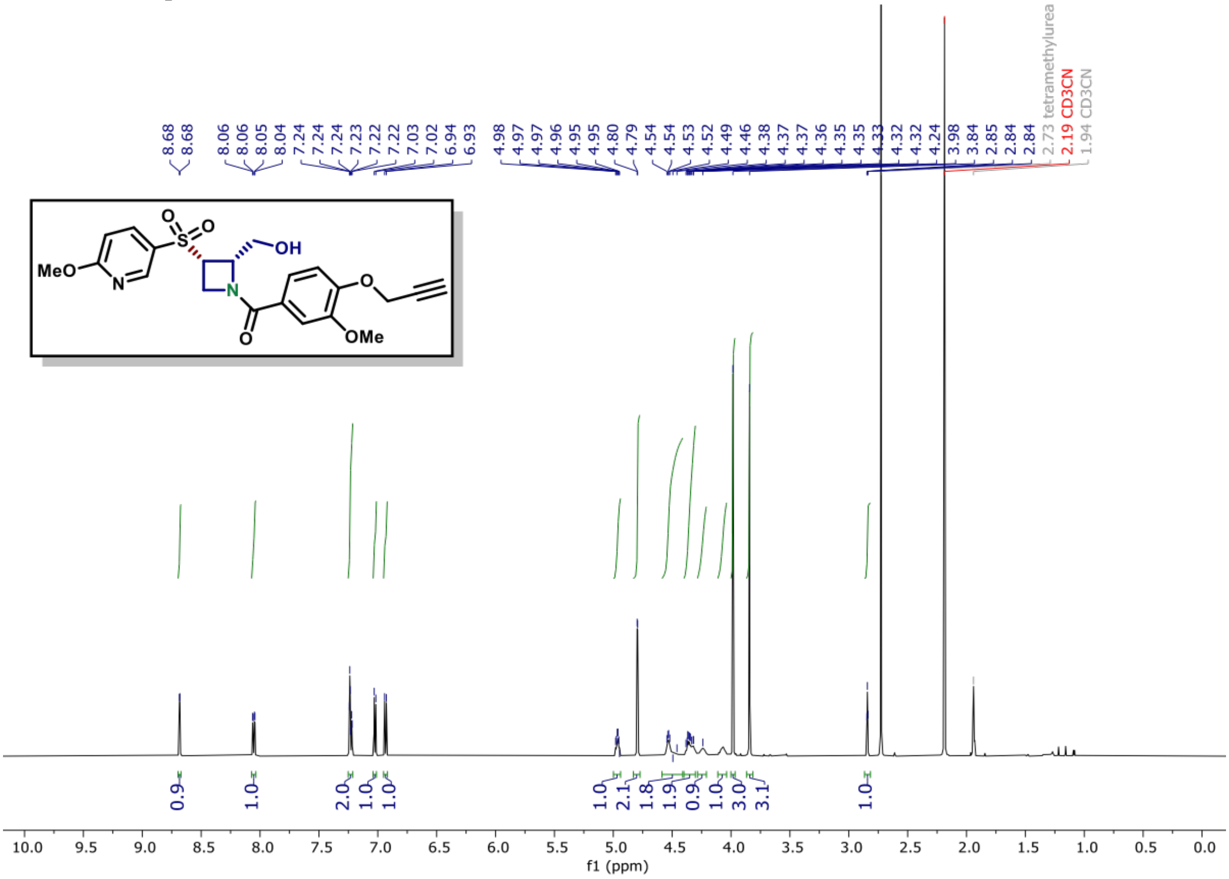

**[13]C NMR of compound (11C) (150 MHz, $CDCl_3$)**

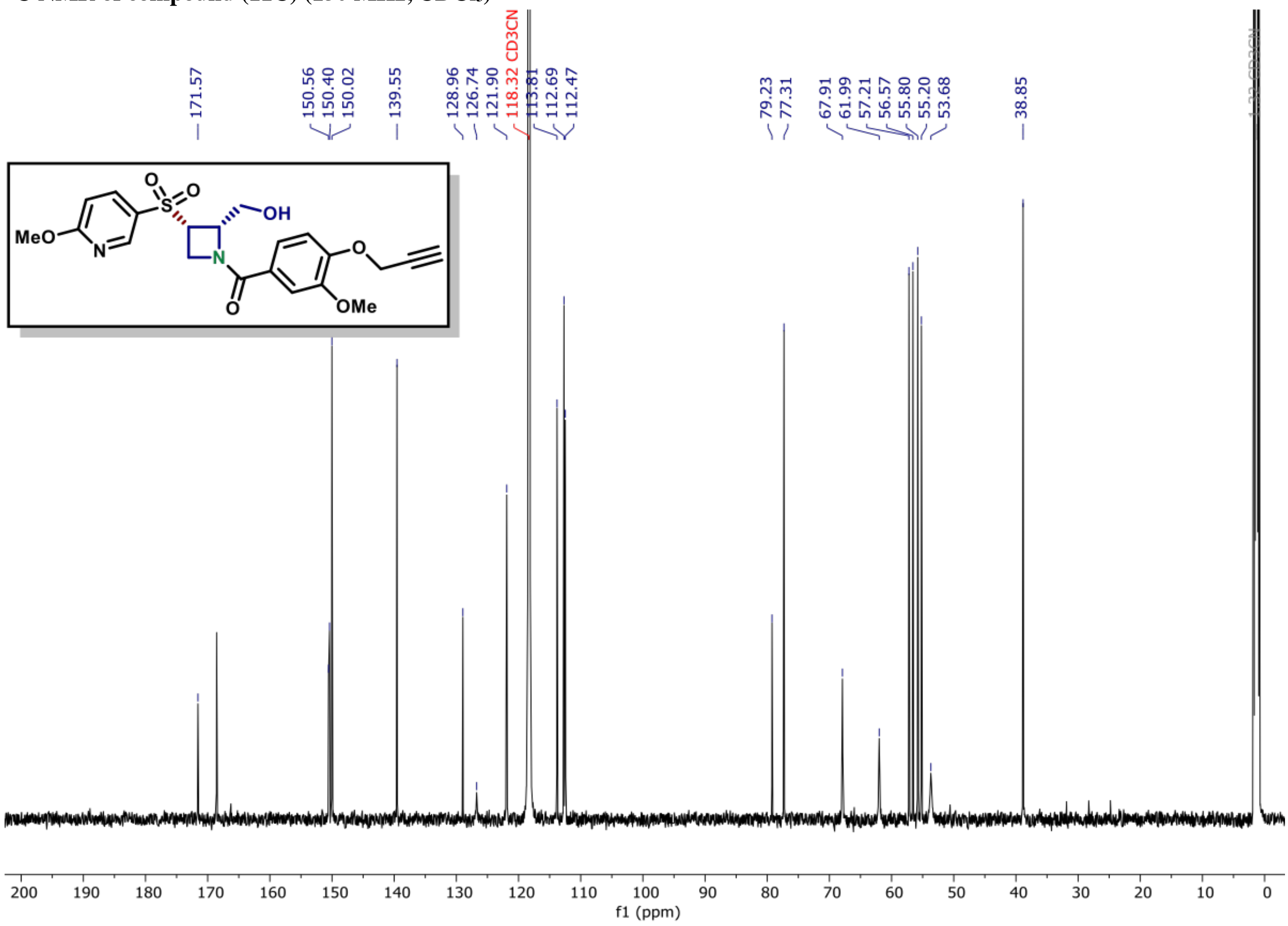

**$^{1}$H NMR of compound (11D) (600 MHz, CDCl$_3$)**

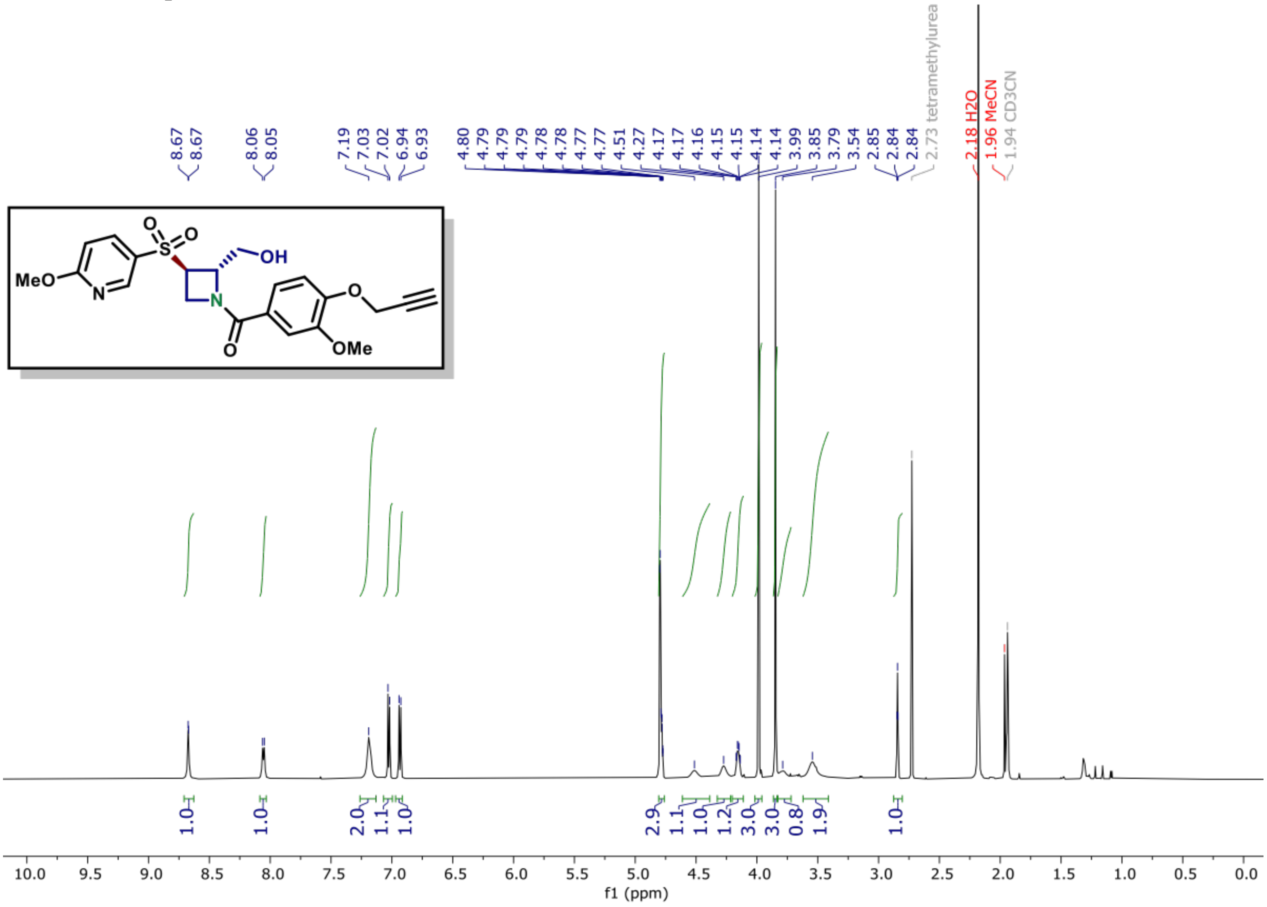

**$^{13}$C NMR of compound (11D) (150 MHz, CDCl$_3$)**

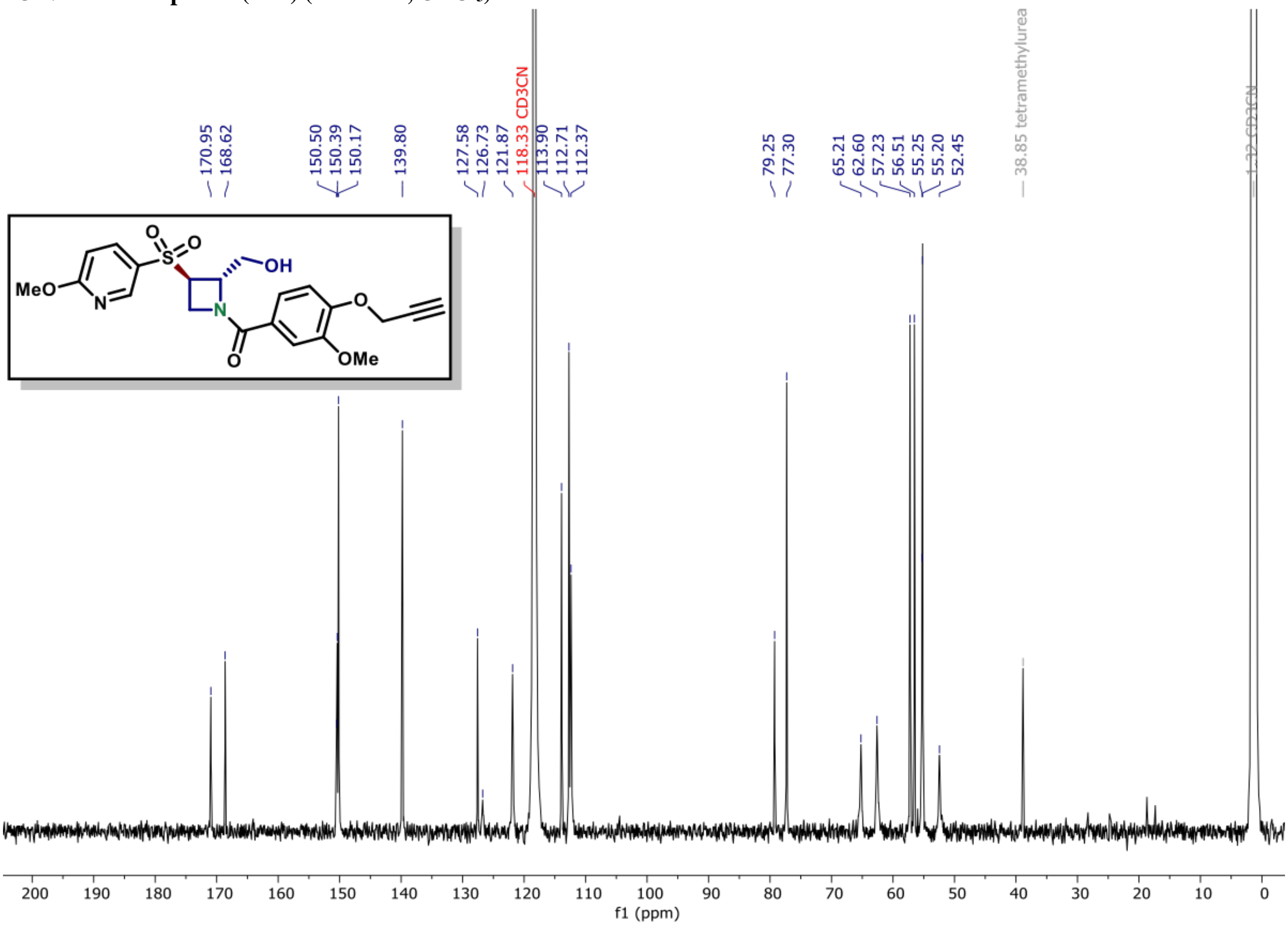

[1]H NMR of compound (12A) (600 MHz, $CDCl_3$)

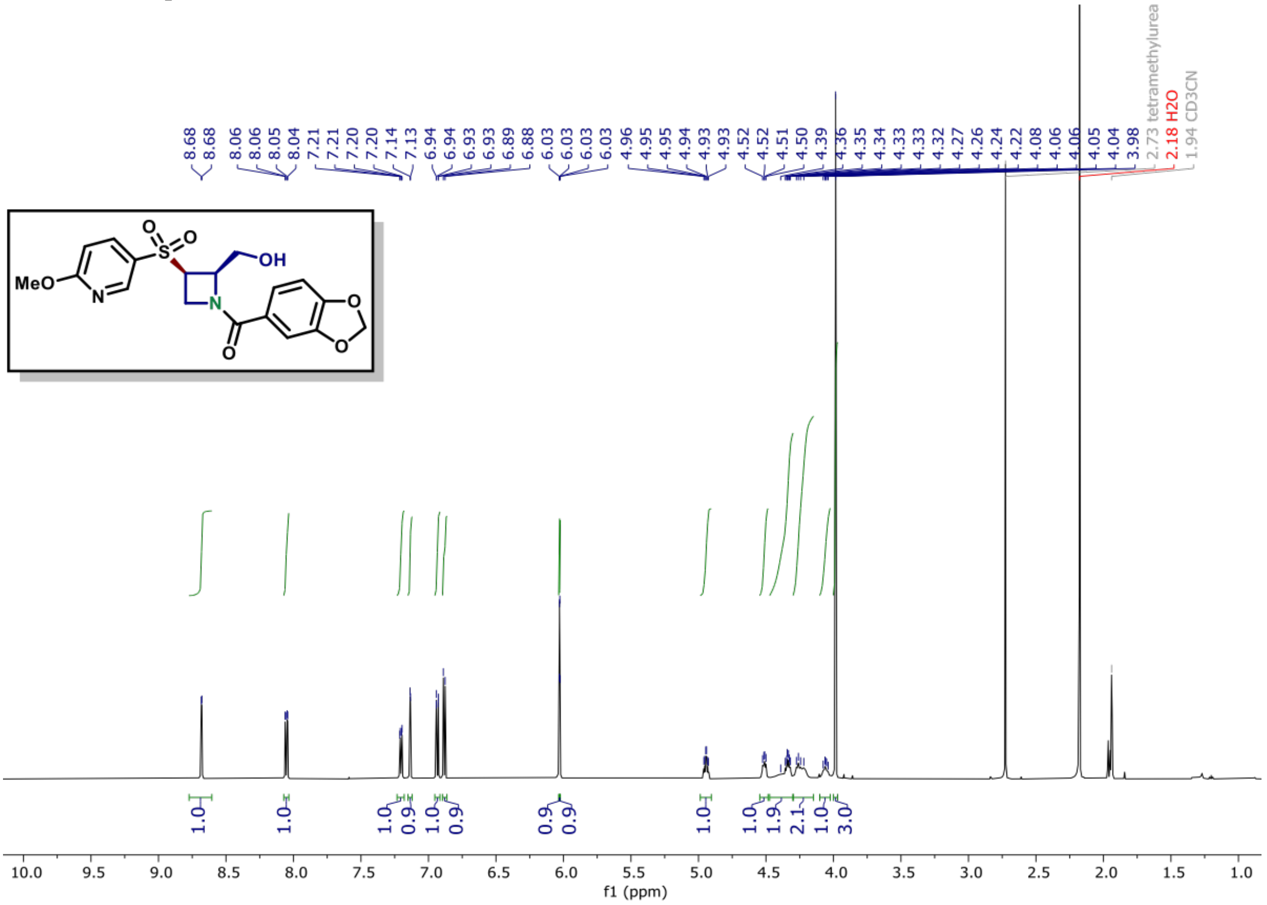

[13]C NMR of compound (12A) (150 MHz, $CDCl_3$)

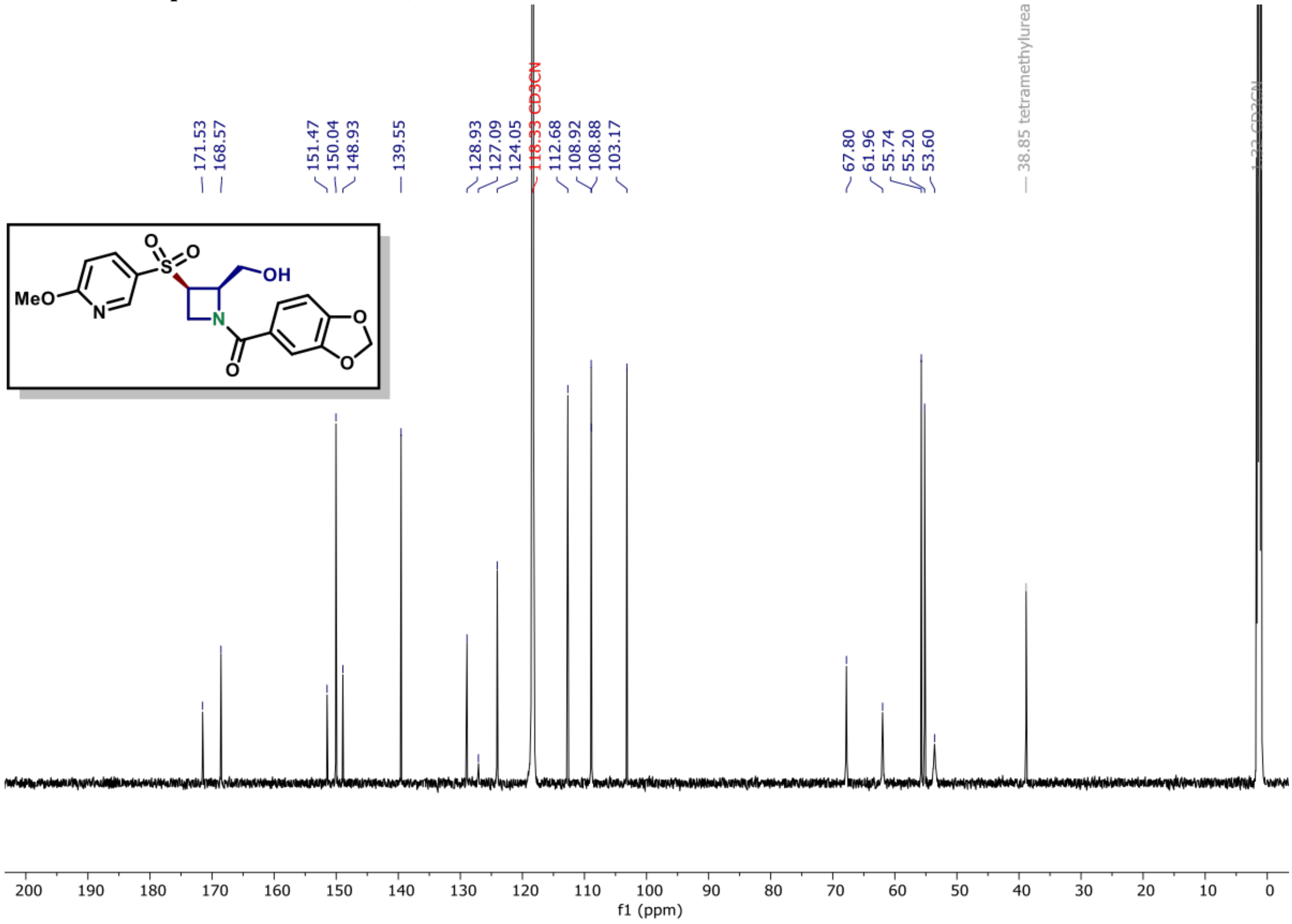

[1]H NMR of compound (12B) (600 MHz, $CDCl_3$)

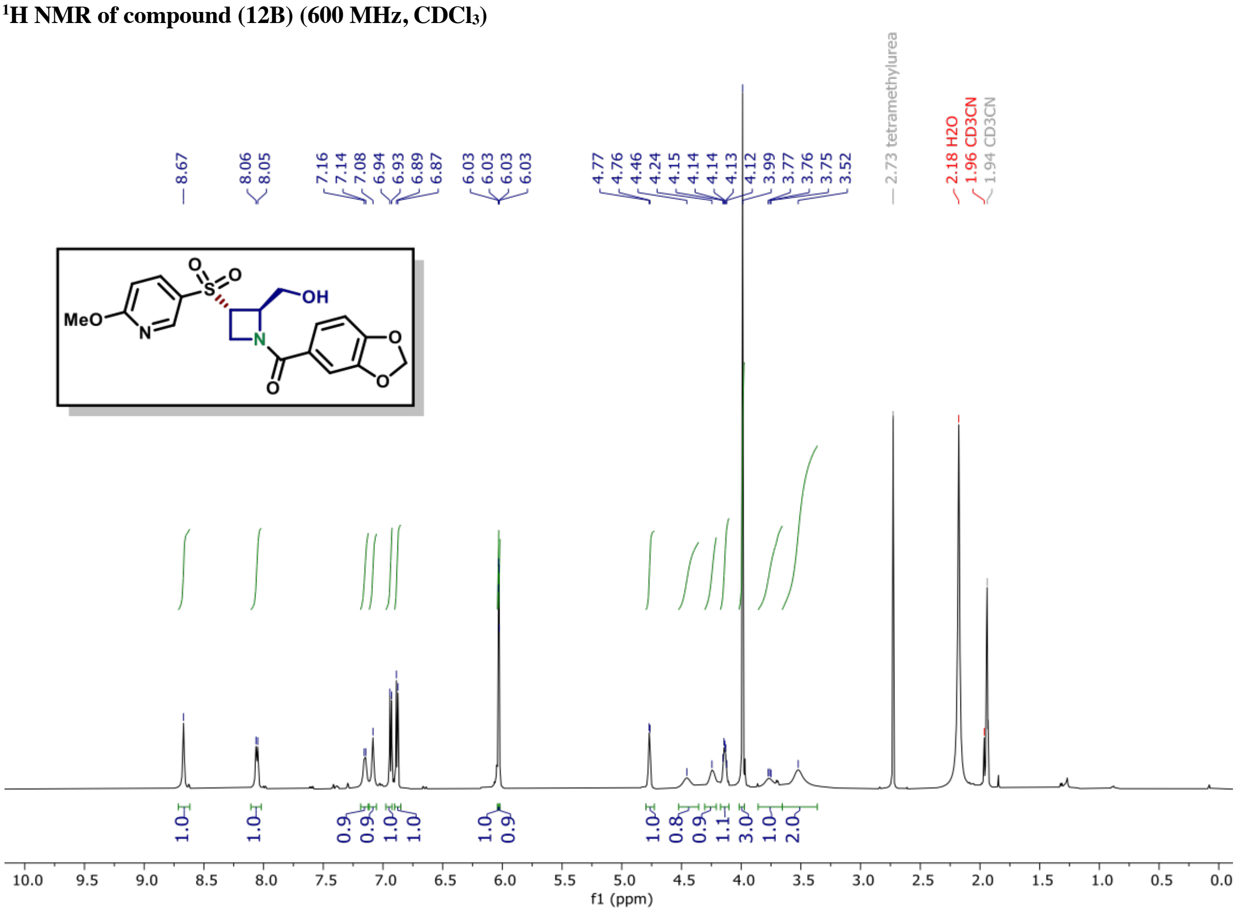

[13]C NMR of compound (12B) (150 MHz, $CDCl_3$)

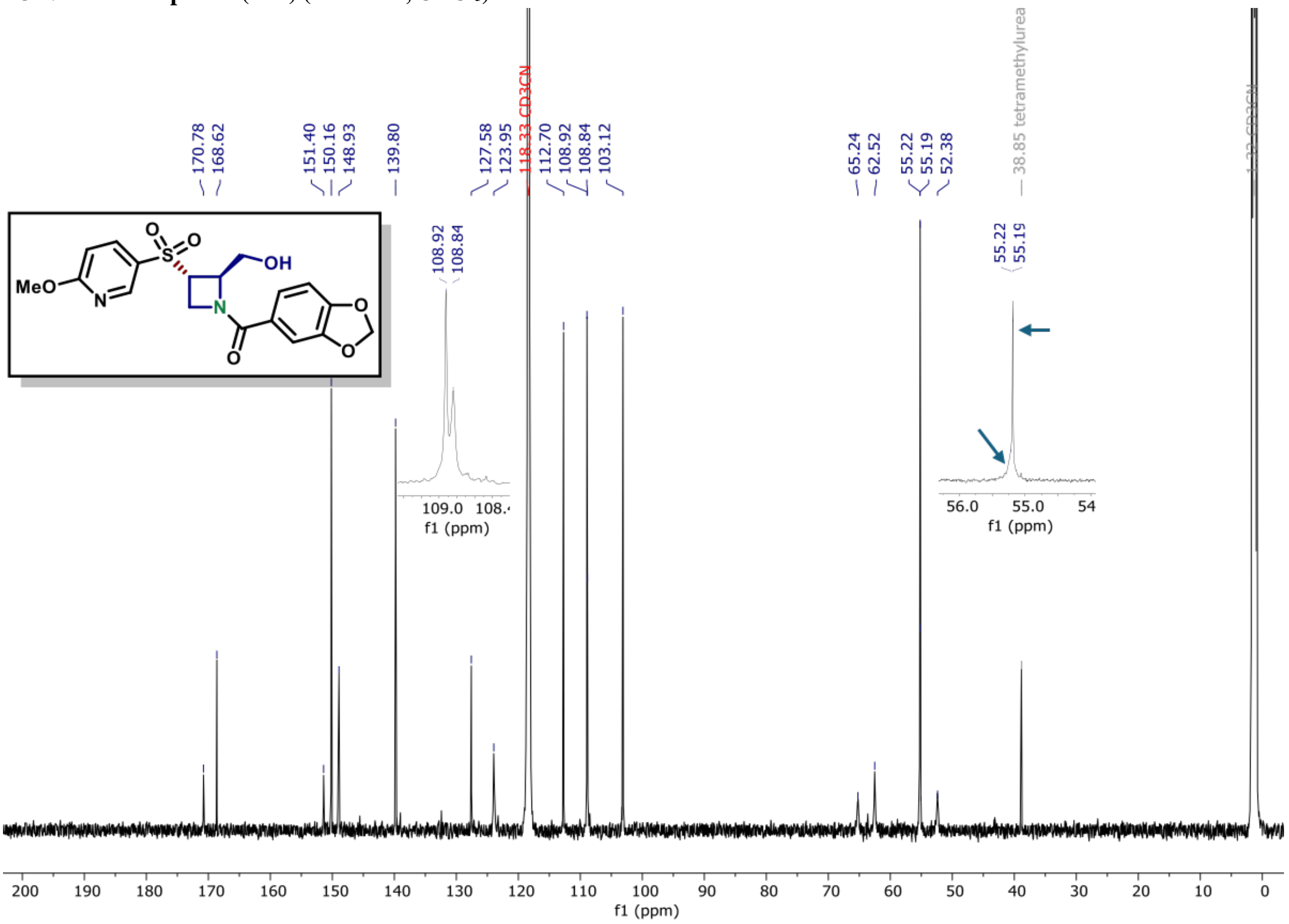

[1]H NMR of compound (12C) (600 MHz, $CDCl_3$)

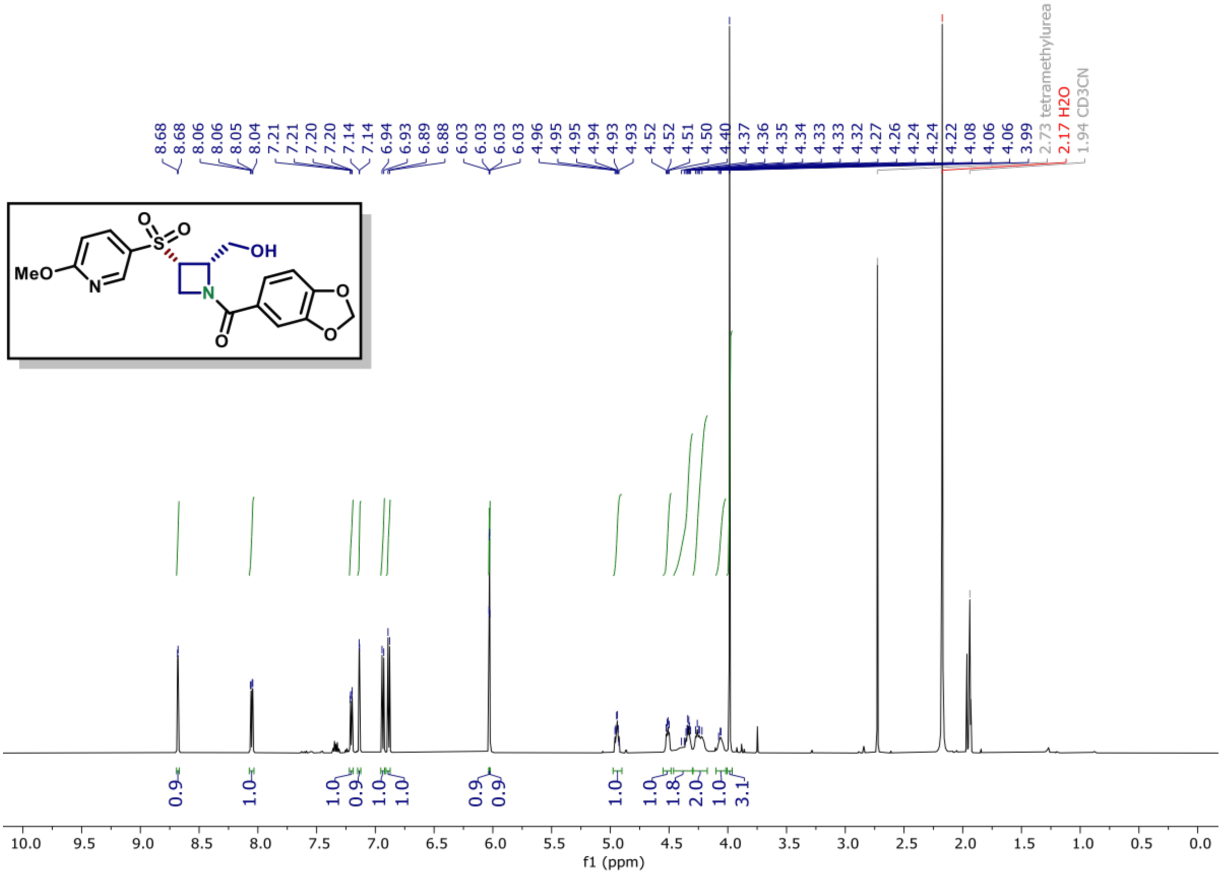

[13]C NMR of compound (12C) (150 MHz, $CDCl_3$)

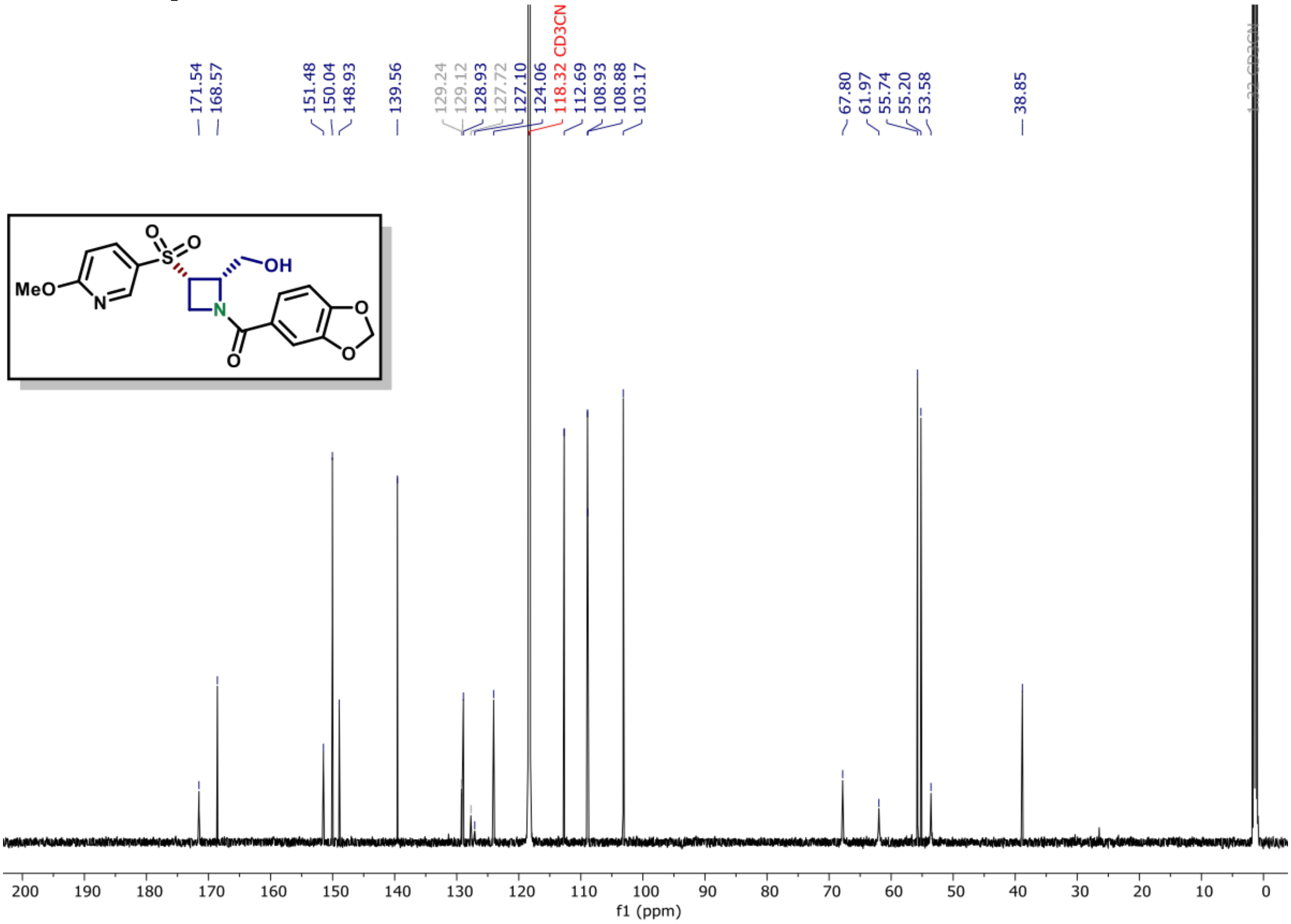

**[1]H NMR of compound (12D) (600 MHz, CDCl3)**

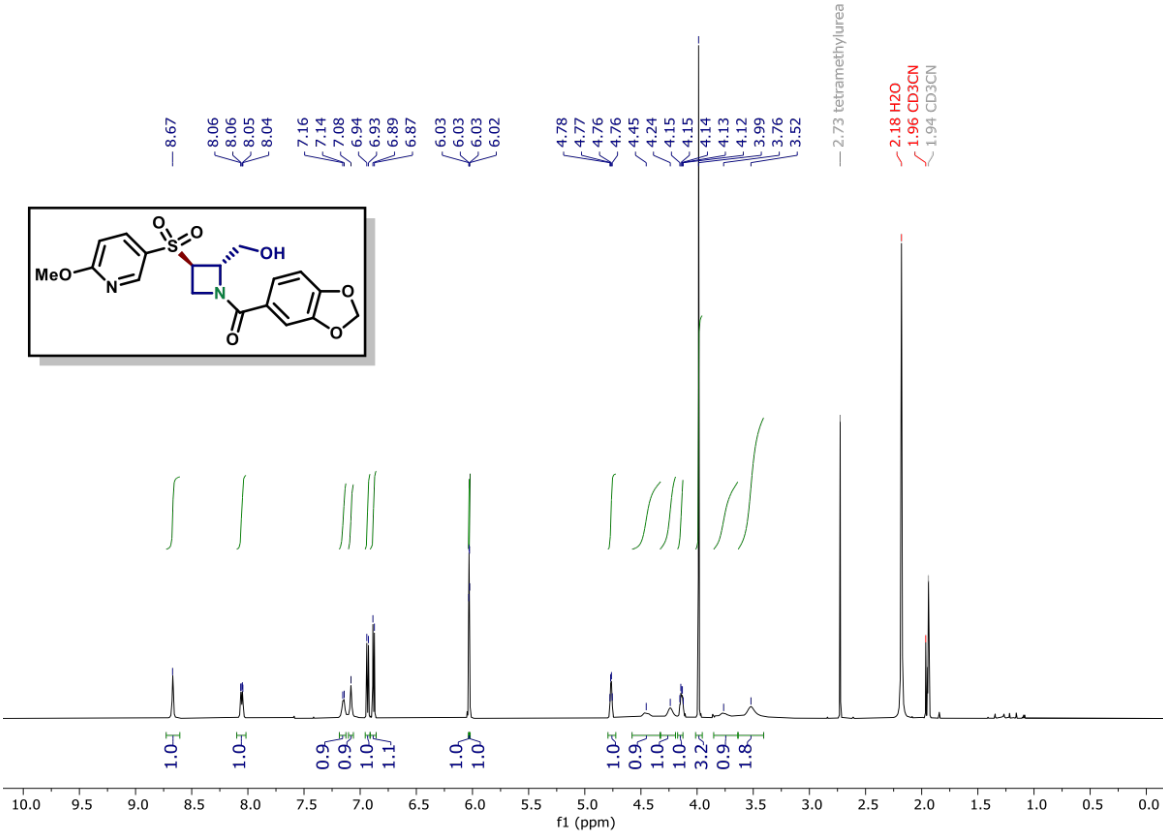

**[13]C NMR of compound (12D) (150 MHz, CDCl3)**

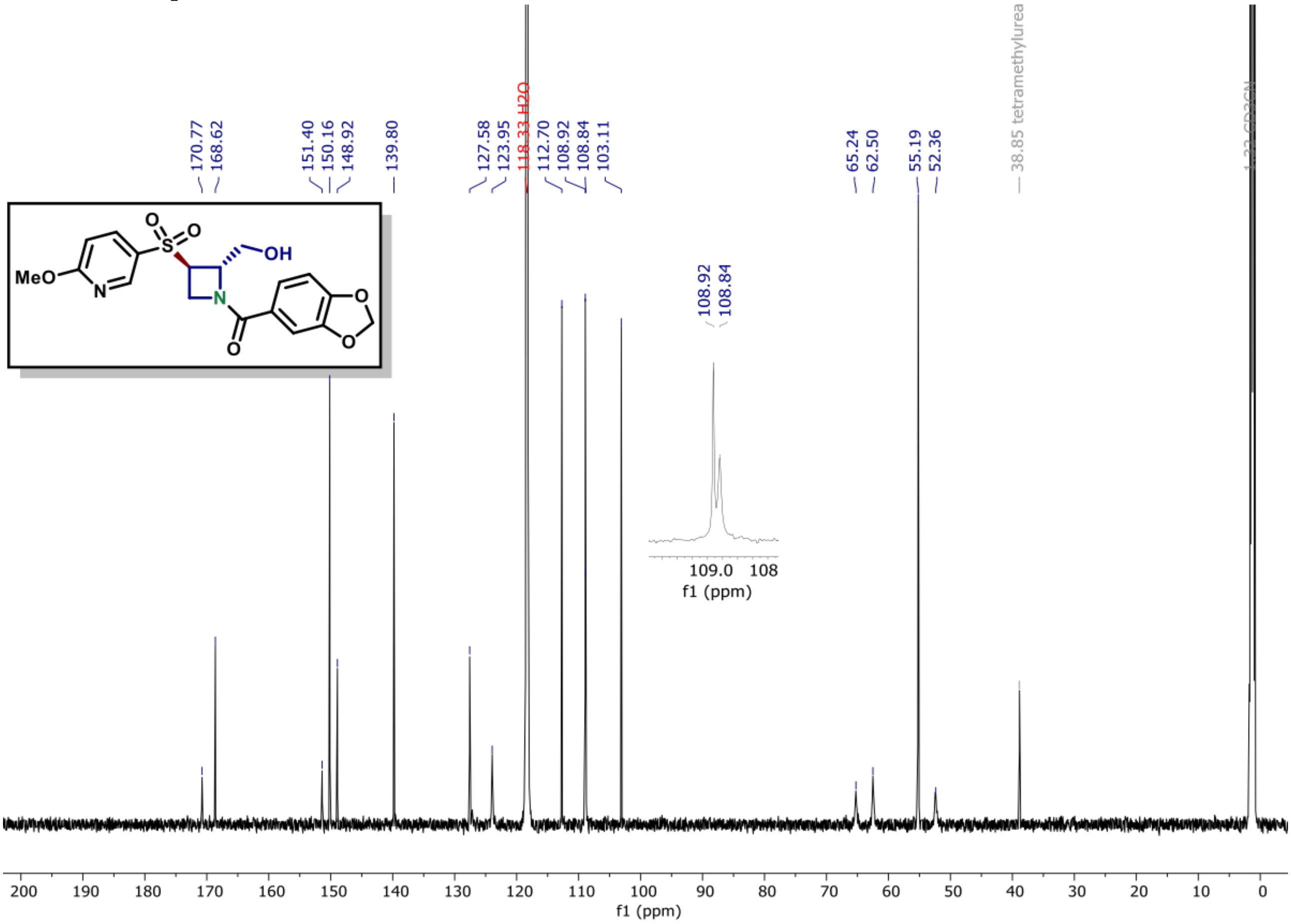

$^{31}P$ NMR of compound (1) (162 MHz, $CDCl_3$)

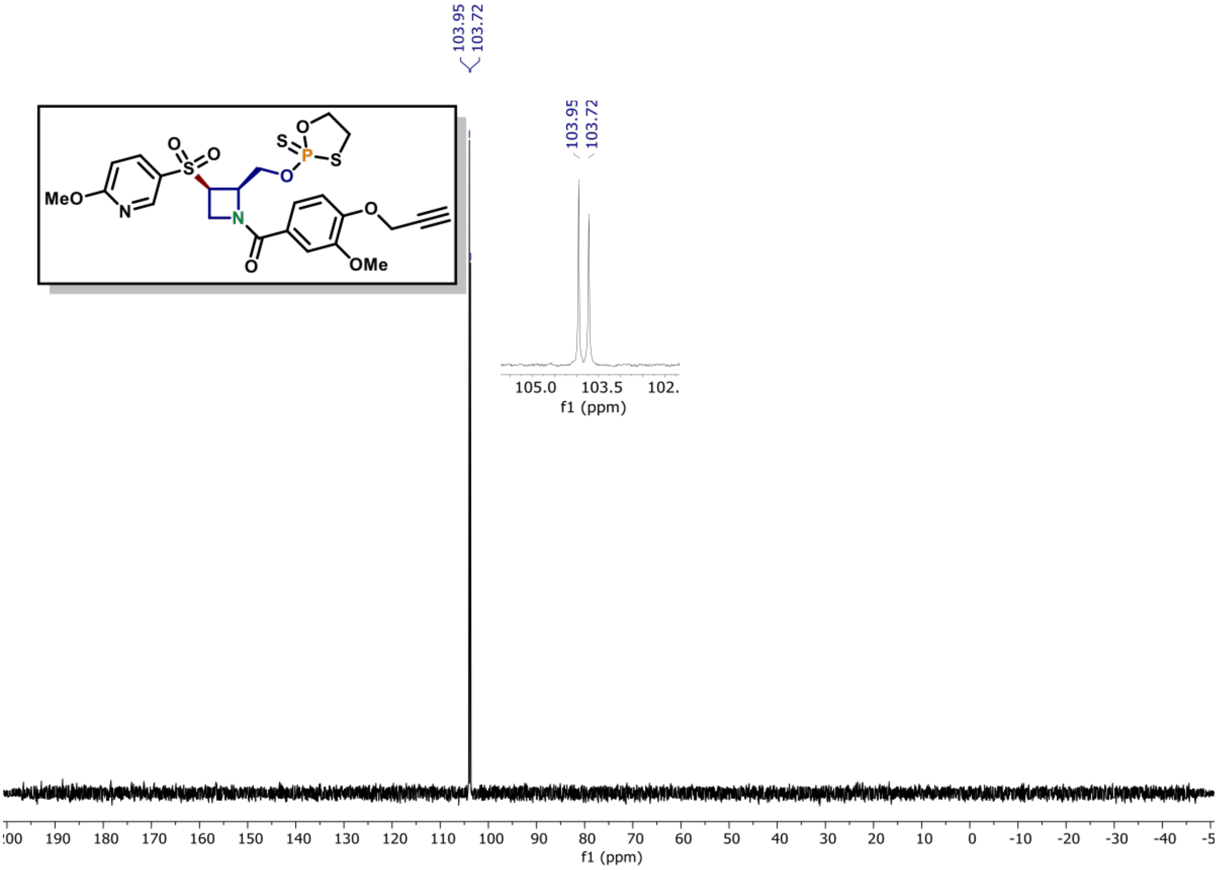

$^{1}H$ NMR of compound (5A) (600 MHz, $CDCl_3$)

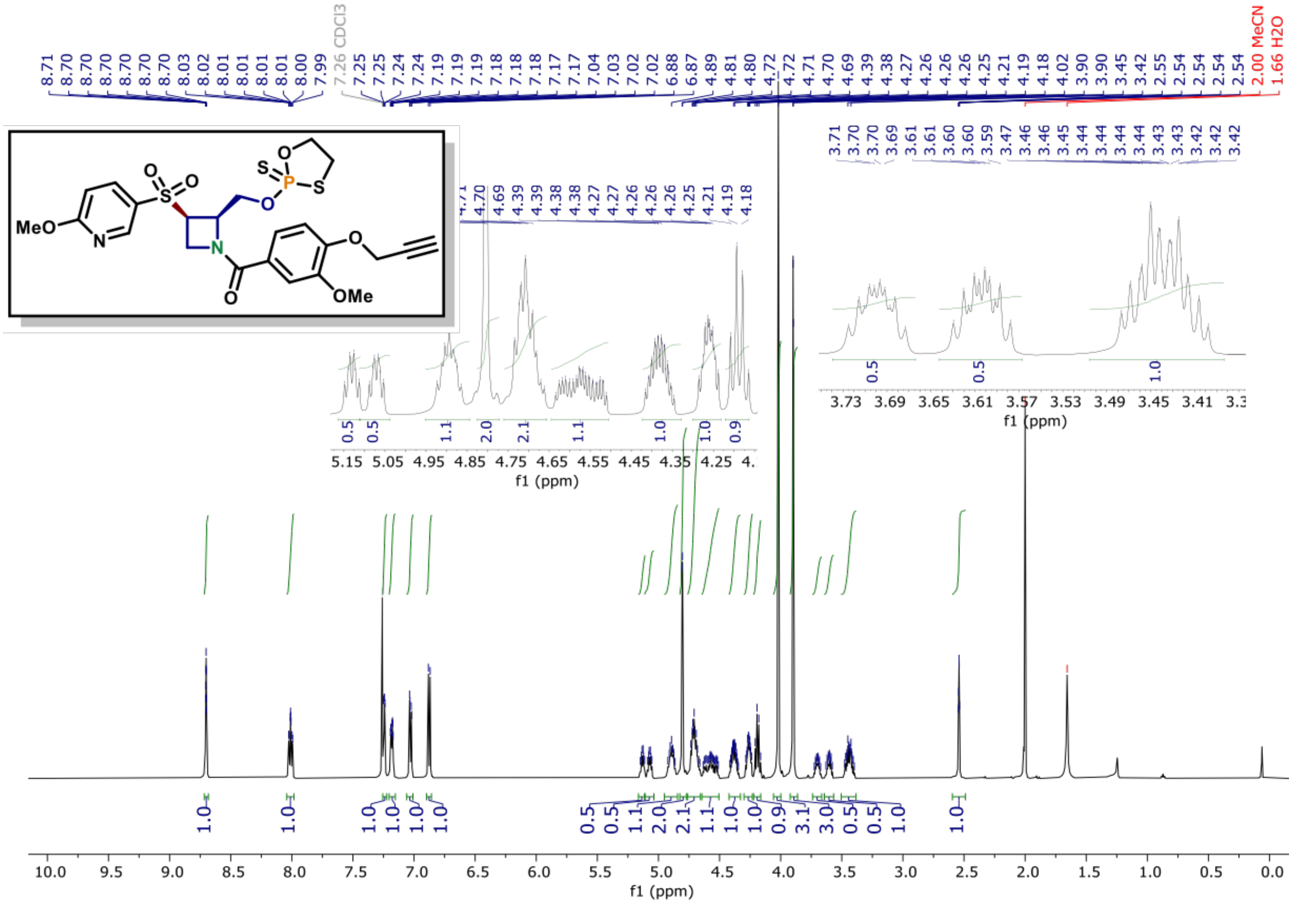

**$^{13}$C NMR of compound (1) (150 MHz, $CDCl_3$)**

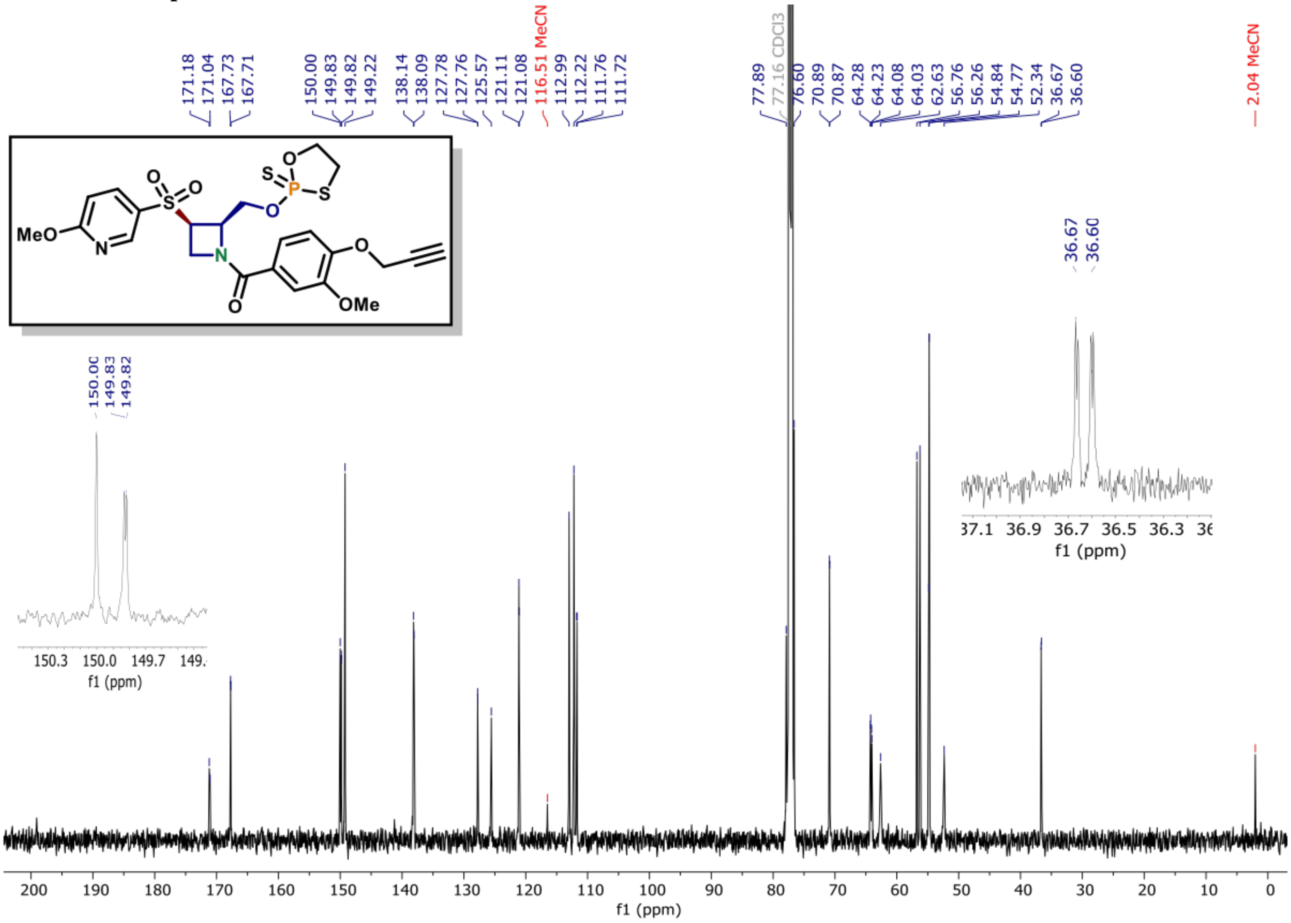

**[31]P NMR of compound (3) (162 MHz, $CDCl_3$)**

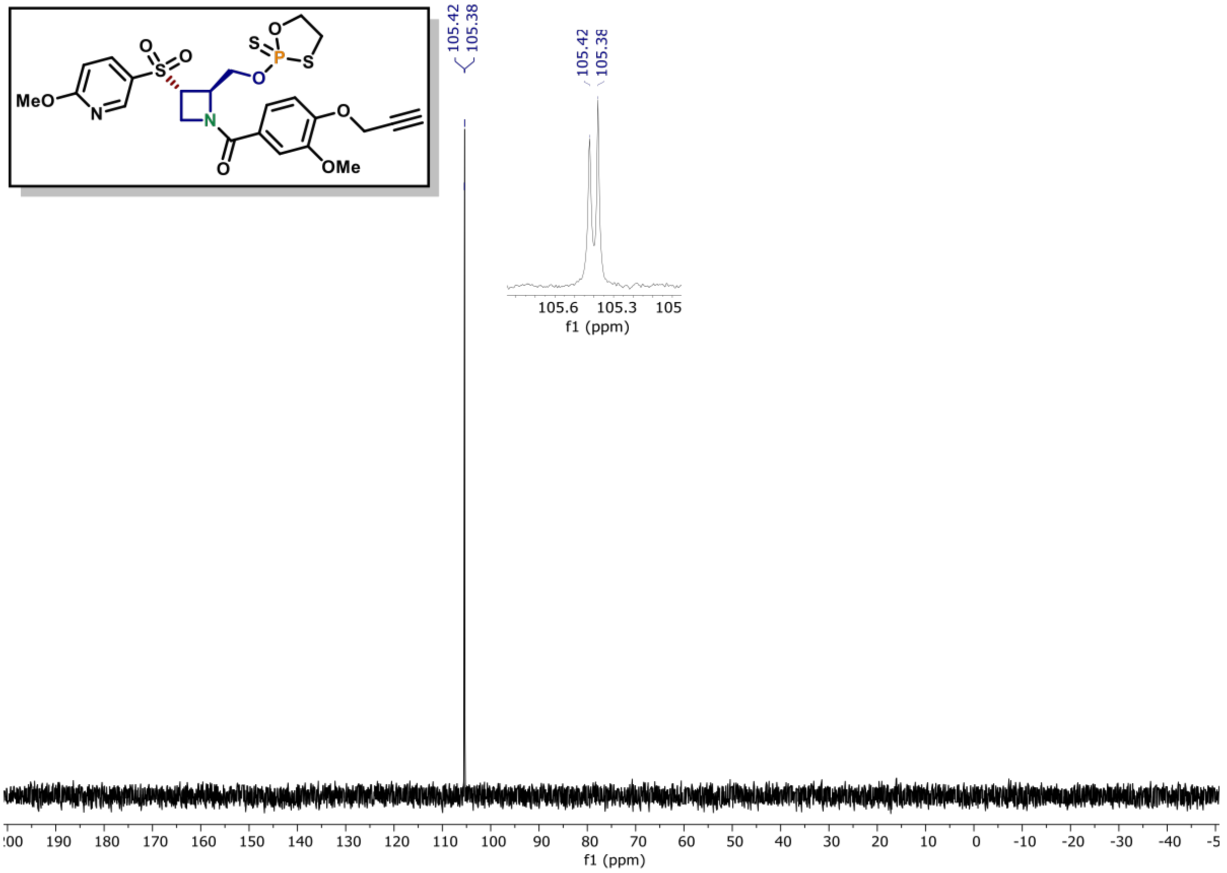

**[1]H NMR of compound (3) (600 MHz, $CDCl_3$)**

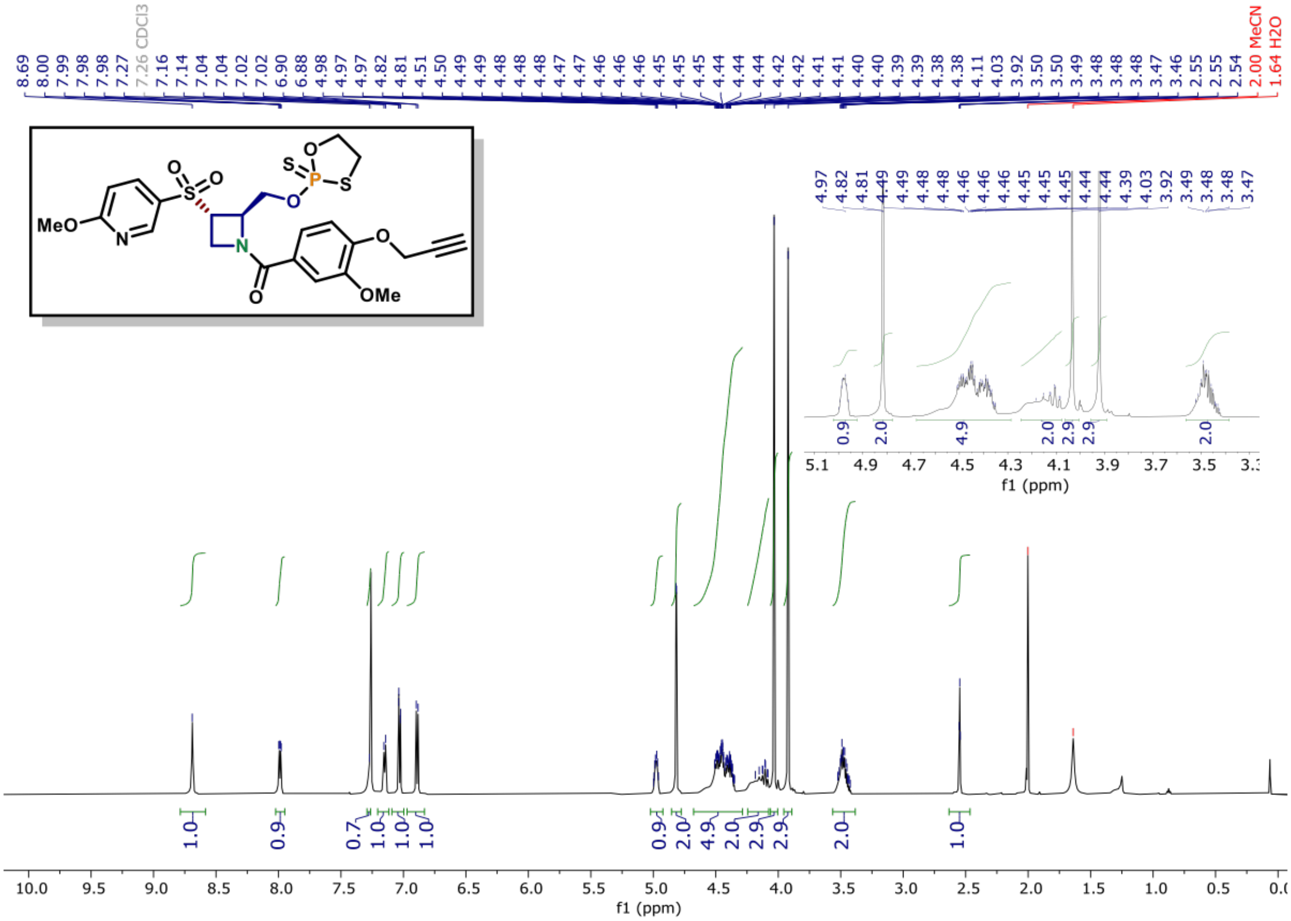

**$^{13}$C NMR of compound (3) (150 MHz, $CDCl_3$)**

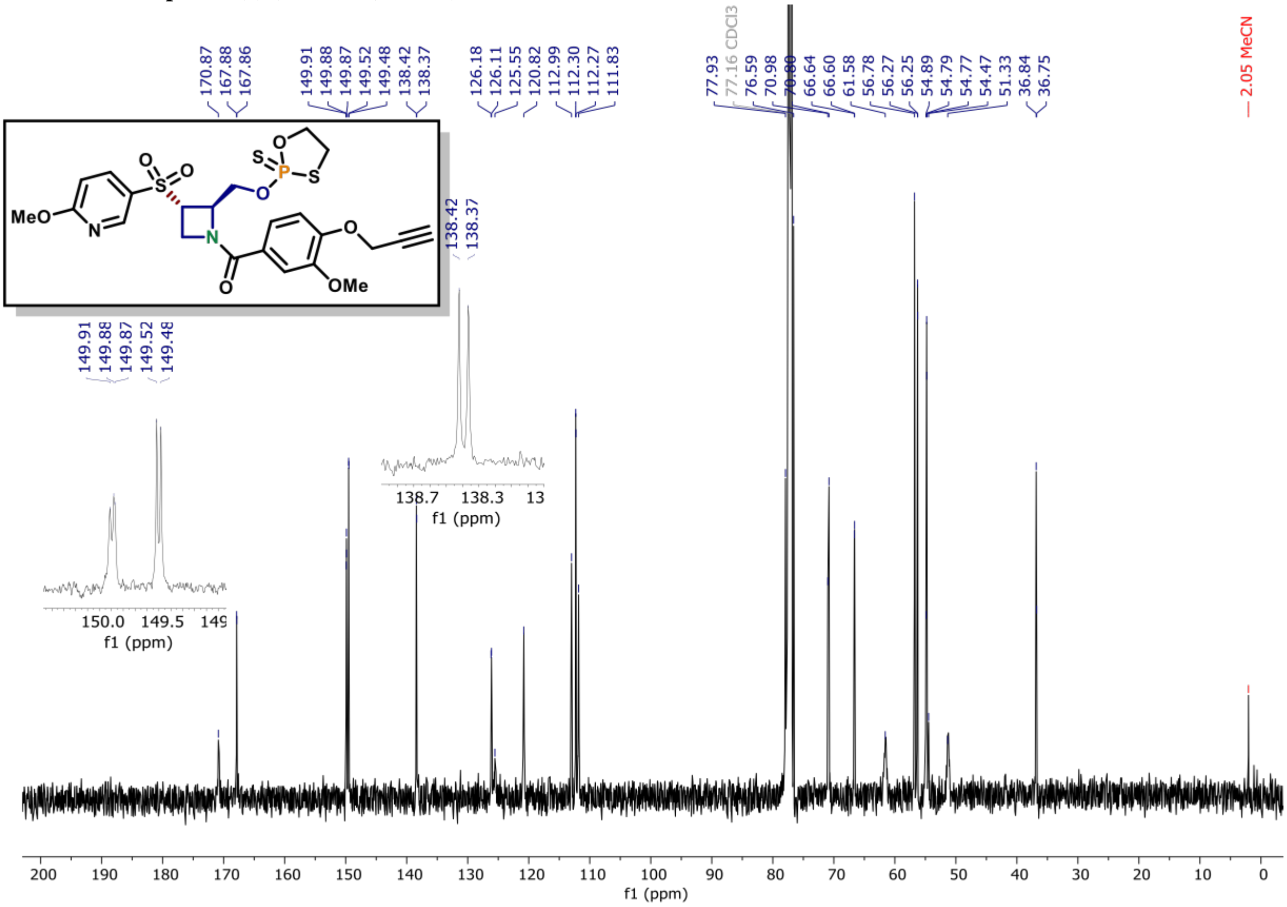

[31]P NMR of compound (2) (162 MHz, CDCl$_3$)

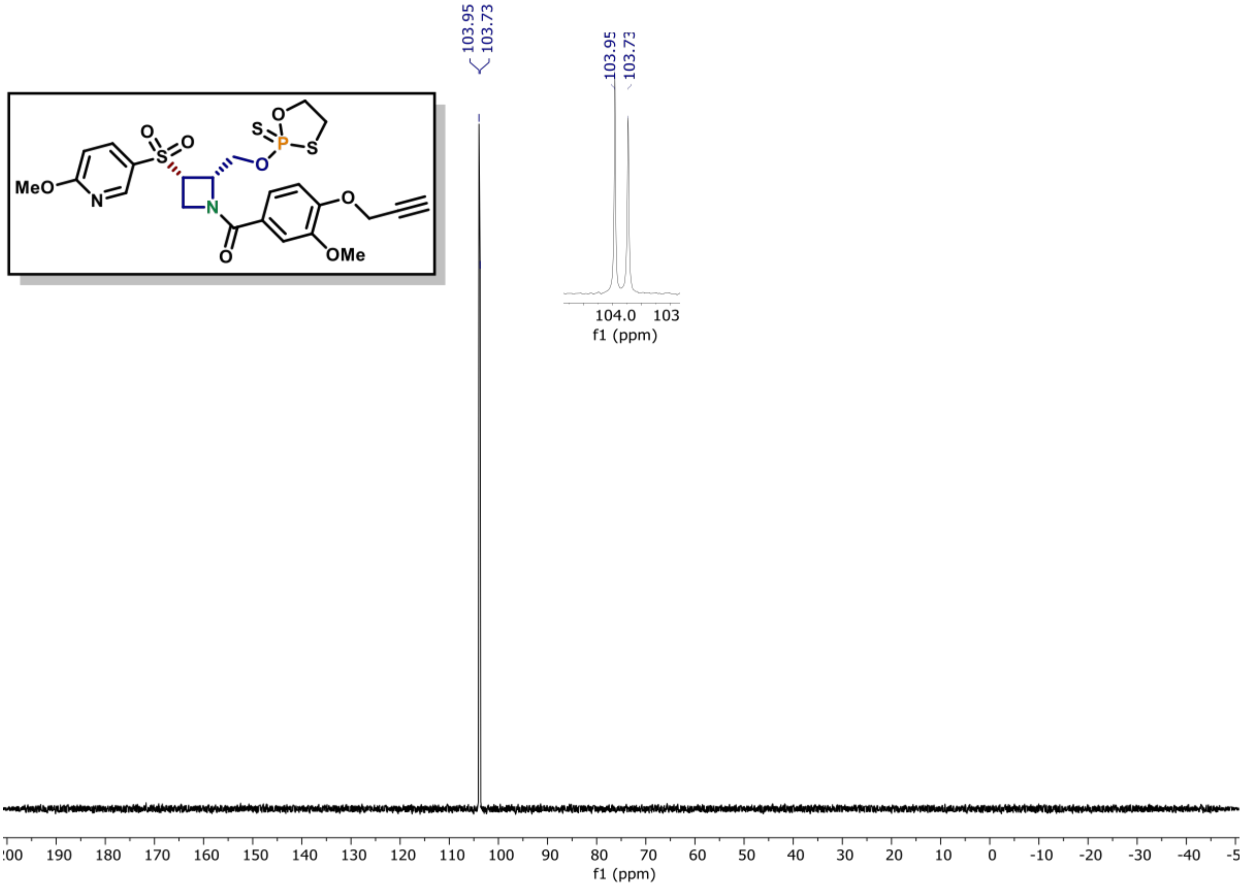

[1]H NMR of compound (2) (600 MHz, CDCl$_3$)

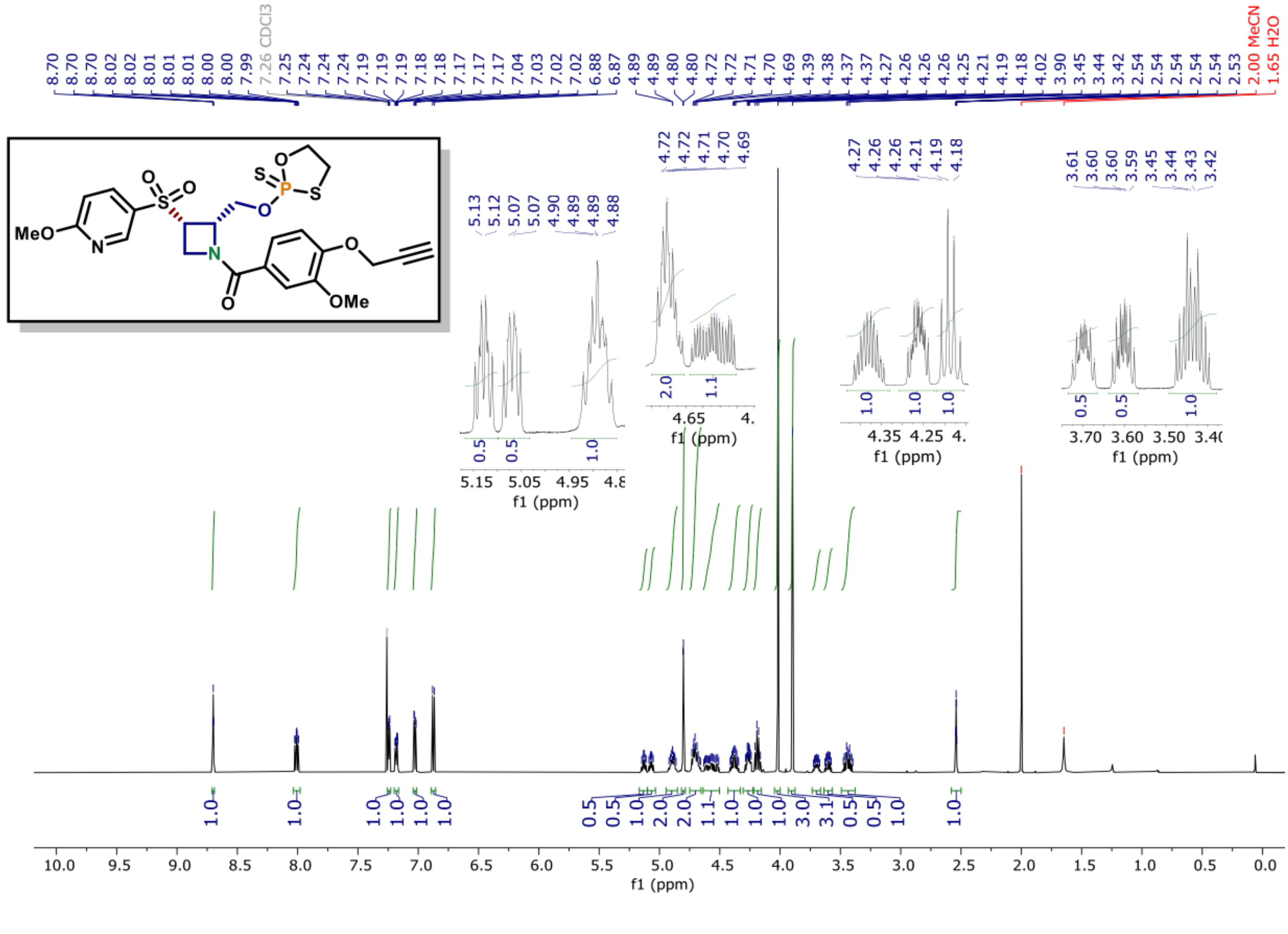

[13]C NMR of compound (2) (150 MHz, CDCl₃)

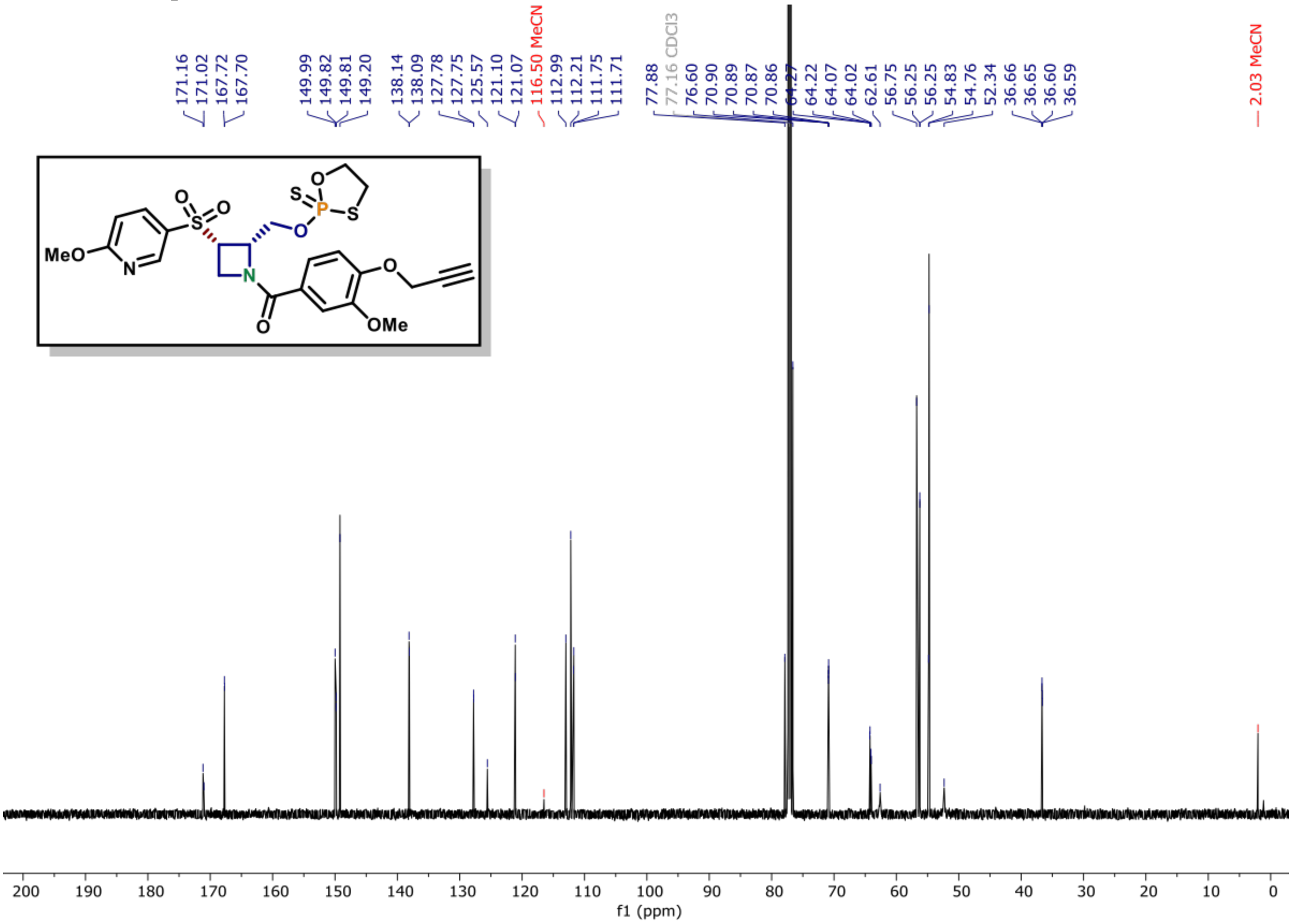

**[31]P NMR of compound (4) (162 MHz, $CDCl_3$)**

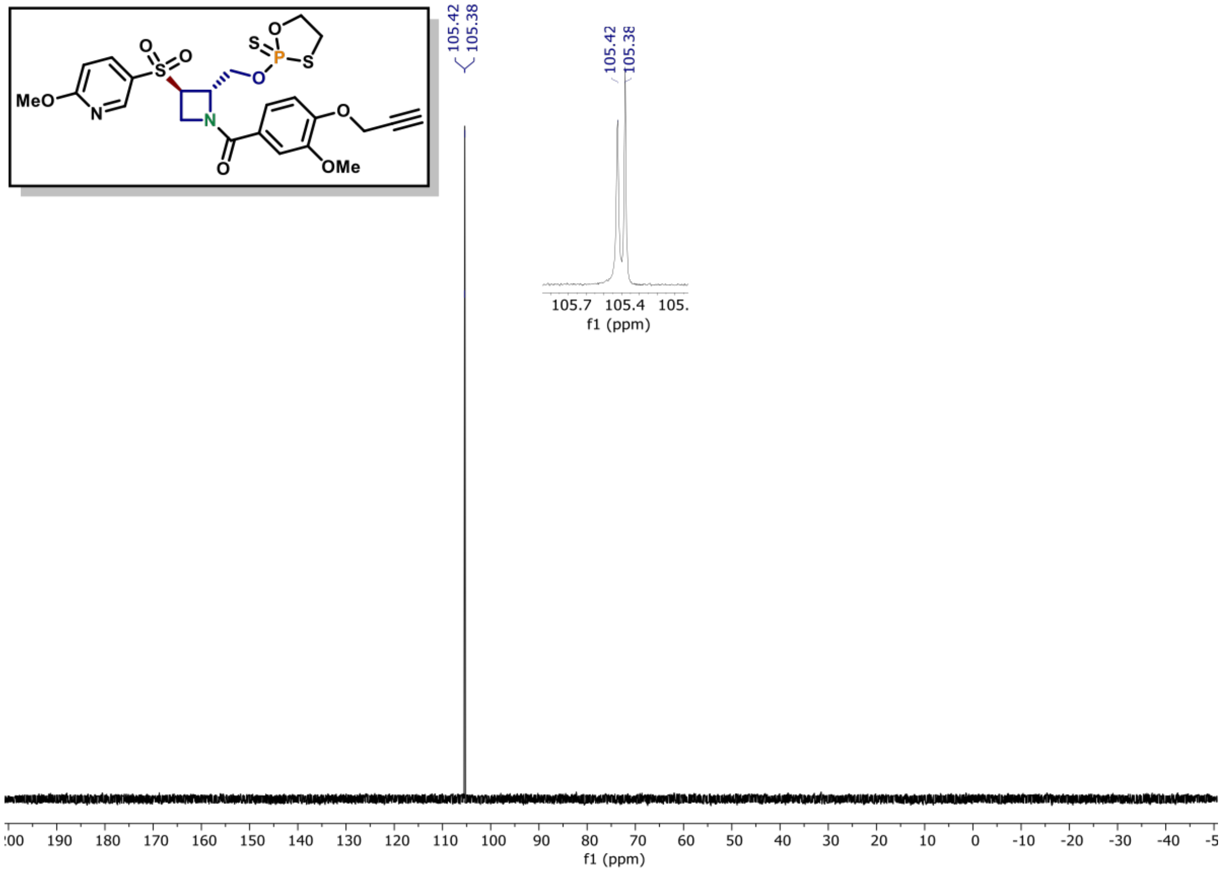

**[1]H NMR of compound (4) (600 MHz, $CDCl_3$)**

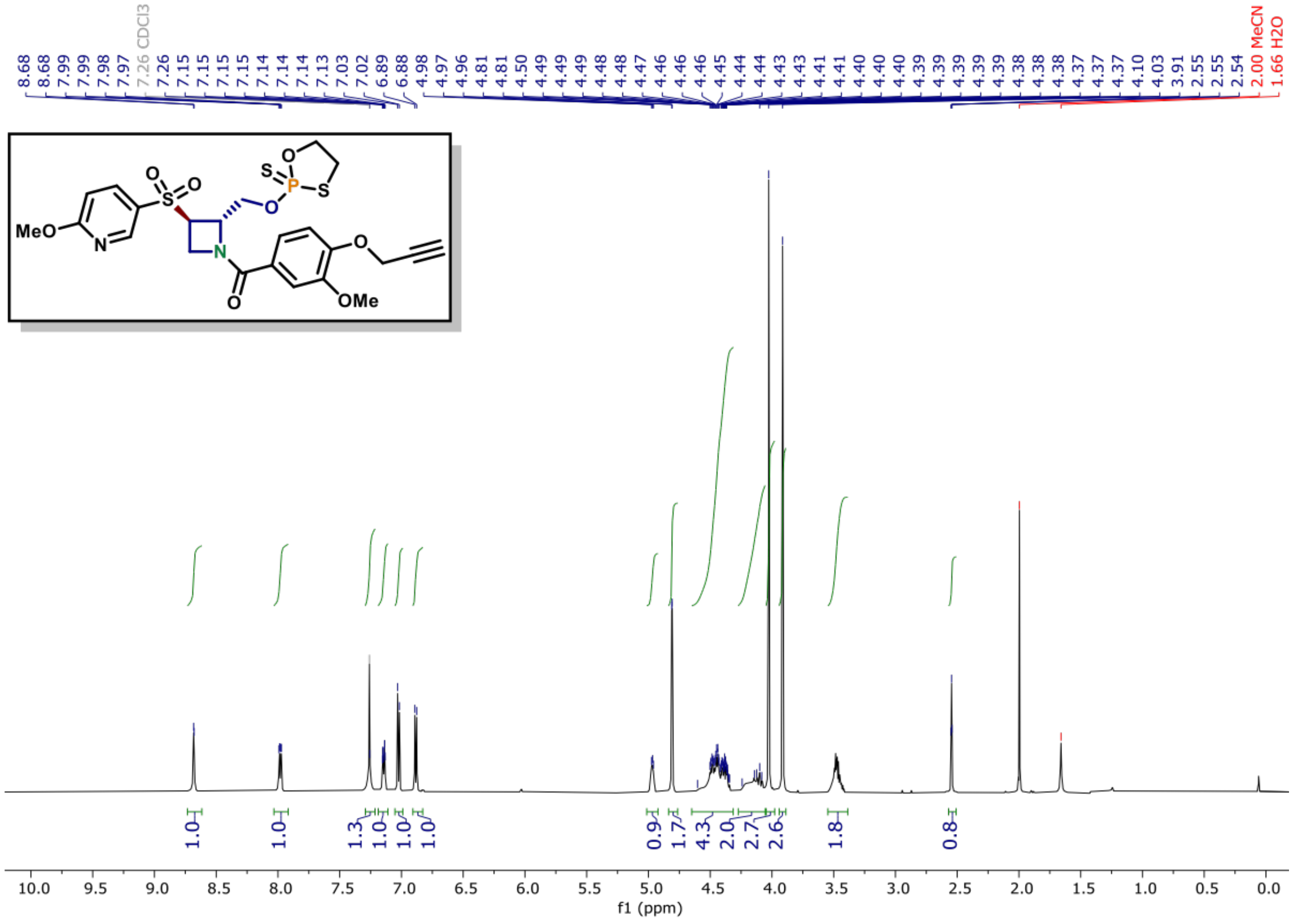

[13]C NMR of compound (4) (150 MHz, CDCl$_3$)

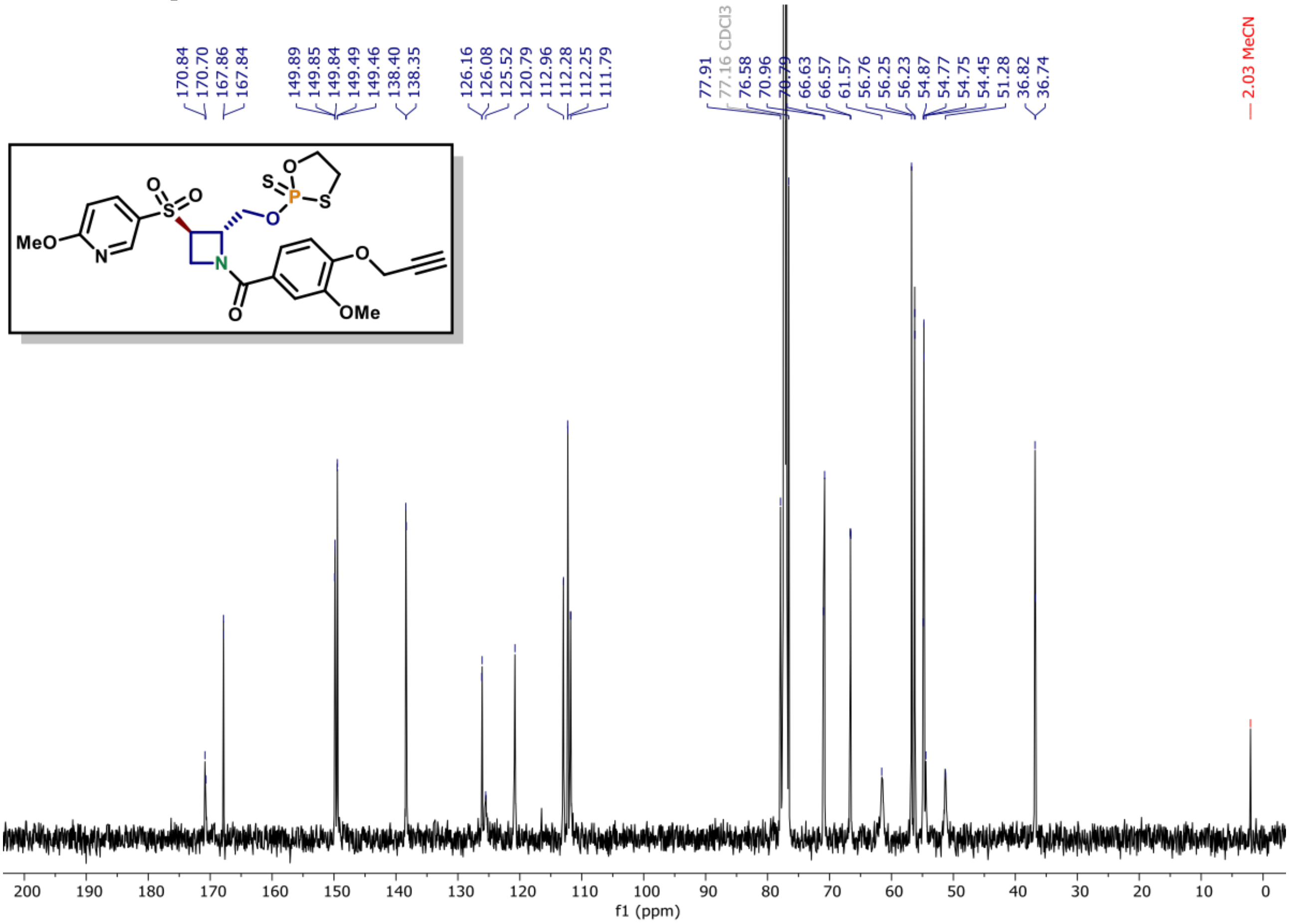

**[31]P NMR of compound (5) (162 MHz, $CDCl_3$)**

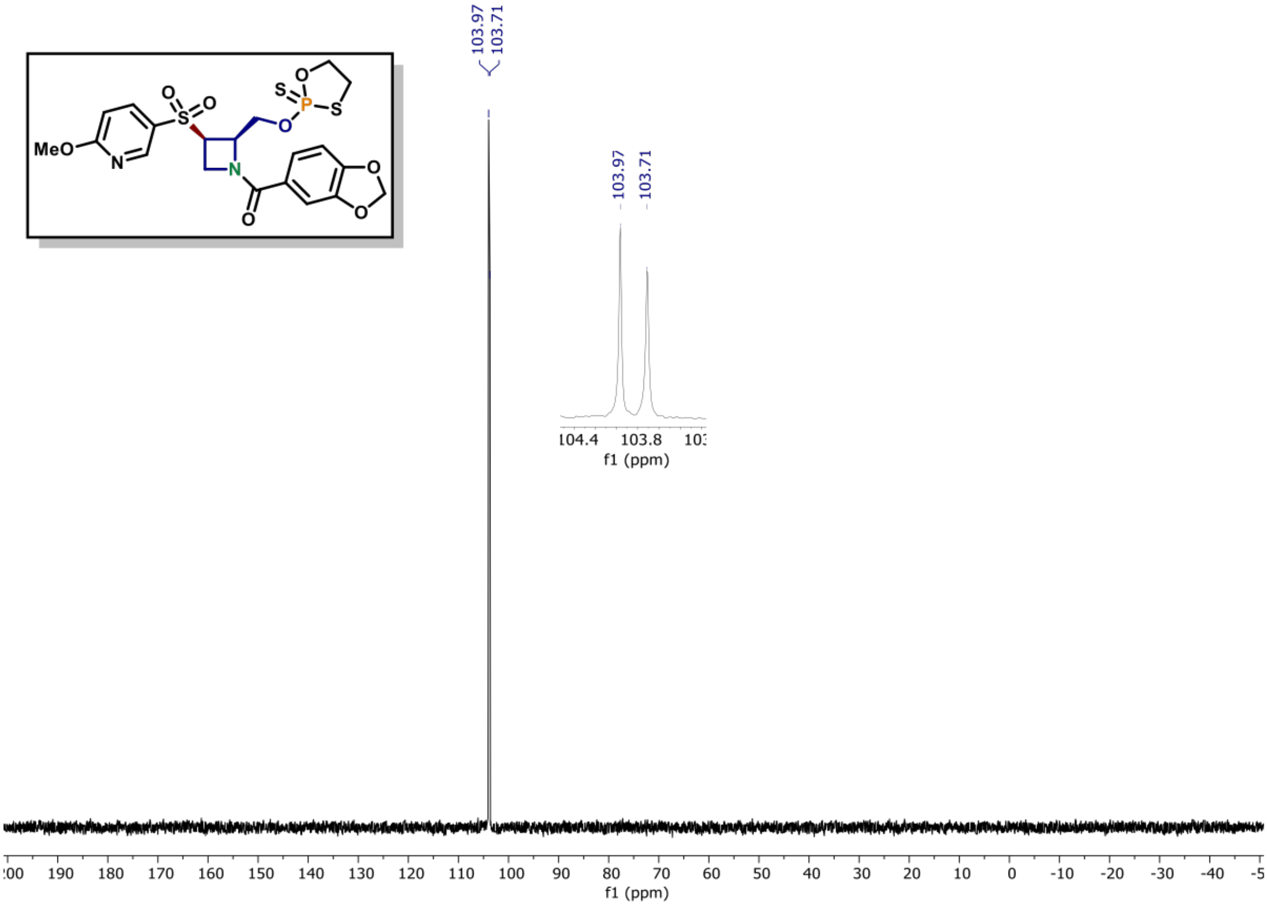

**[1]H NMR of compound (5) (600 MHz, $CDCl_3$)**

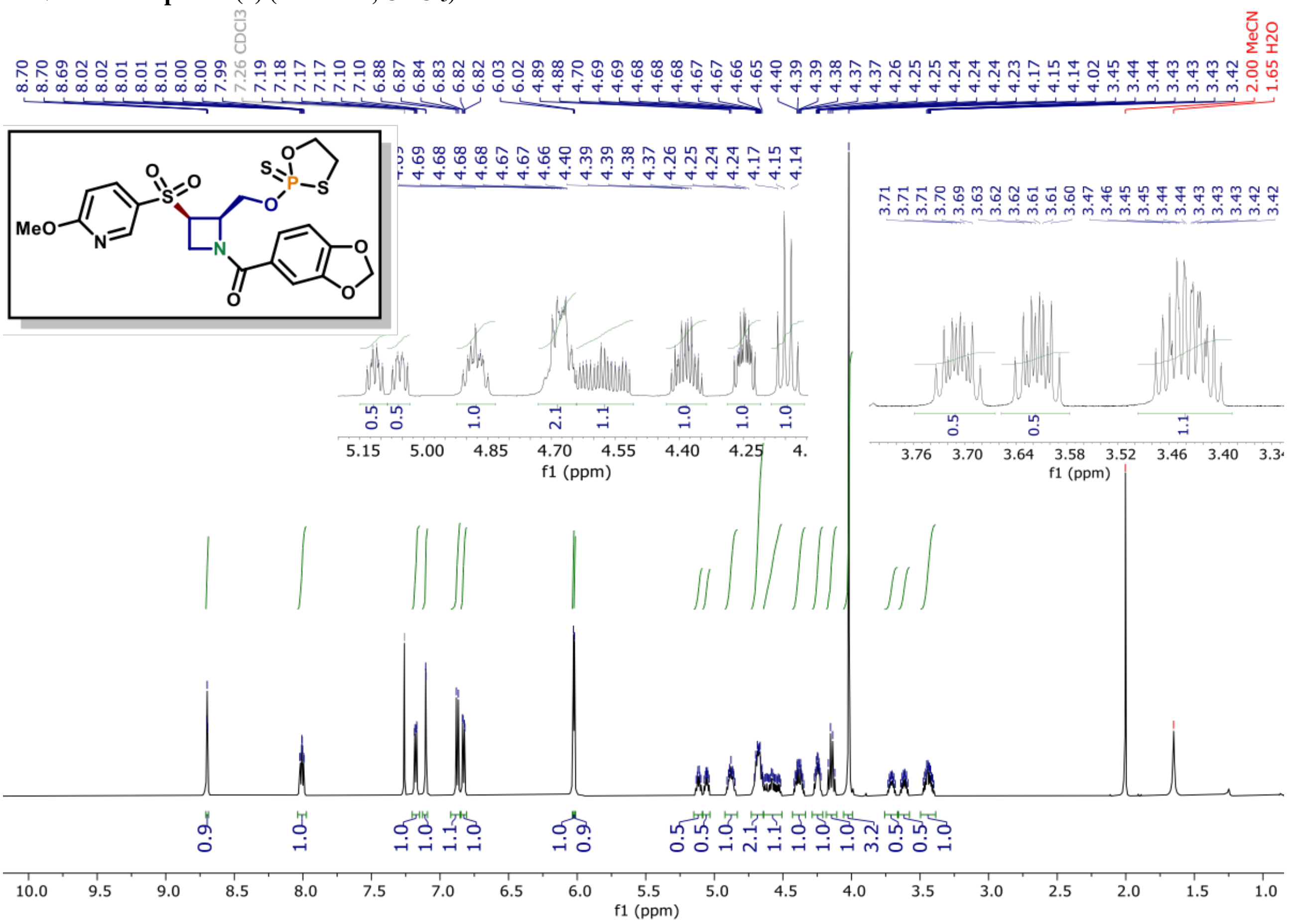

$^{13}$C NMR of compound (5) (150 MHz, CDCl$_3$)

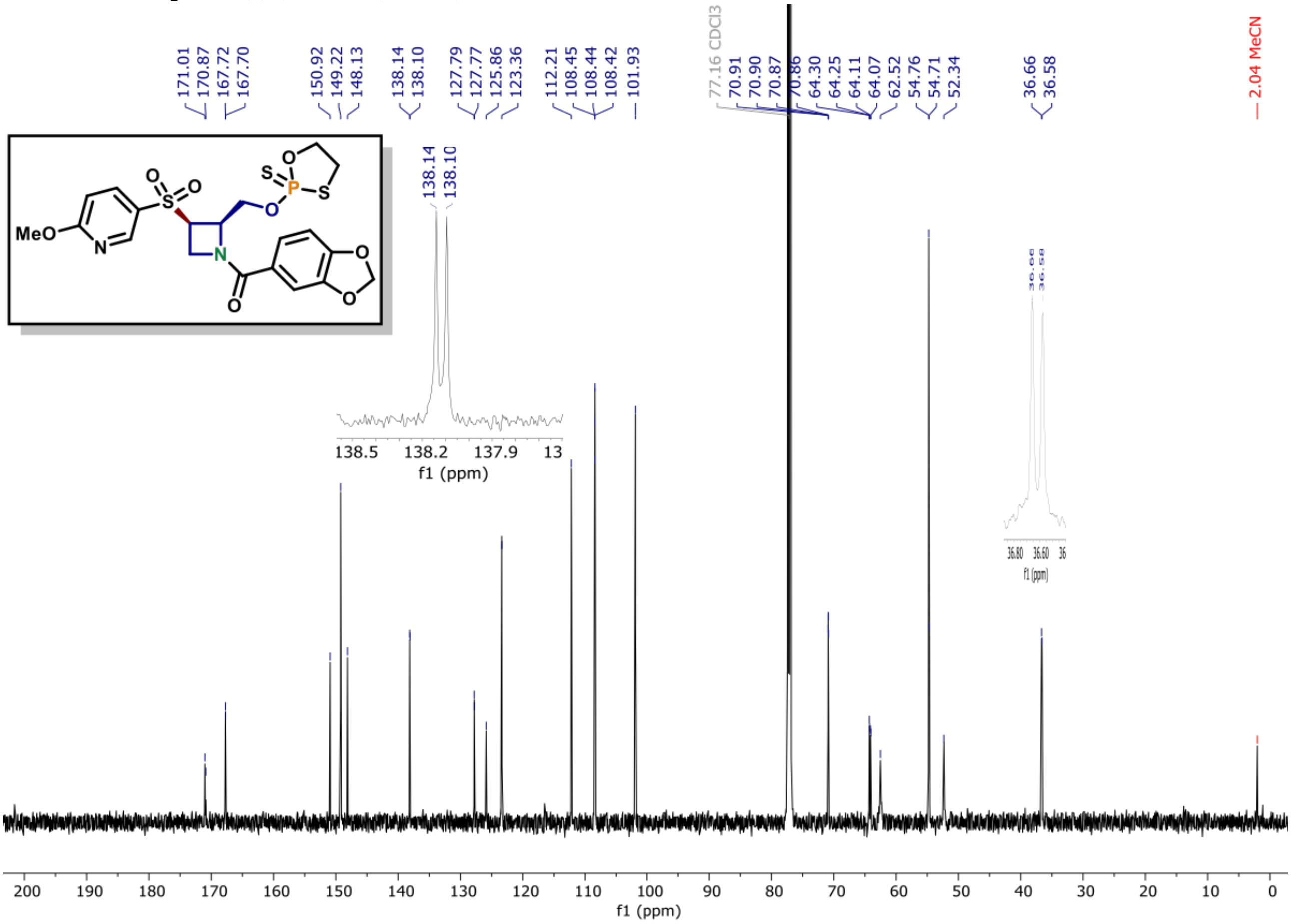

[31]P NMR of compound (7) (162 MHz, CDCl3)

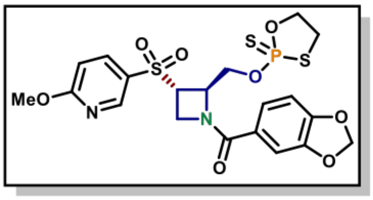

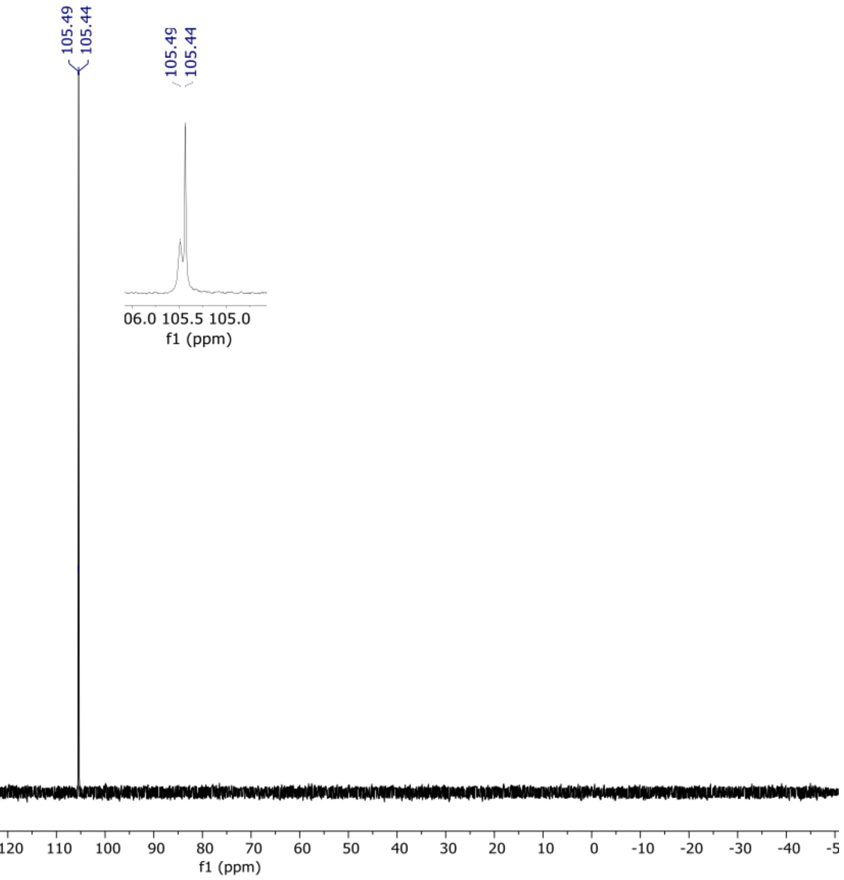

[1]H NMR of compound (7) (600 MHz, CDCl3)

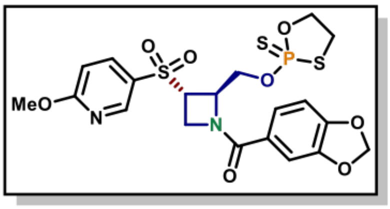

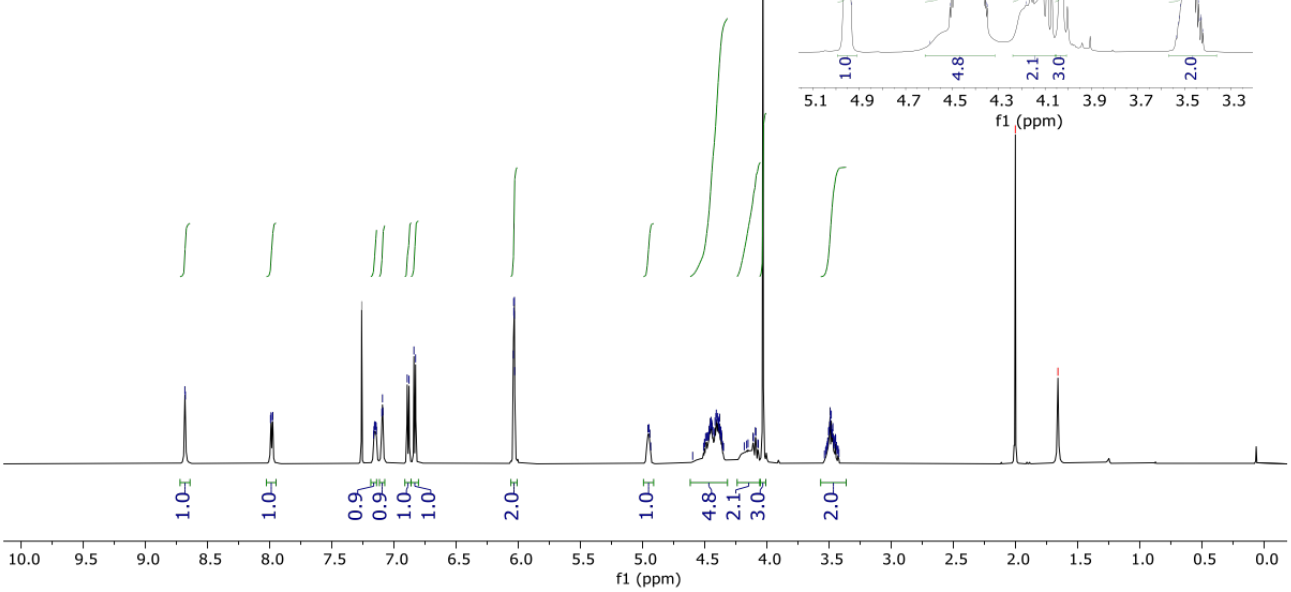

[13]C NMR of compound (7) (150 MHz, $CDCl_3$)

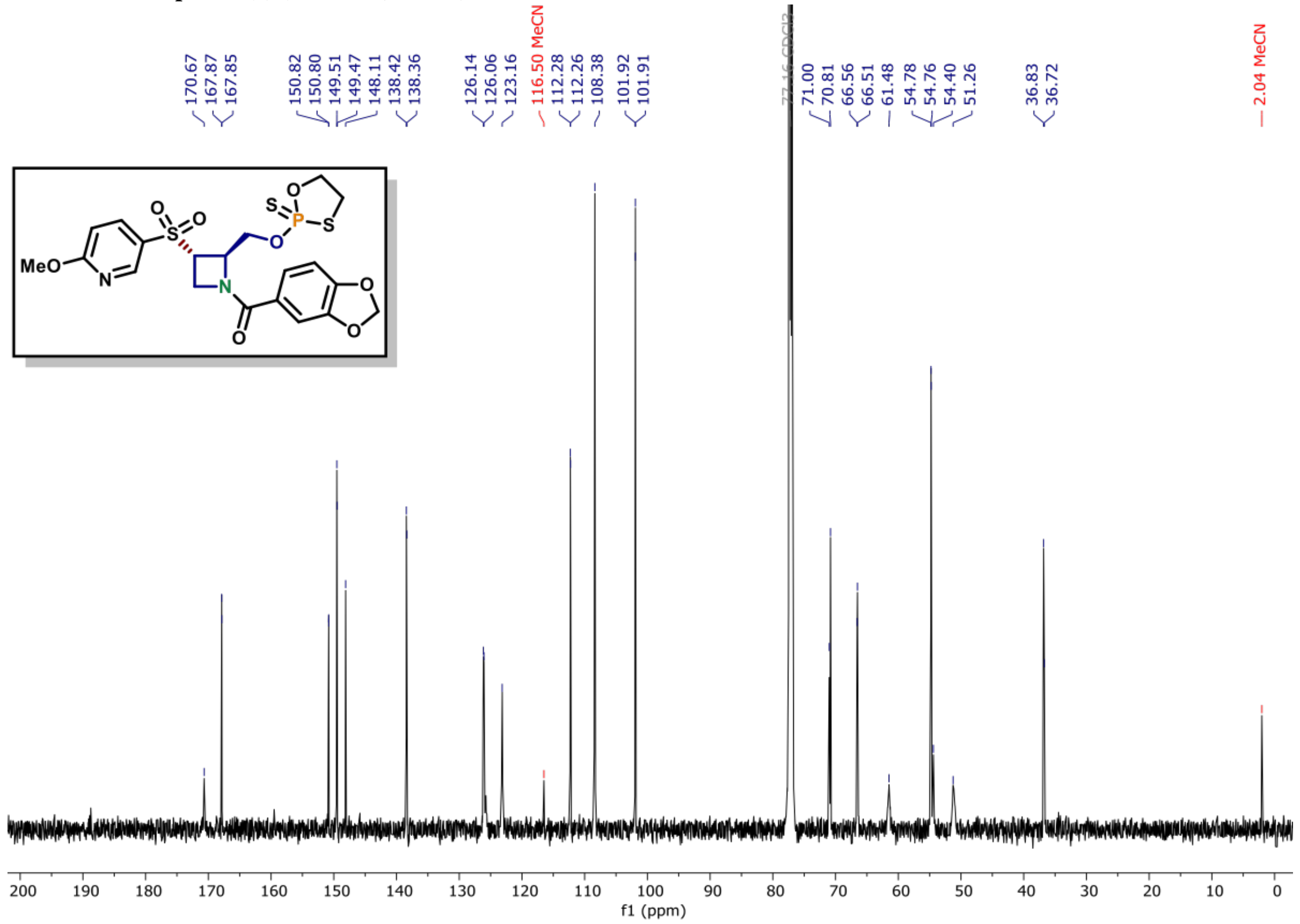

$^{31}$P NMR of compound (6) (162 MHz, CDCl$_3$)

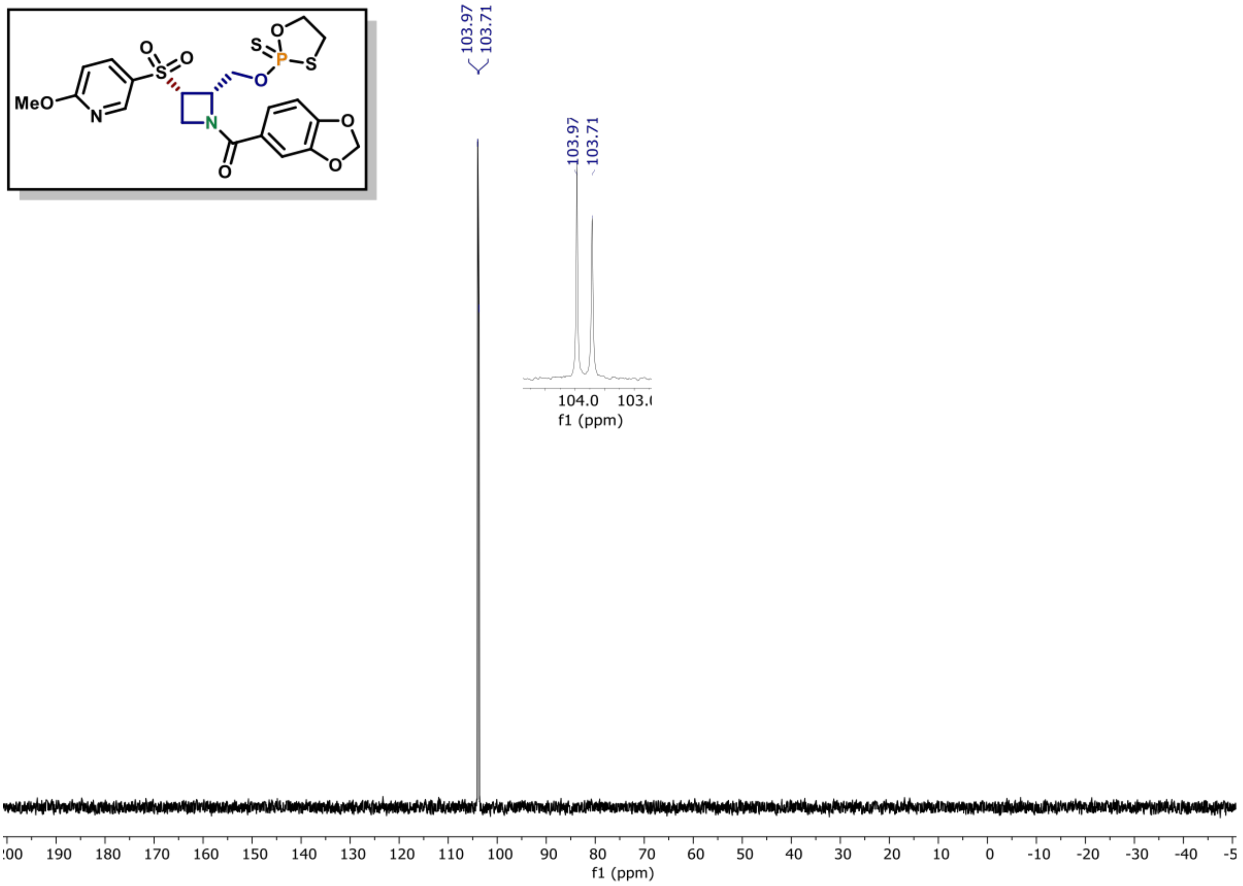

$^{1}$H NMR of compound (6) (600 MHz, CDCl$_3$)

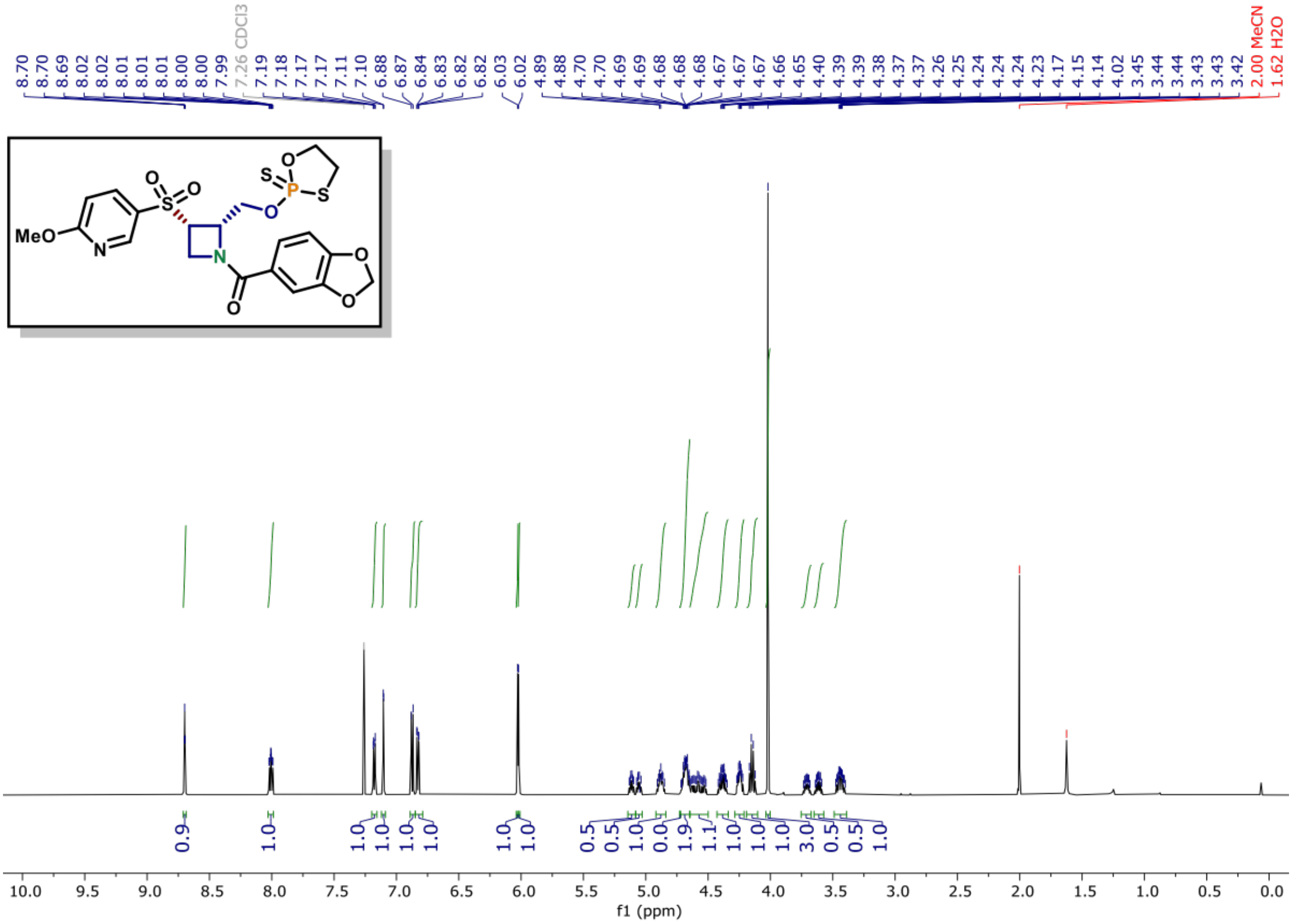

$^{13}$C NMR of compound (6) (150 MHz, CDCl$_3$)

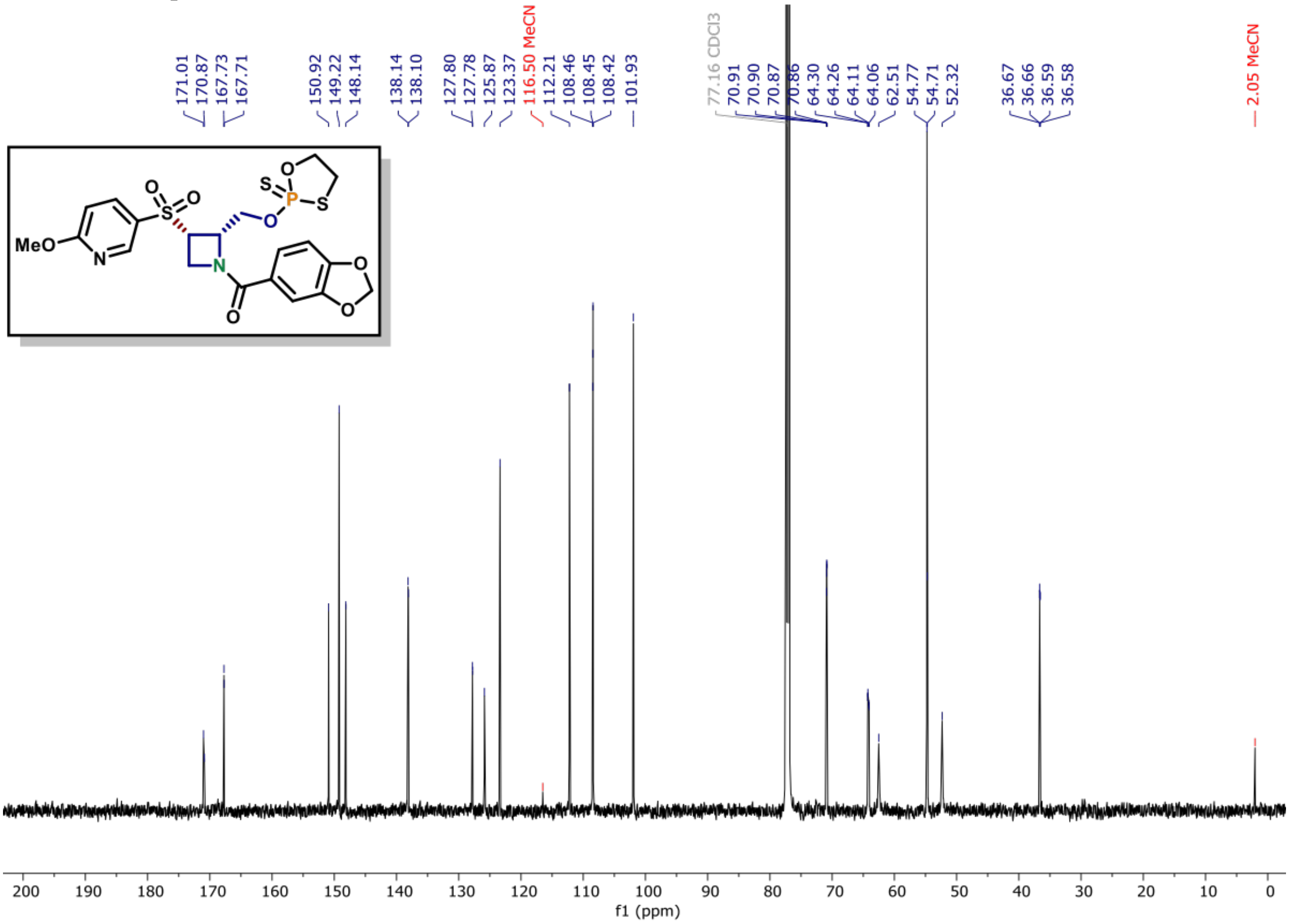

**$^{31}$P NMR of compound (8) (162 MHz, $CDCl_3$)**

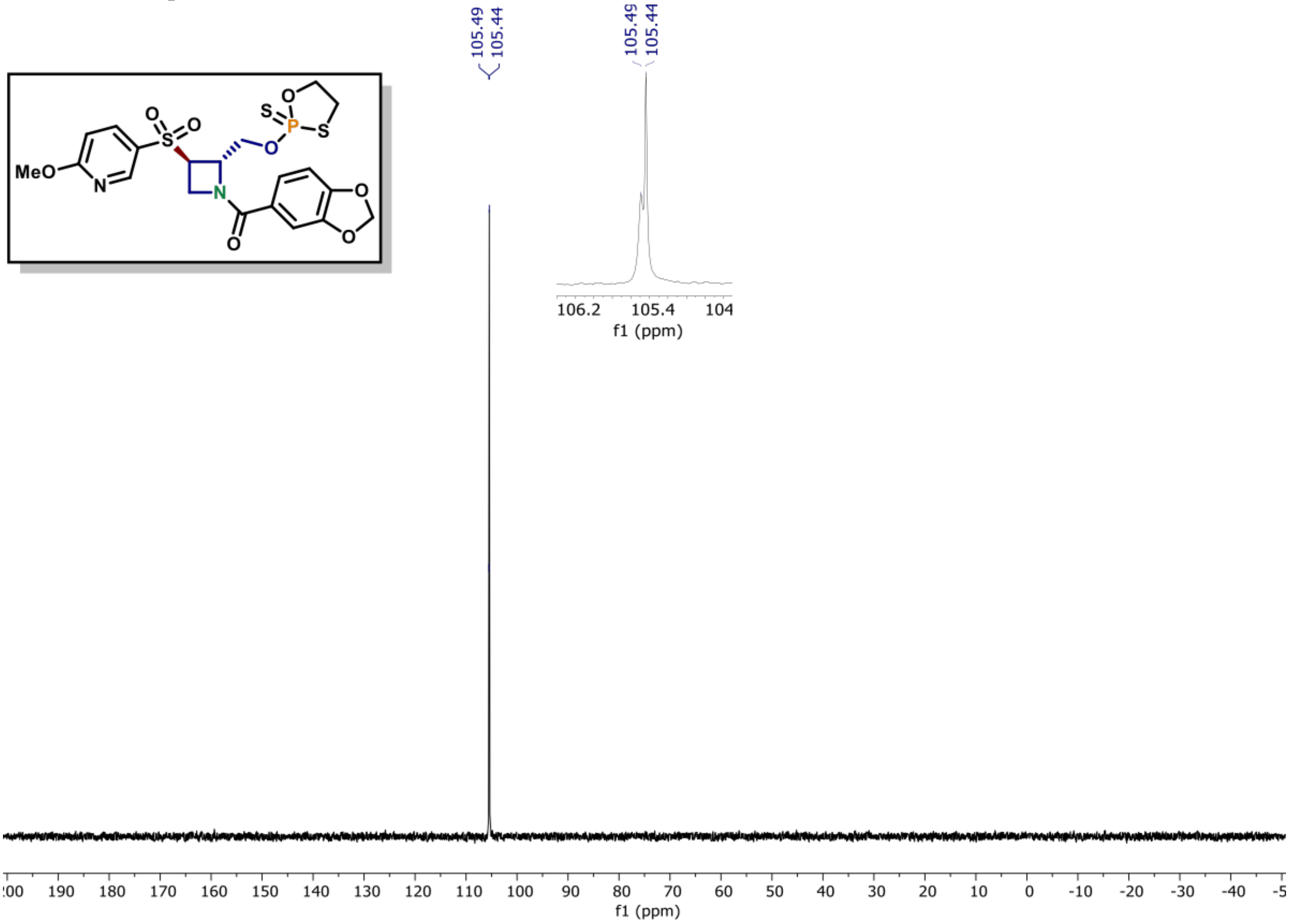

**$^1$H NMR of compound (8) (600 MHz, $CDCl_3$)**

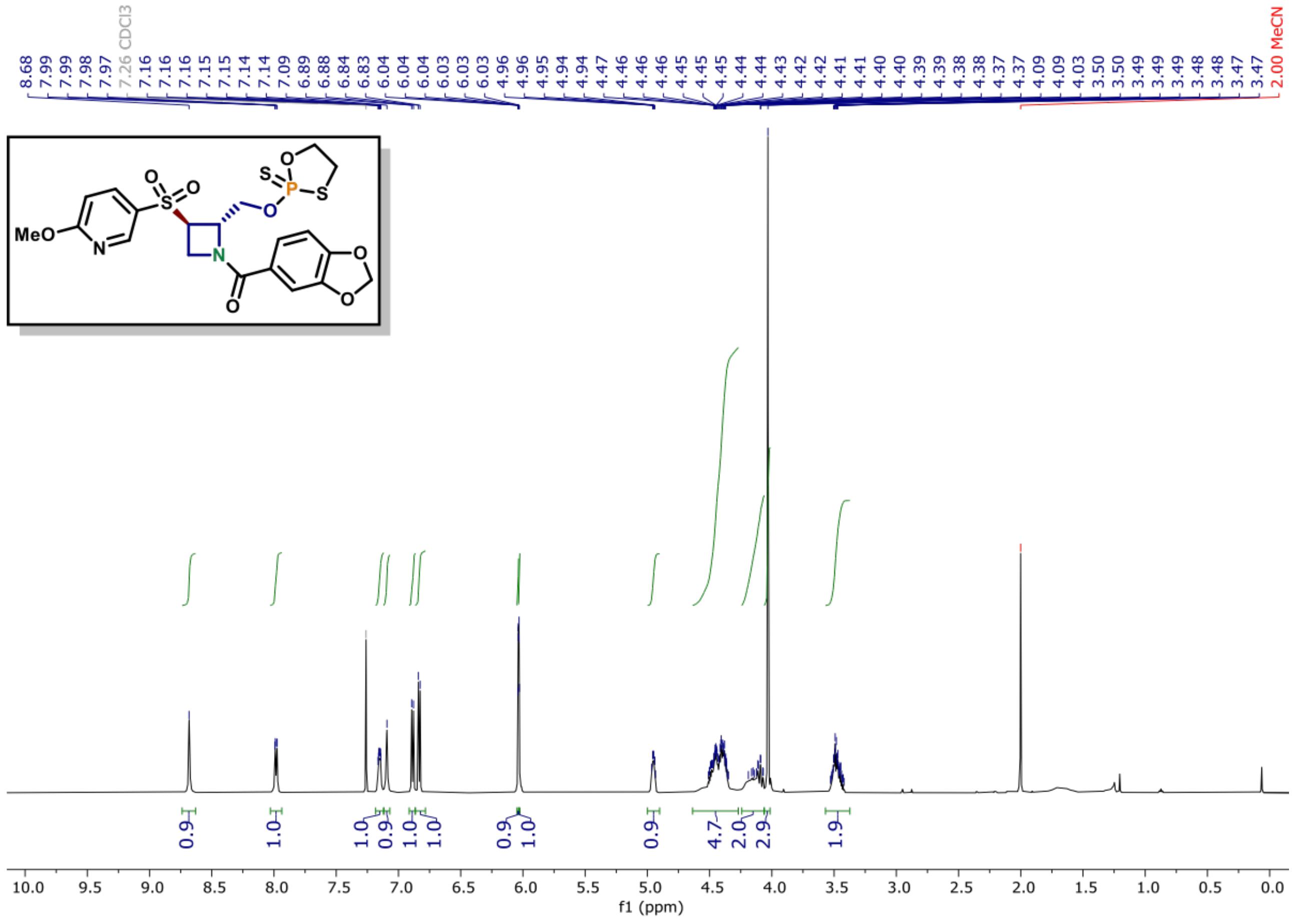

**$^{13}$C NMR of compound (8) (150 MHz, $CDCl_3$)**

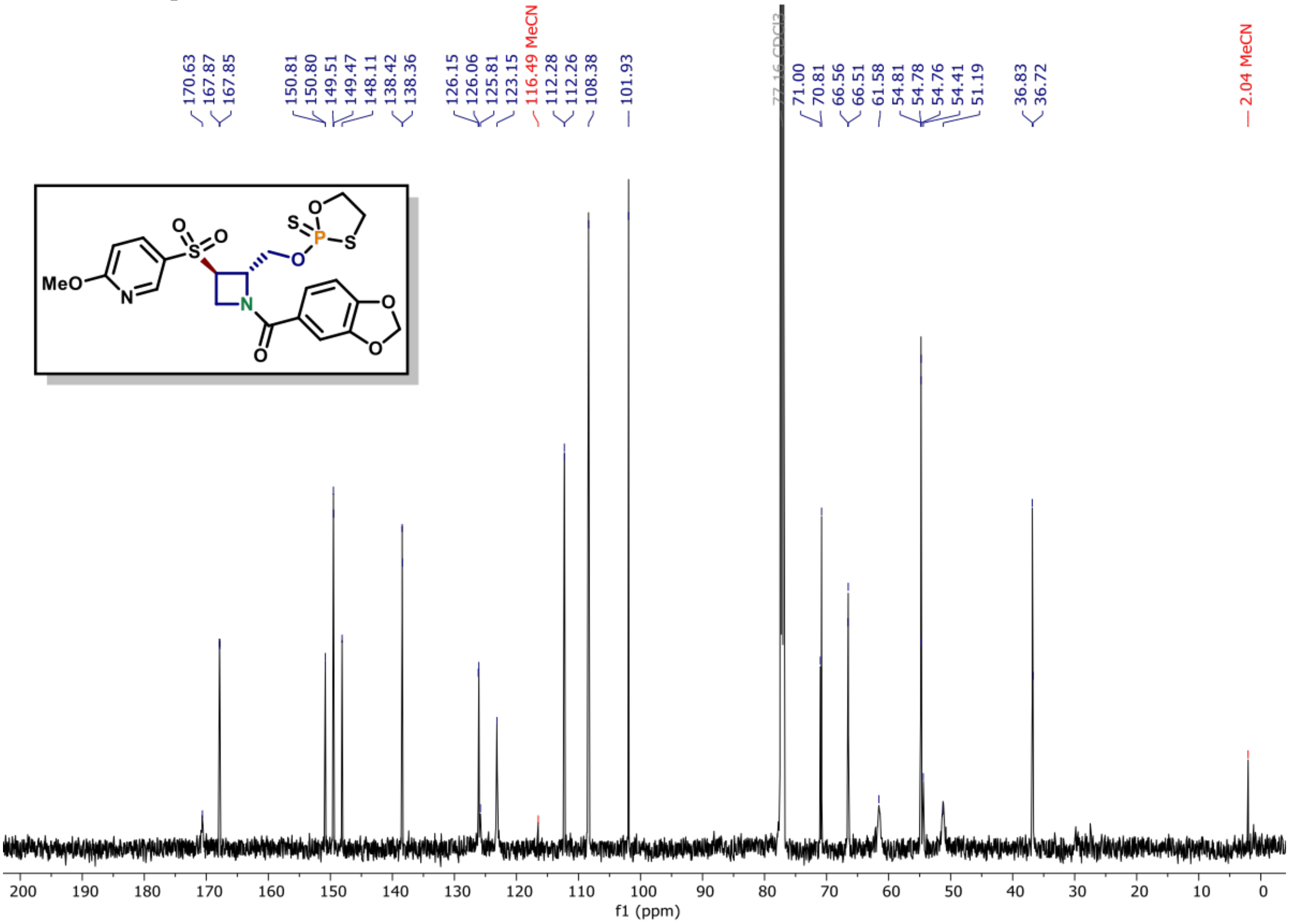

### Chiral Column Analysis

#### Compounds MB-5A & MB-5B

##### Method:

*The analysis was performed on a Waters UPC2 SFC with a* ***Daicel IBN*** *column (3 µm, 4.6x100 mm) under isocratic conditions (3.3 mL/min, 50% MeOH / $CO_2$, 1600 psi backpressure) at 30 °C. The enantiomers were detected by UV light (220 nm).*

##### Summary:

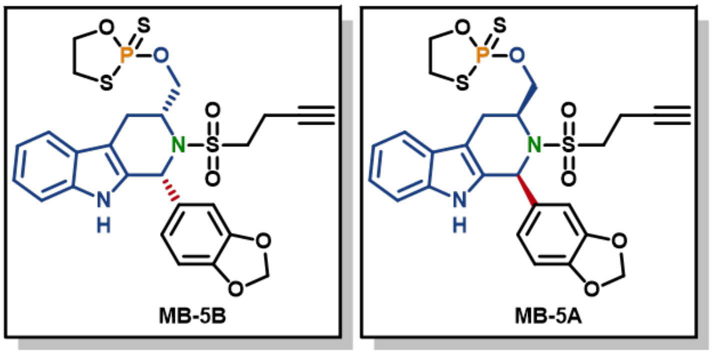

e.e. = 99.6% e.e. = 100%

Peak Areas Summary
Area Summarized by Name

| | SampleName | Vial | Date Acquired | Ent 1_Ep1 (µV*sec) | Ent 1_Ep2 (µV*sec) | Ent 2 (µV*sec) |
|---|---|---|---|---|---|---|
| 1 | MB-5B | 1:C,4 | 5/21/2024 8:03:56 PM PDT | 1053470 | 1226168 | 4242 |
| 2 | MB-5A | 1:C,5 | 5/21/2024 8:08:05 PM PDT | | | 2639538 |

##### Compound MB-5A :

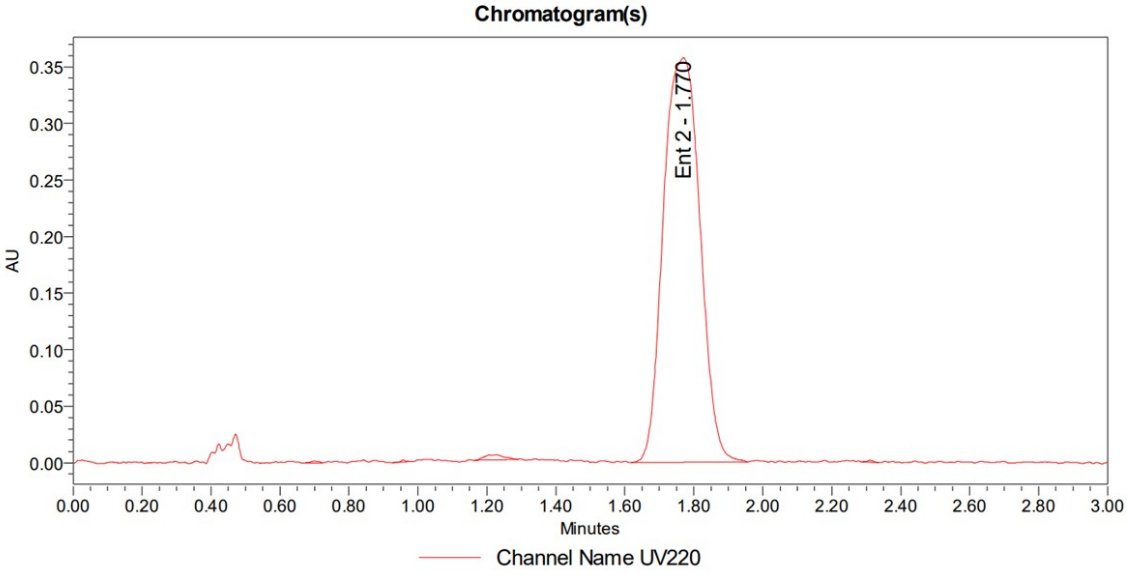

##### Compound MB-5B:

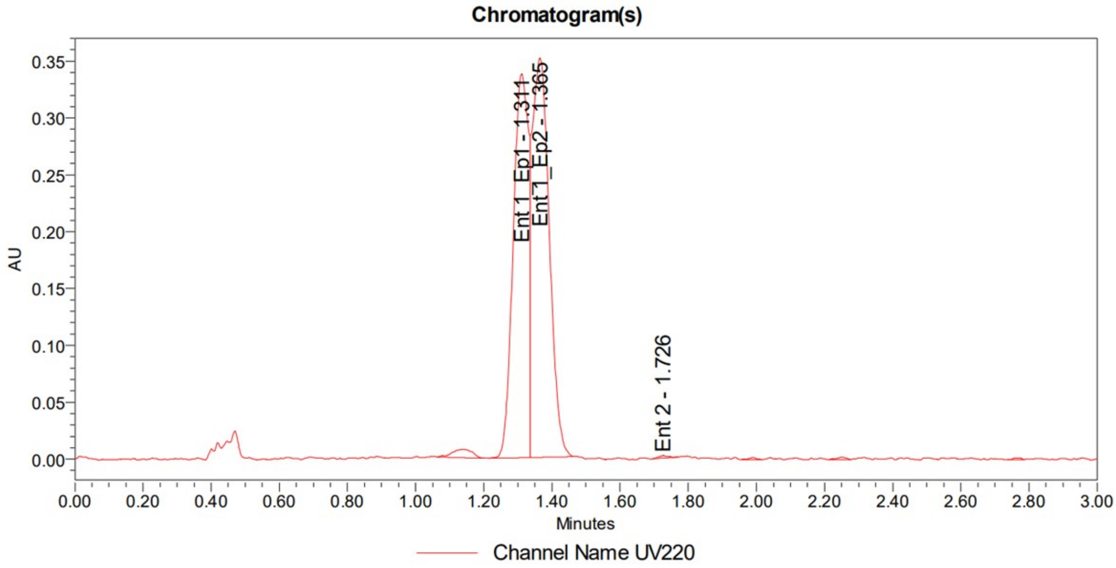

**Mixture (MB-5A + MB-5B ):**

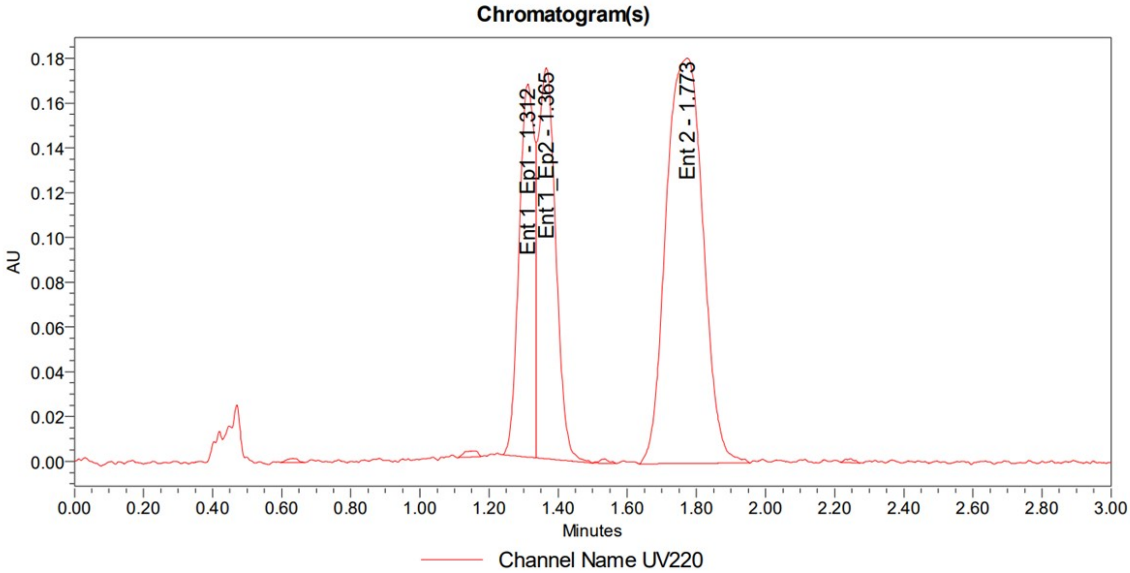

**Compounds MB-6A & MB-6B**

**Method:**

*The analysis was performed on a Waters UPC2 SFC with a* ***Daicel IBN*** *column (3 µm, 4.6x100 mm) under isocratic conditions (3.3 mL/min, 50% MeOH / $CO_2$, 1600 psi backpressure) at 30 °C. The enantiomers were detected by UV light (220 nm).*

**Summary:**

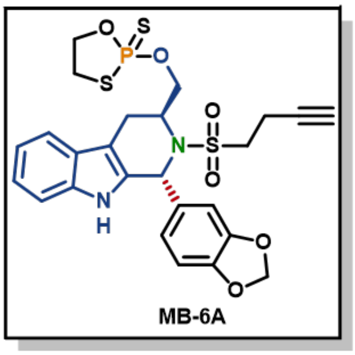

e.e. = 95.9%

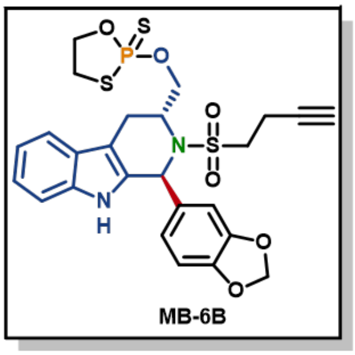

e.e. = 97.6%

Peak Areas Summary
Area Summarized by Name

| | SampleName | Vial | Date Acquired | Ent 1_Ep1 (µV*sec) | Ent 1_Ep2 (µV*sec) | Ent 2_Ep1 (µV*sec) | Ent 2_Ep2 (µV*sec) |
|---|---|---|---|---|---|---|---|
| 1 | MB-6A | 1:C,1 | 5/21/2024 7:51:24 PM PDT | 900818 | 868516 | 37368 | |
| 2 | MB-6B | 1:C,2 | 5/21/2024 7:55:33 PM PDT | | 23142 | 914251 | 969100 |

**Compound MB-6A:**

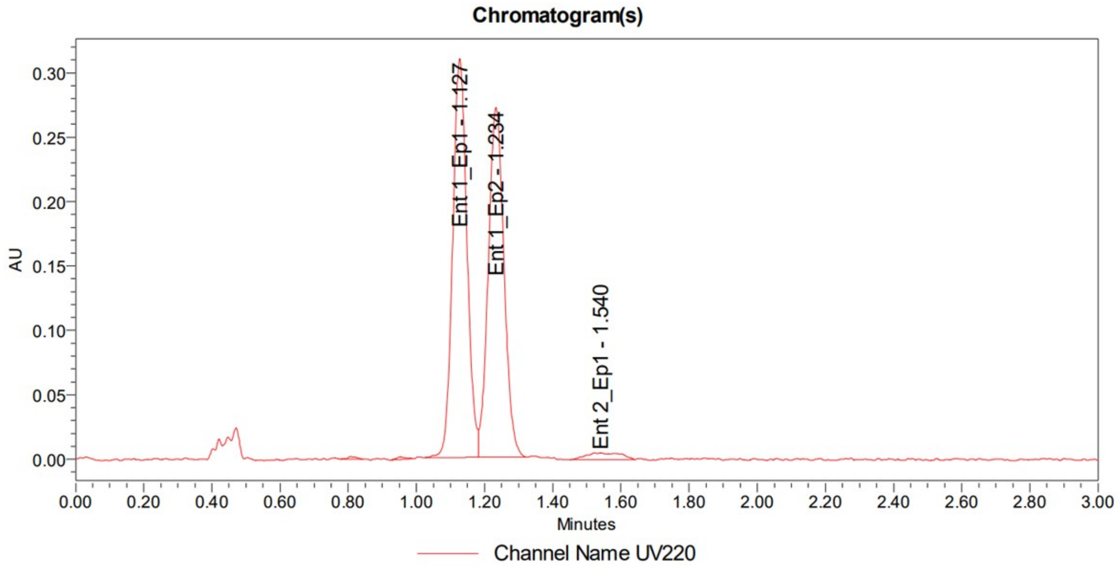

**Compound MB-6B:**

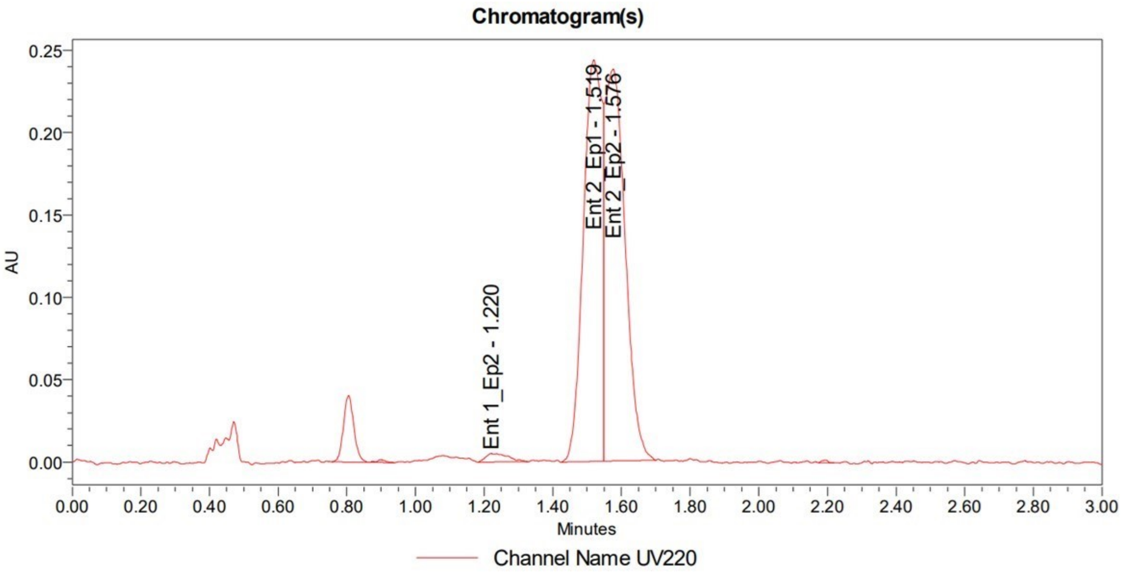

**Mixture (MB-6A + MB-6B):**

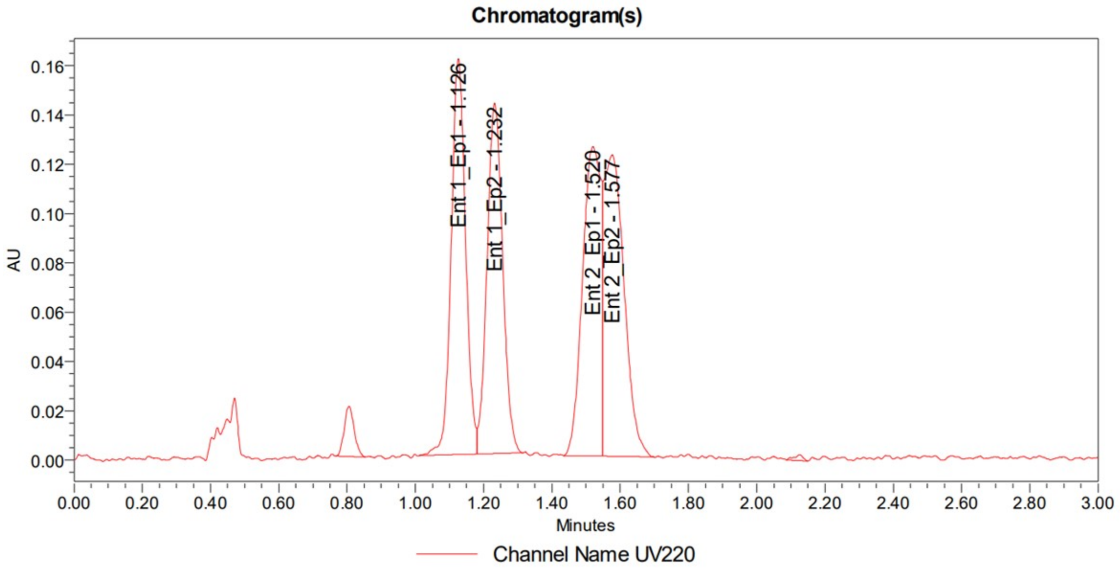

**Compounds MB5A & MB-5B**

**Method:**

*The analysis was performed on a Waters UPC2 SFC with a* ***Daicel IA*** *column (3 µm, 4.6x100 mm) under isocratic conditions (3.3 mL/min, 55% MeOH / $CO_2$, 1600 psi backpressure) at 30 °C. The enantiomers were detected by UV light (218 nm).*

**Summary:**

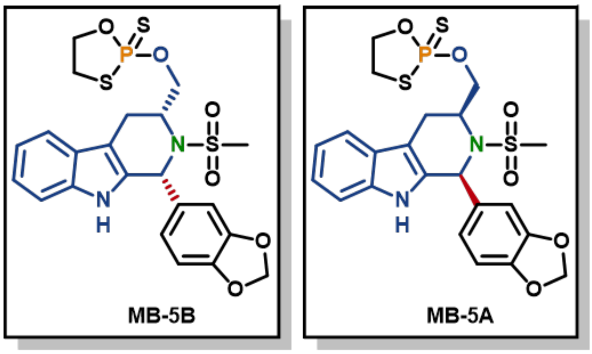

e.e. = 100%    e.e. = 100%

Peak Areas Summary
Area Summarized by Name

| | SampleName | Vial | Date Acquired | Epimer 1 (µV*sec) | Epimer 2 (µV*sec) | Epimer_1 (µV*sec) | Epimer_2 (µV*sec) |
|---|---|---|---|---|---|---|---|
| 2 | MB-5B | 1:B,2 | 5/21/2024 6:45:58 PM PDT | 1322770 | 1459302 | | |
| 3 | MB-5A | 1:B,3 | 5/21/2024 6:50:31 PM PDT | | | 1116522 | 1188082 |

**Compound MB-5A:**

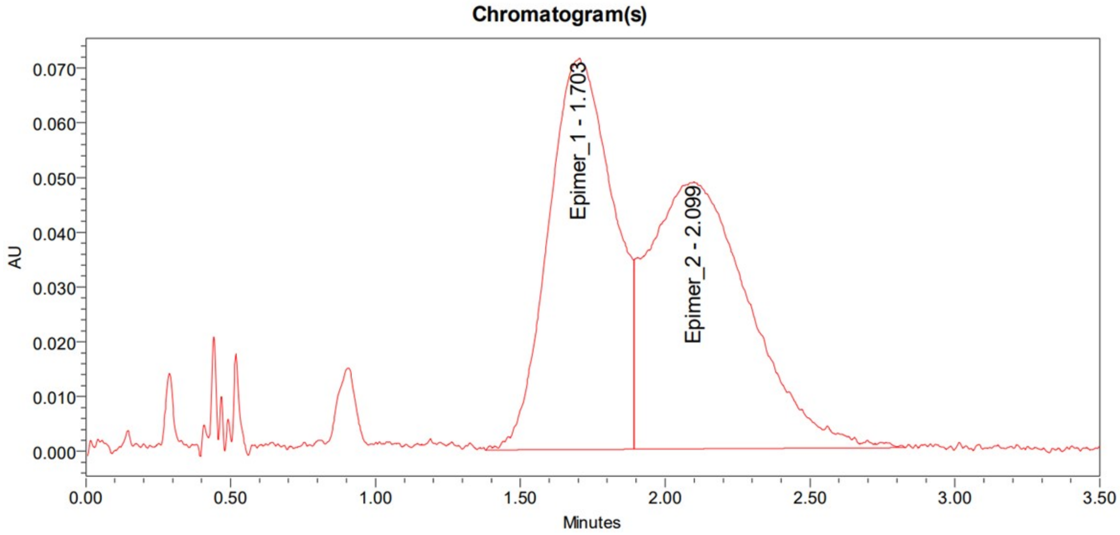

**Compound MB-5B:**

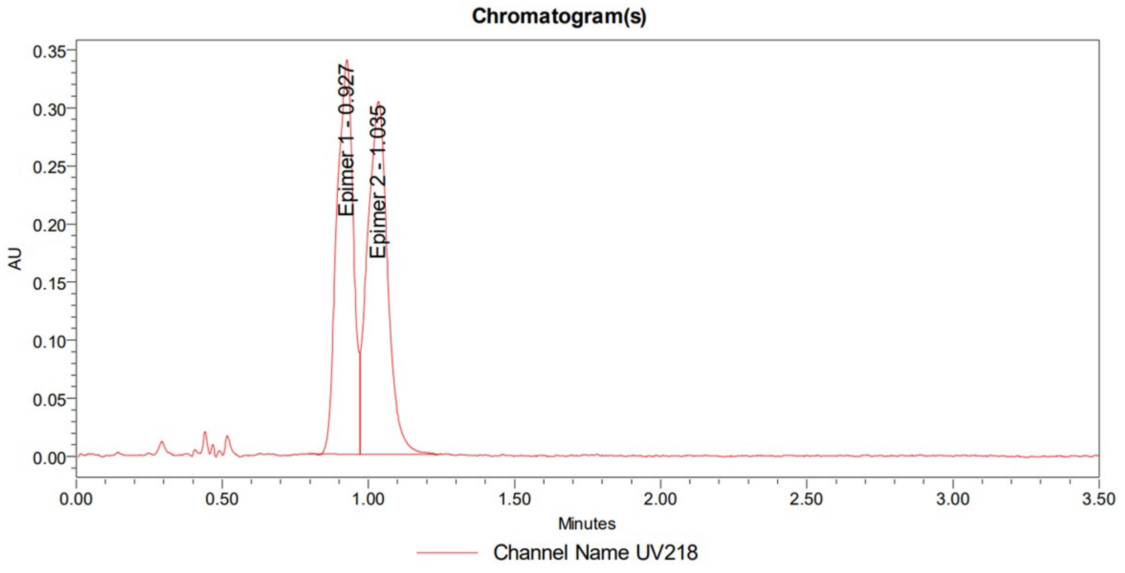

**Mixture (MB-5A + MB-5B):**

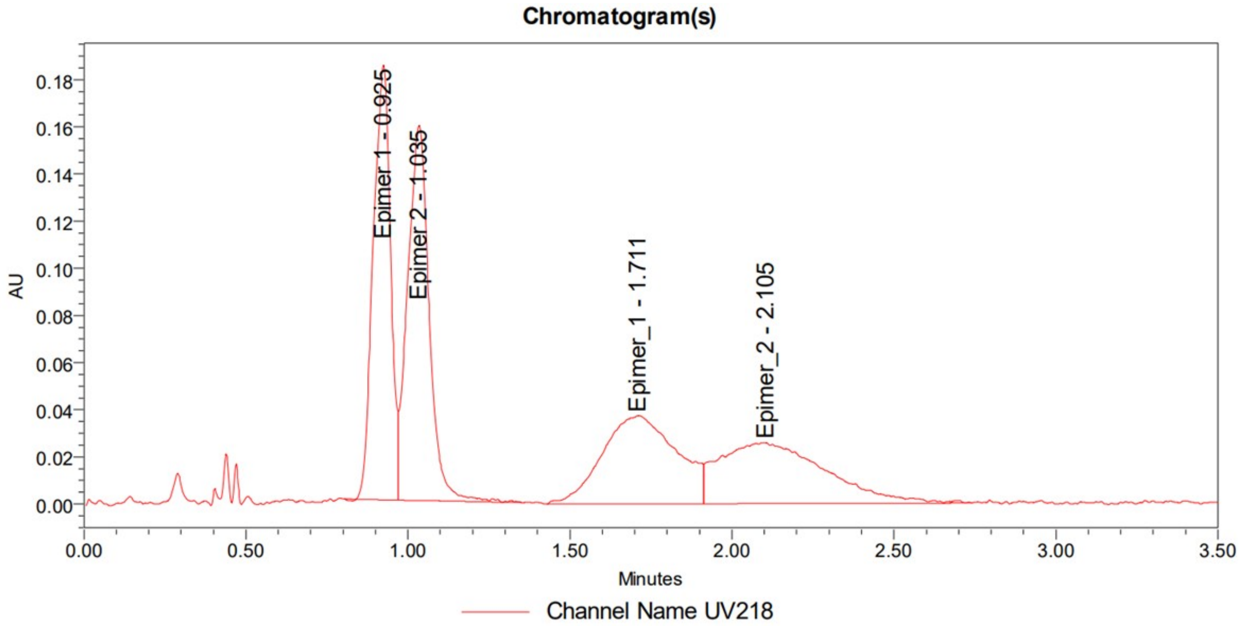

**Compounds MB-8A & MB-8B**

**Method:**

*The analysis was performed on a Waters UPC2 SFC with a **Daicel IH** column (3 µm, 4.6x100 mm) under isocratic conditions (3.3 mL/min, 45% MeOH / $CO_2$, 1600 psi backpressure) at 30 °C. The enantiomers were detected by UV light (212 nm).*

**Summary:**

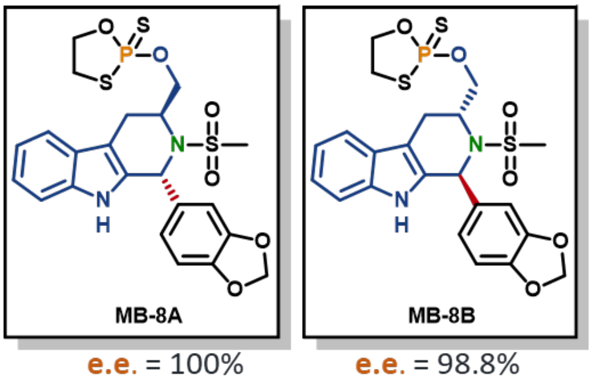

Peak Areas Summary
Area Summarized by Name

| | SampleName | Vial | Date Acquired | Ent 1 (µV*sec) | Ent 2_Ep1 (µV*sec) | Ent 2_Ep2 (µV*sec) |
|---|---|---|---|---|---|---|
| 1 | MB-8A | 1:C,7 | 5/21/2024 8:16:27 PM PDT | 5369668 | | |
| 2 | MB-8B | 1:C,8 | 5/21/2024 8:20:37 PM PDT | 37768 | 3207666 | 3477740 |

**Compound MB-8A:**

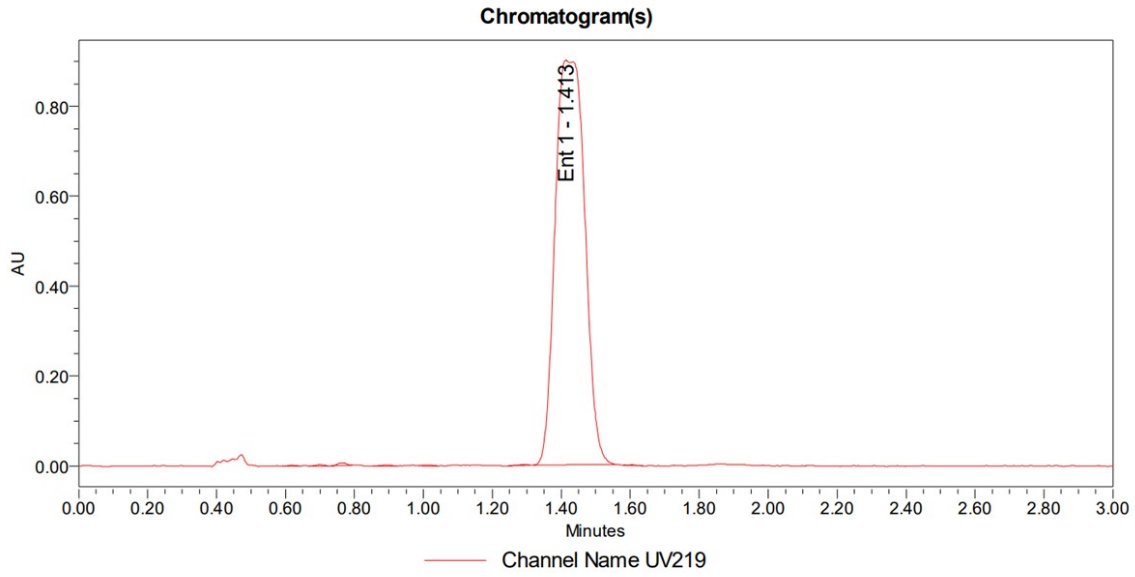

**Compound MB-8B:**

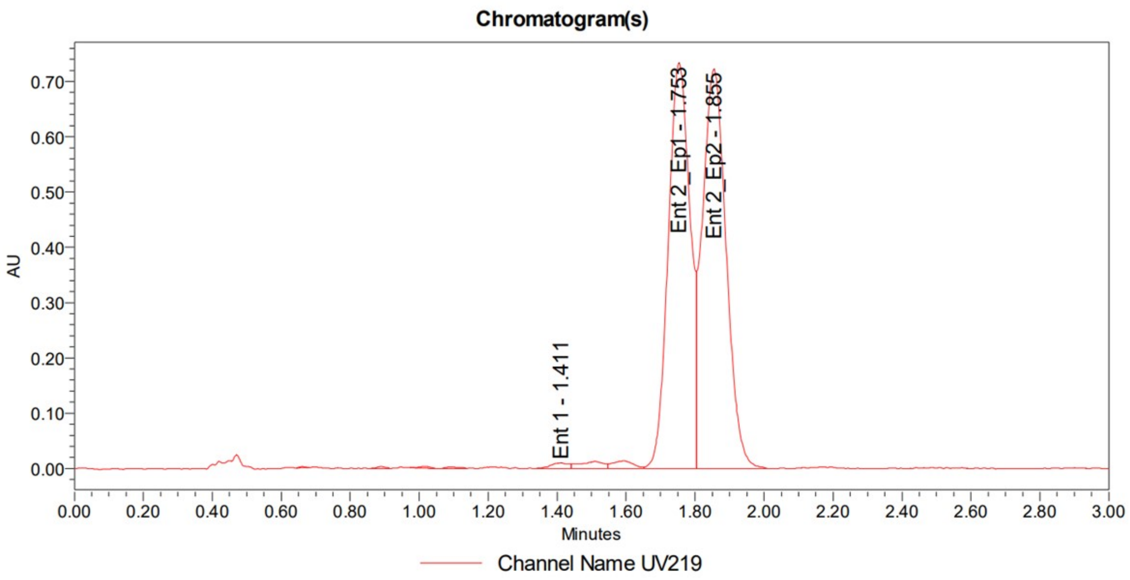

**Mixture (MB-8A + MB-8B):**

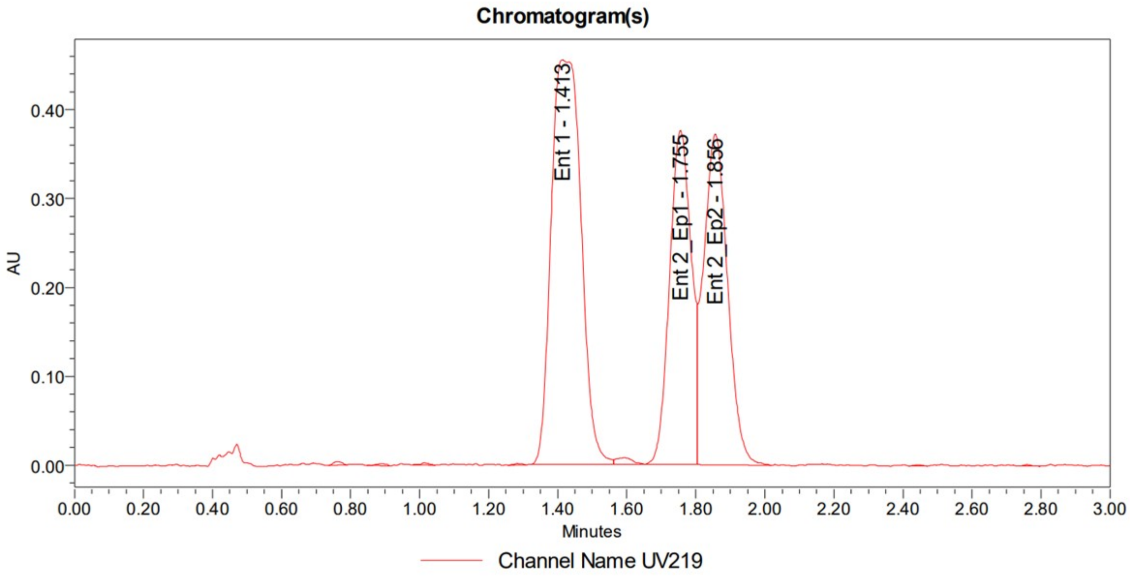

**Compounds MB-1A & MB-1B**

**Method:**

*The analysis was performed on a Waters UPC2 SFC with a* ***Phenomenex CEL-1*** *column (3 µm, 4.6x100 mm) under isocratic conditions (3.3 mL/min, 45% MeOH / $CO_2$, 1600 psi backpressure) at 30 °C. The enantiomers were detected by UV light (212 nm).*

**Summary:**

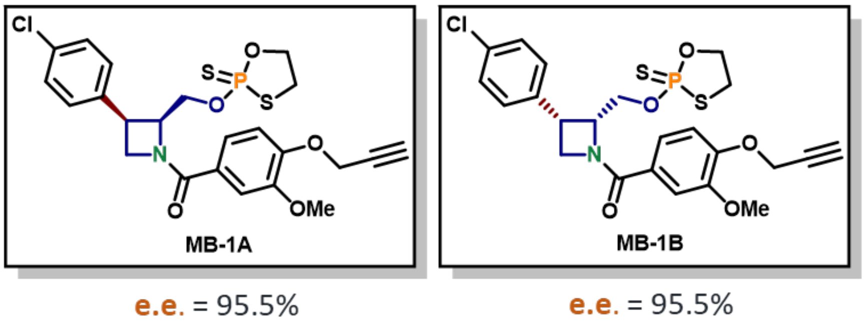

Peak Areas Summary
Area Summarized by Name

| | SampleName | Vial | Date Acquired | Ent1 (µV*sec) | Ent2 (µV*sec) |
|---|---|---|---|---|---|
| 1 | MB-1A | 1:B,1 | 5/14/2024 12:52:30 PM PDT | 1962775 | 44841 |
| 2 | MB-1B | 1:B,2 | 5/14/2024 12:56:34 PM PDT | 26676 | 1169460 |

**Compound MB-1A:**

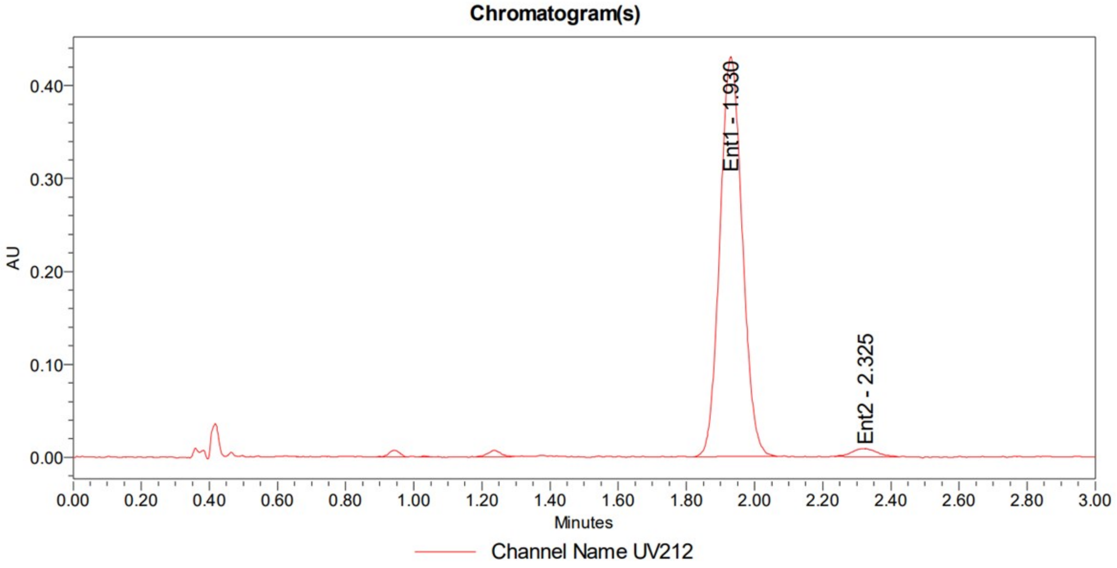

Compound **MB-1B**:

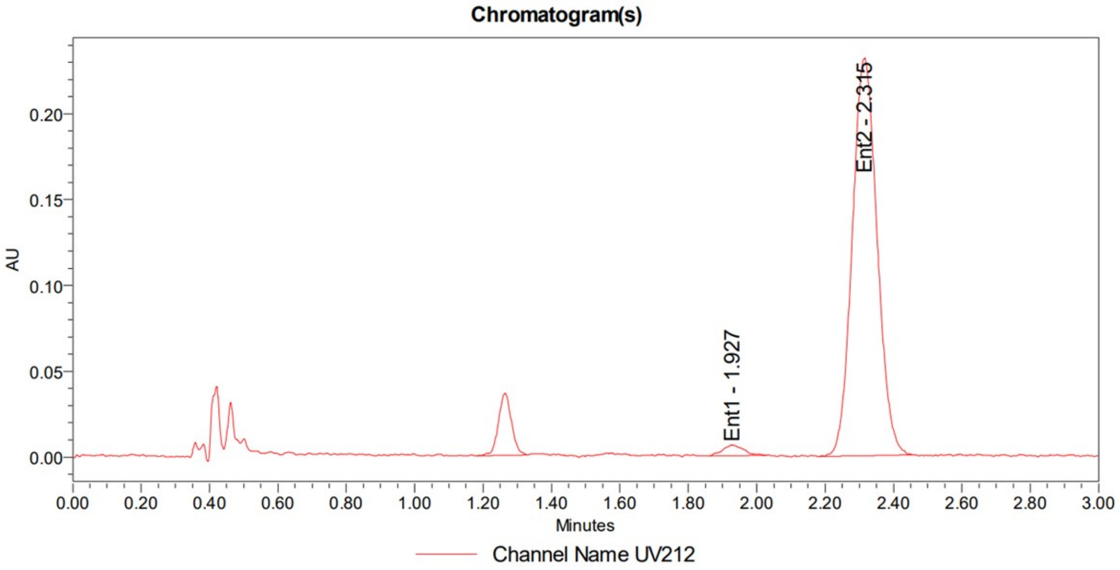

Mixture (**MB-1A** + **MB-1B**):

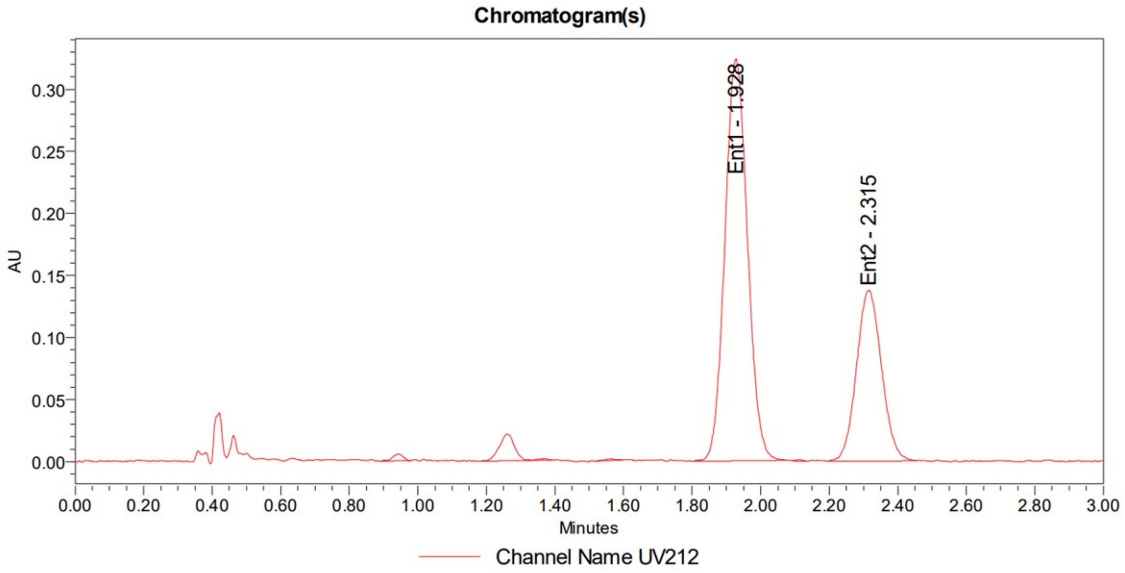

**Compounds MB-2A & MB-2B**

**Method:**

The analysis was performed on a Waters UPC2 SFC with a Daicel IH column (3 µm, 4.6x100 mm) under isocratic conditions (3.3 mL/min, 45% MeOH / $CO_2$, 1600 psi backpressure) at 30 °C. The enantiomers were detected by UV light (212 nm).

**Summary:**

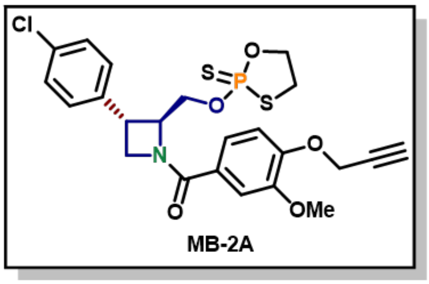

e.e. = 99.3%

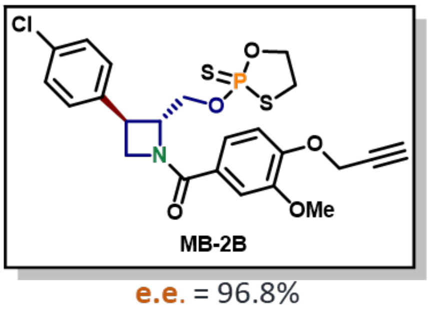

e.e. = 96.8%

Peak Areas Summary
Area Summarized by Name

| | SampleName | Vial | Date Acquired | Ent1_Ep1 (µV*sec) | Ent1_Ep2 (µV*sec) | Ent2_Ep1 (µV*sec) | Ent2_Ep2 (µV*sec) |
|---|---|---|---|---|---|---|---|
| 1 | MB-2A | 1:C,1 | 5/14/2024 11:16:45 AM PDT | 1853 | 2892 | 682909 | 726012 |
| 2 | MB-2B | 1:C,2 | 5/14/2024 11:20:49 AM PDT | 750127 | 857169 | 13084 | 12846 |

**Compound MB-2A:**

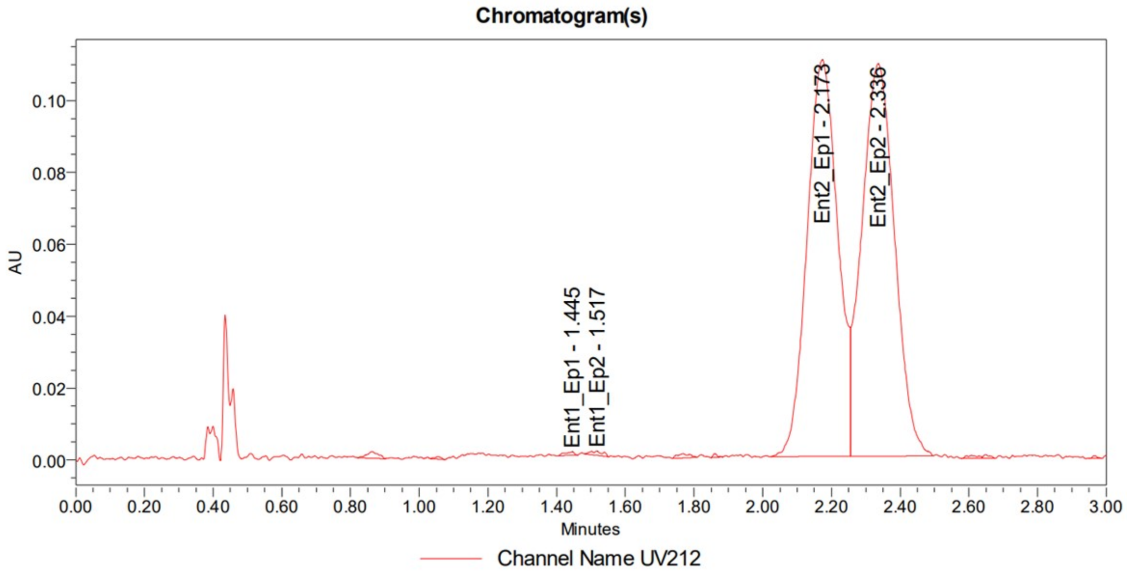

**Compound MB-2B:**

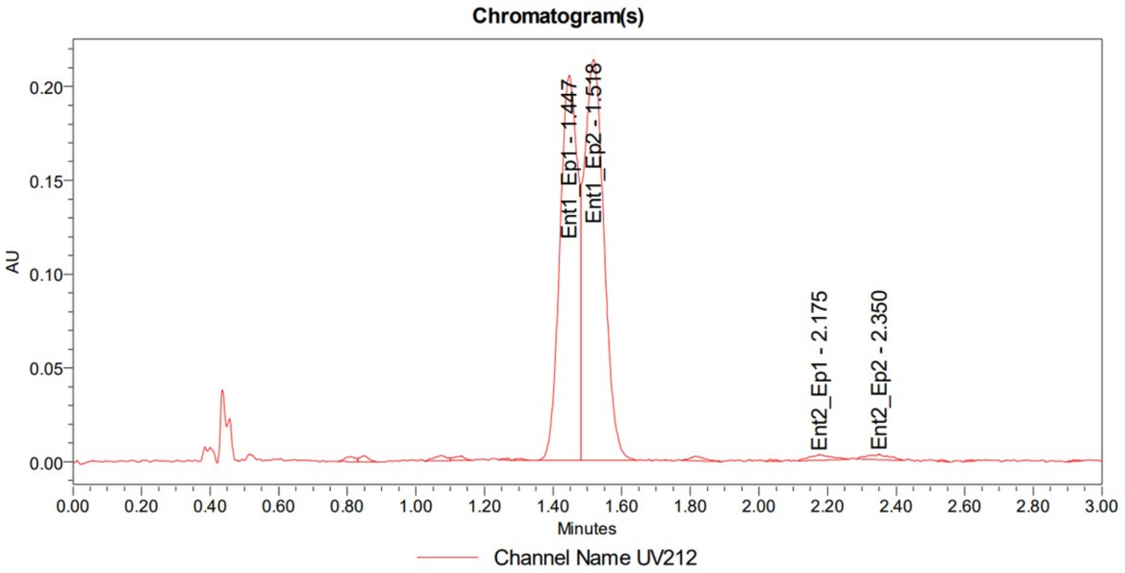

**Mixture (MB-2A + MB-2B):**

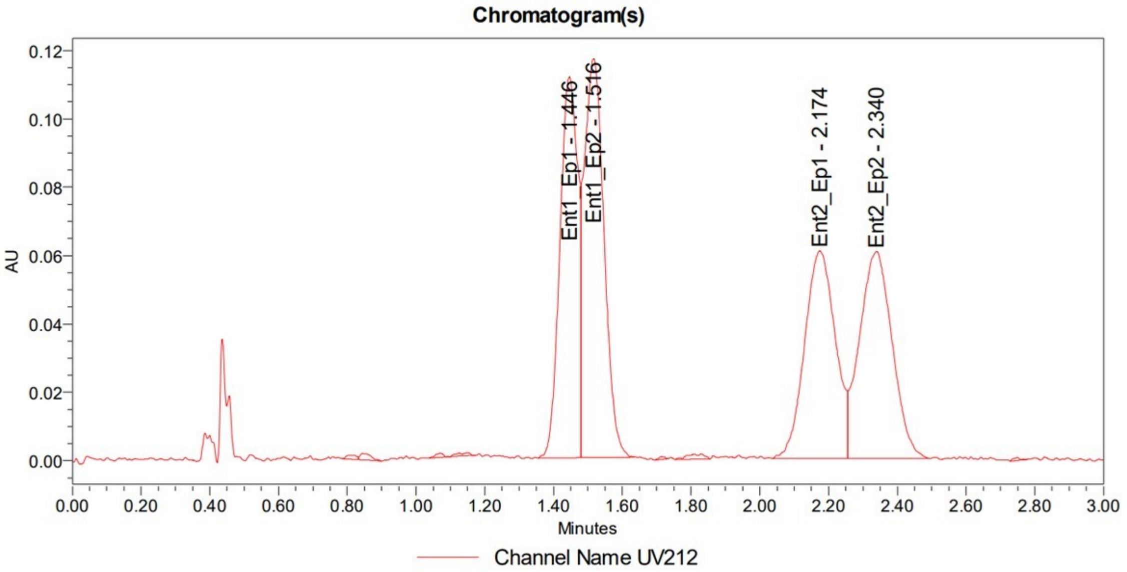

**Compounds MB-3A & MB-3B**

**Method:**

*The analysis was performed on a Waters UPC2 SFC with a Daicel IA column (3 µm, 4.6x100 mm) under isocratic conditions (3.3 mL/min, 45% MeOH / CO2, 1600 psi backpressure) at 30 °C. The enantiomers were detected by UV light (212 nm).*

**Summary:**

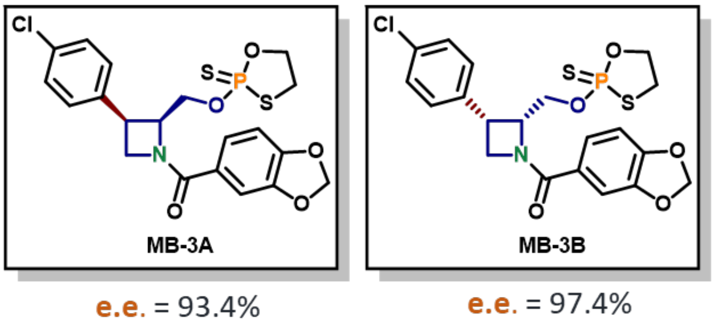

Peak Areas Summary
Area Summarized by Name

| | SampleName | Vial | Date Acquired | Ent1 (µV*sec) | Ent2_Ep1 (µV*sec) | Ent2_Ep2 (µV*sec) |
|---|---|---|---|---|---|---|
| 1 | MB-3A | 1:D,1 | 5/14/2024 1:33:51 PM PDT | 56820 | 784865 | 905336 |
| 2 | MB-3B | 1:D,2 | 5/14/2024 1:40:00 PM PDT | 1560400 | 19976 | |

**Compound MB-3A:**

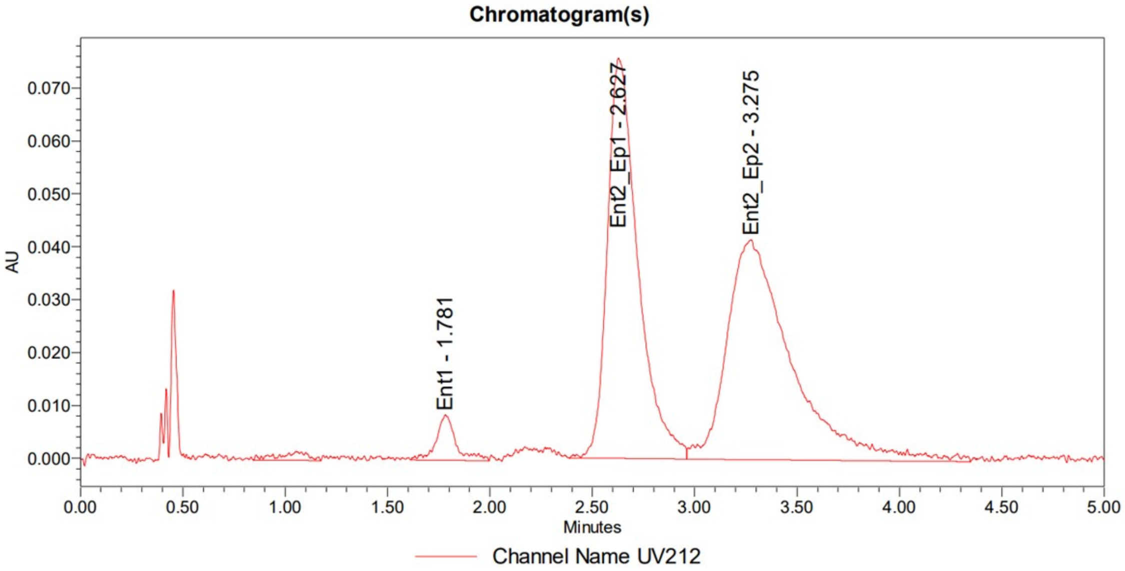

**Compound MB-3B:**

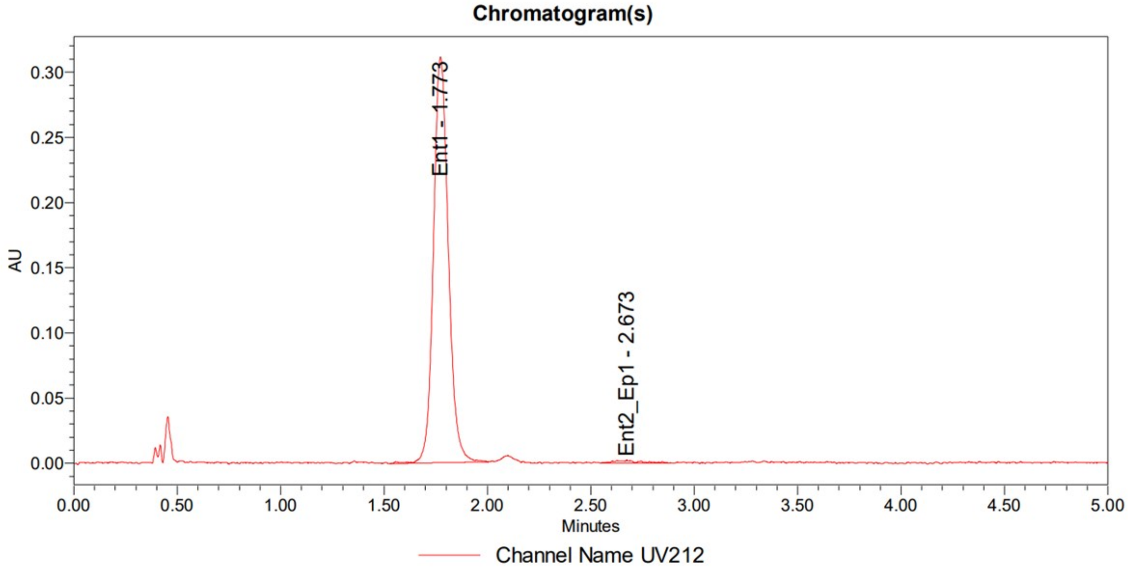

**Mixture (MB-3A + MB-3B):**

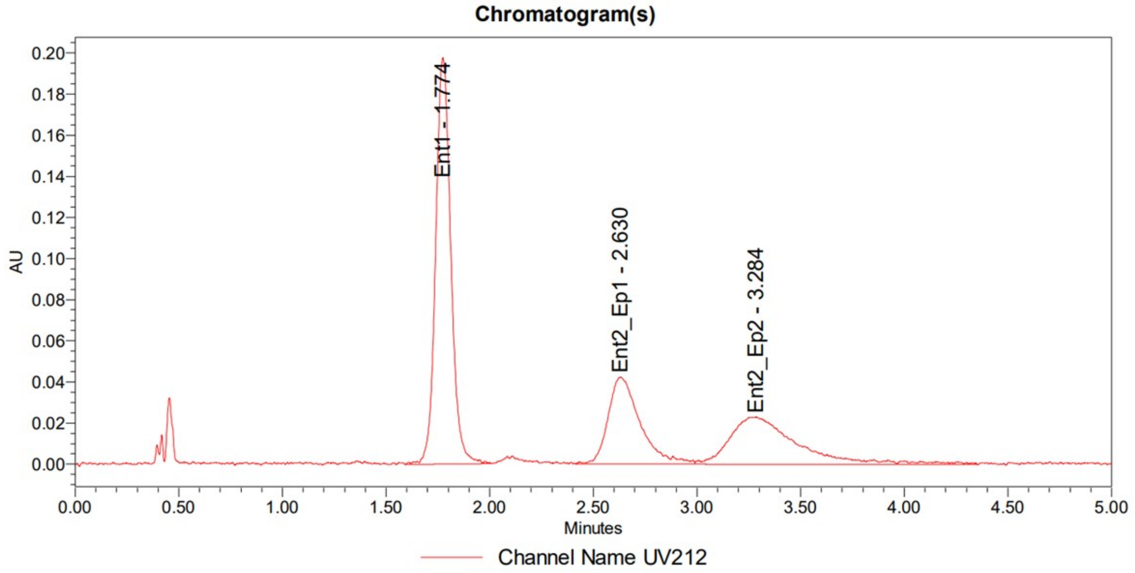

**Compounds MB-4A & MB-4B**

**Method:**

*The analysis was performed on a Waters UPC2 SFC with a Daicel IA column (3 µm, 4.6x100 mm) under isocratic conditions (3.3 mL/min, 45% MeOH / $CO_2$, 1600 psi backpressure) at 30 °C. The enantiomers were detected by UV light (212 nm).*

**Summary:**

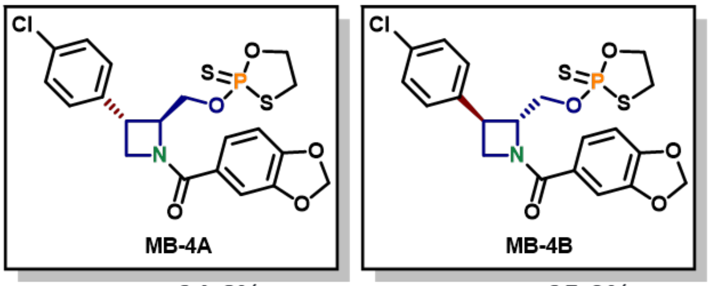

e.e. = 94.6% e.e. = 95.0%

Peak Areas Summary
Area Summarized by Name

| | SampleName | Vial | Date Acquired | Ent1 (µV*sec) | Ent2_Ep1 (µV*sec) | Ent2_Ep2 (µV*sec) |
|---|---|---|---|---|---|---|
| 1 | MB-4A | 1:E,1 | 5/14/2024 1:52:11 PM PDT | 50252 | 864113 | 956065 |
| 2 | MB-4B | 1:E,2 | 5/14/2024 1:56:17 PM PDT | 4067757 | 49660 | 55406 |

**Compound MB-4A:**

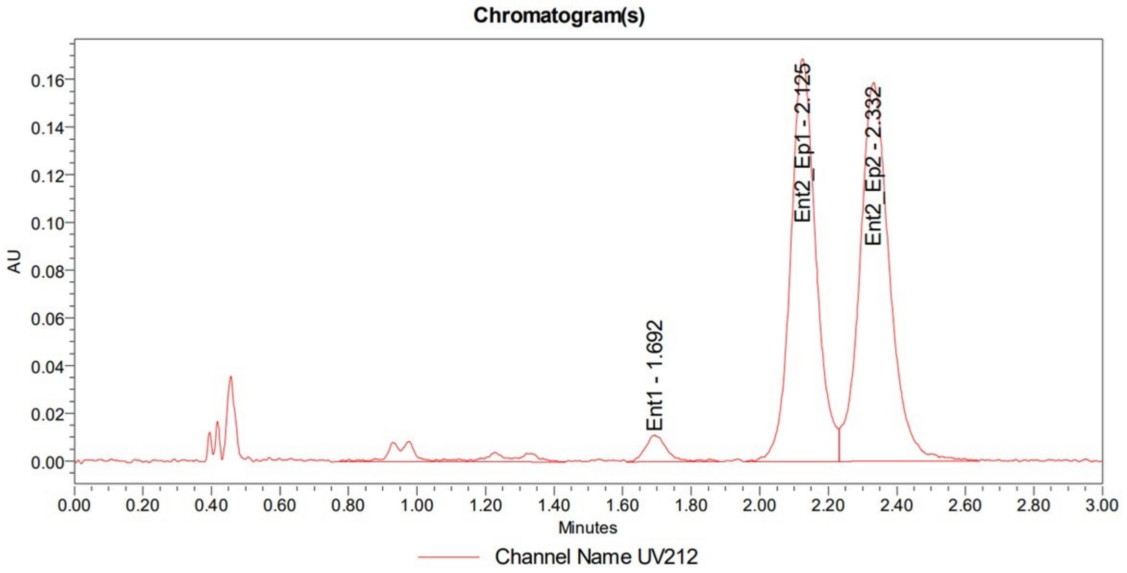

**Compound MB-4B:**

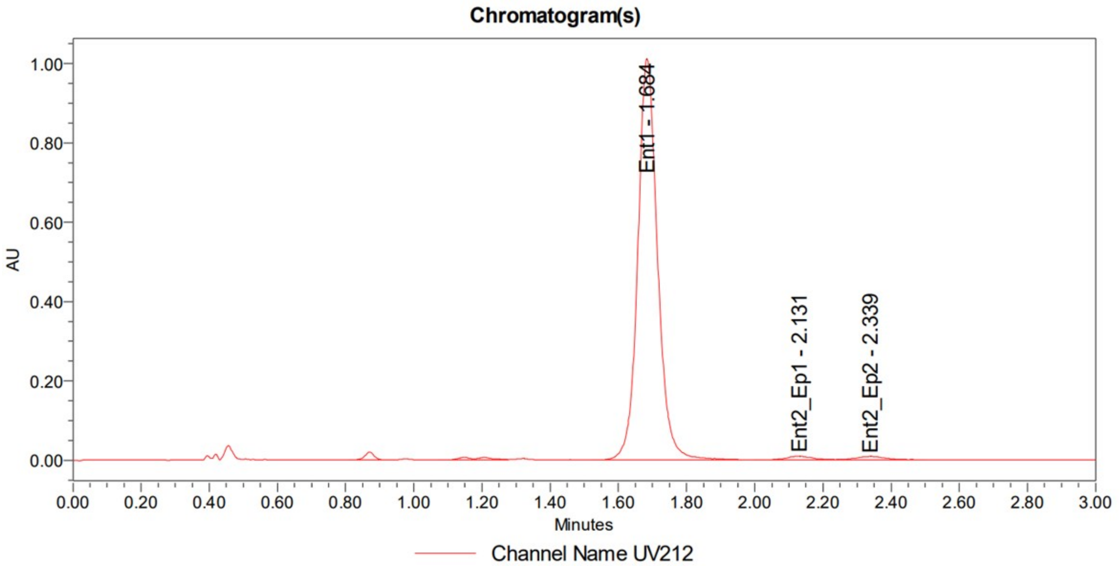

**Mixture (MB-4A + MB-4B):**

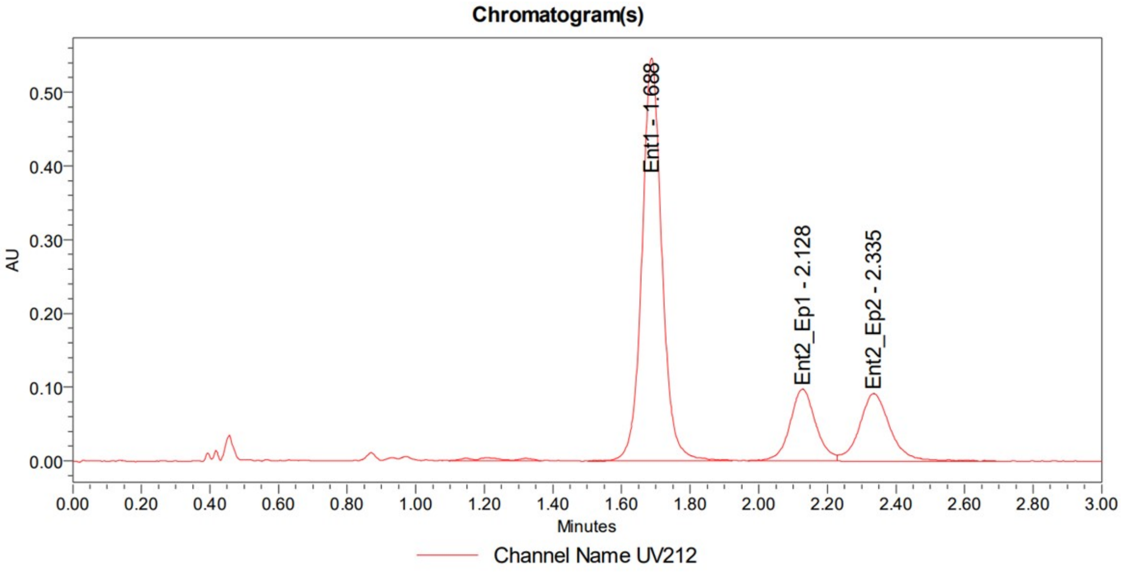

**Compounds MAH-1A & MAH-1B**

**Method:**

*The analysis was performed on a Waters UPC2 SFC with a* ***Daicel IC*** *column (3 µm, 4.6x100 mm) under isocratic conditions (3.3 mL/min, 50% MeOH / $CO_2$, 1600 psi backpressure) at 30 °C. The enantiomers were detected by UV light (218 nm).*

**Summary:**

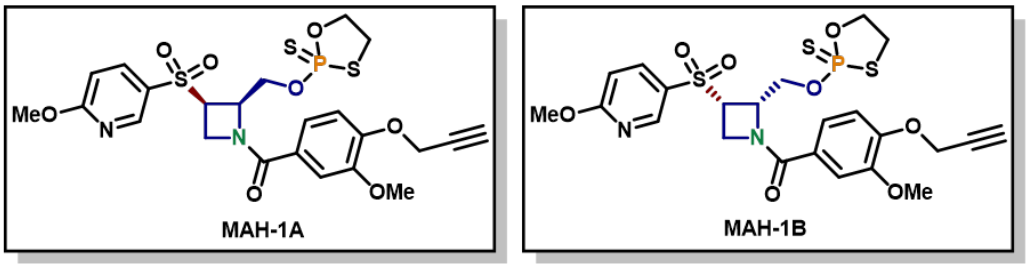

e.e. = 96.8%  e.e. = 95.1%

Peak Areas Summary
Area Summarized by Name

| | SampleName | Vial | Date Acquired | Ent1a (µV*sec) | Ent1b (µV*sec) | Ent2a (µV*sec) | Ent2b (µV*sec) |
|---|---|---|---|---|---|---|---|
| 2 | MAH-1A | 1:F,2 | 5/15/2024 8:30:15 PM PDT | 1220878 | 1256907 | 25861 | 14683 |
| 3 | MAH-1B | 1:F,3 | 5/15/2024 8:37:54 PM PDT | 30640 | 33231 | 1305043 | 1247090 |

**Compound MAH-1A:**

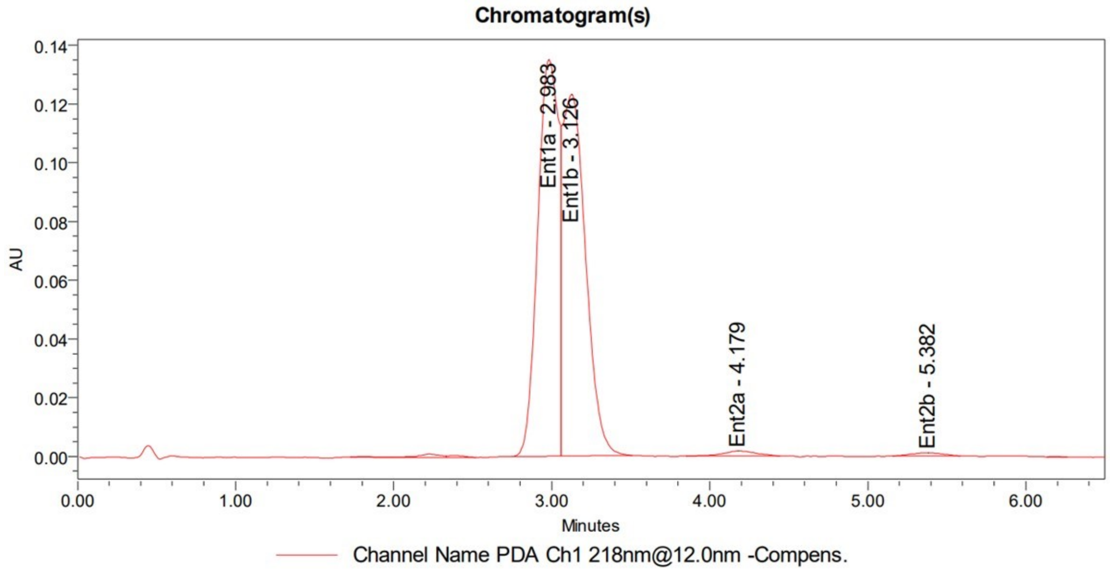

**Compound MAH-1B:**

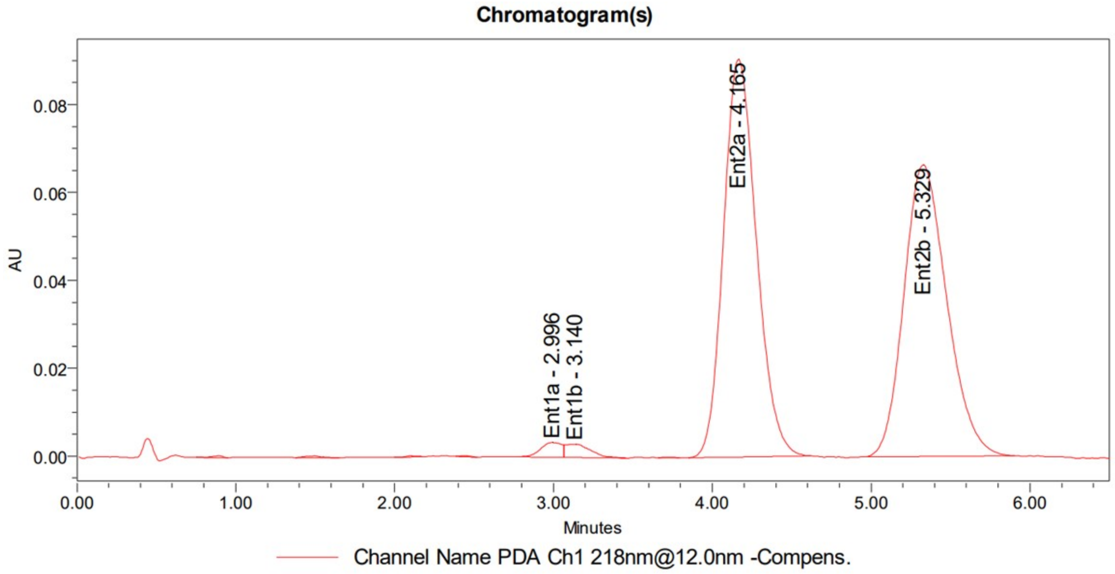

**Mixture (MAH-1A + MAH-1B):**

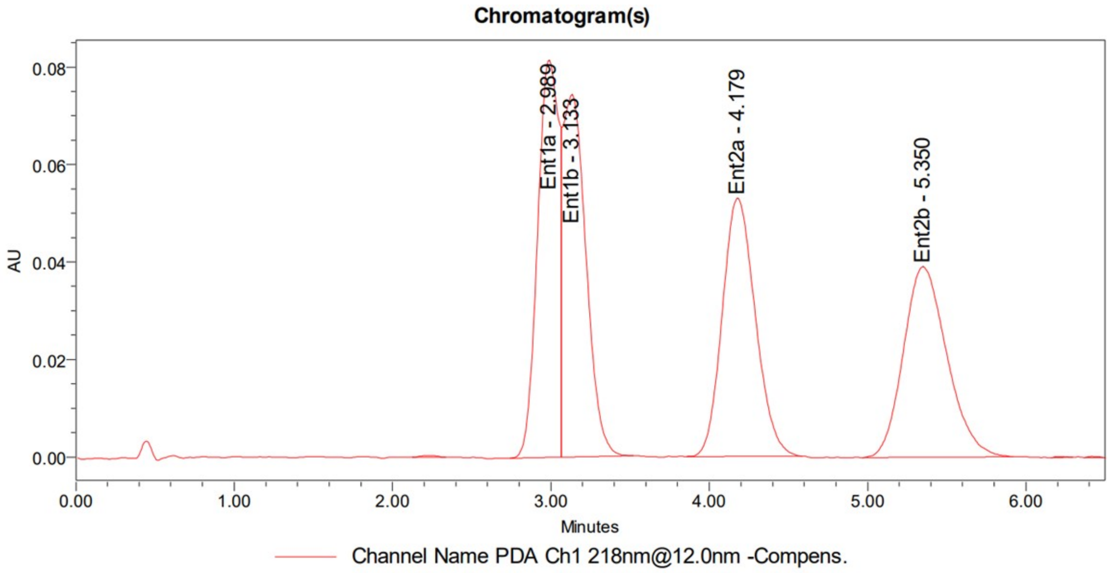

**Compounds MAH-2A & MAH-2B**

**Method:**

*The analysis was performed on a Waters UPC2 SFC with a **Daicel IH** column (3 μm, 4.6x100 mm) under isocratic conditions (3.3 mL/min, 45% MeOH / $CO_2$, 1600 psi backpressure) at 30 °C. The enantiomers were detected by UV light (217 nm).*

**Summary:**

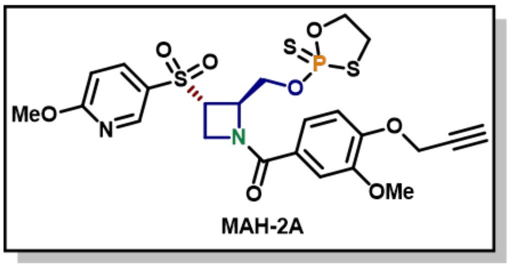

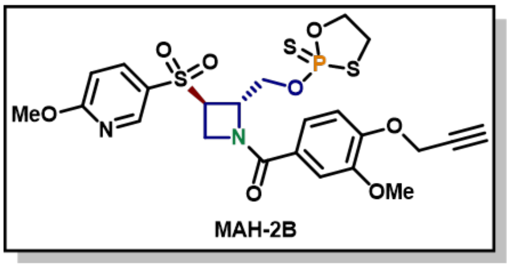

e.e. = 100%

e.e. = 94.3%

Peak Areas Summary
Area Summarized by Name

| | SampleName | Vial | Date Acquired | Ent1_Ep1 (μV*sec) | Ent1_Ep2 (μV*sec) | Ent2_Ep1 (μV*sec) | Ent2_Ep2 (μV*sec) |
|---|---|---|---|---|---|---|---|
| 1 | MAH-2A | 1:B,1 | 5/15/2024 12:27:48 PM PDT | 235164 | 259403 | | |
| 2 | MAH-2B | 1:B,2 | 5/15/2024 12:32:46 PM PDT | 7403 | 13189 | 367436 | 331954 |

**Compound MAH-2A:**

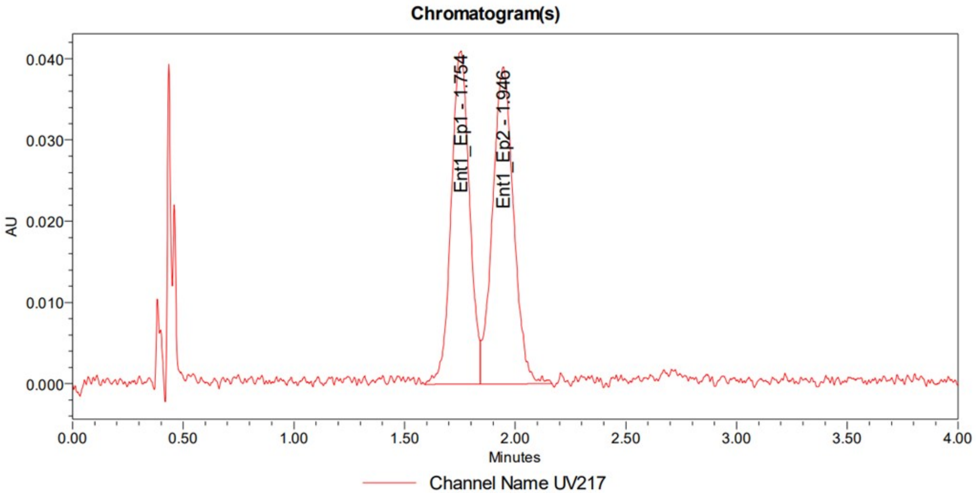

**Compound MAH-2B:**

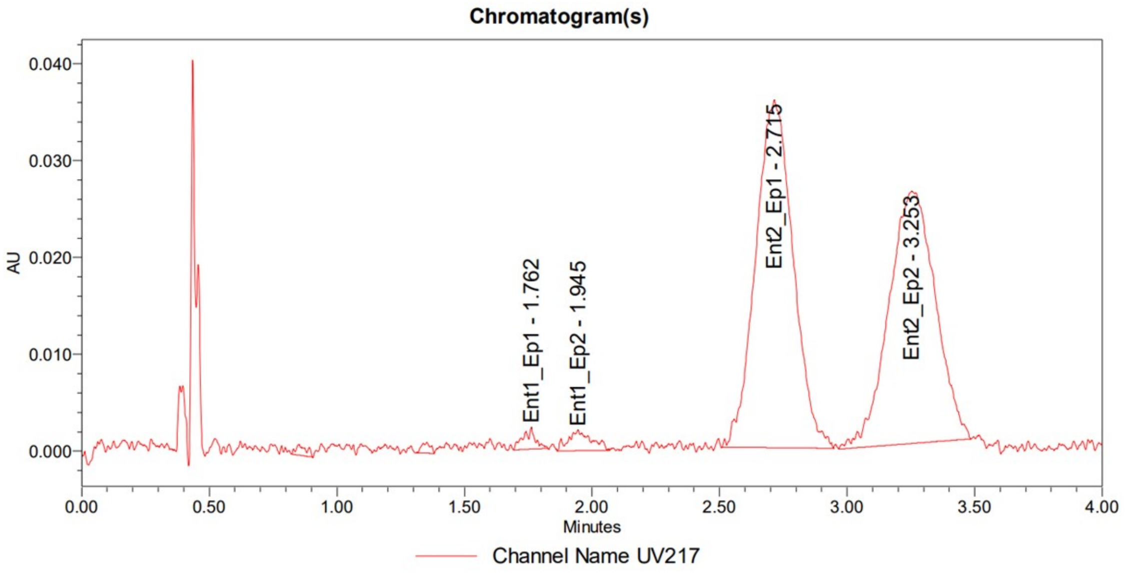

**Mixture (MAH-2A + MAH-2B):**

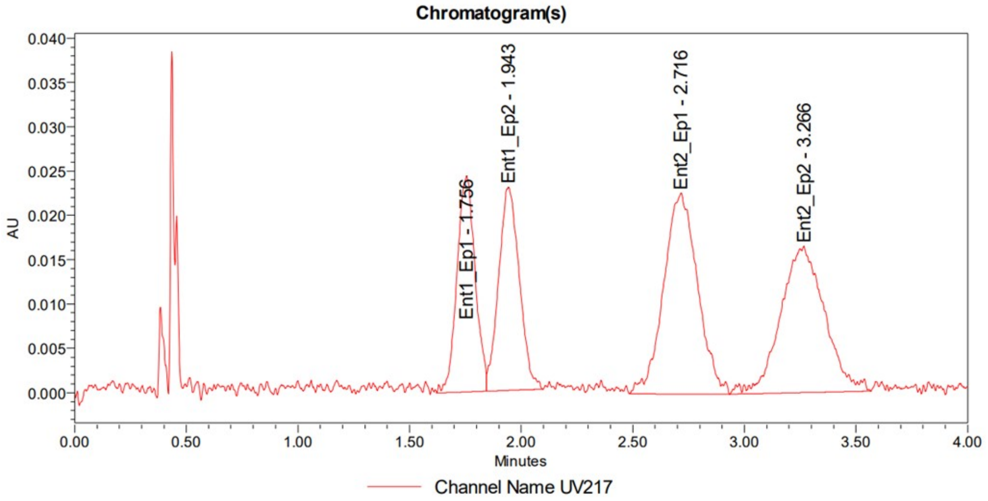

**Compounds MAH-3A & MAH-3B**

**Method:**

*The analysis was performed on a Waters UPC2 SFC with a Daicel IH column (3 µm, 4.6x100 mm) under isocratic conditions (3.3 mL/min, 45% MeOH / $CO_2$, 1600 psi backpressure) at 30 °C. The enantiomers were detected by UV light (218 nm).*

**Summary:**

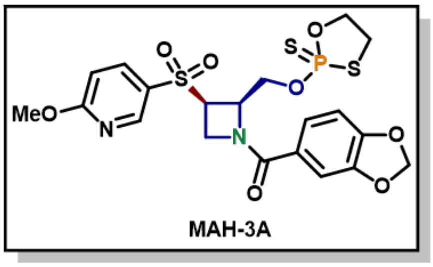

e.e. = 95.8%

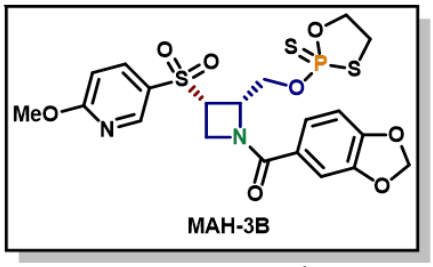

e.e. = 95.7%

Peak Areas Summary
Area Summarized by Name

| | SampleName | Vial | Date Acquired | Ent1_Ep1 (µV*sec) | Ent1_Ep2 (µV*sec) | Ent2_Ep1 (µV*sec) | Ent2_Ep2 (µV*sec) |
|---|---|---|---|---|---|---|---|
| 1 | MAH-3A | 1:C,1 | 5/15/2024 12:43:03 PM PDT | 19491 | 14872 | 790441 | 821582 |
| 2 | MAH-3B | 1:C,2 | 5/15/2024 12:48:45 PM PDT | 732593 | 711869 | 18679 | 13127 |

**Compound MAH-3A:**

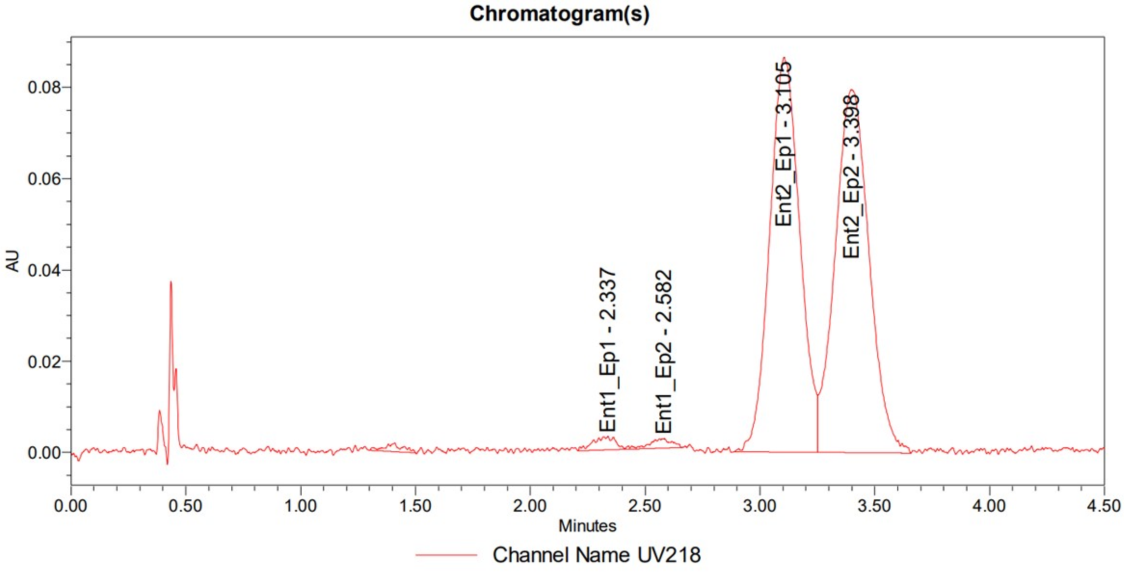

**Compound MAH-3B:**

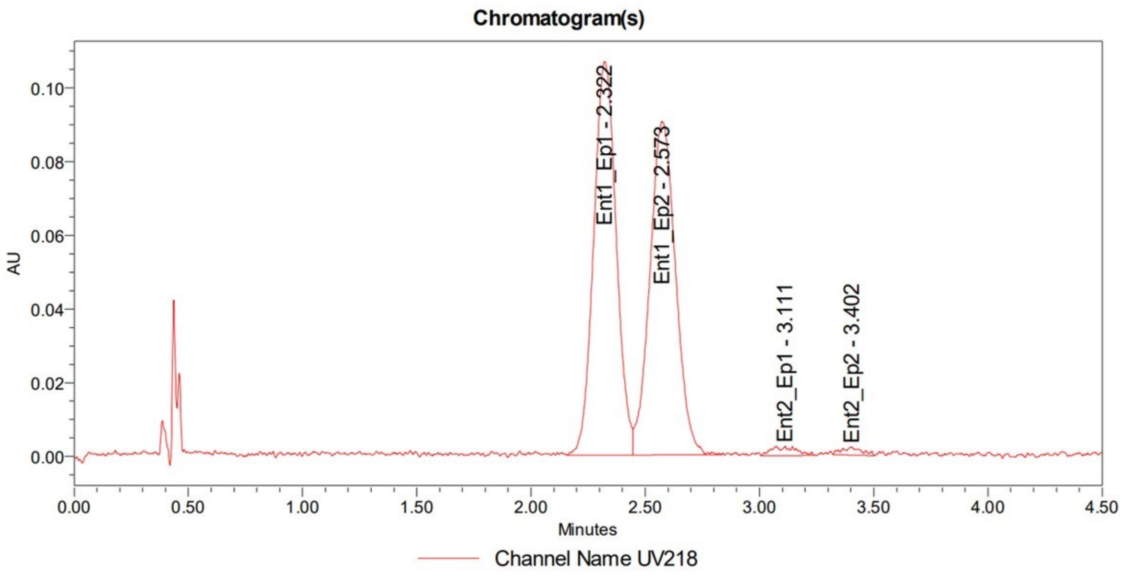

**Mixture (MAH-3A + MAH-3B):**

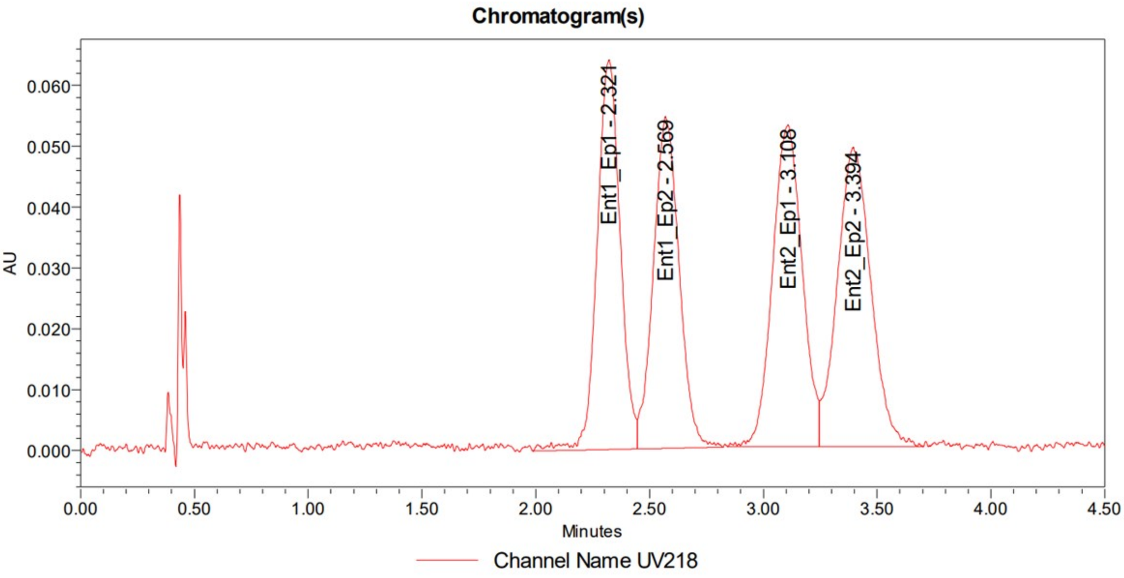

**Compounds MAH-4A & MAH-4B**

**Method:**

*The analysis was performed on a Waters UPC2 SFC with a* ***Daicel IH*** *column (3 µm, 4.6x100 mm) under isocratic conditions (3.3 mL/min, 50% MeOH / $CO_2$, 1600 psi backpressure) at 30 °C. The enantiomers were detected by UV light (217 nm).*

**Summary:**

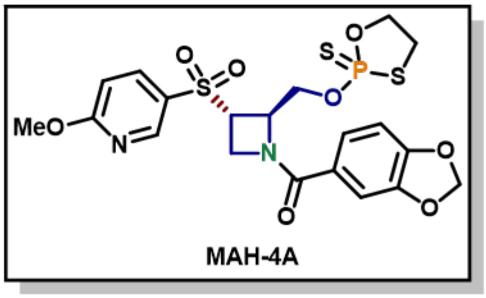

e.e. = 96.8%

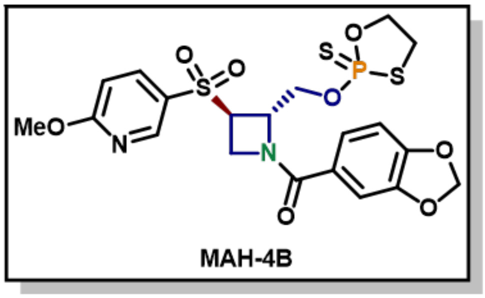

e.e. = 91.0%

Peak Areas Summary
Area Summarized by Name

| | SampleName | Vial | Date Acquired | Ent1 (µV*sec) | Ent2 (µV*sec) |
|---|---|---|---|---|---|
| 2 | MAH-4A | 1:F,5 | 5/16/2024 10:41:37 AM PDT | 2986801 | 45834 |
| 3 | MAH-4B | 1:F,6 | 5/16/2024 10:47:46 AM PDT | 88794 | 1879348 |

**Compound MAH-4A:**

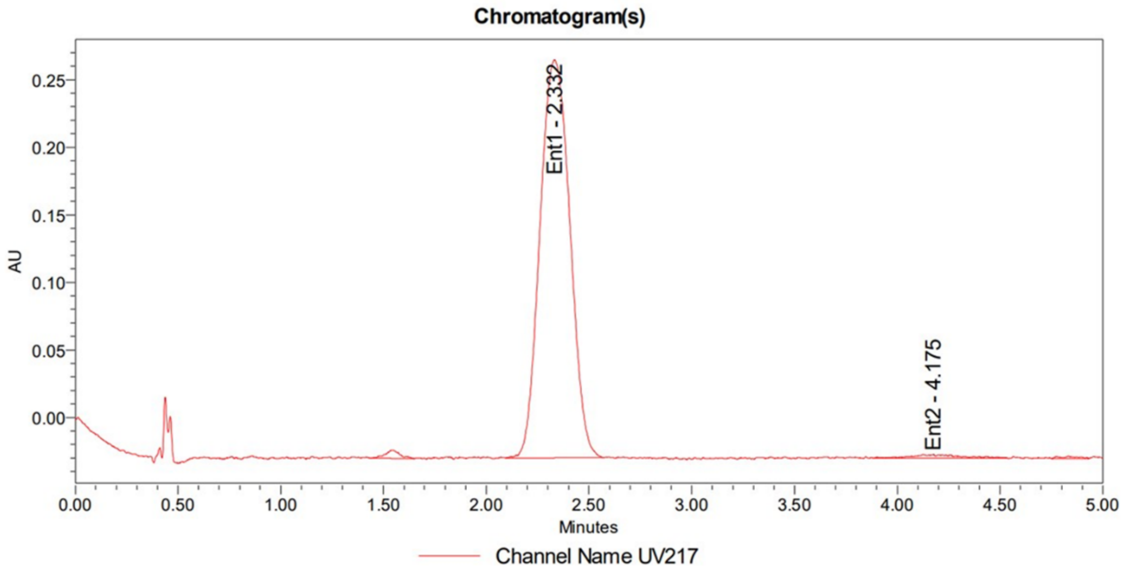

**Compound MAH-4B:**

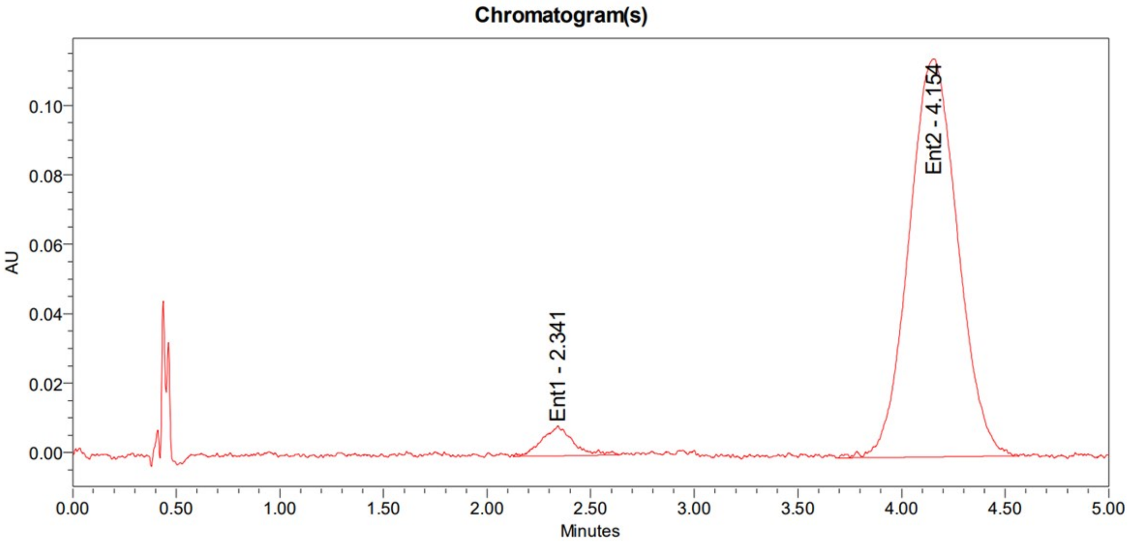

**Mixture (MAH-4A + MAH-4B):**

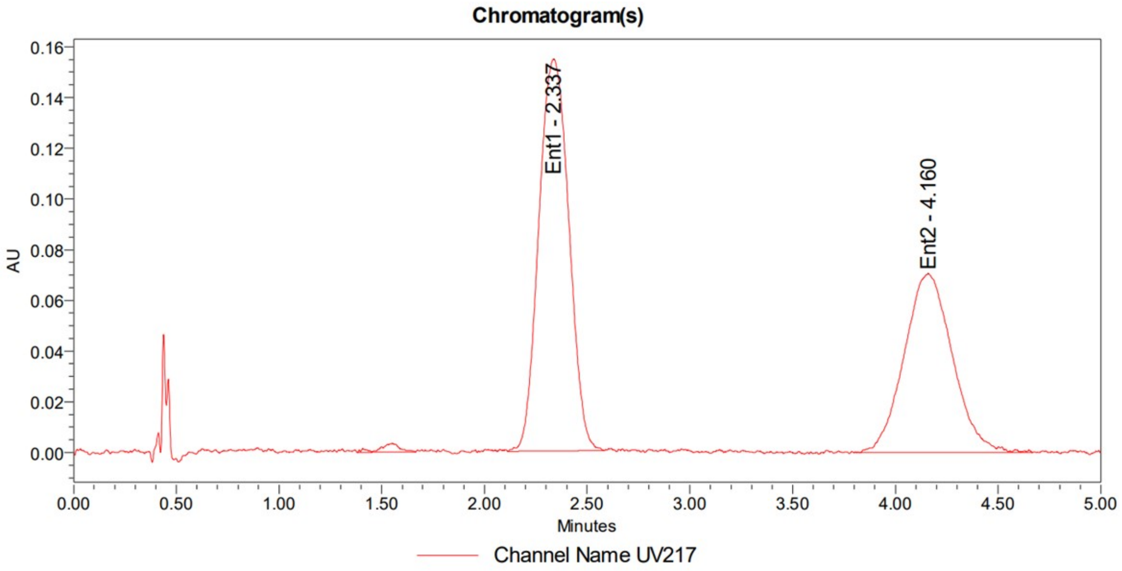
